# Leverage drug perturbation to reveal genetic regulators of hepatic gene expression in African Americans

**DOI:** 10.1101/2022.04.27.489765

**Authors:** Yizhen Zhong, Tanima De, Juan Avitia, Cristina Alarcon, Minoli A. Perera

## Abstract

**Background:** Expression quantitative loci (eQTL) studies have paved the way in identifying genetic variation impacting gene expression levels. African Americans (AAs) are disproportionately underrepresented in eQTL studies, resulting in a lack of power to identify population-specific regulatory variations especially related to drug response. Specific drugs are known to affect the biosynthesis of drug metabolism enzymes as well as other genes. We used drug perturbation in cultured primary hepatocytes derived from AAs to determine the effect of drug treatment on eQTL mapping and to identify the drug response eQTLs (reQTLs) that show altered effect size following drug treatment.

**Methods:** Whole-genome genotyping (Illumina MEGA array) and RNA-sequencing were performed on 60 primary hepatocyte cultures after treatment with 6 drugs (Rifampin, Phenytoin, Carbamazepine, Dexamethasone, Phenobarbital, and Omeprazole) and at baseline (no treatment). eQTLs were mapped by treatment and jointly using Meta Tissue.

**Results:** We found varying transcriptional changes across different drug treatments and identified Nrf2 as a potential general transcriptional regulator. We jointly mapped eQTL with gene expression data for across all drug treatments and baseline which increased our power to detect eQTLs by 2.7-fold. We also identified 2,988 reQTLs (eQTLs with altered effect size after drug treatment), which were more likely to overlap transcription factor binding sites and uncovered a novel reQTL, rs61017966 that increases *CYP3A5* gene expression, a major drug metabolizing enzyme responsible for both drug response and adverse events across several drug classes.

**Conclusions:** Our results provide novel insights into the genetic regulation of gene expression in hepatocytes through drug perturbation and provide insight into SNPs that effect the liver’s ability to respond to transcription upregulation.

## Introduction

Mapping expression quantitative trait loci (eQTL) has revealed important regulatory variants of gene expression and has helped implicate the biological mechanism underlying GWAS associations.(Cookson et al., 2009) These regulatory effects are largely context-specific and can change with stimulation. As an example, genetic regulation of the transcriptional response to bacterial infection in immune cells (Barreiro et al., 2012, Nédélec et al., 2016, Kim-Hellmuth et al., 2017), statins treatment in LCLs (Mangravite et al., 2013), and doxorubicin treatment in iPSC-derived cardiomyocytes (Knowles et al., 2018) have been studied and provide insights into the interindividual variability to stimulus response and disease susceptibility. However, the genetic regulation of liver enzyme after drug exposure is not clearly understood, especially in populations that are largely underrepresented in current genetic studies.

The maintenance of the proper drug concentration is critical to drug efficacy and to avoiding adverse events. Proper concentrations of drug within the body are driven by the conversion of medications to metabolites through Phase I and Phase II enzymes. Hepatic enzymes are responsible for much of this conversion. This process involves two classes of drug metabolizing enzymes (DMEs): Phase I DME reactions which include oxidation, reduction, and hydrolysis through the cytochrome P450 (CYP450) proteins, and Phase II DME reactions which include glucuronidation, sulfation, and acetylation and occur through conjugation enzymes such as uridine 5′-diphosphoglucuronosyltransferases (UGTs), sulfotransferases (SULTs), N-acetyltransferases (NATs), and glutathione S-transferases (GSTs). (Jancova et al., 2010) There are more than 50 CYP enzymes found in human tissues with the CYP1, 2, and 3 families metabolize 70%-80% of drugs currently in clinical use. (Zanger and Schwab, 2013). The expression level of drug metabolism enzymes and their regulatory proteins vary greatly between individuals and populations. (Zanger and Schwab, 2013, Zhang et al., 2021) Differences in the activity and expression of these important genes have already been associated to several severe adverse drug events (ADE) such as opioid toxicity and adverse cardiovascular outcomes in individuals taking the antiplatelet drugs, just to name a few.(de Leon et al., 2003, Biswas et al., 2022) Importantly, most studies of these genes have focused on coding variants and have been conducted in mainly European and East Asian populations. Knowledge of regulatory genetic variants that result in variability to drug metabolism enzymes is an unexplored area in understudied populations like African Americans. (Relling and Klein, 2011), (Perera et al., 2013)

The expression of DMEs can be affected by drug perturbation. It is well known that specific drugs upregulate the transcription of DMEs leading to greater enzyme activity in the liver. During drug enzyme induction, drug molecules bind and activate several nuclear receptors such as the pregnane X receptor (PXR), Constitutive androstane receptor (CAR), Glucocorticoid receptor (GR), and Aryl hydrocarbon receptor (AhR) resulting in increased transcription. (Prakash et al., 2015) The genetic basis underlying interindividual variability in transcriptional response upon drug enzyme induction is largely unknown and is important to inform on the ability of individuals to respond to drug exposure with clinical relevance in multi-drug dosing. It should be noted that African Americans suffer from a higher rate of ADEs and death due to ADE than other populations. (Shepherd et al., 2012)

Lastly paired transcriptomic and genomics data in under-represented populations is scarce, limiting the ability to conduct well-powered eQTL mapping studies. Approximately 13% of GTEx subject self-identify as African American, however, there are only 15 African ancestry subjects with both genotype and gene expression data in liver – leaving discovery of population-specific eQTLs woefully underpowered. Moreover, as we and others have shown, population specific regulatory variation may uncover the biological underpinning of GWAS associated loci. (Zhong et al., 2020, Gay et al., 2020)

Here we investigated the transcription and genetic regulation changes in response to multiple drug perturbations in hepatocytes, a highly clinically relevant cell type, that are derived from African American donors. Our analysis addresses the three questions 1) what are the commonalities and specificity of the transcriptional response across different drug inducer exposures. 2) what are the potential genetic regulators of drug-induced transcriptional changes, and 3) what are the regulatory variants underlying response to drug perturbation. Together, we comprehensively characterize the genetic regulation in hepatocytes after drug purtubation and leveraging gene induction to identify novel genetic variants that affect drug response in African Americans.

## Results

### Drug metabolizing enzymes are differentially expressed (DE) after treatment with enzyme inducers

To investigate the perturbation effect of drugs on the transcriptome, we compared the expression profiles of drug-treated conditions with the baseline gene expression (**Figure 1B**). DMEs were among the most differentially expressed genes (**Figure 1B**), including phase1 DME genes *CYP3A4* and *CYP2B6*, phase 2 DME genes *UGT1A4* and *UGT1A5* and the transporter gene, *SLC5A12*. The average number of up-regulated genes was 332 and the average number of down-regulated genes was 289 across 6 drug treatment conditions (FDR<0.01, log(FC)>=0.5). Phenobarbital (458 up-regulated genes and 832 down-regulated genes) and Omeprazole treatment (597 up-regulated gene and 673 down-regulated gene) had the largest number of DE genes, suggesting these two drugs induced a stronger transcriptional response compared to other treatments. Carbamazepine treatment (80 up-regulated genes and 20 down-regulated genes) had the least number of DE genes. We observed the same pattern when using a less stringent threshold to identify DE genes (FDR<0.01 with no log(FC) threshold, **Sup Figure 1**).

**Figure 1:**
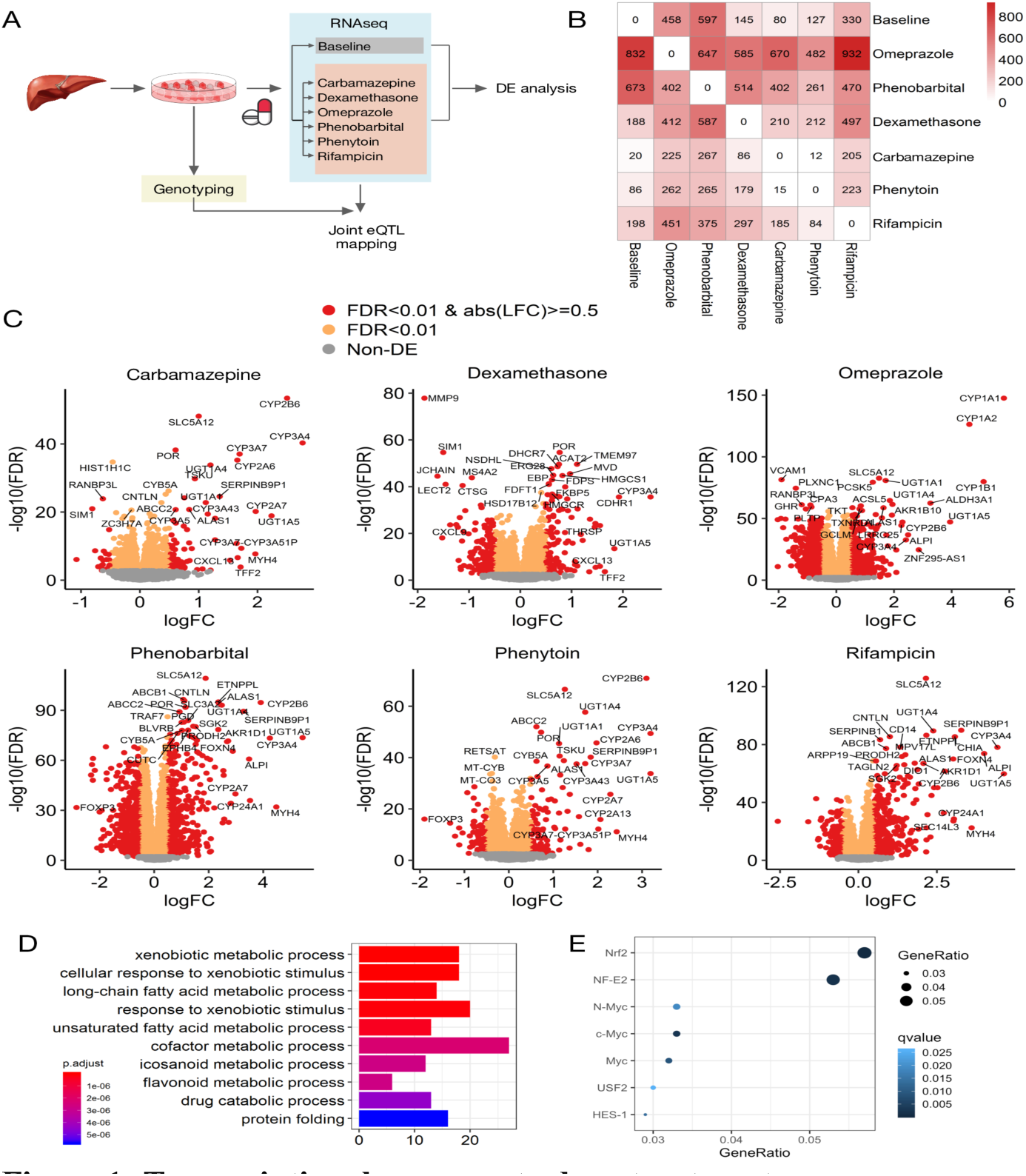
Transcriptional response to drug treatments. **A**) 60 livers from African American donors were obtained and cultured primary hepatocytes extracted. Genome-wide genotyping and RNA sequencing at baseline (no treatment) and after treatment with 6 drugs known to induce drug metabolizing enzymes were obtained to measure gene expression. The transcriptional change with drug treatments were assessed with differential gene expression analysis and the genetic regulation change by jointly mapping eQTLs across all conditions. **B**) Number of DE genes with FDR<0.01 and abs[Log (FC)] >=0.5. The values above the diagonal show the number of up-regulated genes and the values below the diagonal show the number of down-regulated genes. **C**) Volcano plots show the DE genes of each drug treatment comparing with baseline. Drug metabolizing enzymes including *CYP2B6, CYP3A4, UGT1A5* and *UGT1A4* were some of the top DE genes. **D**) Pathways represented by commonly upregulated genes (genes that were significantly upregulated in all conditions at FDR<0.01) The x-axis shows the number of genes in each ontology category with the color scale representing the adjusted p-value of the enrichment. **E**) Enrichment of transcriptional binding sites in commonly upregulated genes. The x-axis shows the GeneRatio for each significant transcriptional binding factor (number of commonly upregulated genes in each category divided by the total number of genes in each category).

We further sought to identify the commonalities across different drug treatments which may implicate the common mechanism of regulation. Using a threshold of FDR<0.01, we identified 200 commonly up-regulated DE genes and 196 commonly down-regulated genes across all drug treatments. Commonly upregulated genes include: *CYP3A5, CYP2B6, UGT1A1, UGT1A4* and *UGT1A5* (Supplementary file1). Gene ontology (GO) analysis showed commonly up-regulated genes were strongly enriched in metabolic process, xenobiotic stimulus and drug metabolic process (**Figure 1D**). Commonly up-regulated genes were also enriched for genes with Nrf2 transcription factor binding sites (TFBS) (**Figure 1E**). The enrichment remains when using a less stringent threshold to identify DE genes (FDR<0.05). Nrf2 is the key regulatory of anti-oxidative response and its role in the coordinated regulation of drug metabolism is gaining attention(Shen and Kong, 2009a). Nrf2 is also upregulated in most drug treatment conditions with the exception of Dexamethasone (Log fold change: Omeprazole=0.44, FDR=4.04e-31, Phenobarbital=0.41, FDR=3.46e-28 Dexamethasone=-0.03, FDR=0.39, Carbamazepine=0.16, FDR=6.77e-6, Phenytoin=0.27, FDR=3.05e-14, Rifampicin=0.53, FDR=3.92e-42). Commonly down-regulated genes have fewer specific functions than commonly up-regulated genes with the top enriched GO terms being nucleotide biosynthetic process and developmental process (Supplementary File 2, Sup Figure 2).

### Inter-individual variability is the major source of transcriptome variability

We used a linear mixed model to investigate the sources of variation in gene expressions (**Fig 2A**). The largest variance was attributed to inter-individual differences (mean ± sd = 0.612±0.162) which may be due to genetics, epigenetics, physiological conditions, pathological factors and environmental influences. Genes with large inter-individual variance were enriched in immune response pathways. The inter-treatment difference (labeled drug in **Figure 2A**) was the second largest tested factor contributing to the transcriptome variability (mean ± sd = 0.041±0.041, **Figure 2B**). Genes with high inter-treatment differences include *CYP1B1, CYP1A1* and *CYP1A2* which were significantly upregulated only with Omeprazole treatment (**Figure 1C**). These genes belong to the “Ahr gene battery” and are induced through the activation of the Ahr nuclear receptor. This may suggest the drug-specific transcriptional response is mostly due to the selective binding to nuclear receptors.

**Figure 2:**
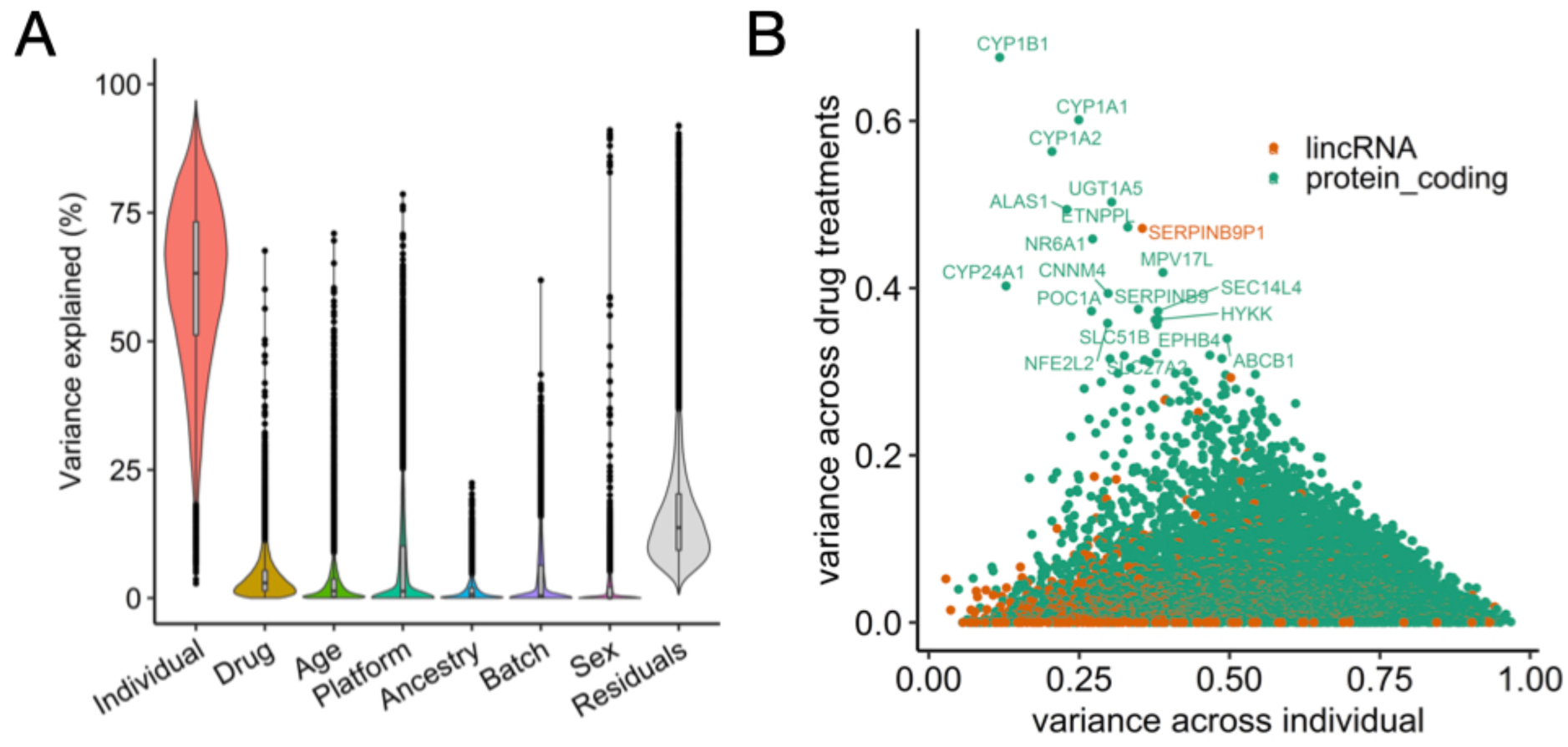
Factors contributing to the variability in cross-drug gene expression. We used a linear mixed model to partition variability in expression data across multiple drug treatment conditions. **A**) Distribution of the percentage of variability explained by each factor. The most variability is attributed to inter-individual variability and then inter-condition variability (labeled “Drug”). **B**) Comparison of inter-individual variability with inter-condition variability of each gene. *CYP1B1, CYP1A1* and *CYP1A2* have high inter-condition variability.

Genes with large proportions of variability explained by other factors can also provide interesting insights. Genes with large contribution from sex were sex chromosome-linked such as *EIF1AY, UTY* and *KDM5D* as expected. The majority of the variability in *ZNF518B* (71.0%), *TSPYL5* (69.6%), *ZIC1* (65.2%) were explained by age. Notably, these genes were previously found to be associated with age in other tissue types (Martella et al., 2014, Mimura, 2019, Bacos et al., 2016). This suggests that this age-dependent expression pattern remains in African American hepatocytes and our method was robust in identifying contributing factors of gene expression.

### Joint-mapping of eQTLs across drug treatments increases detection power

We first performed eQTL mapping separately in each drug treatment condition (TBT, Treatment-By-Treatment). We consistently detected more eGenes with drug treatment as compared with baseline. The enrichment of TBT eQTLs in functional annotations did not differ between drug treatments and baseline (**Sup Figure 3**).

To leverage the repeated measurements of gene expression of each primary hepatocyte cell culture in different conditions, we used a linear mixed model to jointly map eQTLs while accounting for the correlation of expression in different drug treatment conditions implemented in Meta-Tissue (Sul et al., 2013). Meta-Tissue estimates the intercept and coefficient of genotype on gene expression for each condition and performs meta-analysis after uncorrelation of coefficient estimates. Meta-Tissue also estimates the posterior probability that the effect is present in each condition (represented as an m-value).

With the Meta-Tissue fixed effect (FE) and random effect (RE) meta-analysis of uncorrelated coefficients, we increased the number of identified eGenes (Meta-Tissue eGene) and eQTLs (Meta-Tissue eQTLs) compared with that of baseline TBT eQTL analysis (**Table 1**). The number of unique eGenes (n=665) combined across all TBT conditions was still less than that of Meta-Tissue FE model (n=818) or RE model (734). The fixed effect model assumes the eQTL effect is the same across different conditions, while the random effect model allows the eQTL effects to vary from condition to condition. 694 eQTLs are unique to the RE analysis and 6,042 eQTLs are unique to the FE analysis, suggesting eQTLs have both fixed effect or random effect across drug treatment conditions.

**Table 1:** eGene and eQTL discovery with different mapping methods.

|  | **TBT baseline**<br>**(eGenes/eQTLs)** | **TBT all**<br>**conditions**<br>**(eGenes/eQTLs)** | **Meta-Tissue fixed**<br>**effect**<br>**(eGenes/eQTLs)** | **Meta-Tissue**<br>**random effect**<br>**(eGenes/eQTLs)** |
| --- | --- | --- | --- | --- |
| FDR < 0.05 | 299/24,799 | 665/48,027 | 818/53,487 | 734/48,134 |

The number of eGenes (genes with at least one significant eQTL) and eQTLs identified with each method using an adjusted p-value threshold of less than 0.05. **TBT baseline:** Significant eGenes and eQTLs in hepatocytes after multi-testing correction on nominal p-values calculated from the Meta-Tissue linear regression in the baseline condition only. **TBT all conditions:** We applied multi-testing correction on nominal p values calculated from the Meta-Tissue linear regression in each condition separately (6 drug treatment conditions and baseline) and identified the intersection of unique eGenes and eQTLs that are significant in any conditions. **Meta-Tissue fixed effect:** Significant eGenes and eQTLs in the fixed effect model. **Meta-Tissue random effect:** Significant eGenes and eQTLs in the random effect model.

The Meta-Tissue FE eGenes are significantly enriched in the glutathione derivative metabolic process (q value = 0.00358) and xenobiotic catabolic process (q value = 0.03) within the biological process ontologies while the baseline eGenes were not (**Figure 3B**). This demonstrates that drug treatment with enzyme inducers uncovered important genetic regulation of drug metabolism that was undetectable at baseline. We also increased the replication with eGenes found in GTEx liver (n=153). The number of overlapping eGenes between Meta-Tissue FE model and GTEx liver was 501 compared with 220 of baseline eGenes mapped through a TBT approach (**Figure 3A**).

**Figure 3.**
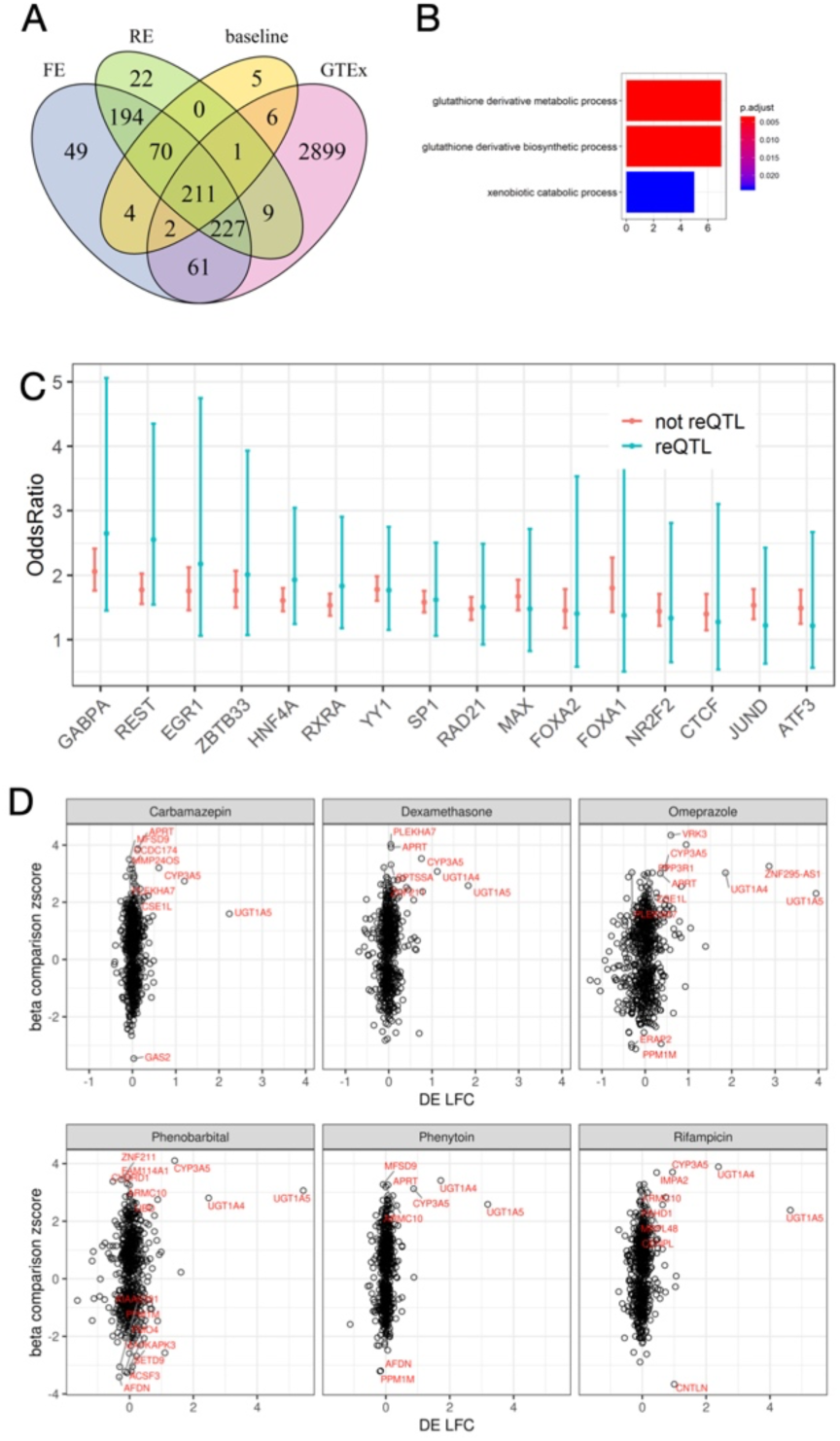
Identification of reQTLs. **A**) The Venn Diagram comparing the eGenes identified in the Meta-Tissue FE model, Meta-Tissue RE model, baseline TBT and the GTEx liver dataset. Using Meta-Tissue increased the replication in GTEx liver as compared with TBT baseline results. **B**) Enrichment analysis of Meta-Tissue FE eGenes. The x axis is the number of eGenes in each category with the color of each bar representing the adjusted enrichment p-value. The baseline eGenes were not enriched in metabolic processes while the FE eGenes showed enrichment in glutathione derivative metabolic process. **C**) Enrichment of reQTLs and non-reQTLs within TFBS annotations. reQTLs show larger OR for overlap with GABPA, REST, ERG1 ZBTB33, RXRA and HNF4A binding peaks than non-reQTLs. **D**) Comparison of log fold change of eGenes expression (DE LFC) and the highest absolute z score among eQTLs for each eGene. This analysis showed independence between the differential gene expression and the effect size of eQTLs for those genes that are differentially expressed between treatment and baseline.

### Genetic regulation change upon drug response

We examined the heterogeneity of eQTLs effects across conditions with I^2^ statistics which estimates the percentage of variation in eQTL effect size explained by heterogeneity. We found TBT eQTLs had more extreme I^2^ than the meta-tissue FE model and the meta-tissue RE model eQTLs (**Sup Figure 4**), suggesting that the TBT model might be more powerful in identifying drug-specific eQTLs than meta-analysis based methods. Therefore, we combined the Meta-Tissue FE eQTLs and TBT eQTLs (n=59169) to identify if any of these eQTLs were drug response specific (reQTL). Note that heterogeneity measured by I^2^ captures the variance across all conditions instead of the difference between drug treatment and baseline.

We defined reQTLs as eQTLs with altered effect size after drug treatment. We used the Z-score method (Kim-Hellmuth et al., 2017) to test if the eQTL effect size after drug treatment was significantly different from the effect size at baseline while accounting for the correlation of beta estimates from linear mixed model. The magnitude of changes in the eQTL effect size is measured as a z score. We computed the significance of z score by comparing it with null z scores calculated from random permutations. We identified 2,988 reQTLs for 194 genes using q value < 0.05 across 6 drug treatment conditions [Supplementary File 3]. The remaining Meta-Tissue FE eQTLs and TBT eQTLs were classified as non-reQTL.

The z scores were mostly correlated between drug treatment conditions for reQTLs, suggesting the shared genetic regulation changes across drug treatments (**Sup Figure 5**). Interestingly, the highest correlation of z scores was for Carbamazepine and Phenytoin (Pearson correlation=0.880, p-value<2.2e-16) which is consistent with the high similarity of their transcriptome profiles. In particular, the highest z scores were found for *UGT1A5, UGT1A4* and *CYP3A5* eQTLs across all drug treatment conditions (**Figure 3D**). We also identified reQTLs that were specific to one drug treatment (N = 2,541). For example, regulatory variants for *VRK3* were only seen after omeprazole treatment (**Figure 3D**; **Sup Figure 6**). *VRK3* belongs to the vaccinia-related kinases family and affects cell cycle progression as a serine/threonine kinase (Lee et al., 2017). Omeprazole has also been shown to restrict cell proliferation and induce cell cycle arrest (Hou et al., 2018, Paz et al., 2020). This suggests genetic regulation of *VRK3* might play a role in the response to Omeprazole treatment by regulating the cell cycle.

The enrichment of reQTL and non-reQTL in TFBS showed that while both groups were enriched for several TFBS, reQTLs have a larger OR for overlap with GABPA (also known as Nrf2), REST, ERG1, ZBTB33, RXRA and HNF4A binding peaks as compared to non-reQTLs (**Figure 3C**). As described previously, Nrf2 was enriched in commonly up-regulated genes across 6 drug treatments as previously stated. The enrichment of reQTLs in Nrf2 binding peaks further highlights the role of Nrf2 in modulating drug response. These results suggest the importance of these TFBS in the appropriate transcriptional response to drug exposure, such as upregulating drug clearance mechanisms to prevent toxicity. This response may be blunted or enhanced by reQTLs.

We compared the population differentiation in allele frequency between reQTLs and non-reQTLs. We used Wright’s fixation index (Fst) to measure the extent of variation in allele frequency between Africans and Europeans. The reQTLs have higher Fst than non-reQTLs (**Sup Figure 7**, Wilcoxon rank sum one-sided test p=3.62e-10, median Fst of reQTL = 0.118; median Fst of non-reQTL: 0.111). This finding suggests allele frequency differences in regulatory variants link to transcriptional response may underly population differences in drug response or ADEs, though functional assays are needed validate these findings.

We then investigated the mechanism underlying the reQTLs with drug treatment. For each eGene, we compared the log fold change with the largest absolute z score among its eQTLs for each drug treatment (**Figure 3D**). There was an independent relationship between the differential expression of each gene after drug treatment and the change of eQTL effect size. For example, *PLEKHA7* is an adherens junction protein and has been associated with blood pressure in previous GWAS (Lin et al., 2011). Here we found *PLEKHA7* was not differently expressed with drug treatments (LFC; Omeprazole=-0.31, Phenobarbital= −0.10, Dexamethasone=-0.05, Carbamazepine=-0.06, Phenytoin=-0.07, Rifampicin=-0.06), but had large difference in eQTL beta estimates (maximum z score: Omeprazole=-3.04, Phenobarbital= 2.37, Dexamethasone=4.02, Carbamazepin=3.14, Phenytoin=1.67, Rifampicin=2.67) as shown in **Sup Fig 8**. We also identified lfc QTL where we tested the association between genotype with gene expression log fold change. There is a low correlation between lfc eQTL z score and reQTL z score, which further suggests the independence between gene expression change and regulatory effect change (**Sup Fig 9**).

### Analysis of *CYP3A5* expression regulation through rs61017966

Among the eGenes with reQTLs, we found one DME, *CYP3A5*, which had 59 reQTLs. *CYP3A5* is labeled as a Very Important Pharmacogene (VIP gene) within PharmGKB(Gong et al., 2021), and metabolizes ∼37% of commonly used drugs together with other *CYP3A* family genes. *CYP3A5* genotypes have been linked to interindividual variability in response to the lipid-lowering drugs, statins, and diseases such as blood pressure/hypertension (Lamba et al., 2012). CY3A5 expression is highly differentiated across populations with low expression in Europeans (due to the presence of a splice variant in *CYP3A*5 in this populations) but significantly higher expression in African populations.

When comparing the m-values of all *cis* SNPs of *CYP3A5*, we found that reQTLs were not statistically significant in baseline, suggesting that drug perturbation enhances the discovery of these regulatory variants. One of the drug-specific eQTLs, rs61017966 (Indel: CCA>A), showed that the A allele was positively associated with CYP3A5 expression (**Fig 4A**) in all treatments. Previous eQTL mapping efforts, both in GTEx and in our baseline data, show no significant eQTLs for this gene. The effect size of rs61017966 for CYP3A5 increased from 0.38 in baseline to 0.77-0.96 with drug treatments, with the largest effect size (0.96) seen with phenobarbital treatment (**Fig 4B**). rs61017966 is 15kb upstream of *CYP3A5* and is located in a strong enhancer region (based on ENCODE ChromHMM prediction) (**Fig 4C**). This SNP is positioned under multiple active histone marker peaks (H3K27ac, H3K9ac, H3K4me1 and H3K4me3) mapped in ENCODE liver ChIP-seq data. Moreover, this SNP overlaps many ChIP-seq peaks for TFBS, such as JUND, ERG1 and SP1. This SNP is also highly differentiated across human populations with the allele frequency of the alternative allele at 69% in Africans and 7% in Europeans. Based on these results, this SNP may be a key regulator of *CYP3A5* gene expression, though further functional validation and fine mapping is needed.

**Figure 4:**
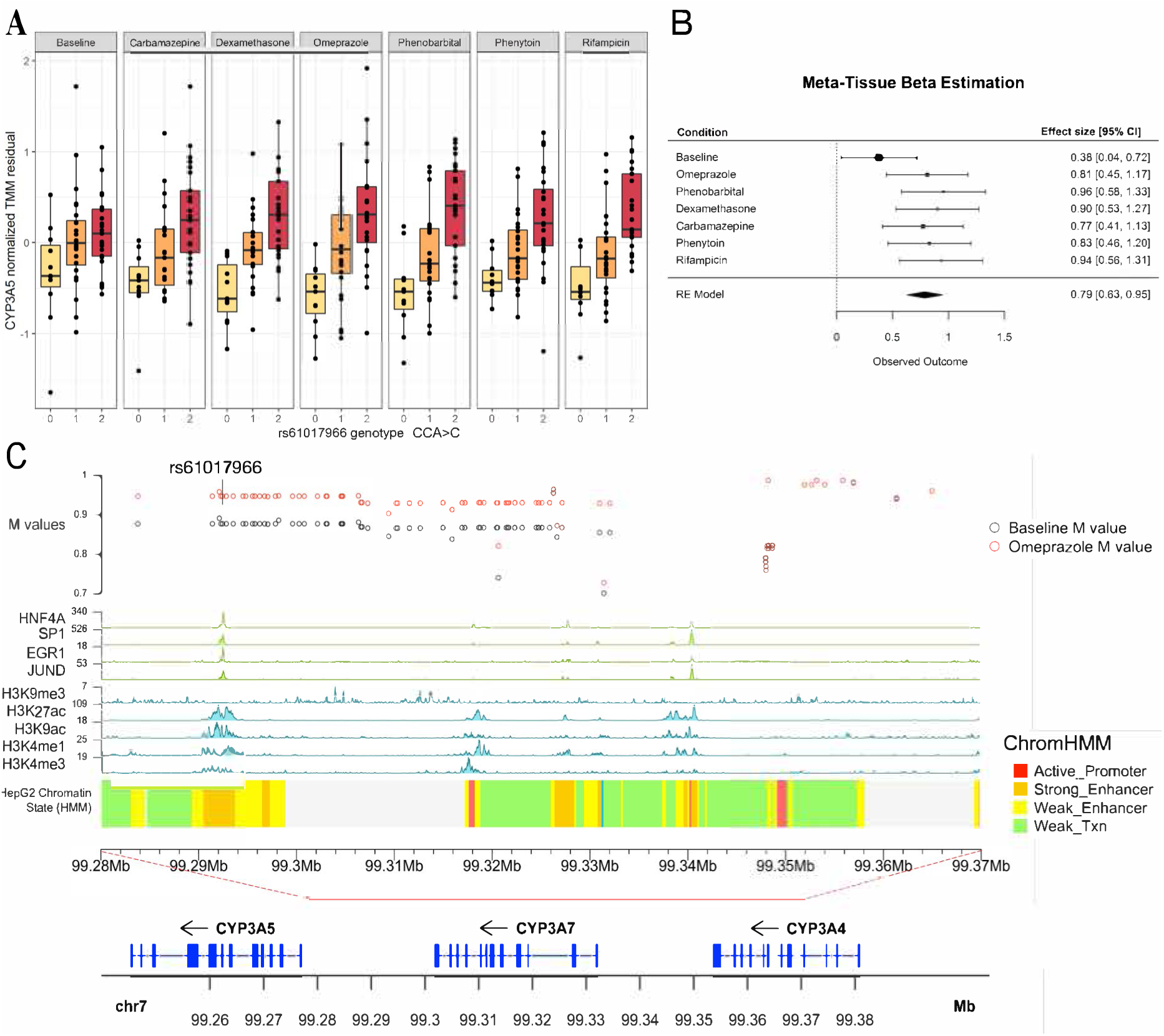
Regulation of *CYP3A5* gene expression through the reQTL, rs61017966. A) Boxplots of rs61017966 genotype and *CYP3A5* gene expression in all drug treatment and baseline. The y-axis is the normalized TMM value after regressing covariates B) Forrest plot of rs61017966 effect size and standard error estimates from Meta-Tissue linear mixed model in each drug condition. C) Genomic context of rs61017966 (annotation shown are from just upstream of the *CYP3A5* TSS to within *CYP3A4*). rs61017966 has different m-values between baseline (black points in the top panel) and after Omeprazole treatment (red points in the top panel). rs61017966 overlaps with TF binding sites (green tracks) and active histone markers (blue tracks). This SNP is 15kb upstream of CYP3A5 and locates in the strong enhancer region based on ENCODE ChromHMM prediction.

Notably, CYP3A5 activity is known to be regulated by a splicing variant, rs776746 (C/T), which is also highly differentiated between Africans and Europeans (T: 82% in AFR and 6% in EUR). We found moderate LD between rs776747 and rs61017966 in African ancestry populations (R^2^: AA hepatocyte: 0.59718, ASW: 0.608969, YRI: 0.508190,). We fitted a baseline model of CYP3A5 expression with rs776746 and a full model of CYP3A5 expression with rs776746 and rs61017966 together. We performed likelihood ratio test showing that the addition of rs61017966 significantly increased the model goodness-of-fit in the phenytoin and rifampicin treatment conditions (Baseline: p=0.866, Omeprazole: p=0.192, Phenobarbital: p=0.083, Dexamethasone: p=0.053, Carbamazepine: p=0.536, Phenytoin: p=0.045, Rifampicin: p=0.027). The finding of significance in two drug treatment conditions as oppose to all, may be due to the different degrees of upregulation of CY3A5 by the different drug treatments. Given the small sample size of our cohort, the larger the transcriptional difference the more power we have to detect the association. Therefore, rs61017966 may confer a non-coding regulatory effect in addition to the effect of the splicing variant on *CYP3A5* gene expression.

## Discussion

Understanding how genetic variation affects genes that are important for drug response promises to help deliver safe and effective drug treatment and therapeutics. Unfortunately, the variation resulting from drug exposure and the underlying genetic regulatory mechanisms are less understood. Furthermore, minority populations such as African Americans are largely underrepresented in current genetic studies even though they have repeatedly shown differences in drug response to a wide variety of medications.(Mehanna et al., 2017, Gong et al., 2016, Liu et al., 2021, Kranzler et al., 2021, Samedy-Bates et al., 2019) Studies also showed African Americans are disproportionately underrepresented in clinical trials which raises concern on the safety and efficacy of new therapies. (2018) Here, we present the first comprehensive evaluation of transcriptional response and its genetic regulation across 6 drug treatments in hepatocytes derived exclusively from African American donors.

We evaluated expression profiles with drug exposure and found varying transcriptional changes with different drugs. We found commonly upregulated genes are enriched in Nrf2 binding and eQTLs discover with both TBT and joint modelling were more likely to overlap with Nrf2 binding sites. Nrf2 plays a central role in regulating the response to antioxidative stress which is a common effect of drug treatment (Deavall et al., 2012) and there is also evidence of the crosstalk between Nrf2 and drug-activated nuclear receptors (Gong et al., 2006, Klotz and Steinbrenner, 2017, Shen and Kong, 2009a). Animal studies have shown that Nrf2 to may regulate Phase I and II enzymes.(Shen and Kong, 2009b, Aleksunes and Klaassen, 2012) Here, we further strengthen the role of Nrf2 as the master regulator of gene expression in human. Meanwhile, we found a distinct expression profile of Omeprazole treatment through the activation of its unique nuclear receptor. The mechanism of the crosstalk and coordination of shared and drug-specific transcriptional responses warrants future investigation.

We showed that inter-individual variability explains the most variance in the drug-induced transcriptome data. However, we found limited genetic regulation for most DMEs which were differentially expressed. For example, we did not find eQTLs for *CYP2B6* or *CYP3A4* which are important DMEs. This suggests other factors besides genetics may play an important role in the dynamic response to drug treatment and are left to be discovered or we lack the power to detect these smaller signals. There is a growing interest in using multi-omics approaches to decipher the variability in phenotypic traits.(Hasin et al., 2017) This might also apply to molecular traits, gene expression in our case, where the integration of genetics and epigenetics could better explain and predict inter-individual variation in transcriptional response.

In our analysis, we identified significantly more eQTLs with joint modeling across multiple drug treatment conditions. This is important as tissues from minority populations, in our case African Americans, are rare in biomedical research, leading to much smaller cohort sizes than seen with other populations. Hence the ability to increase the power to detect associations in minority populations is crucial to bring equity to precision medicine. We demonstrated that we can increase the ability to detect eQTLs by introducing additional variation into the transcriptome, leveraging drug perturbation thus maximizing the information we can gather from a relatively small cohort. This learned lesson encourages inclusion of minority population samples with a relatively small sample size. Many times, non-European samples are removed from large scale multi-omic studies due to limited power, further limiting the transferability of many genomic discoveries.

Drug treatment also reveals novel regulators of important DMEs, such as *CYP3A5* and *UGT1A4*, which do not have any eQTLs identified in the baseline condition or within GTEx liver. In general, we found these eQTLs have increased effect sizes after drug treatments. This first suggests that using eQTLs mapped in the baseline condition may miss important regulatory variation. Second, the increased effect size indicates an enlarged difference in gene expression among individuals with different genotypes after drug exposure. In other words, individuals carrying a certain genotype have greater induced gene expression compared to individuals with the alternative genotype. Given the common occurrence of multiple medications to treat co-morbid conditions, drug exposures may amplify the regulatory effect of some genetic variants. We showed that reQTLs showed higher ORs for overlap with several TFBS, such as GABPA and REST, as compared to non-reQTLs. This suggests one of the potential mechanisms underlying differential gene induction is through recruitment of transcription factor binding. More analysis and types of molecular profiling are needed to examine other potential mechanisms such as histone modification, enhancer activity and DNA methylation (Pai et al., 2015).

In our analysis, we identified a putative regulatory variant of *CYP3A5*. The CYP3A enzymes are thought to metabolize approximate 50% of drugs on the market. The amount of CYP3A5 protein varies in human liver, and the low or absent levels of this enzyme in most European ancestry populations. Because of the overall lack of *CY3A5* expression in European populations, the impact of CYP3A5 has not been comprehensively studied. CYP3A5 is the main metabolizing enzyme of tacrolimus, which is the drug used to suppress the immune system after solid organ transplantation. Because of the drug’s narrow therapeutic index and high between-patient variability, it is important to individualize initial tacrolimus dose based on CYP3A5 genotypes. The current known *CYP3A5* variants, *CYP3A5*3 (*a splice variant which results in a non-functional enzyme) has been shown to explain only 45% of the variability of tacrolimus dose (Birdwell et al., 2015). African Americans require a higher dose of tacrolimus mostly due to the lower frequency of the *CYP3A5*3* allele in African populations. (Liu et al., 2019). We found a novel variant, rs61017966, located within an enhancer region which also overlaps with active histone markers and various TFBS. Adding rs60107966 to the known splice variant explain significantly more variability in *CYP3A5* expression, suggesting that this variant might improve genotype-guided tacrolimus dosing, but may have broader implication as few studies have explored how increase expression of CYP3A5 may impact drug dosing and adverse events. This is especially relevant to African Ancestry populations that express more CYP3A5 enzyme.(Lamba et al., 2002)

Our study has some limitations. First, we did not provide experimental validation of the regulatory effect of identified reQTLs. More functional analysis is needed to confirm the causal effect of rs61017966 within a relevant cellular or animal model. Second, we were restricted to a limited sample size of hepatocytes due to the rarity of livers from African American donors. It should be noted that this study quadrupled the amount of matched genotype and gene expression data available in African ancestry individuals as compared to large databases such as GTEx (15AA in GTEx). Regulatory variants of other important DMEs might be revealed with larger sample size. Finally, we used cultured primary hepatocytes. This *in vitro* model may not represent gene expression of hepatocytes *in vivo*.

In summary, our study comprehensively characterized the biological response to drug treatment at transcriptional and genetic regulatory levels across different enzyme inducers in African Americans. Leveraging drug treatments provides an advantage to identify novel regulatory variants of important DMEs. We uncovered putative regulatory variants that might improve drug dosing with population differentiated response. Our work enhances the understanding of genetic regulation of gene expression and contributes to increasing equality in personalized genomic medicine.

## Methods

### Drug treatment of primary hepatocyte cultures

The procuration of 62 hepatocyte cultures was described in Zhong *et al* (Zhong et al., 2020). Hepatocyte suspensions (0.6 x 10^6 cells in 0.5 mL of CP medium) were added to collagen-coated wells of a 24-well plate followed by incubation at 37° C for 4-5 hours to allow cells to adhere. The CP medium was replaced with 0.5 mL of 0.25 mg/mL matrigel solution in HI medium, and plates were incubated overnight at 37° C. The following day, matrigel-medium was replaced with 0.5 mL of induction drug in HI medium with antibiotics and the step was repeated for the next two consecutive days after every 24 hours. We performed the induction experiments using the following six drugs: Omeprazole (50 μM), Phenobarbital sodium (1 mM), Dexamethasone (50 μM), Carbamazepine (50 μM), Phenytoin (75 μM), Rifampicin (25 μM). Baseline hepatocytes were treated with DMSO.

### Differential gene expression (DGE) analysis

Gene expression and genotyping data were prepared as described in Zhong *et al* (Zhong et al., 2020). Here we used expression data for baseline and six unique drug treatments. We calculated transcript per million (TPM) by first normalizing the counts by gene length (HGCR38) and then normalizing by read depth (Wagner et al., 2012). Samples were clustered through PCA to determine outliers and 2 samples were removed after this QC step. Gene expression values were filtered to retain genes with > 0.1 TPM in at least 10% of samples and ≥ 6 read counts in at least 10% of samples across seven conditions. We retained only protein coding genes and long non-coding RNAs, resulting in a total of 15,579 genes that were used in the subsequent analyses.

We used weighted trimmed mean of M-values algorithm (TMM) implemented in edgeR (Robinson et al., 2010) to estimate the RNA sequencing library size. Then we calculated the counts per million (CPM) and performed log transformation with voom function implemented in Limma which accounts for estimated library sizes (Smyth et al., 2011). The estimated mean-variance trend was incorporated as precision weights in the model fitting by lmFit in Limma. To identify differentially expressed (DE) genes, we used a paired sample design to compare the expression before and after drug treatments for each individual. We used the empirical Bayes method implemented in Limma to test for DE between all pairs of conditions, with significance prespecified as by FDR<0.01 and log fold change<=0.5.

### Gene set enrichment analysis

Enrichment of DE genes in biological process ontologies was performed using clusterProfiler (Yu et al., 2012) using all expressed genes in all conditions (N = 15,579) as background. We used the FDR method for multiple test correction. We used gProfileR (Raudvere et al., 2019) to test whether regions surrounding the DE genes are enriched for specific TF binding.

### Variance partition analysis

We combined all 7 gene expression datasets and utilized a linear mixed model to partition the variation in gene expression to various contributing factors using the normalized gene expression data after voom transformation. We modeled the effects of the condition, individual, sex, and potential confounding factors (batch and sequencing platform) as random effects and age and ancestry as fixed effects. This analysis was performed using the VariancePartition R package (Hoffman and Schadt, 2016).

### Cross-treatment eQTL analysis

We used Meta-Tissue developed by Sul *et al*. to jointly estimate eQTLs across conditions by integrating the mixed model and meta-analysis (Sul et al., 2013). Because of our experimental design, expression data was measured for the same hepatocyte culture under different conditions. Therefore, expression measurements are correlated across conditions which violates the assumption of independence of studies in meta-analysis. Meta-Tissue removes this correlation between estimates of effect size before meta-analysis. First, Meta-Tissue estimates the effect size of each SNP in each condition using a linear mixed method to account for the sample sharing.

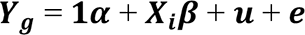

Here ***Y***_***g***_ is the expression vector of dimension NT×1 where N is the number of samples and T is the number of conditions, ***α*** ∈ *R^T^* is the intercept vector, ***X***_***i***_ is the genotype for variant *i*, ***β*** ∈ *R^T^* is the coefficient vector, 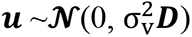 is the random effect of dimension NT×NT where D corresponds to the sample sharing and 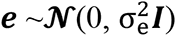 is the random error of dimension NT×NT. Meta-Tissue uses the generalized least square to estimate ***β*** and var(***β***). Next, the Ω = var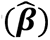 is replaced by *var_new_* 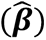 = diag(Ω^-1^e)^-1^ to “uncorrelated” the 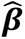 where e is a vector of ones. Third, the “uncorrelated” 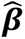 are used in the regular meta-analysis.

Meta-Tissue uses Metasoft (Han and Eskin, 2011) for meta-analysis which can perform with both fixed effect (FE) and random effects (RE) models. Fixed effect model assumes that there is one true effect size that underlies all the studies in the analysis. Random effect model assumes the effect sizes can vary by studies. We performed both analysis since we hypothesized eQTLs may have either fixed or random effect. Metasoft provides betas and standard error estimates. Meta-Tissue estimates the heterogeneity with I^2^, where I^2^=100%×(Q-df)/Q where Q is the chi-squared statistics and df is the degrees of freedom. Finally, Meta-Tissue computes an m-value which estimates the posterior probability of an eQTL being present in each condition by taking information across conditions to explicitly reveal the sharing pattern of eQTLs.

For each treatment condition, we filtered gene expression to retain genes with > 0.1 TPM in at least 20% of samples and ≥ 6 reads in at least 20% of samples. We normalized expression values across samples by TMM implemented in edgeR. After TMM normalization, we performed the inverse normal transformation for each gene expression profile. We estimated probabilistic estimation of expression residuals (PEER) variables on normalized expression in order to remove any potential confounding factors. (Stegle et al., 2012)

We performed eQTL mapping on 60 hepatocyte cultures that passed both genotype and RNA- sequenicing QCs. The genotype data was processed as described in Zhong et al. with 7,180,502 SNPs in total.(Zhong et al., 2020) There were 415 expression profiles across all conditions including baseline. Five gene expression profiles were not included in the analysis due to insufficient hepatocytes to conduct all drug treatments or inadequate/poor quality mRNA to conduct sequencing. We used 17,949 genes that were expressed in any conditions. We first regressed the top 10 PEER variables from the gene expression profiles leaving the expression residual. The number of PEER variables was chosen to optimize the number of detected eGenes in the baseline condition as described in Zhong et al.(Zhong et al., 2020) We then used Meta-Tissue linear mixed model to obtain the estimates of effects and standard error for each gene residual-cis SNP pair (restricted to SNP with 1Mb for the gene start position). We included additional covariates: sex, platform, batch and the first principal component calculated from genotype data to correct for potential known confounding effects following Zhong et al.

### Treatment-by-treatment eQTL analysis

We also preformed treatment-by-treatment (TBT) analysis - which was conducted to find eQTLs in each condition separately. The p values were calculated from fitting multivariate linear regression of gene residual ∼ genotype + sex + platform + batch + PC1. We used a hierarchical correction method to adjust the nominal p values and to call the eGenes and eQTLs as described in Zhong *et al*. Firstly, p values of all cis SNPs are adjusted for multiple testing for each gene using Benjamini and Yekutieli (BY) method as the locally adjusted p values. Secondly, the minimum BY-adjusted p values for all genes are corrected using Benjamini and Hochberg method (BH) as the globally adjusted p values (BY-BH p values). Lastly, for a chosen threshold (here we use, 0.05), we found the largest BY-BH p values under that threshold and the corresponding BY-adjusted p value. This BY-adjusted p-value is used as the threshold to call significant eQTLs. These results were compared to those found in the Cross-treatment eQTL analysis.

### Identification of response QTLs (reQTLs)

We used a beta-comparison approach to identify reQTLs which are eQTLs with altered effect size after drug treatment. We used the beta estimates (***β***), standard deviation of beta estimates (var(***β***)) and correlation of beta estimates (var(***β***)) that were computed in the Meta-Tissue linear mixed model to compare the beta estimates in drug treatment conditions with the baseline condition. We compared the beta estimates in two conditions using a z test similar to Kim-Hellmuth et al. Here beta2, sd2 are the estimates with drug treatment; beta1, sd1 are the estimates in the baseline condition; and cor12 is the correlation of beta1 and beta2 estimates from var(***β***) of Meta-Tissue. We then calculated the Z statistics as:

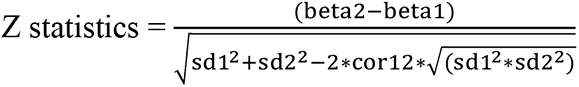

We calculated the Z statistics for all eQTLs mapped in the Meta-Tissue fix effect model, random effect model and TBT analysis. To identify significant reQTLs, we randomly permute the expression across different conditions for each individual. This method maintains the relationship between genotype and expression. However, the effect size of the genotype is now randomly assigned with respect to each condition. We then compute the null z statistics after each permutation. We repeat the permutation 100 times and for each eQTL association test and calculate the empirical p value based on the null z statistics distribution using the qvalue R package (Dabney et al., 2010). We then calculate the q value across all empirical p values and used q value < 0.05 as the cutoff to select significant reQTLs. We performed the multiple-testing correction separately for each drug treatment condition.

### Mapping (log fold change QTL) lfc_eQTL

Here we used the log(CPM) values calculated with limma voom. We first adjusted the effects of batch and the sequencing platform in the data. We estimated any remaining confounding factors with SVA and regressed out these surrogate variables from the log(CPM) values.(Leek et al., 2012) We then calculated the difference of log(CPM) between drug each treatment and baseline representing the log fold-change (lfc) of gene expression and performed inverse normal transformation. We used normalized lfc as the gene expression phenotype in eQTL analysis. We used Matrix eQTL to map lfc_eQTL after adjusting for the first genotype PCs and sex as covariates. We used hierarchical multiple test correction as described previously in the TBT eQTL analysis.

### Enrichment of eQTLs in transcription factor binding sites (TFBS)

Using ENCODE transcription factor ChIP-Seq data from human liver tissue, we identified the number of eQTLs that overlap with the ChIP-Seq narrow peaks (representing transcription factor binding). We randomly sampled 1000 sets of null SNPs that were matched for LD score, minor allele frequency and distance to the nearest gene for each of the eQTL. We then calculated the average overlap between these null SNPs and ChIP-Seq narrow peaks across 1000 different null sets. We used the Fisher exact test to compute the odds ratio and p-value comparing the overlap of eQTLs with the overlap of null SNPs.

### Fixation index (Fst) analysis

We calculated the Fst for eQTLs using 1000 Genomes Yoruba people of Ibadan, Nigeria (YRI) and Utah Residents (CEPH) with Northern and Western European Ancestry (CEU) populations with the GCTA software (Yang et al., 2011). Fst measures the population differences in of SNP allele frequency, with higher values denoting large differences in frequency. We compared the Fst of reQTLs to the Fst of non-reQTLs with Wilcoxon rank sum test.

## Acknowledgements

We would like to thank Drs. Alfred L. George, Jr, Eric Gamazon and Barbara Stranger for many useful discussions and comments. This work was supported by the National Institute of Health (NIH)/National Institute on Minority Health and Health Disparities (NIMHD) grant R01 MD009217 and U54 MD010723. Competing Interest There are no competing interest for the any author.

## Author Contributions

M.A.P. and Y.Z. conceived this study and wrote the paper. Y.Z. performed the analysis. T.D., M.A.P, J.A. and C.A. curated the data.

---

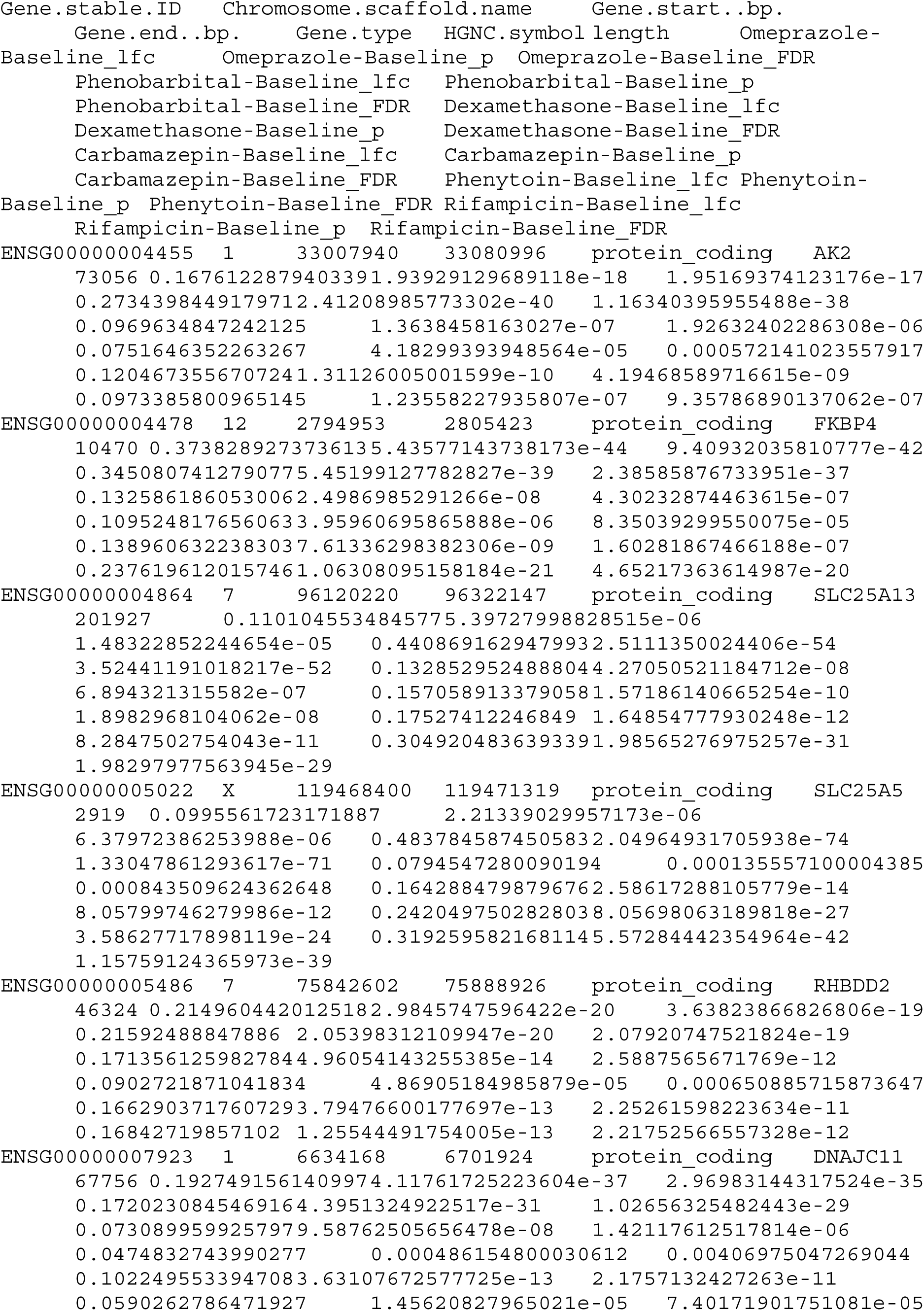

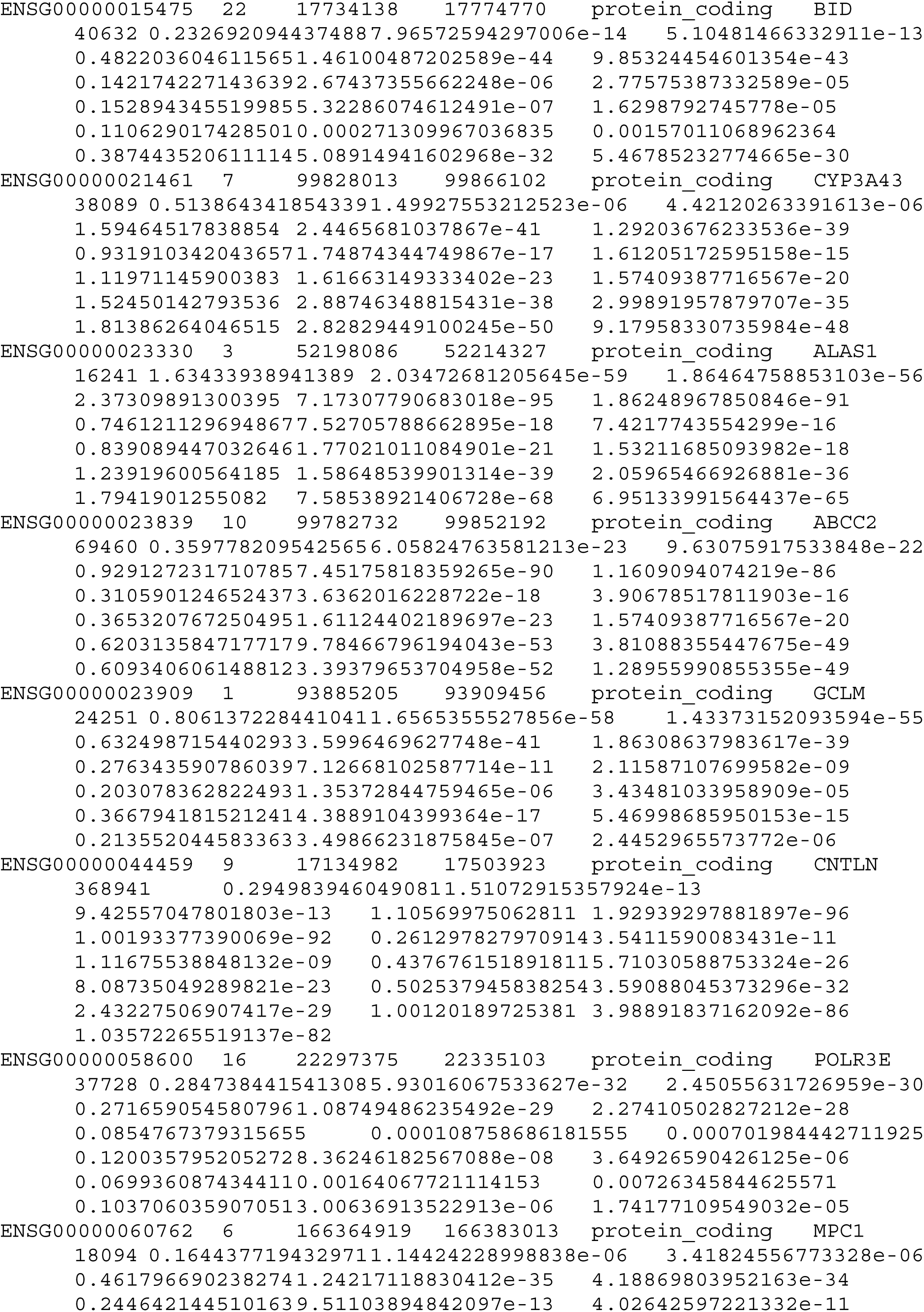

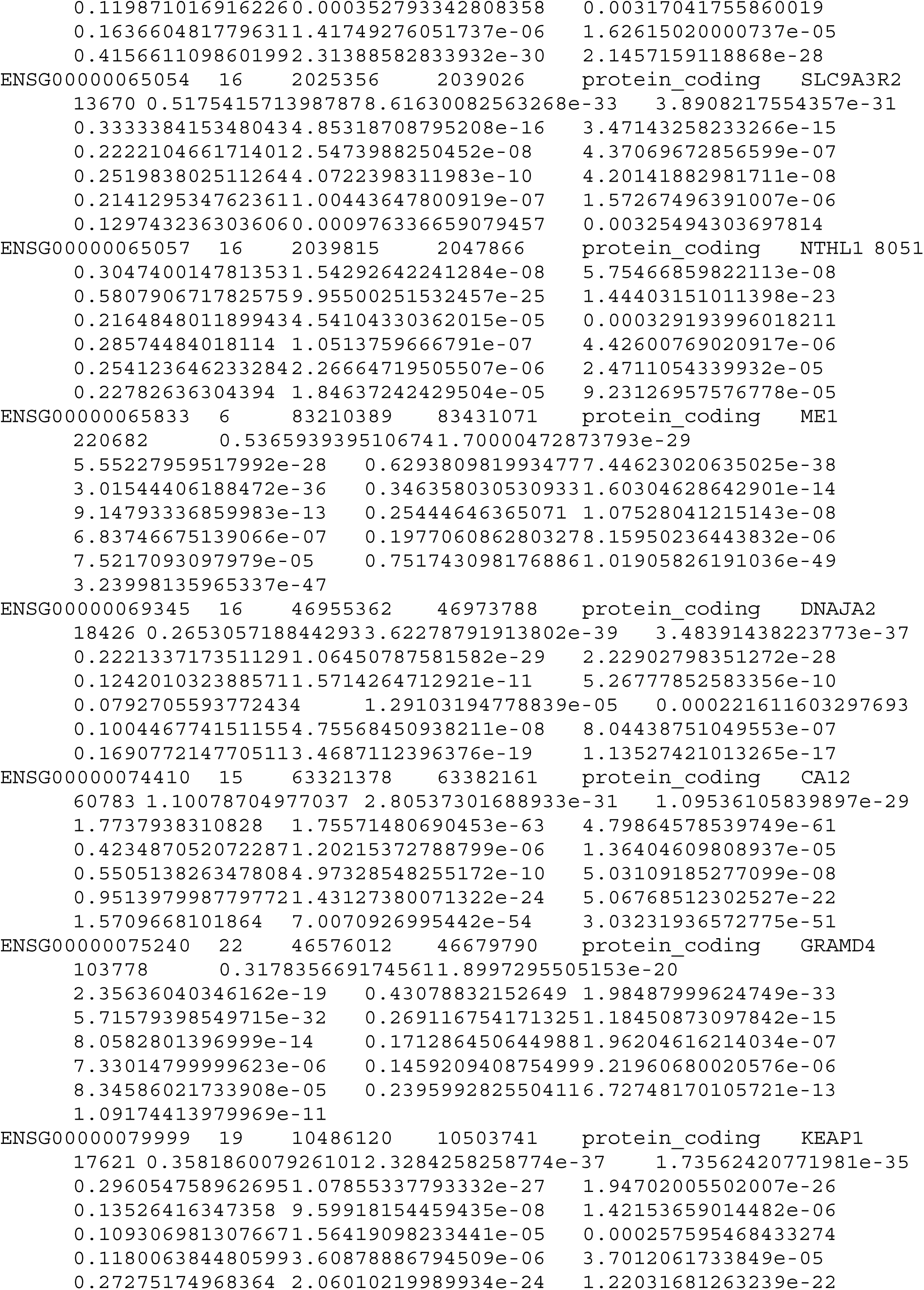

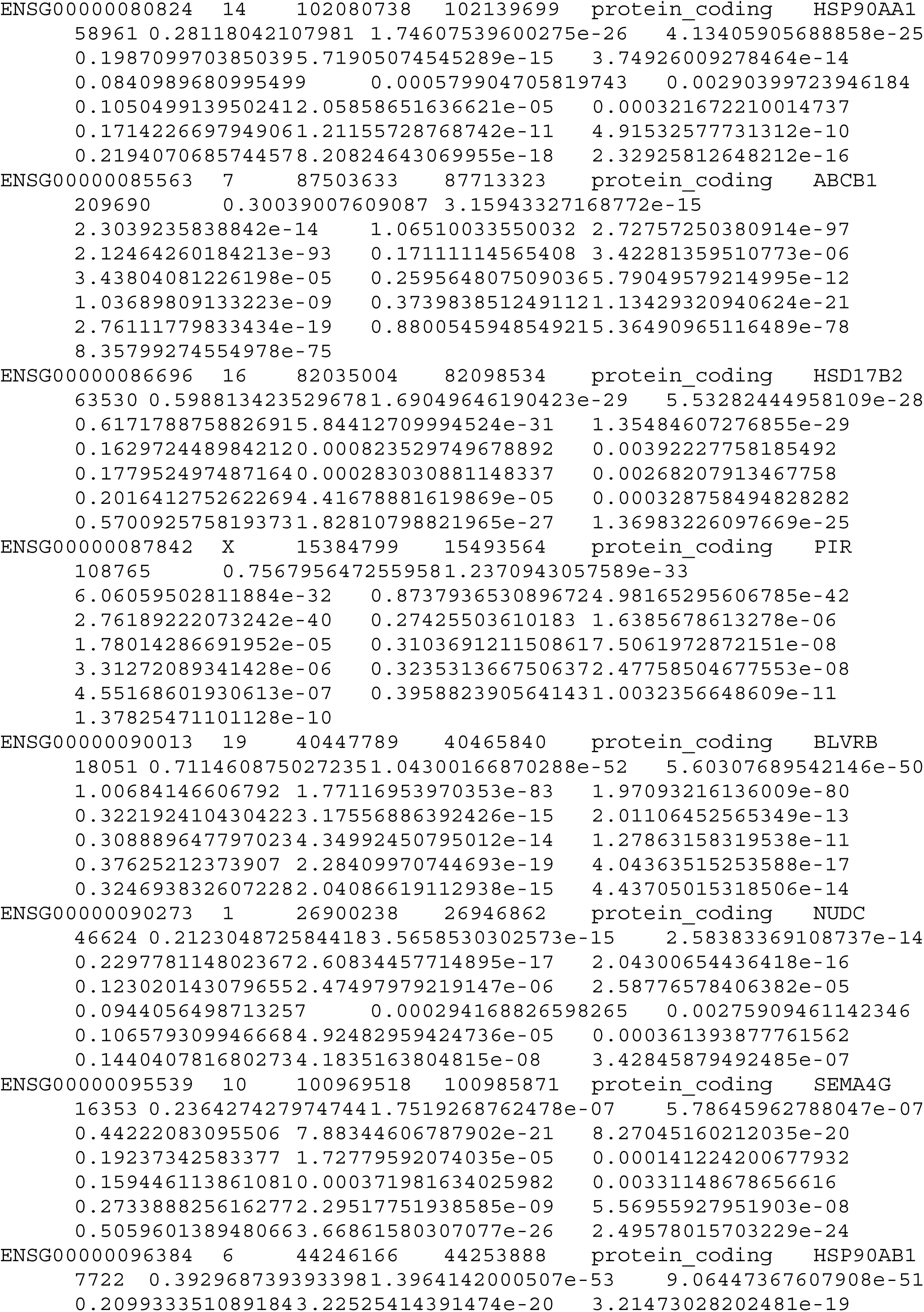

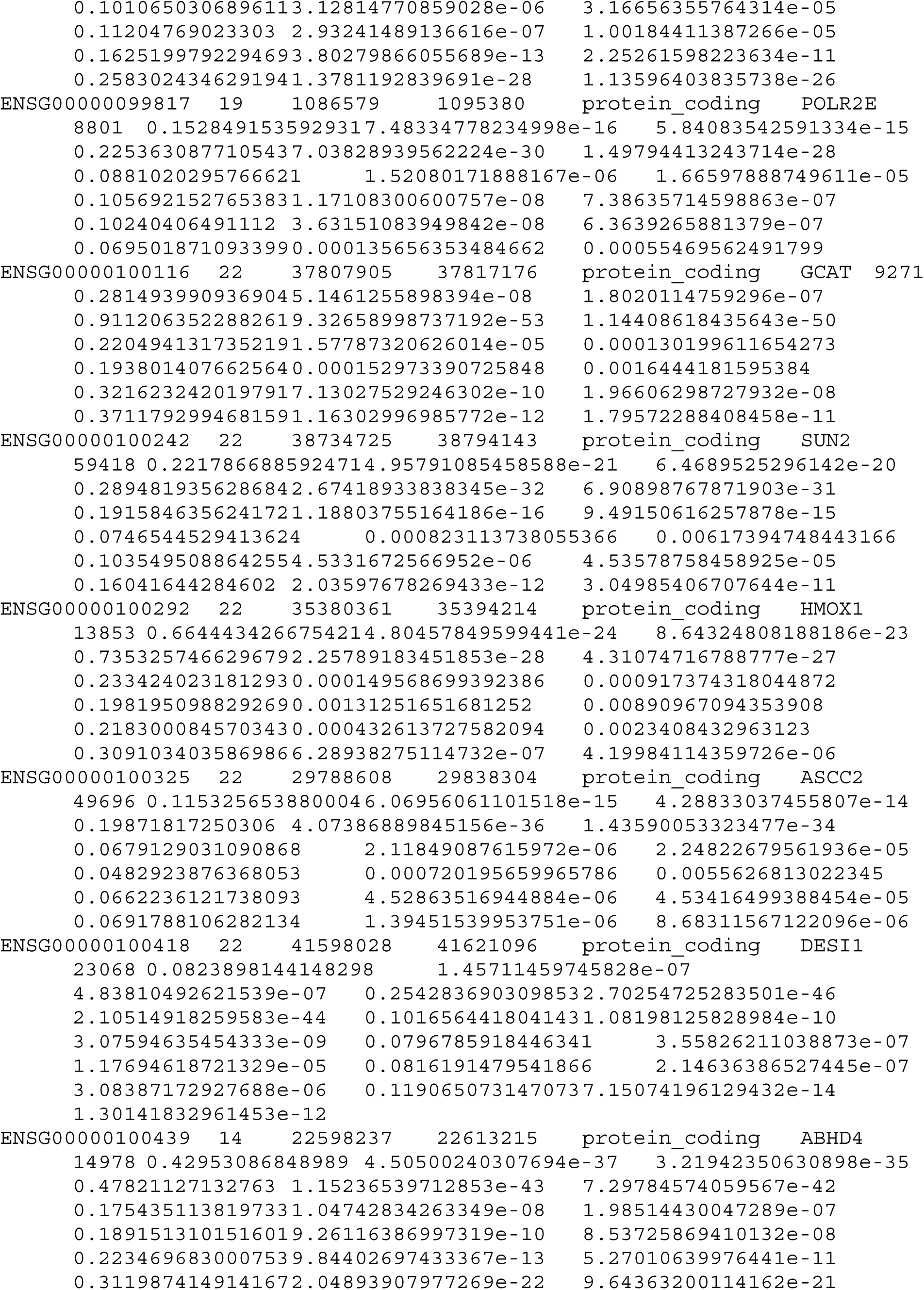

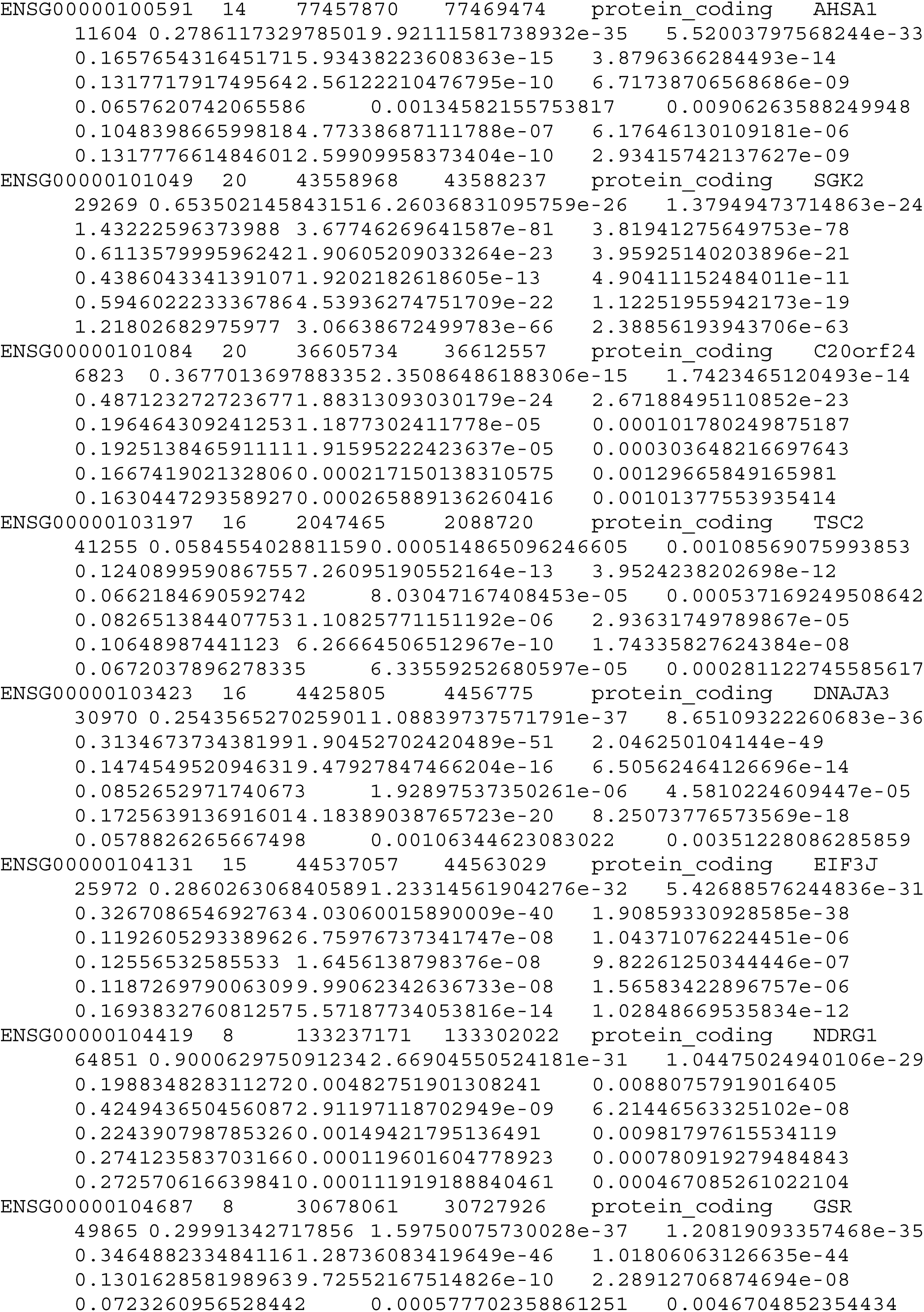

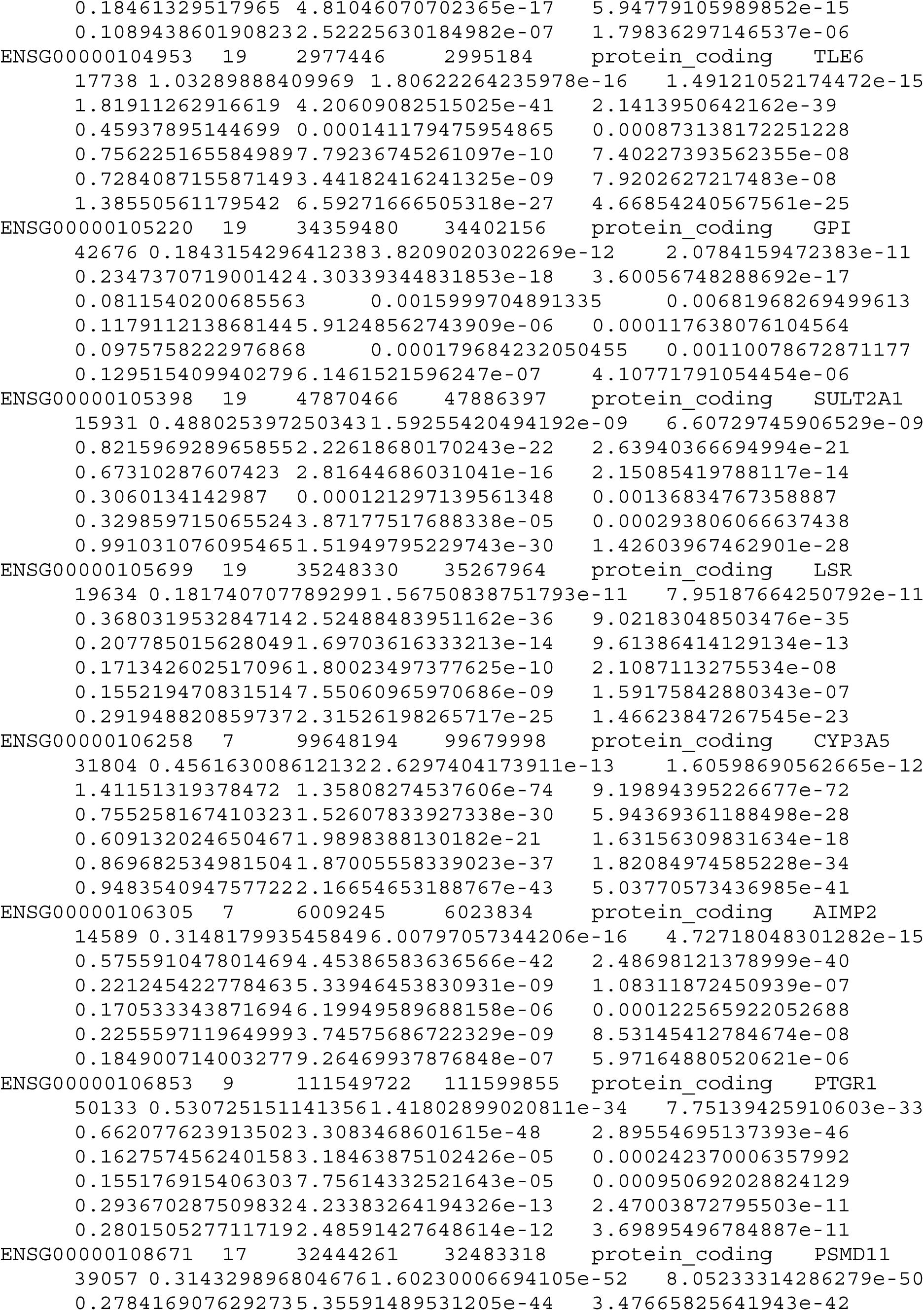

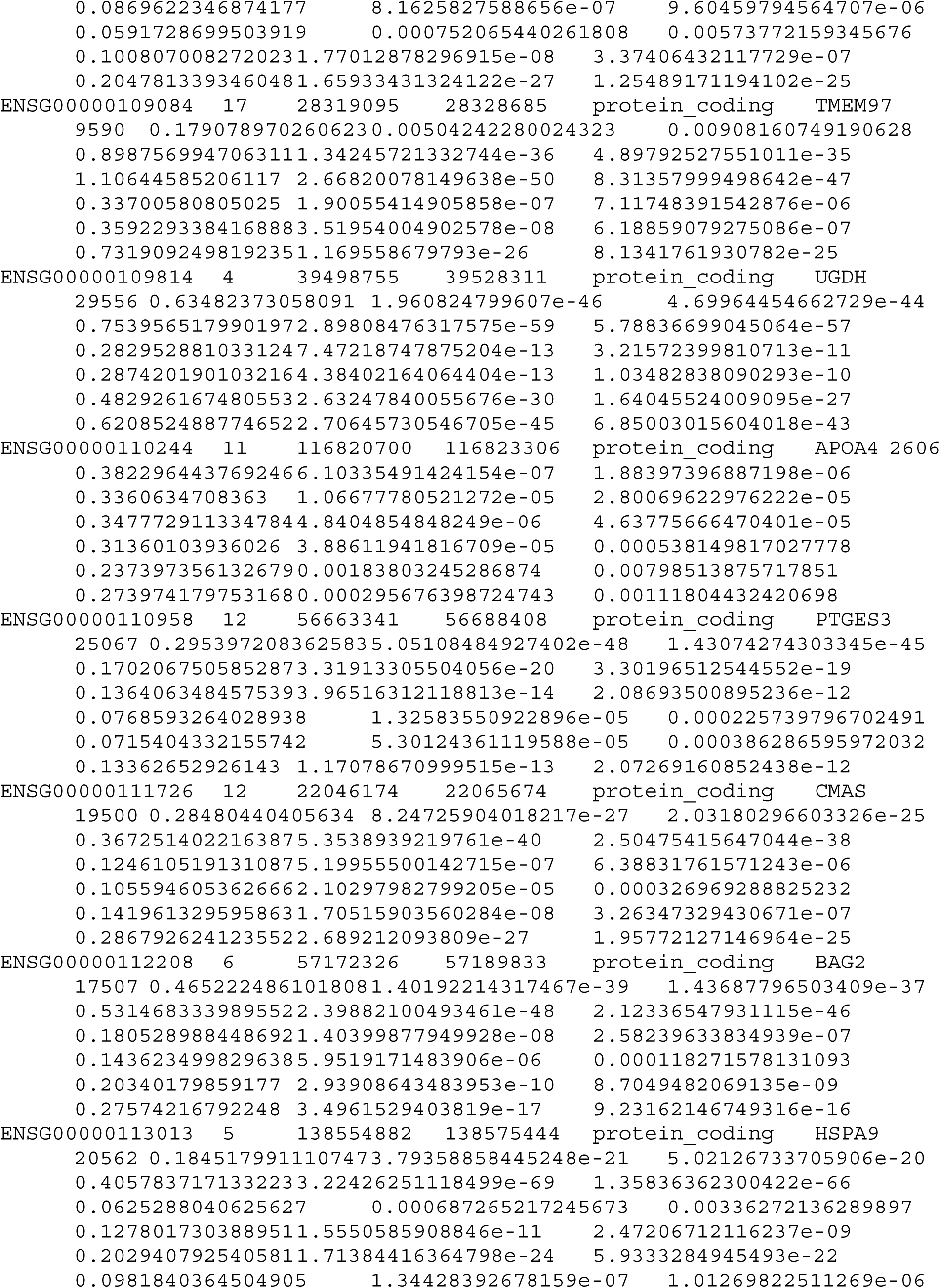

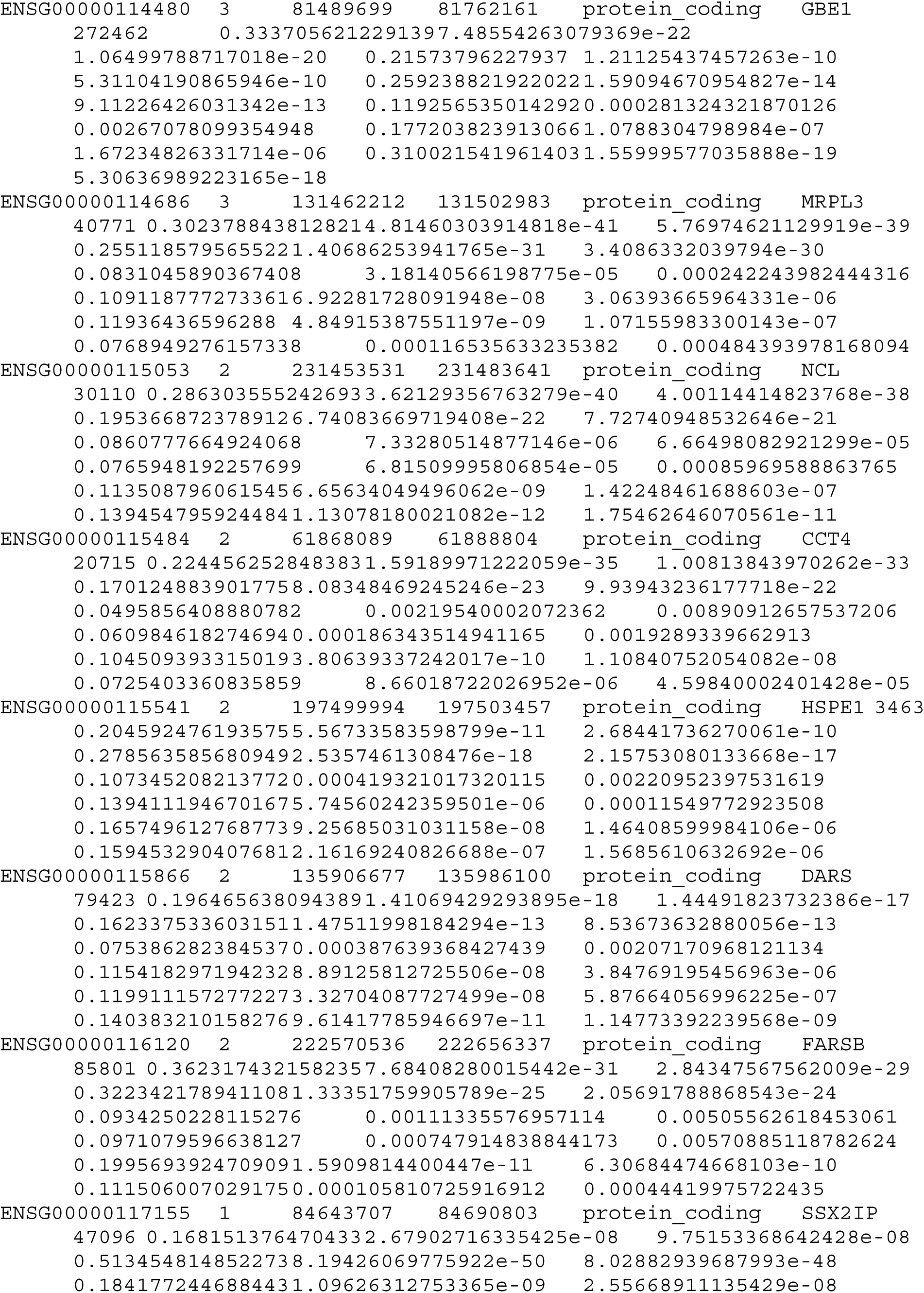

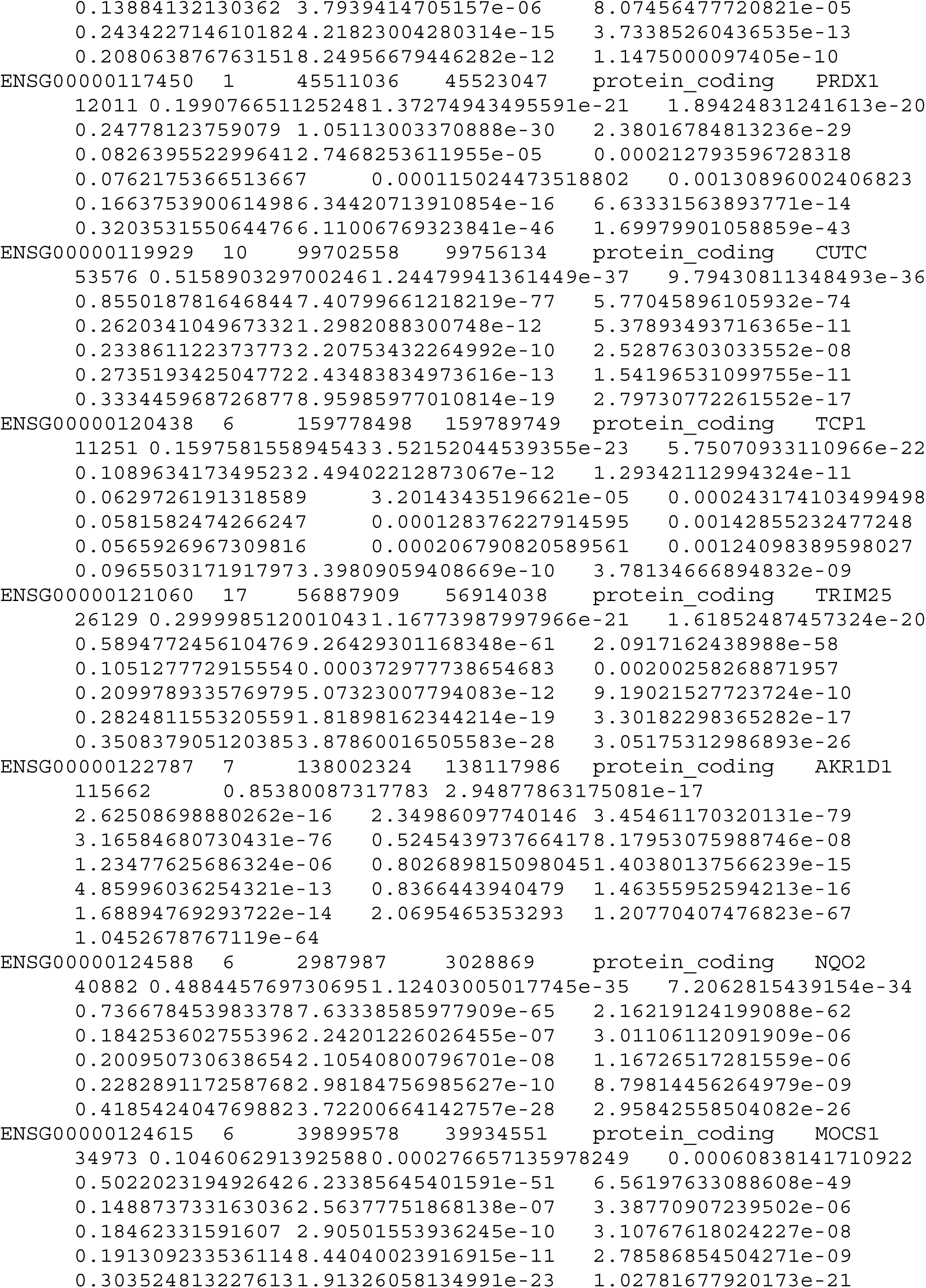

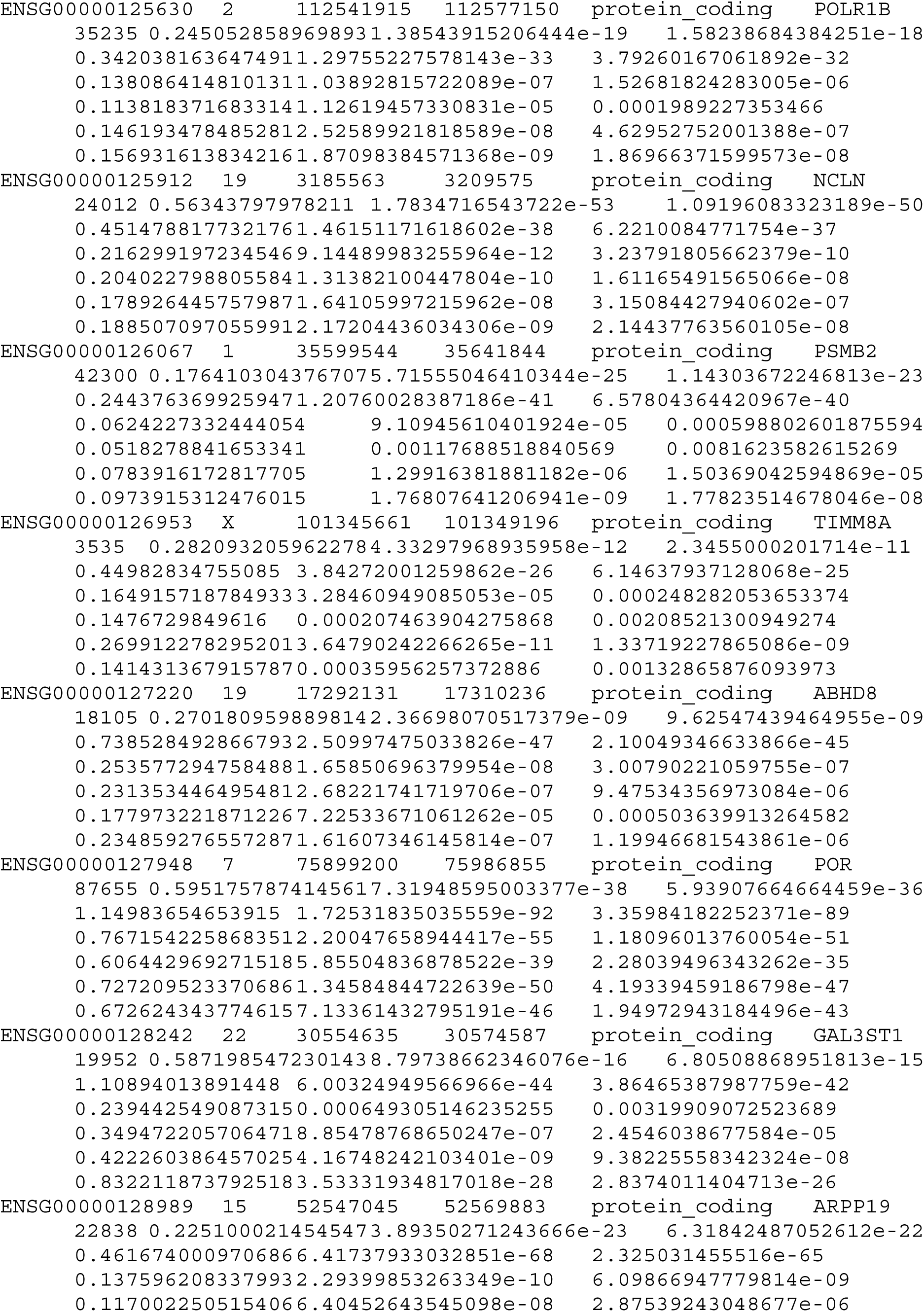

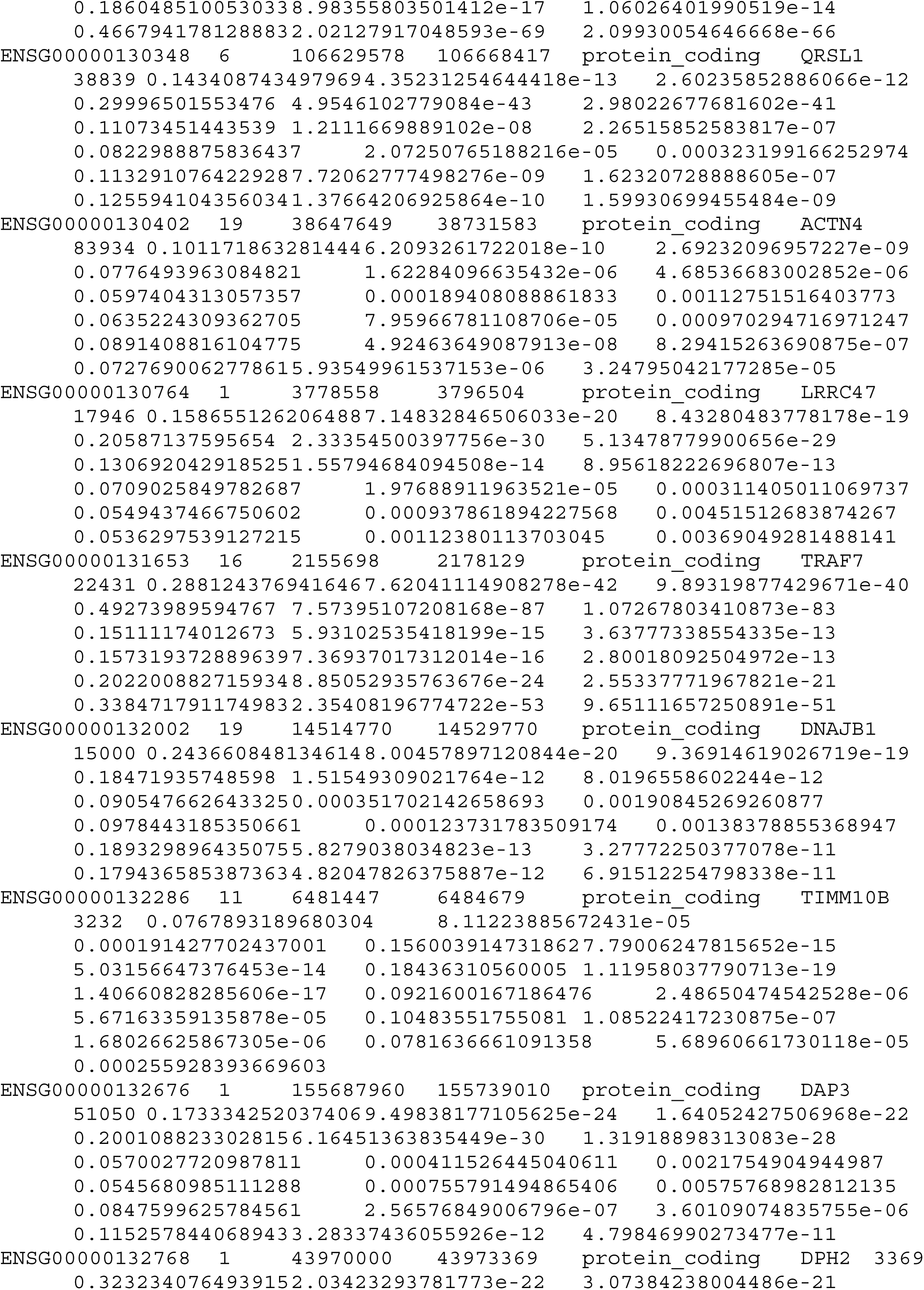

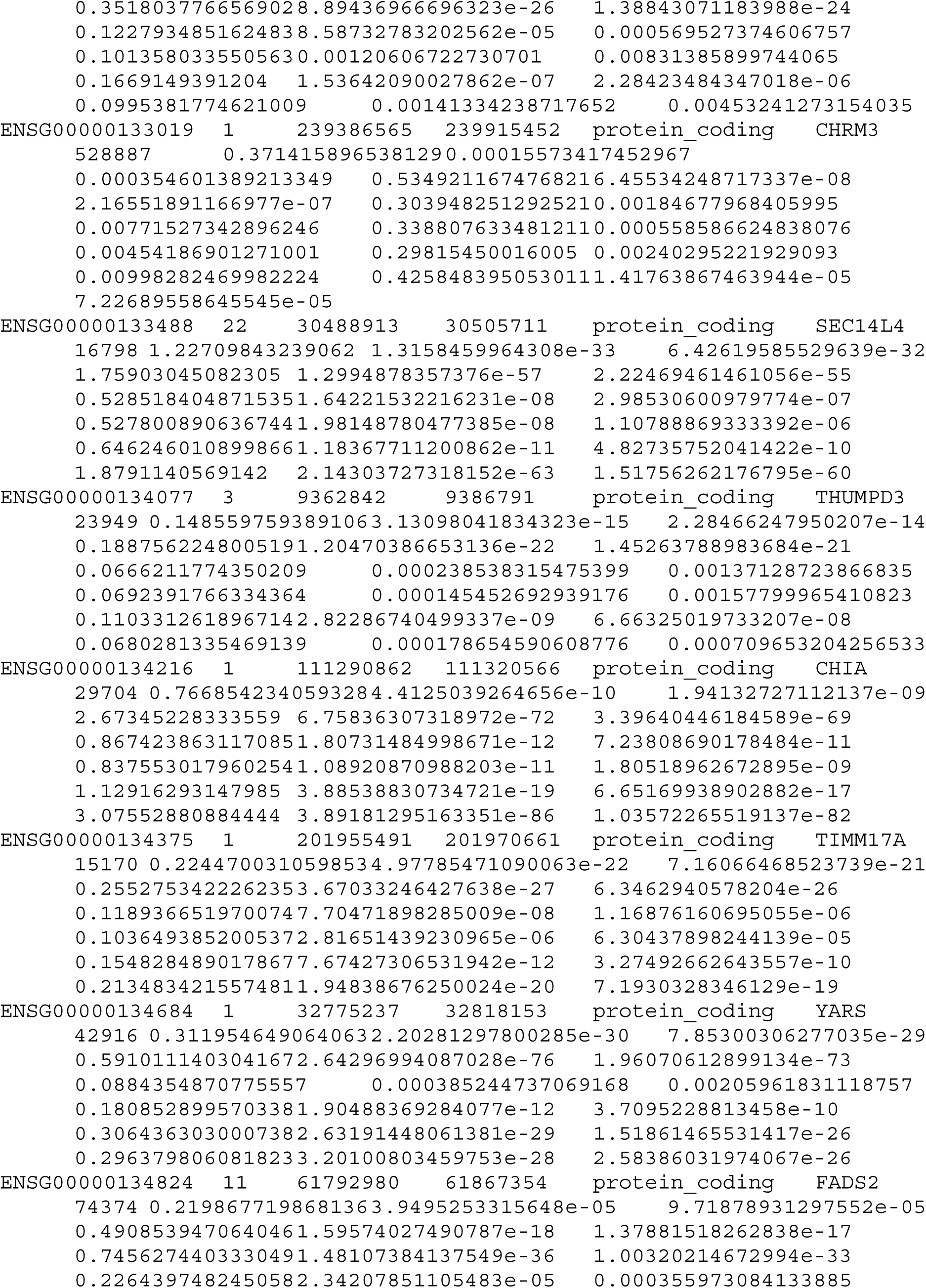

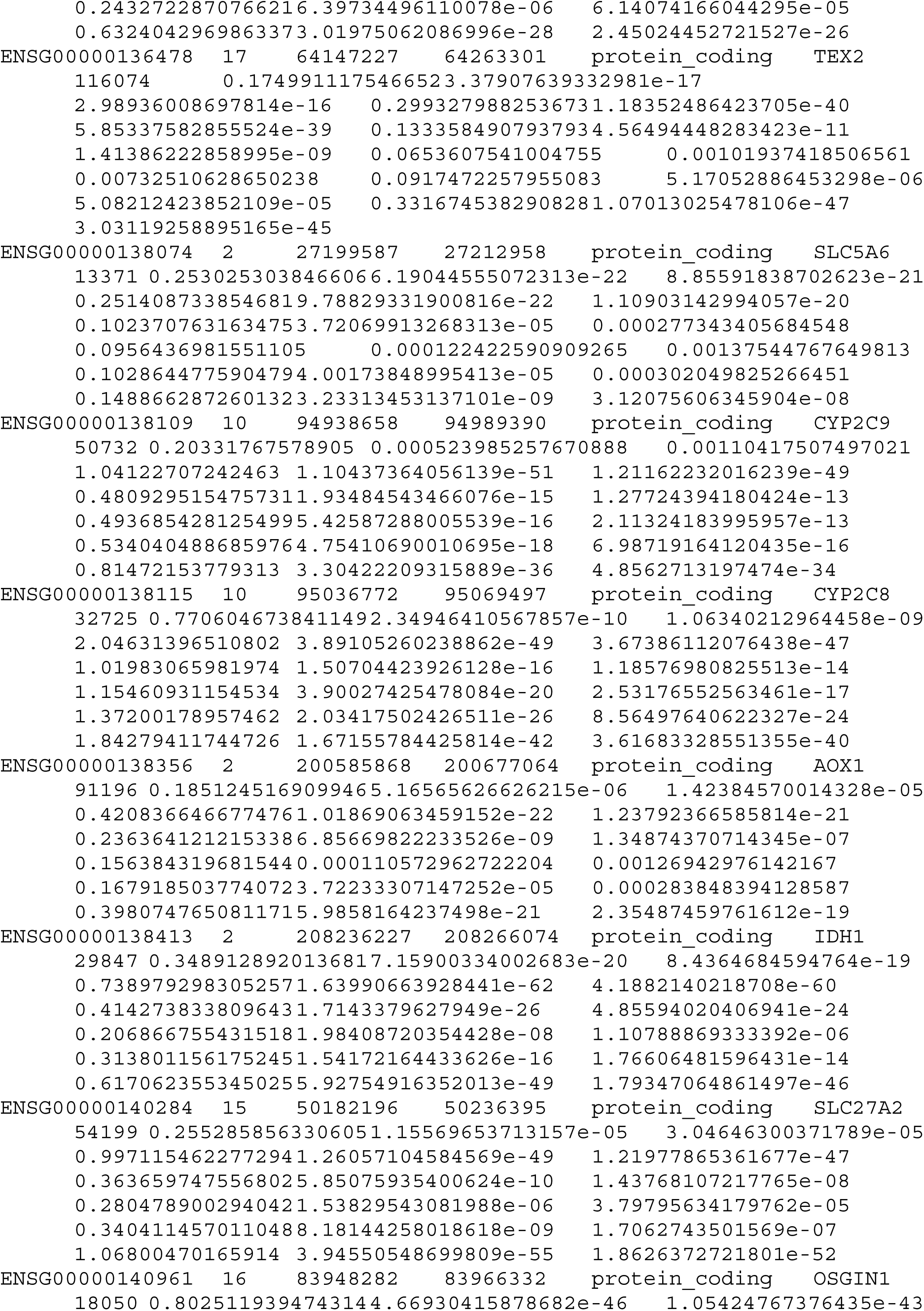

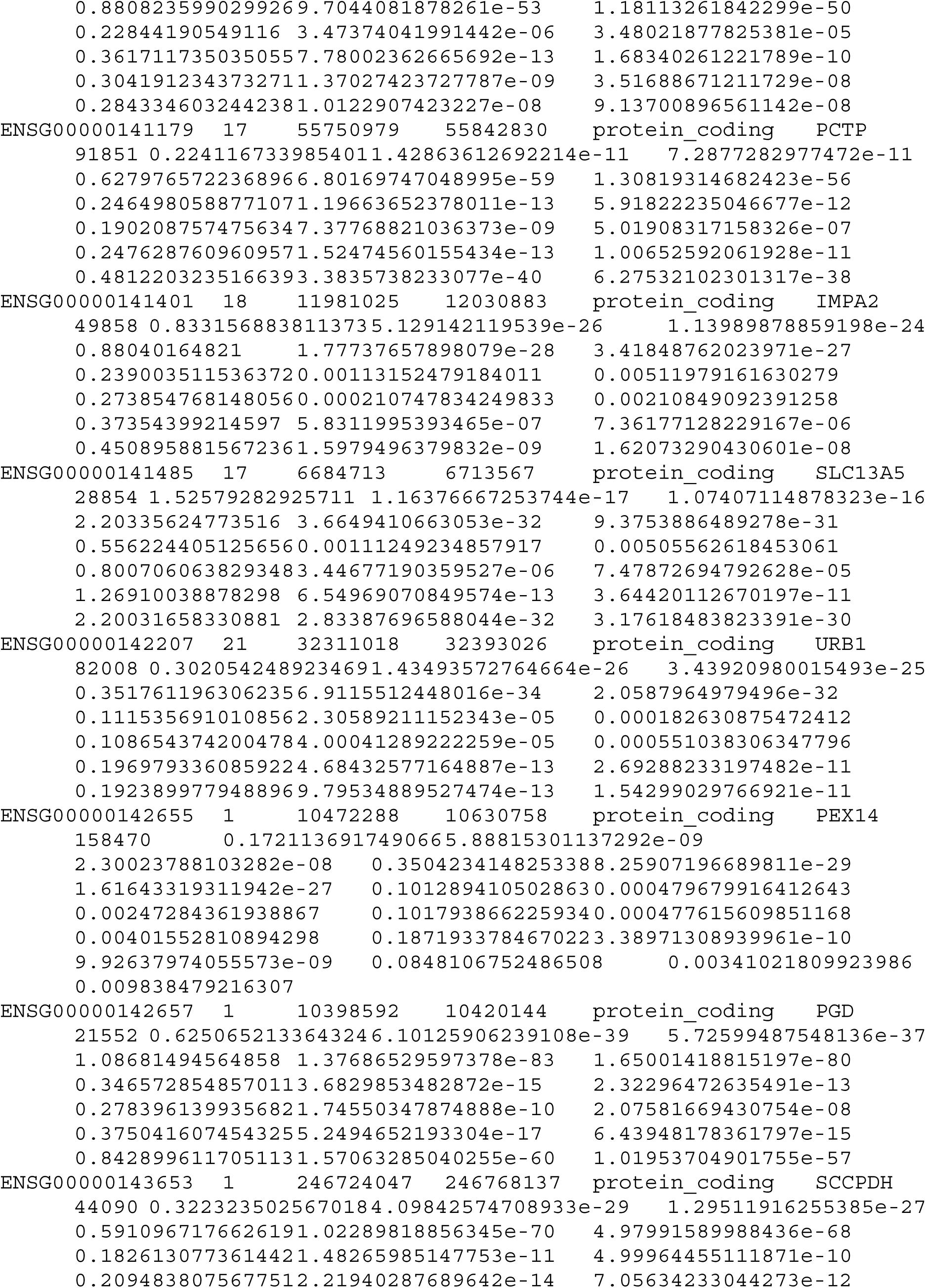

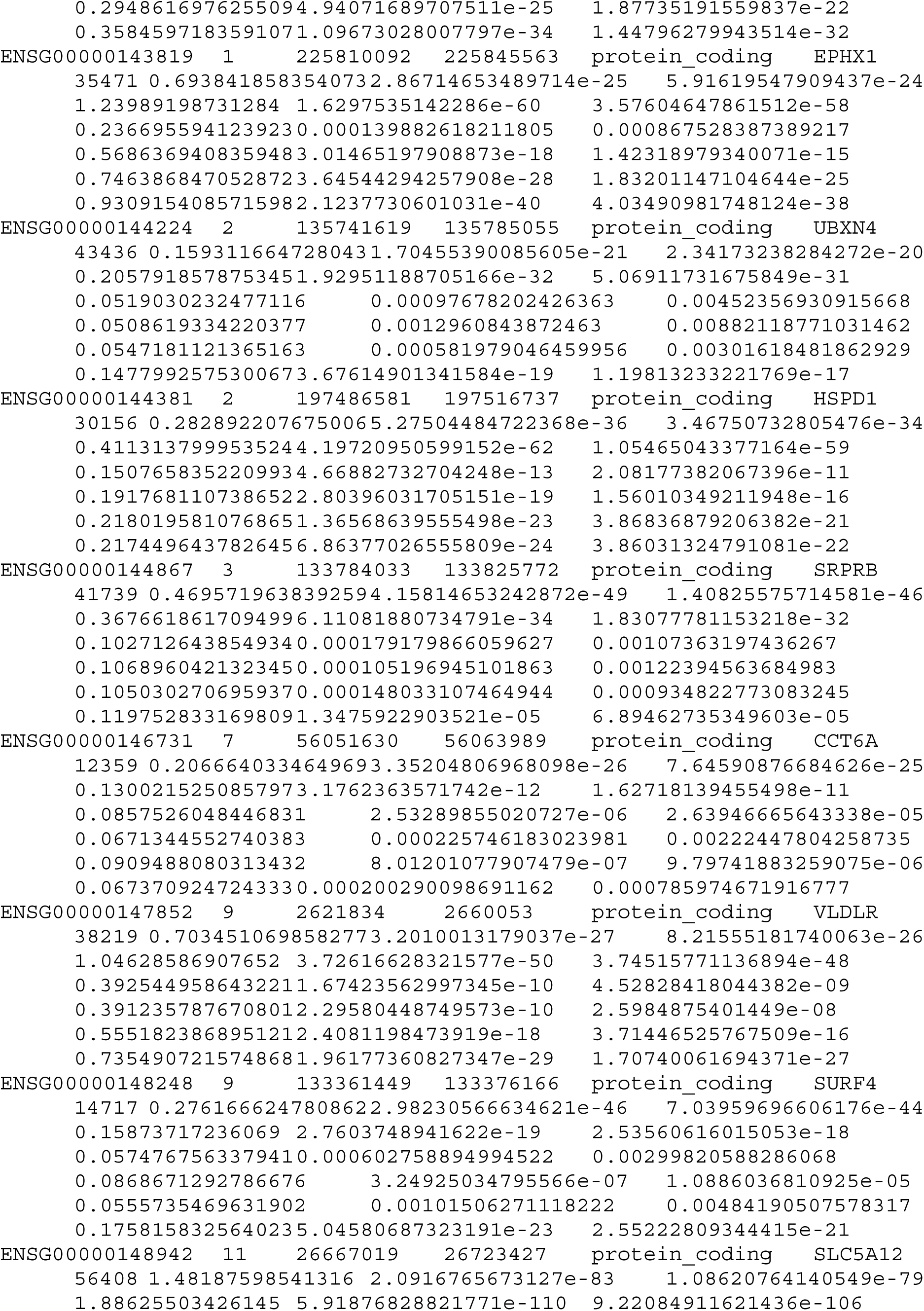

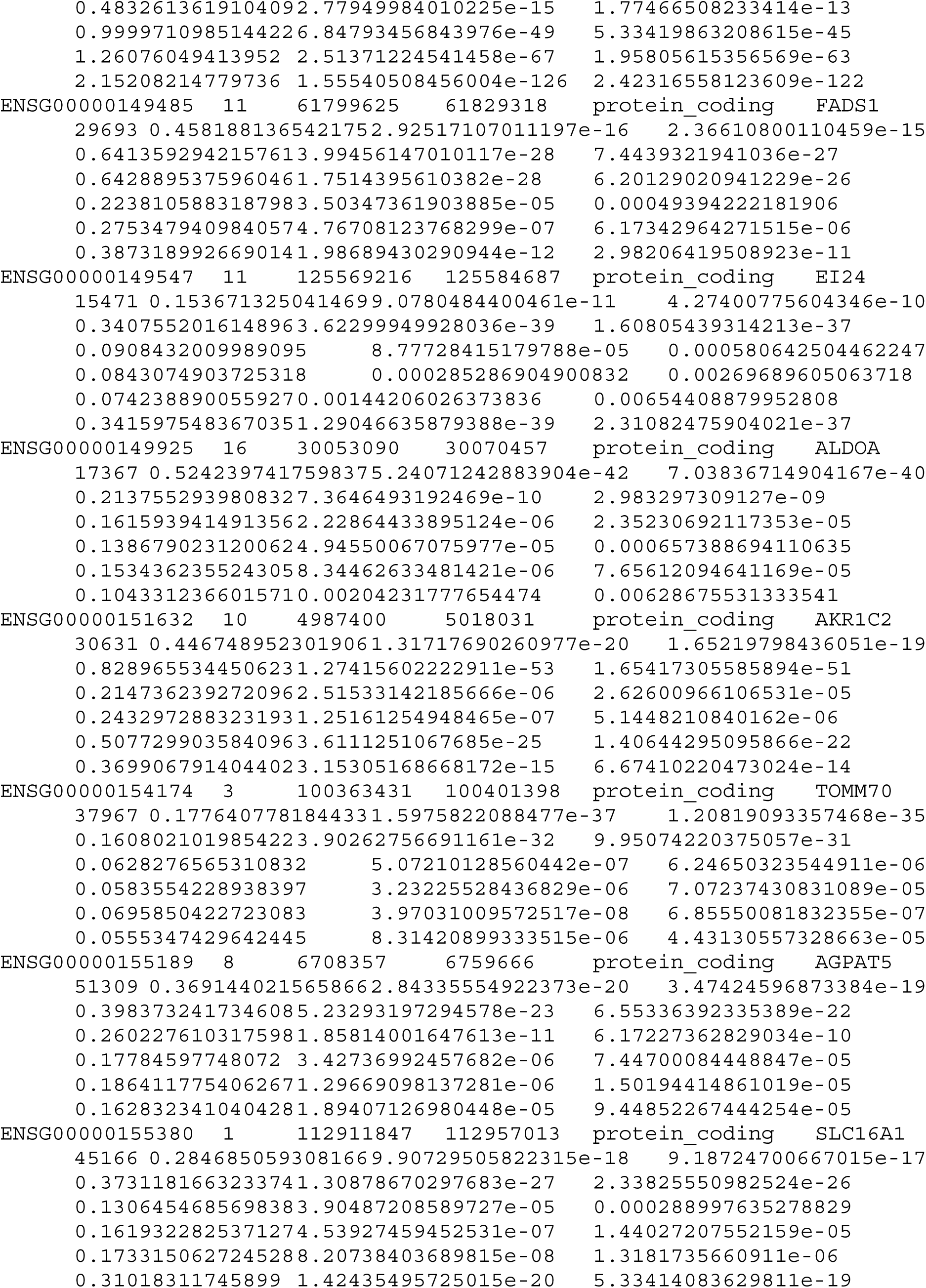

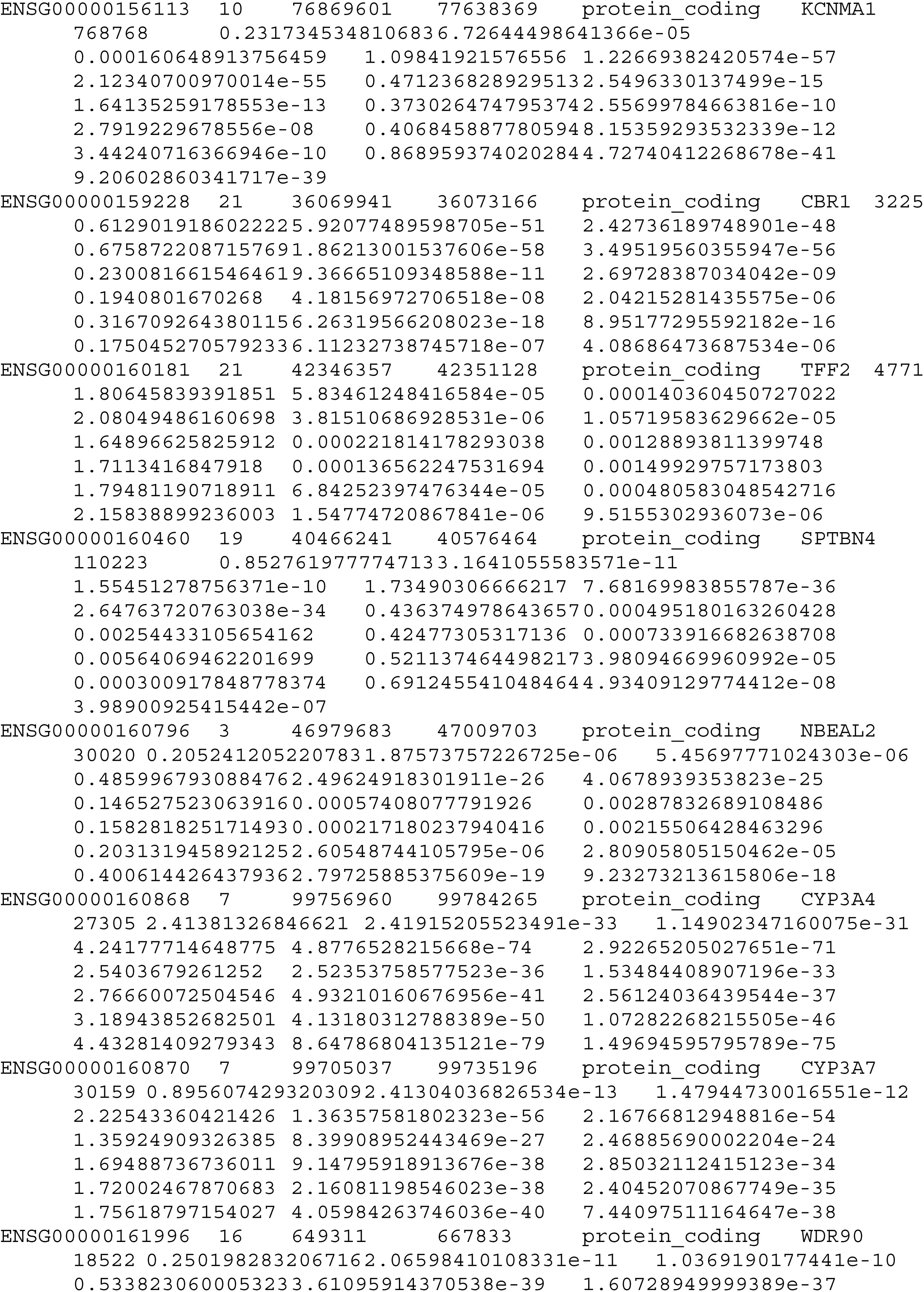

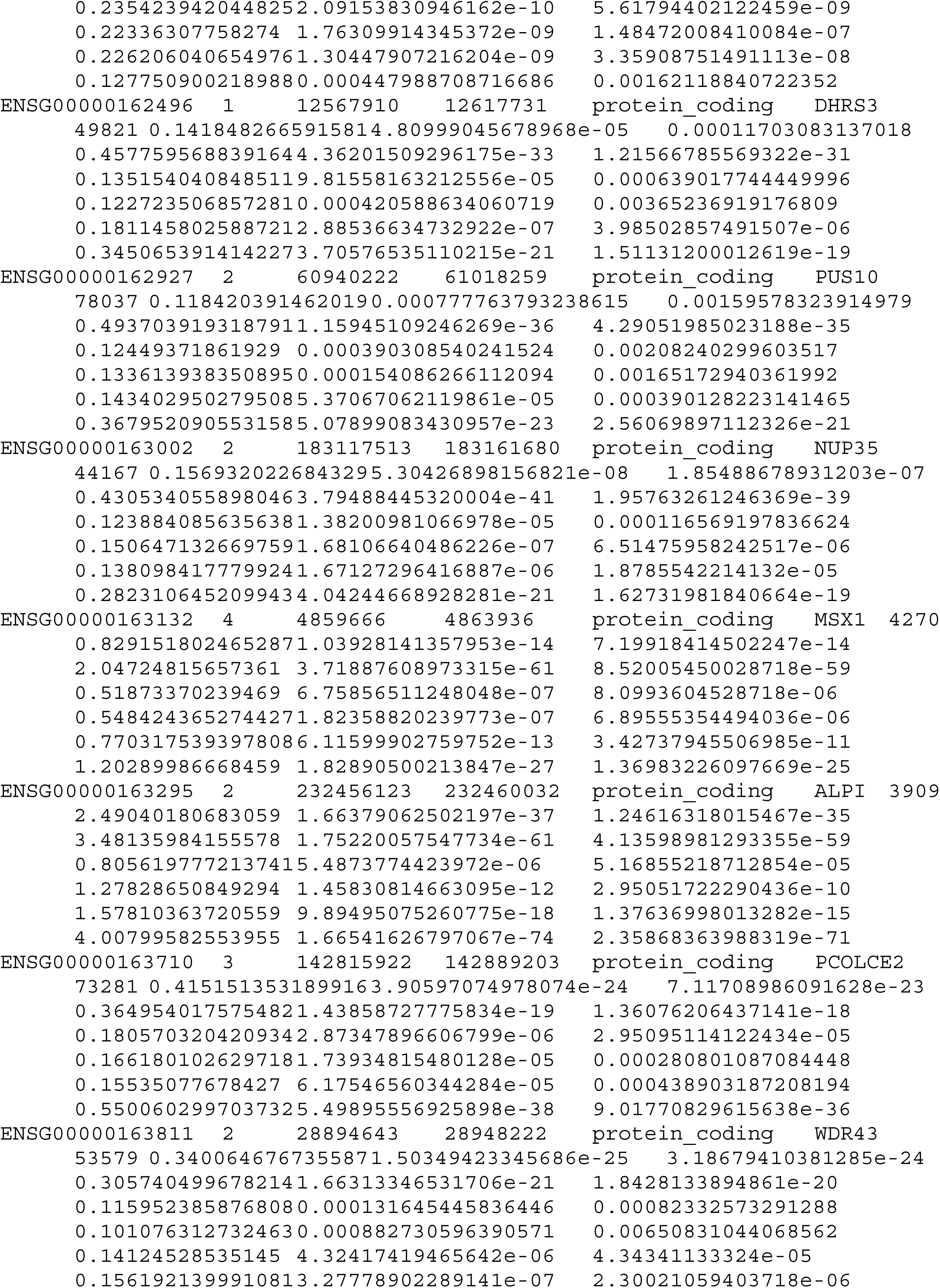

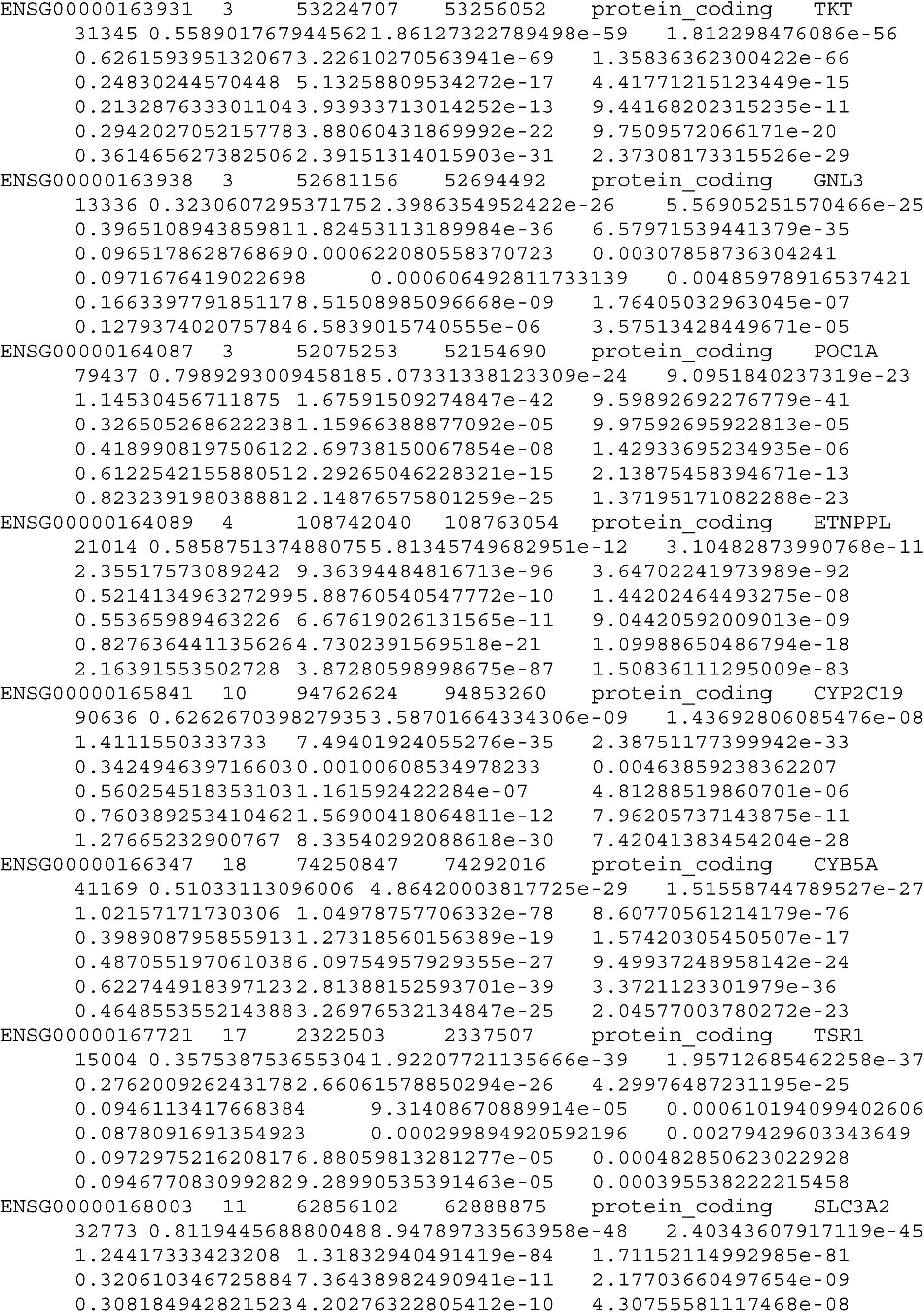

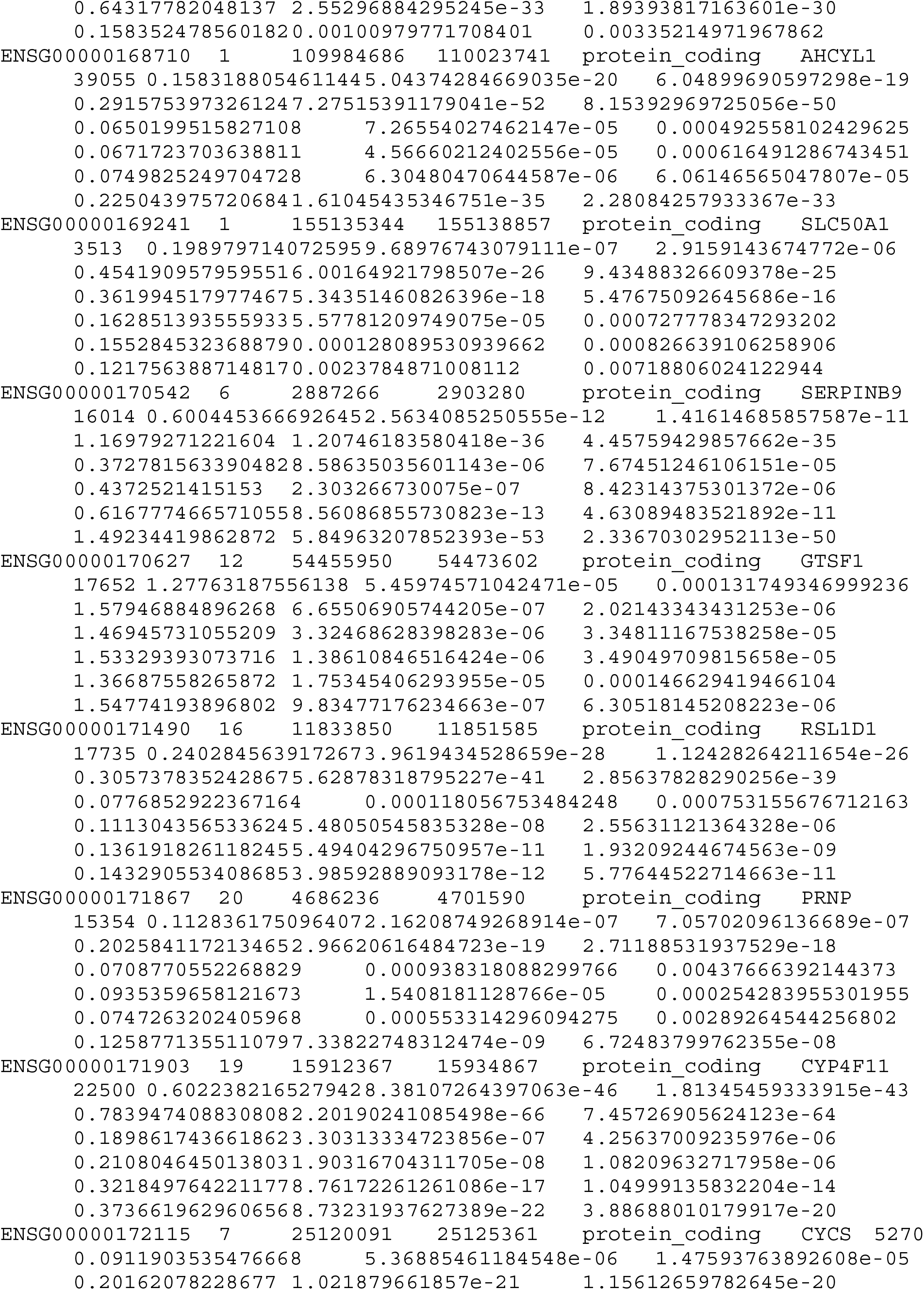

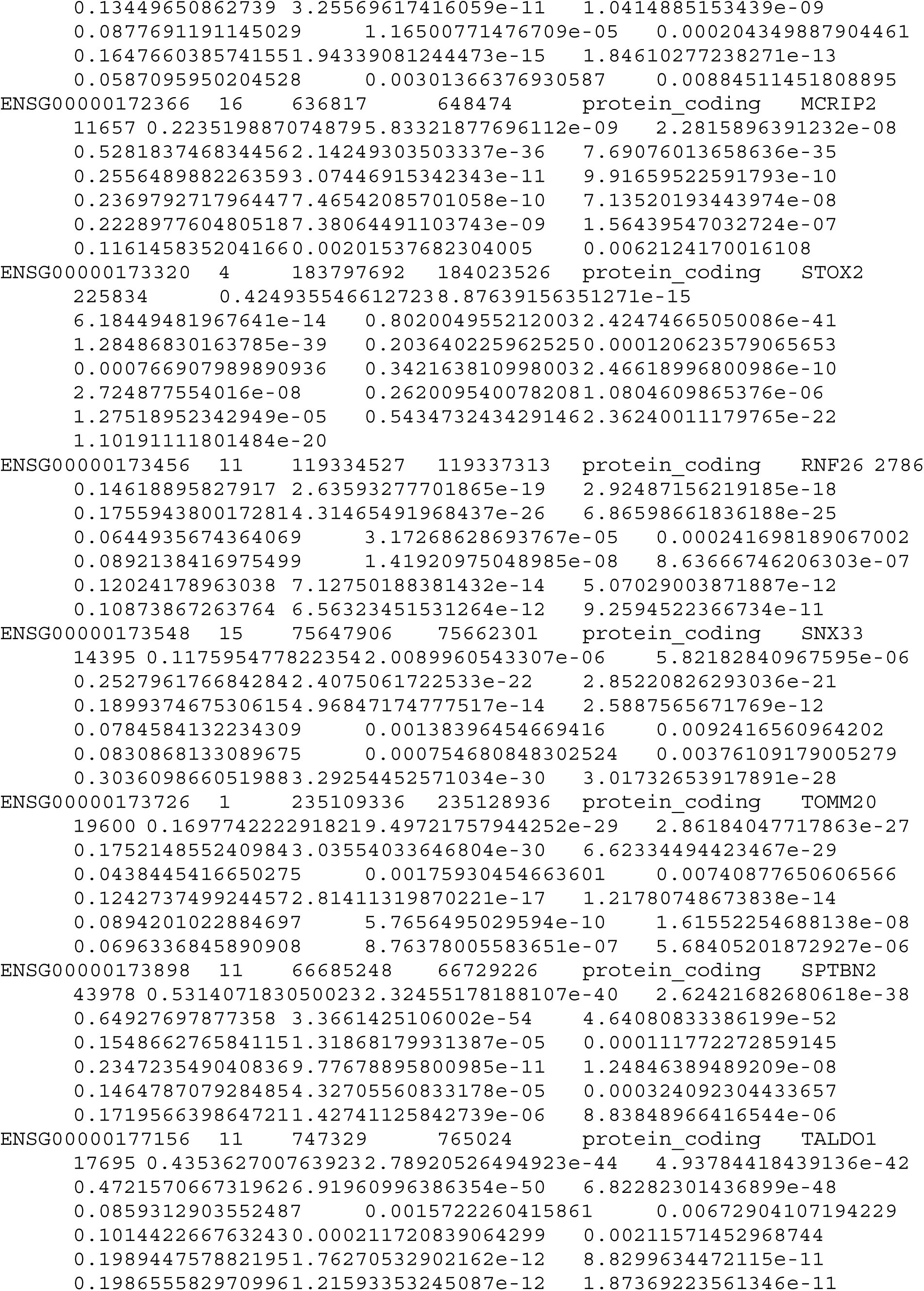

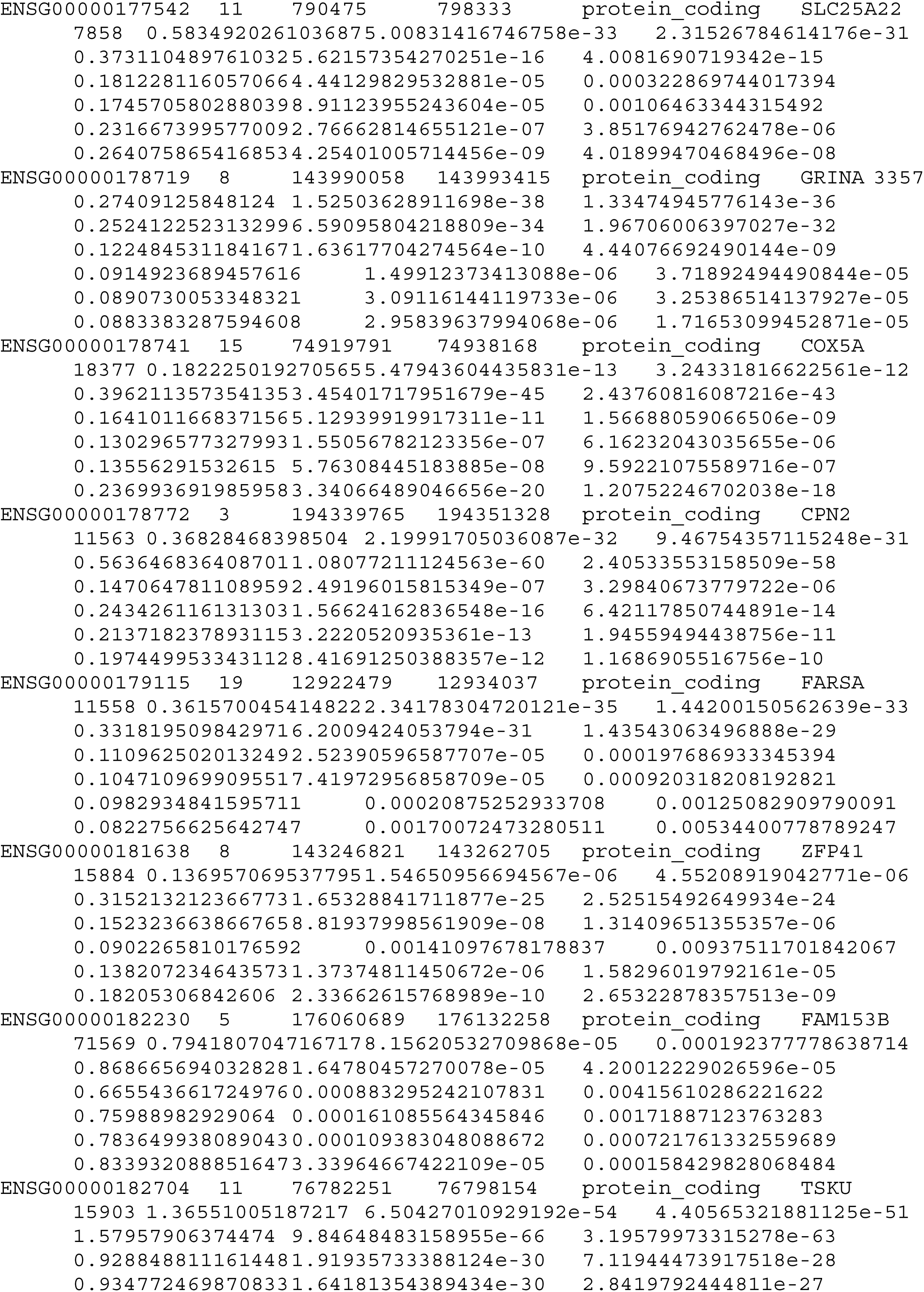

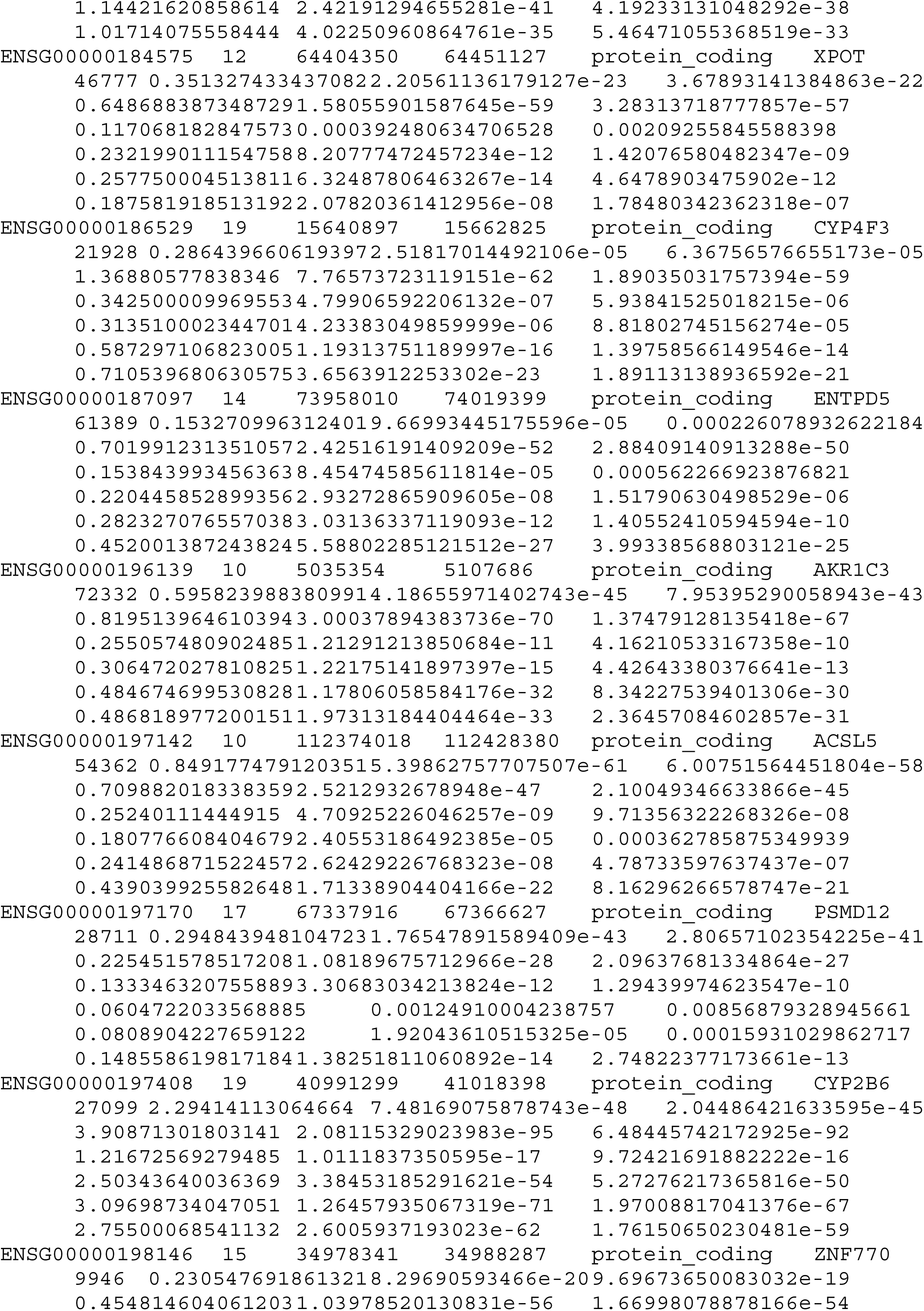

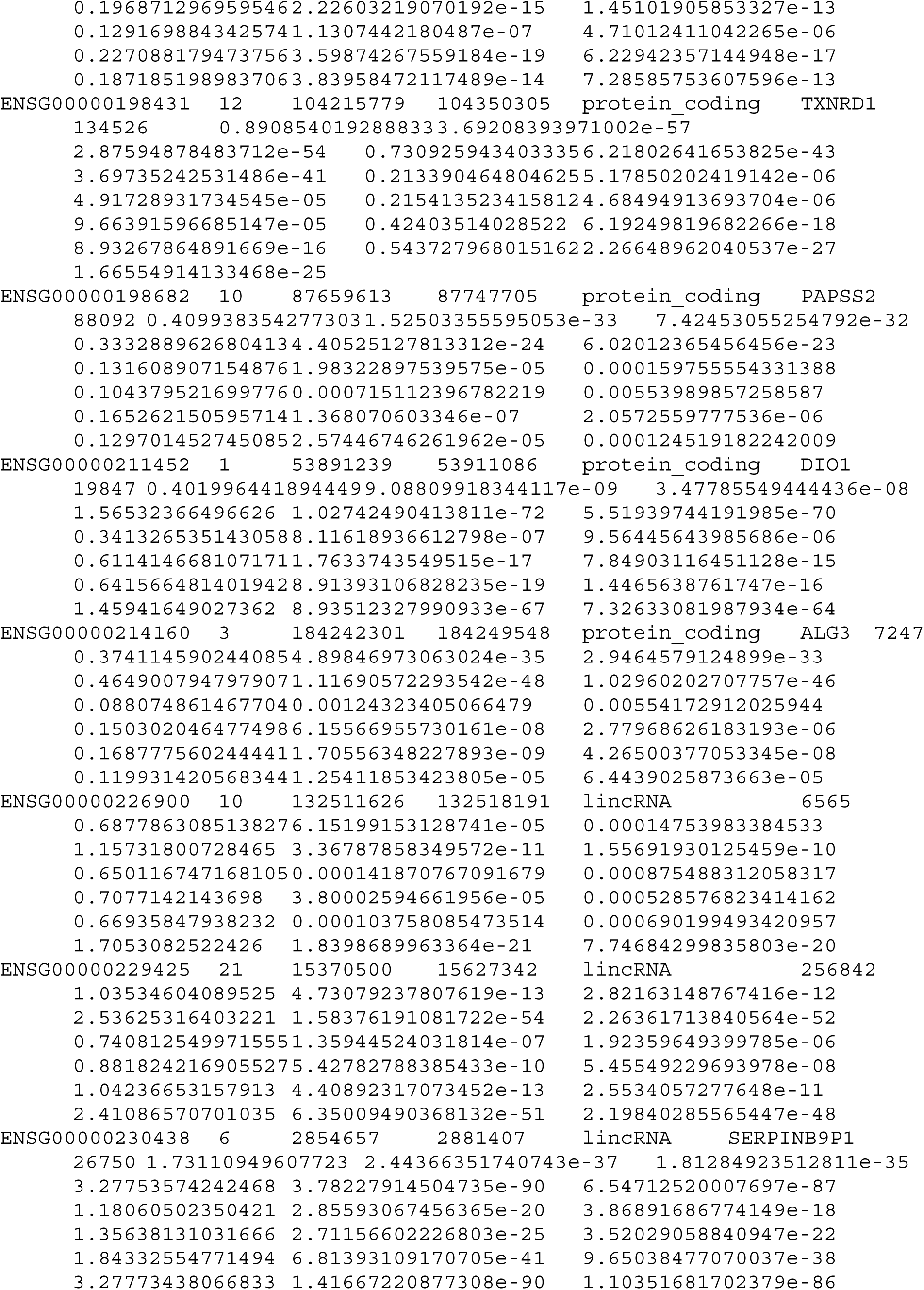

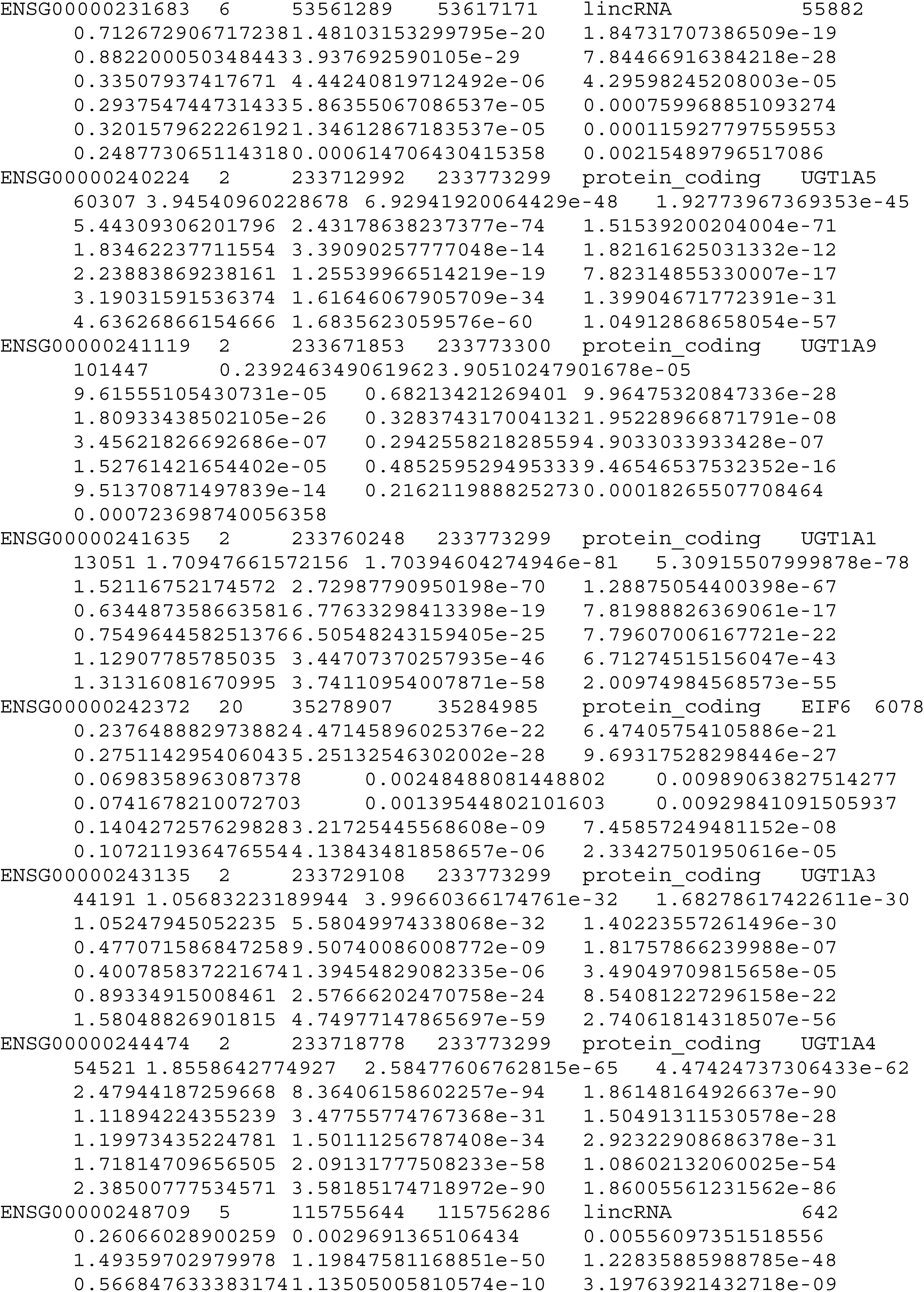

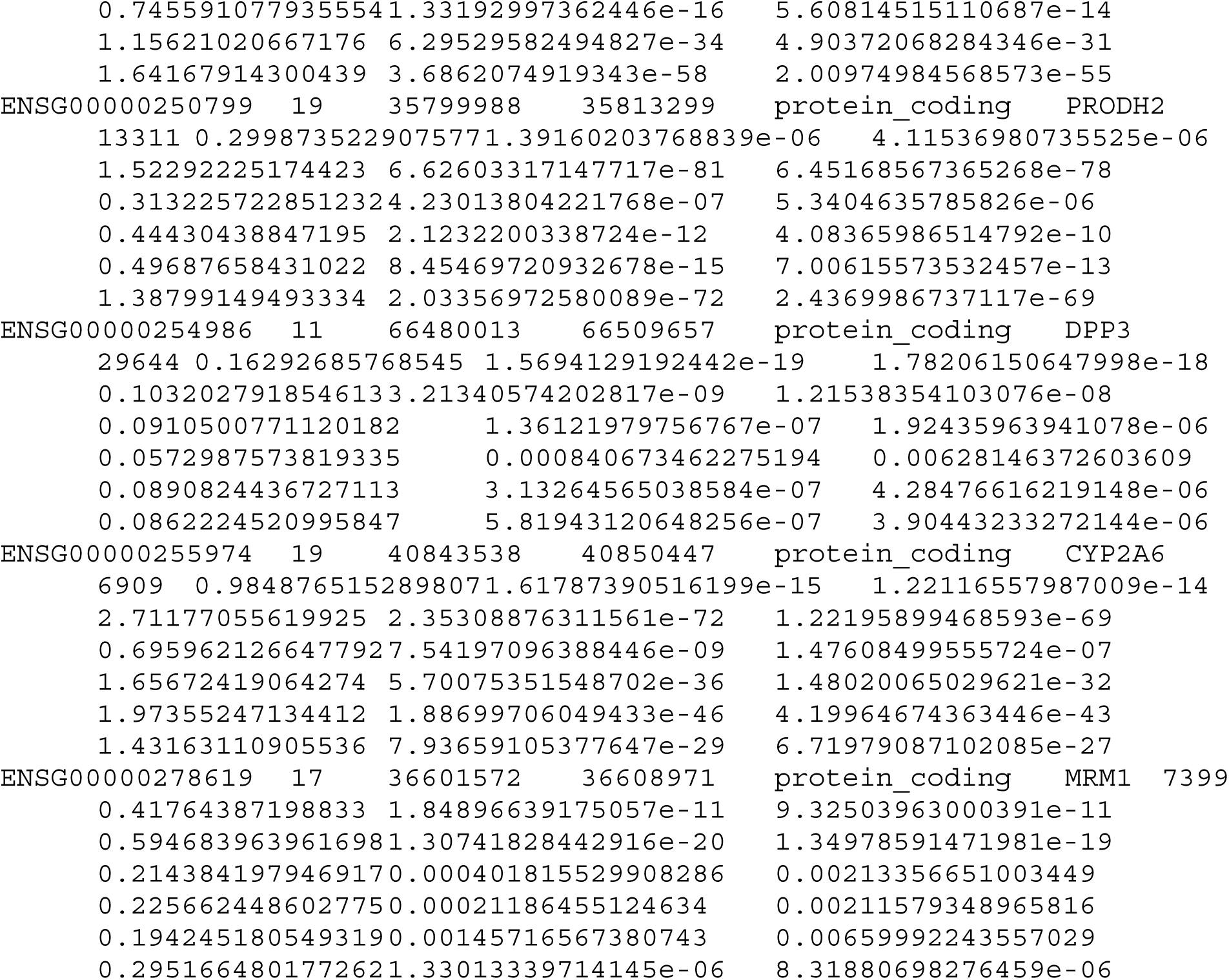

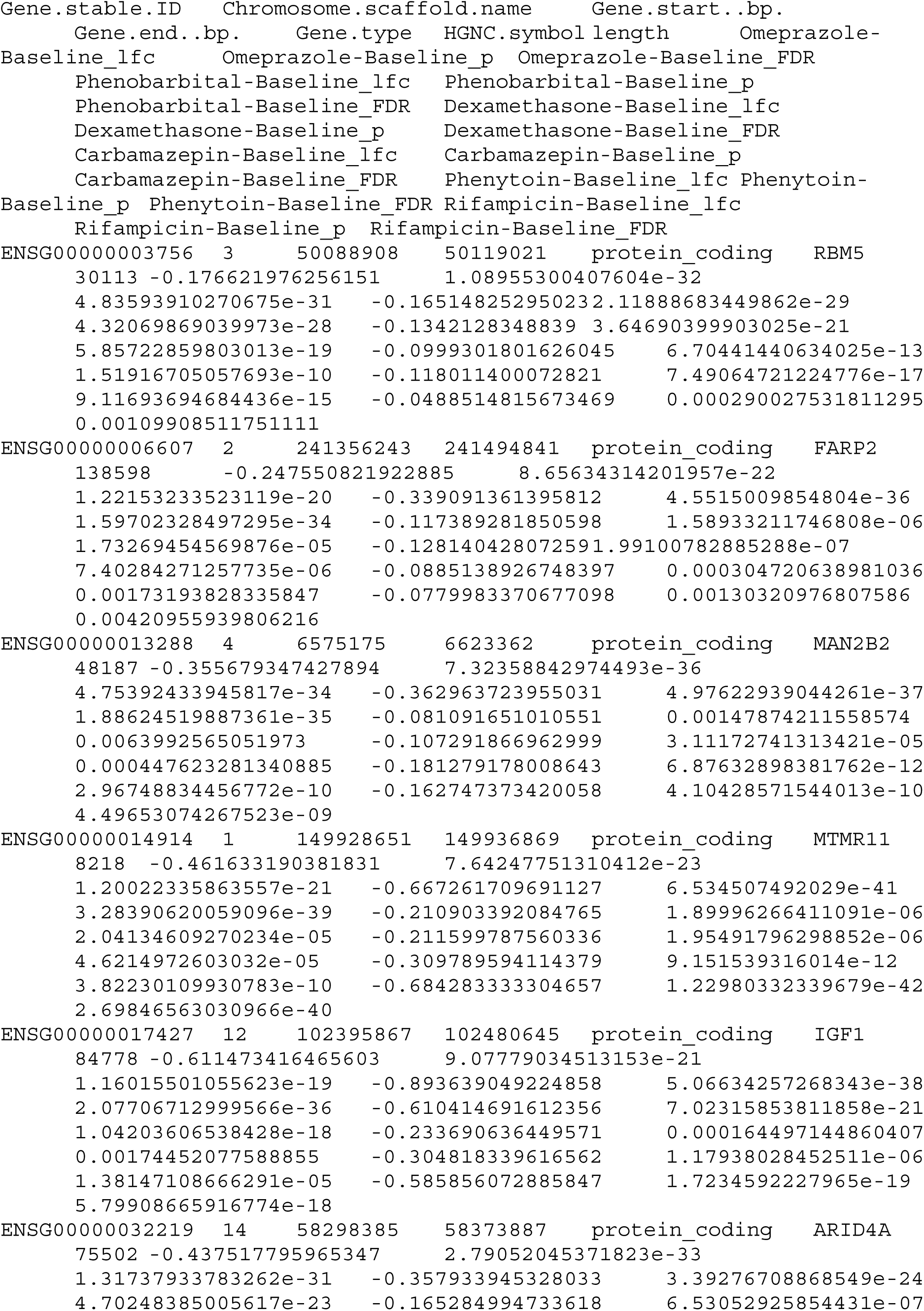

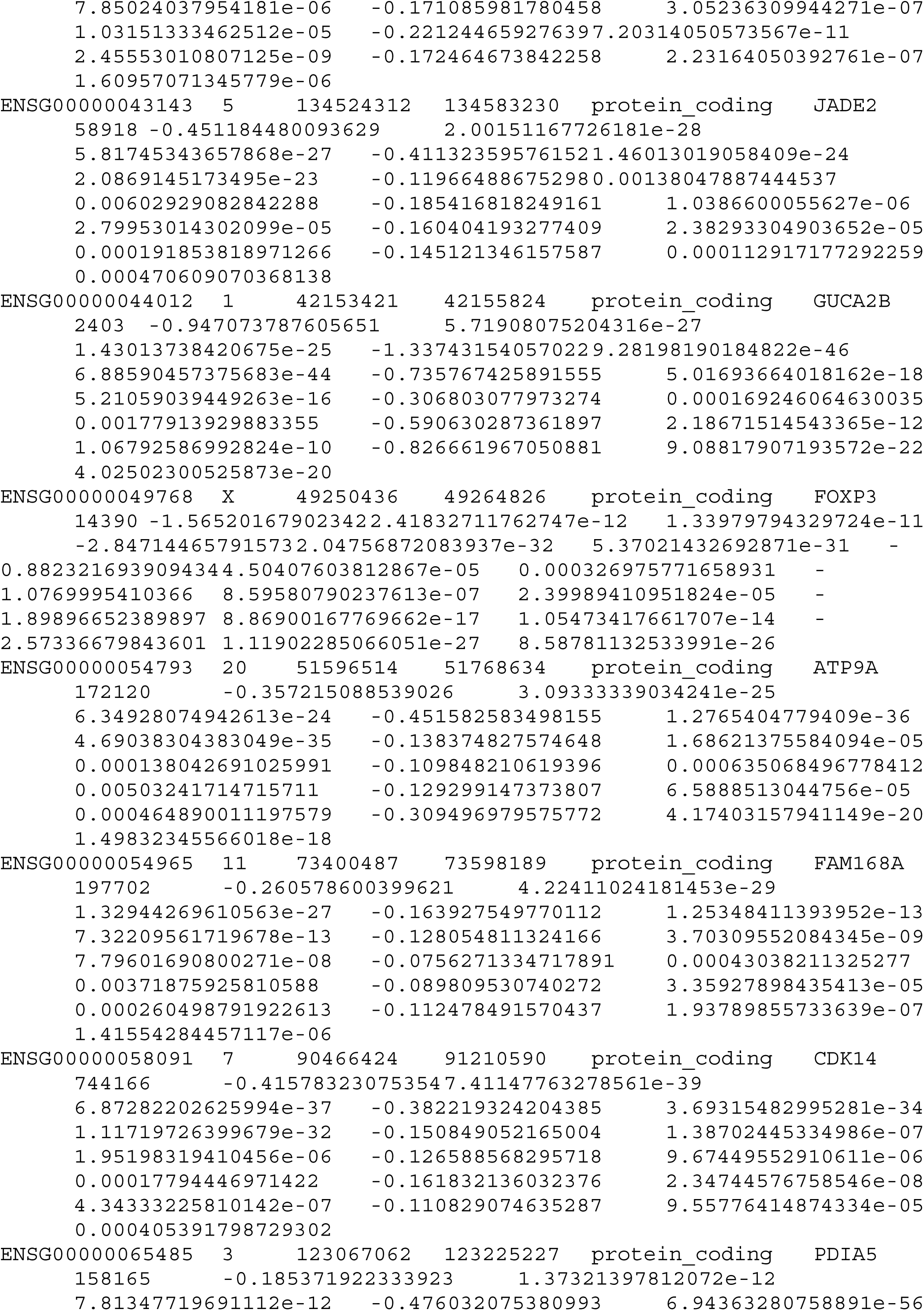

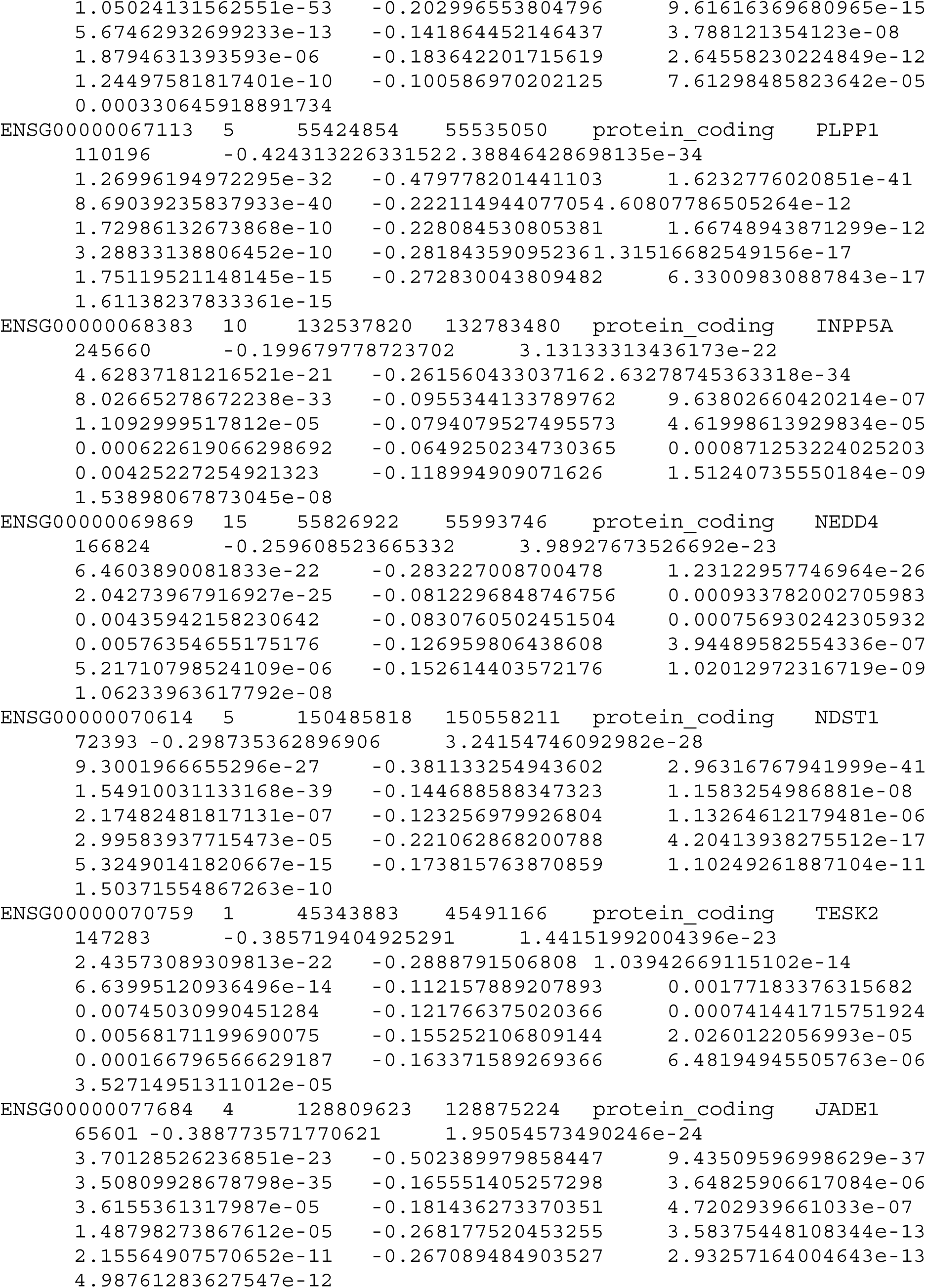

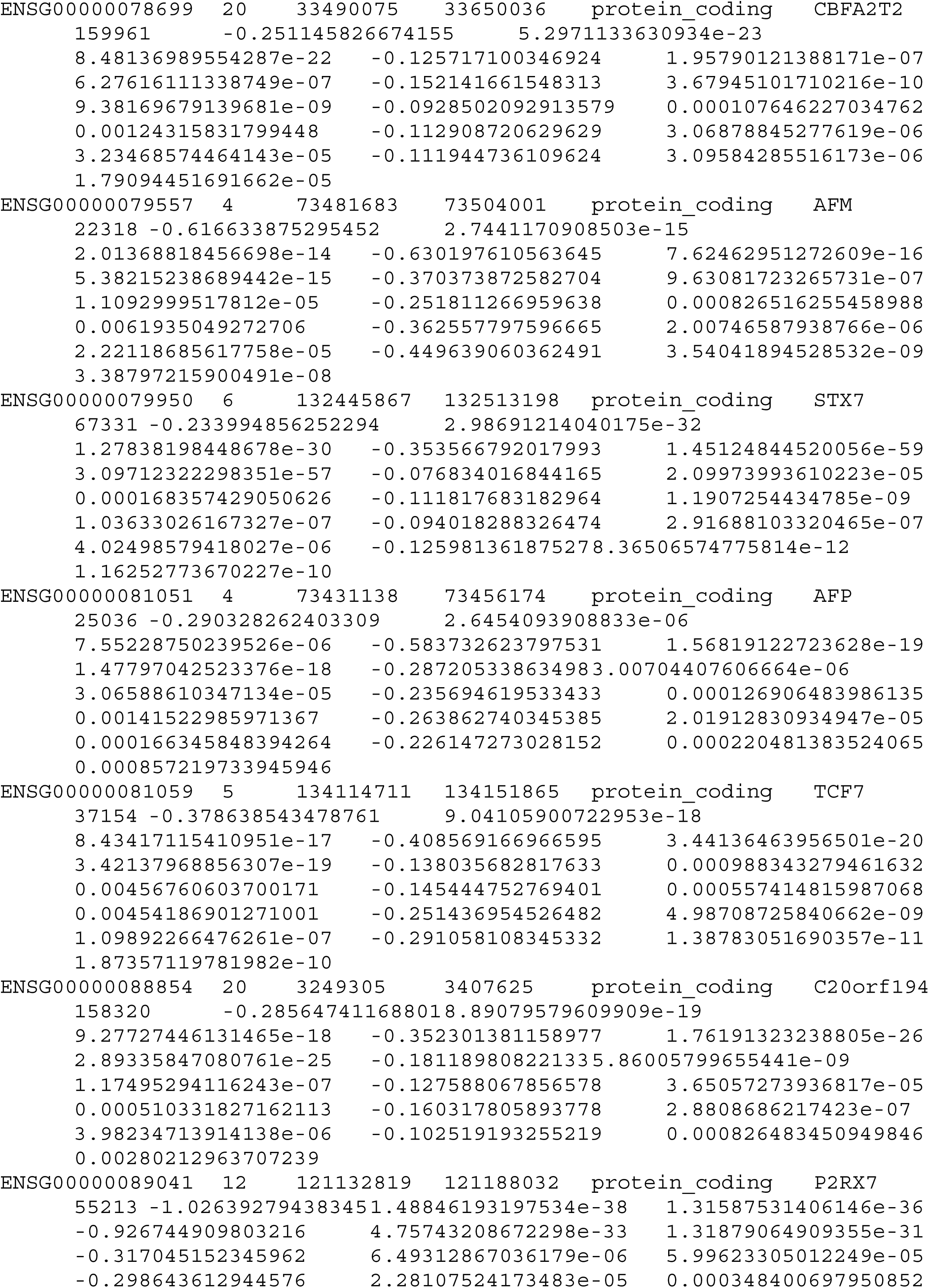

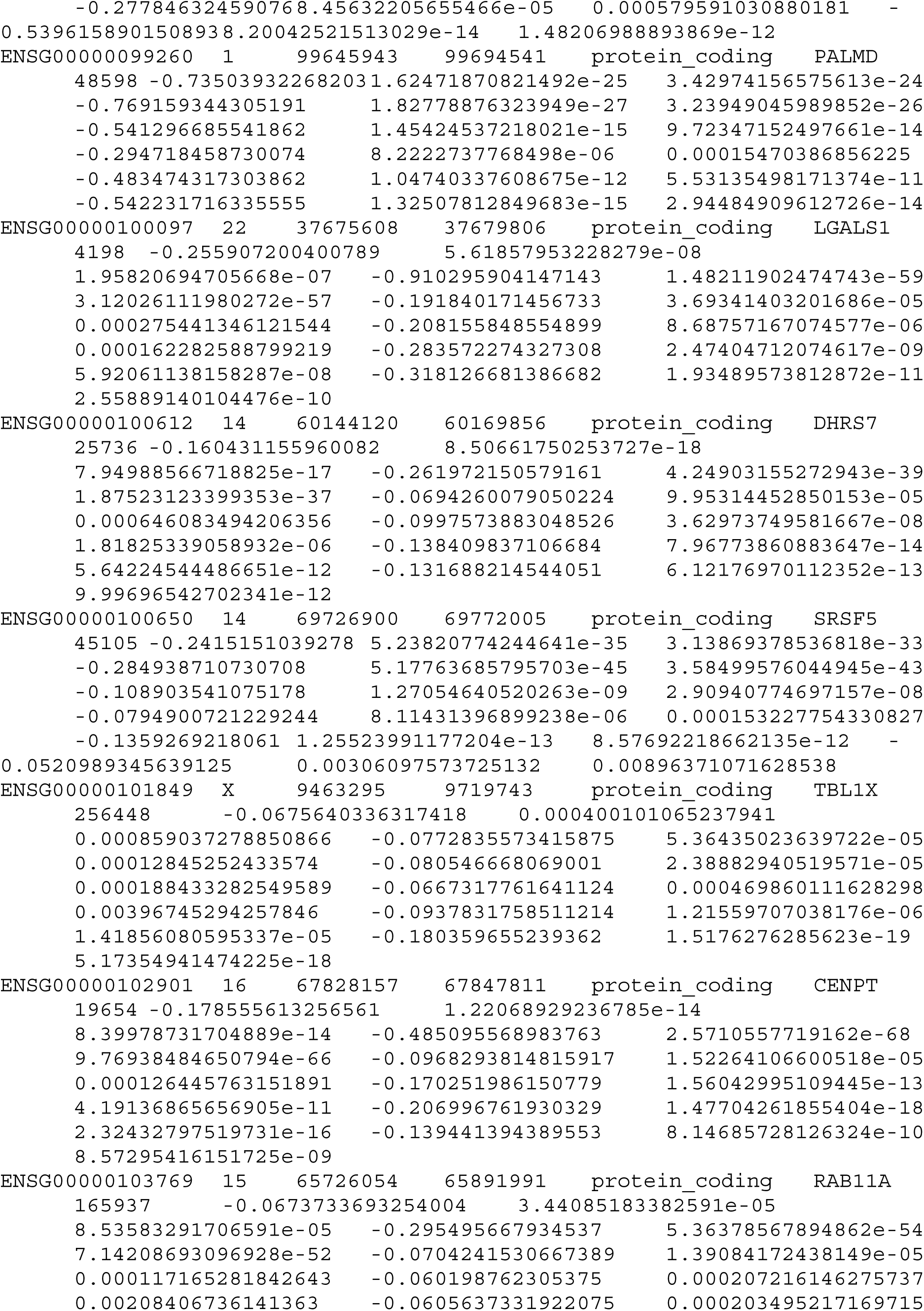

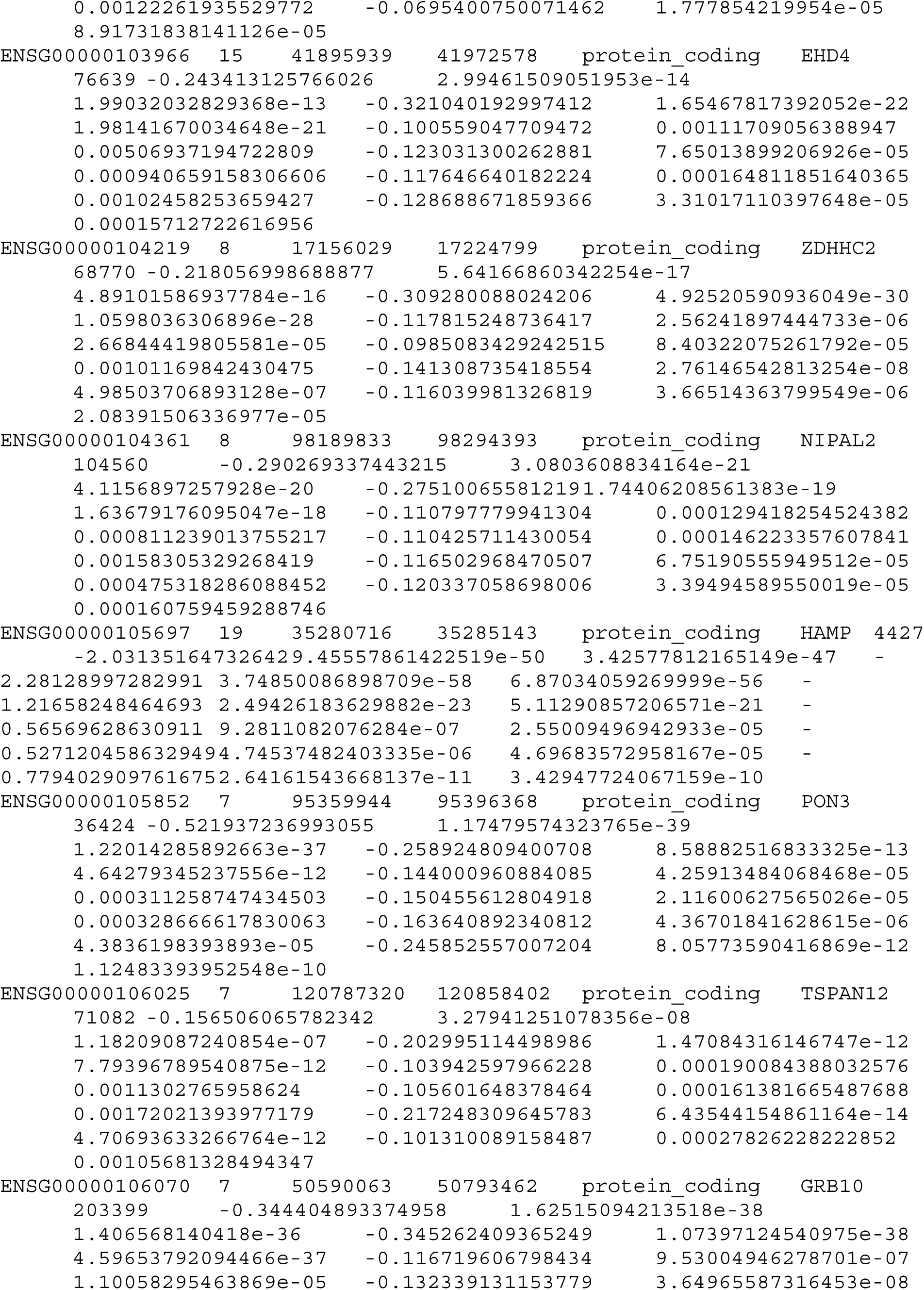

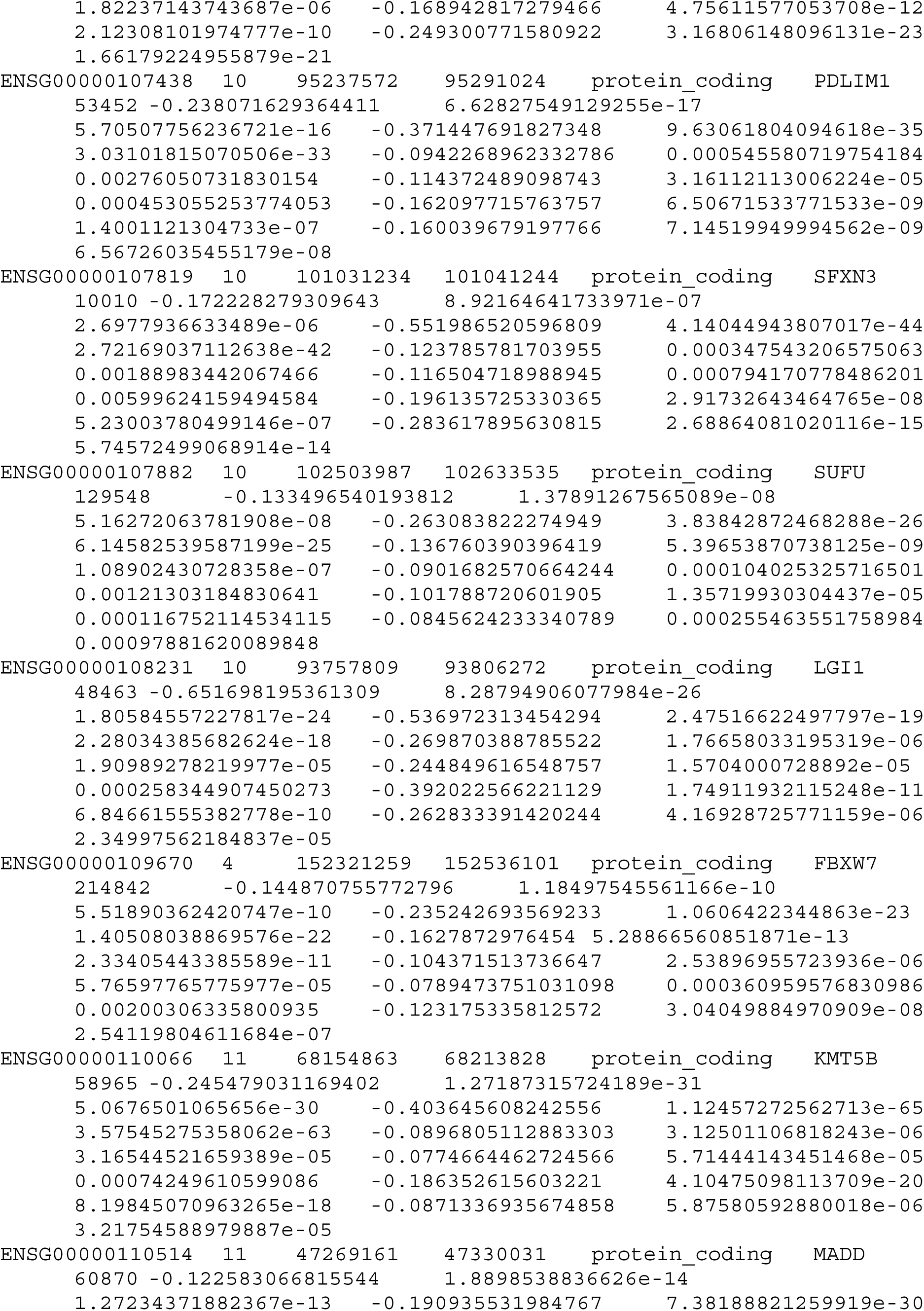

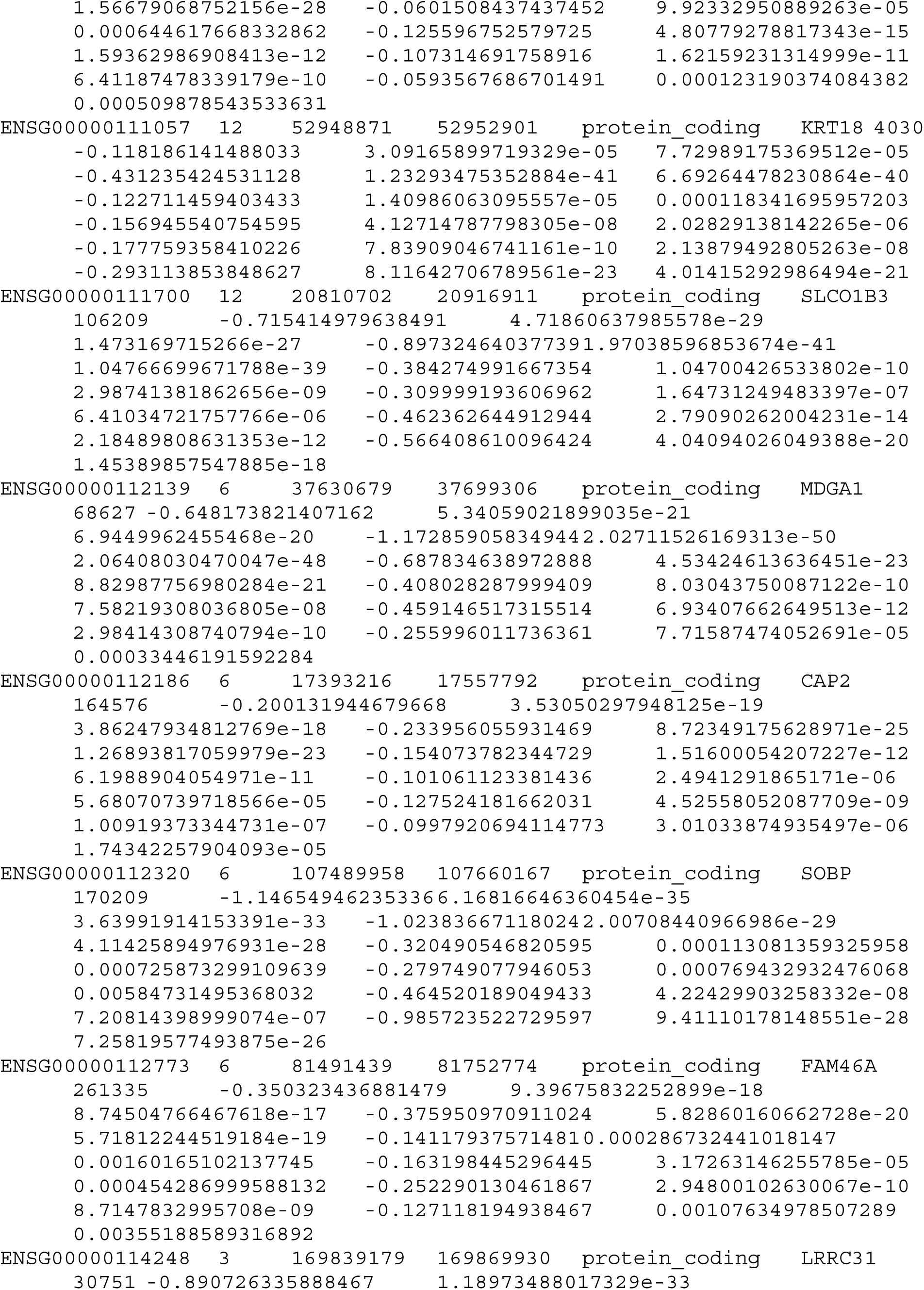

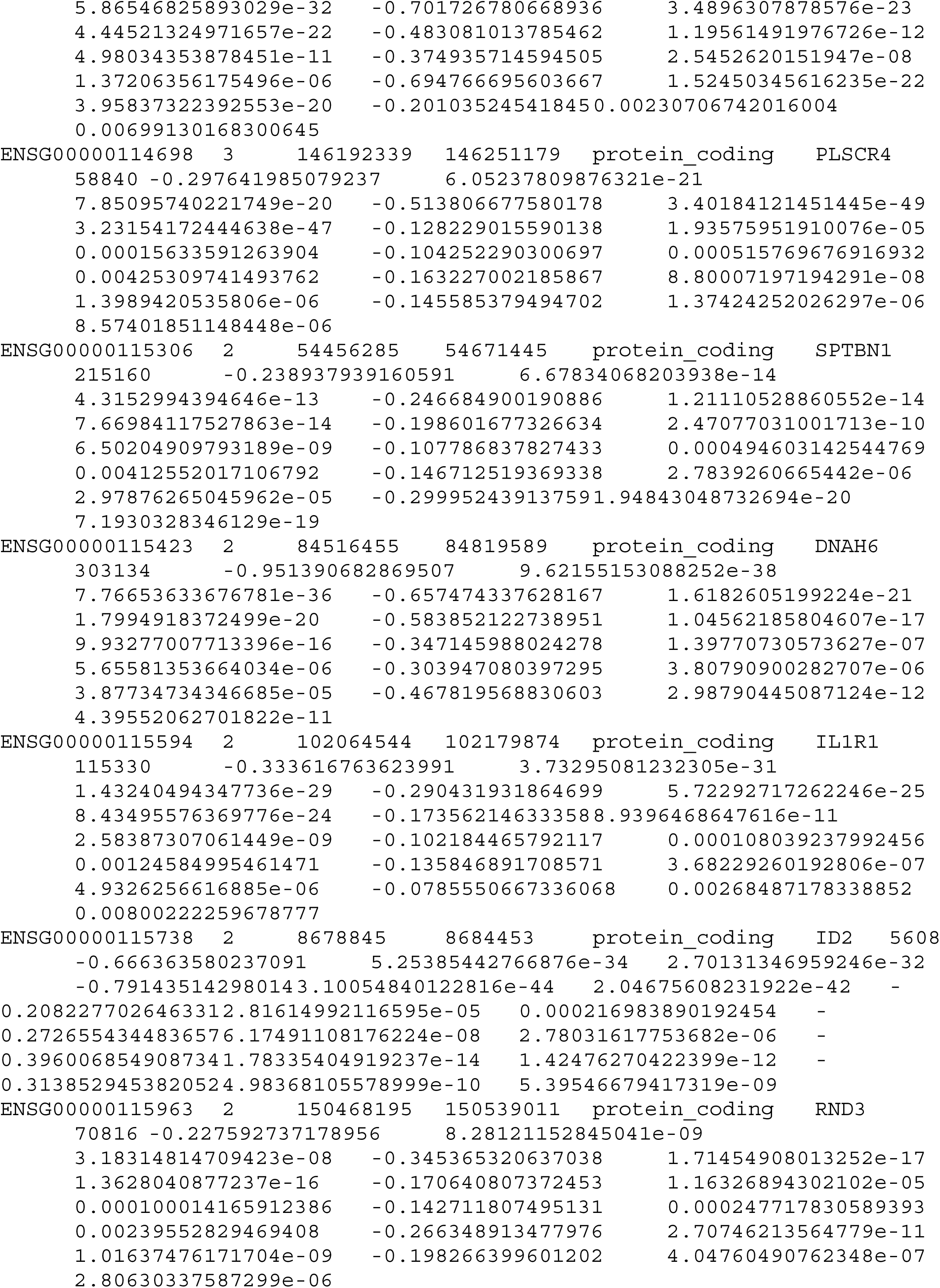

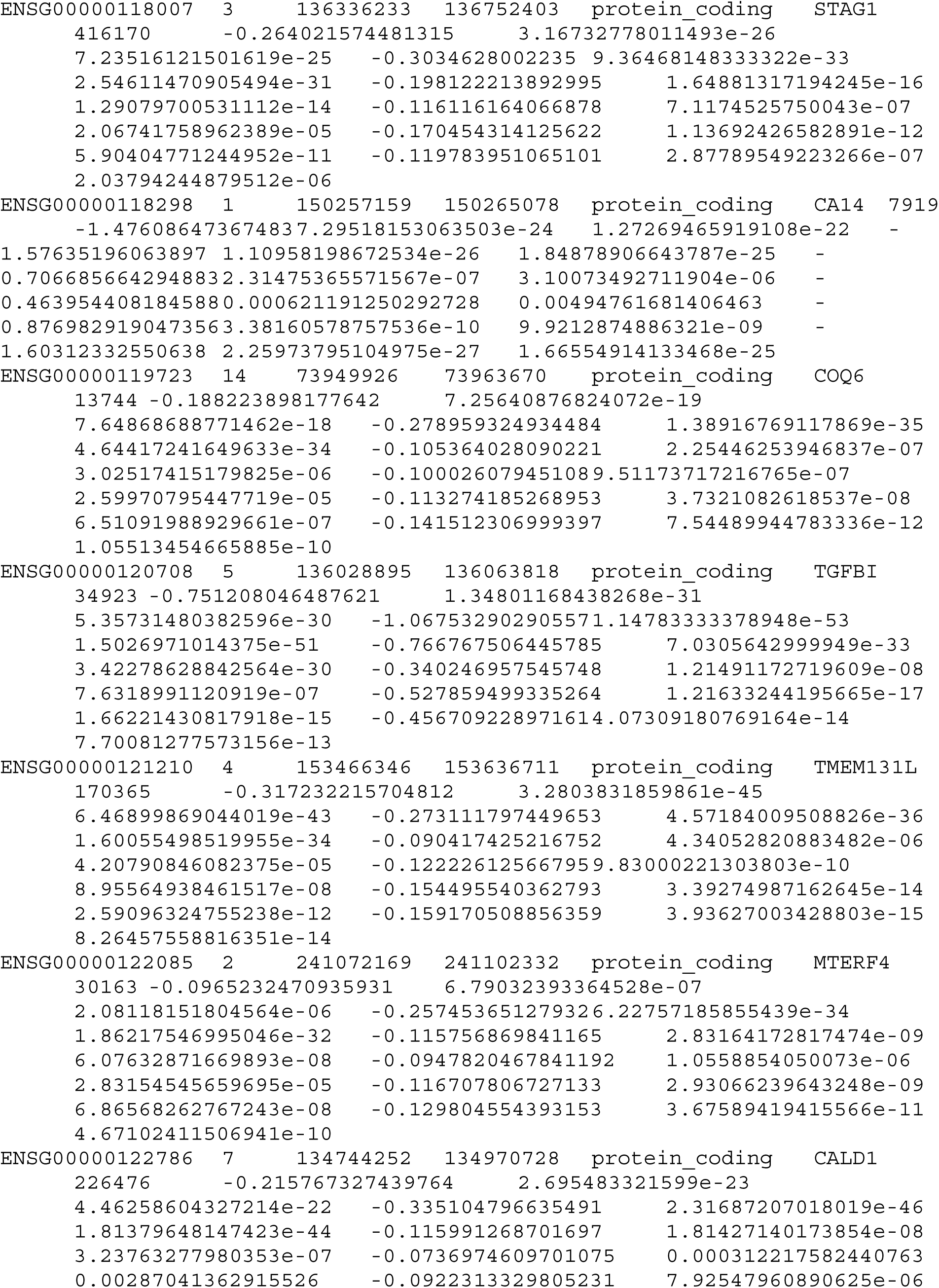

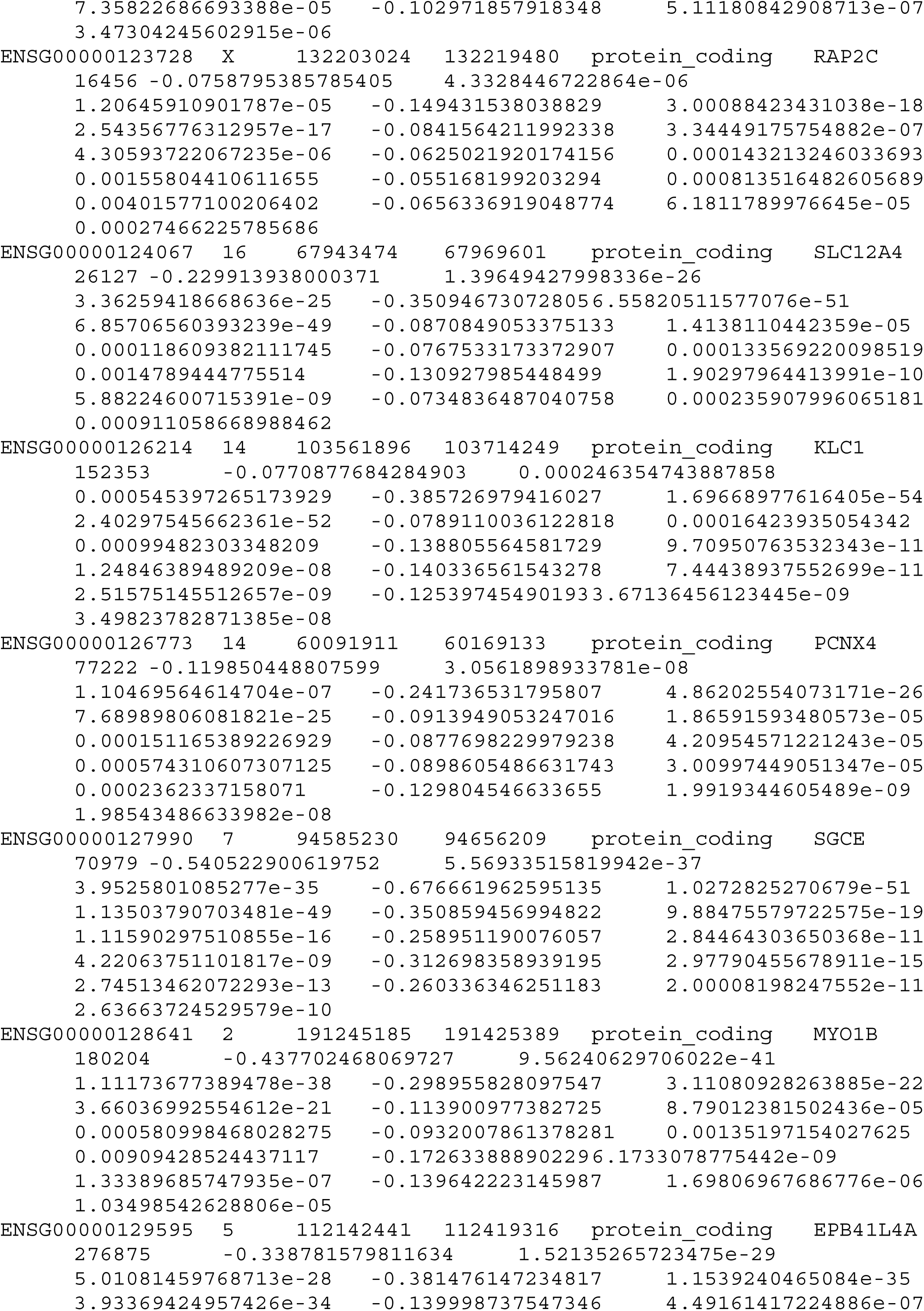

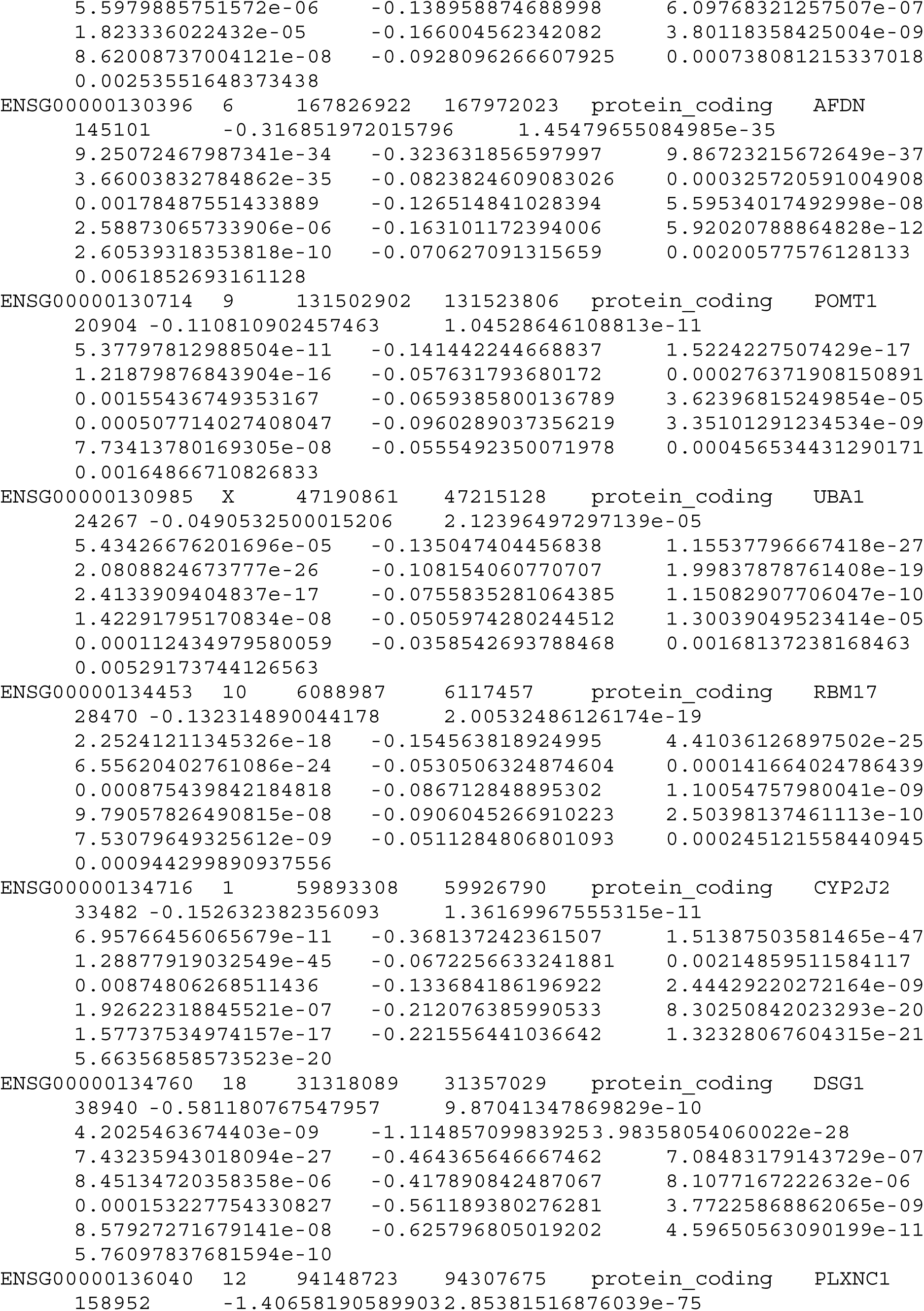

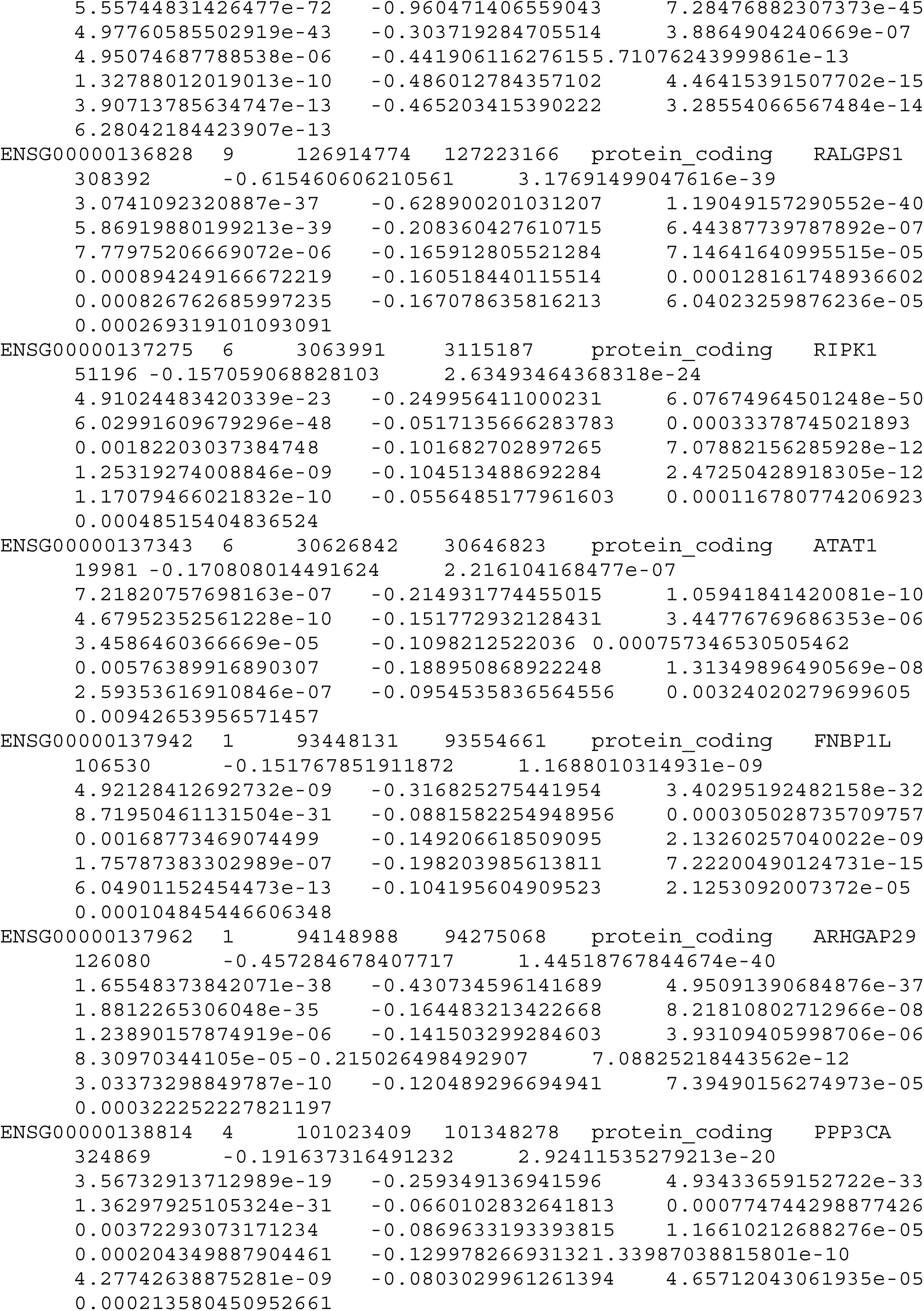

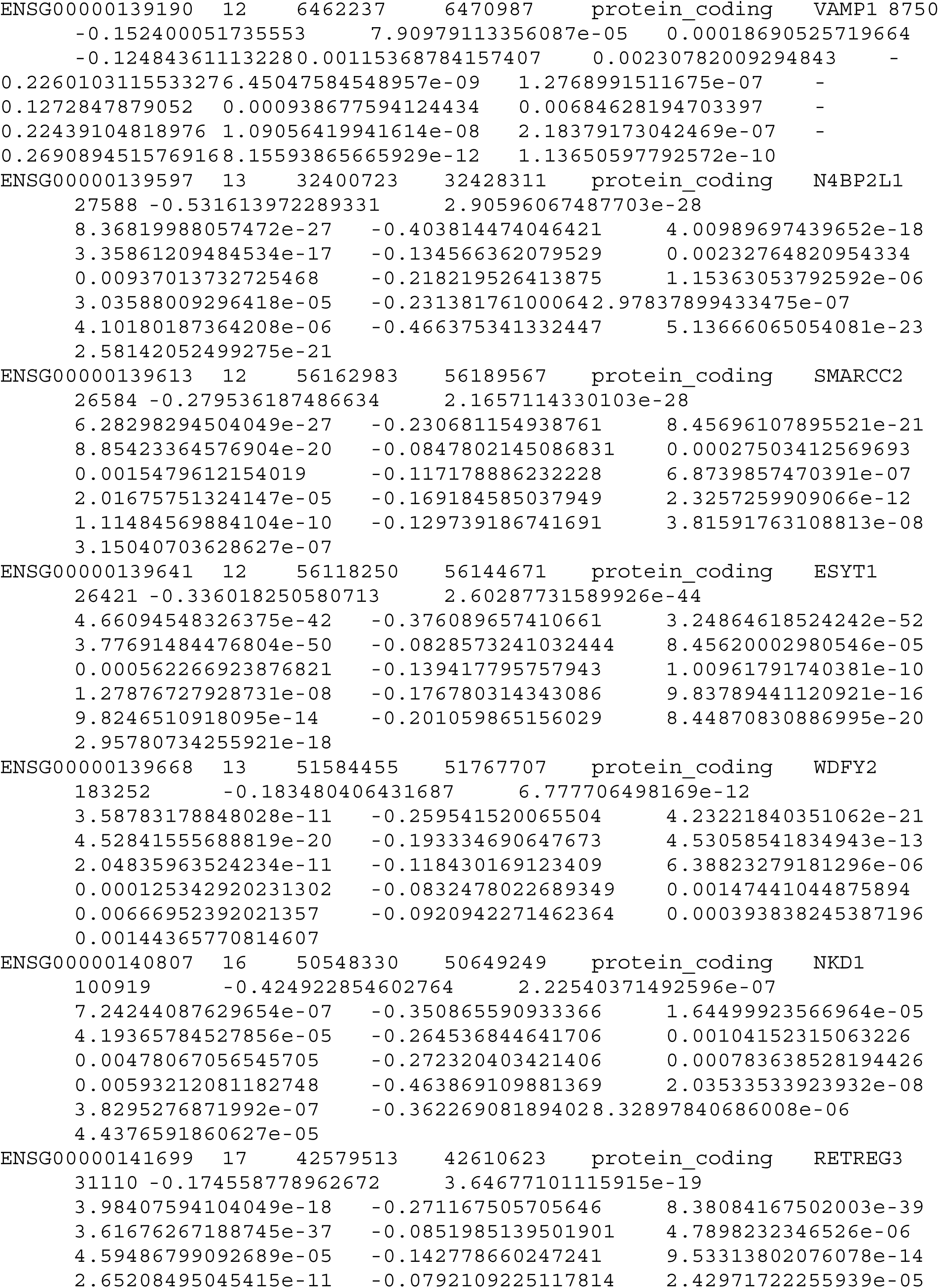

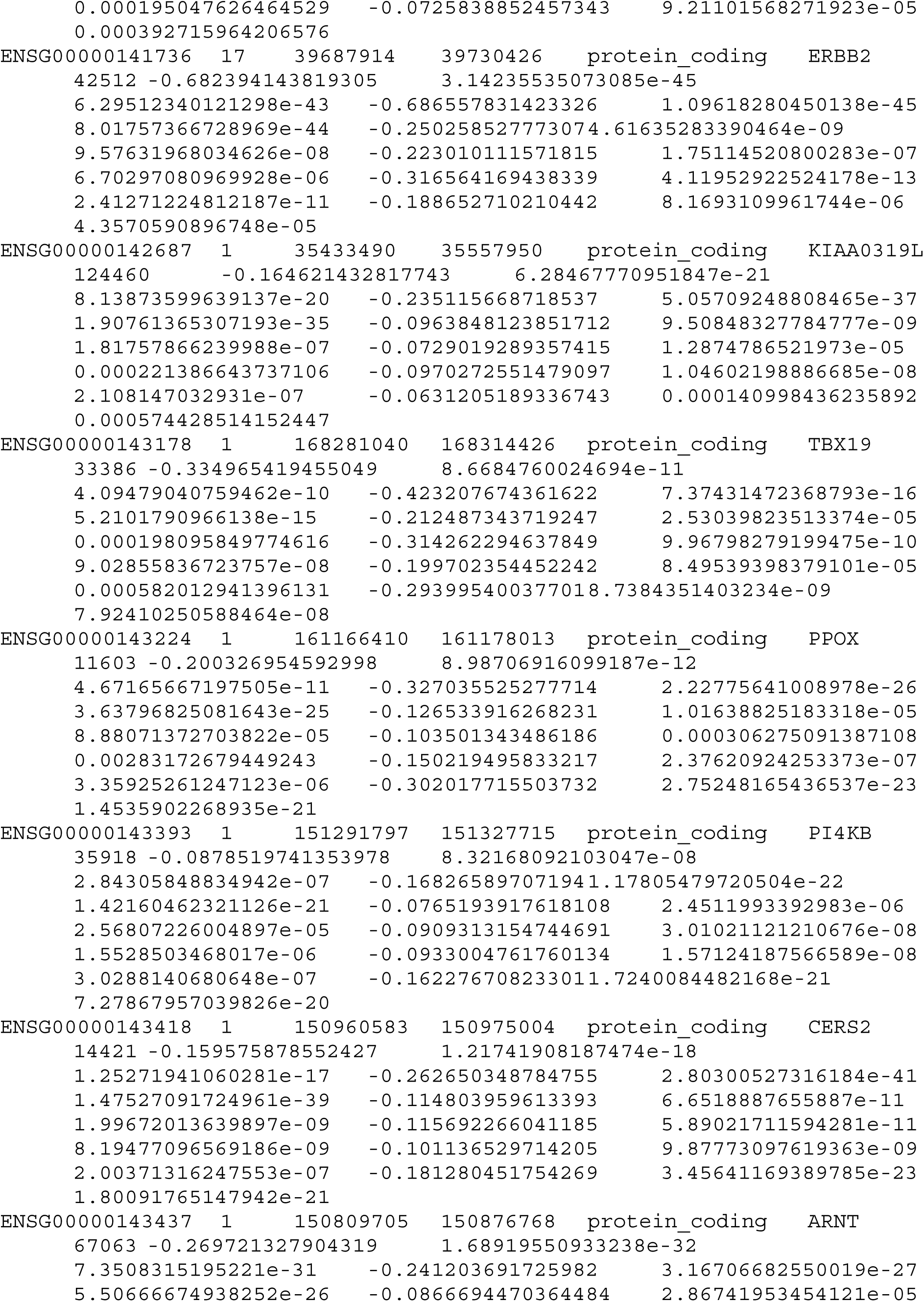

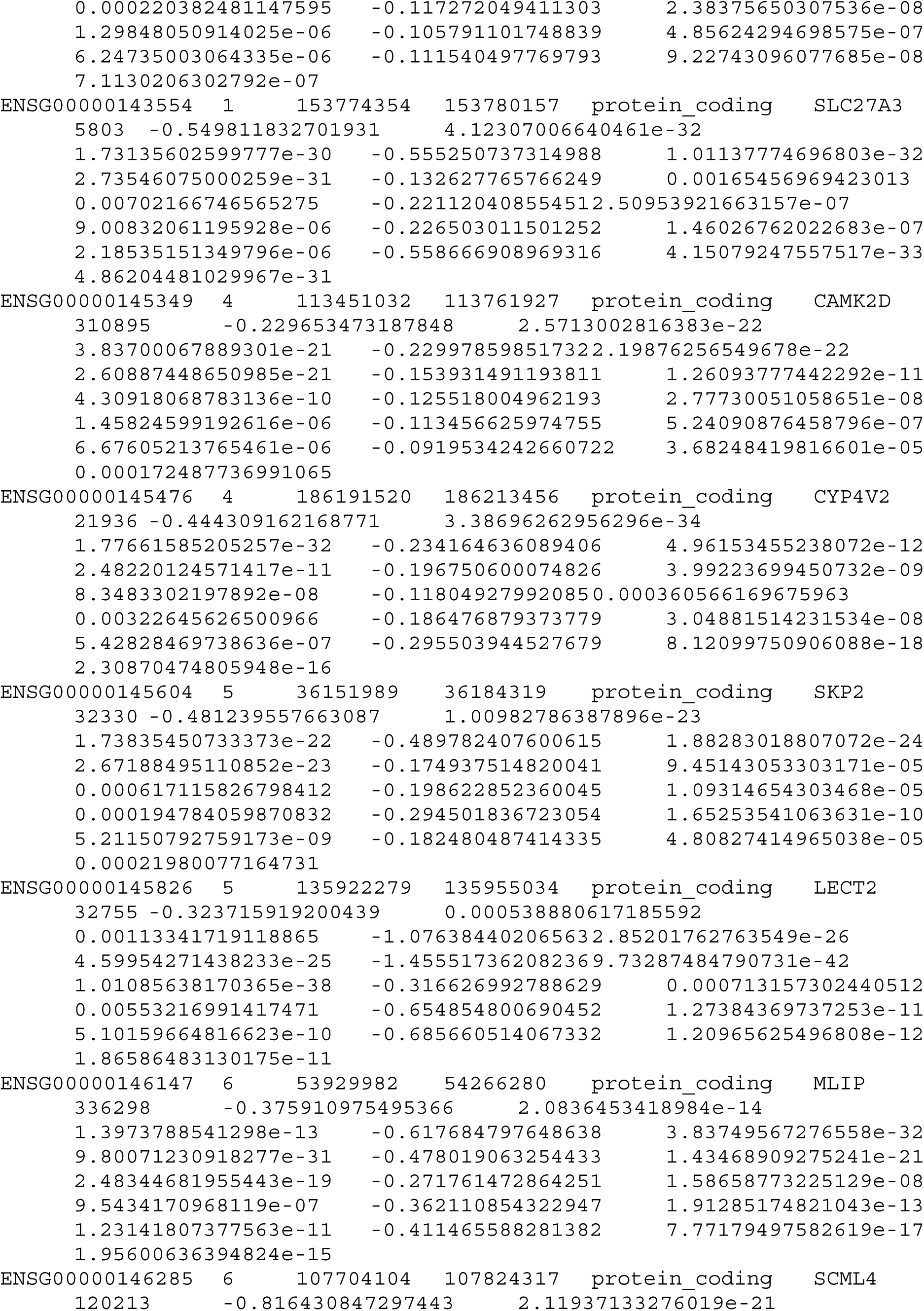

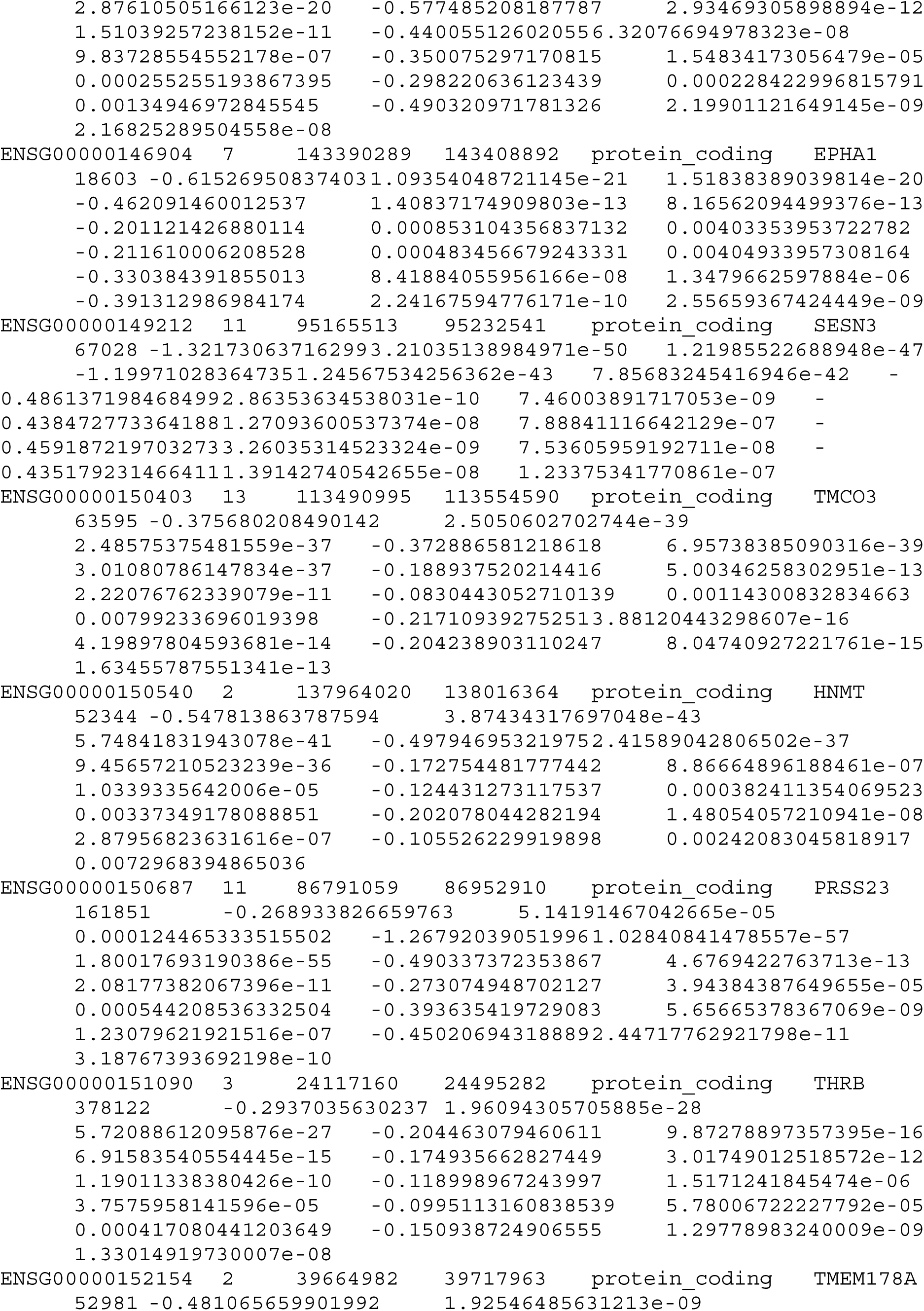

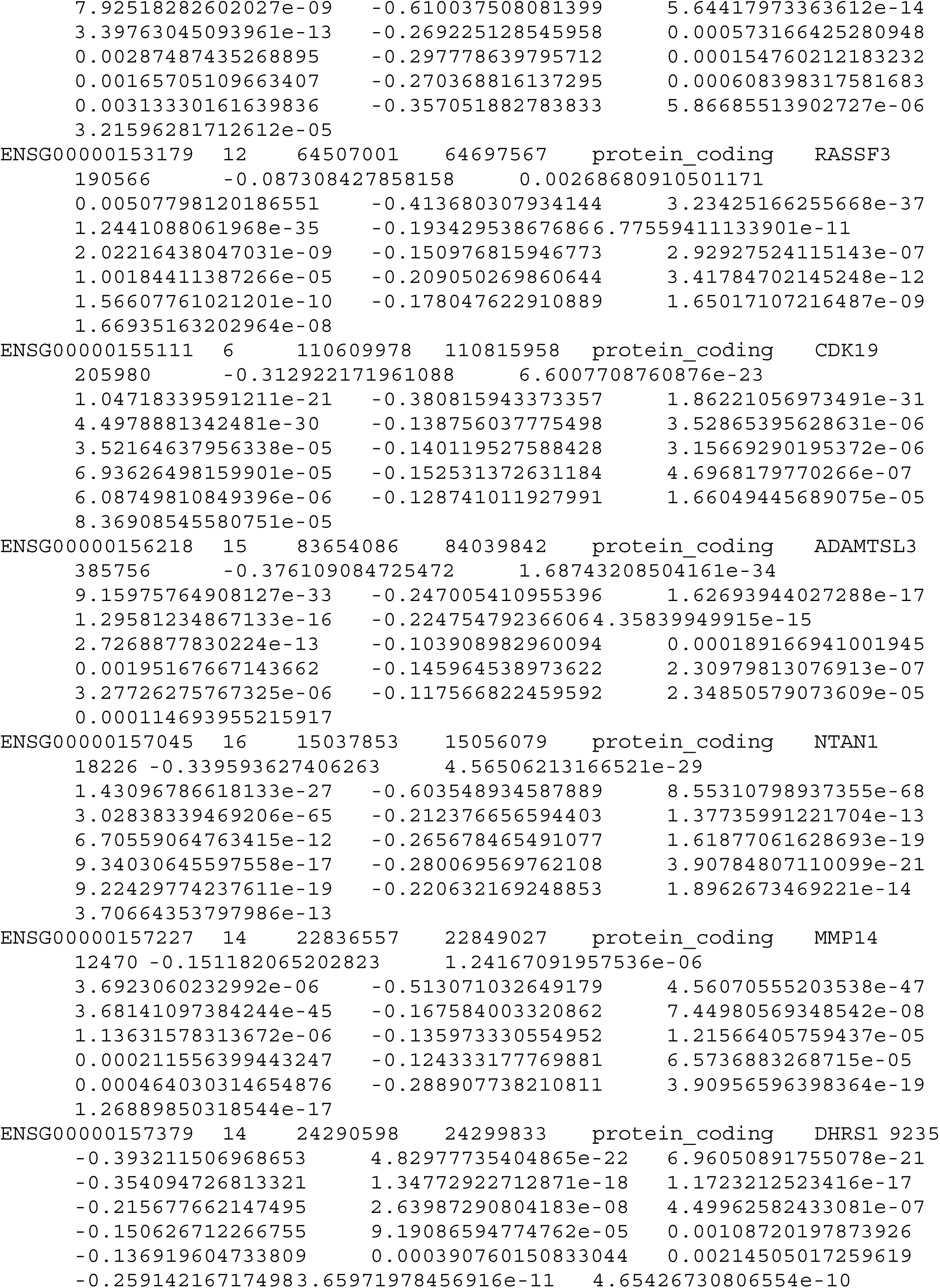

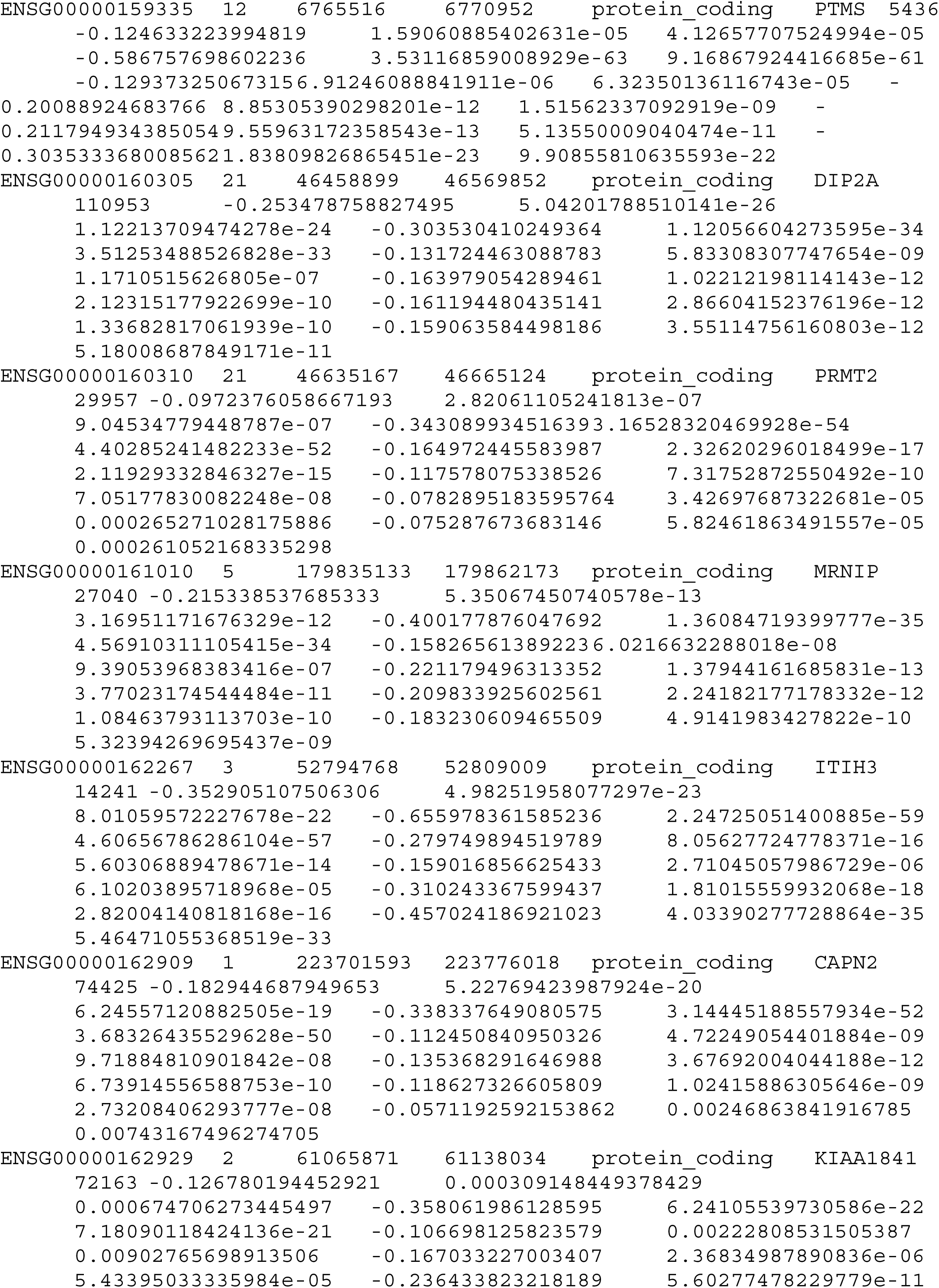

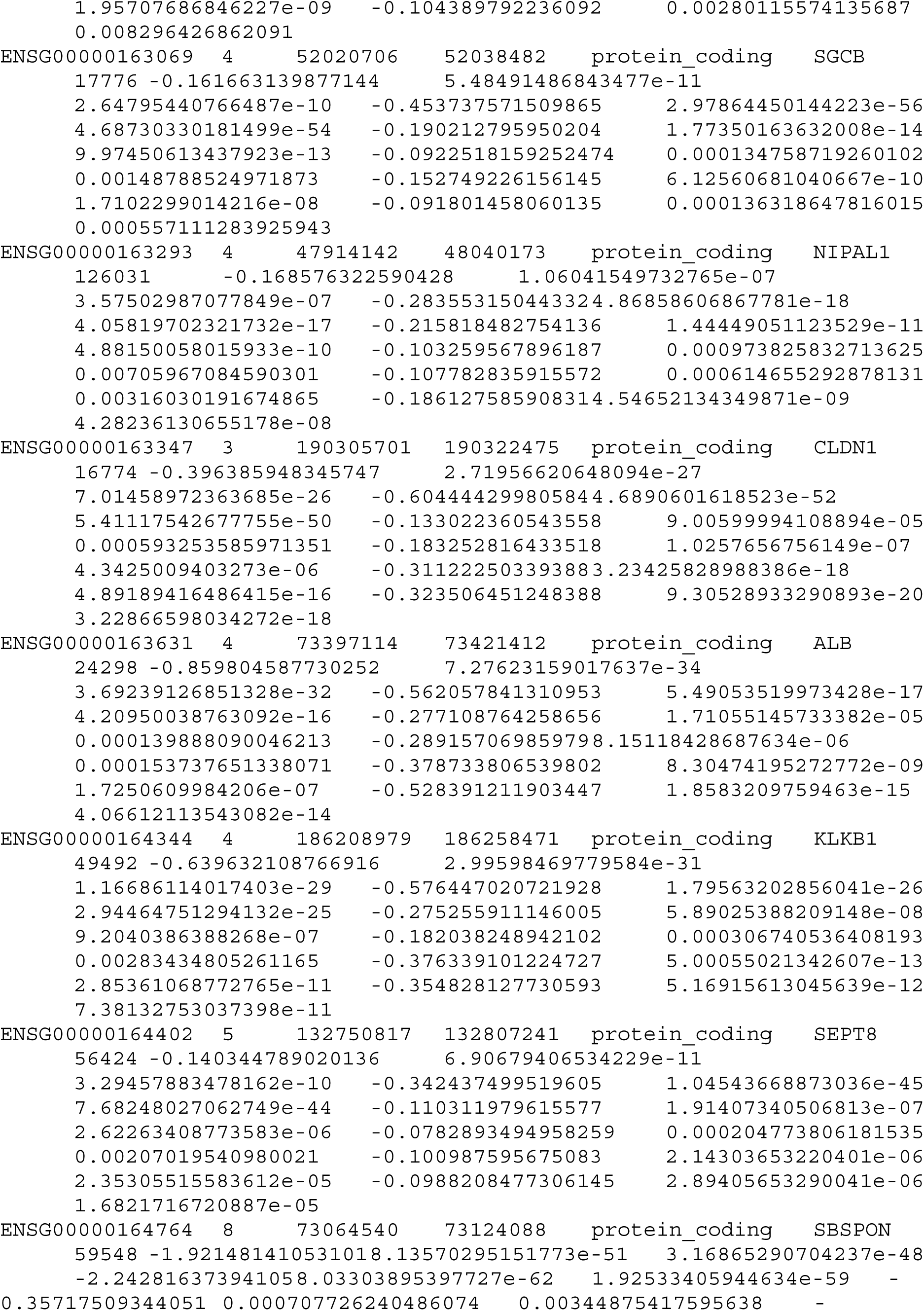

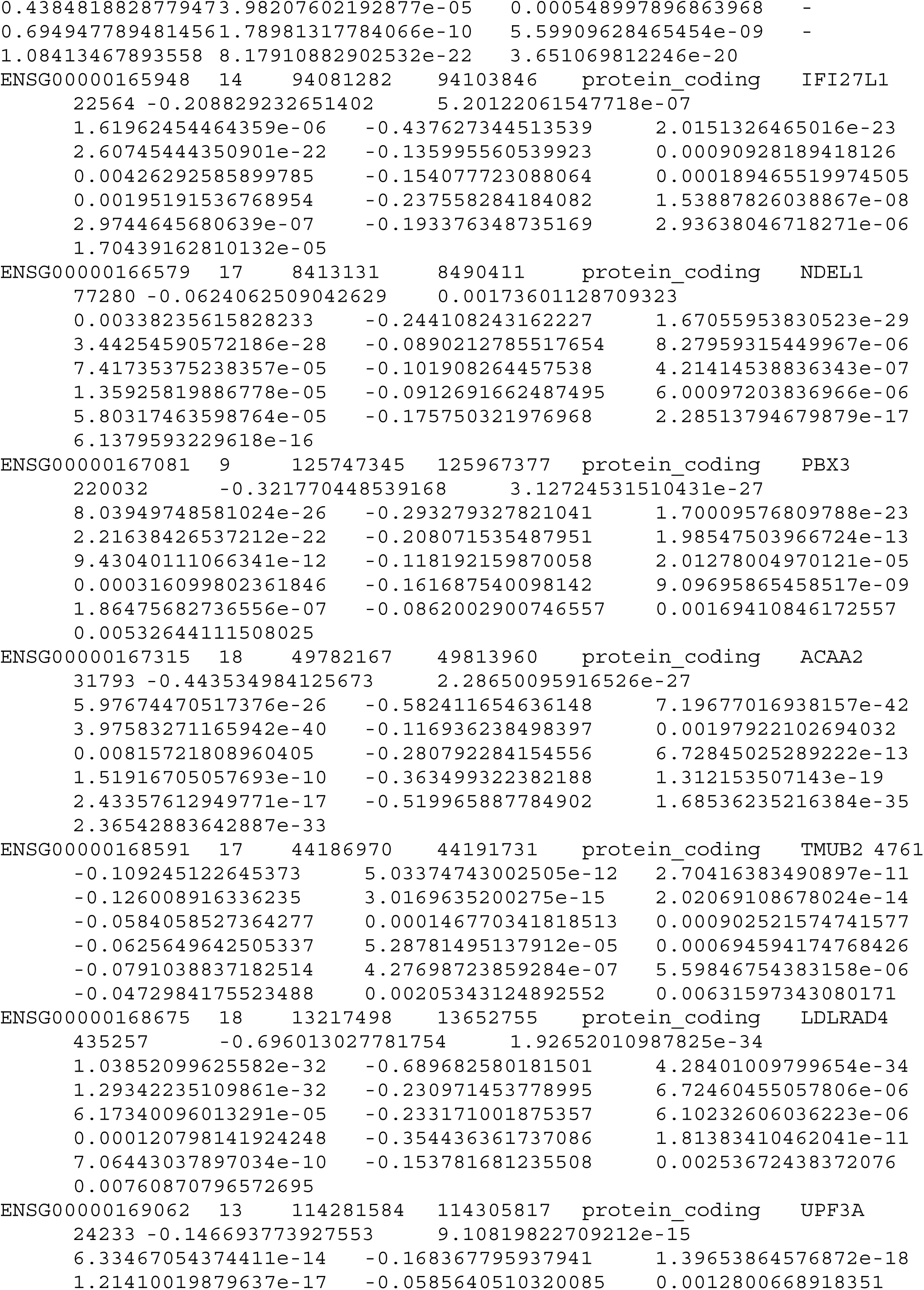

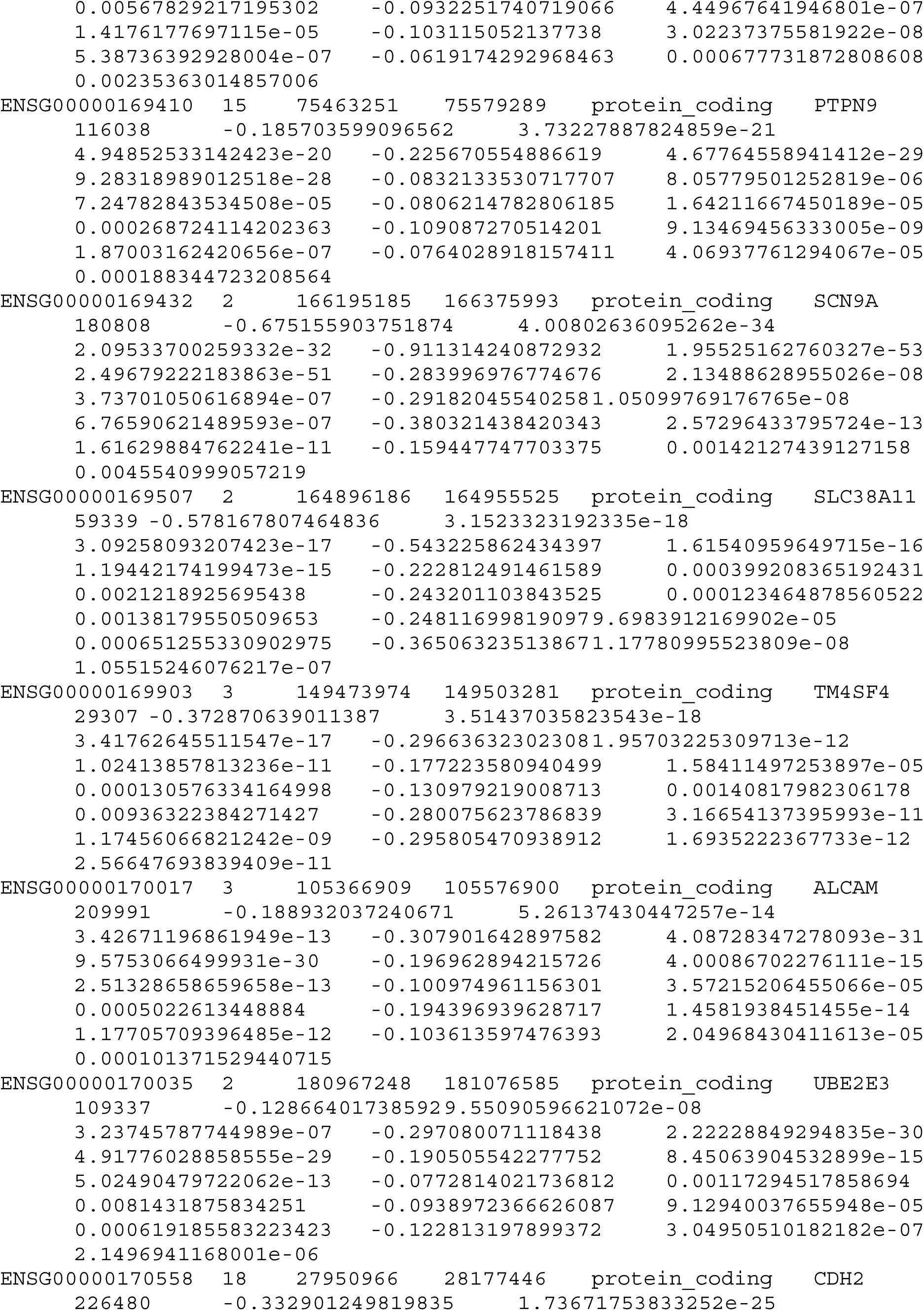

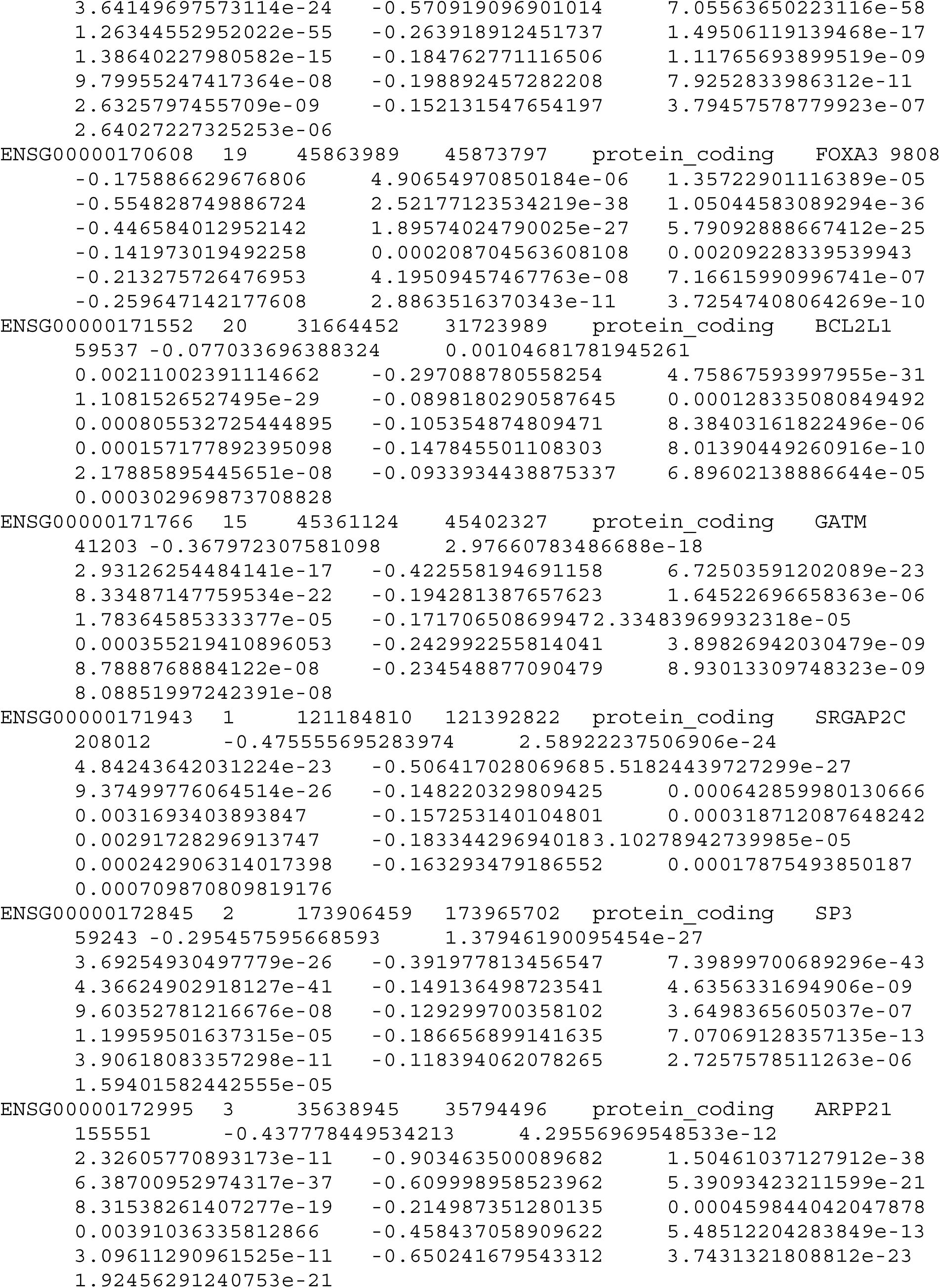

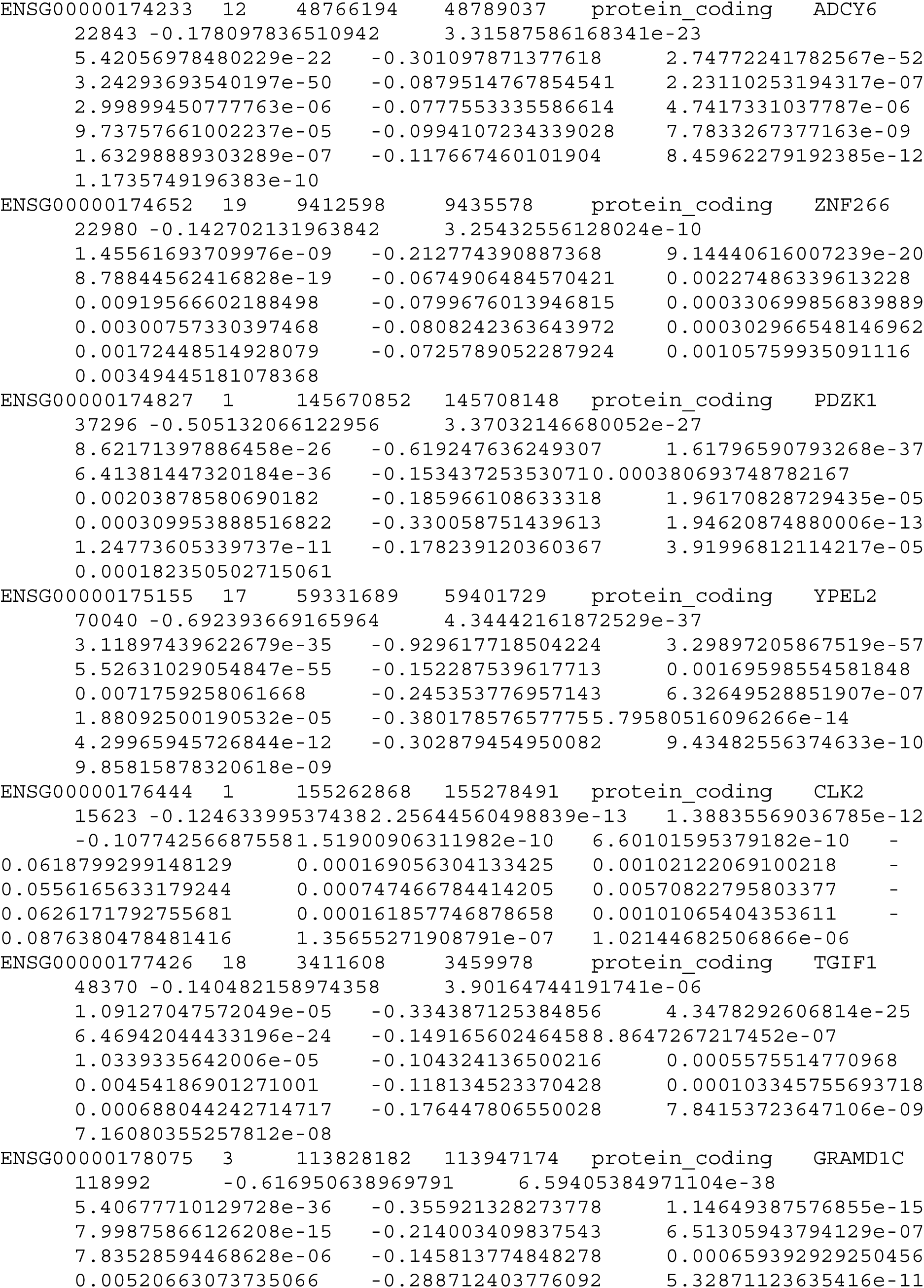

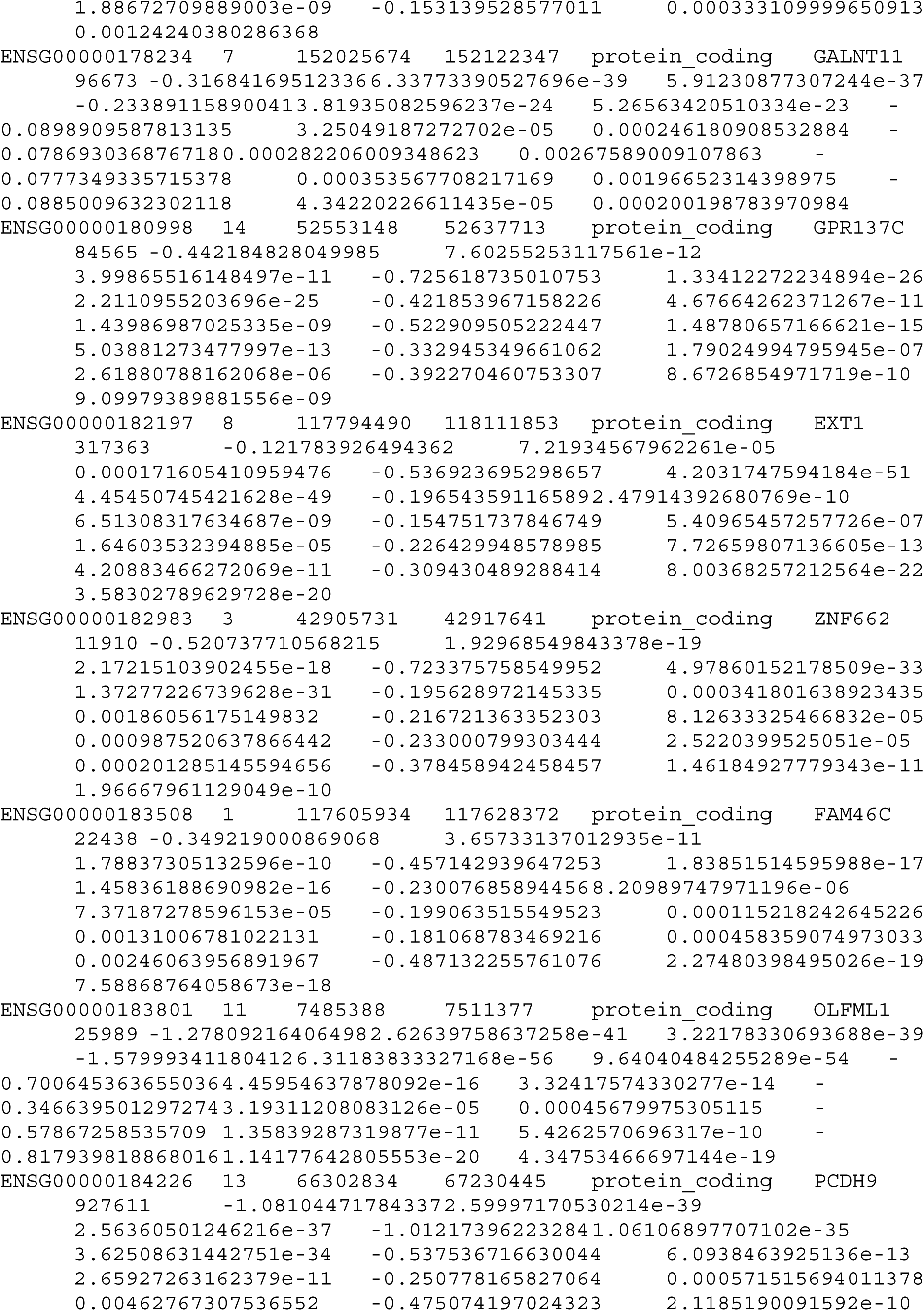

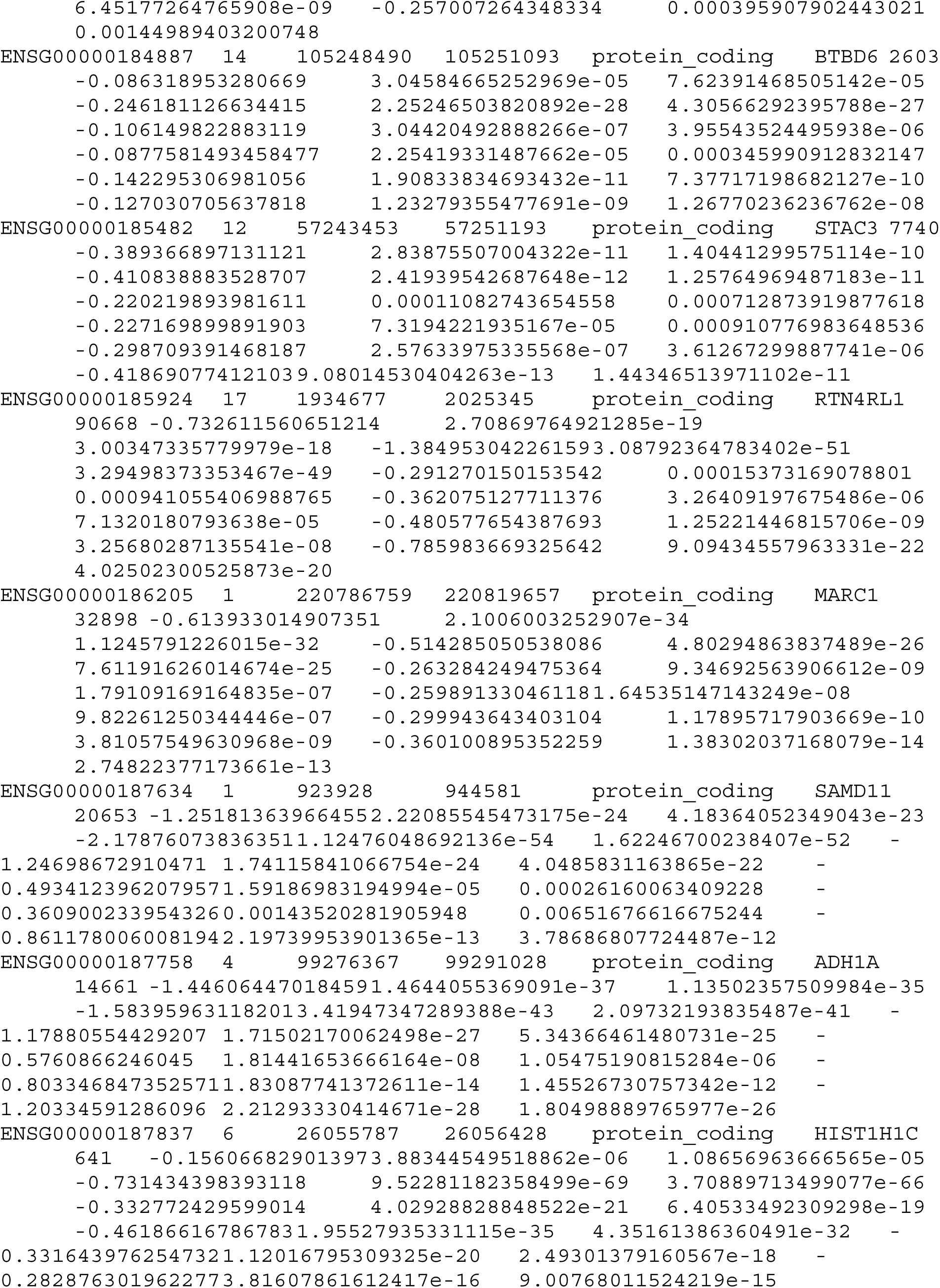

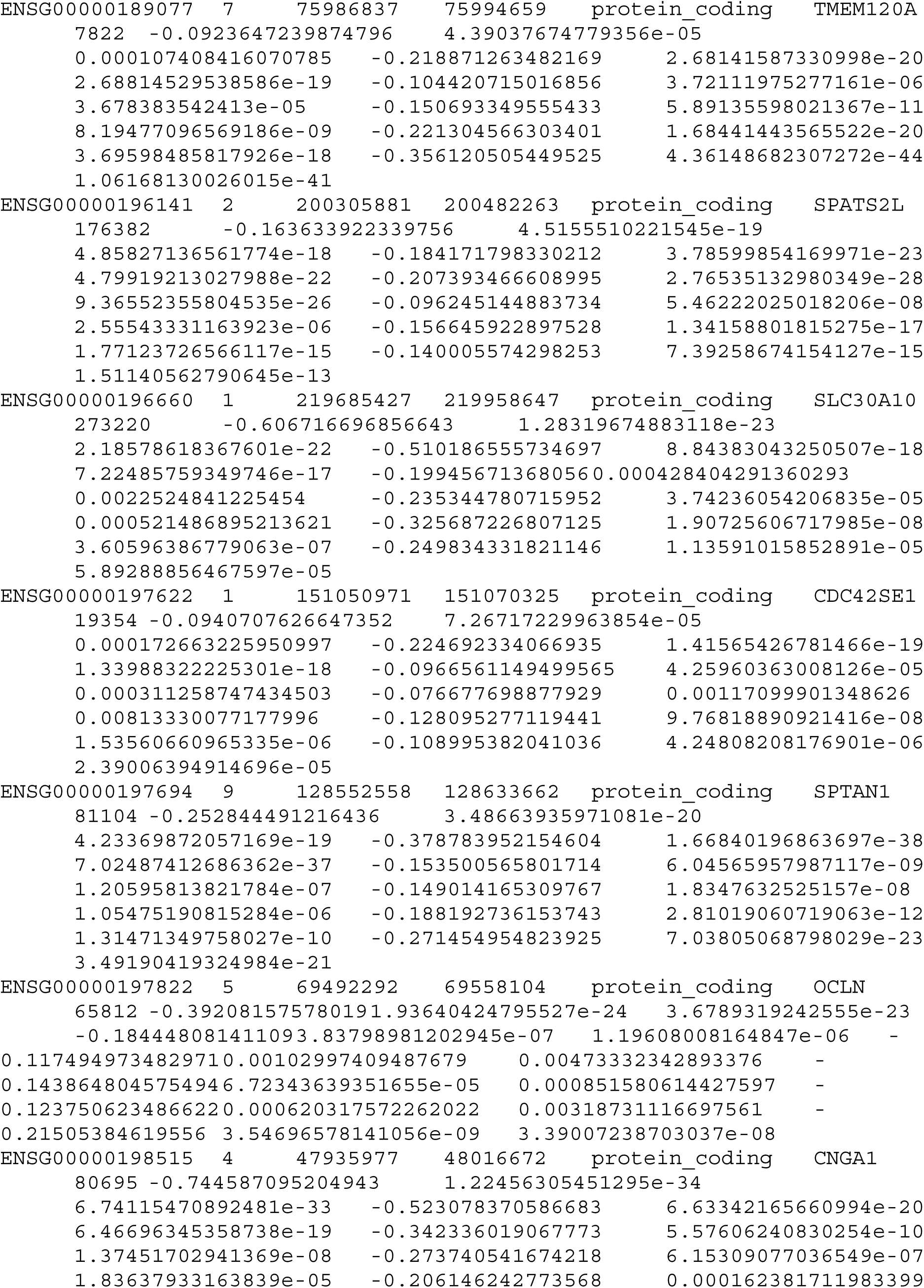

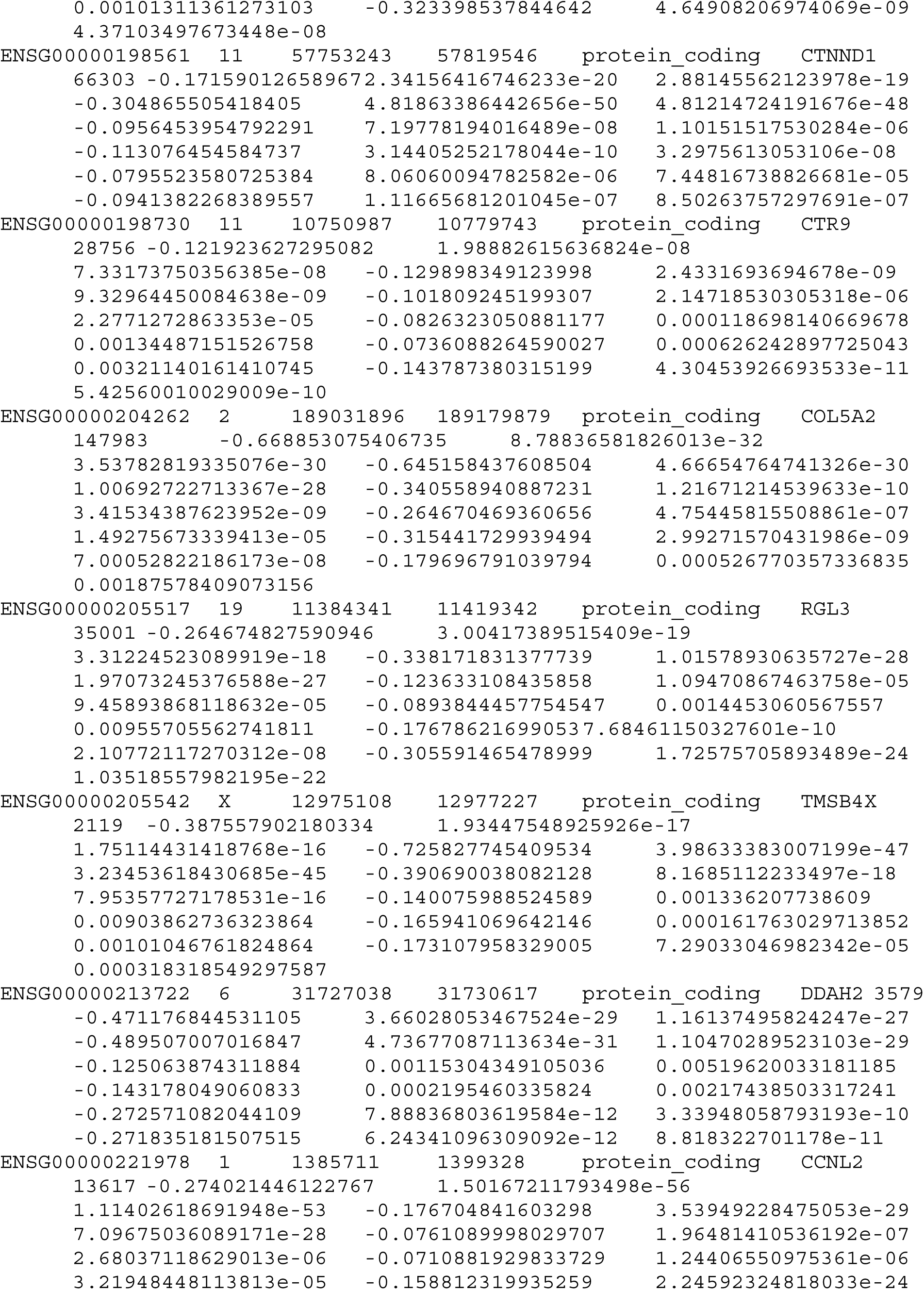

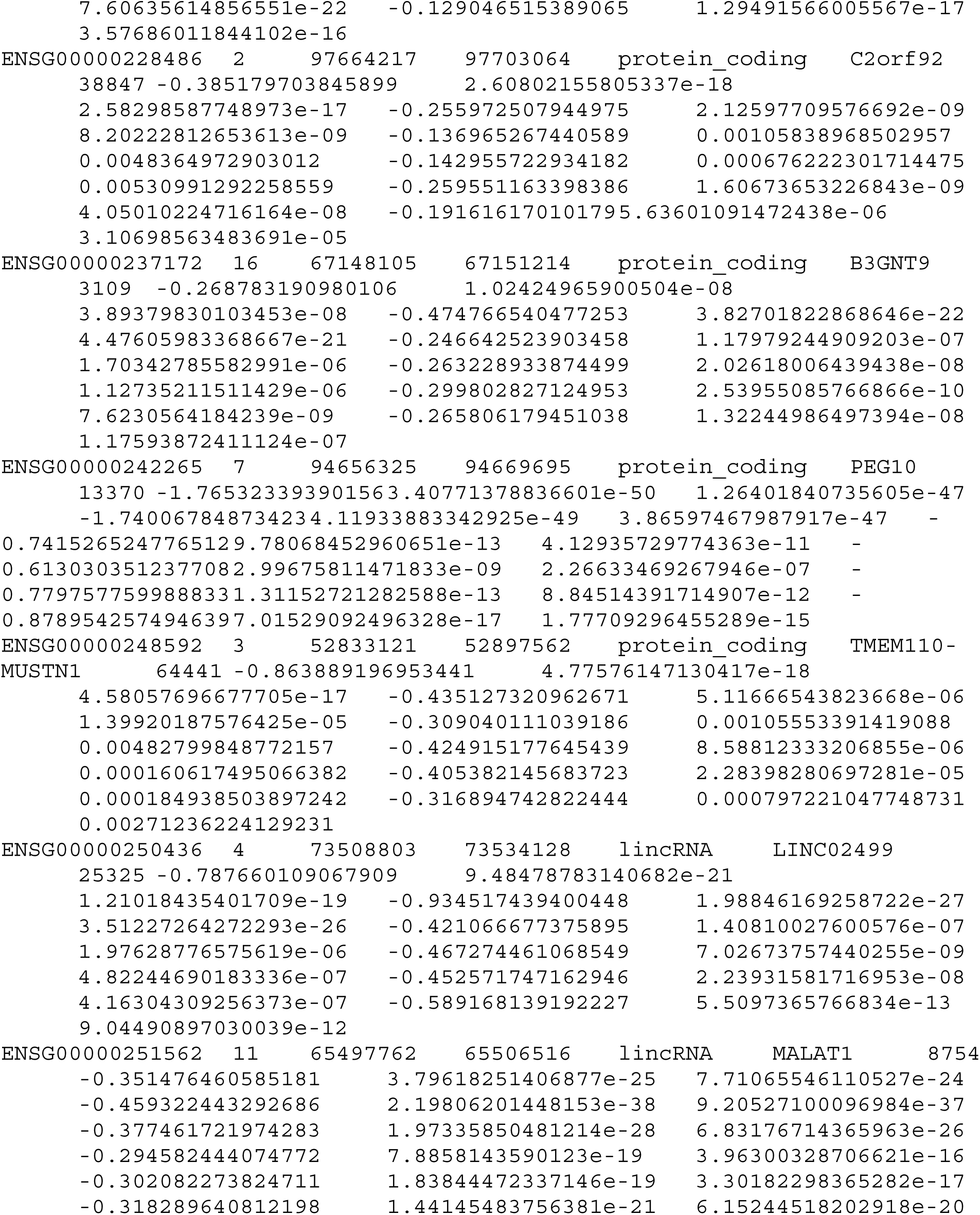

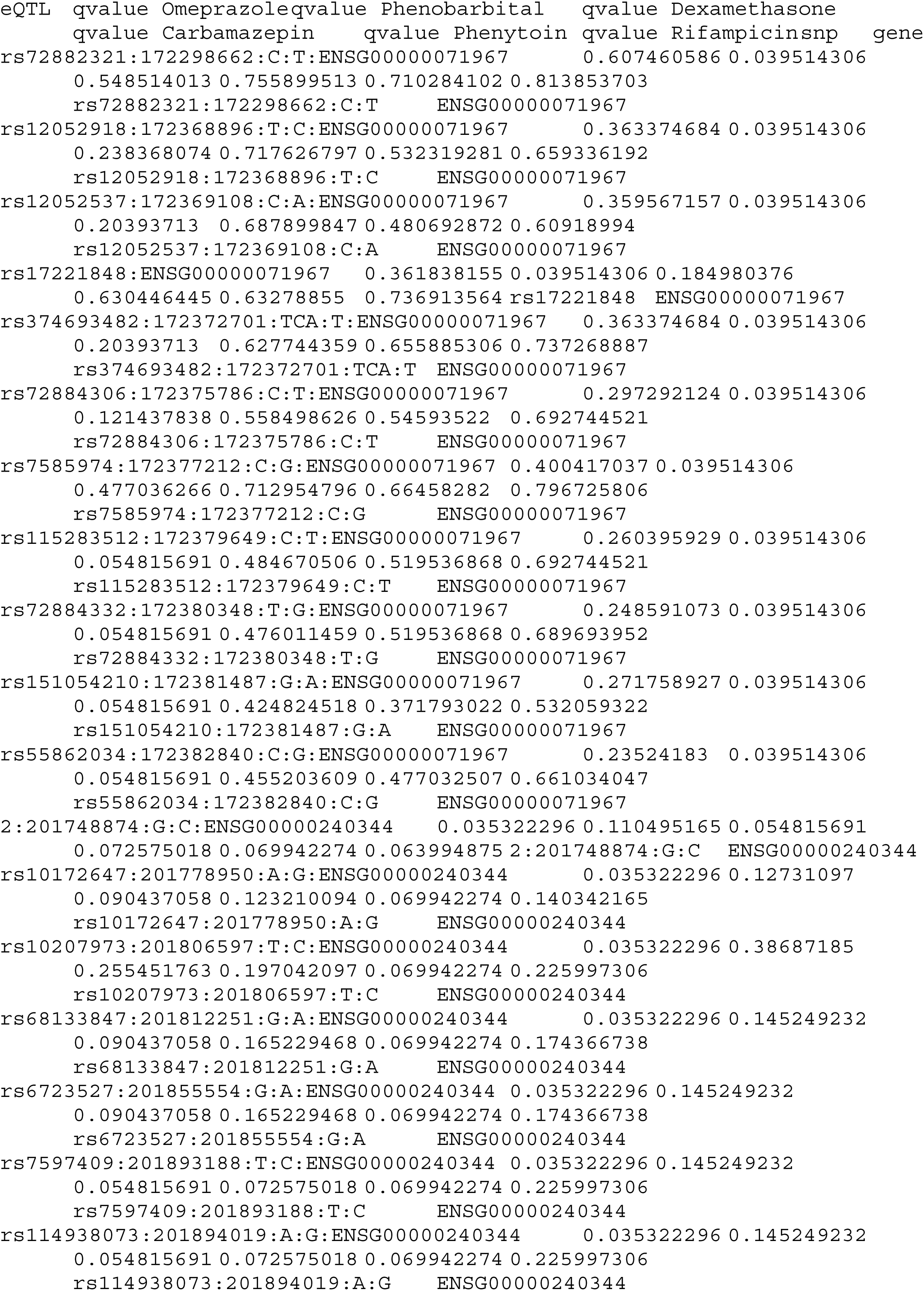

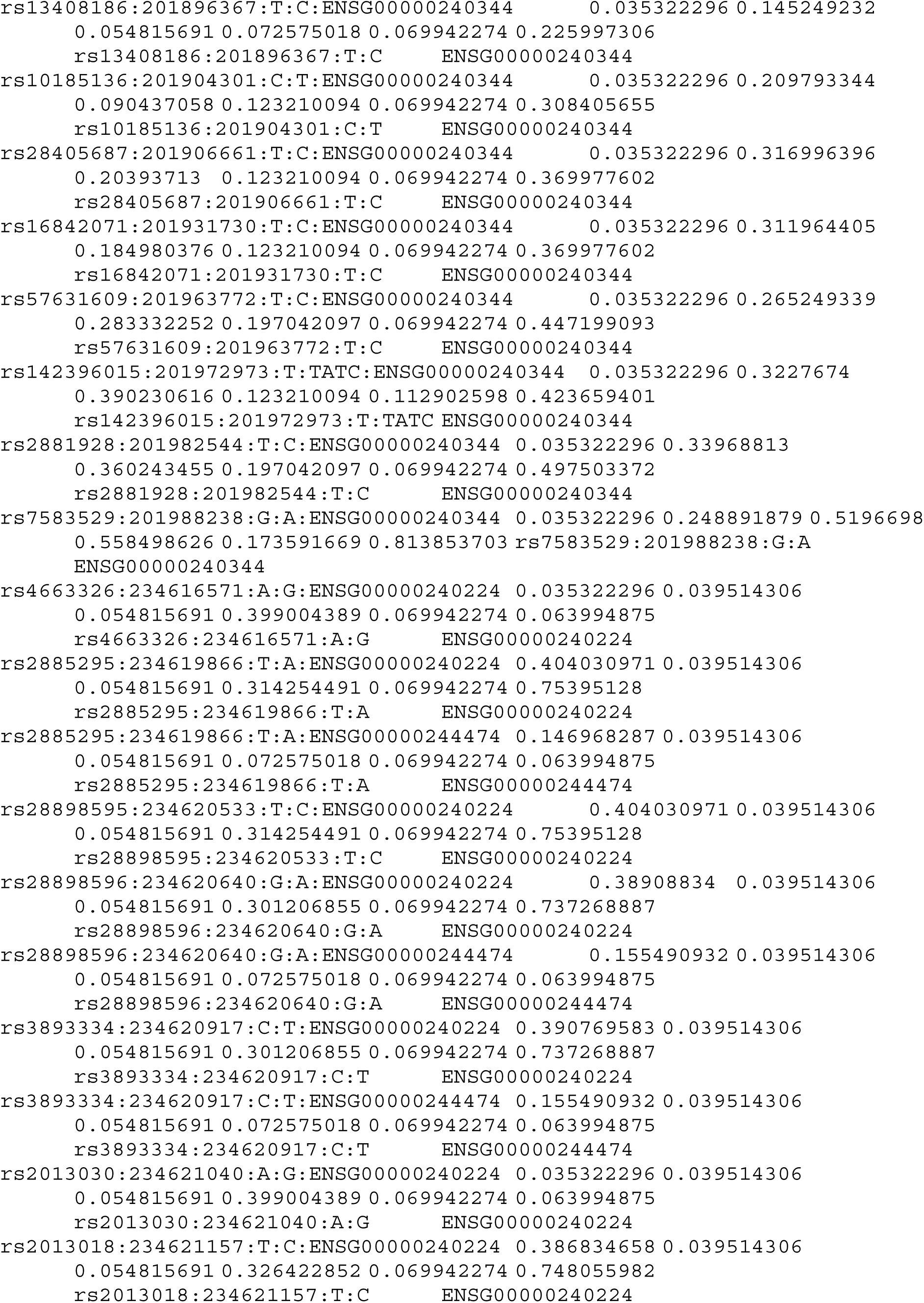

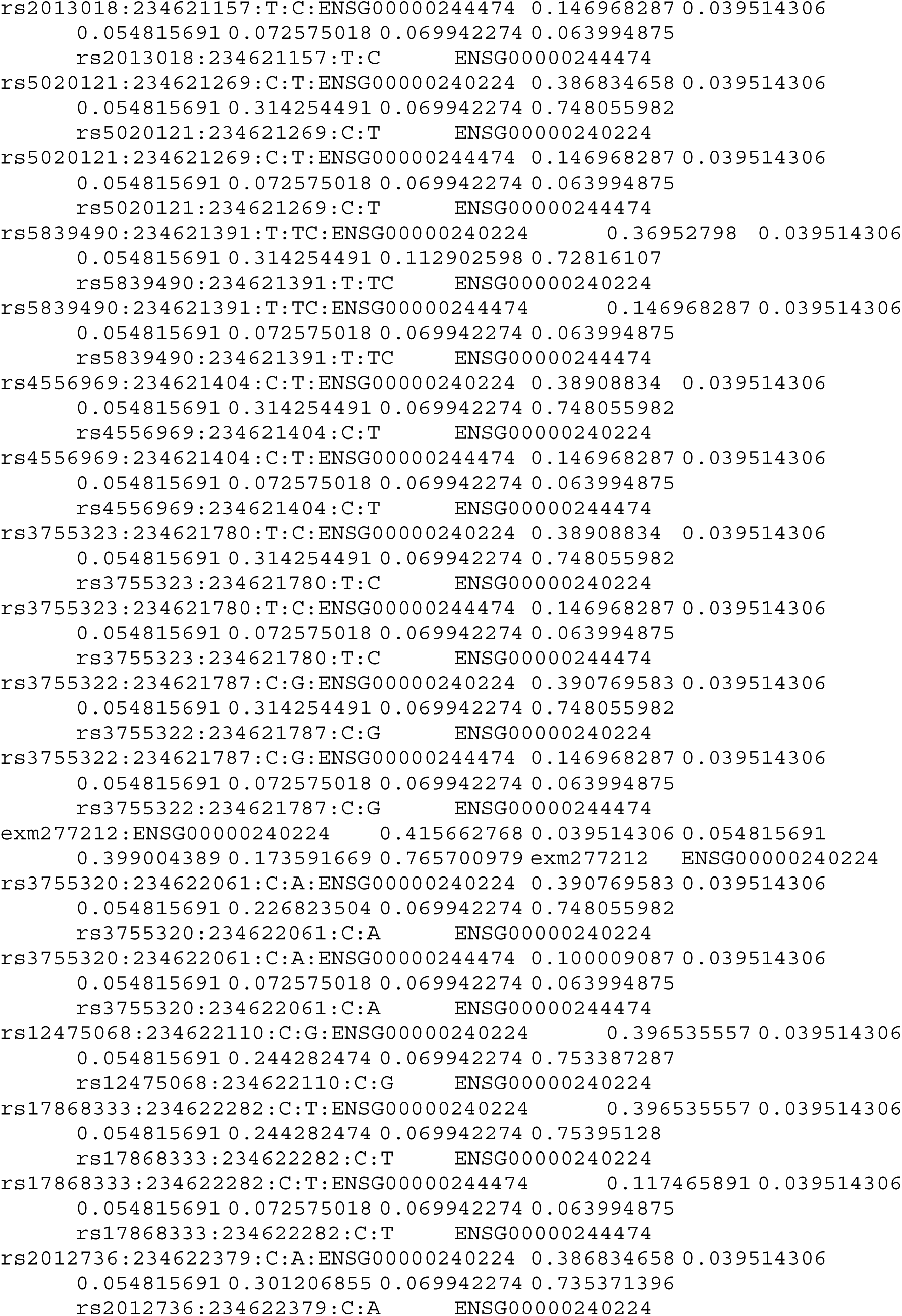

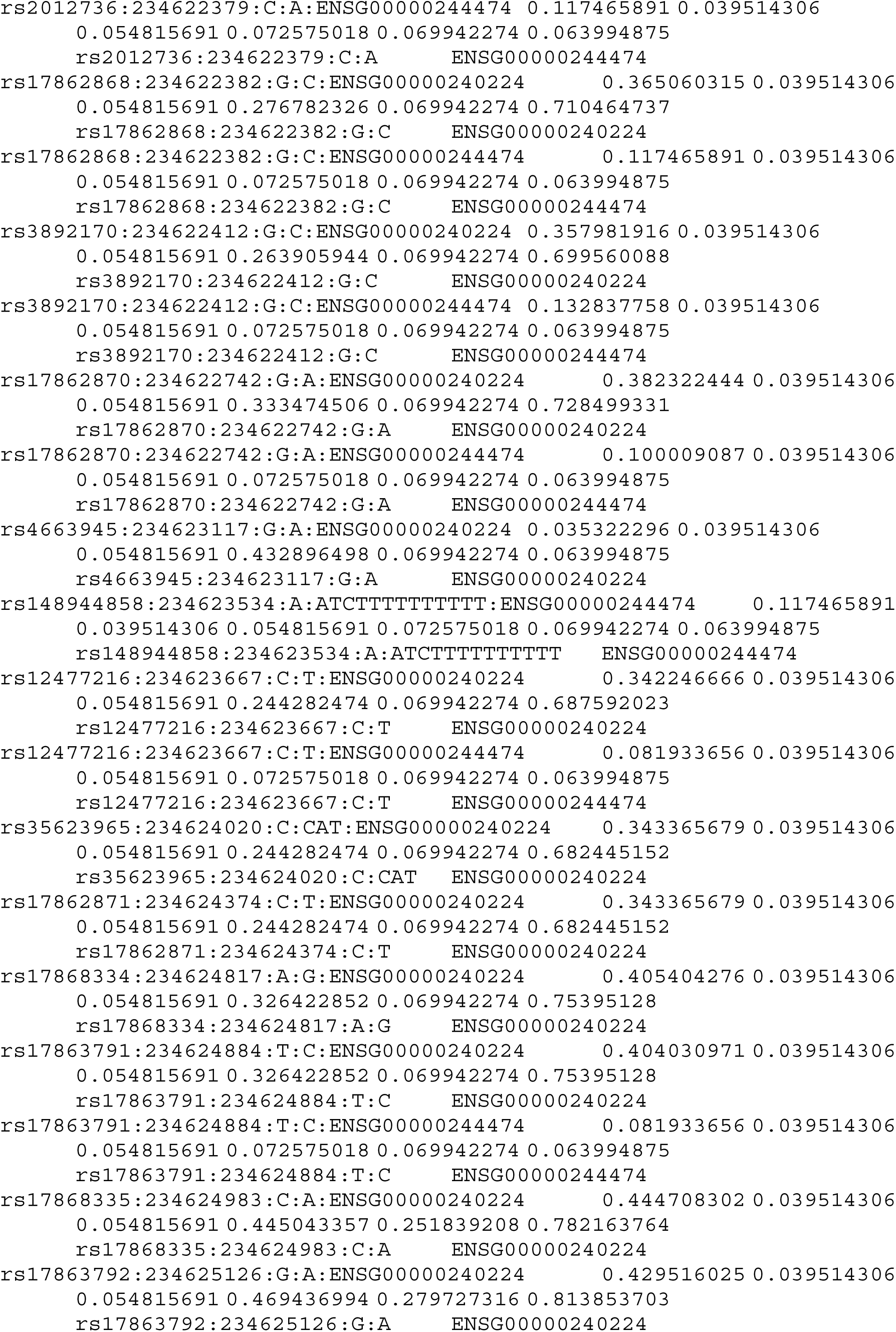

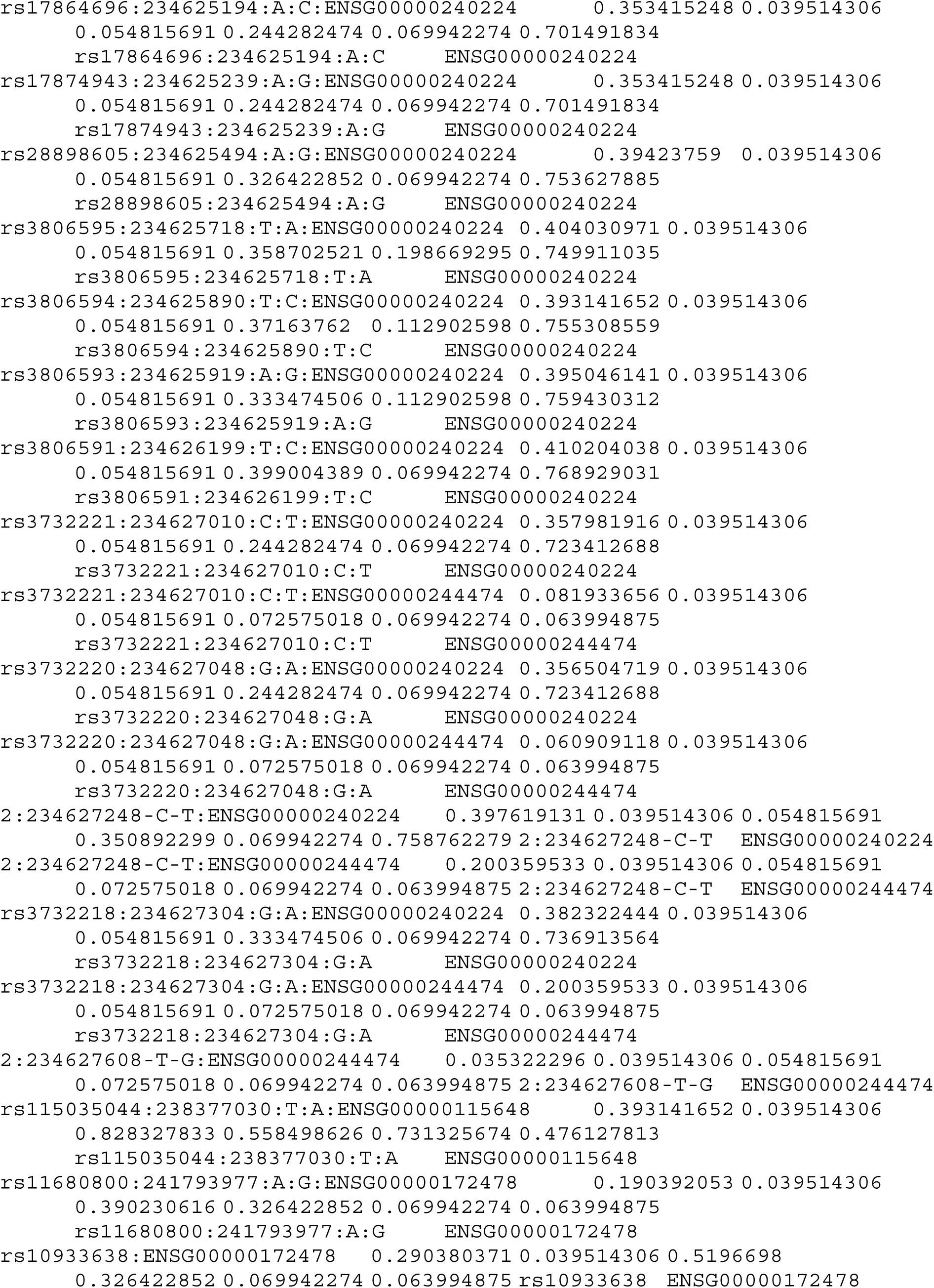

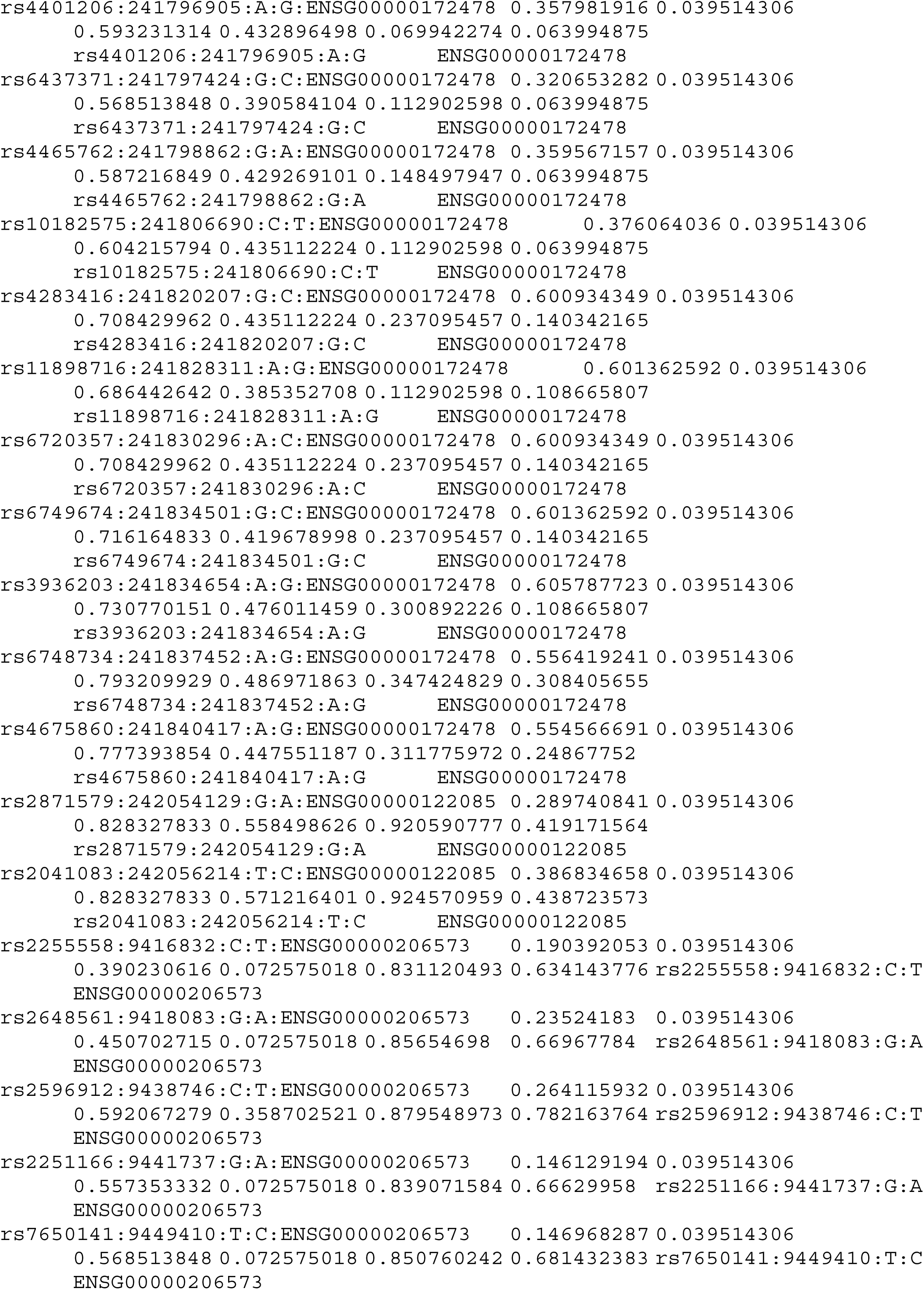

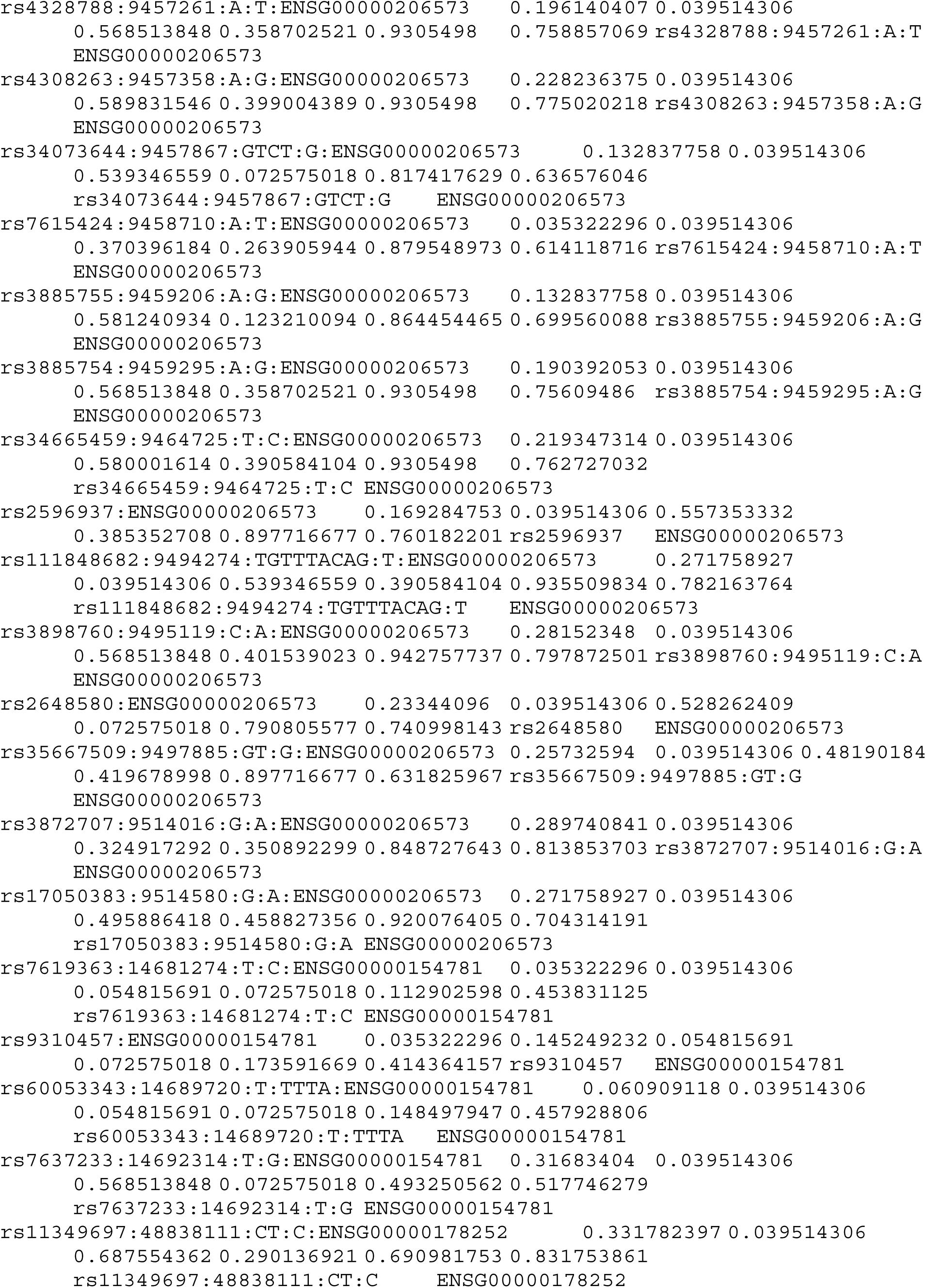

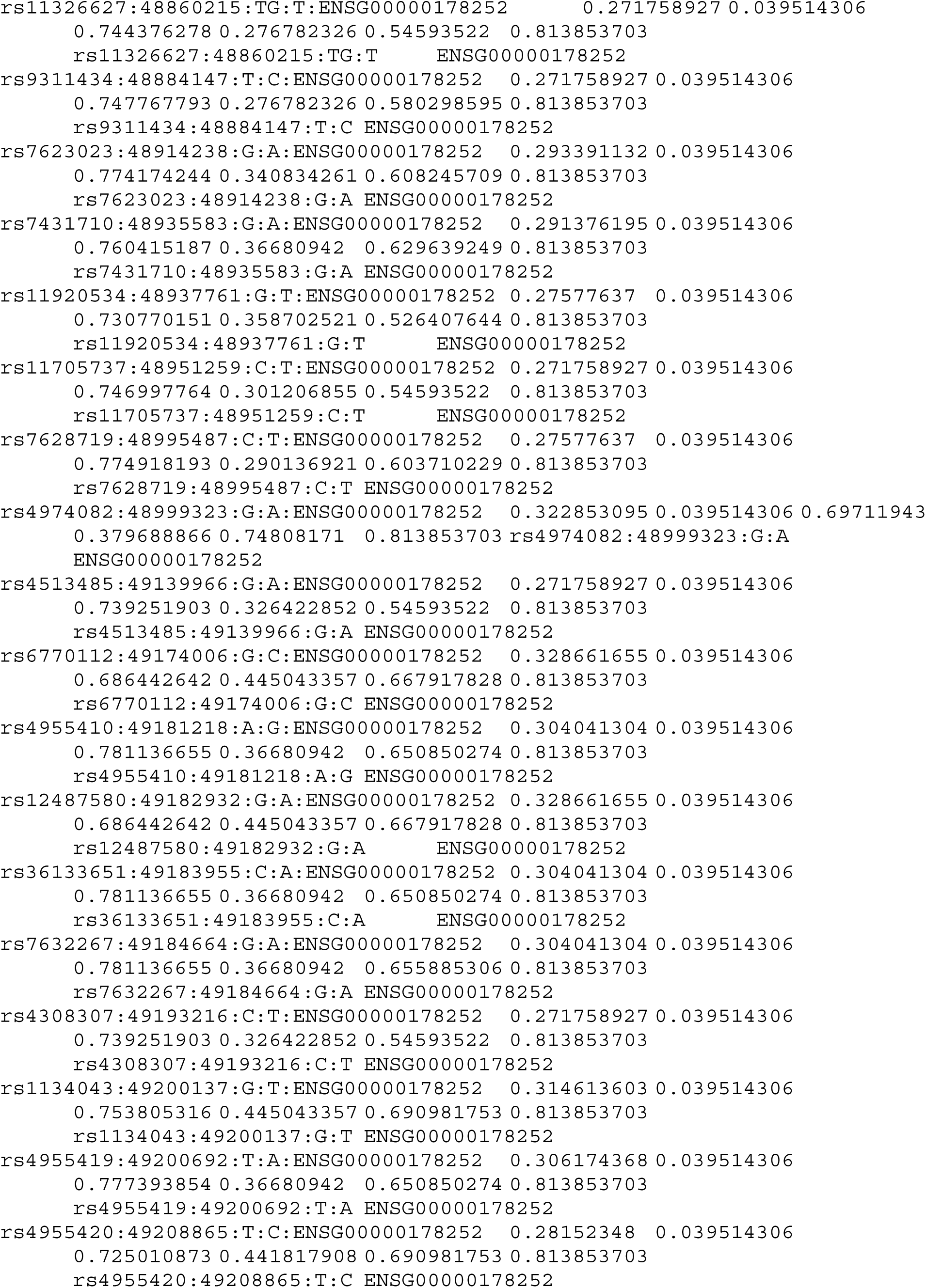

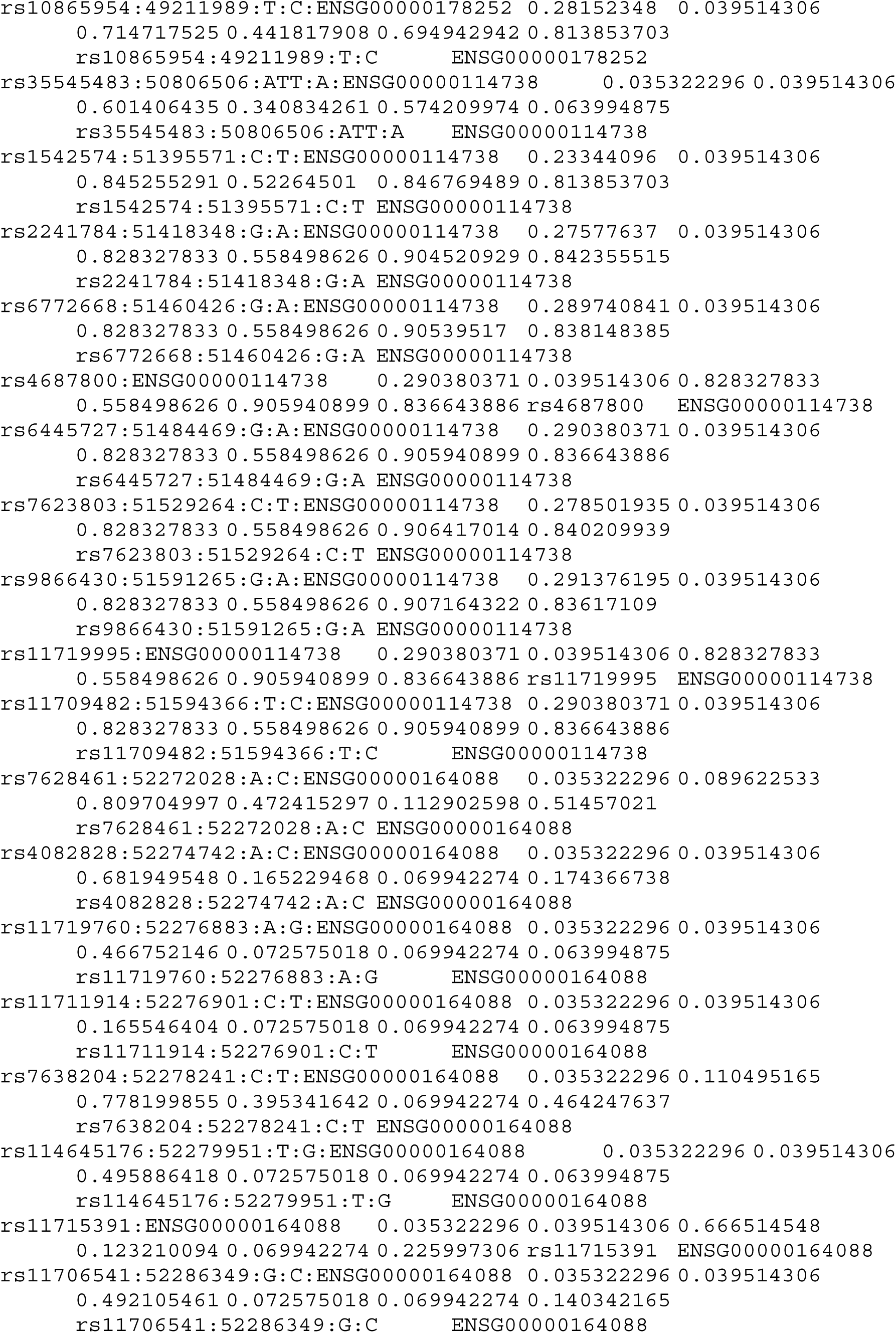

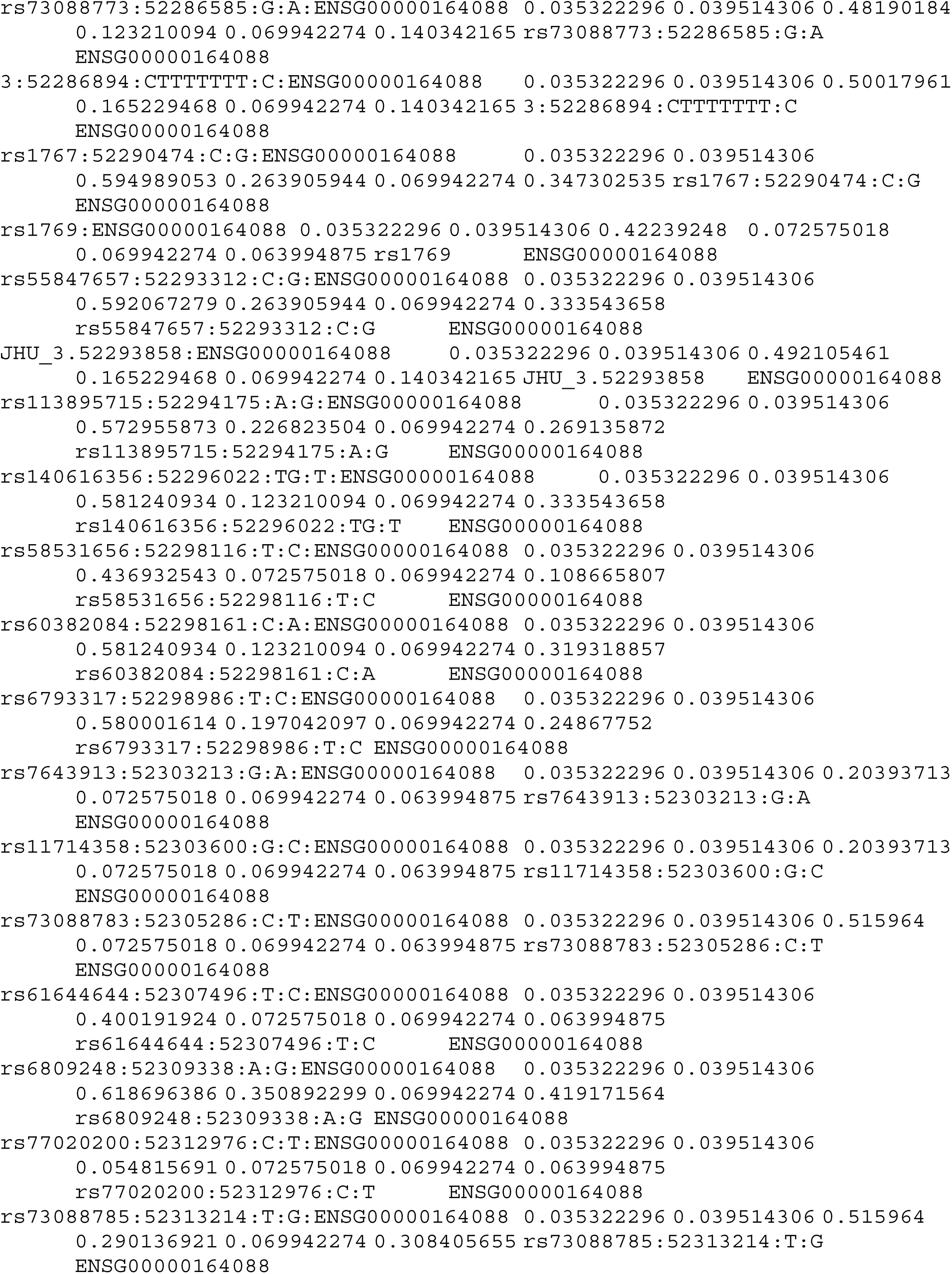

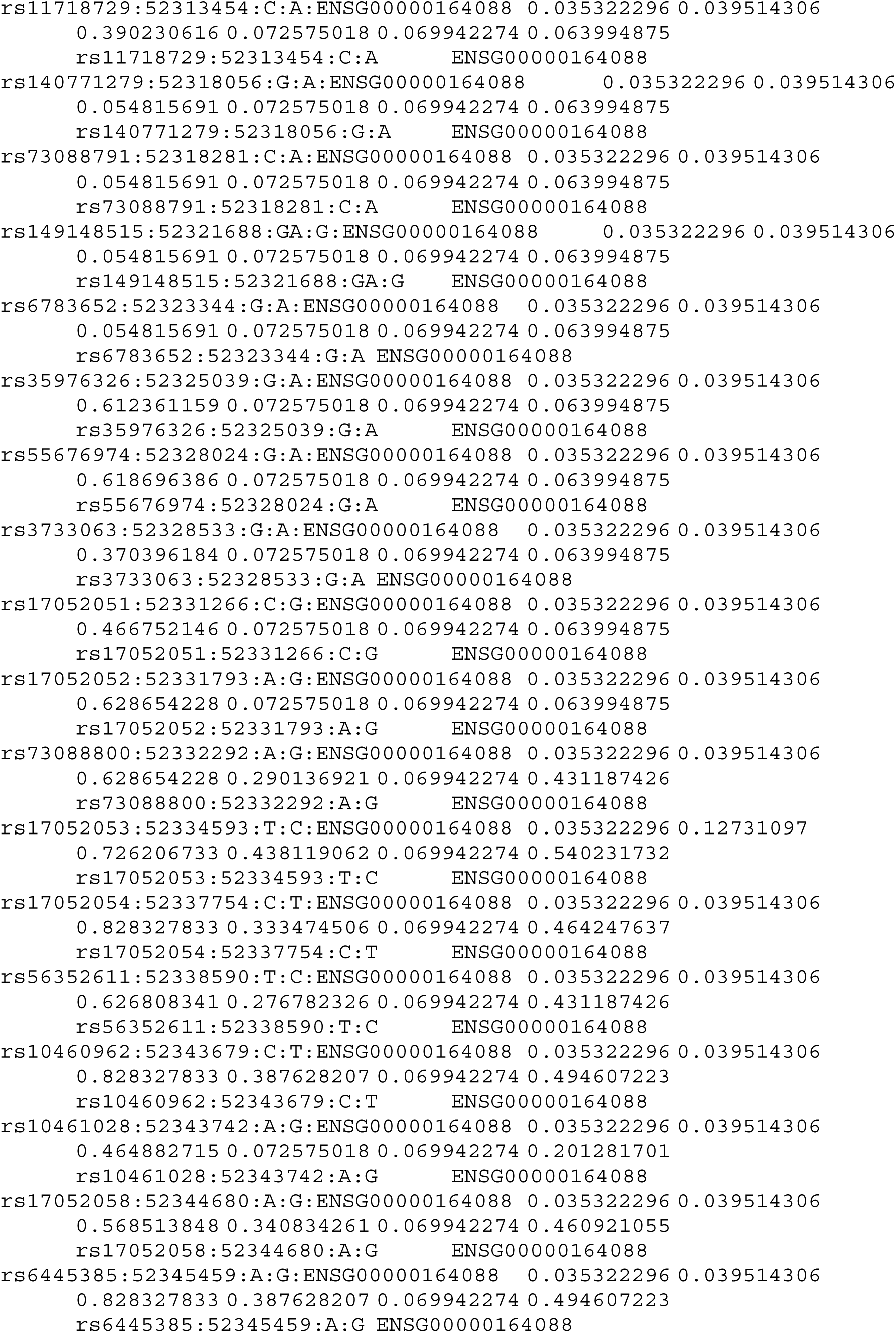

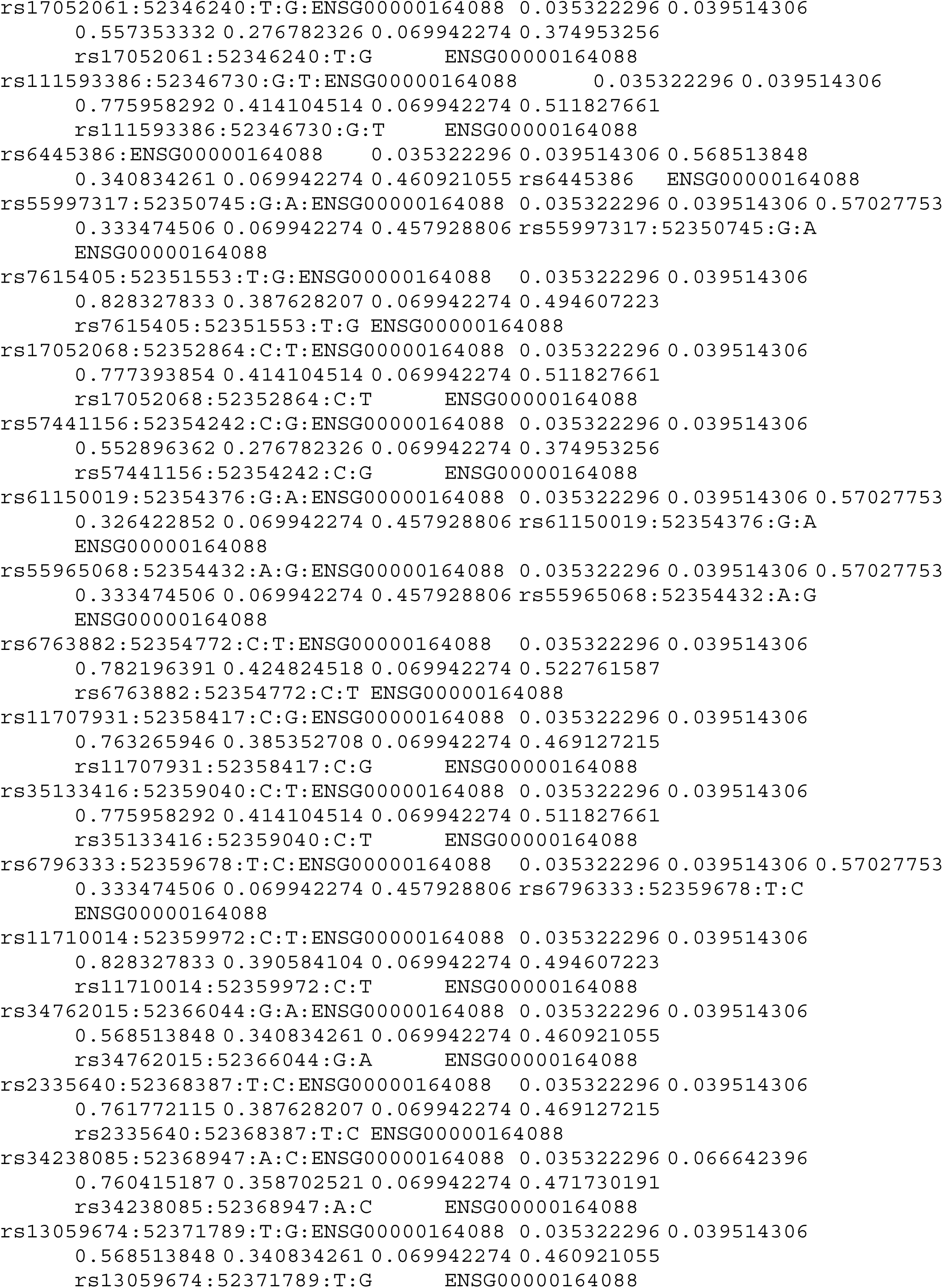

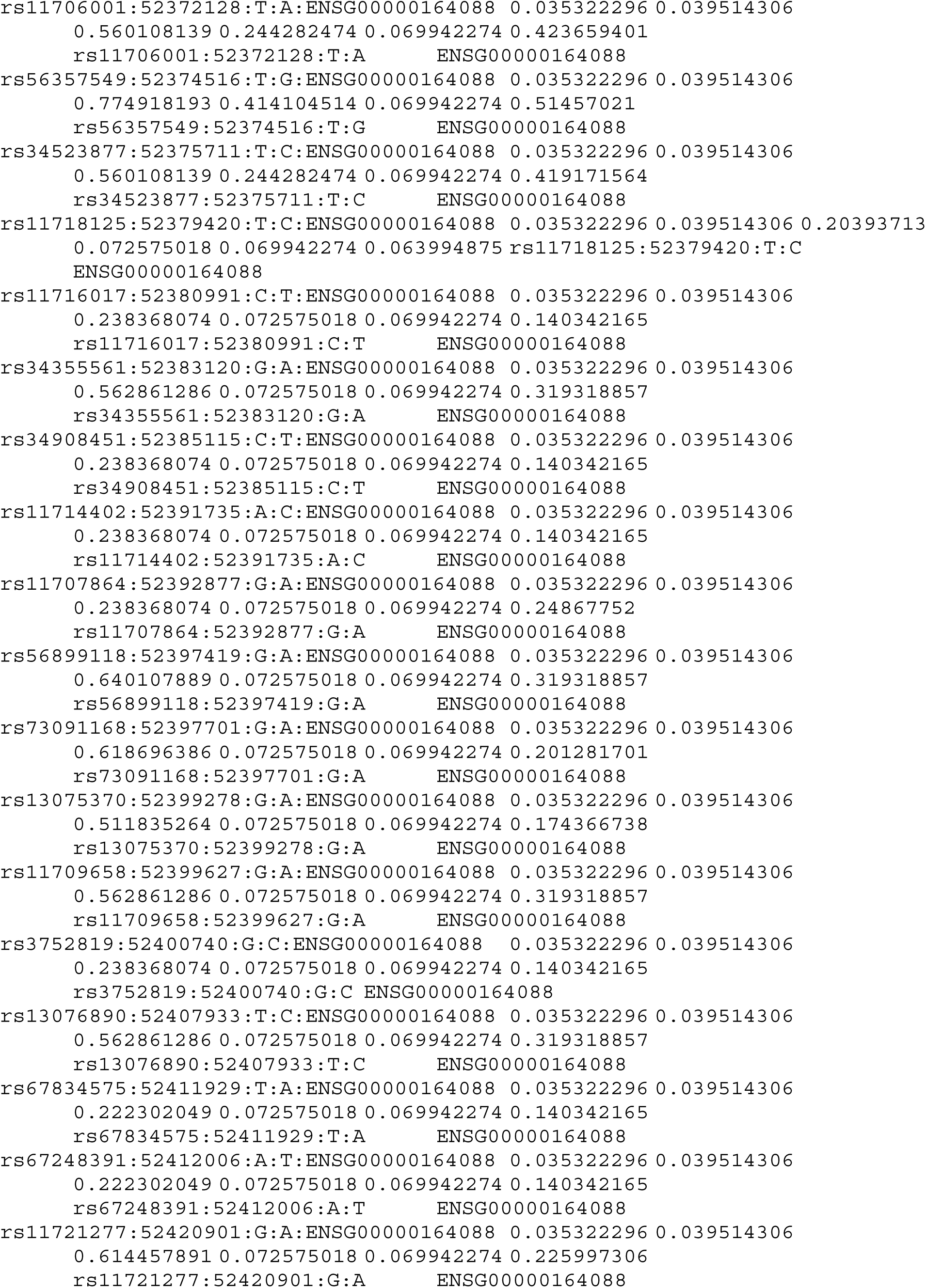

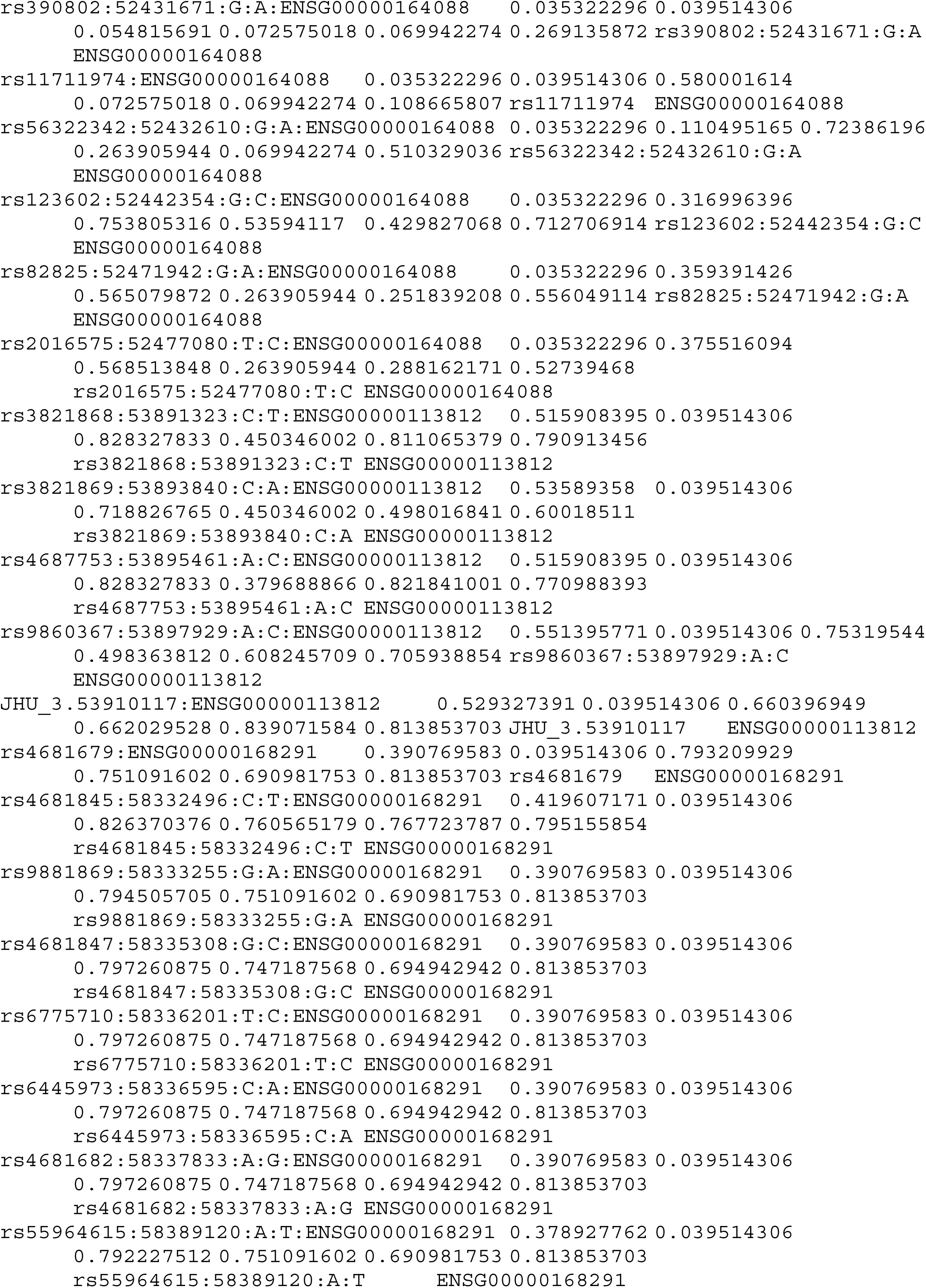

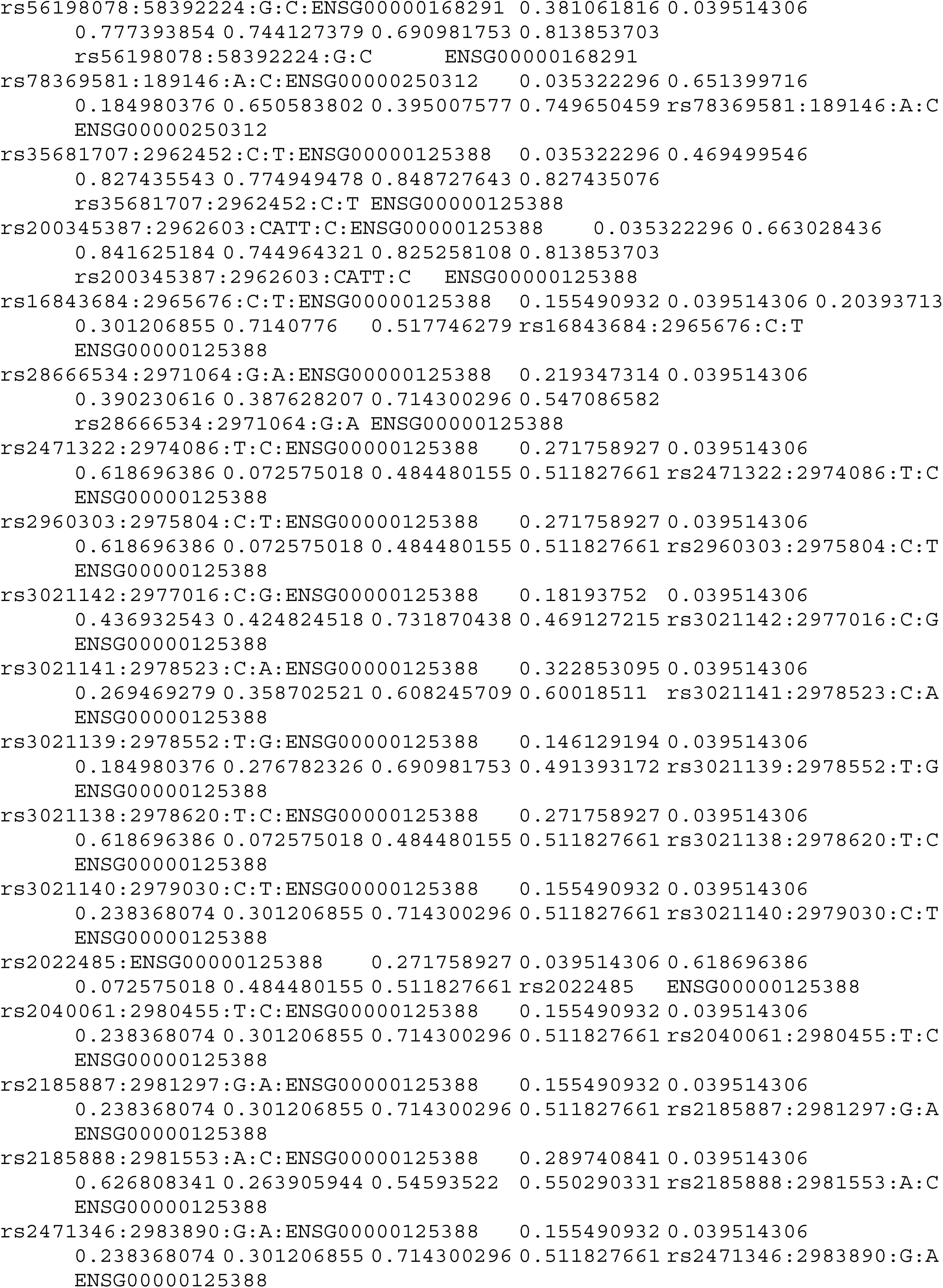

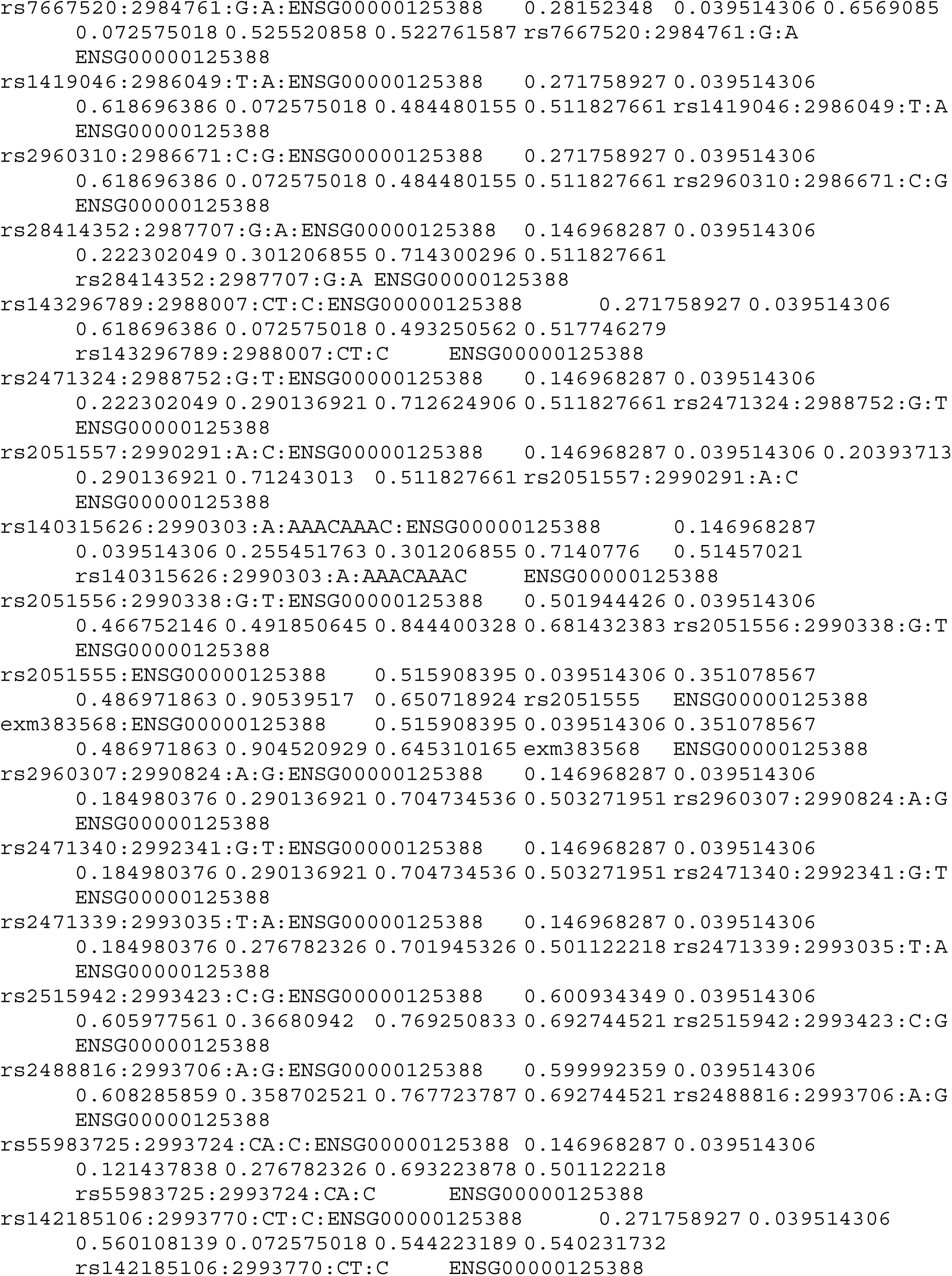

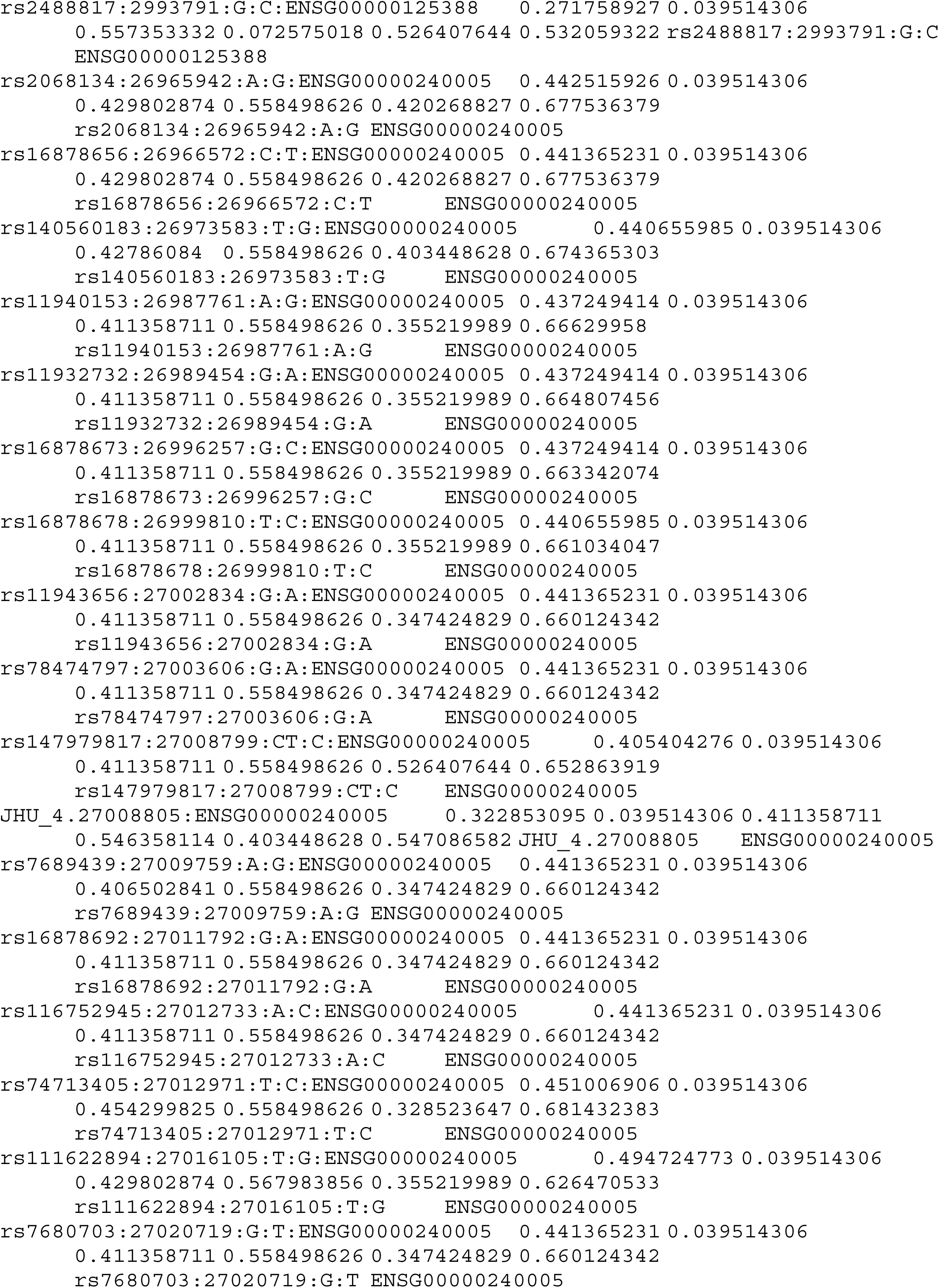

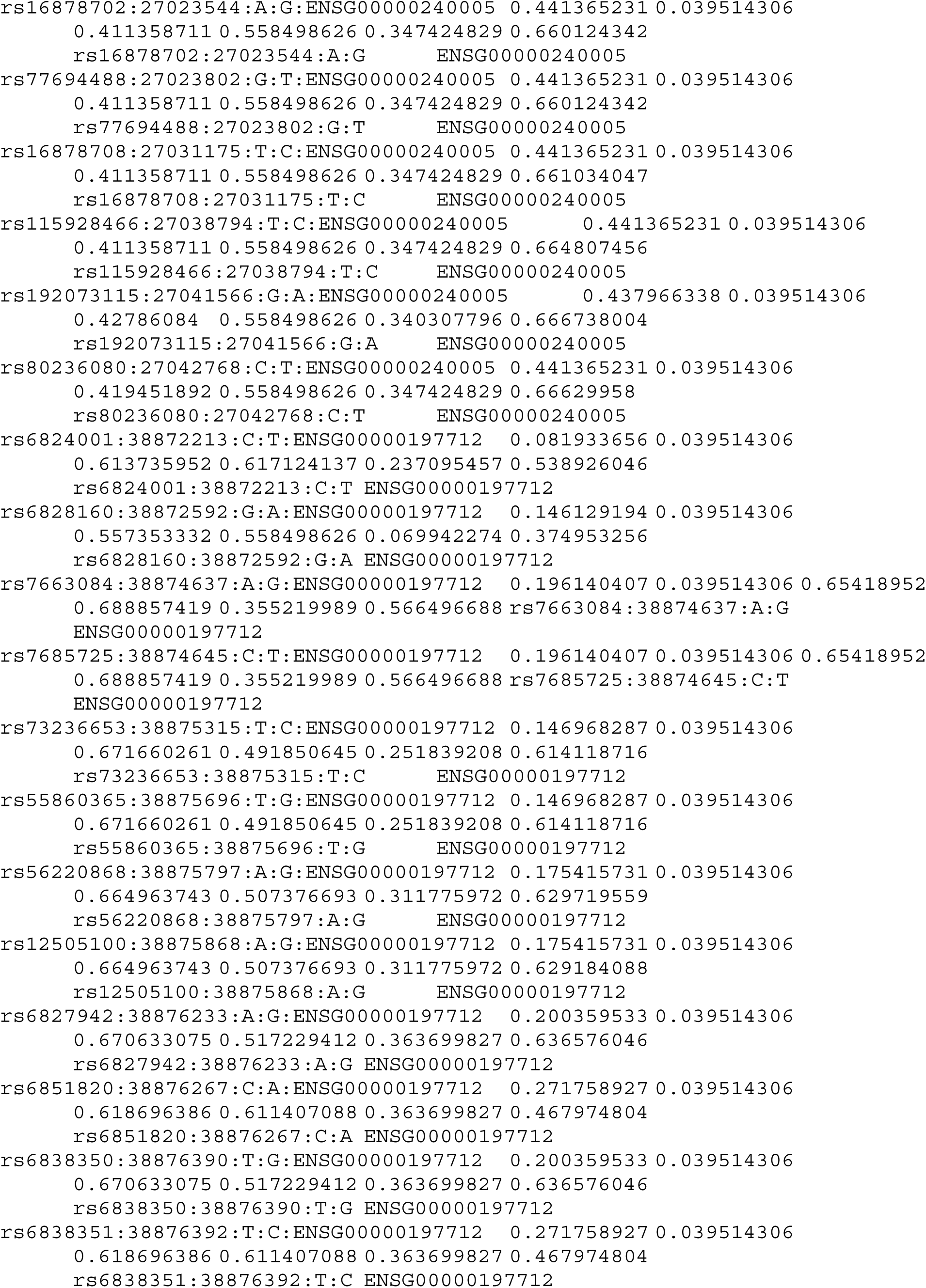

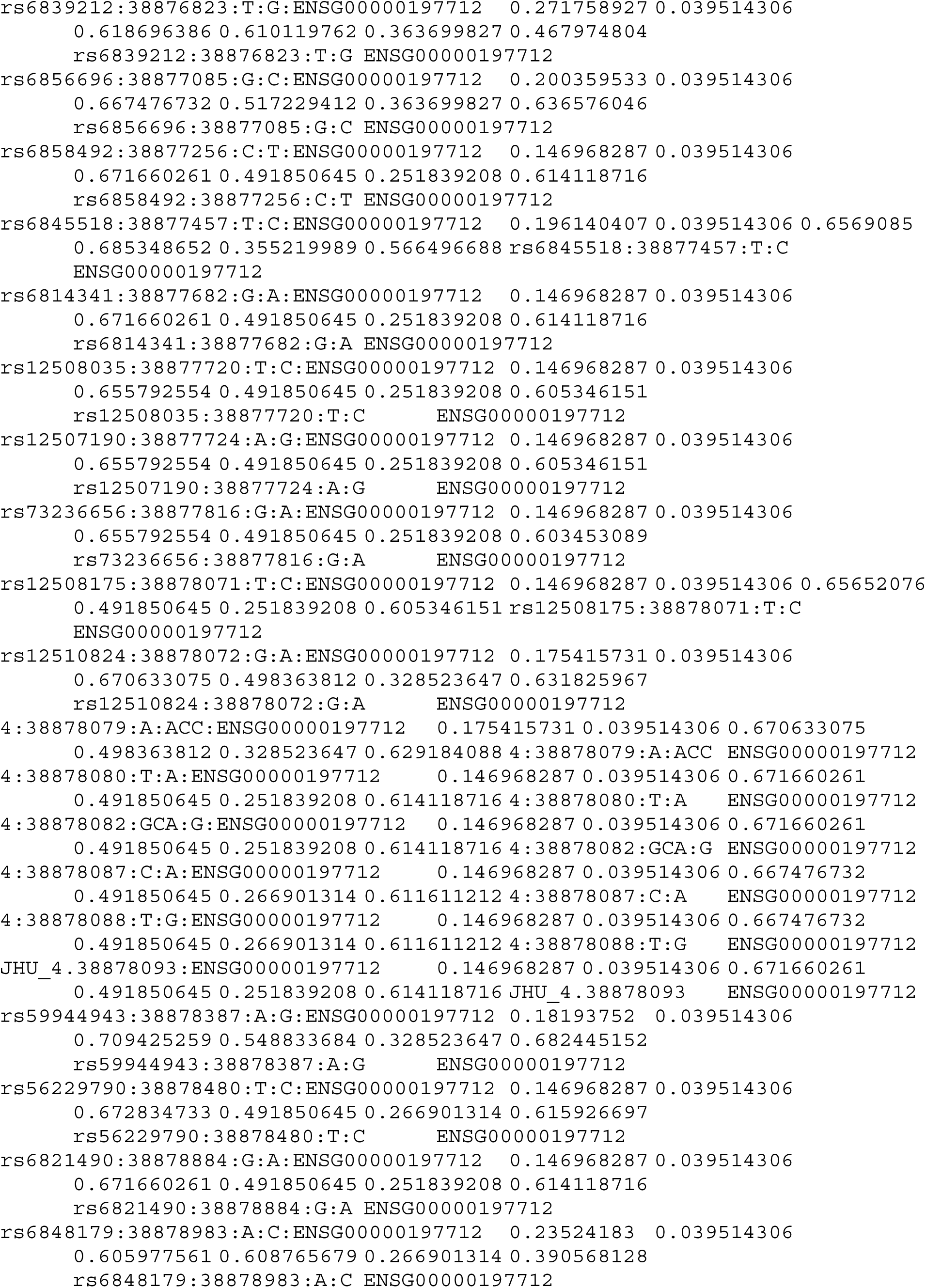

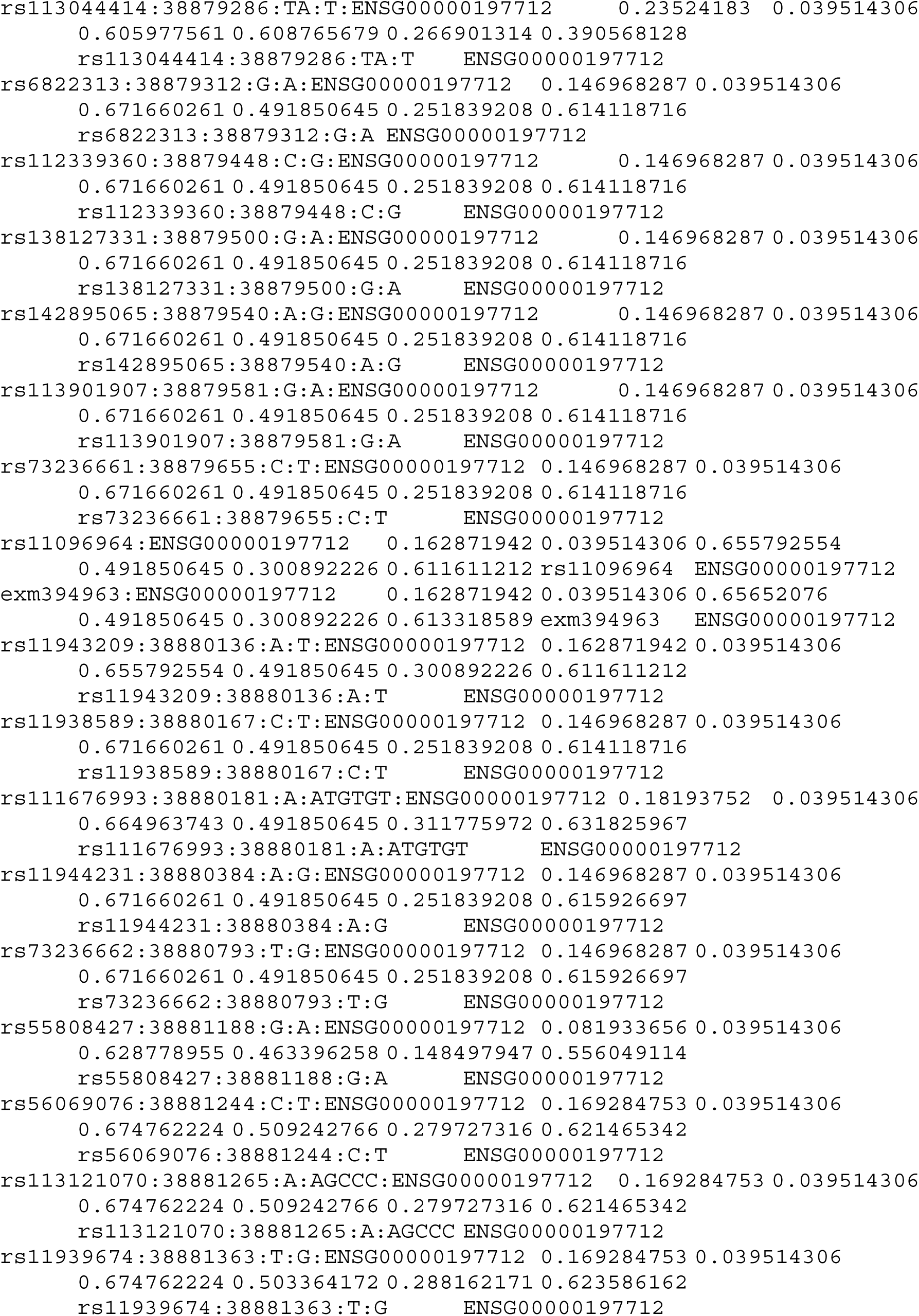

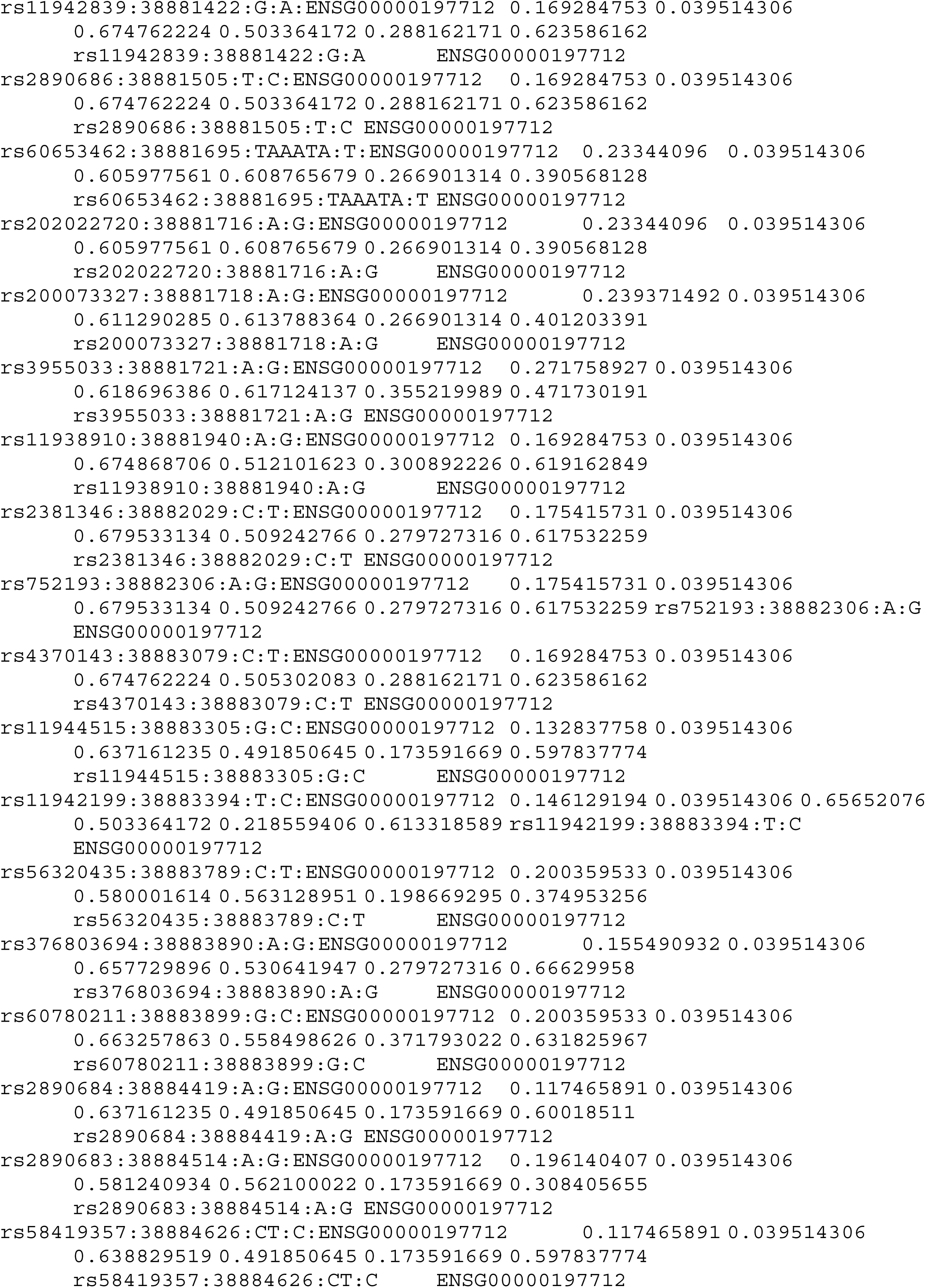

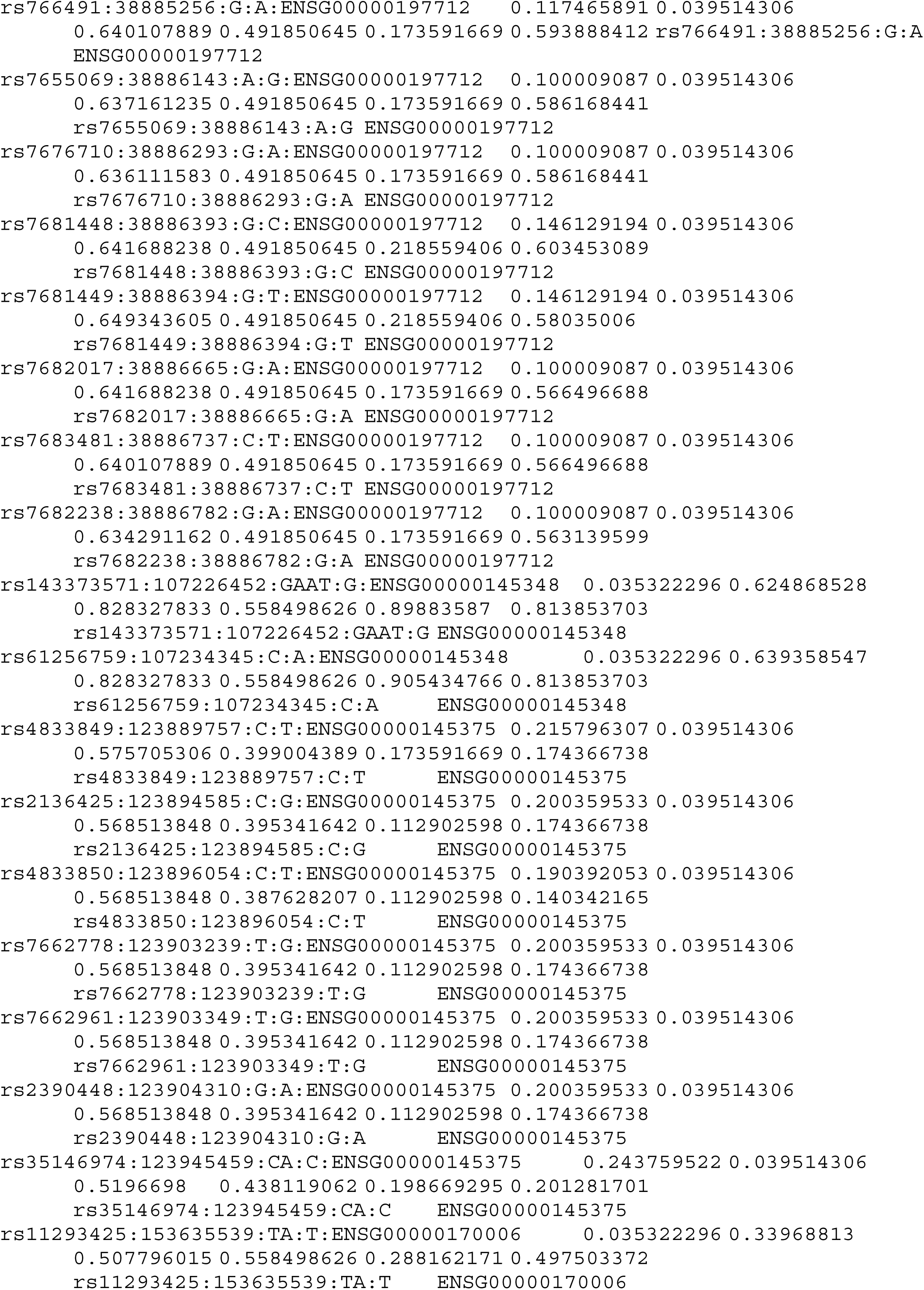

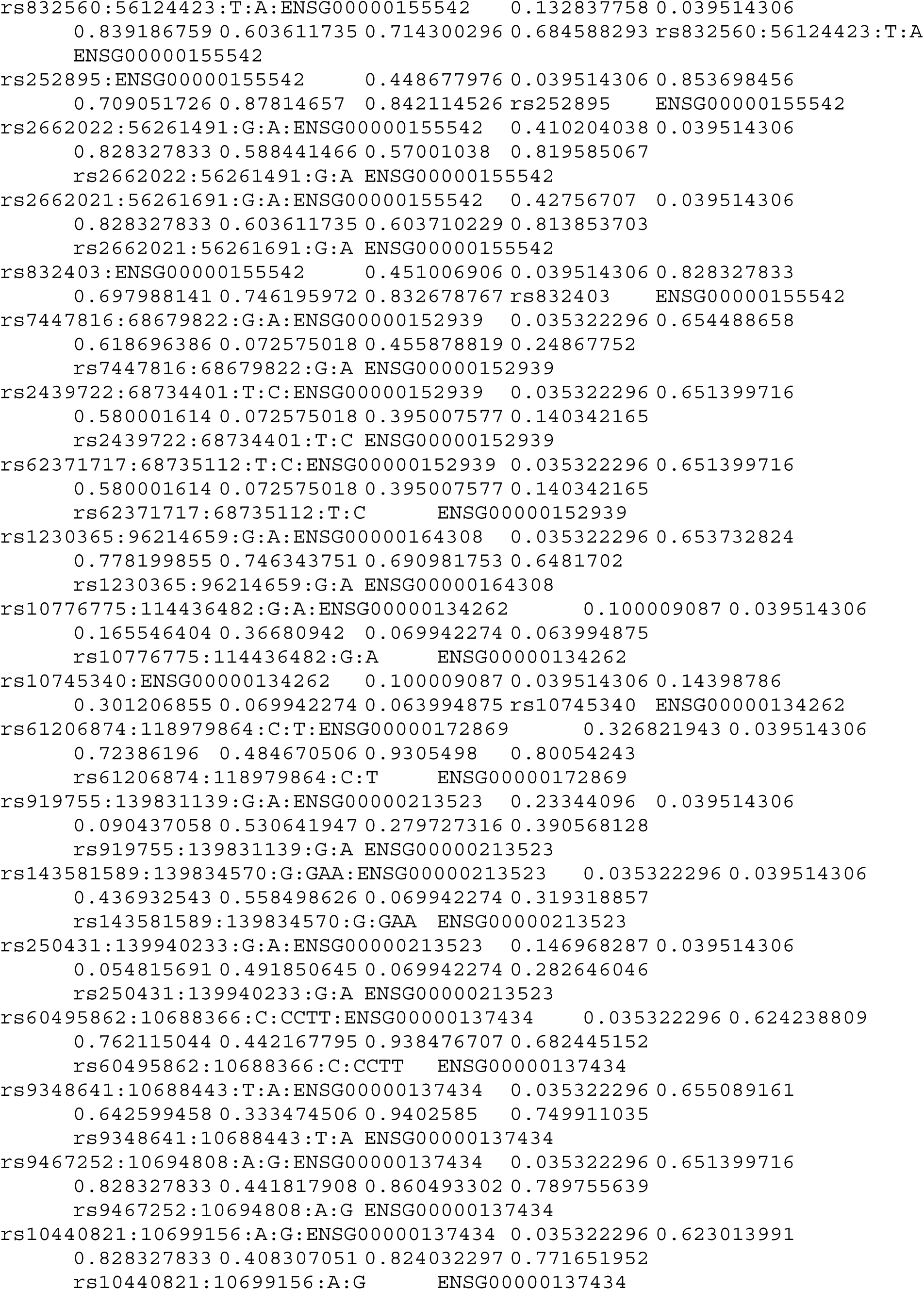

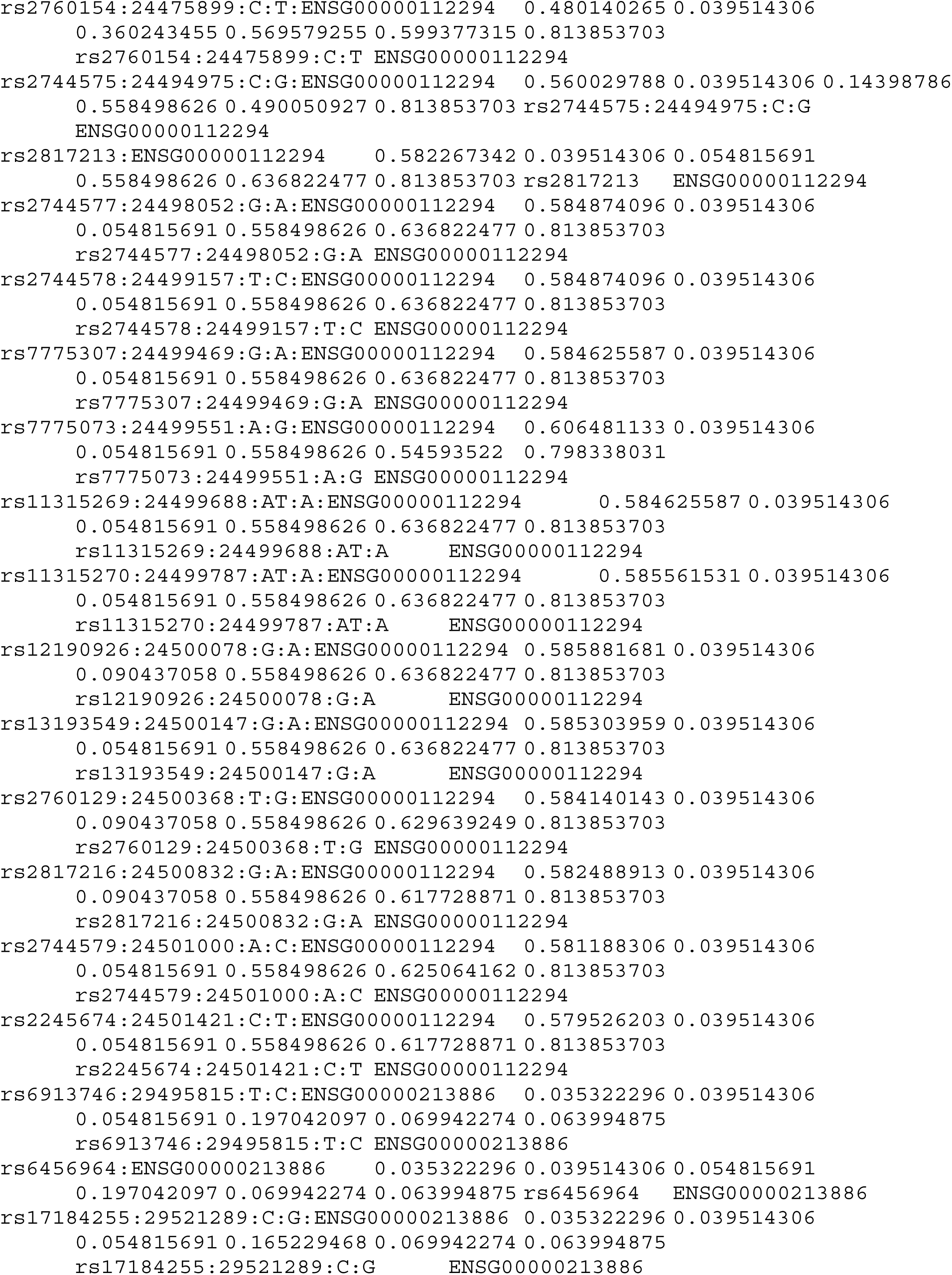

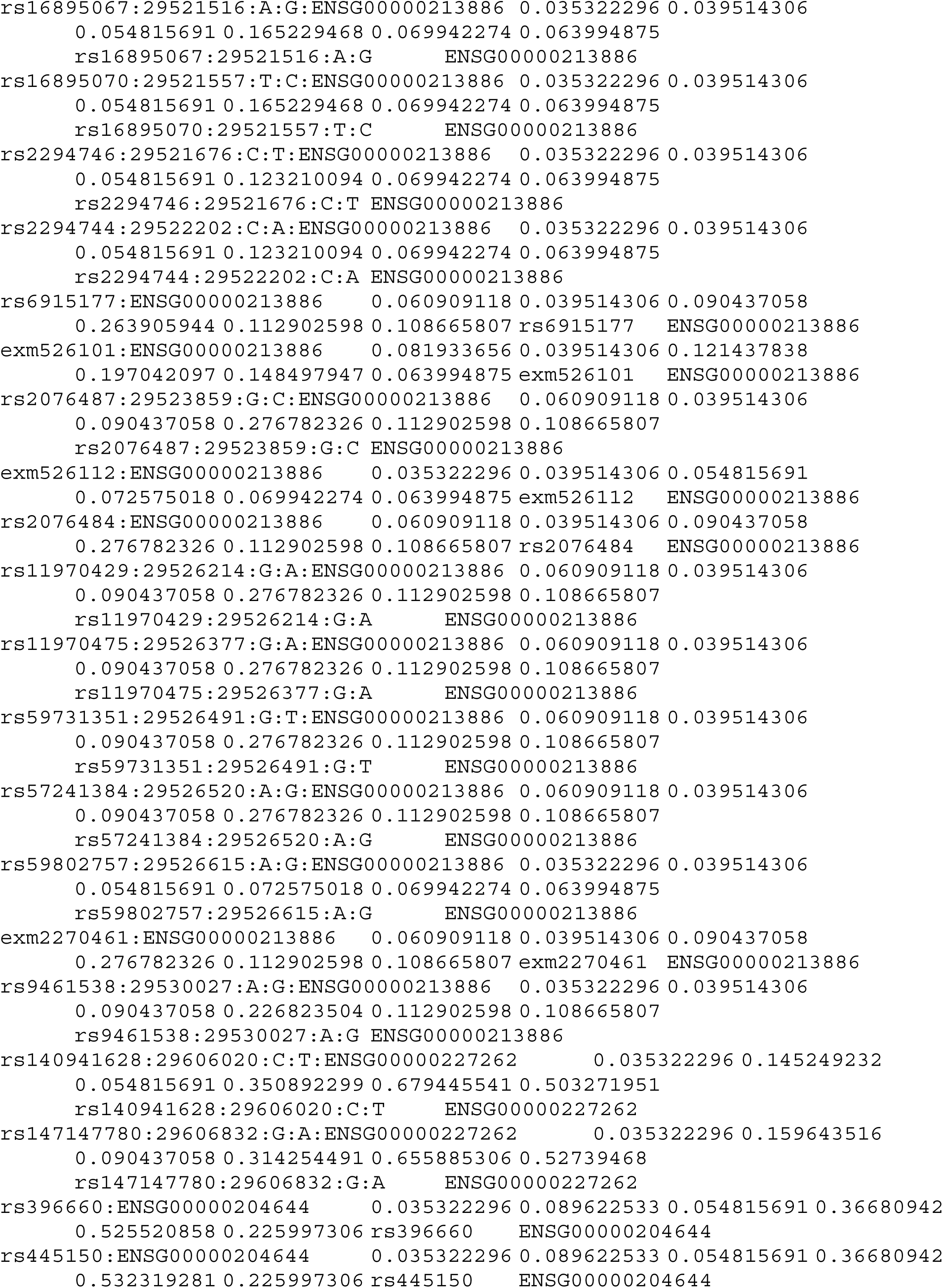

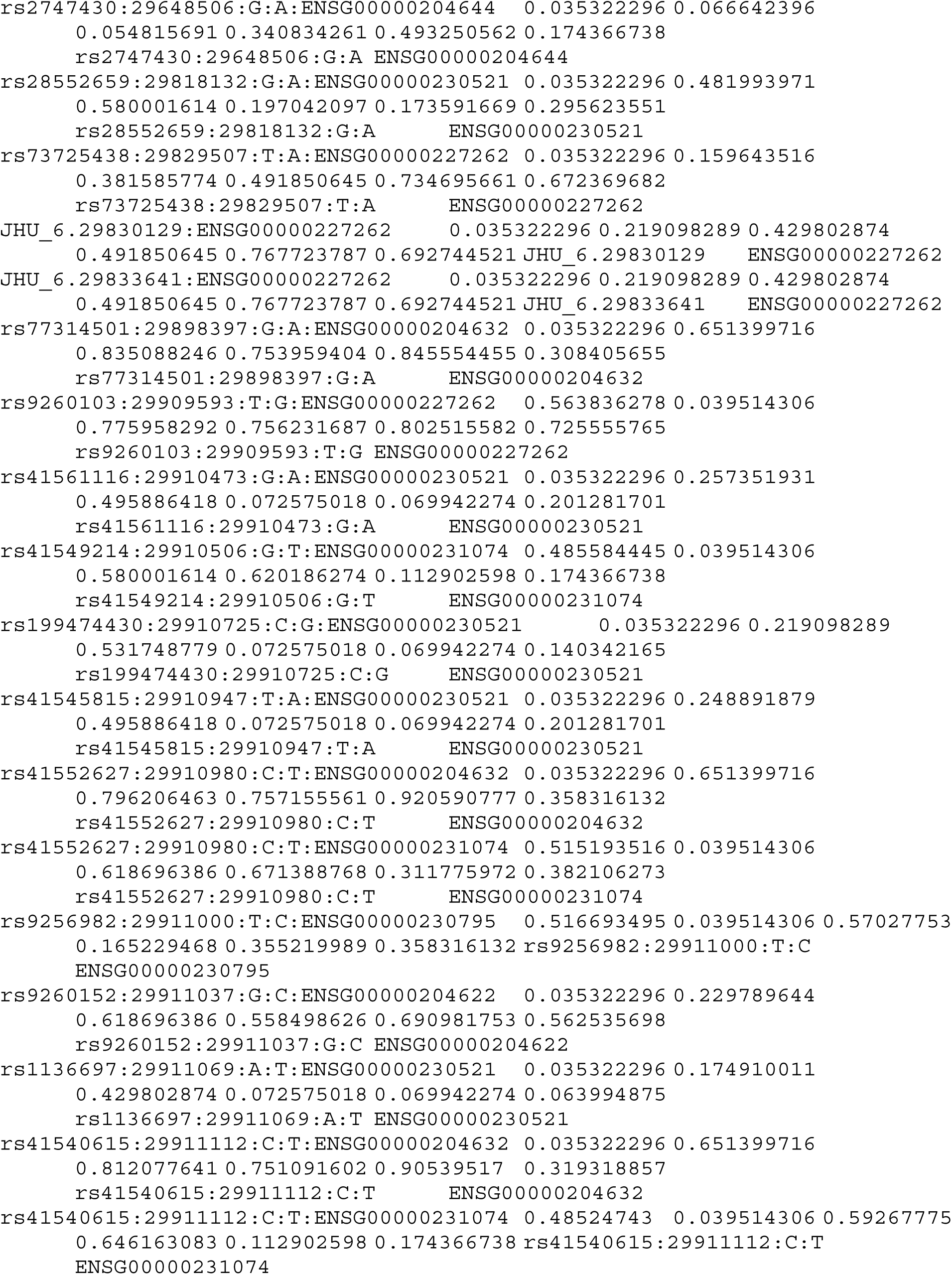

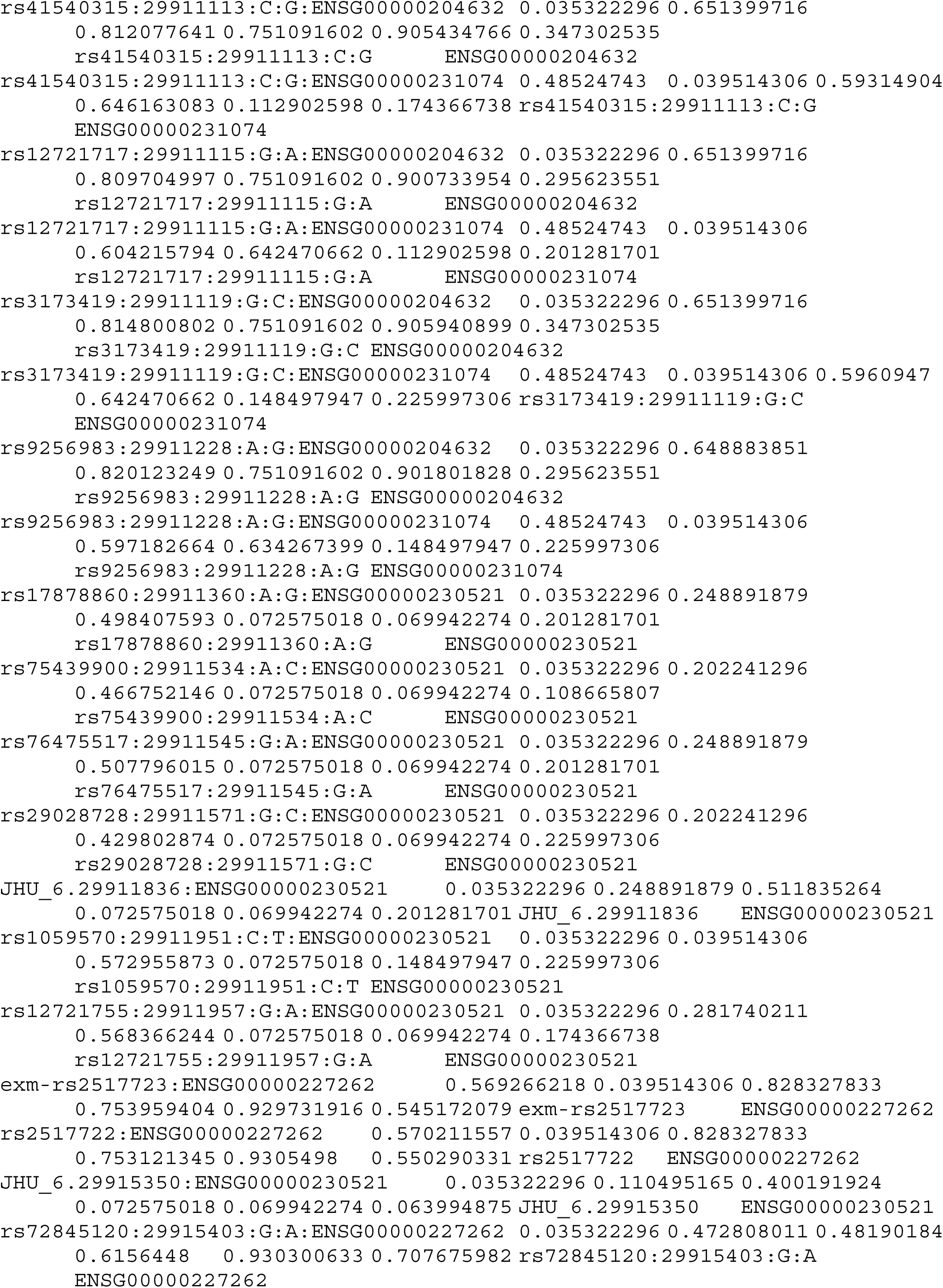

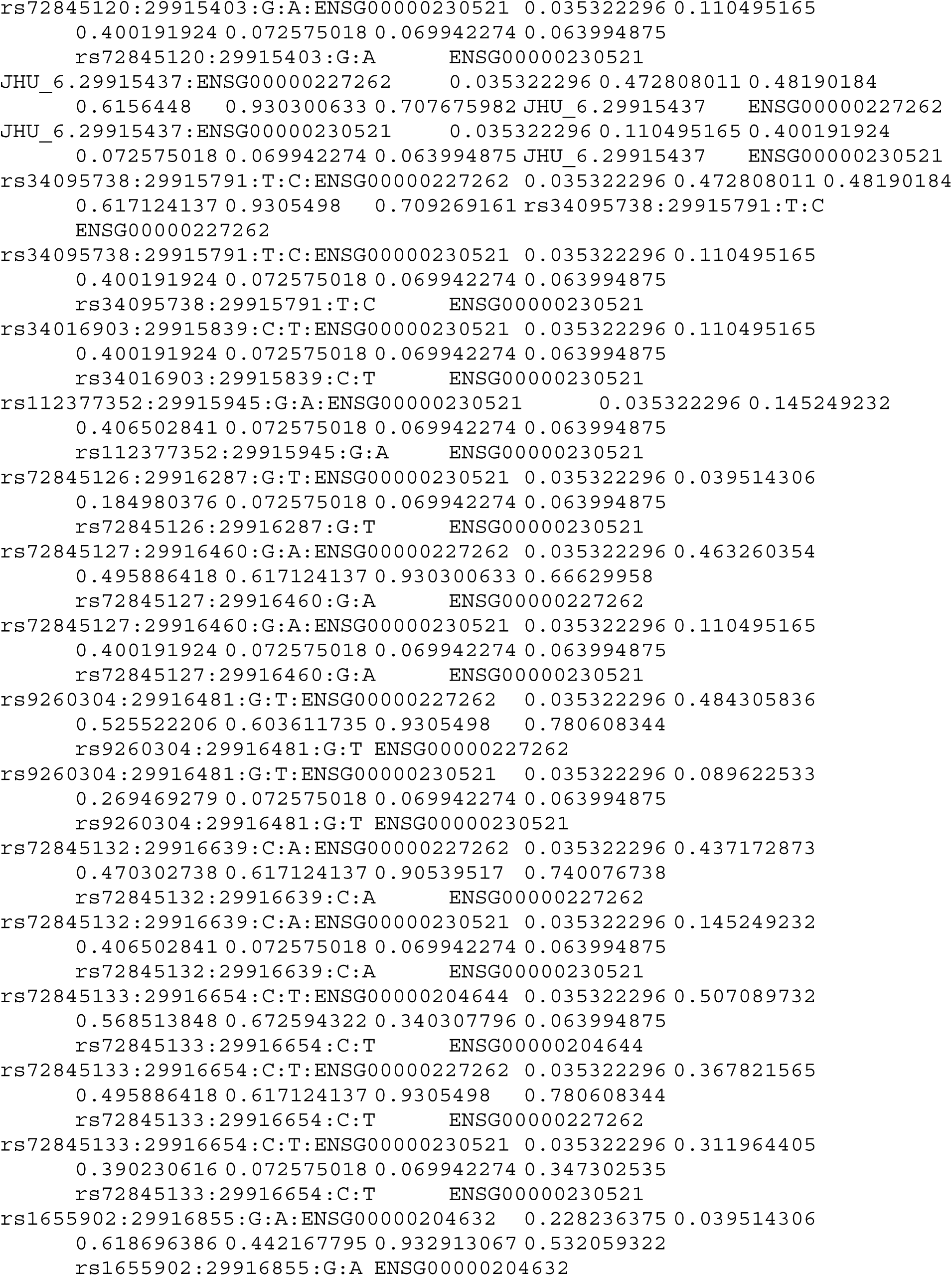

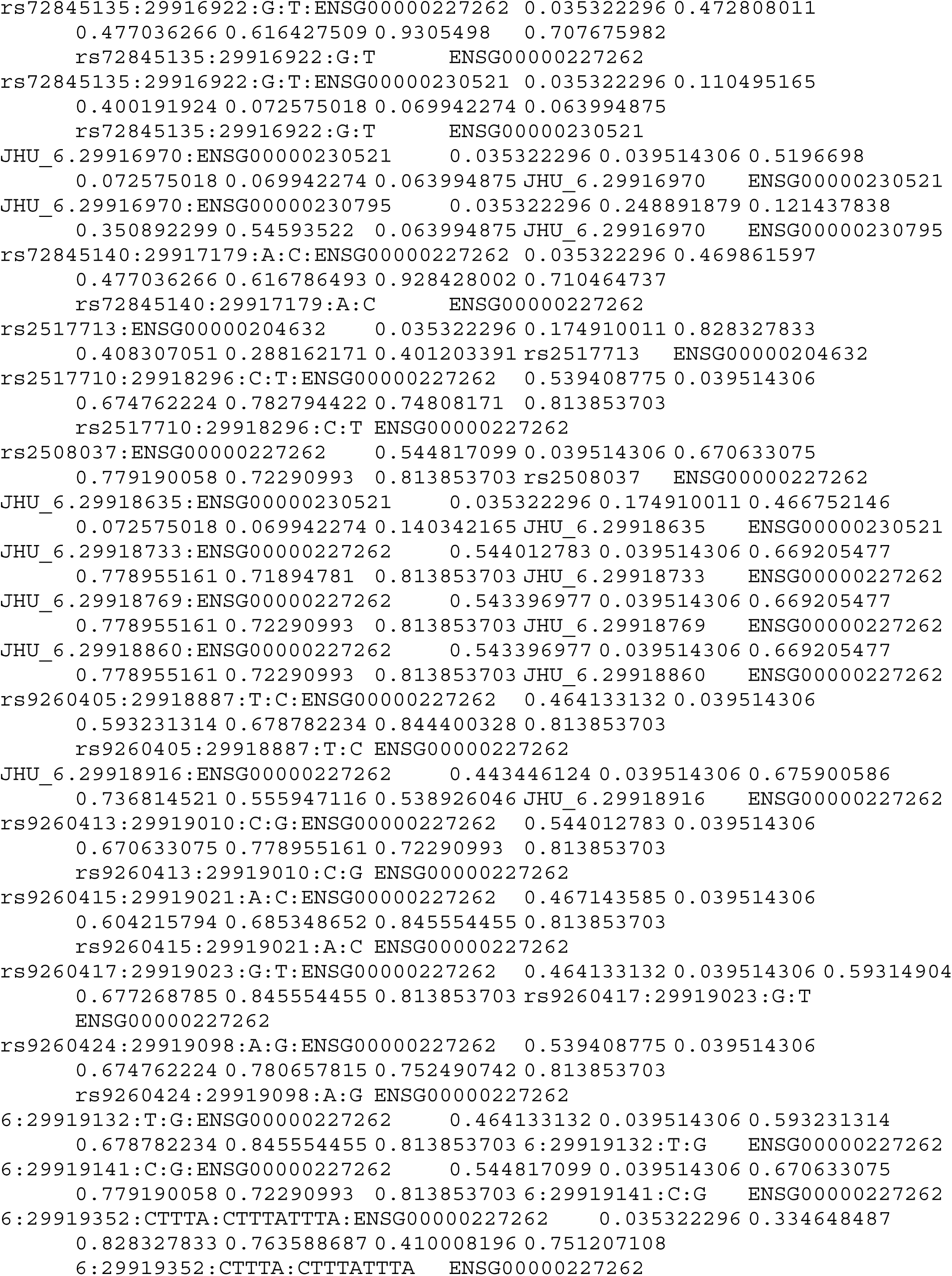

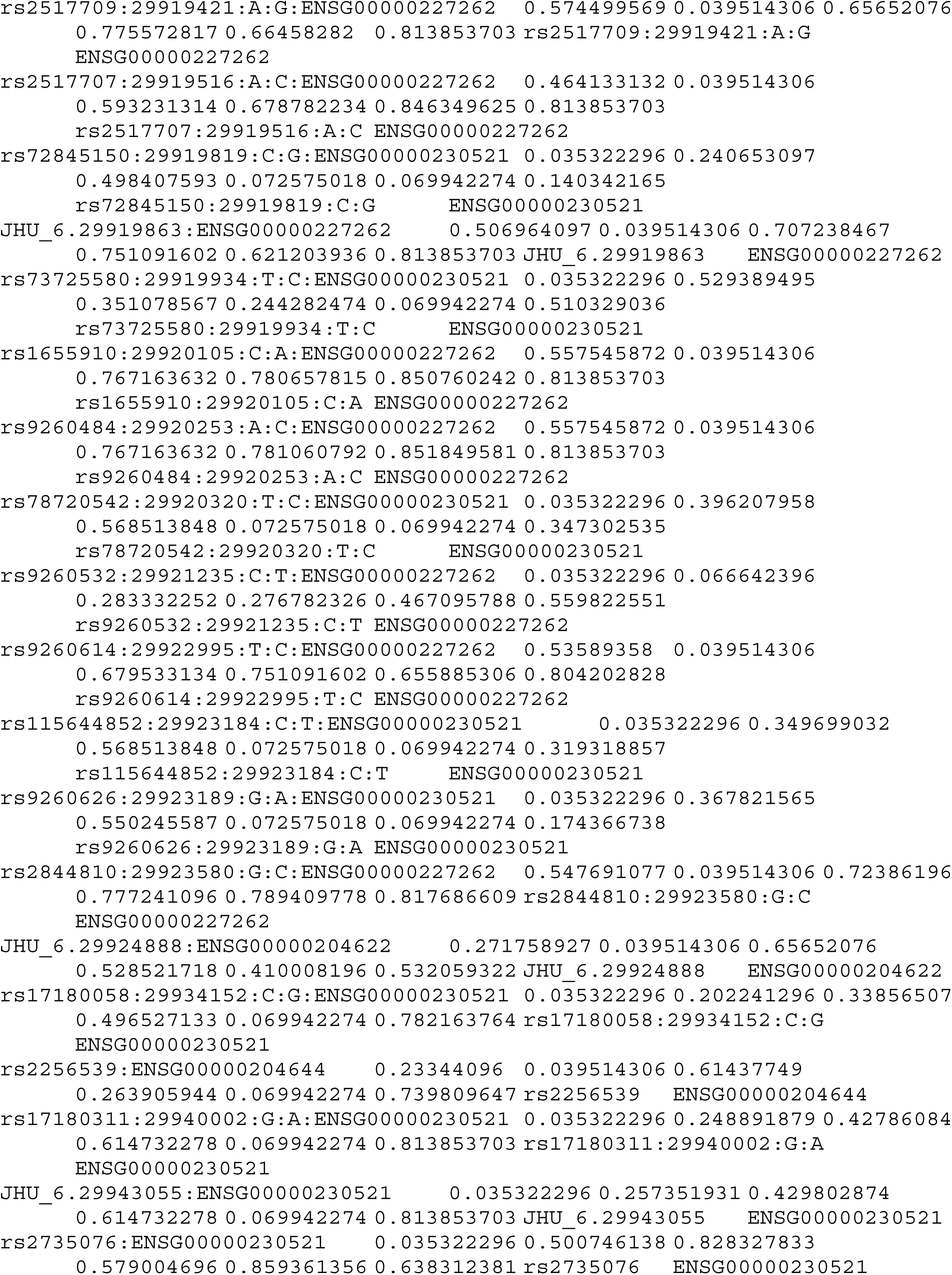

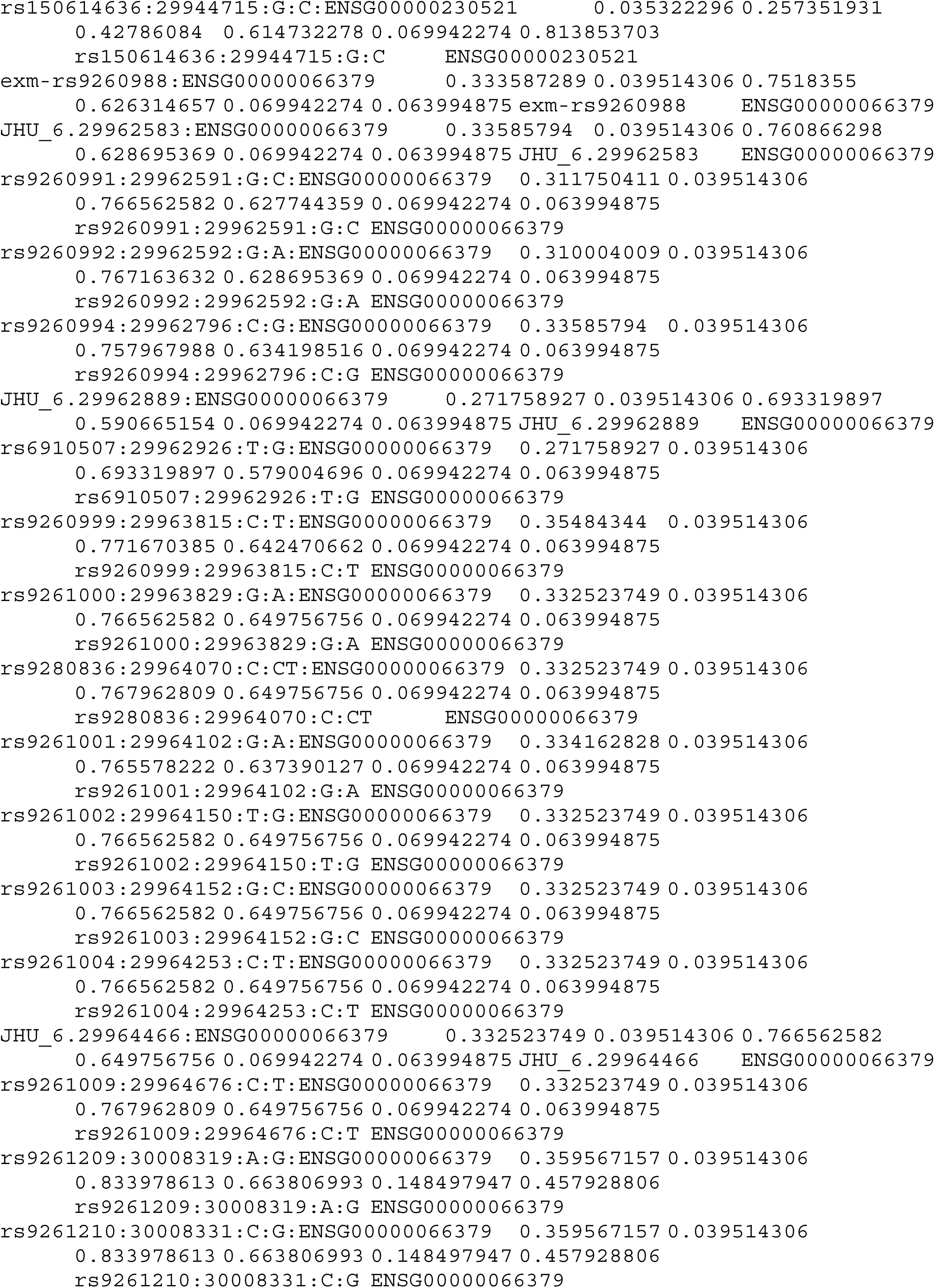

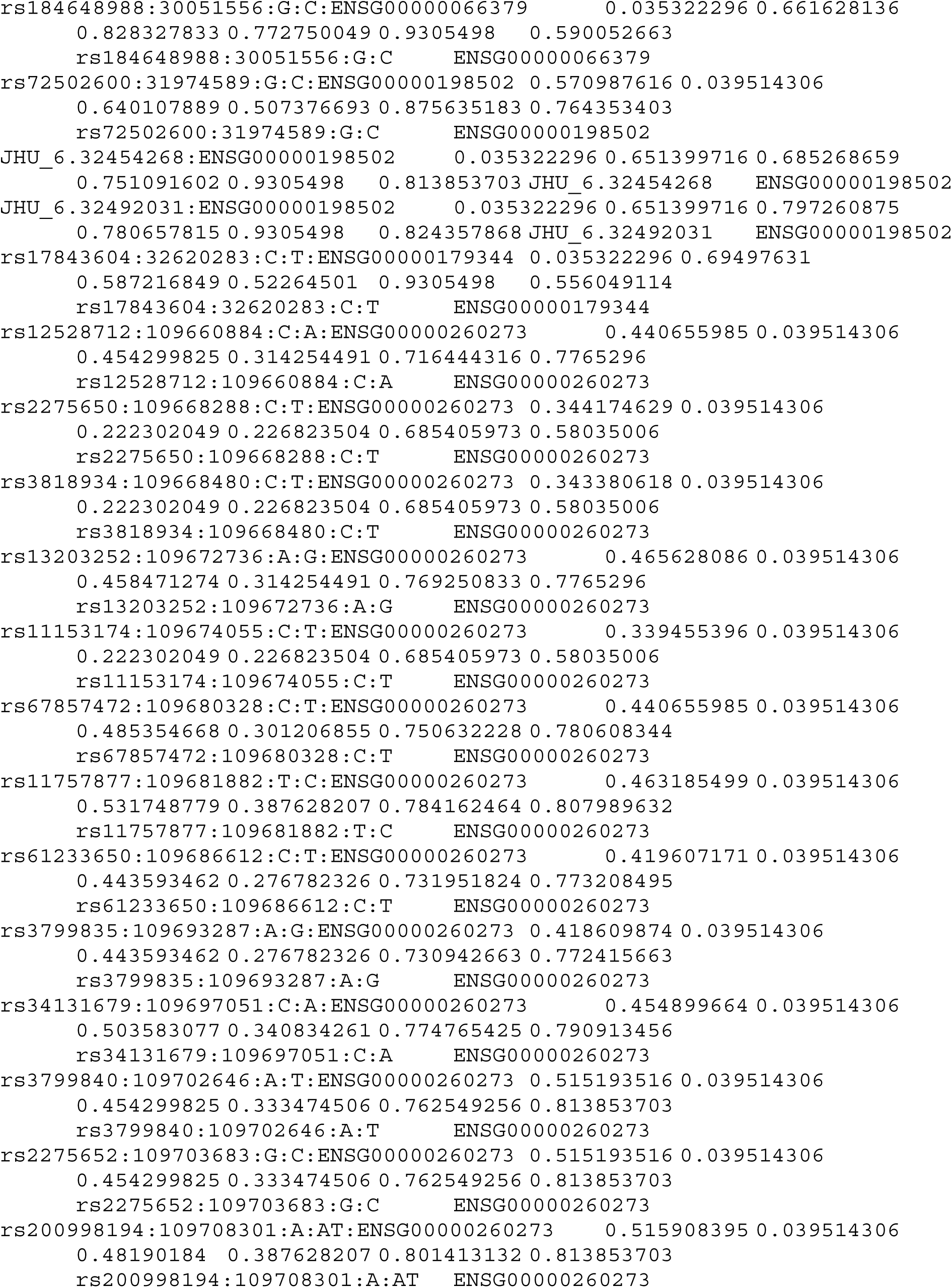

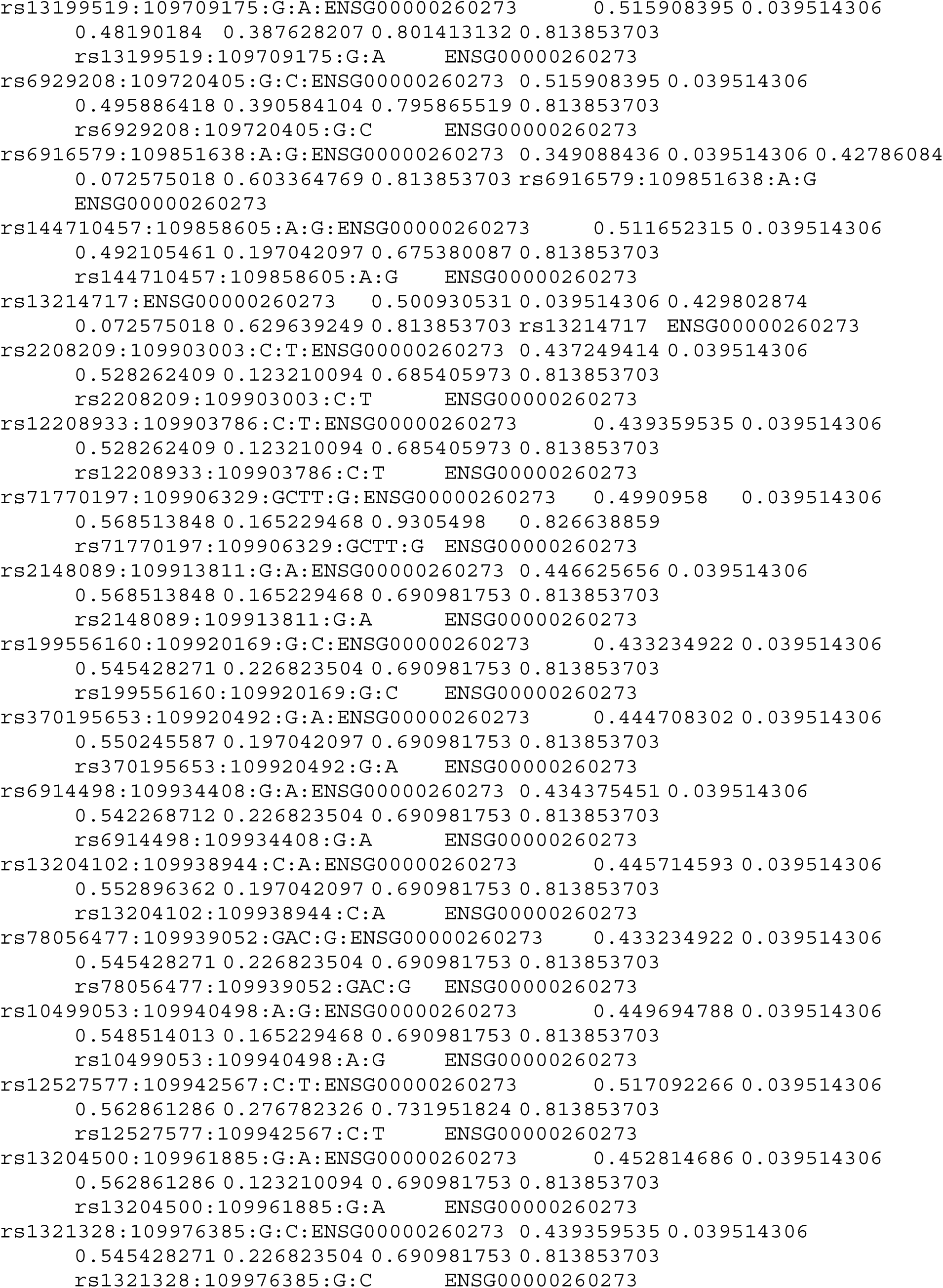

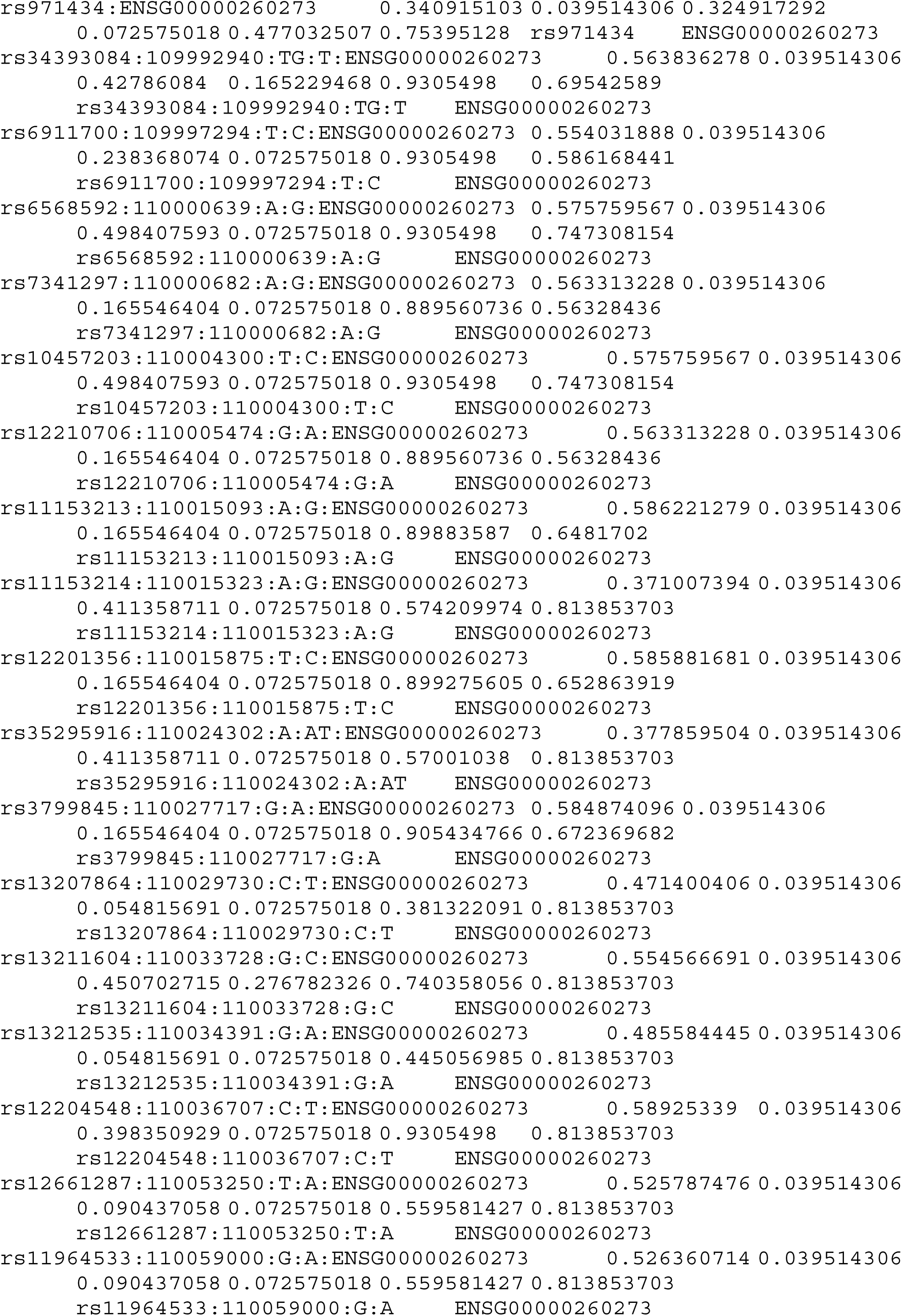

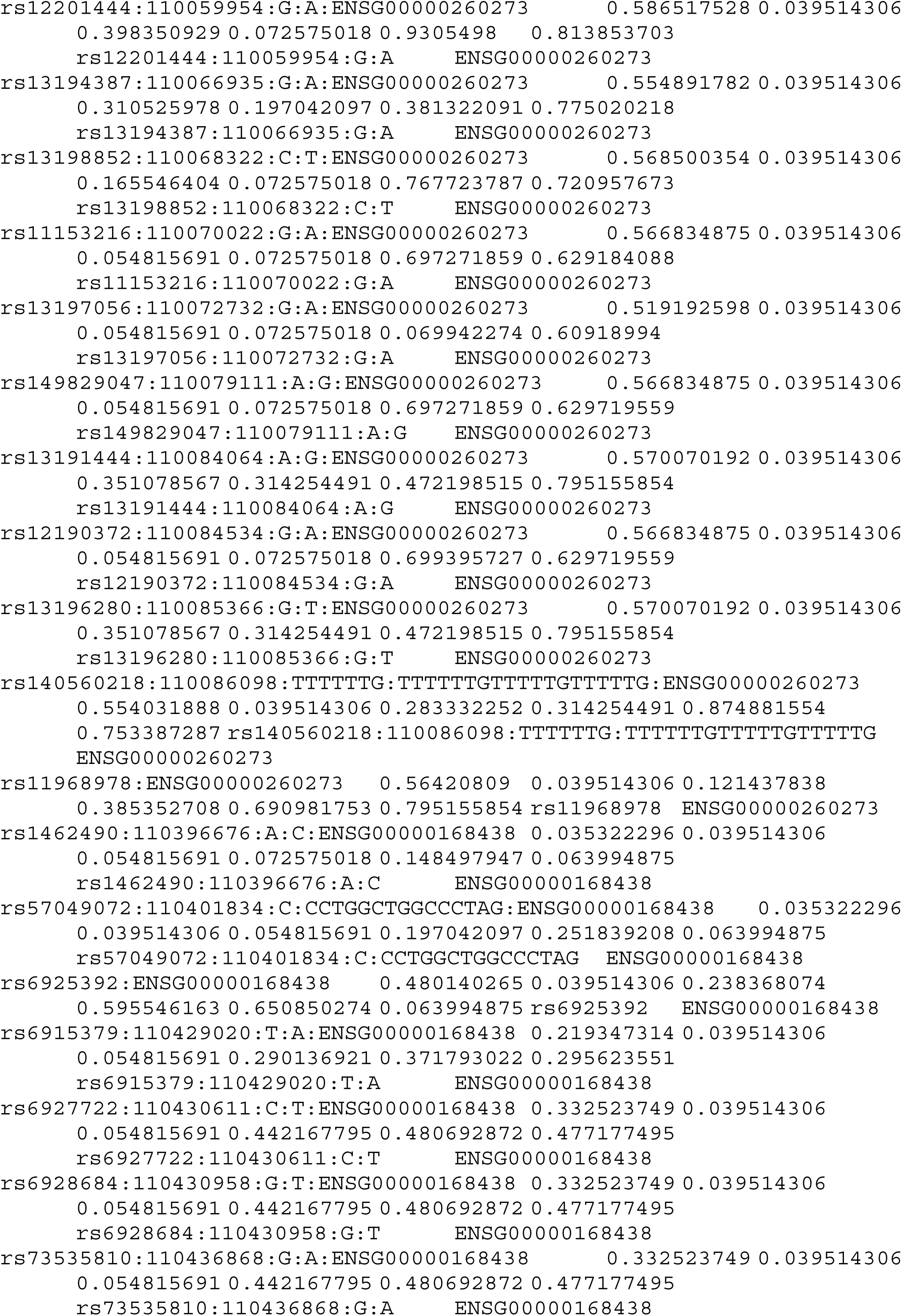

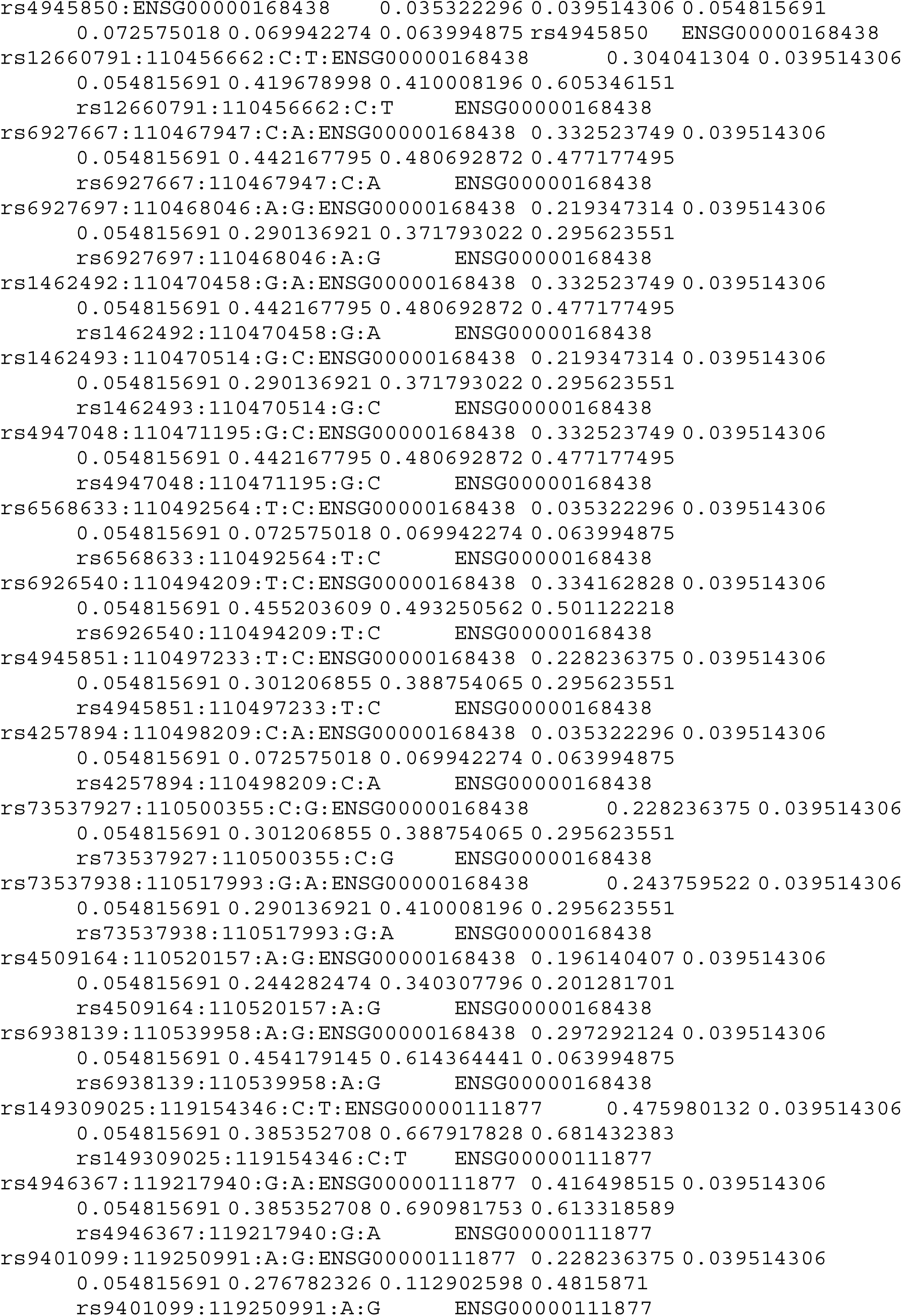

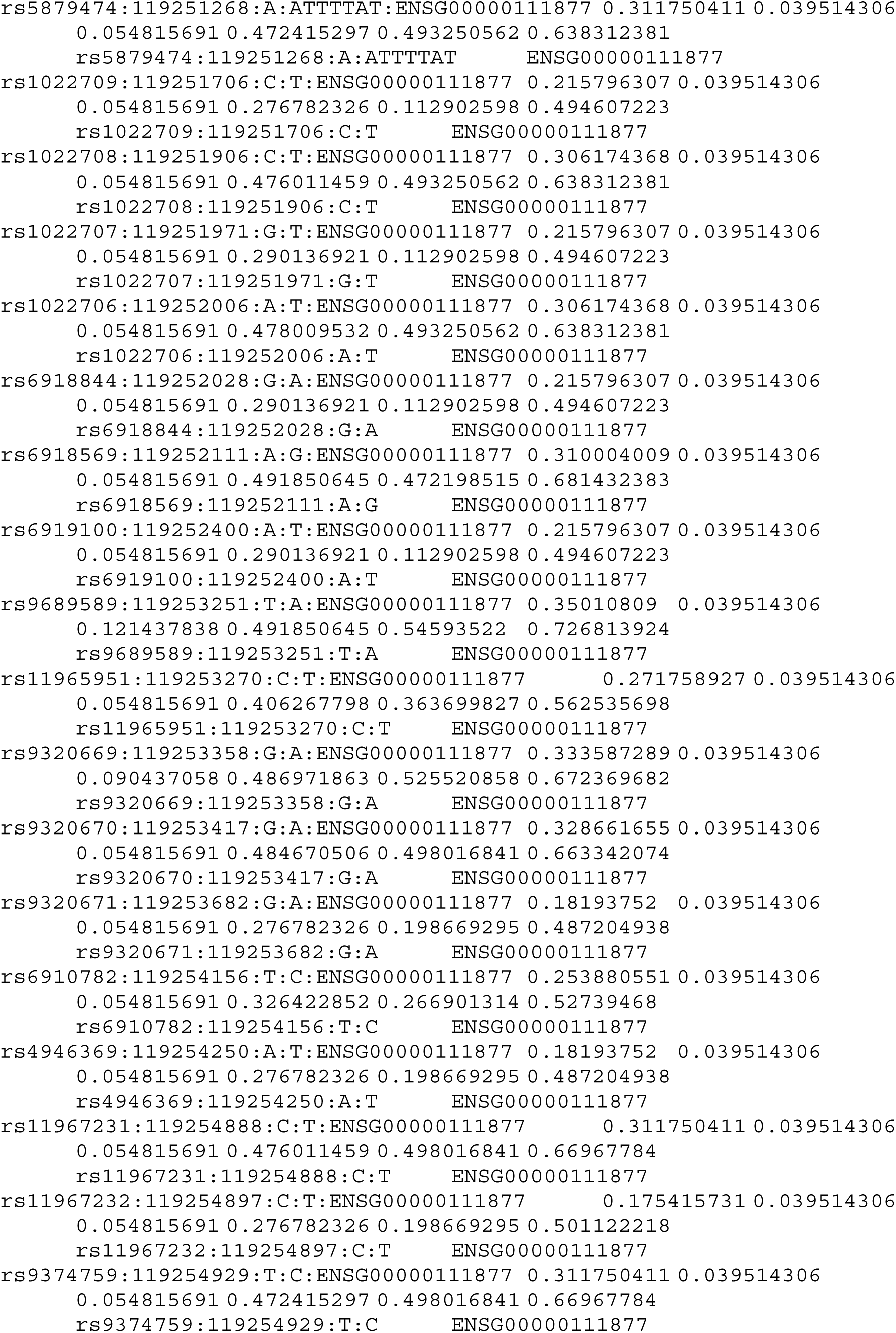

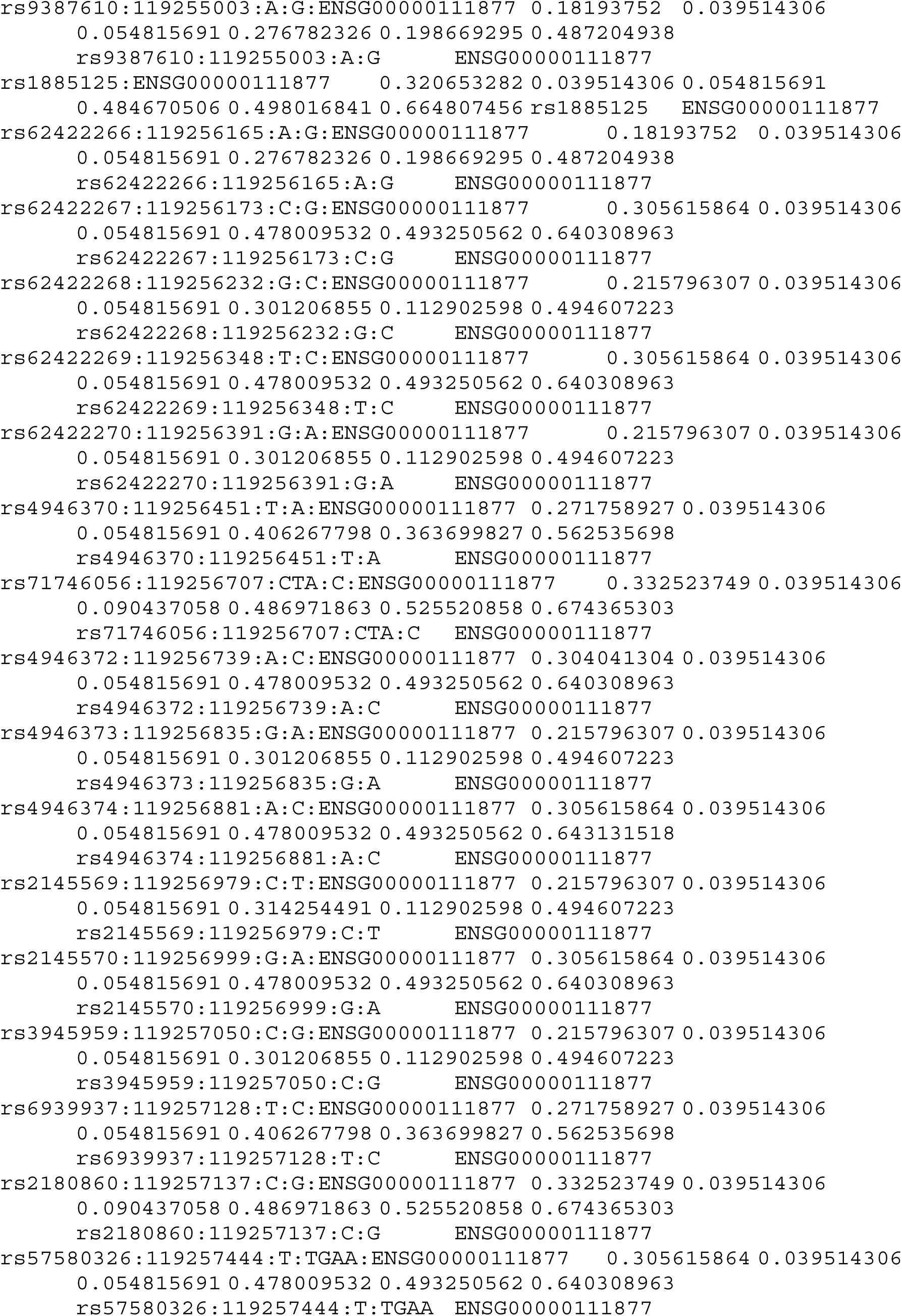

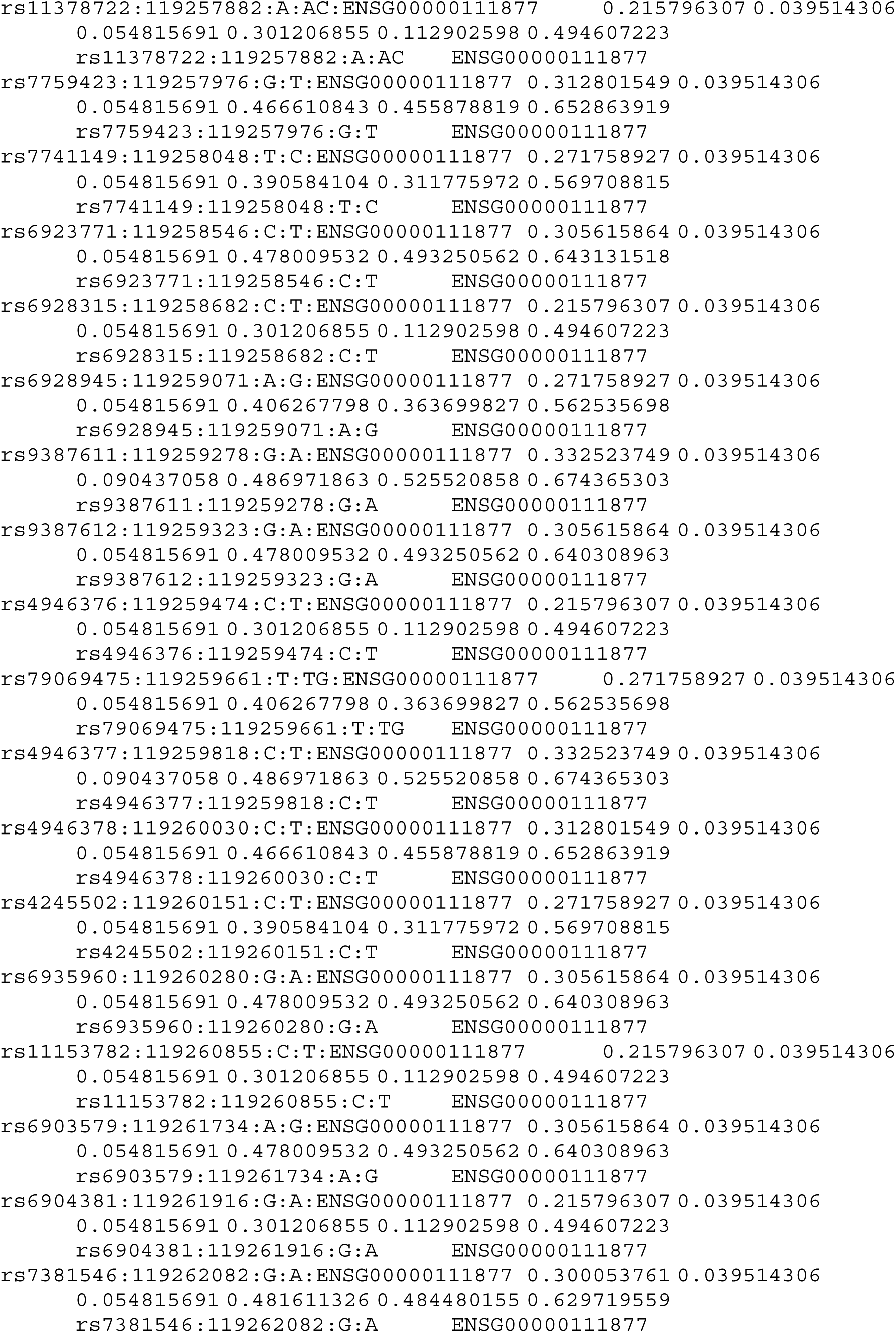

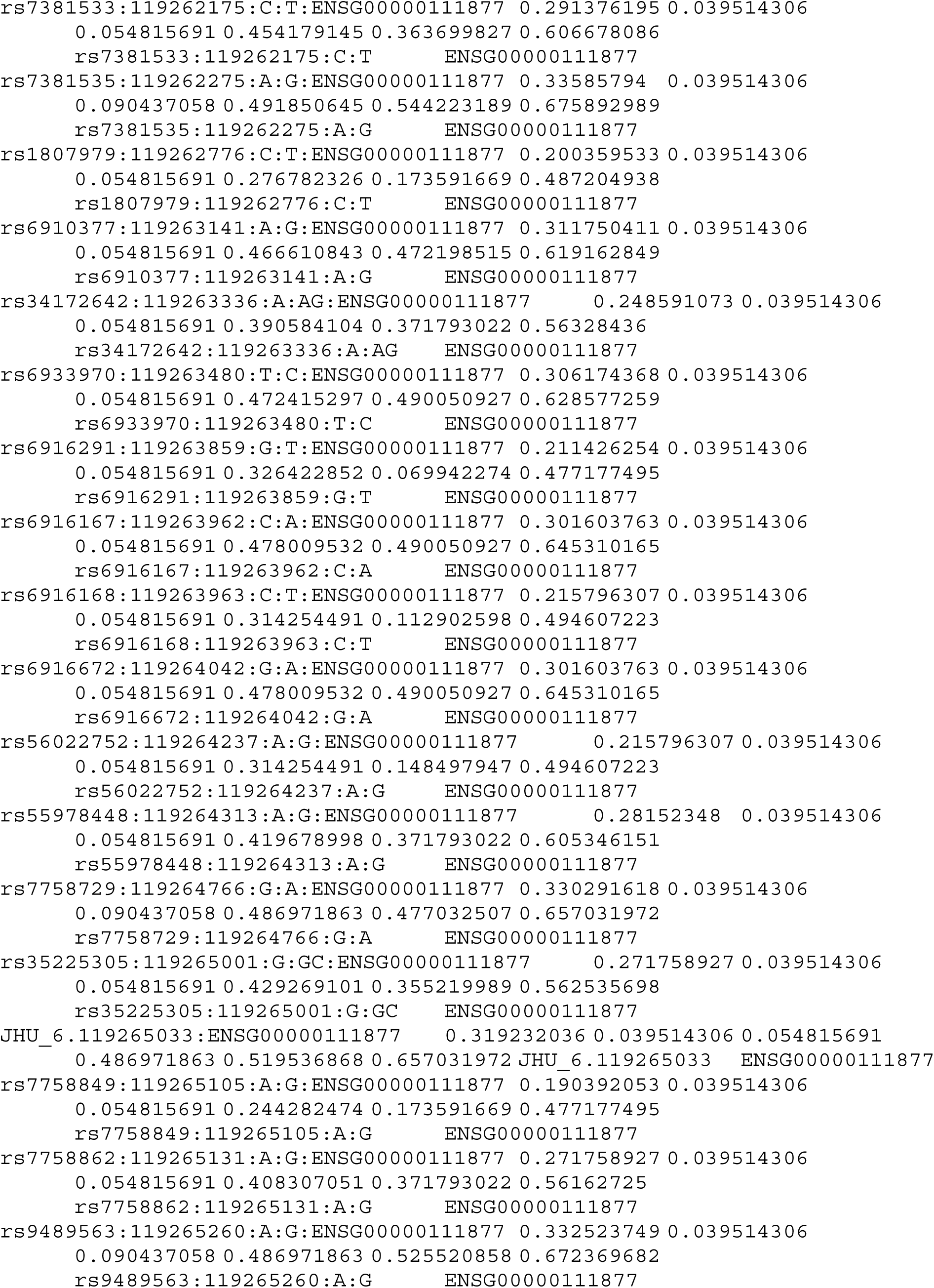

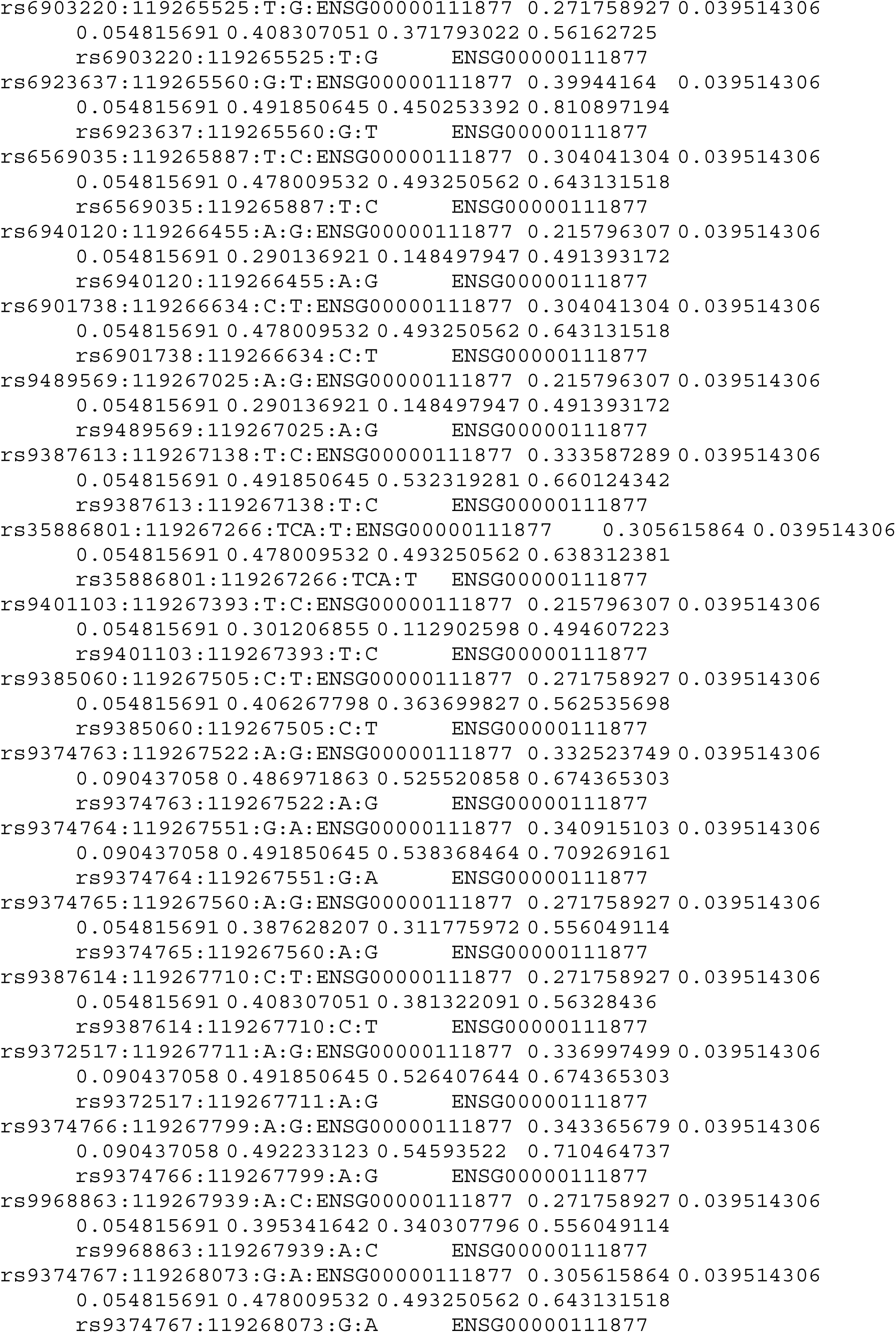

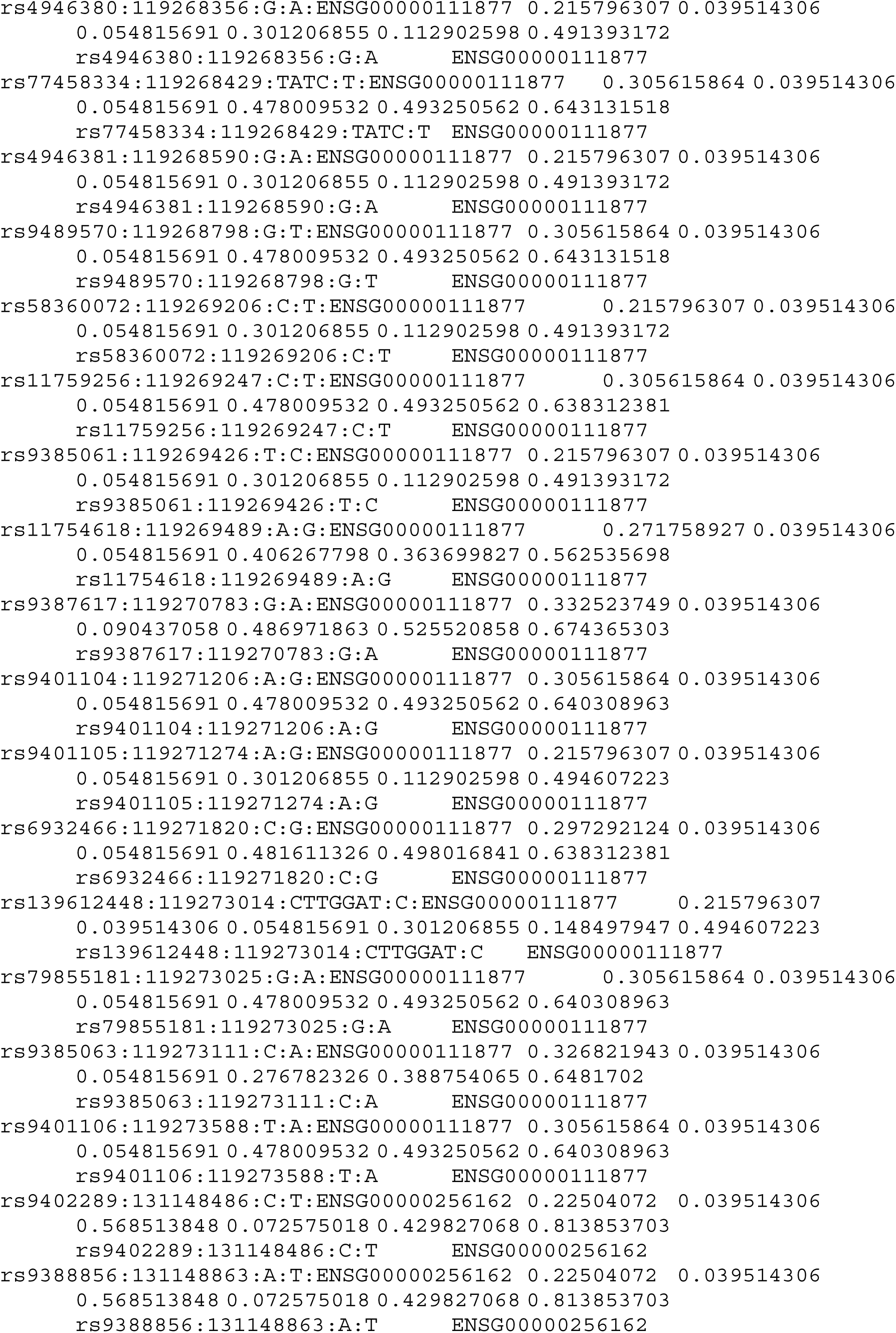

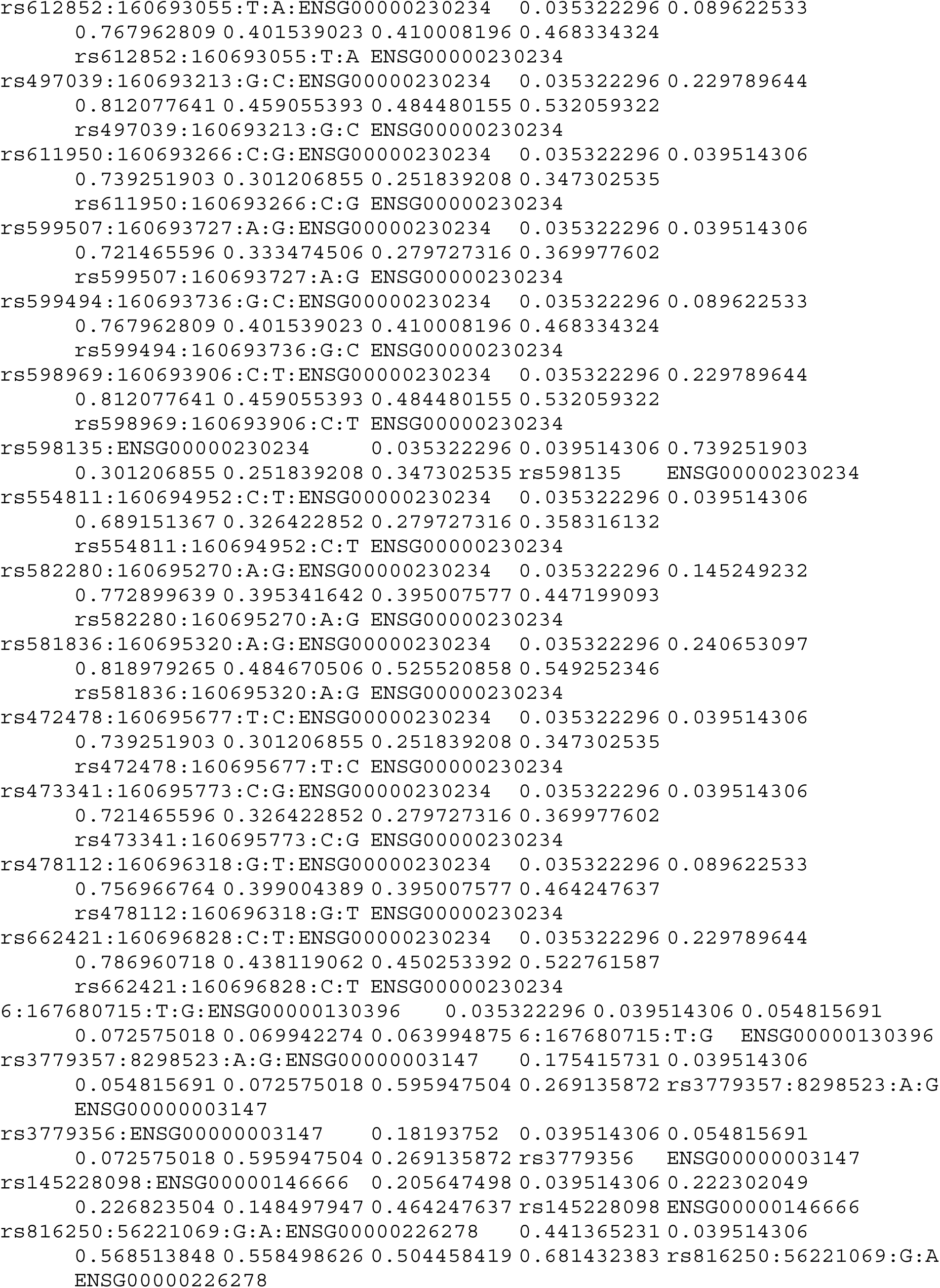

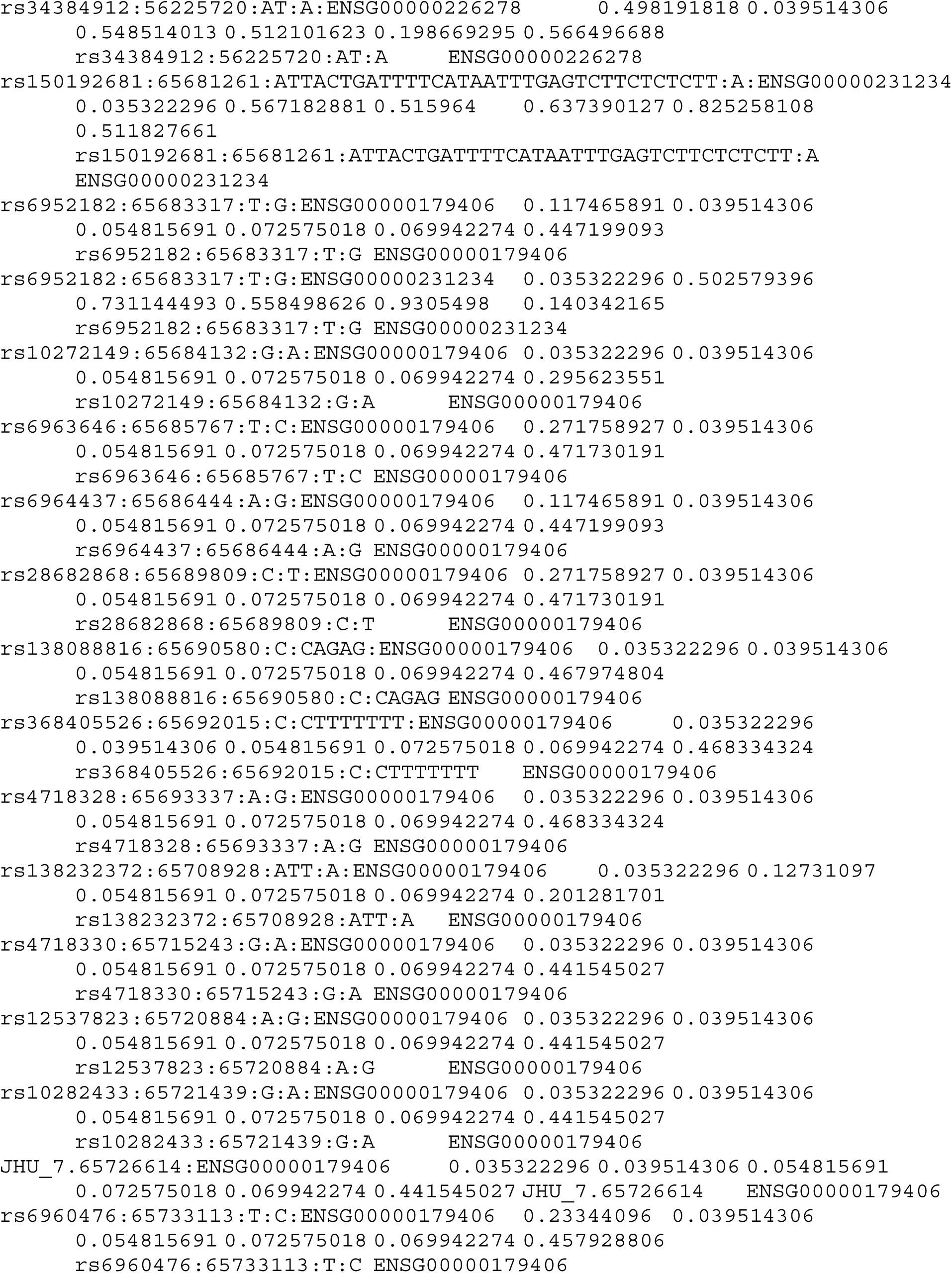

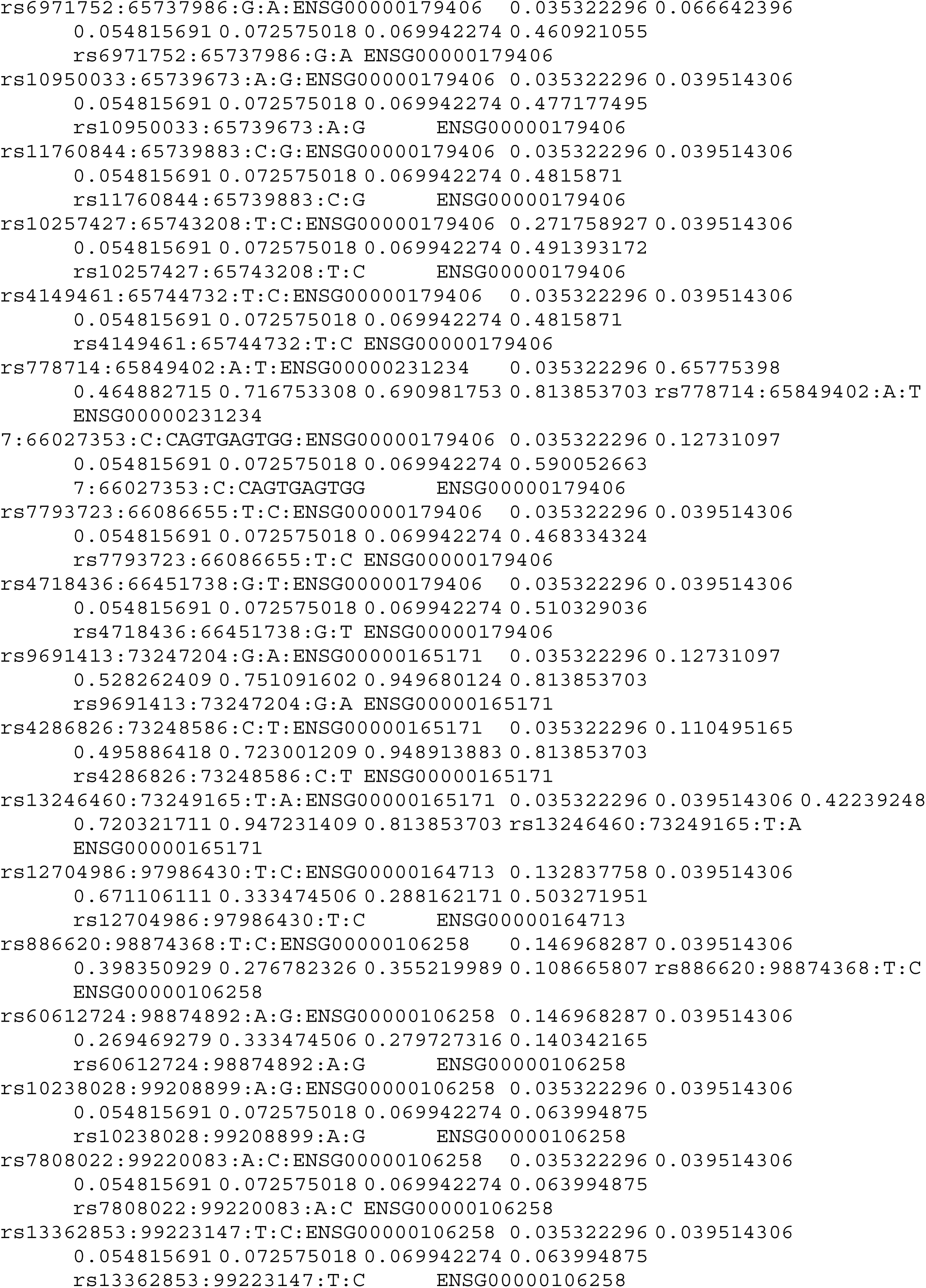

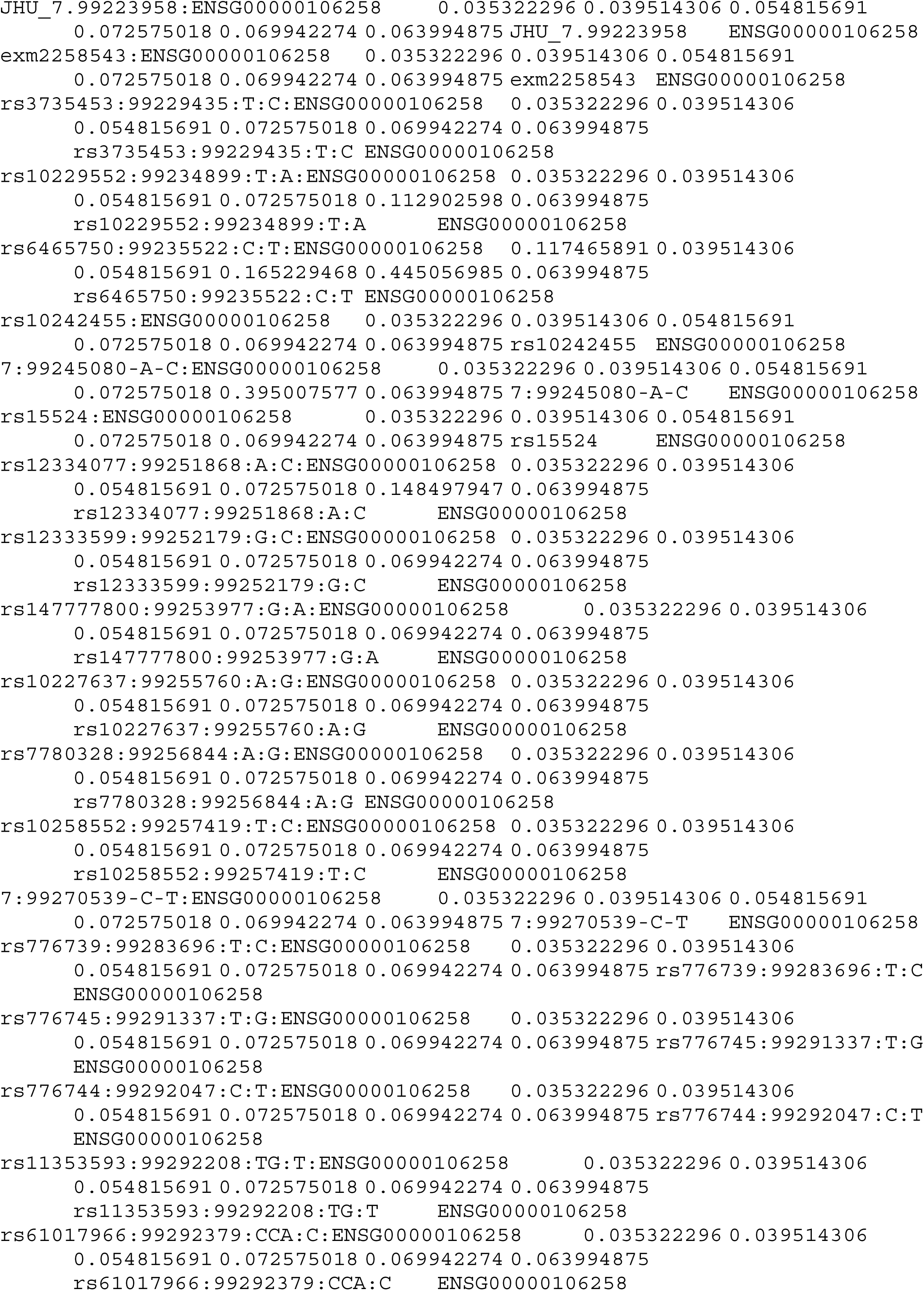

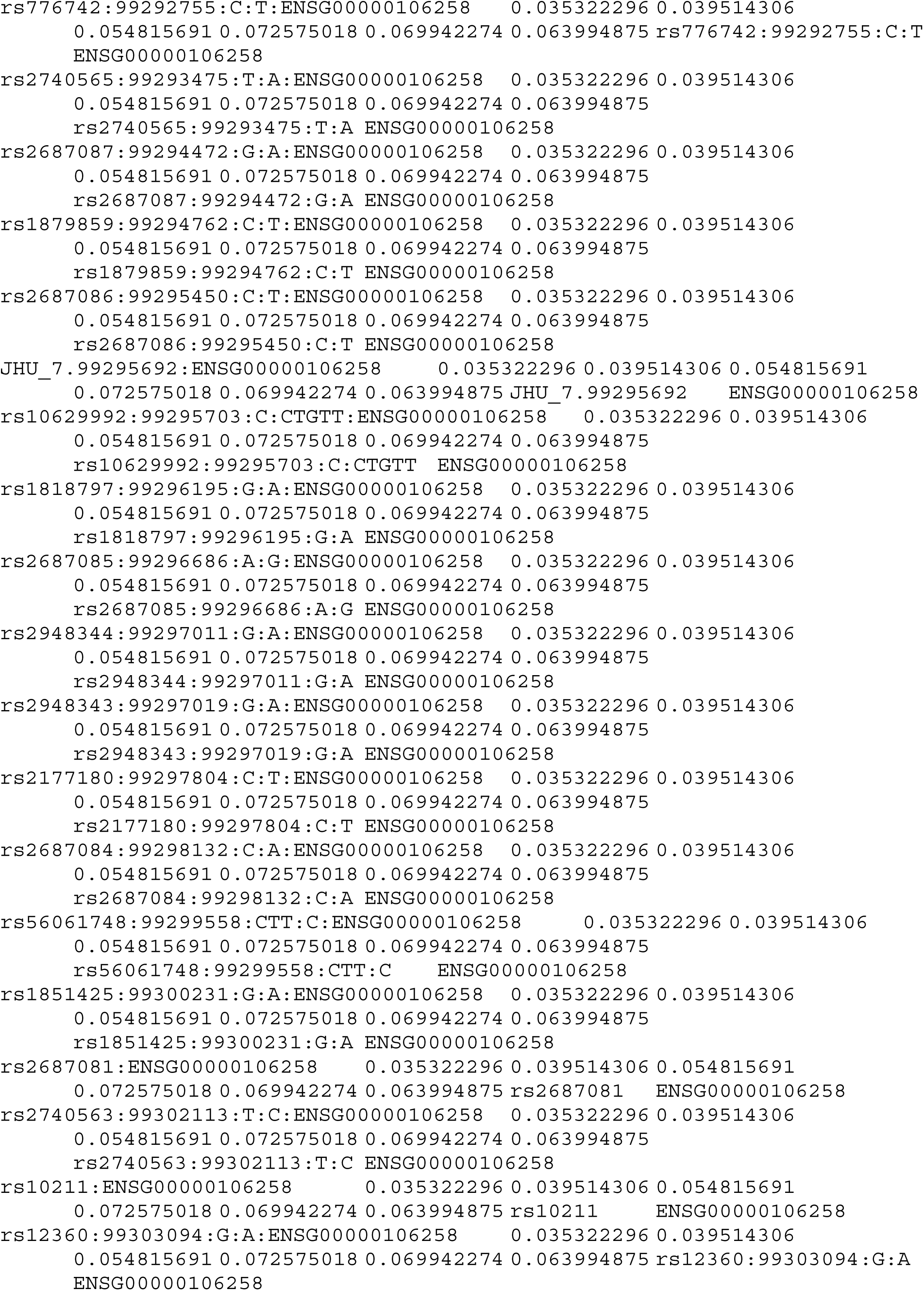

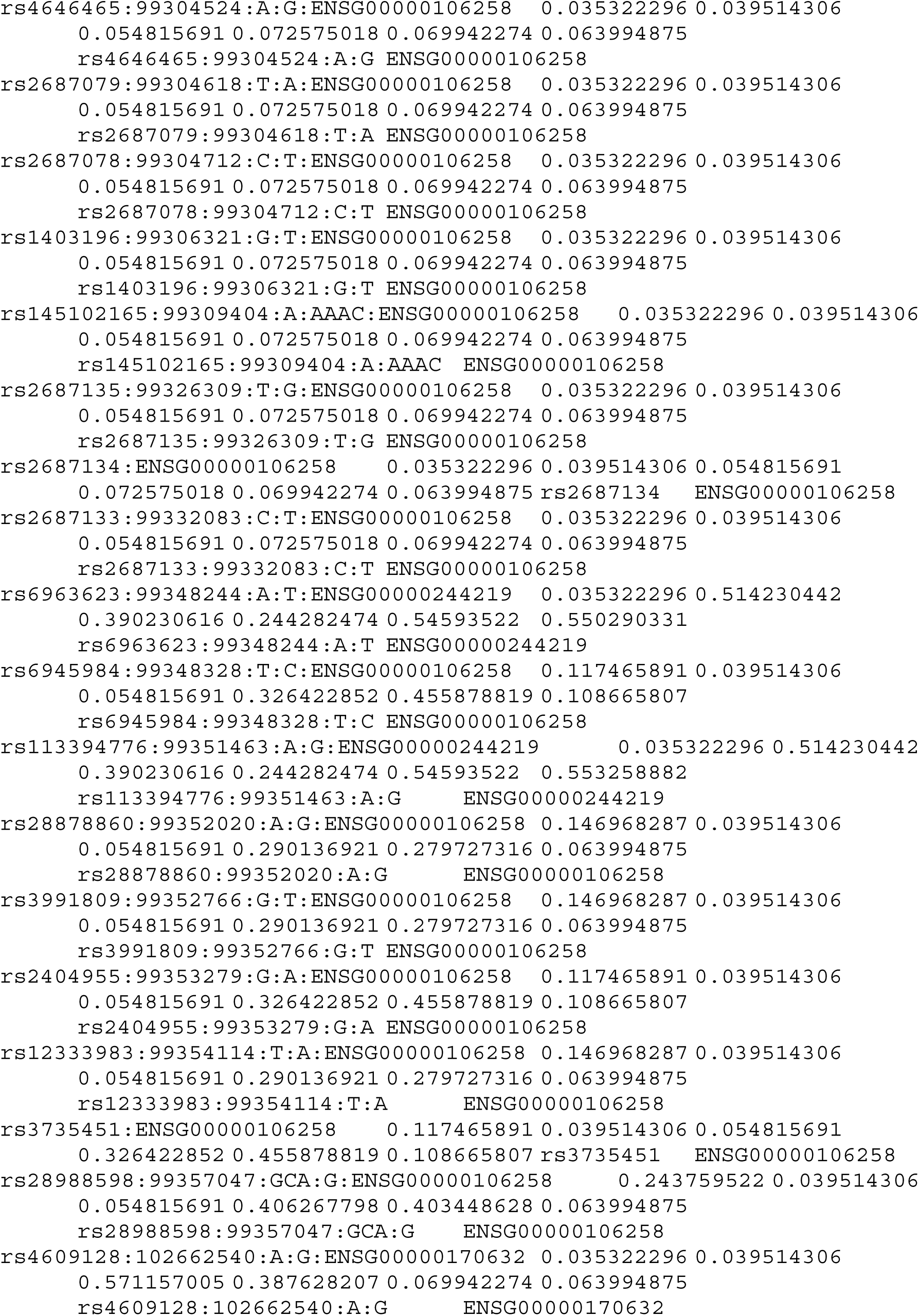

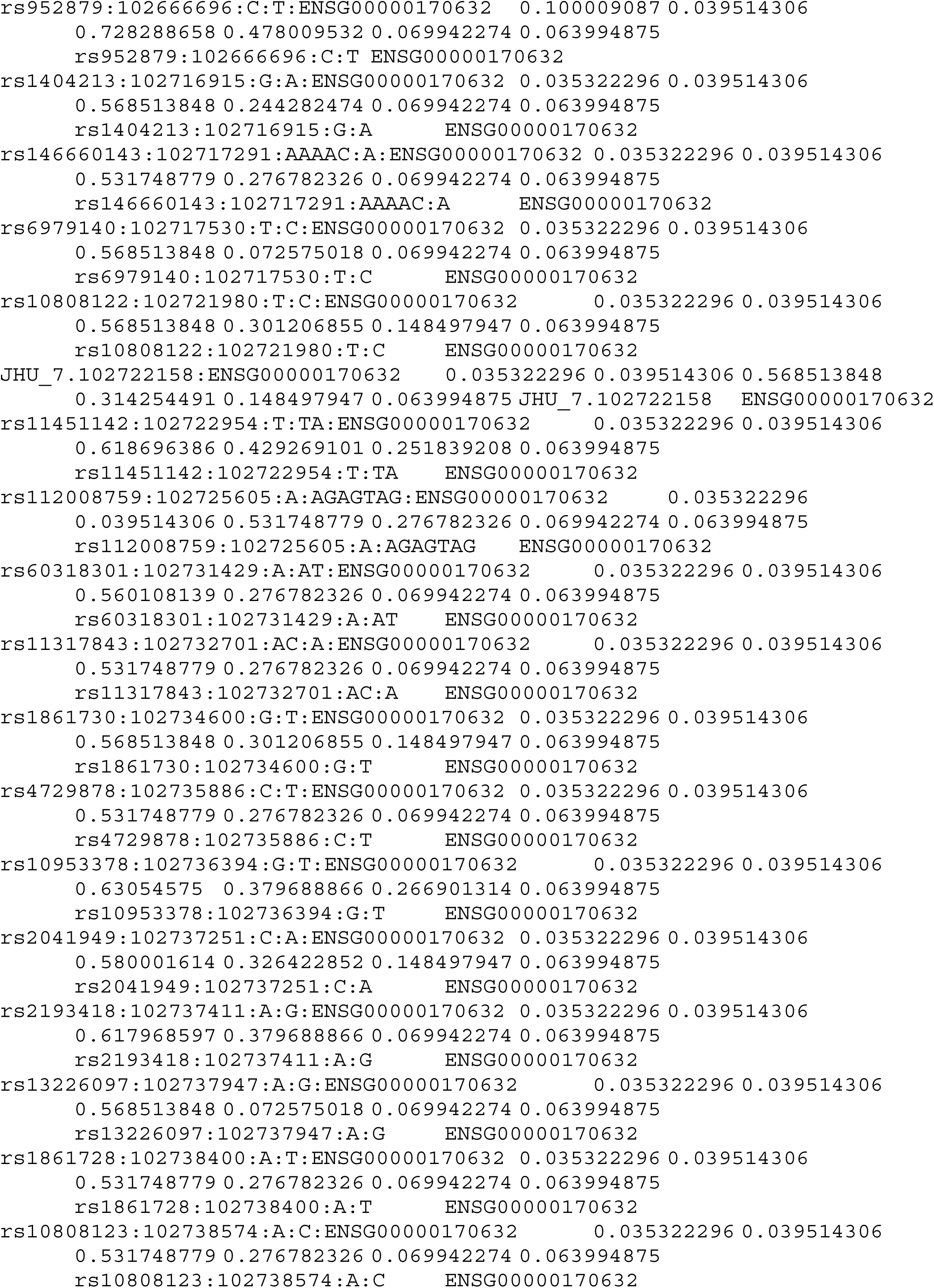

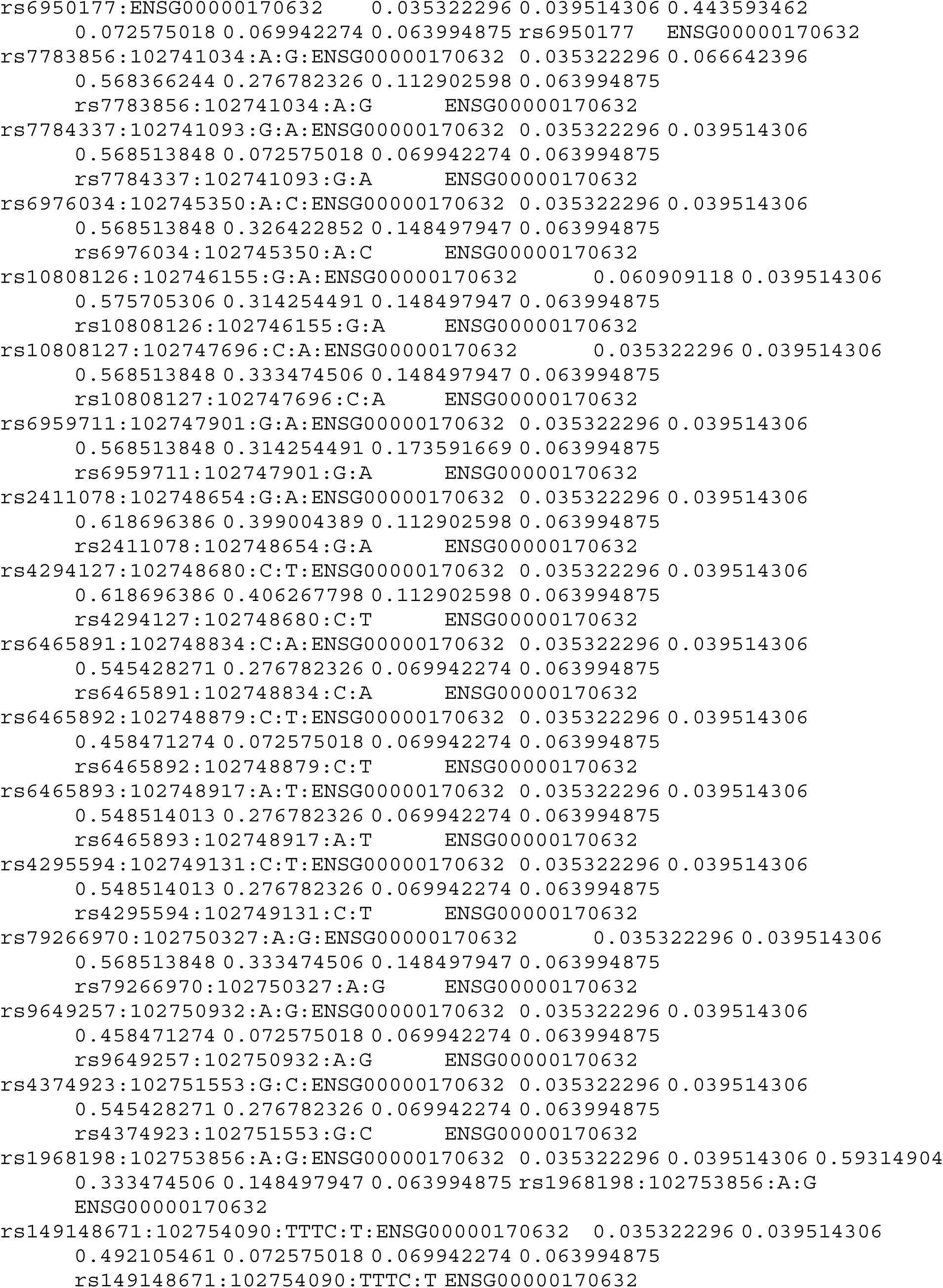

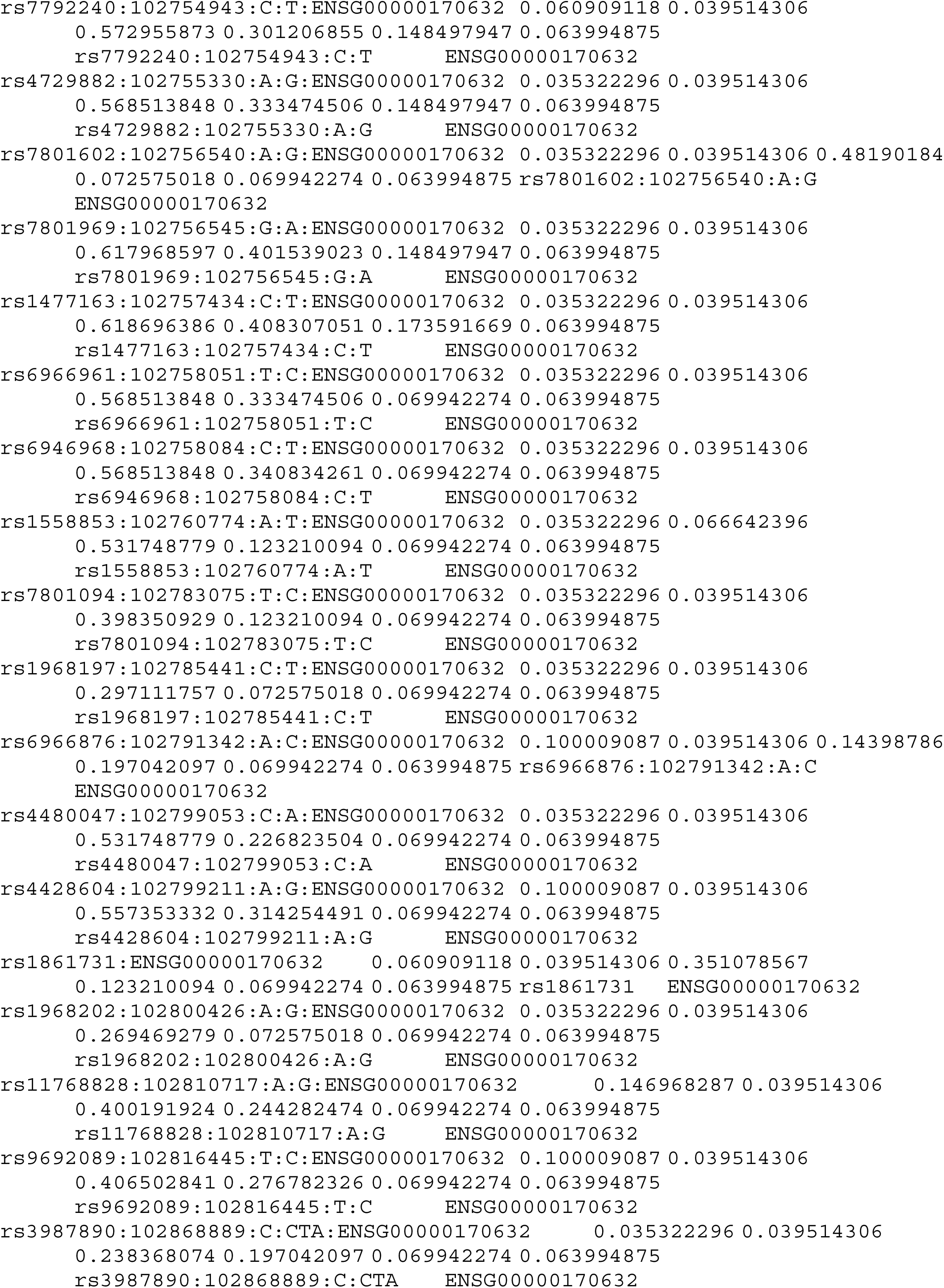

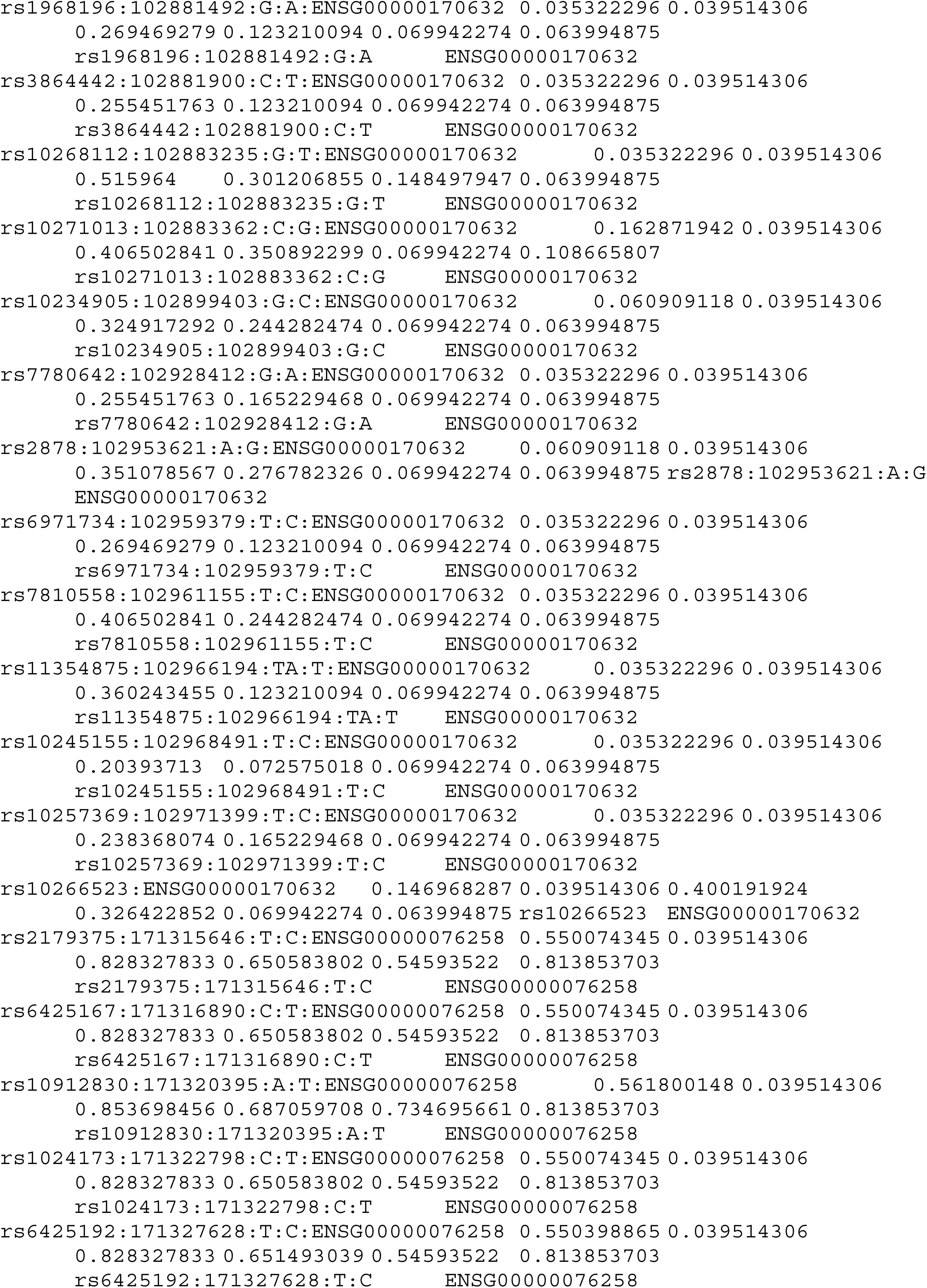

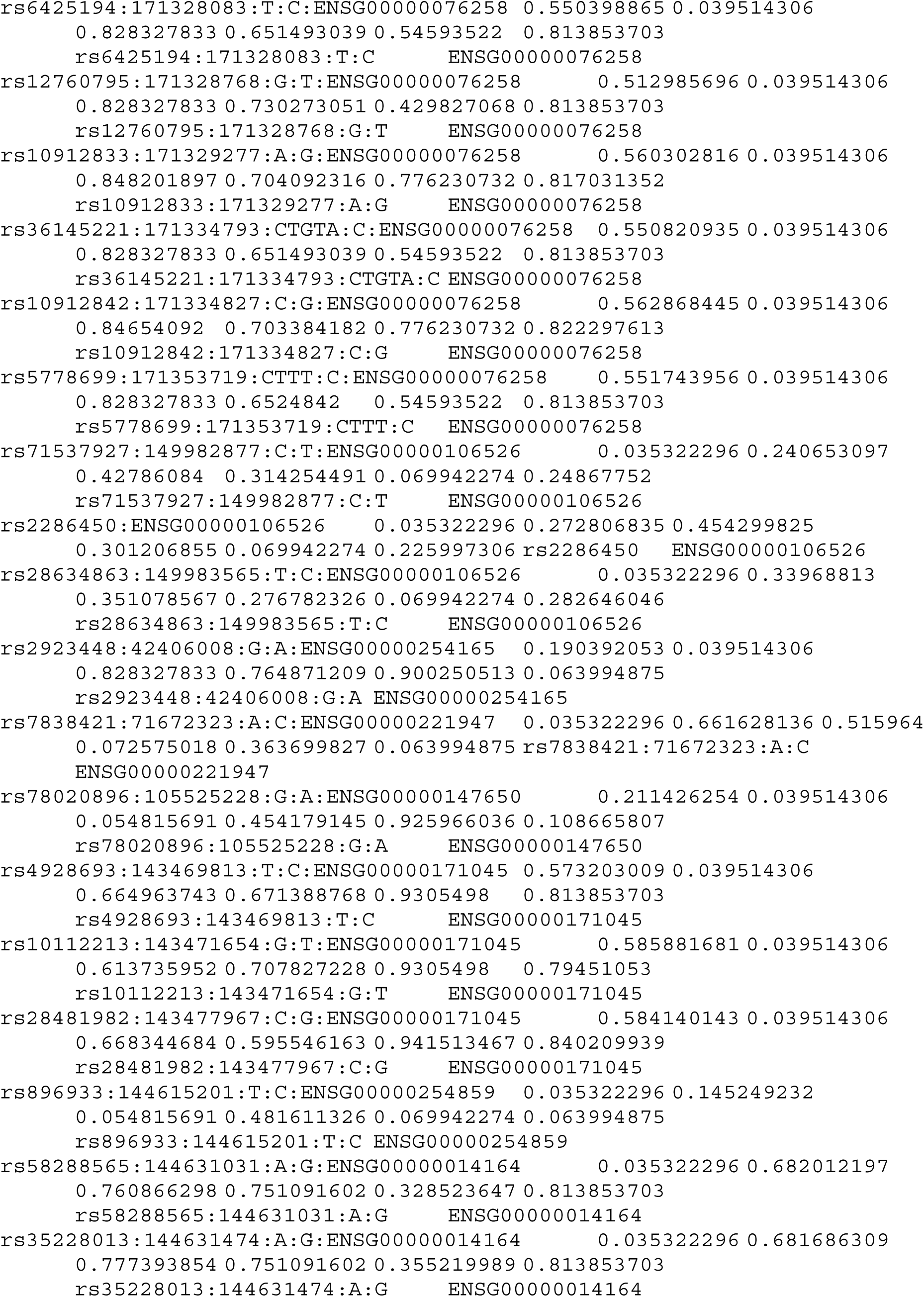

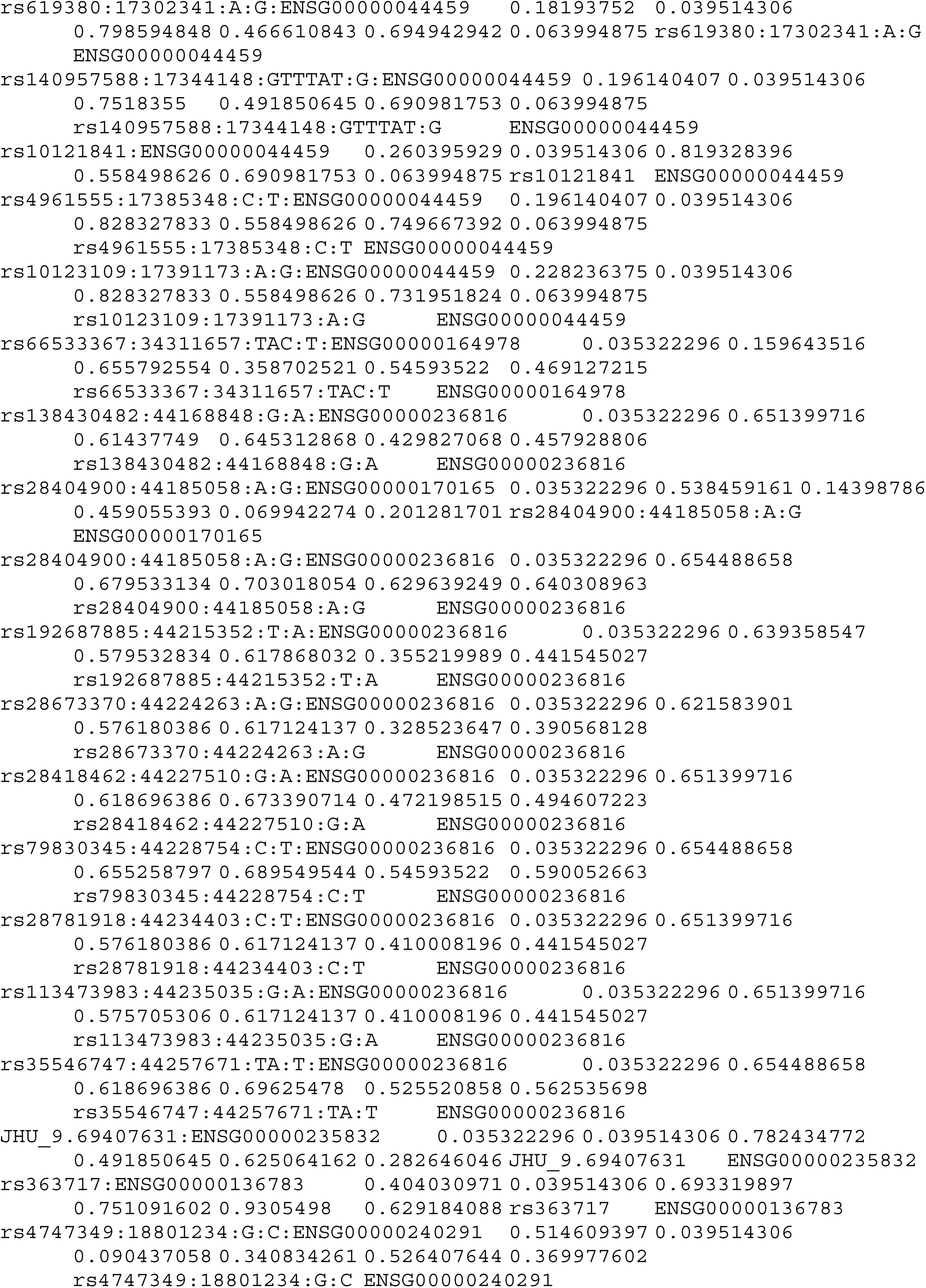

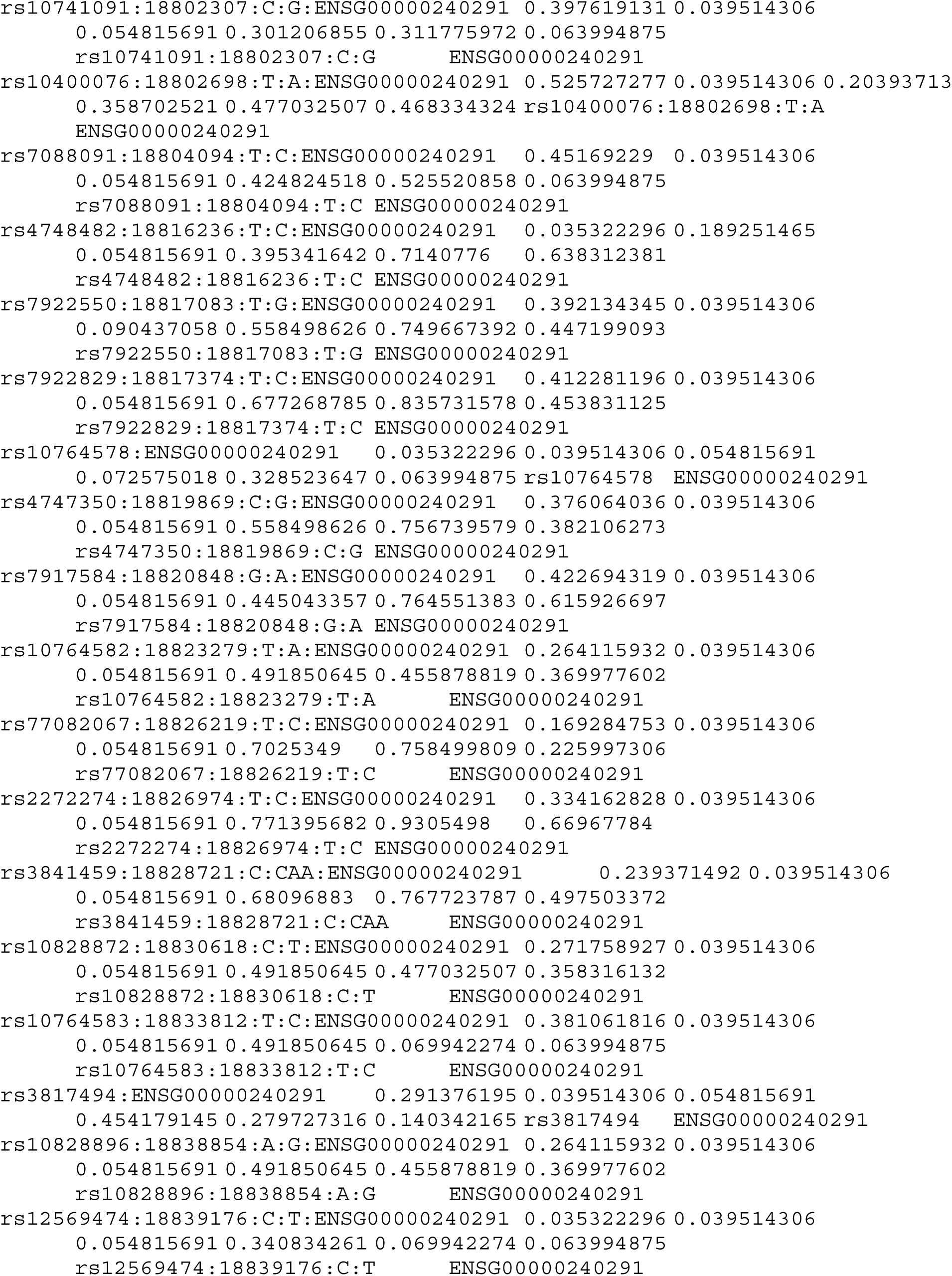

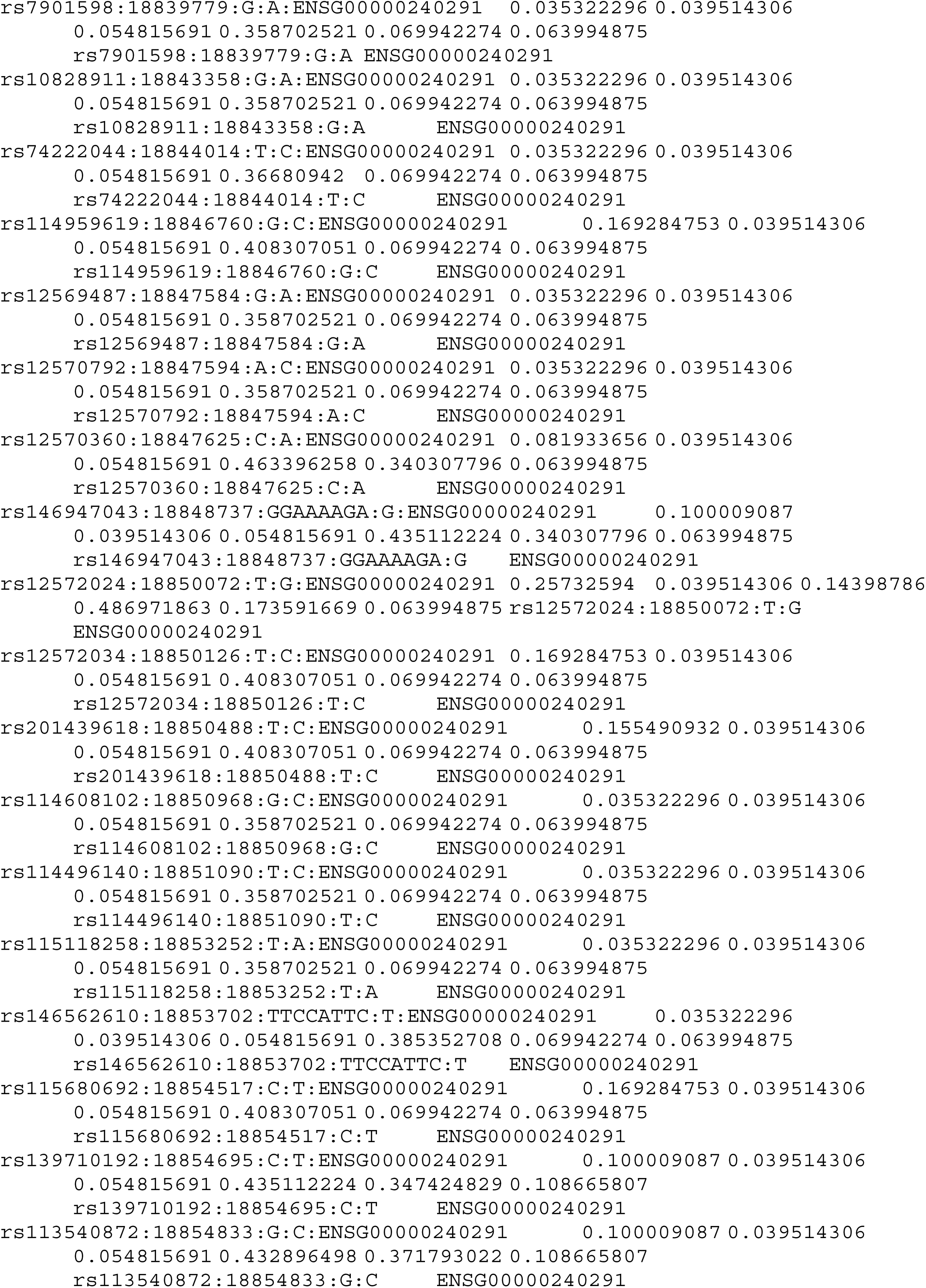

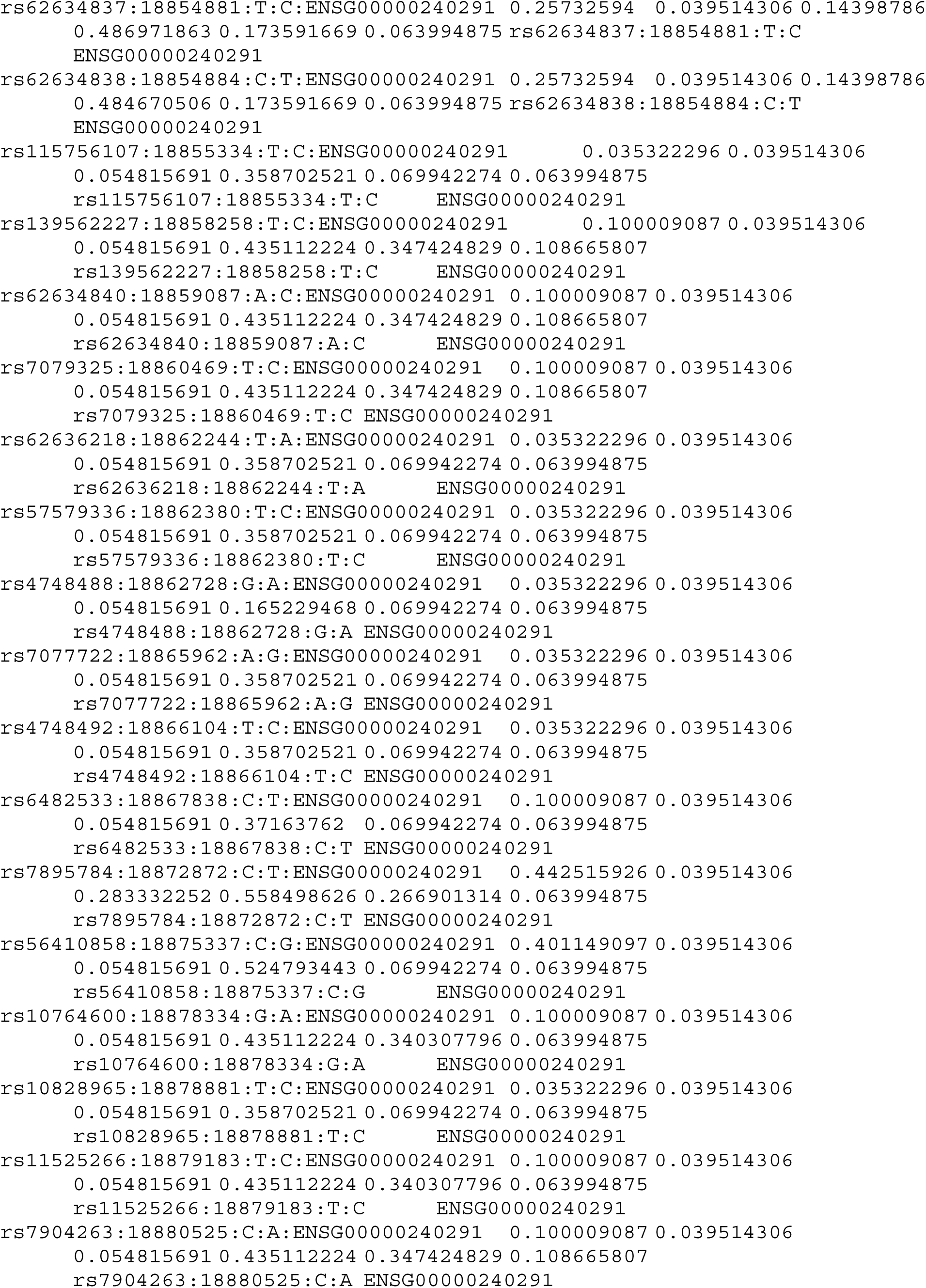

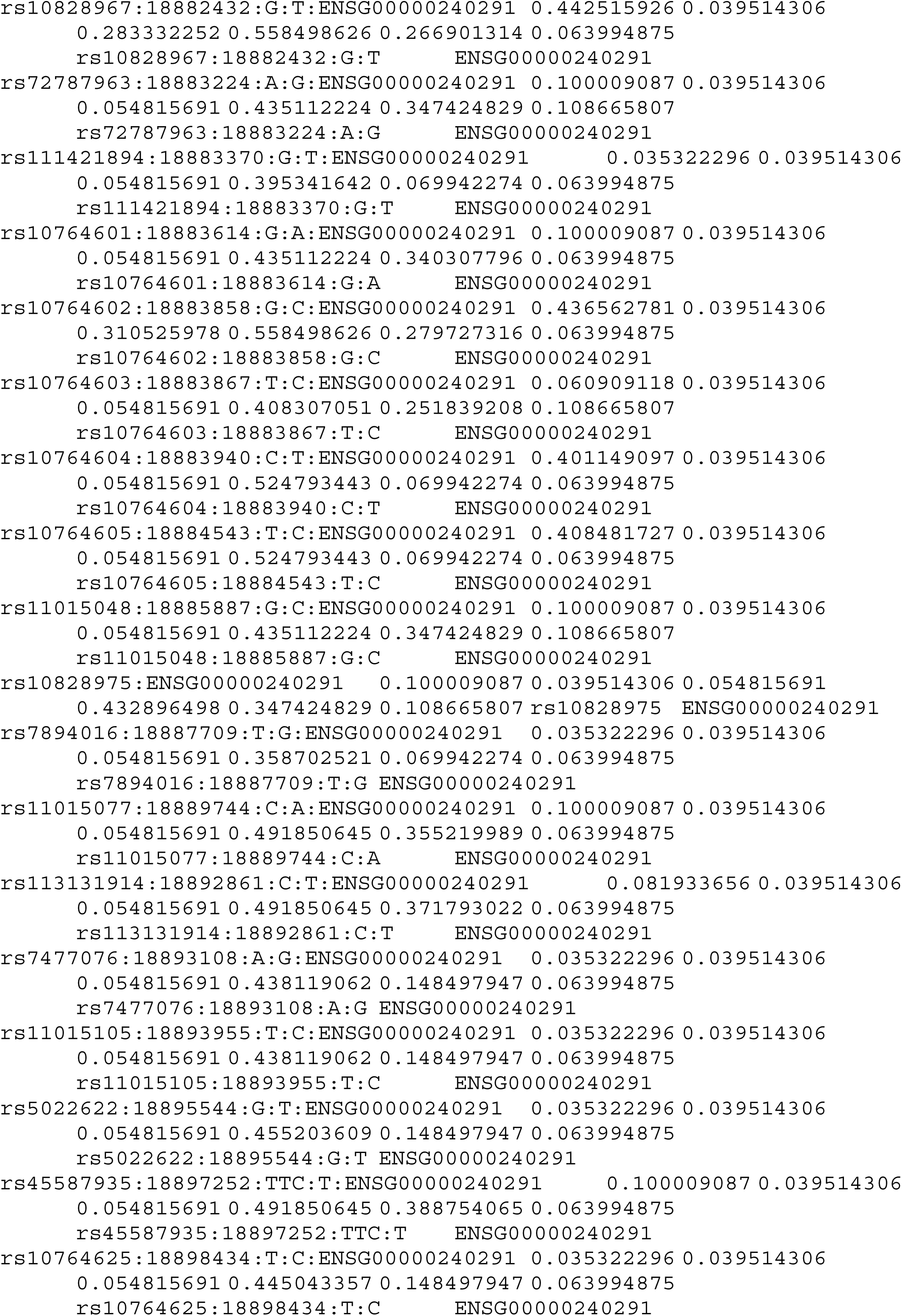

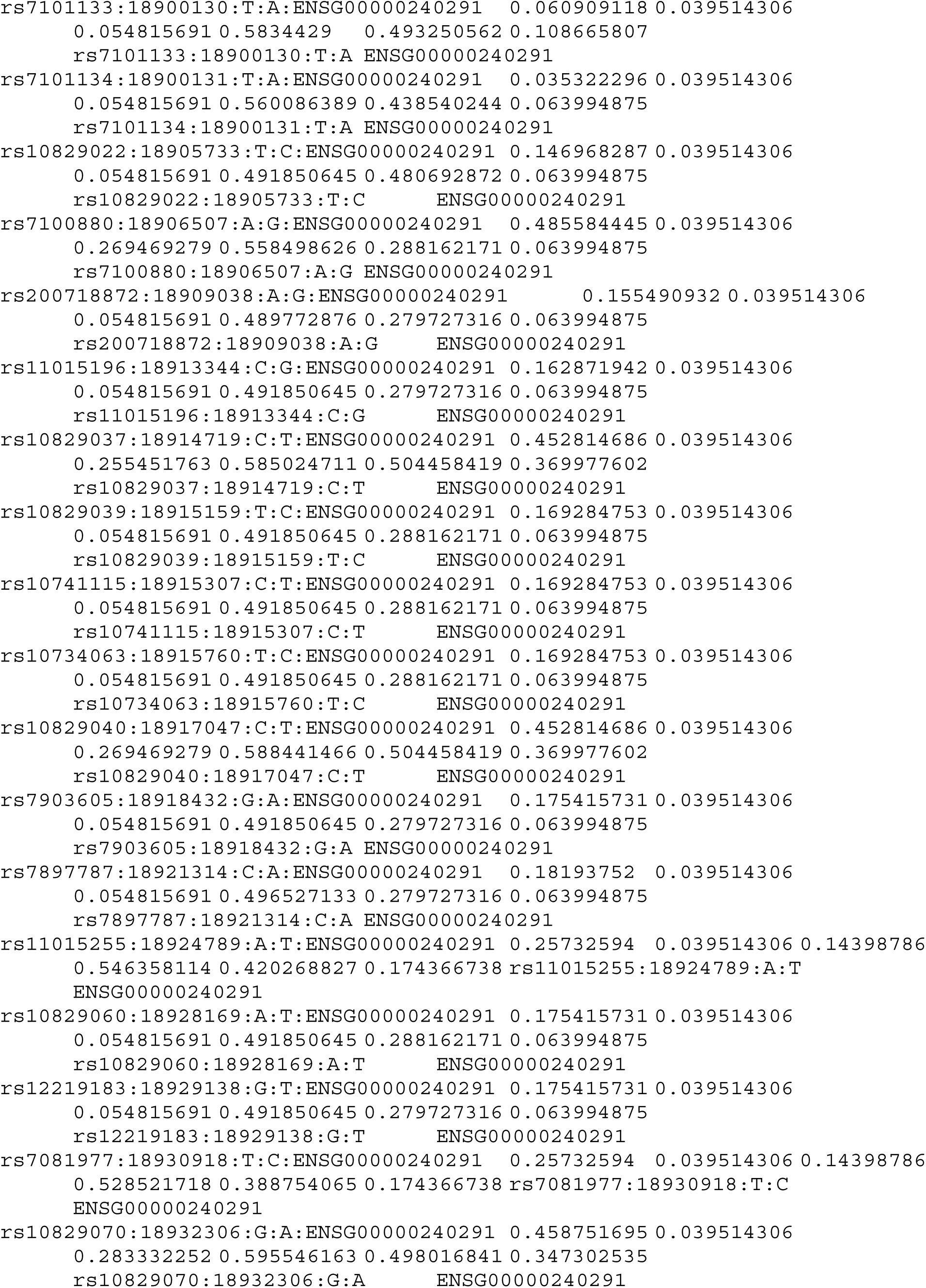

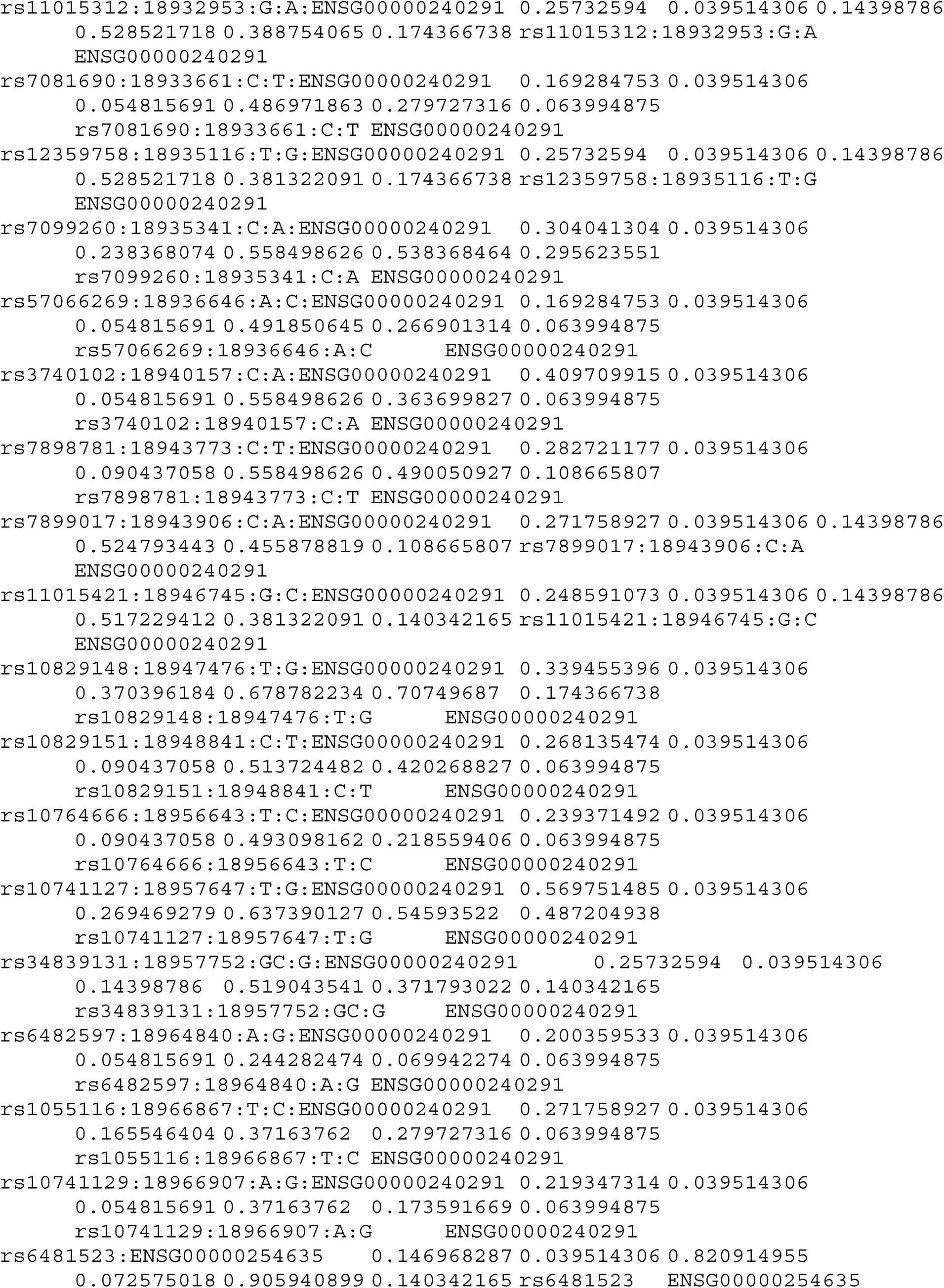

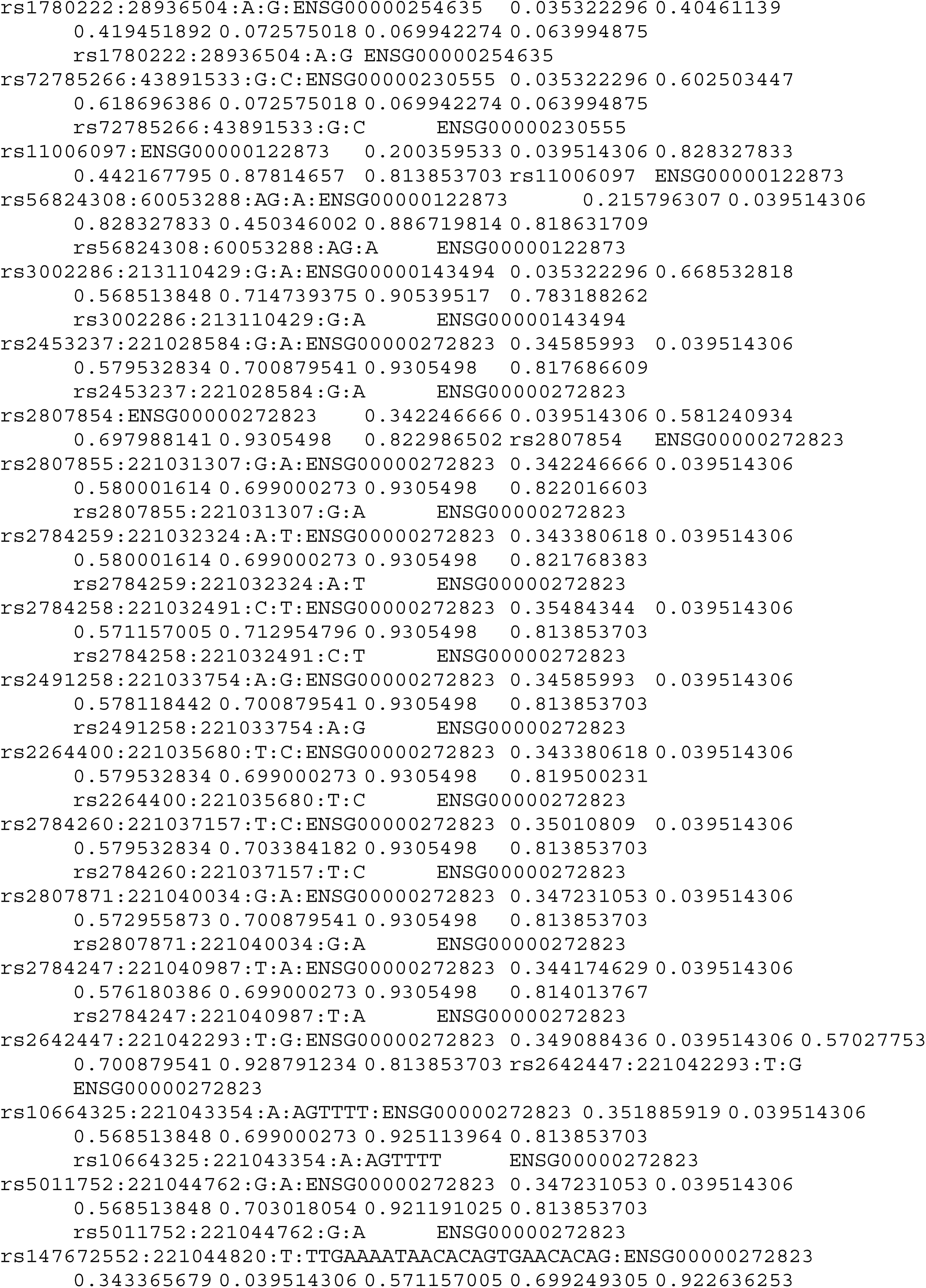

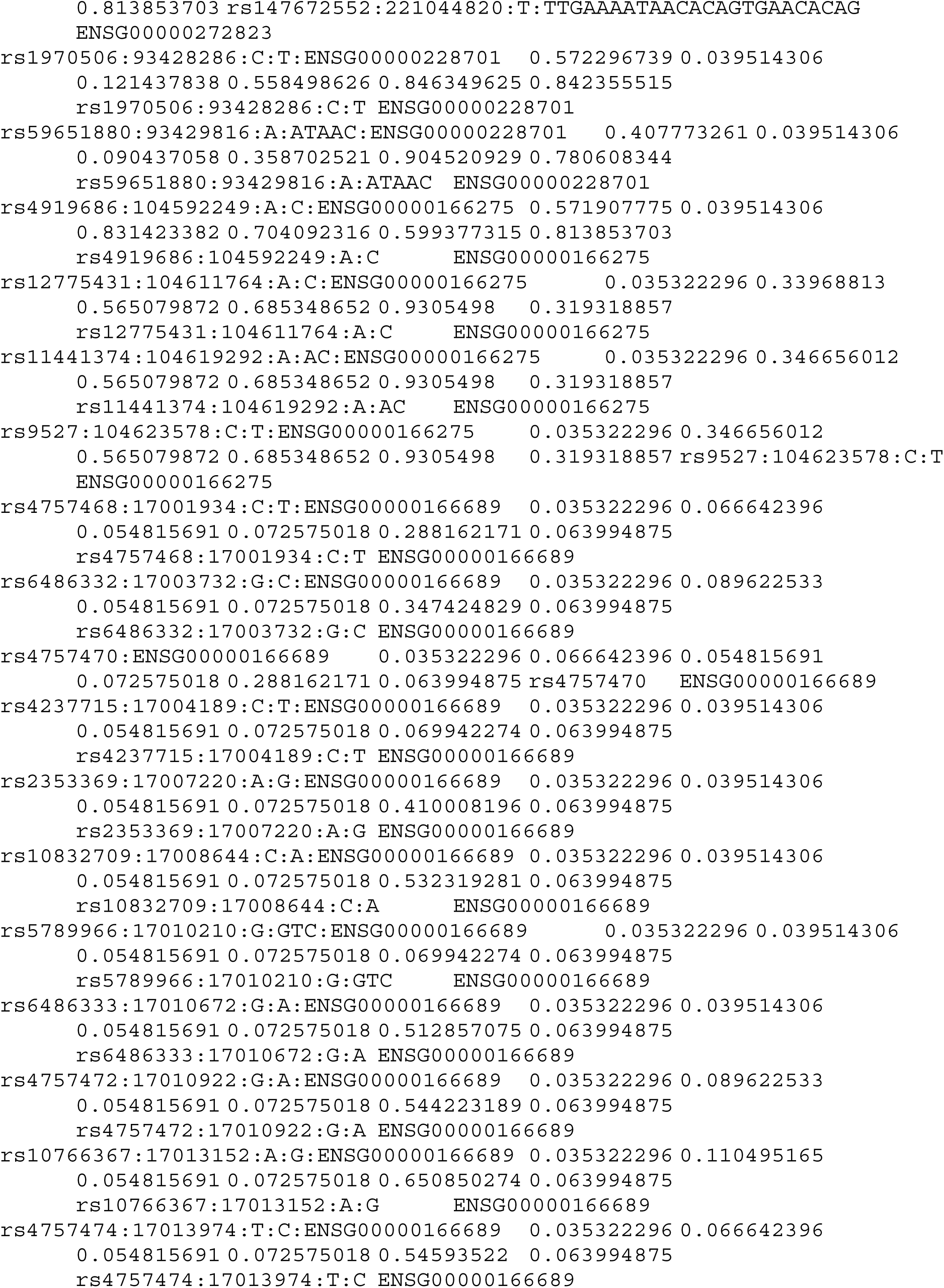

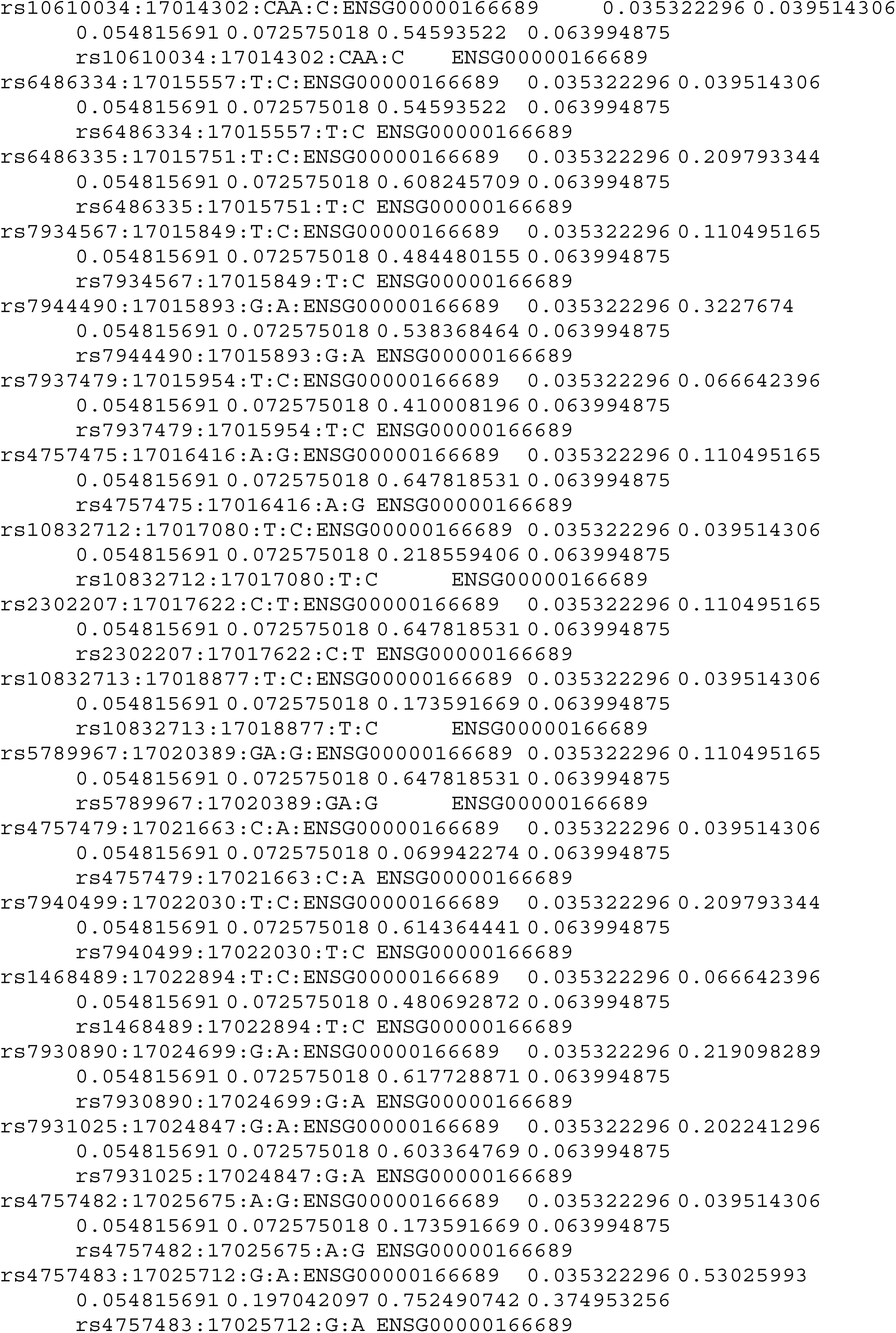

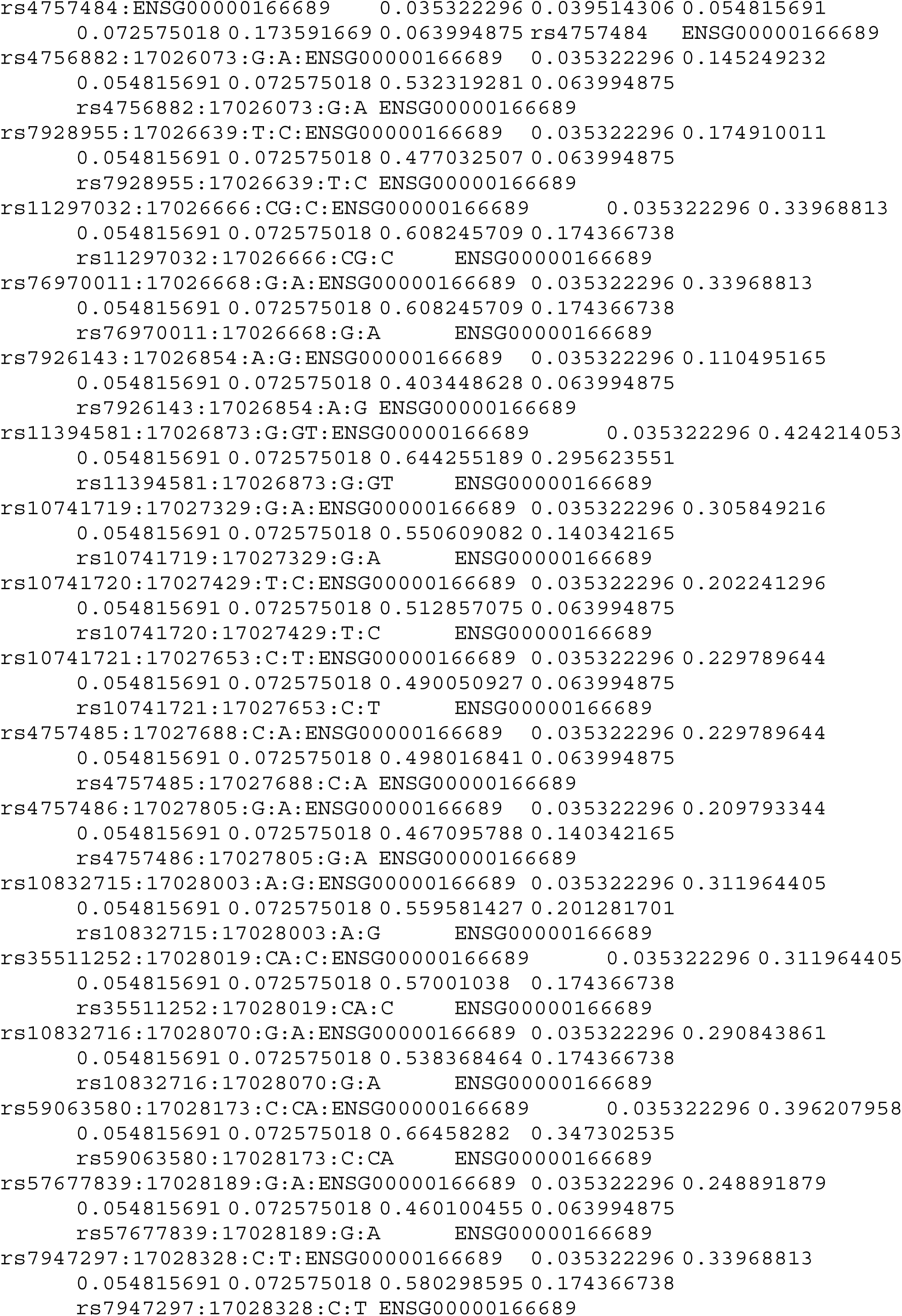

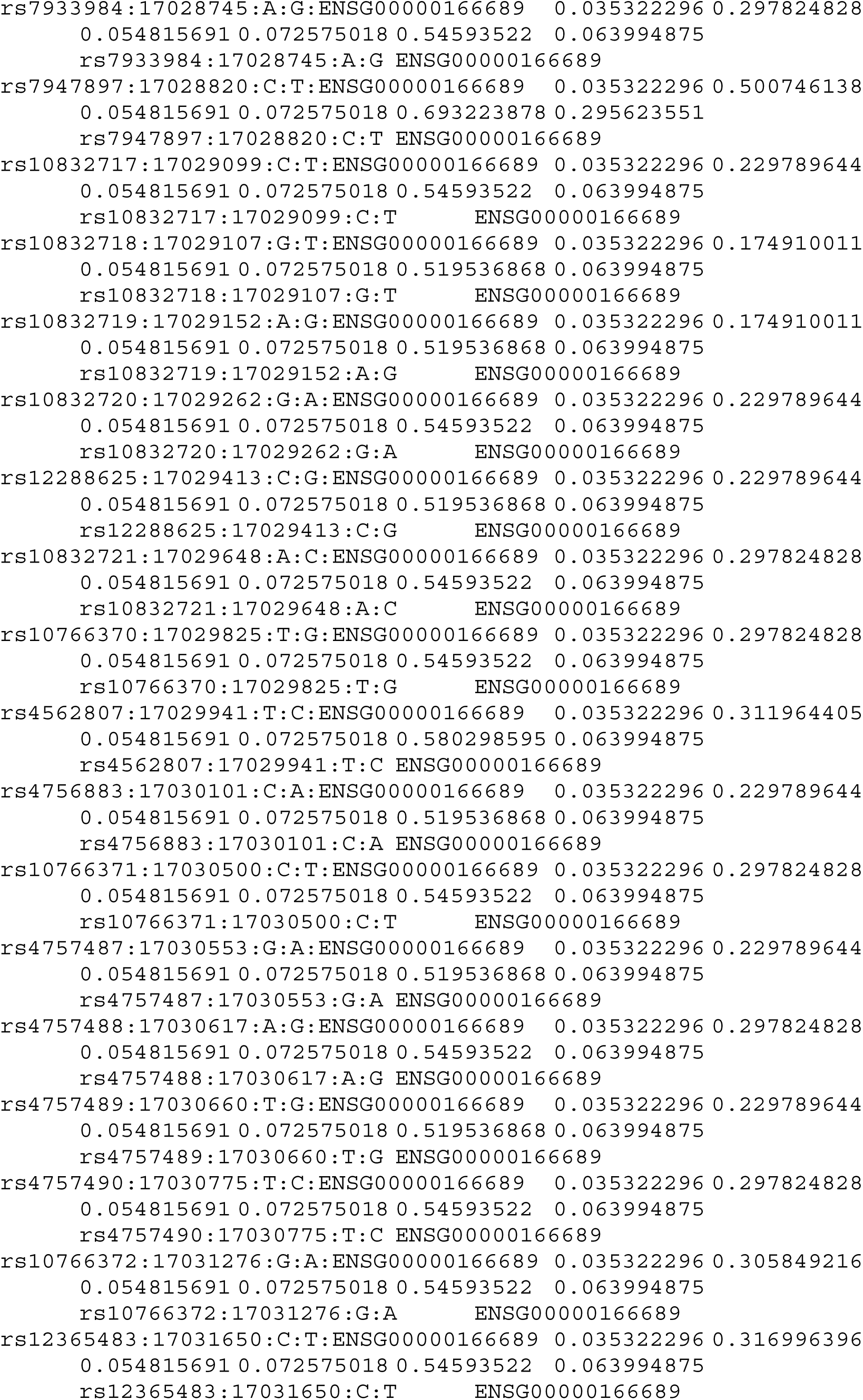

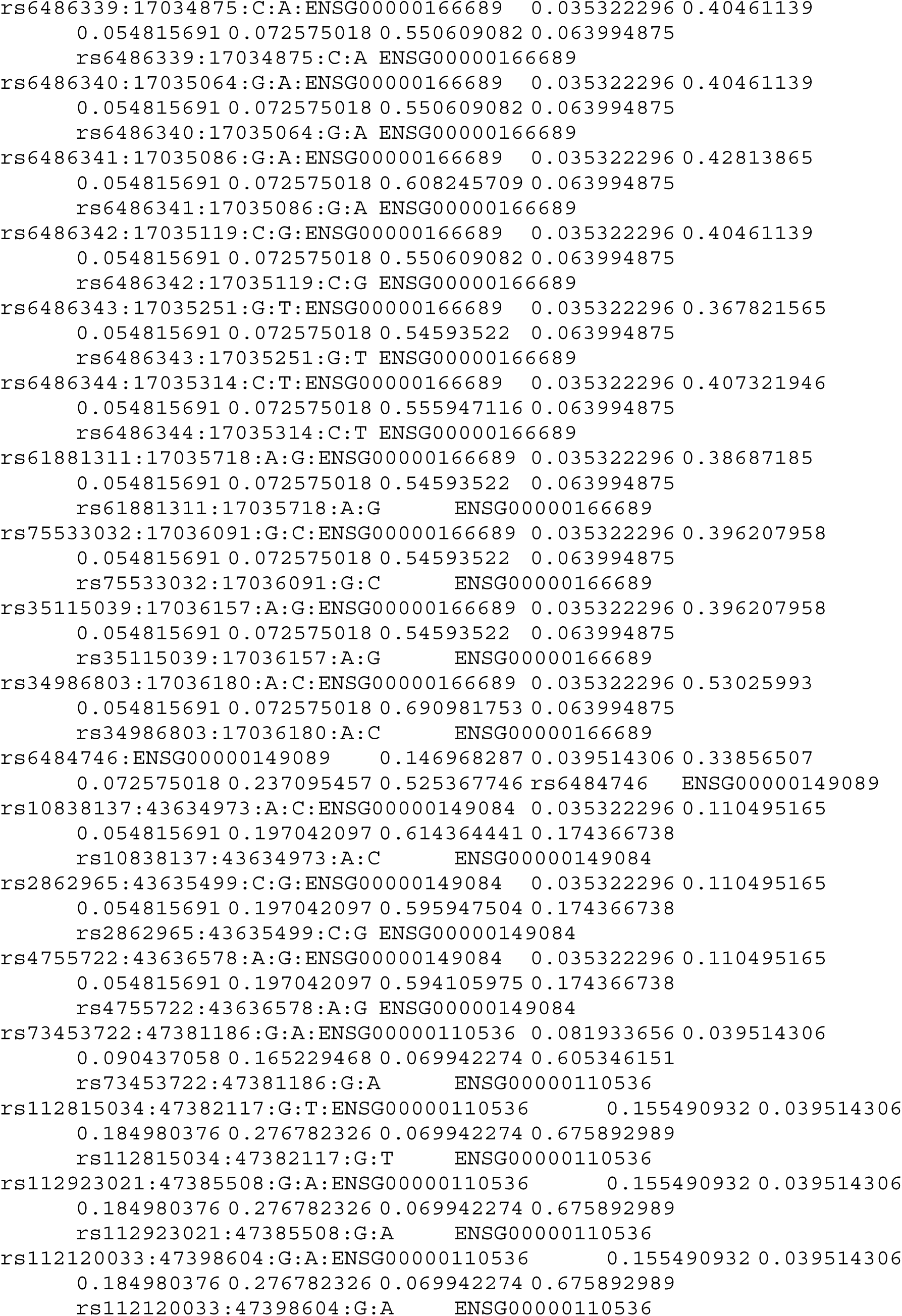

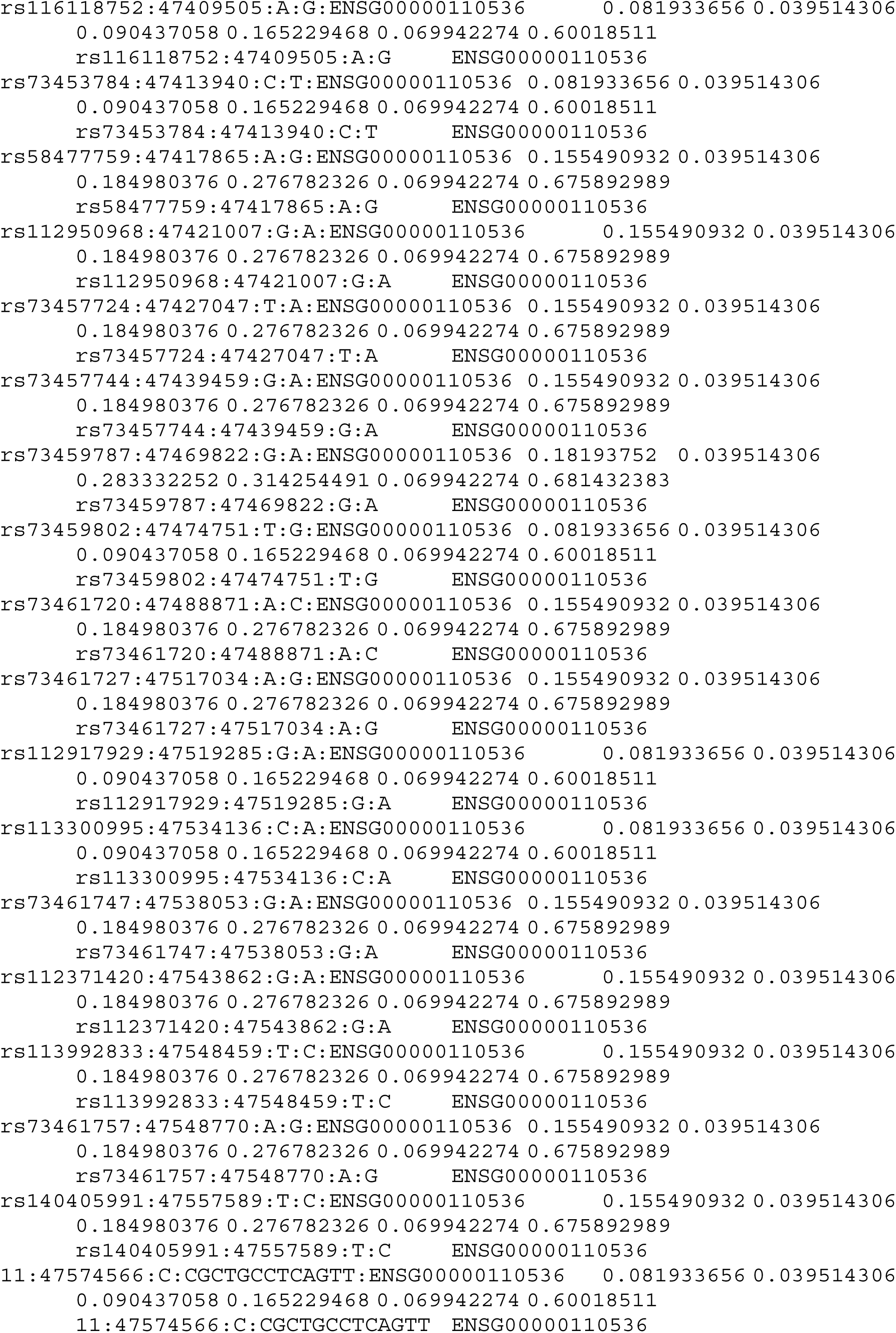

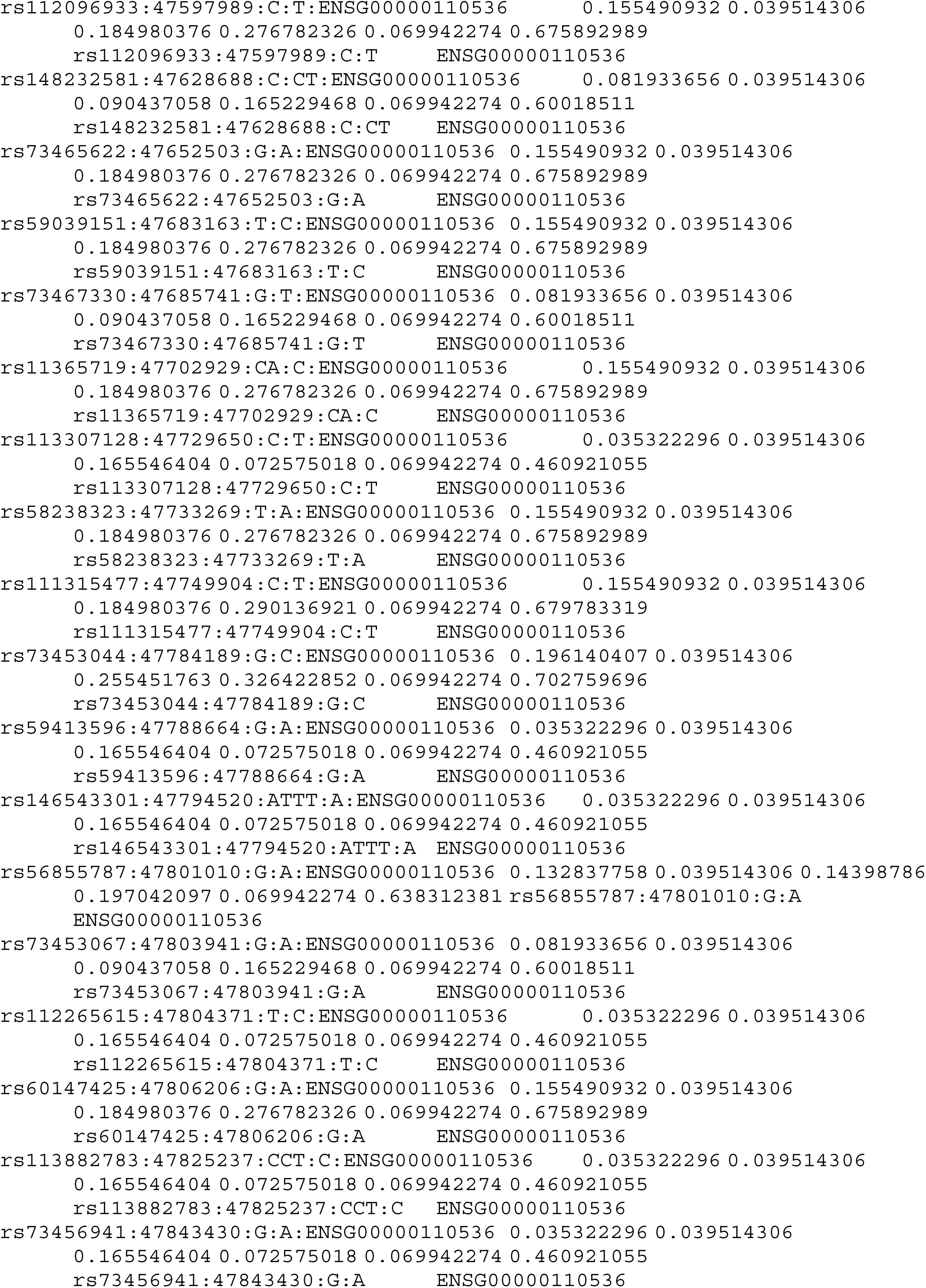

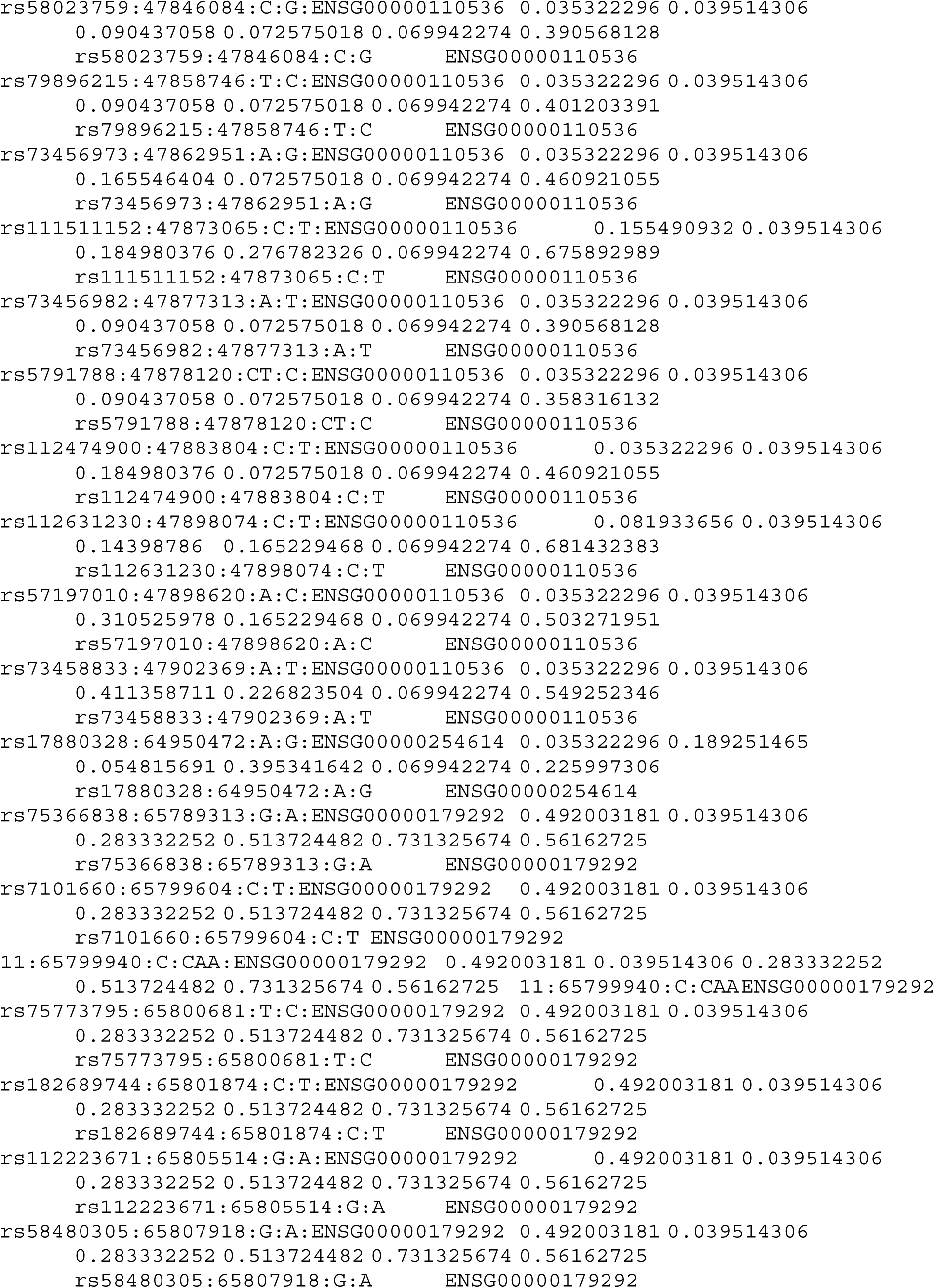

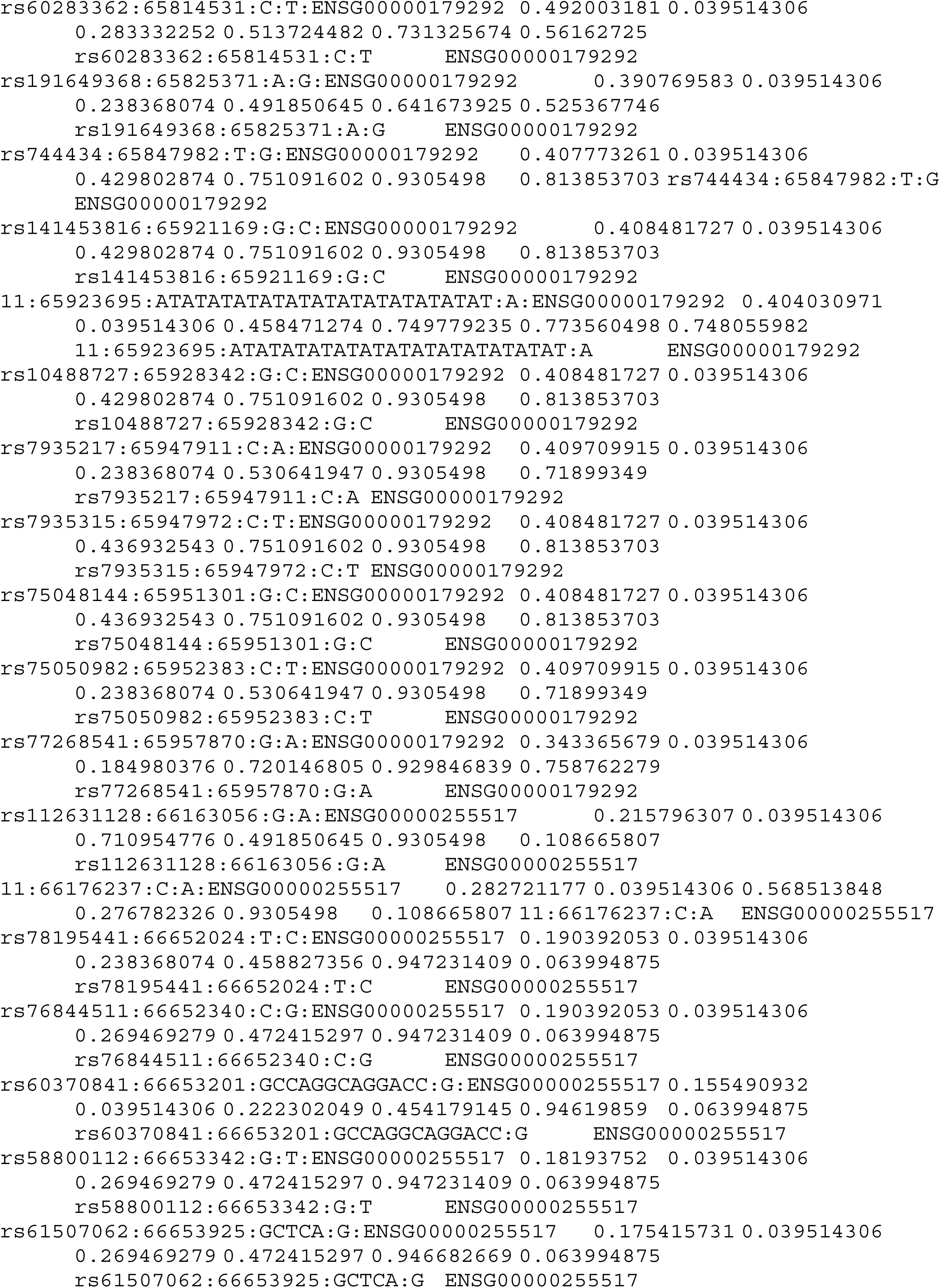

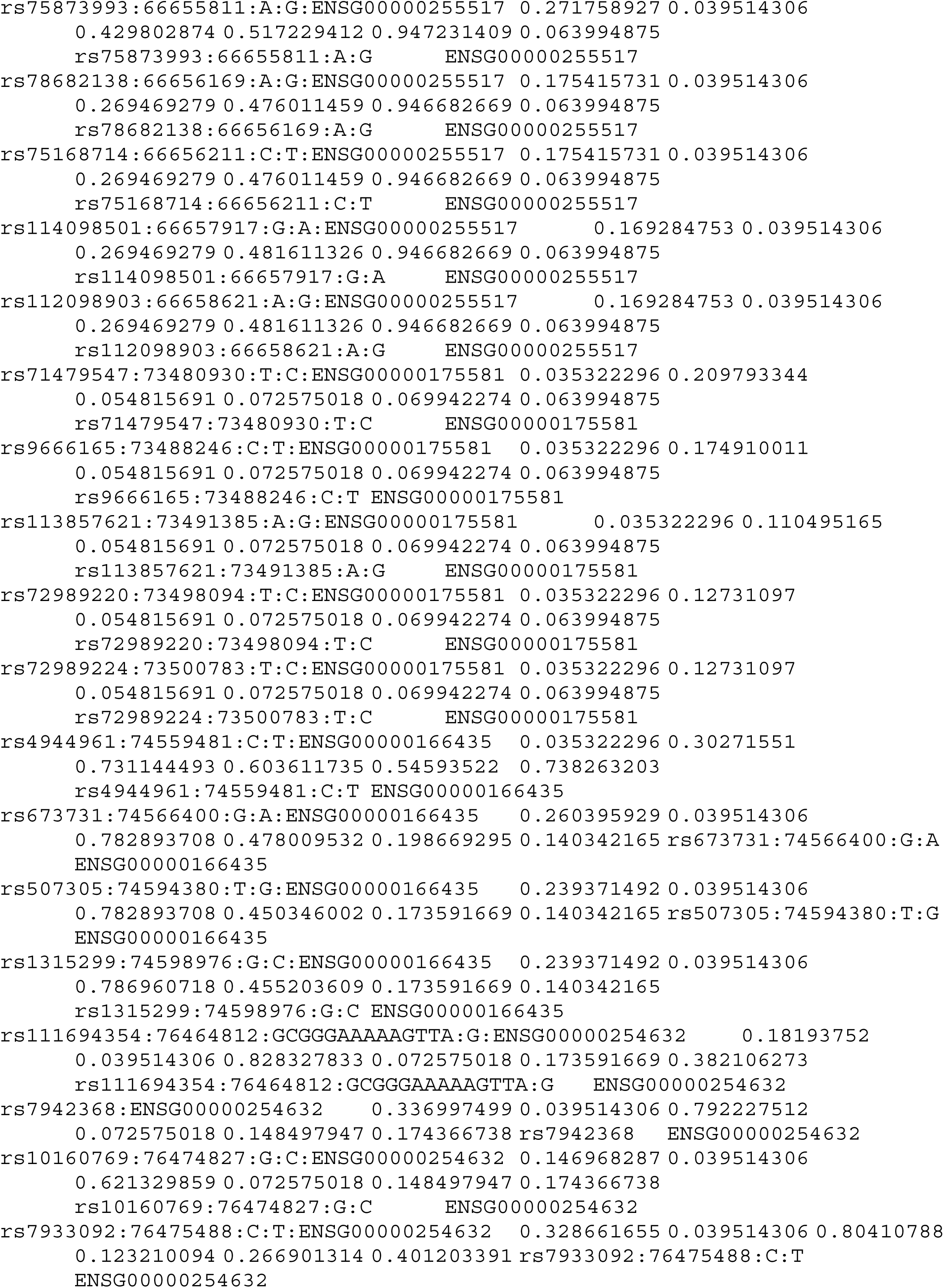

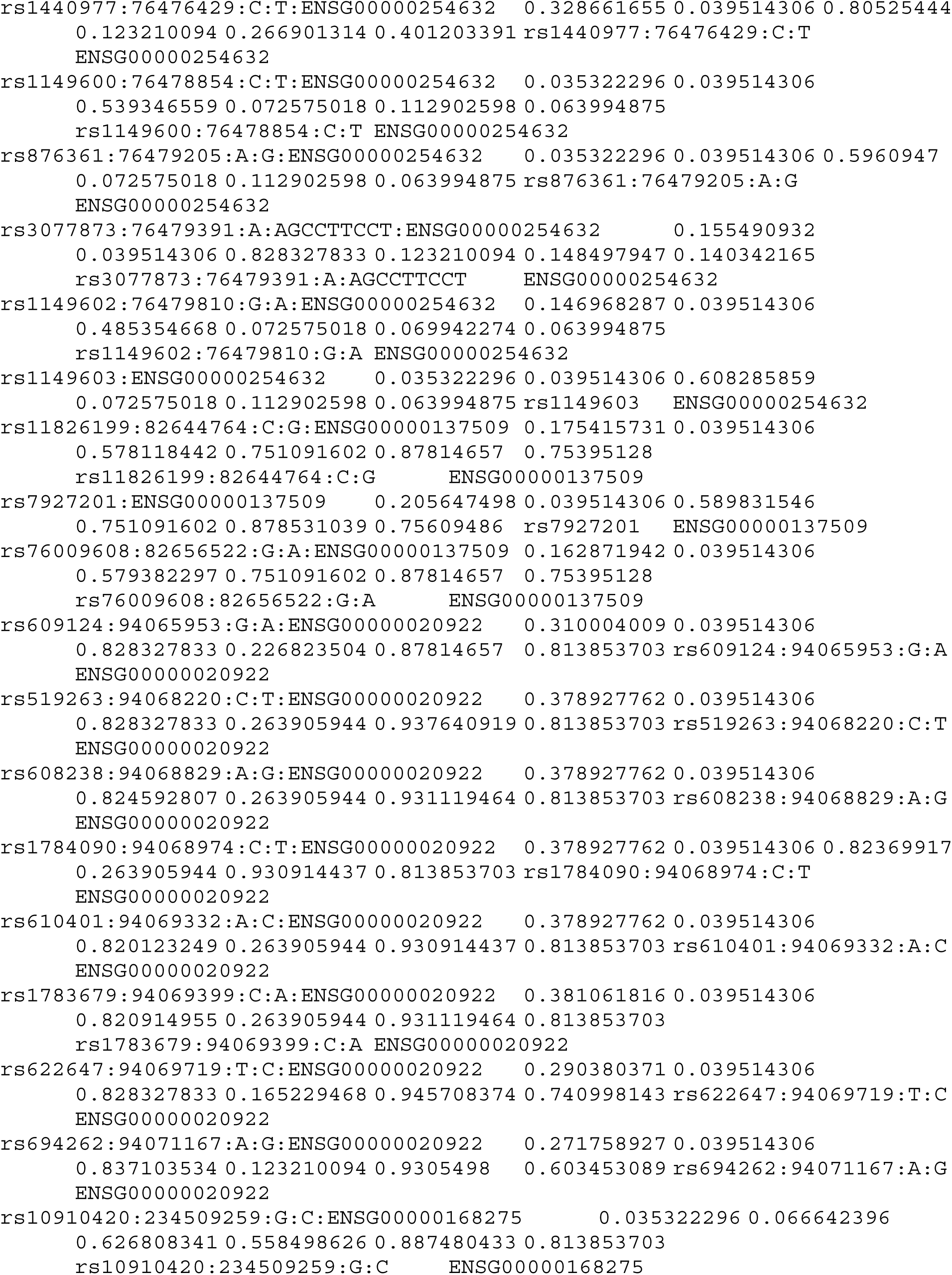

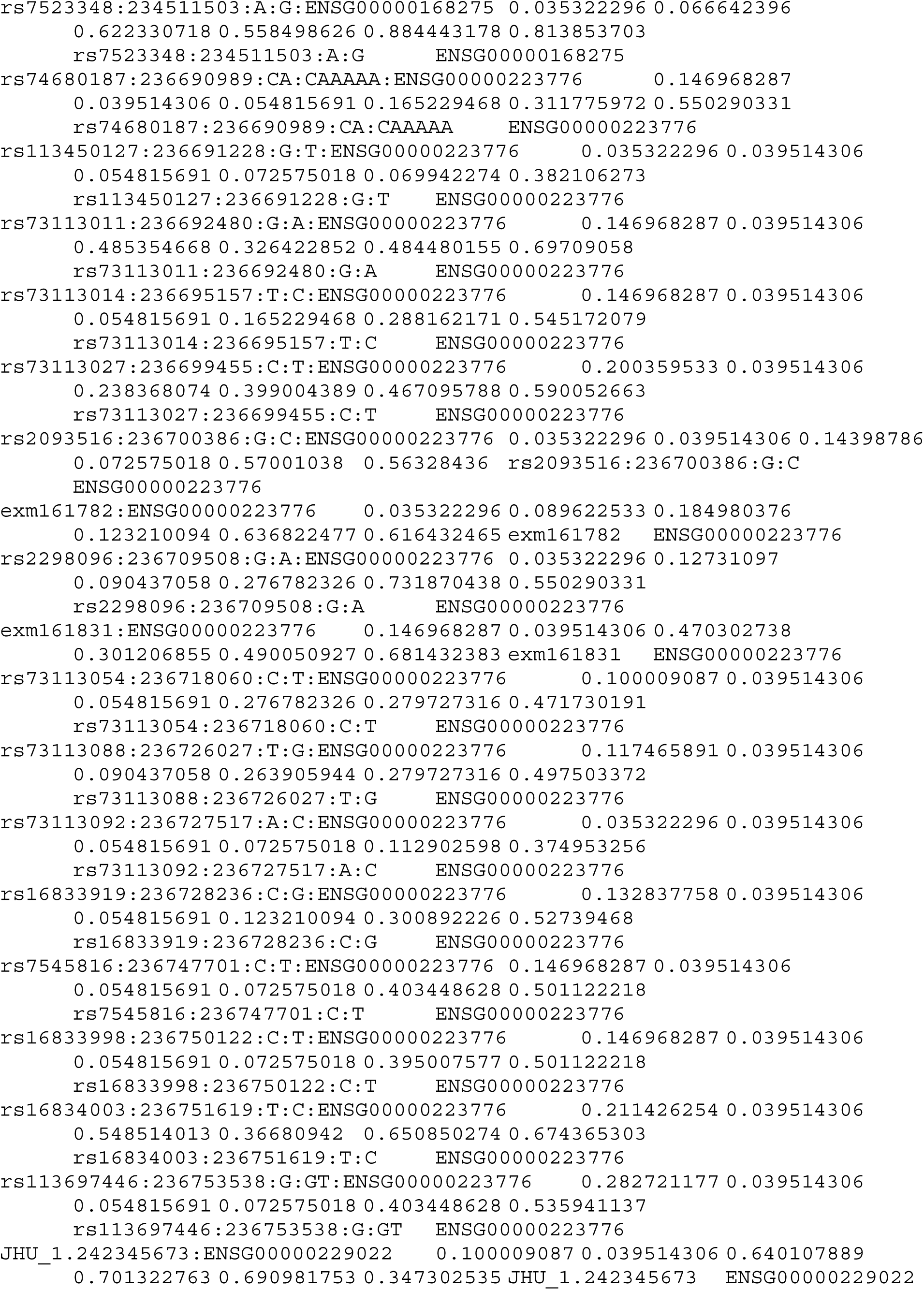

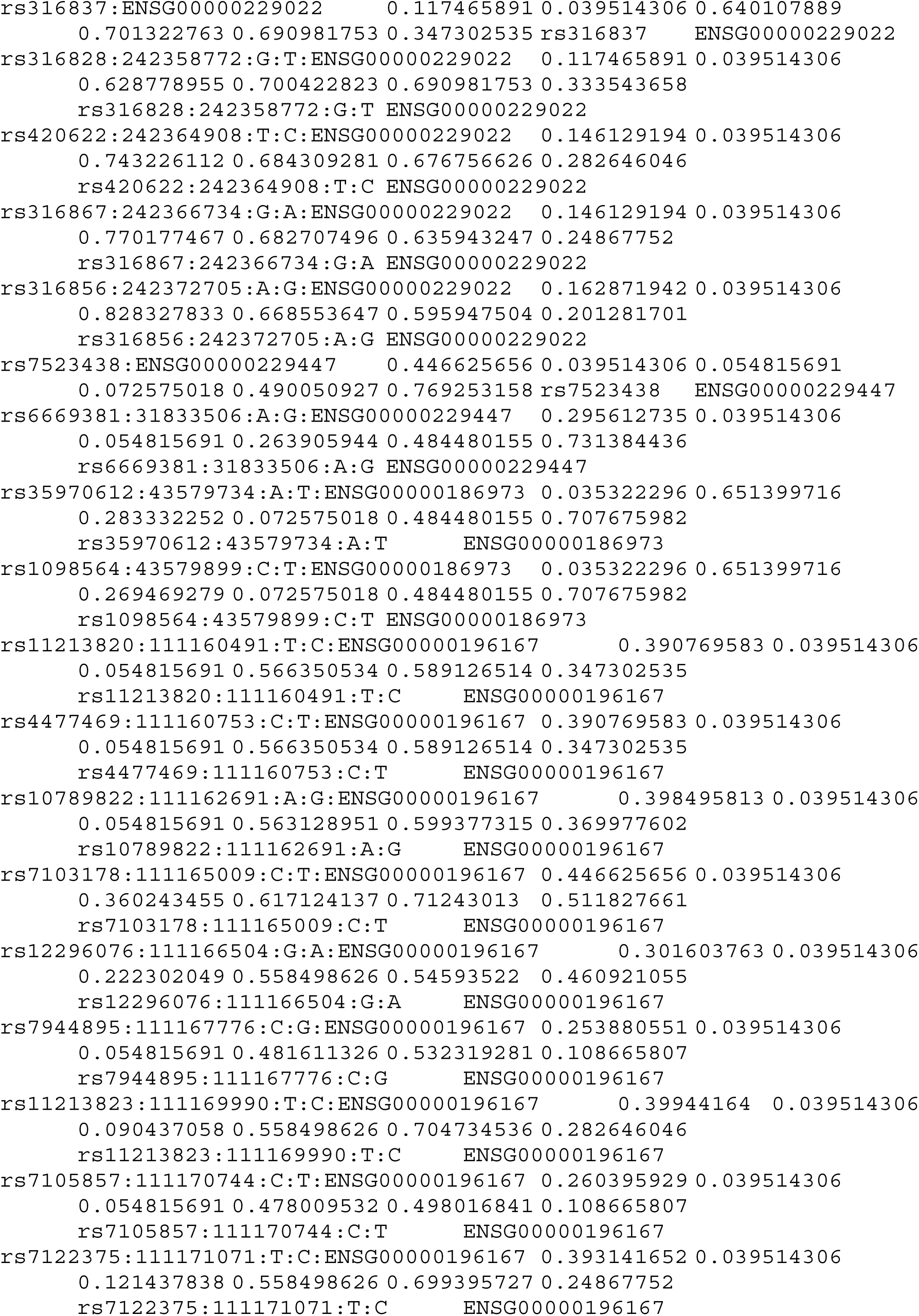

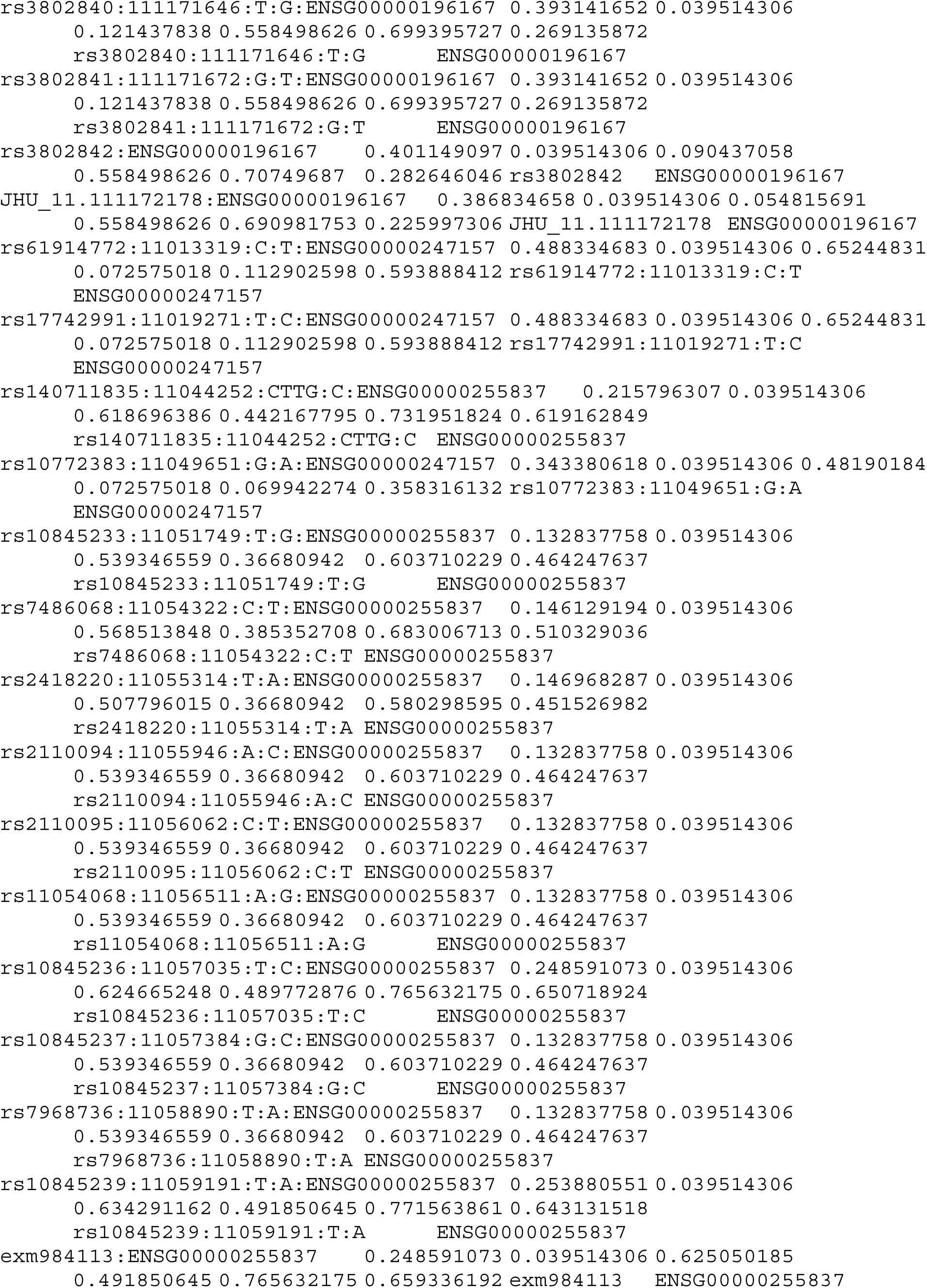

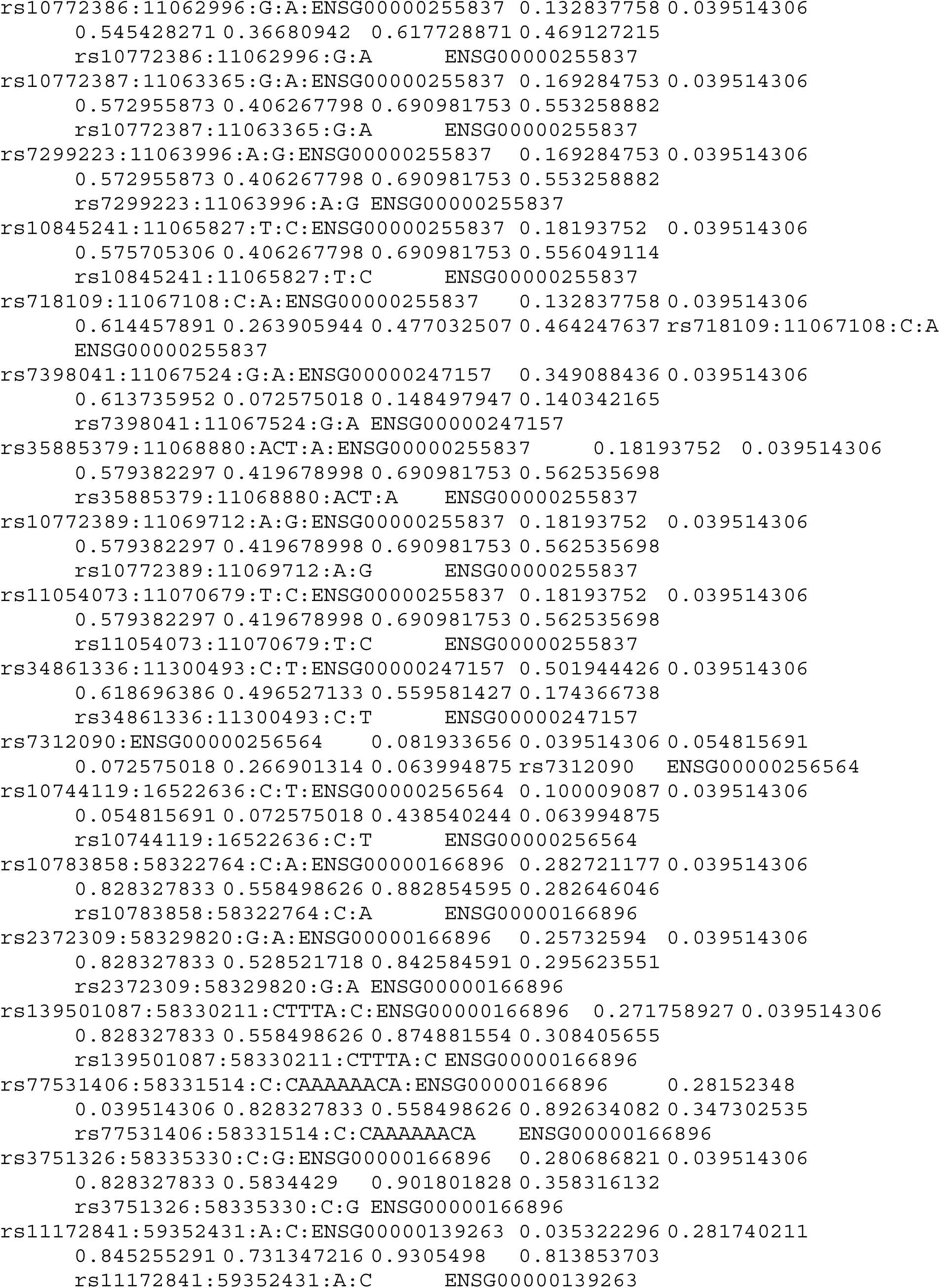

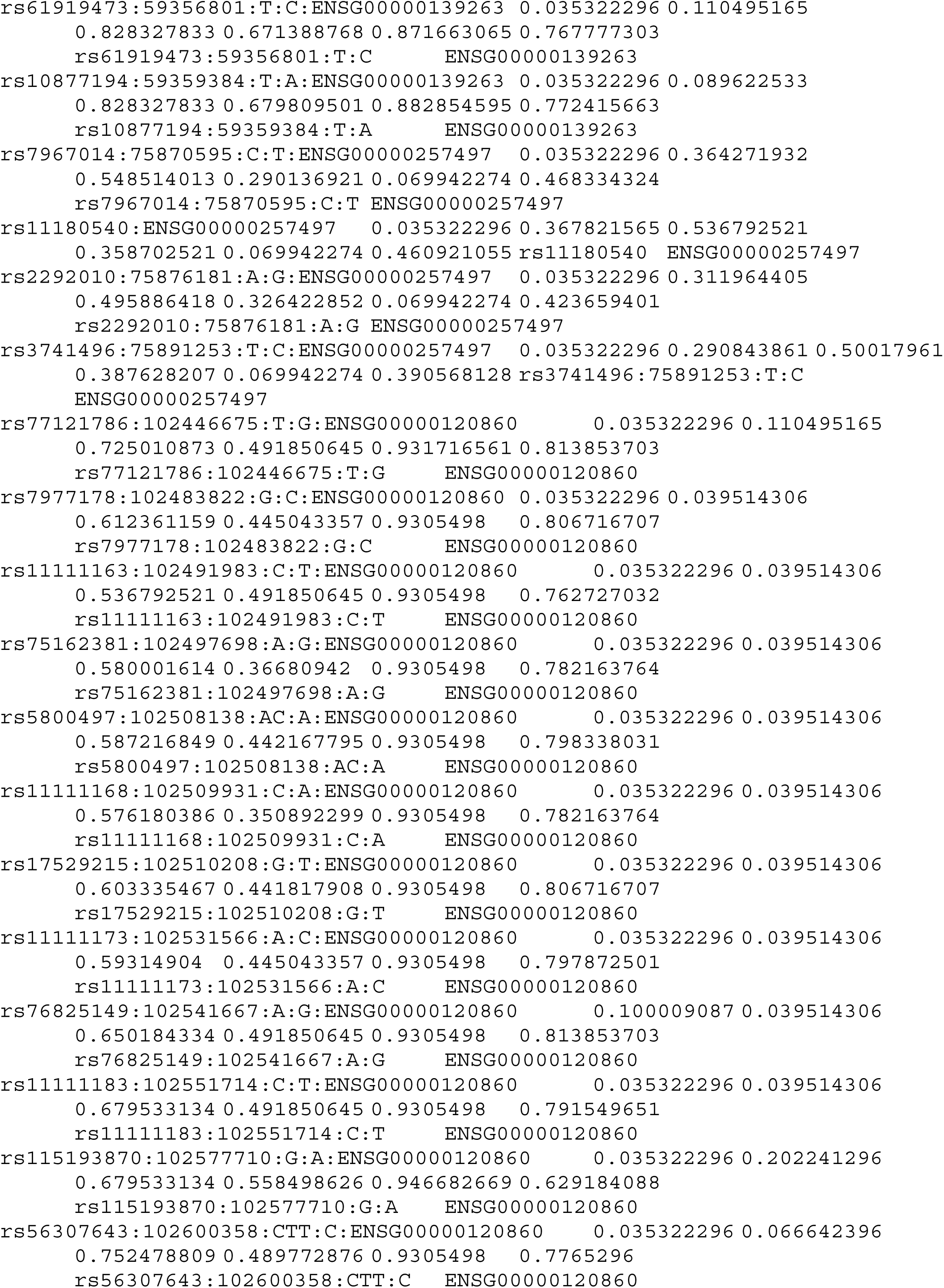

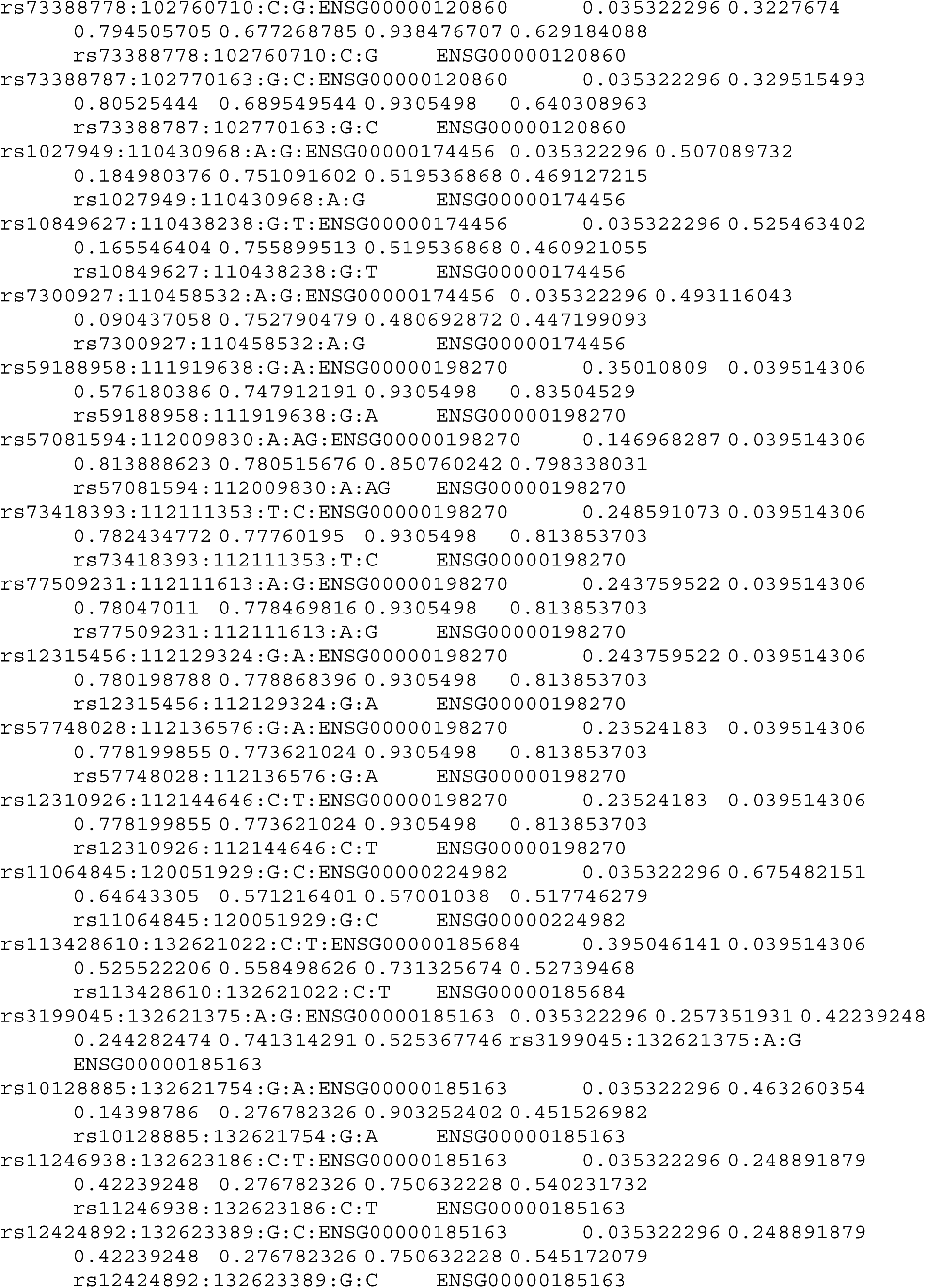

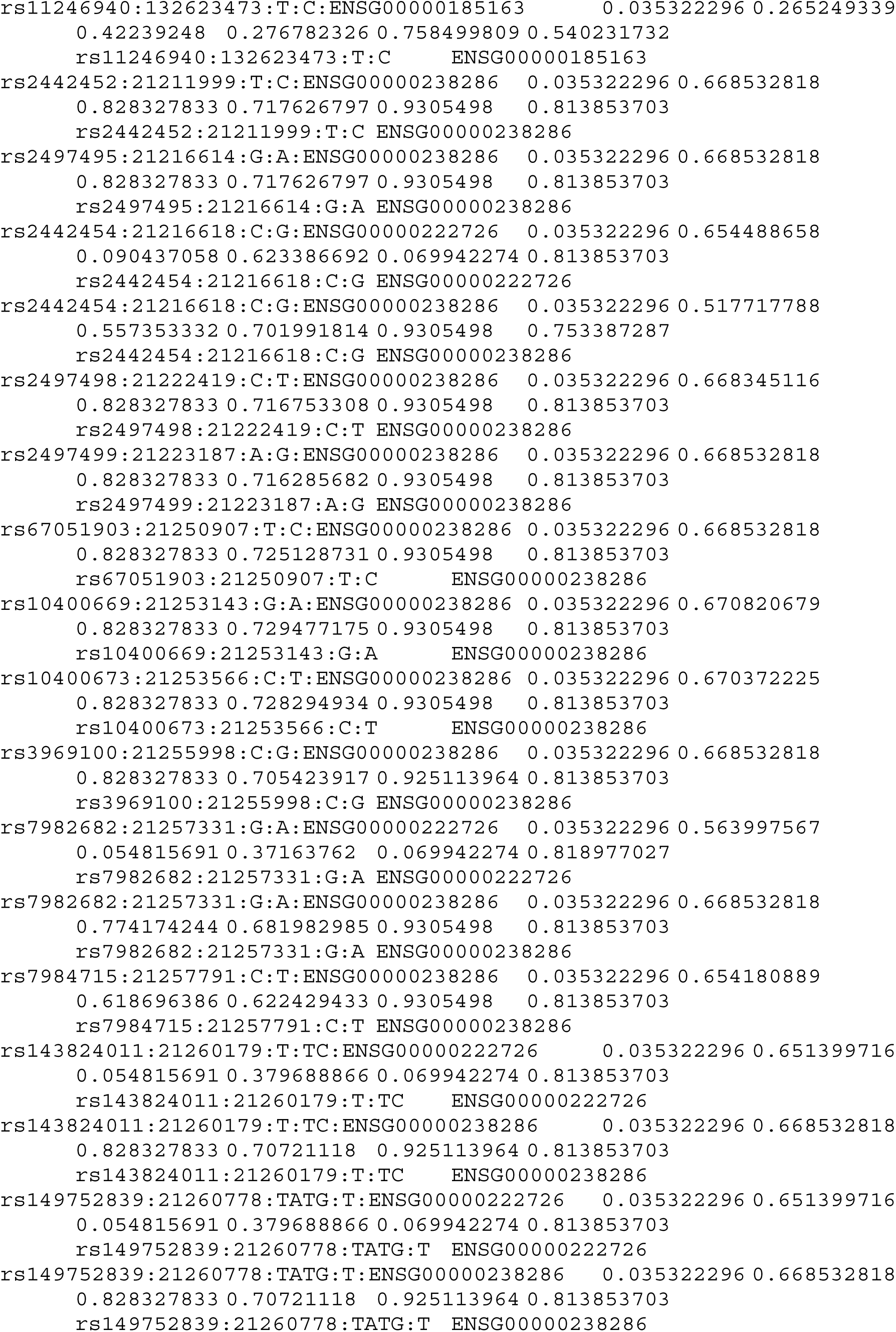

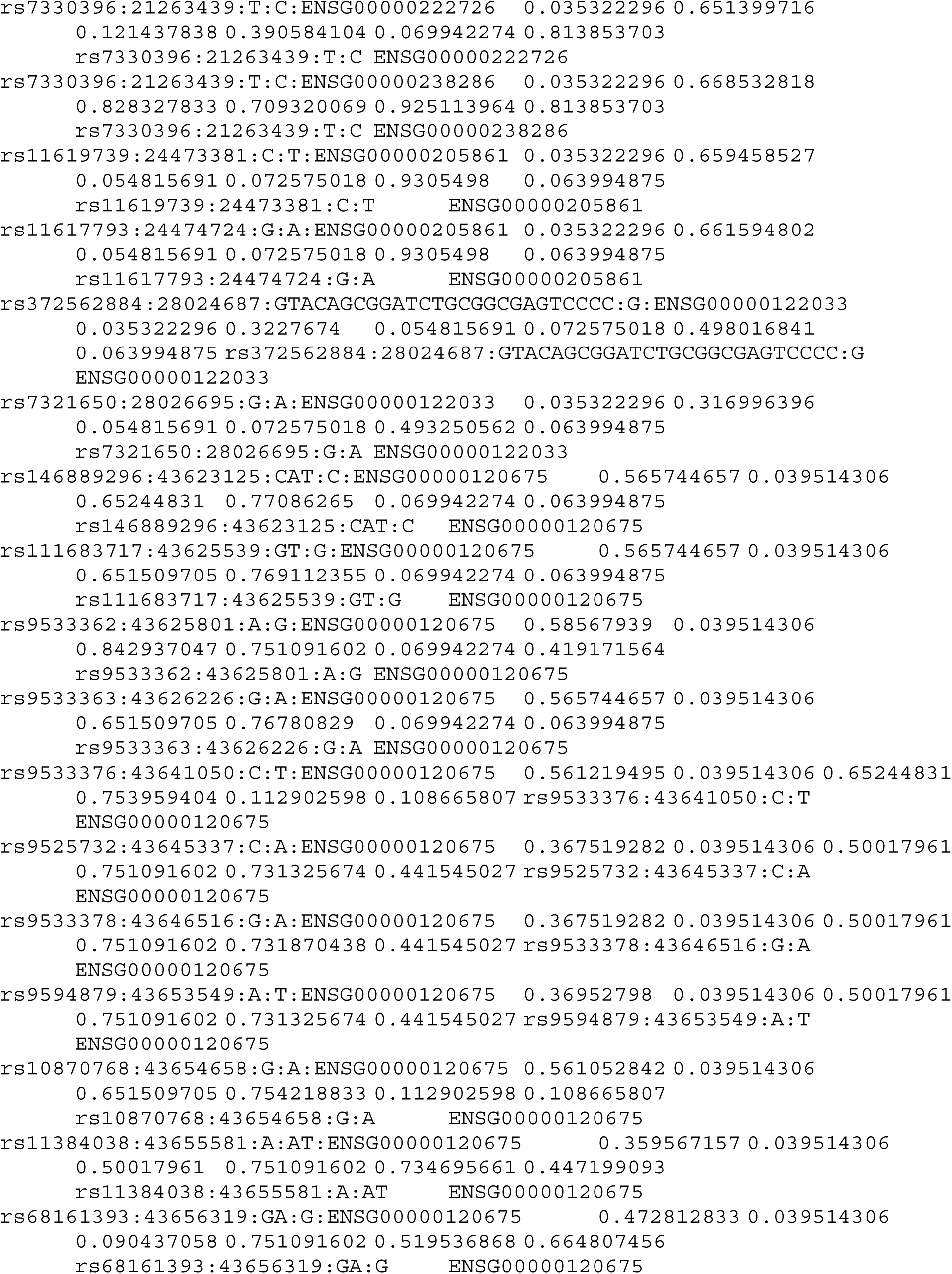

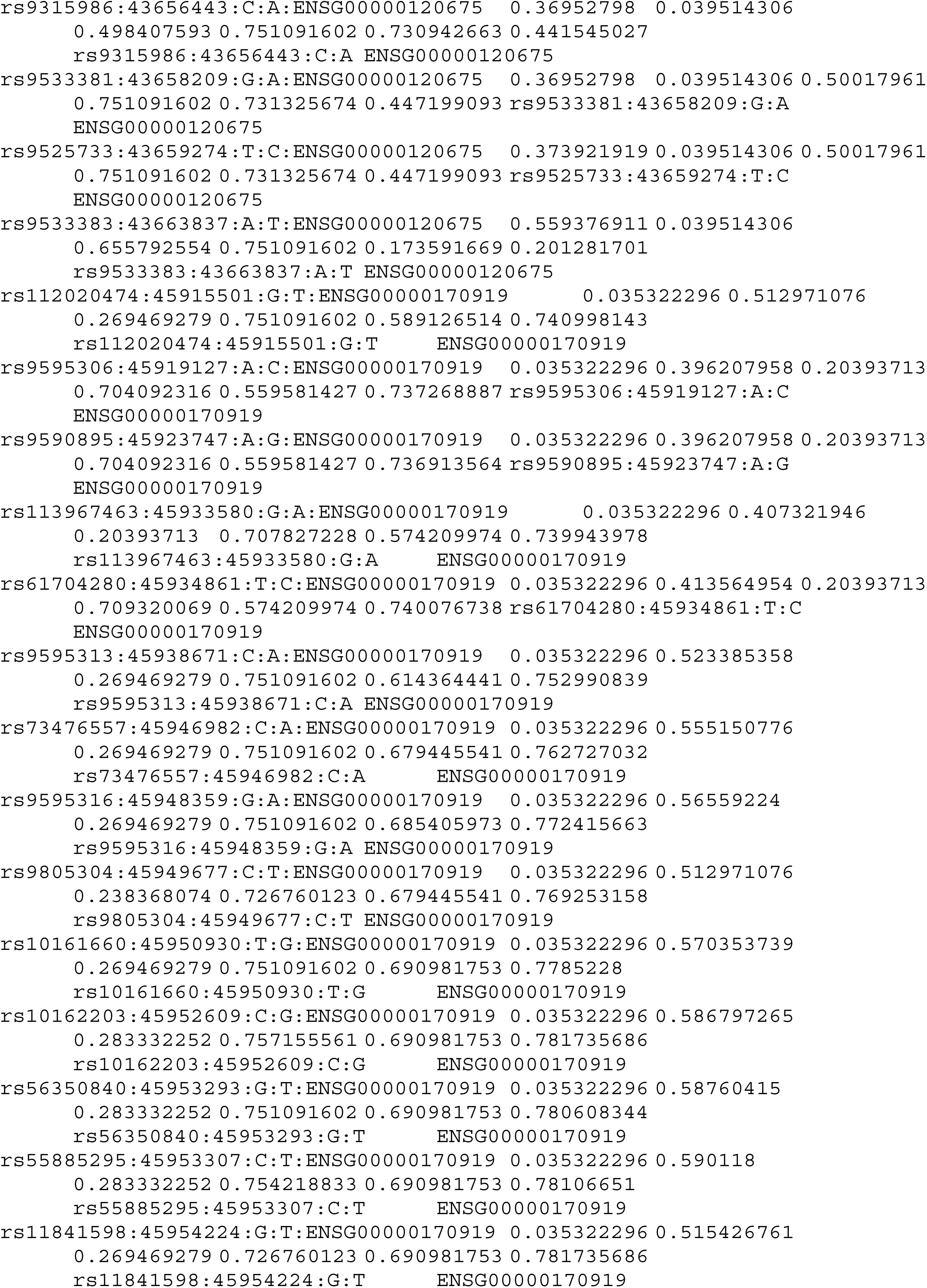

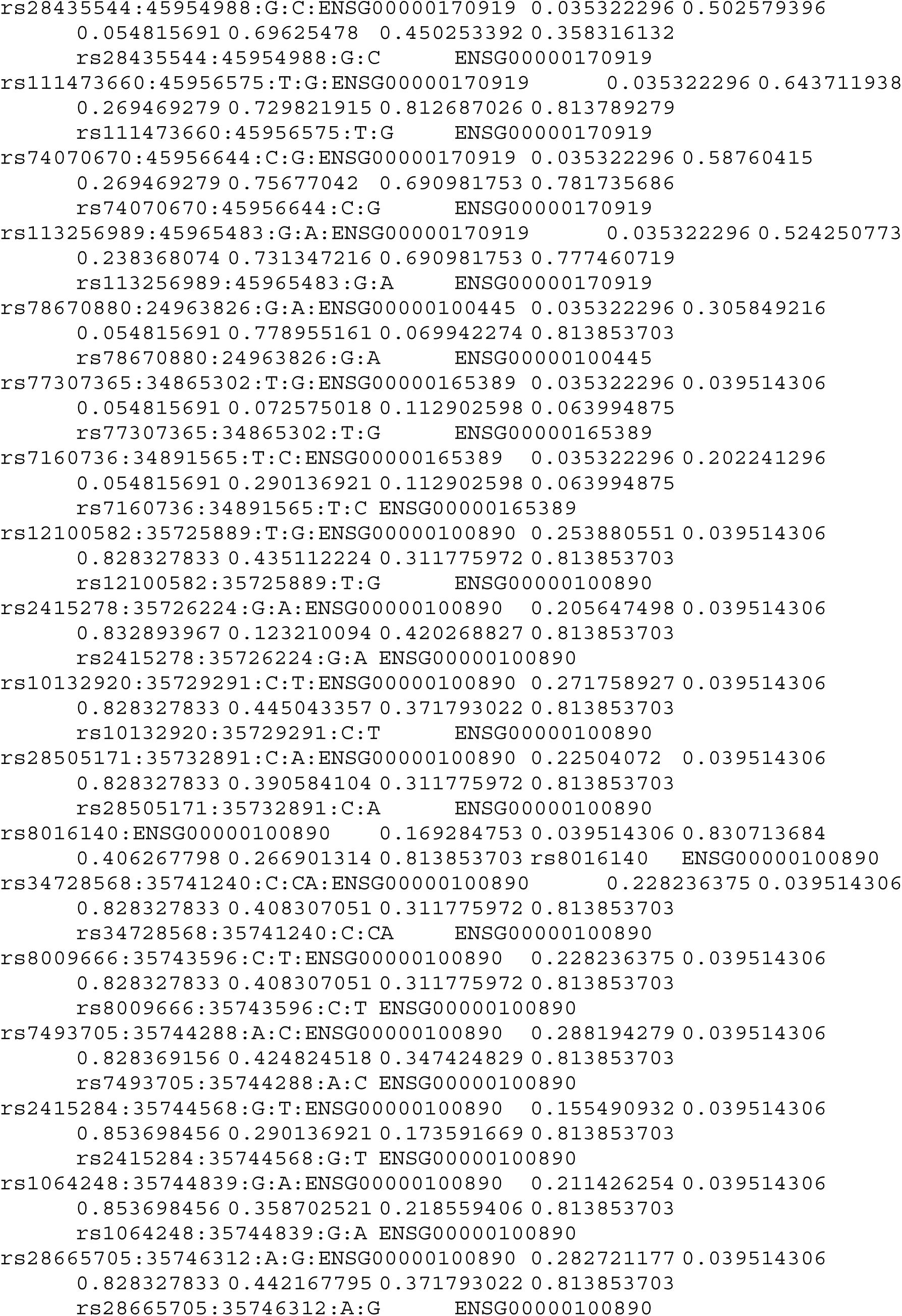

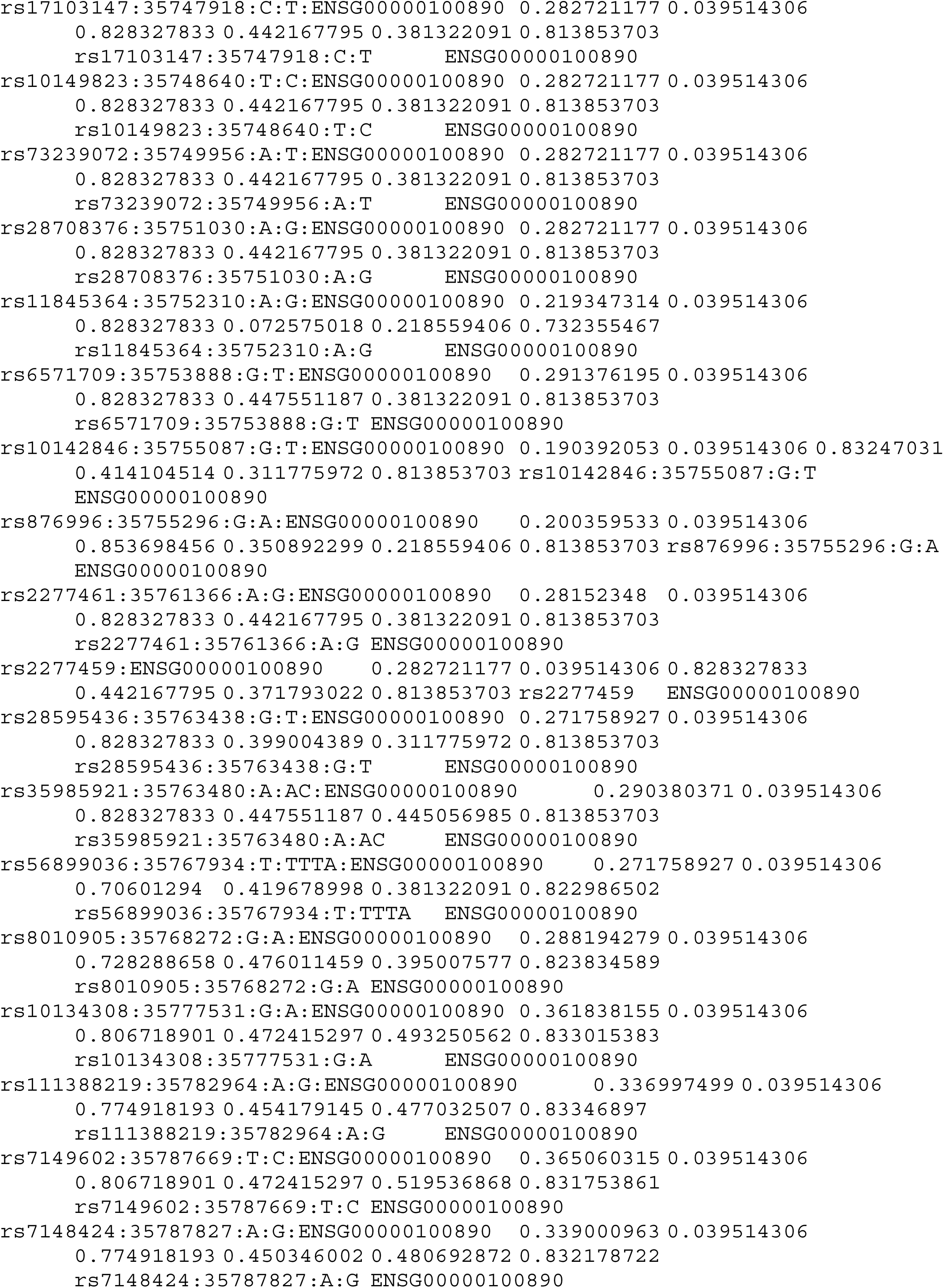

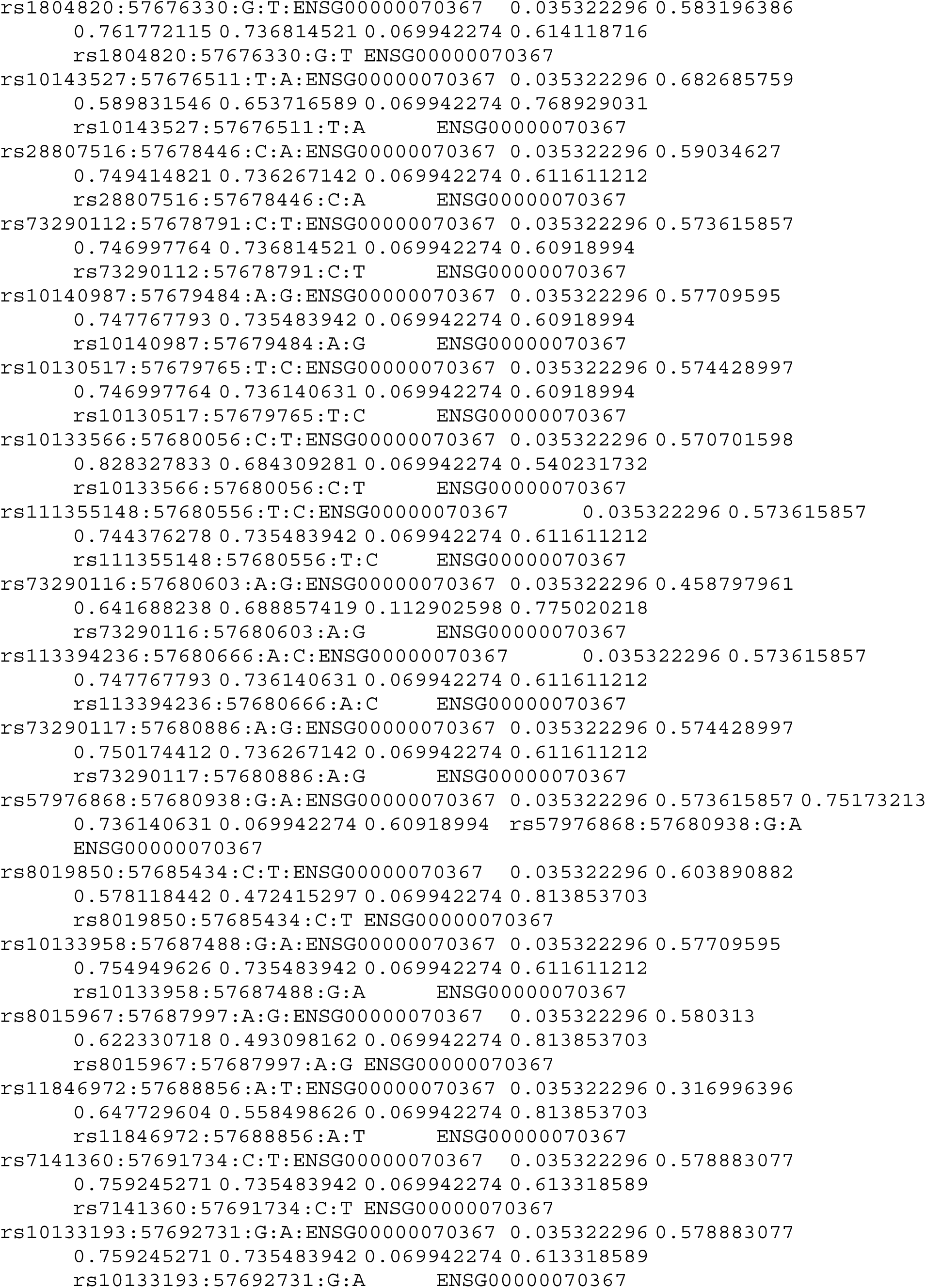

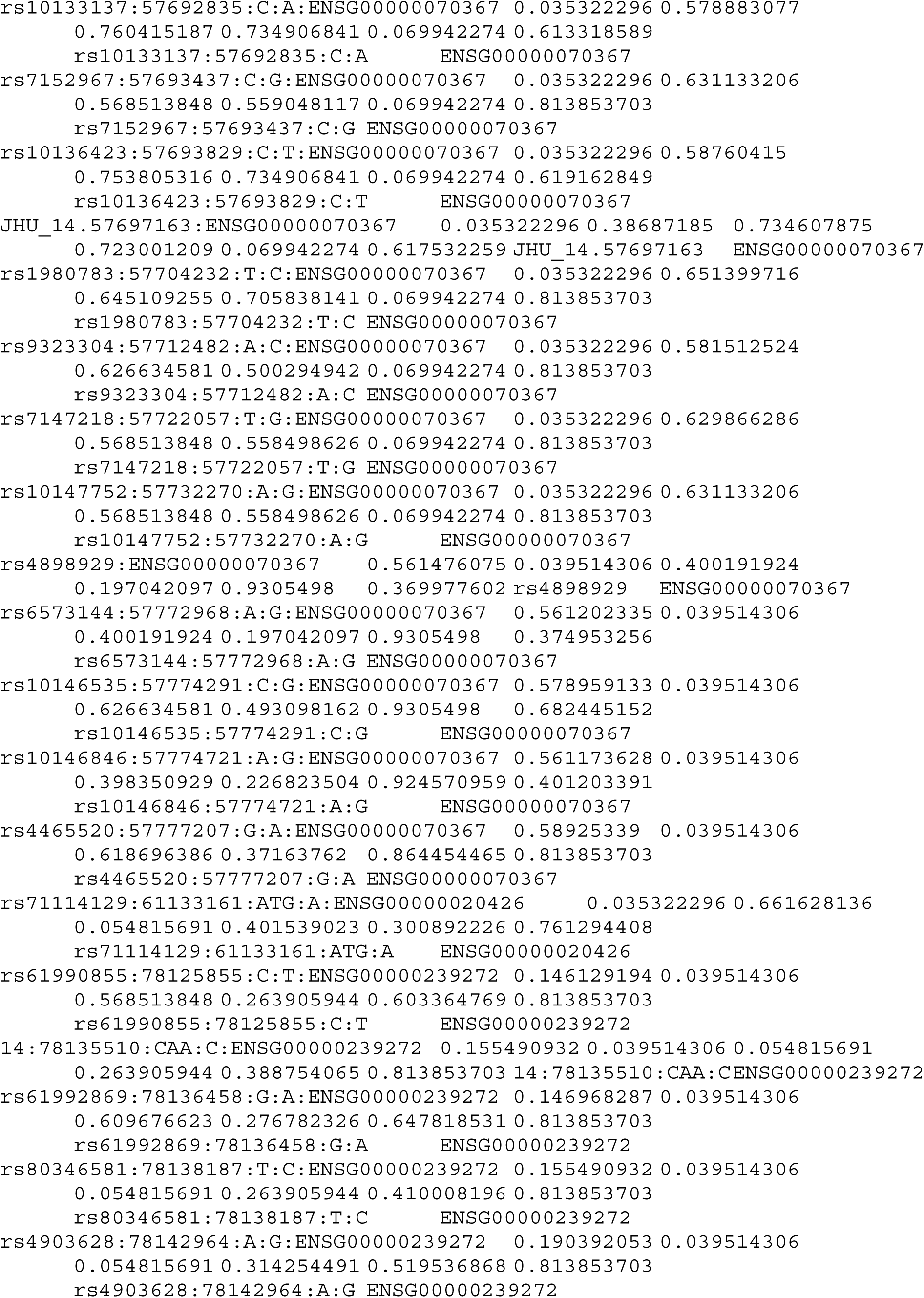

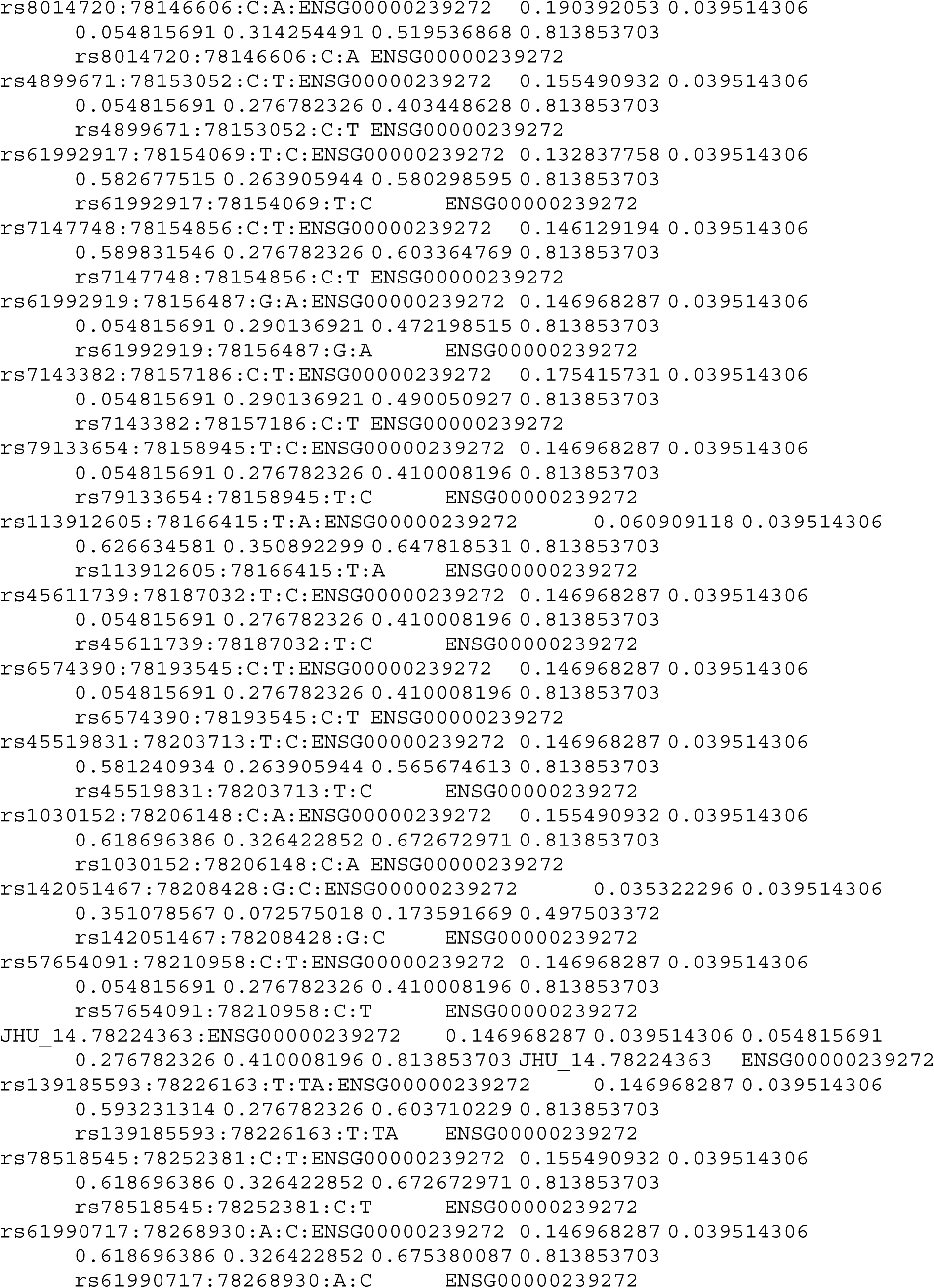

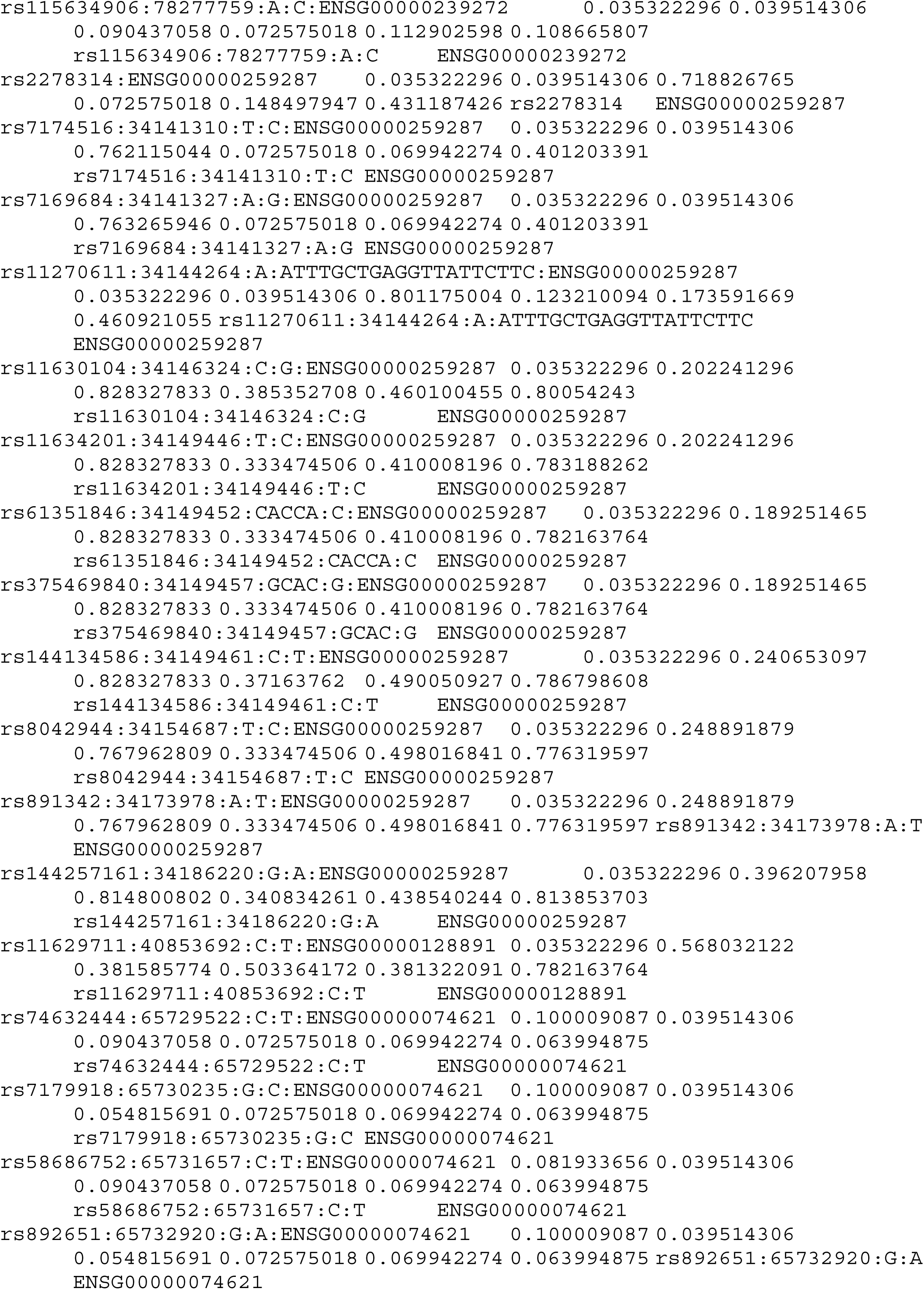

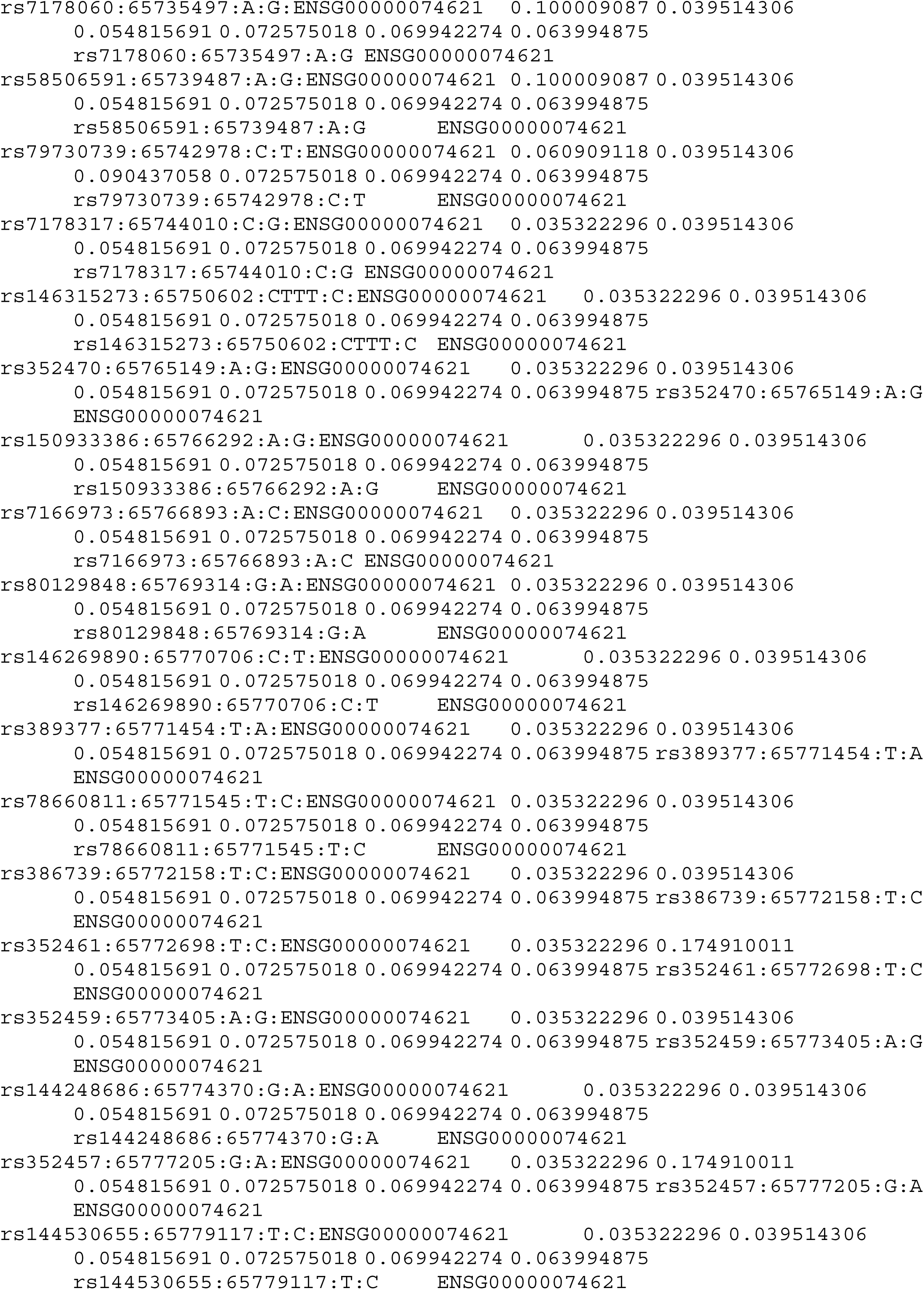

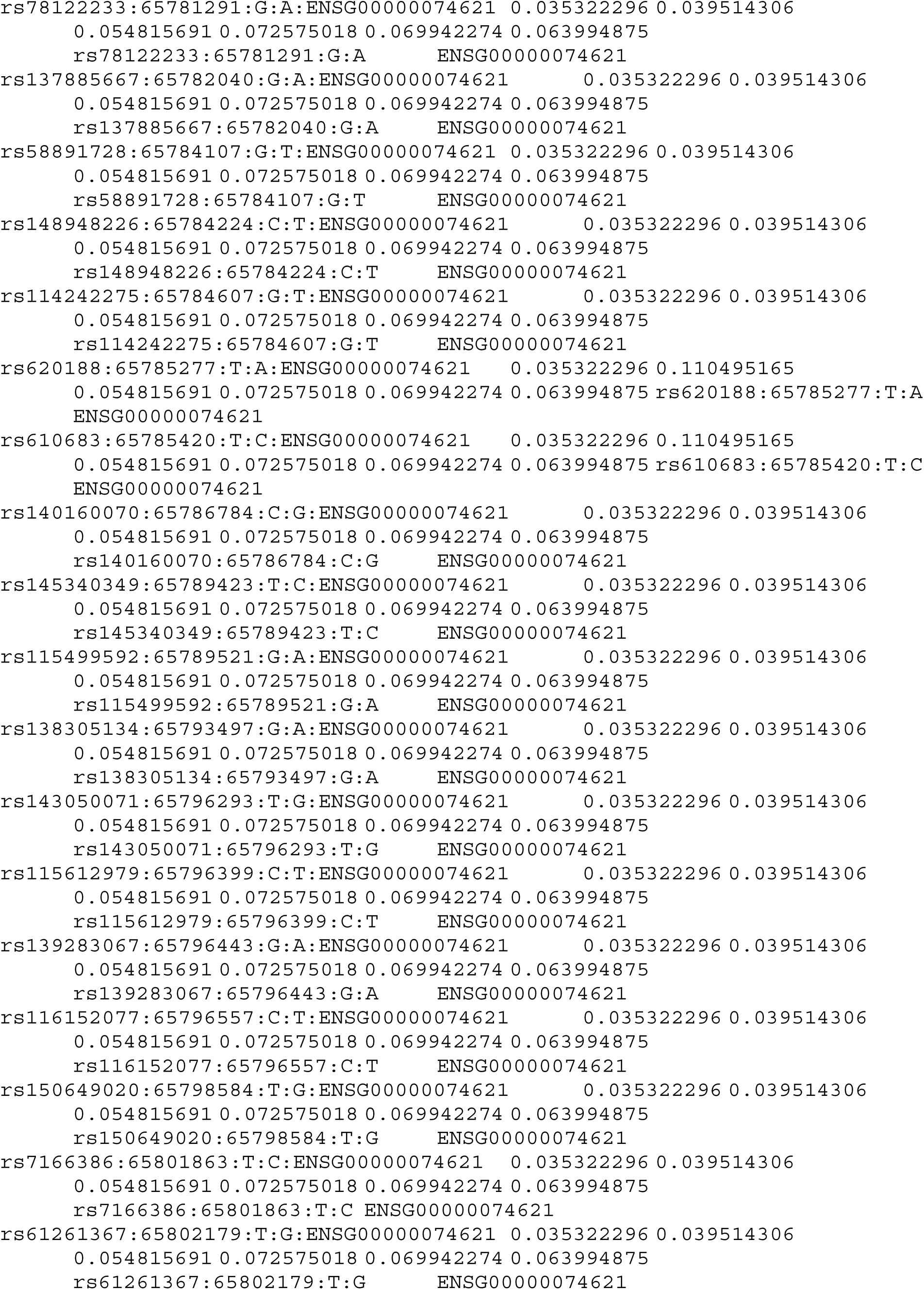

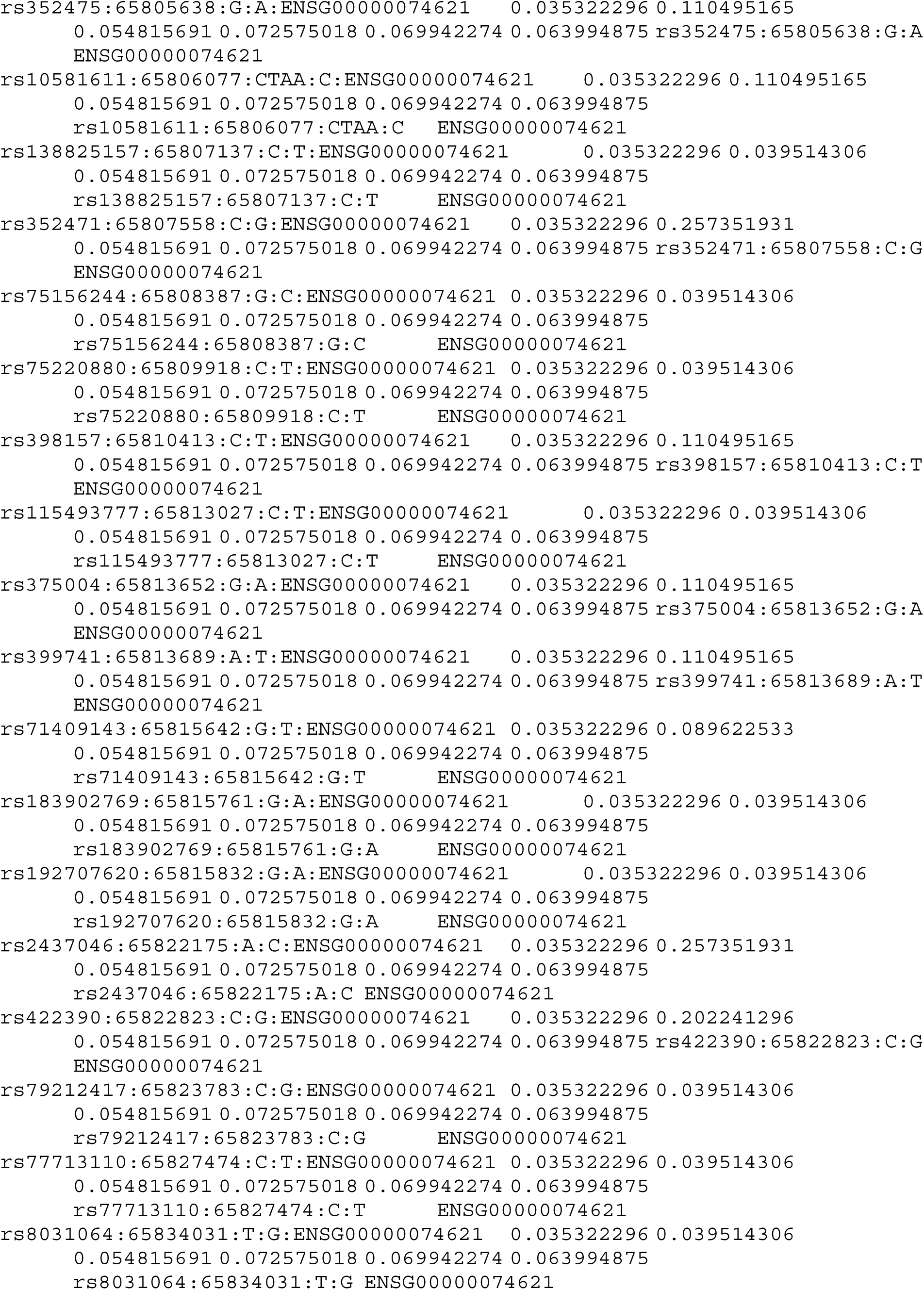

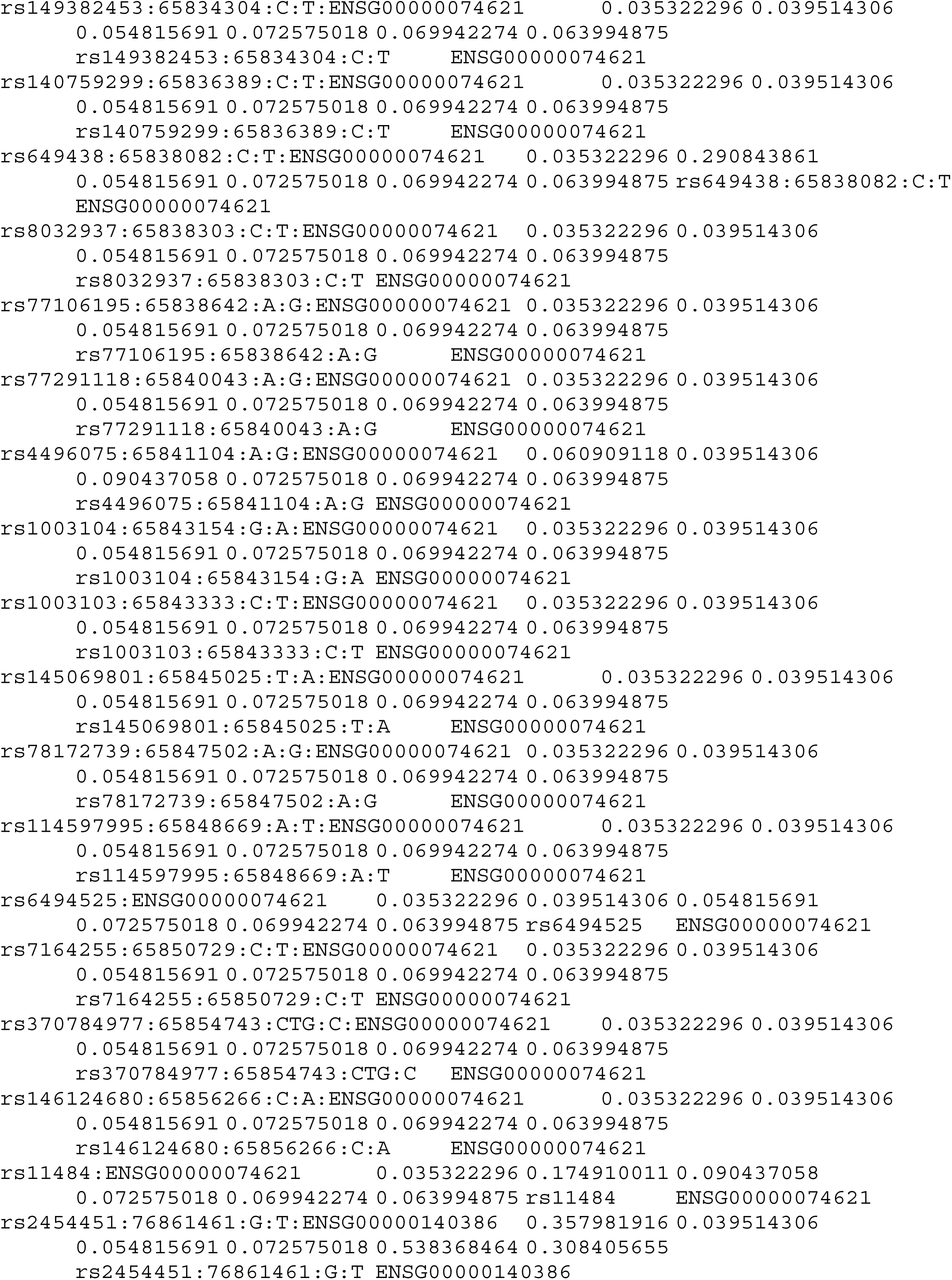

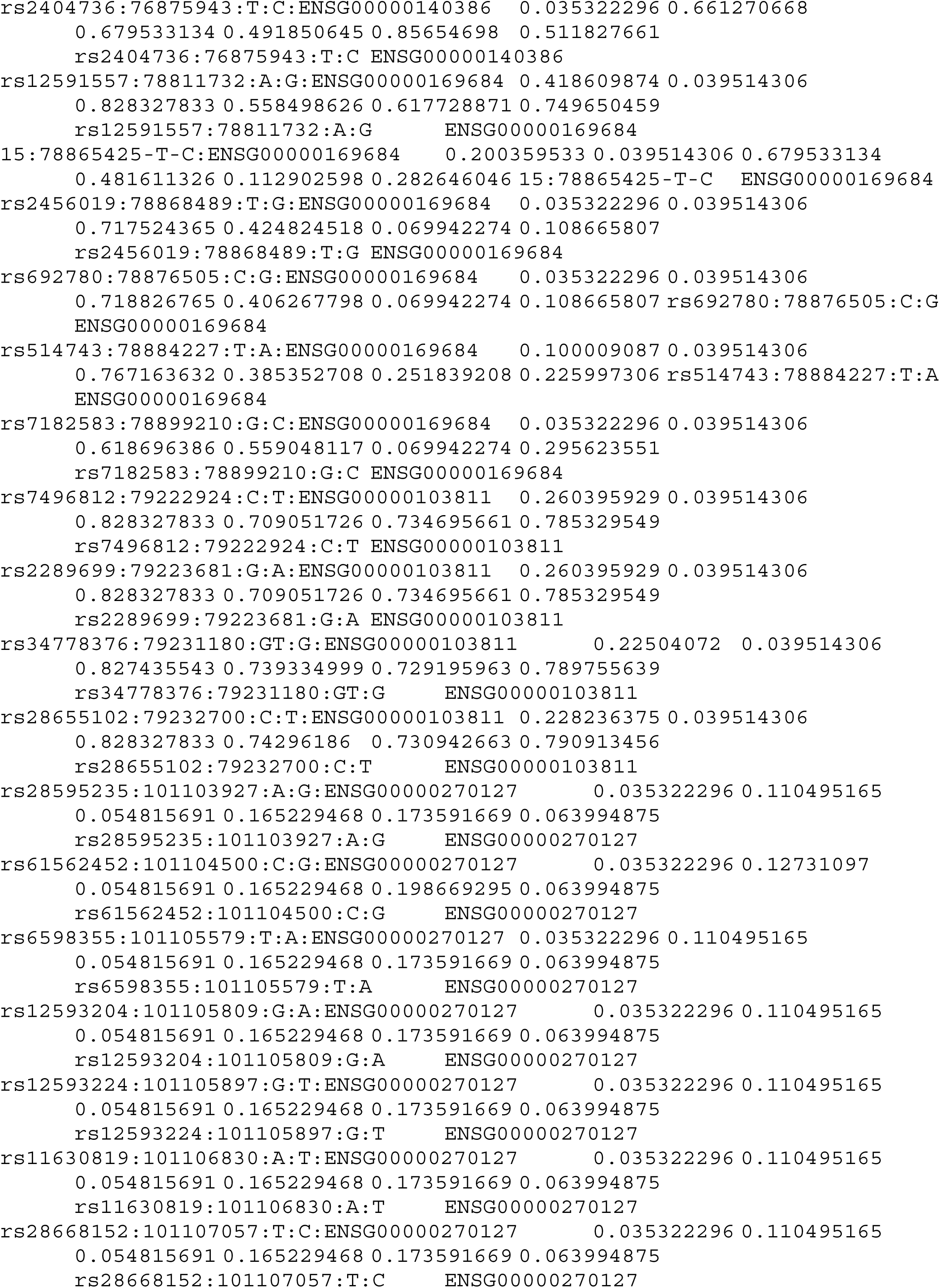

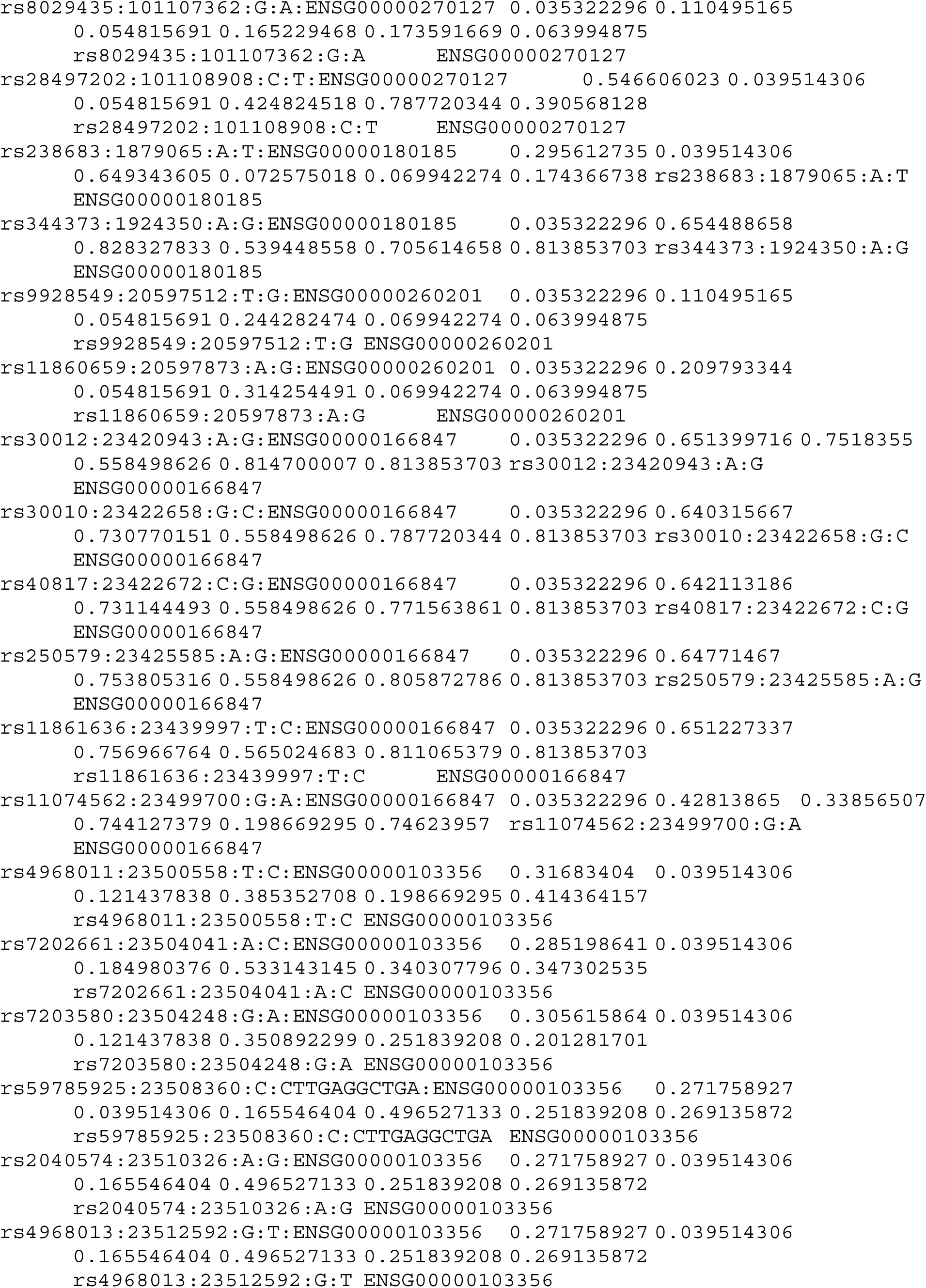

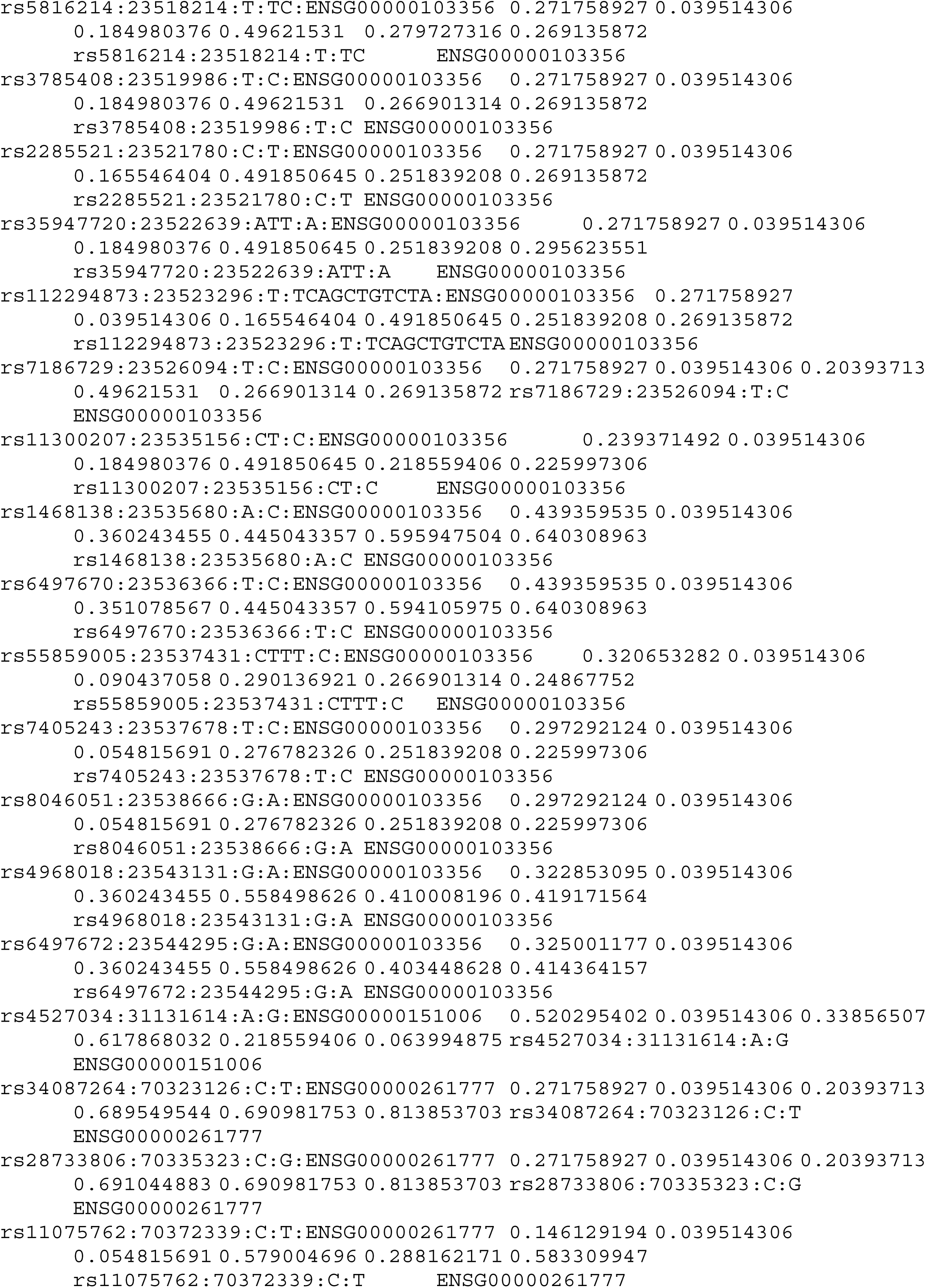

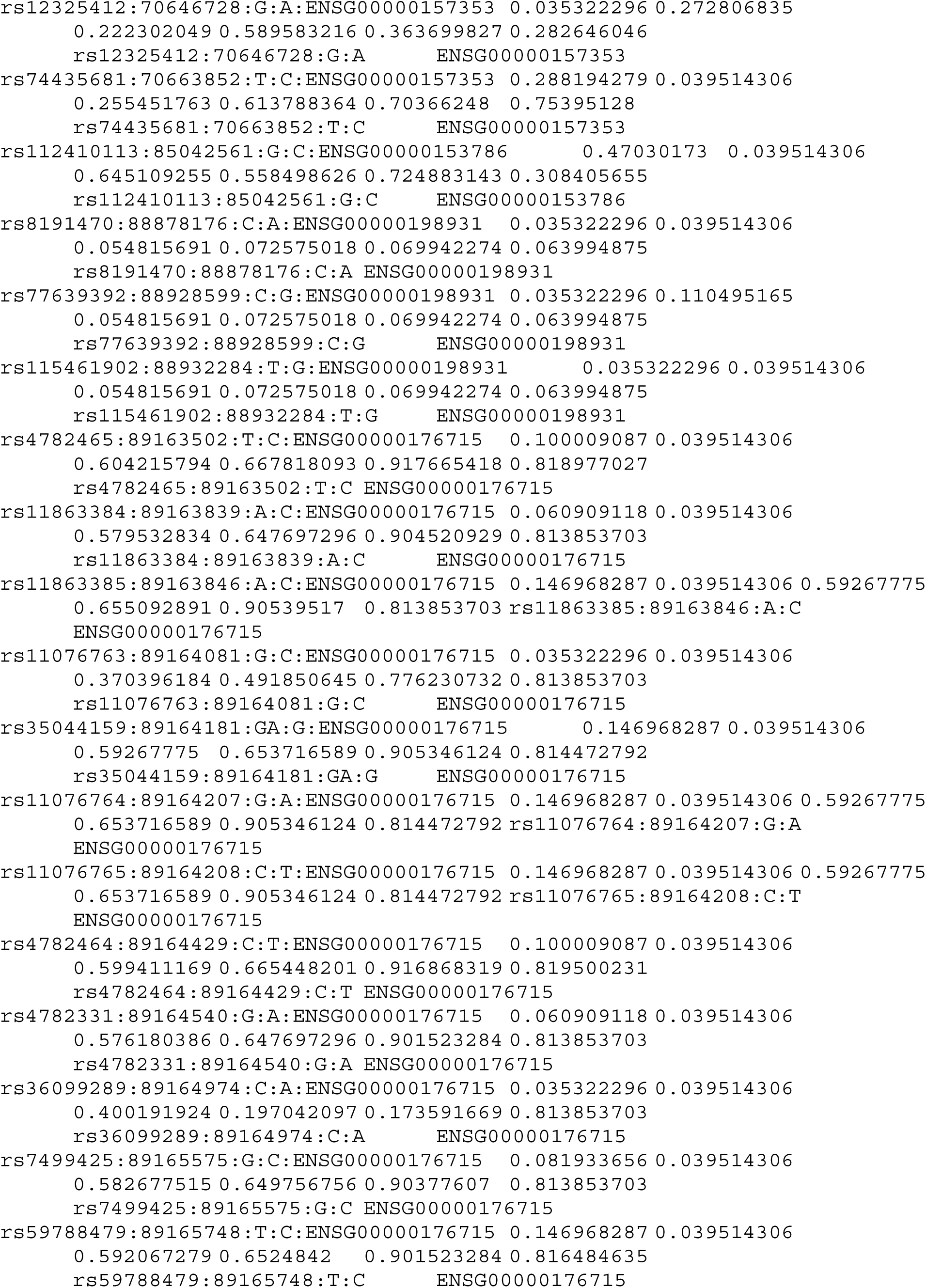

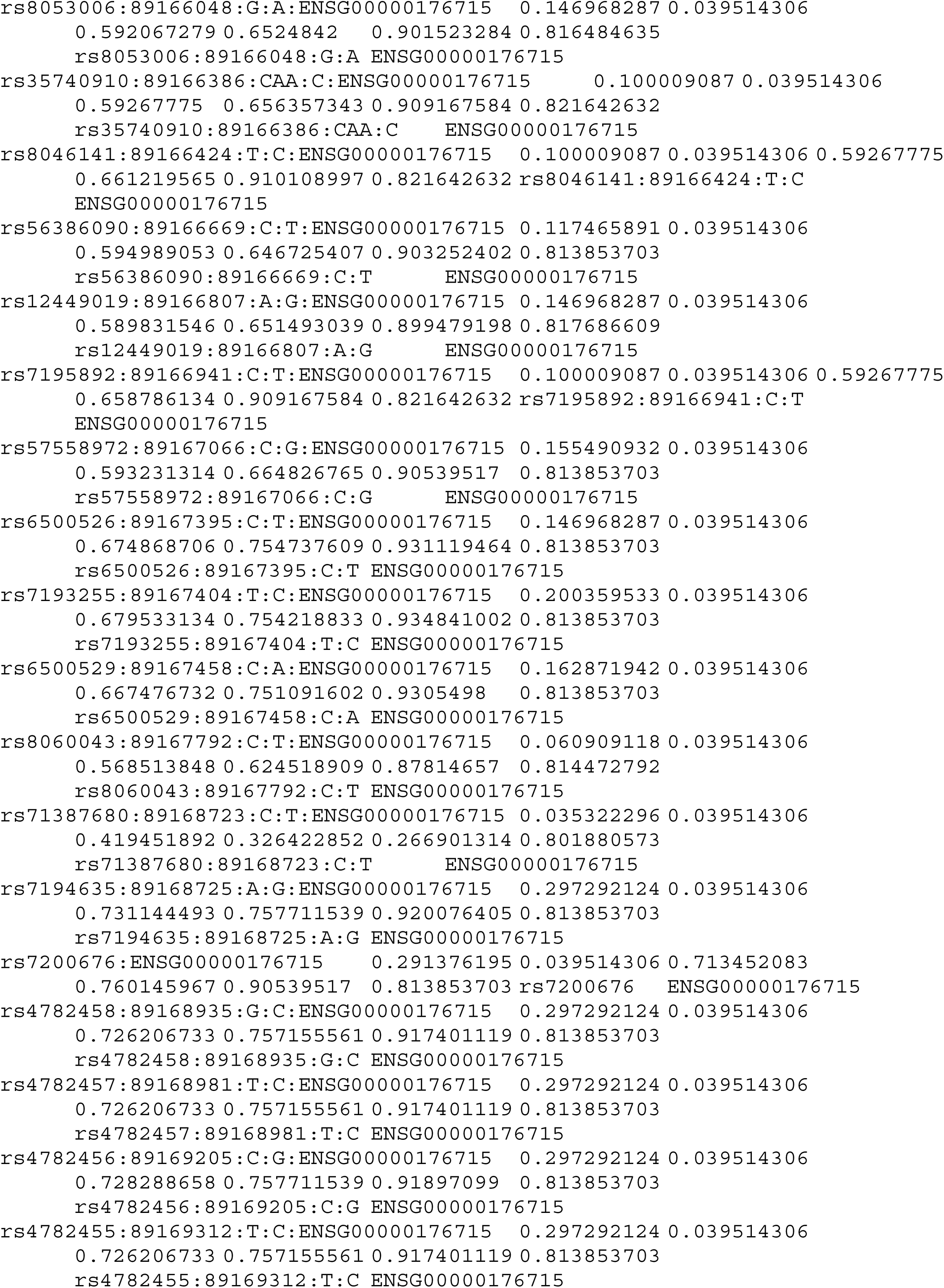

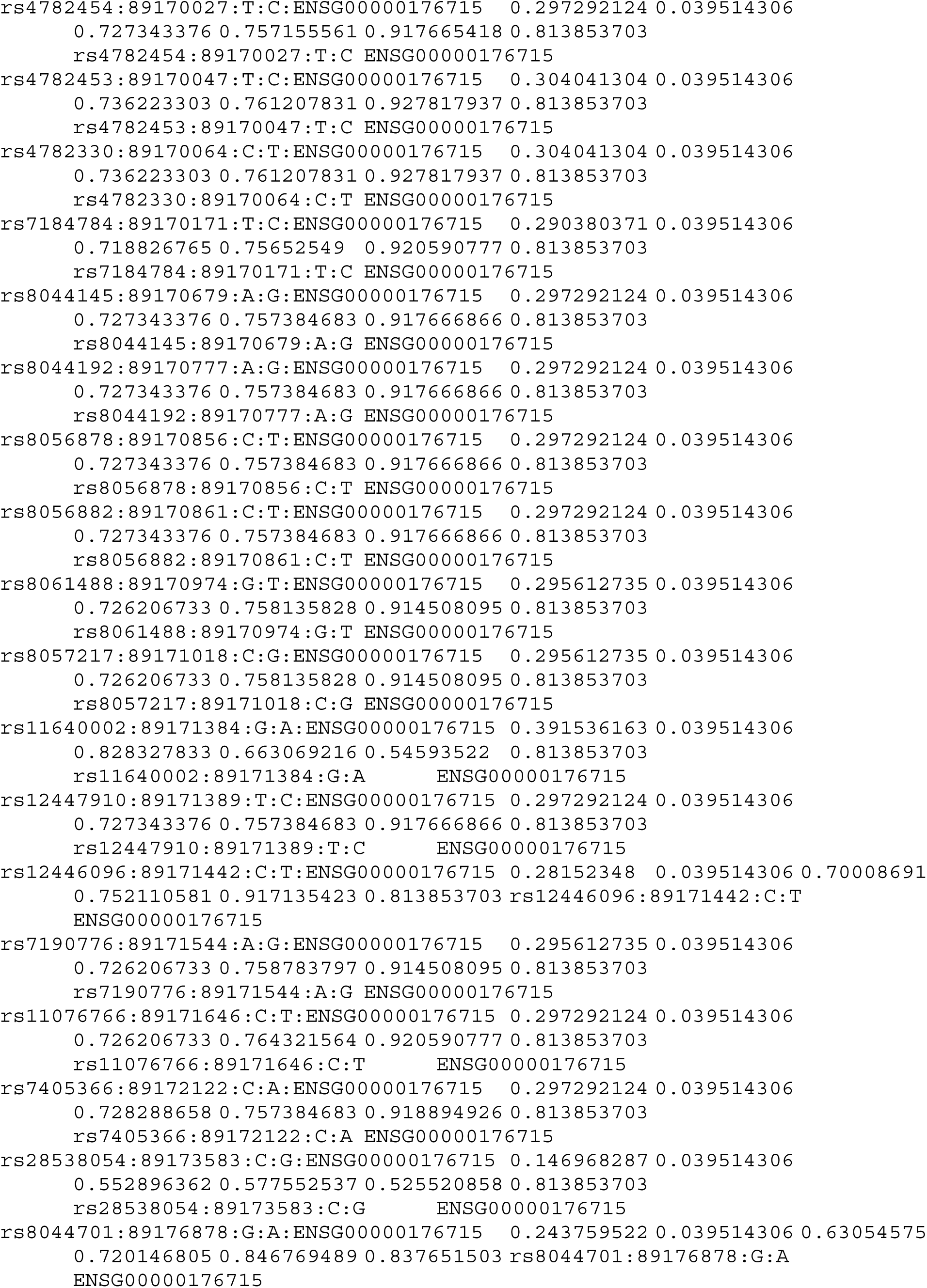

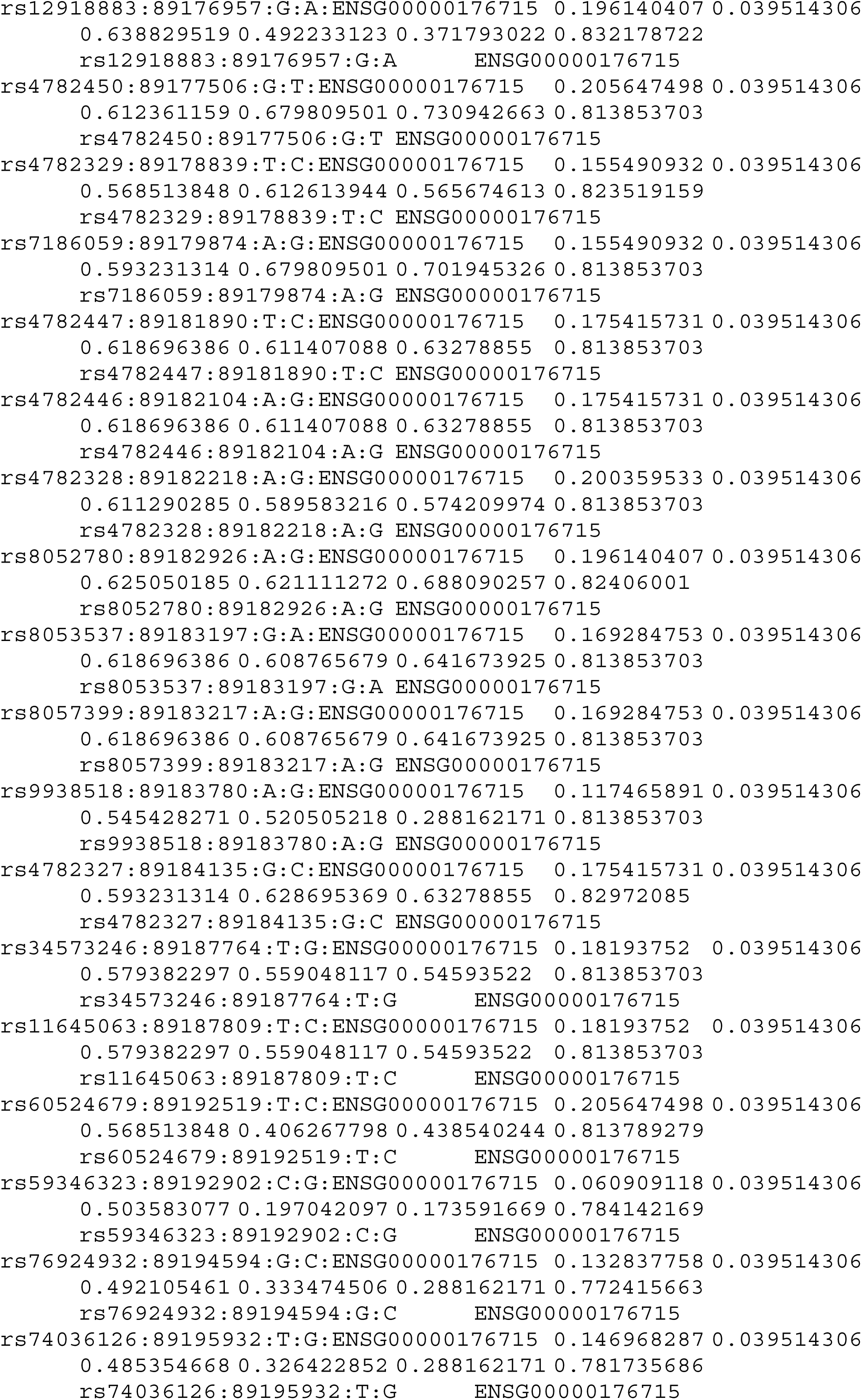

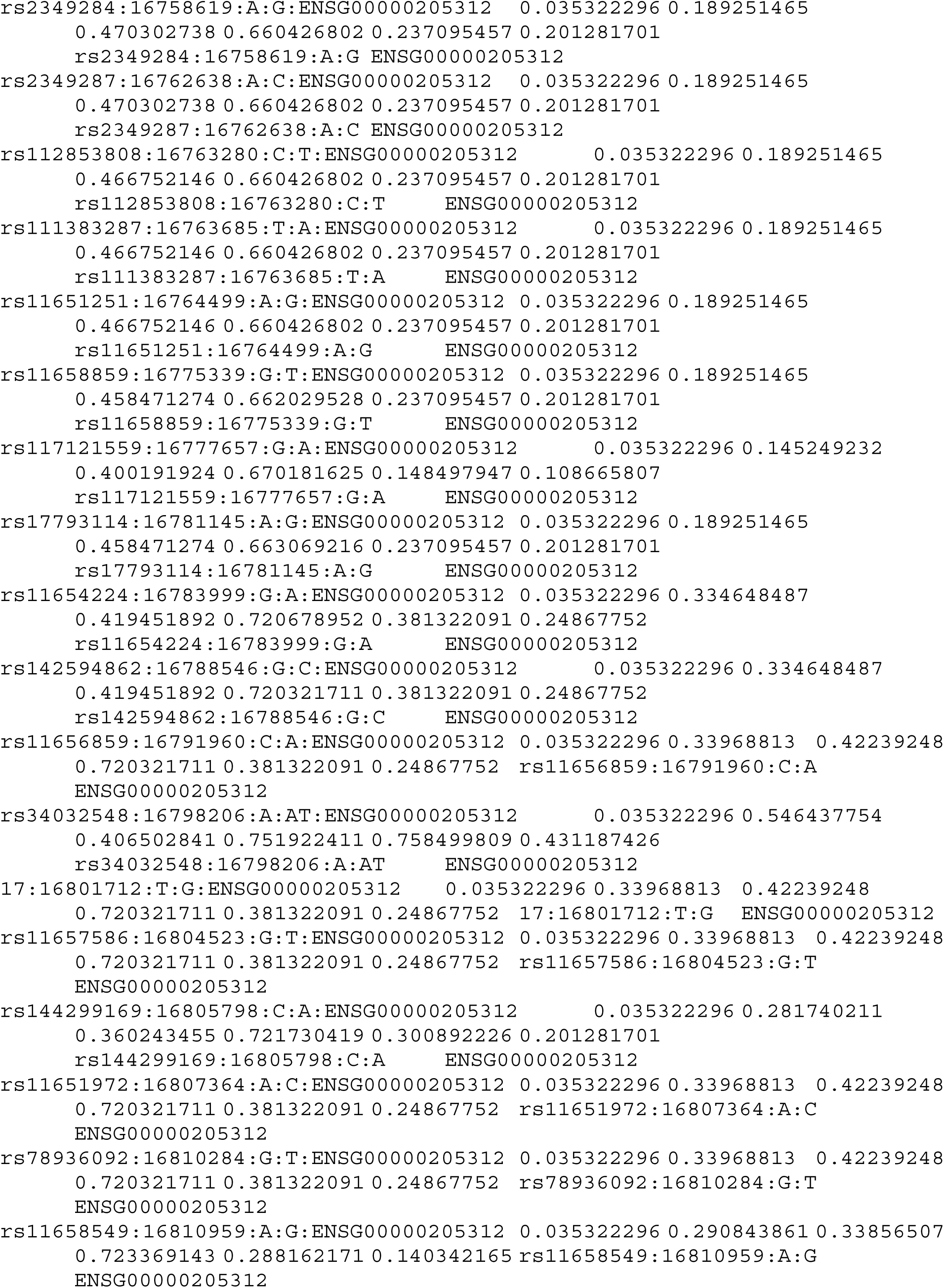

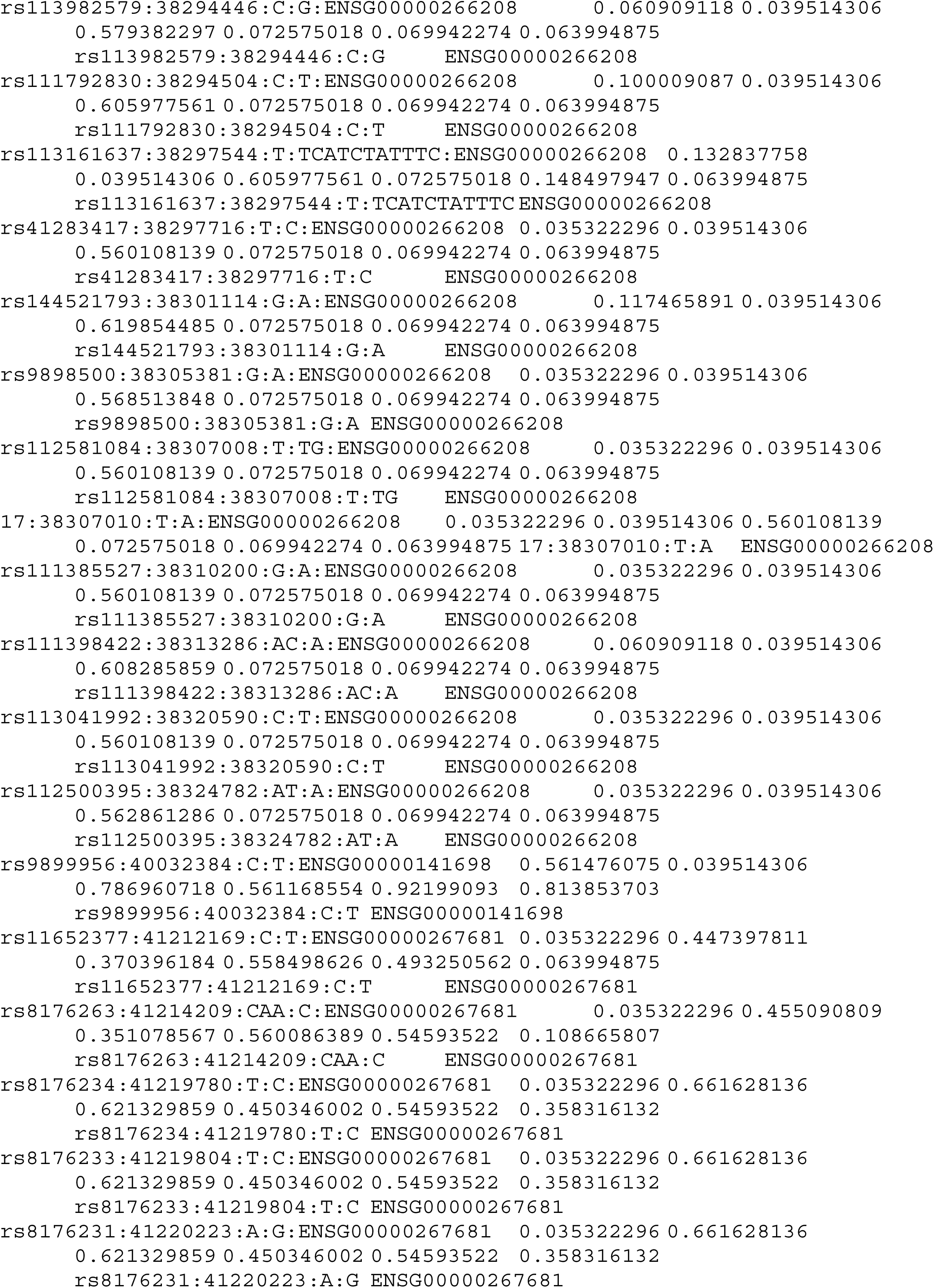

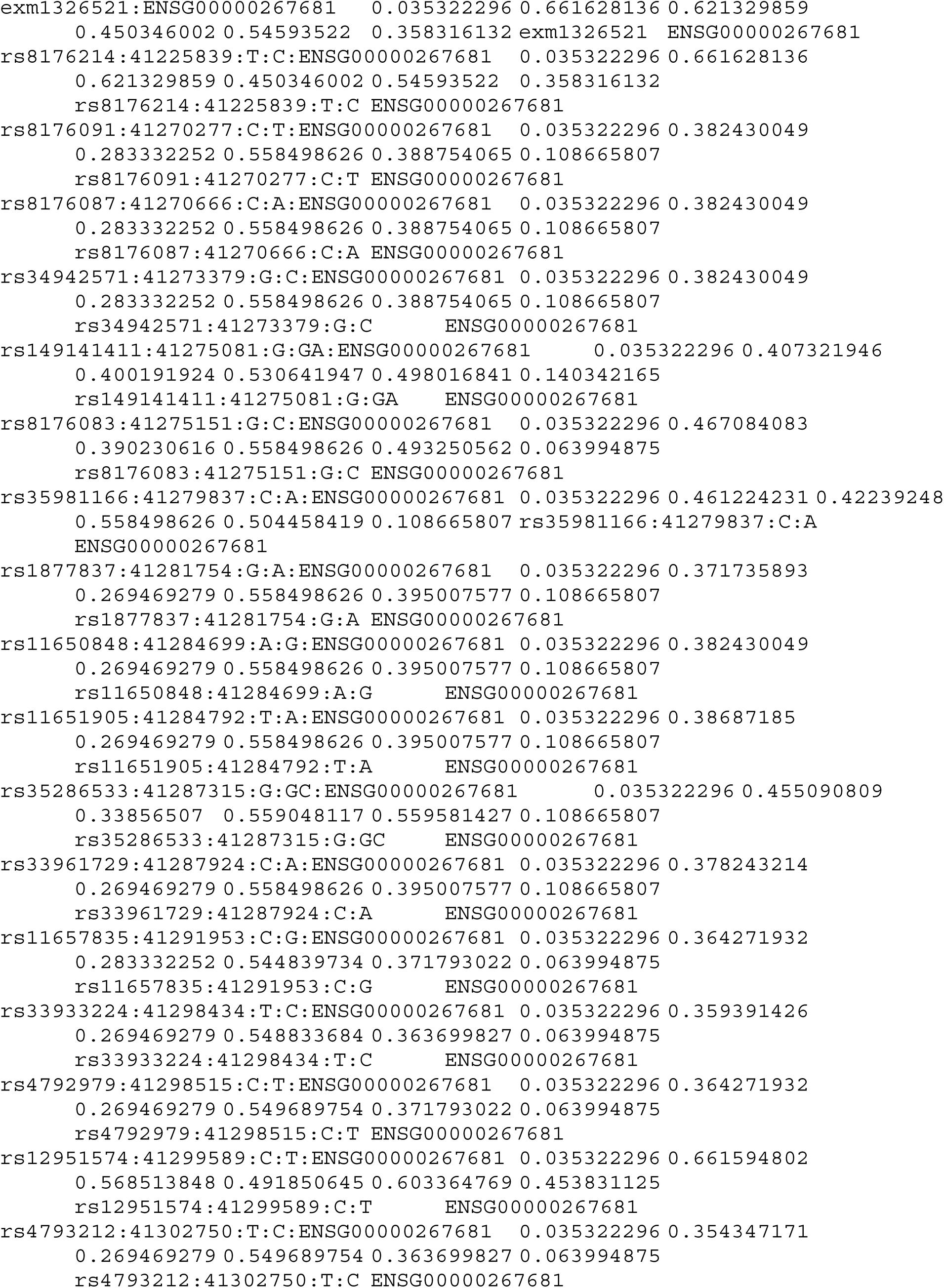

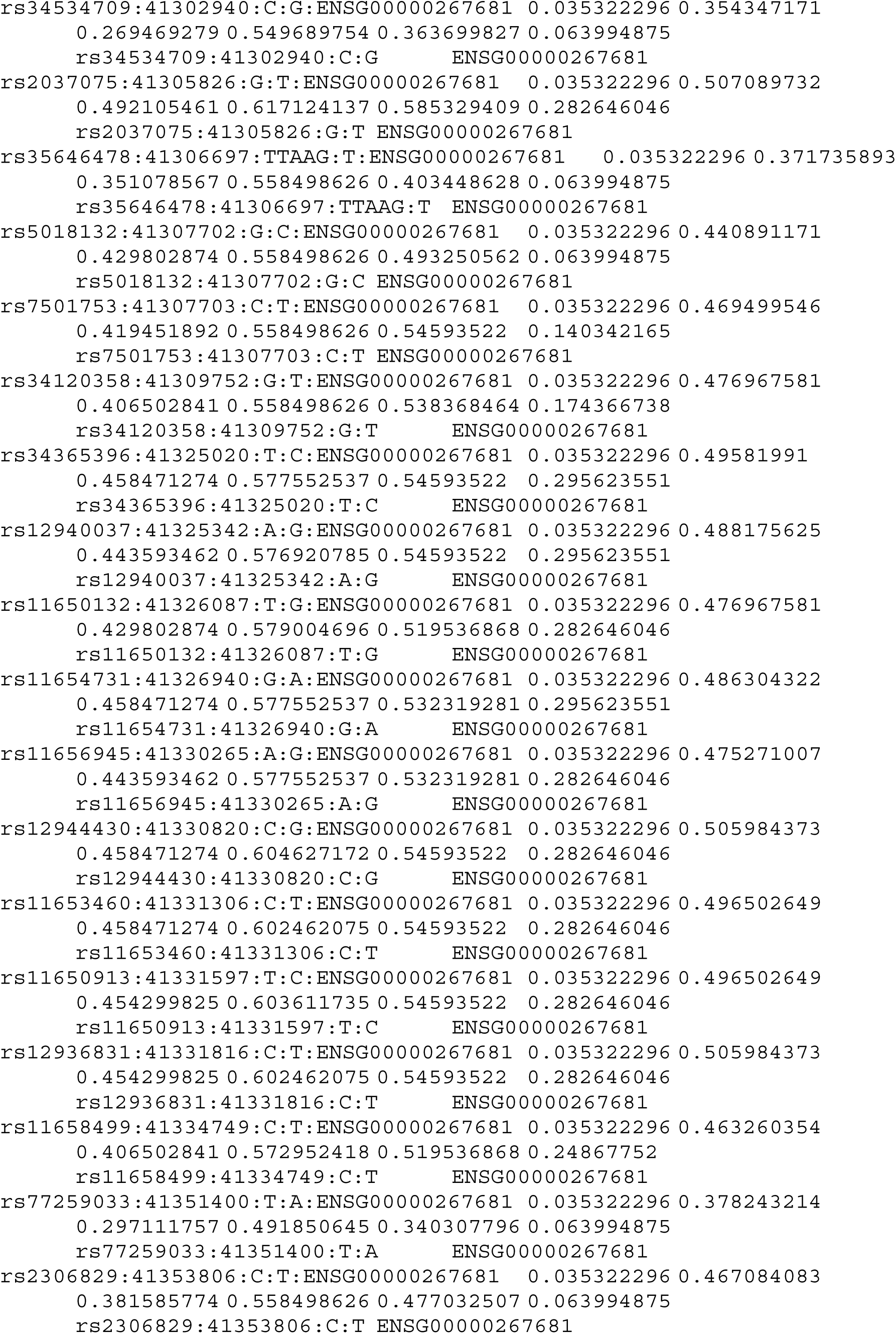

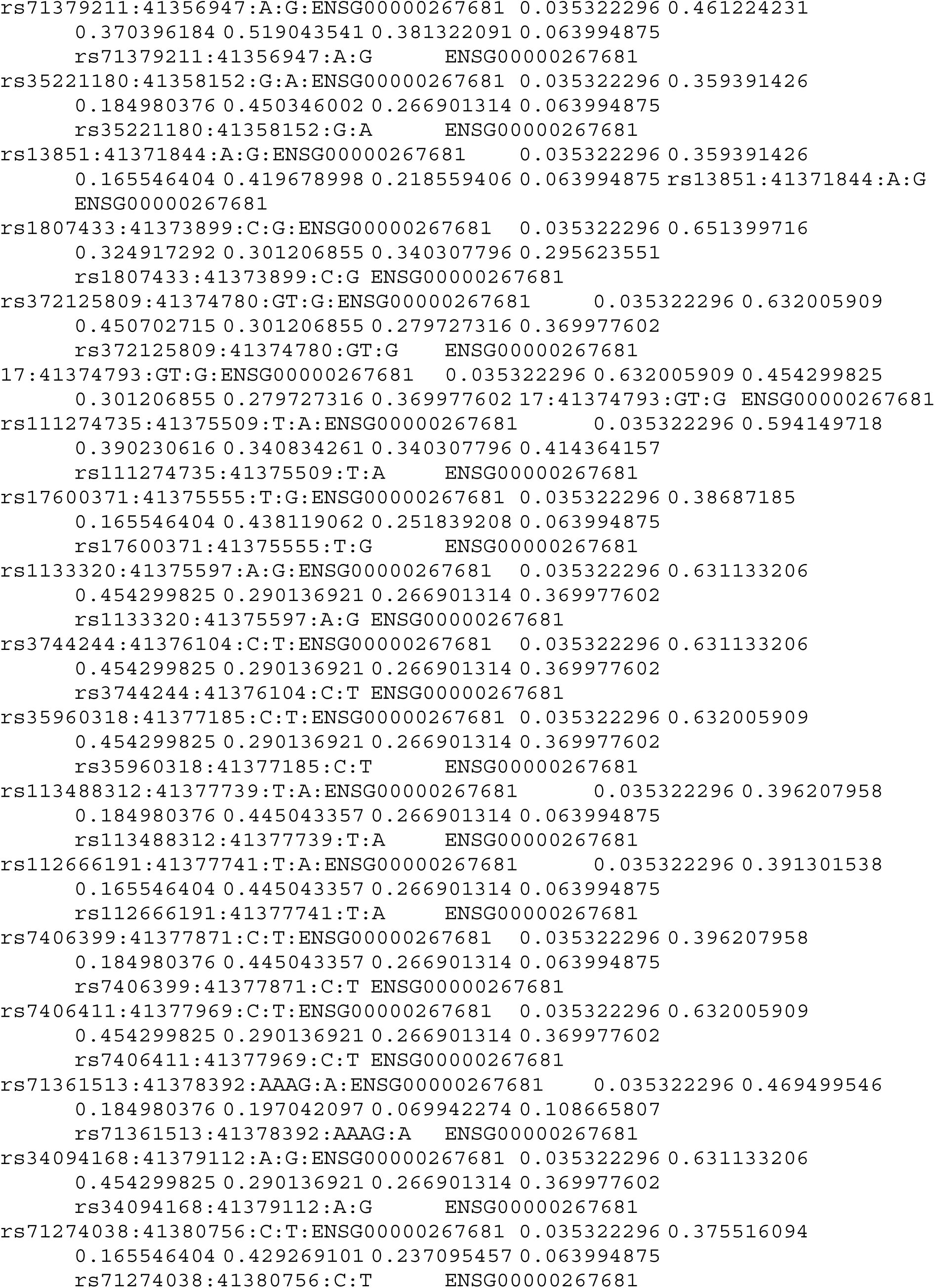

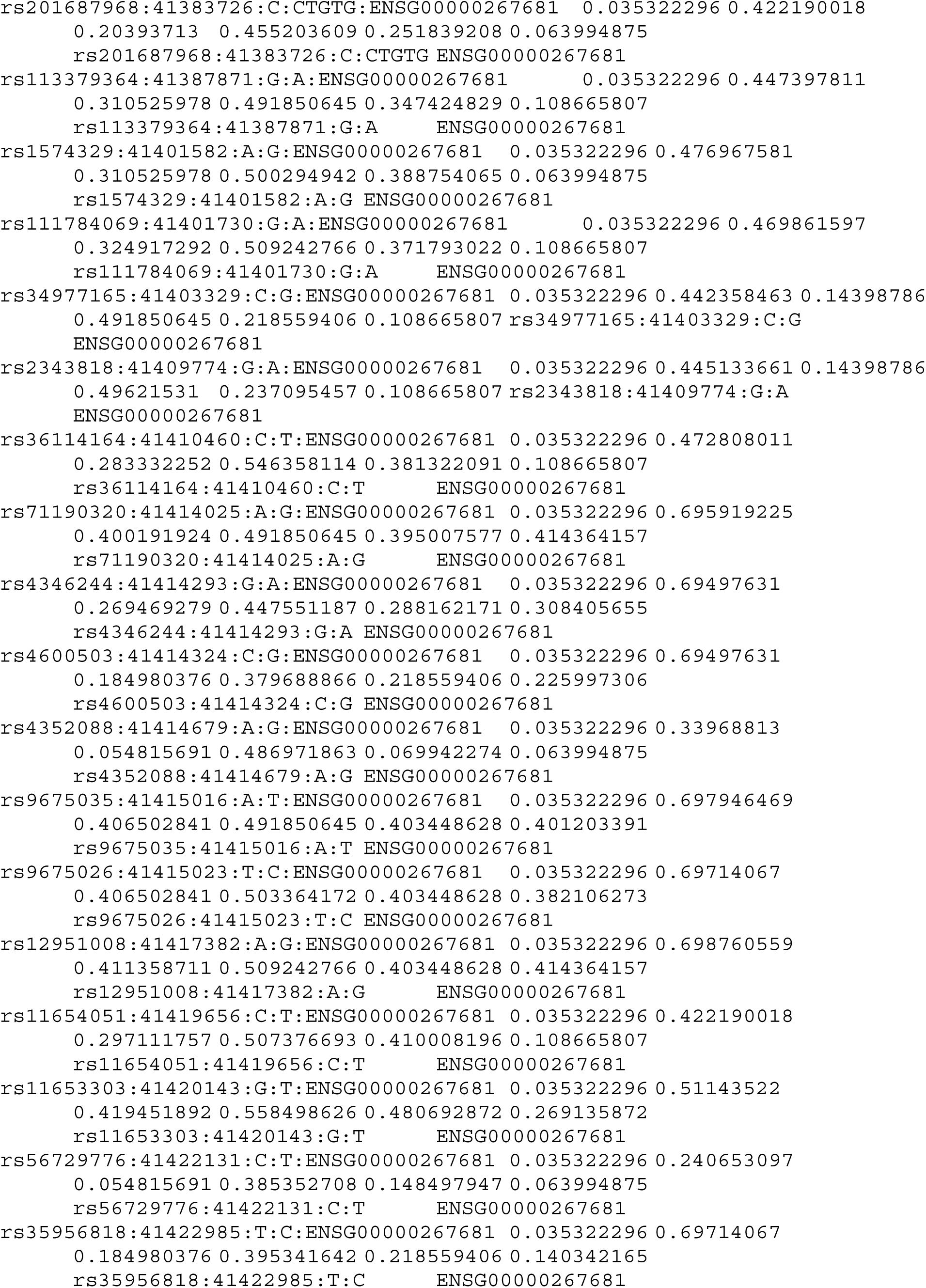

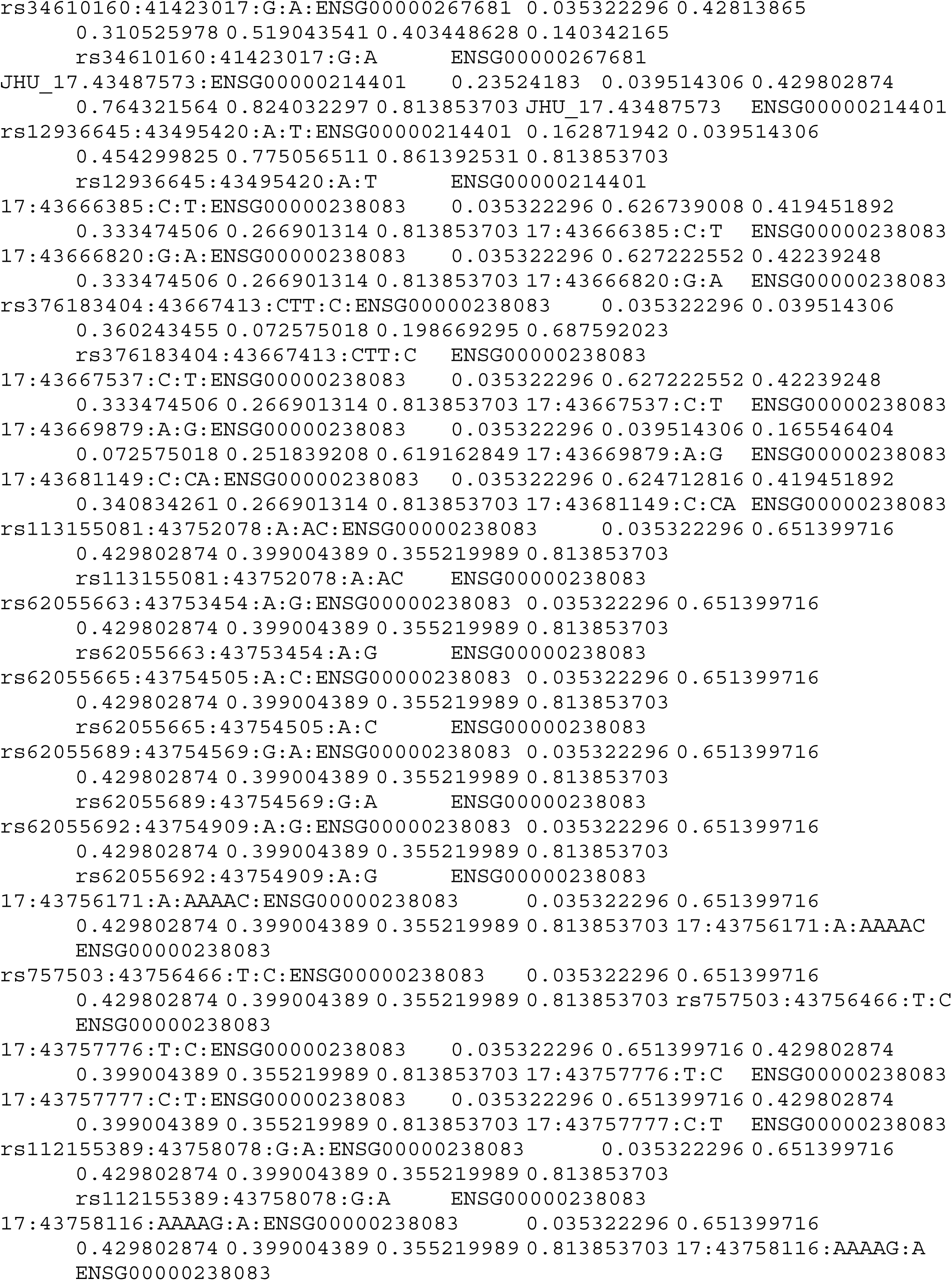

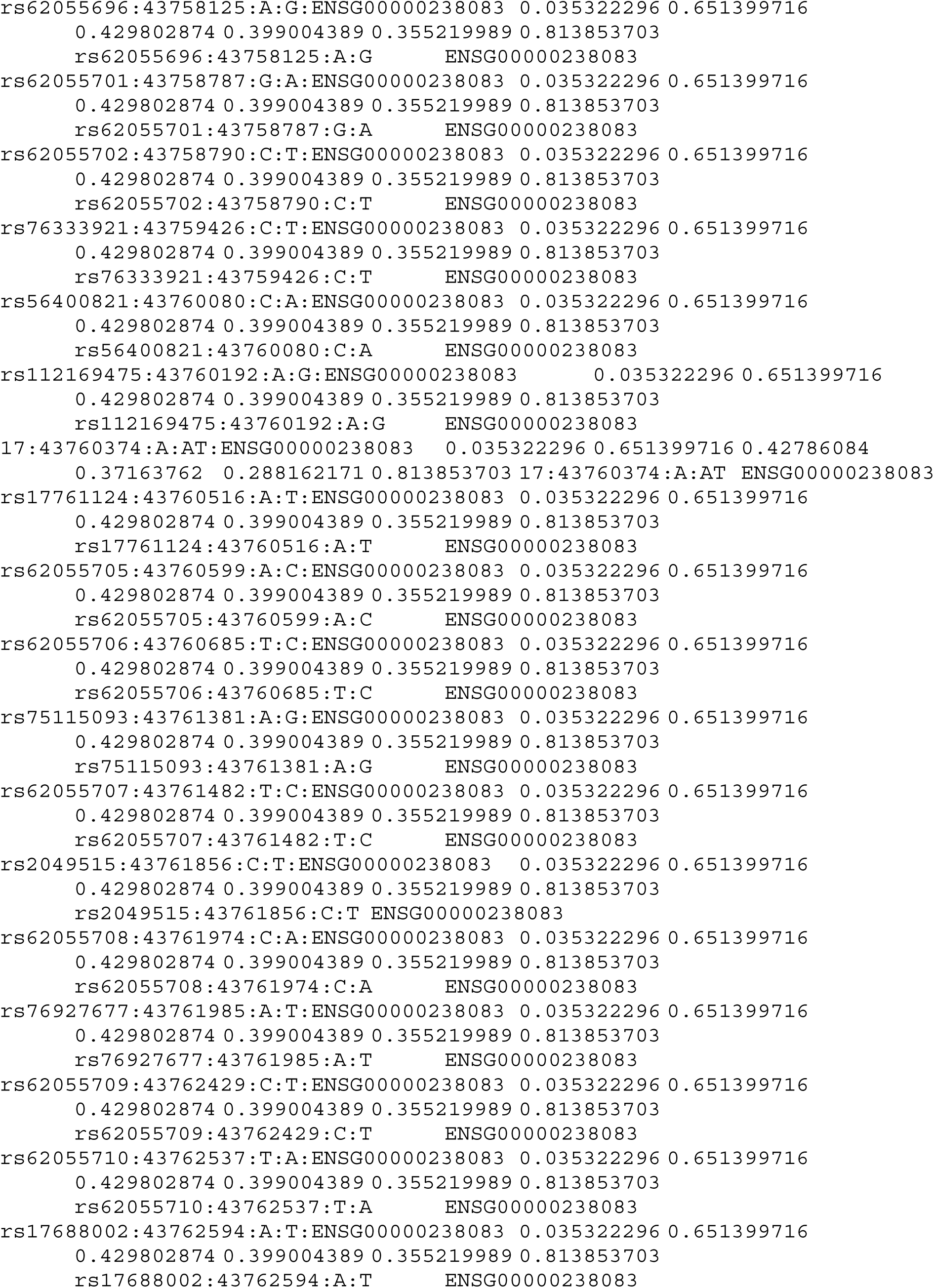

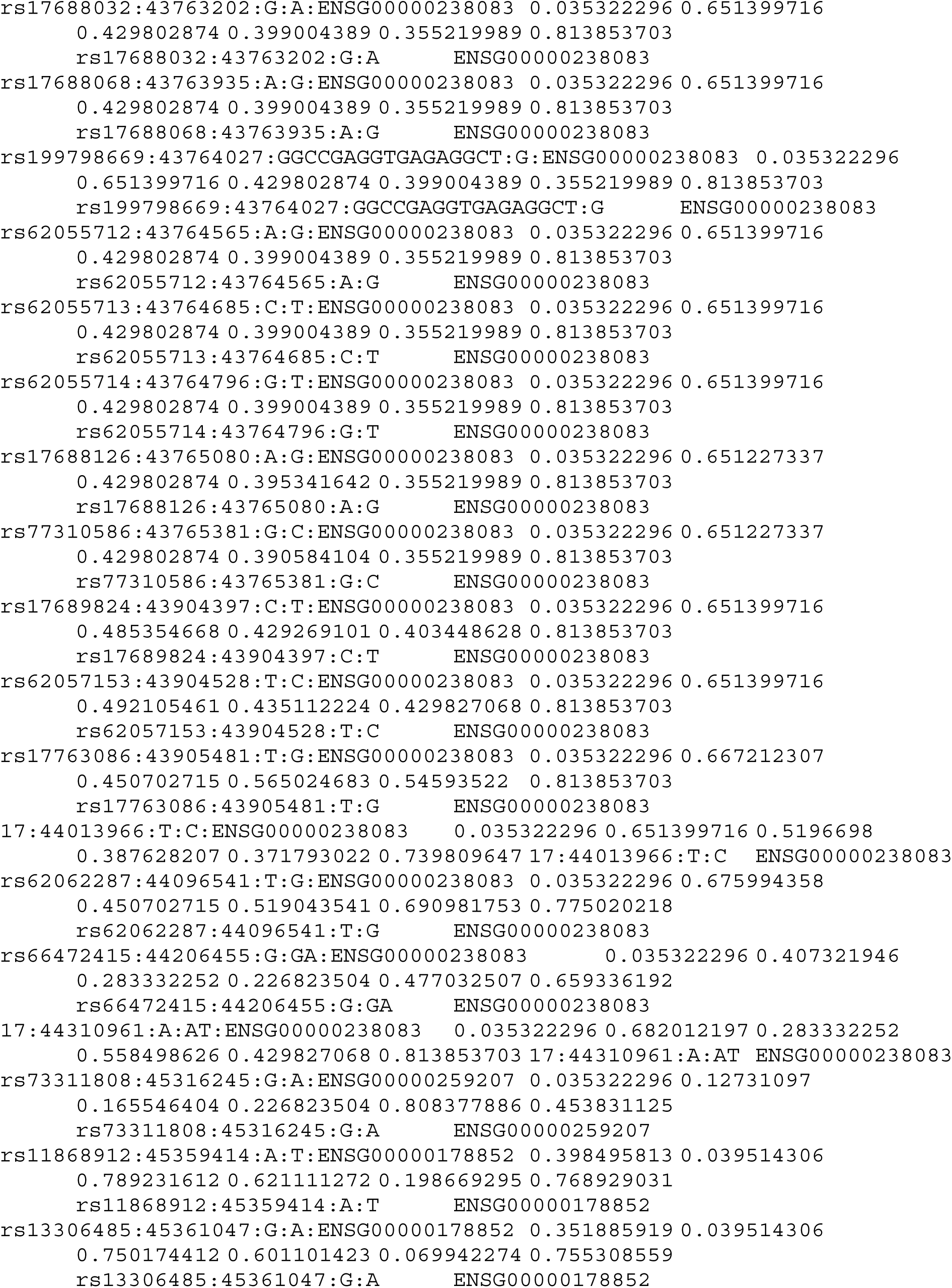

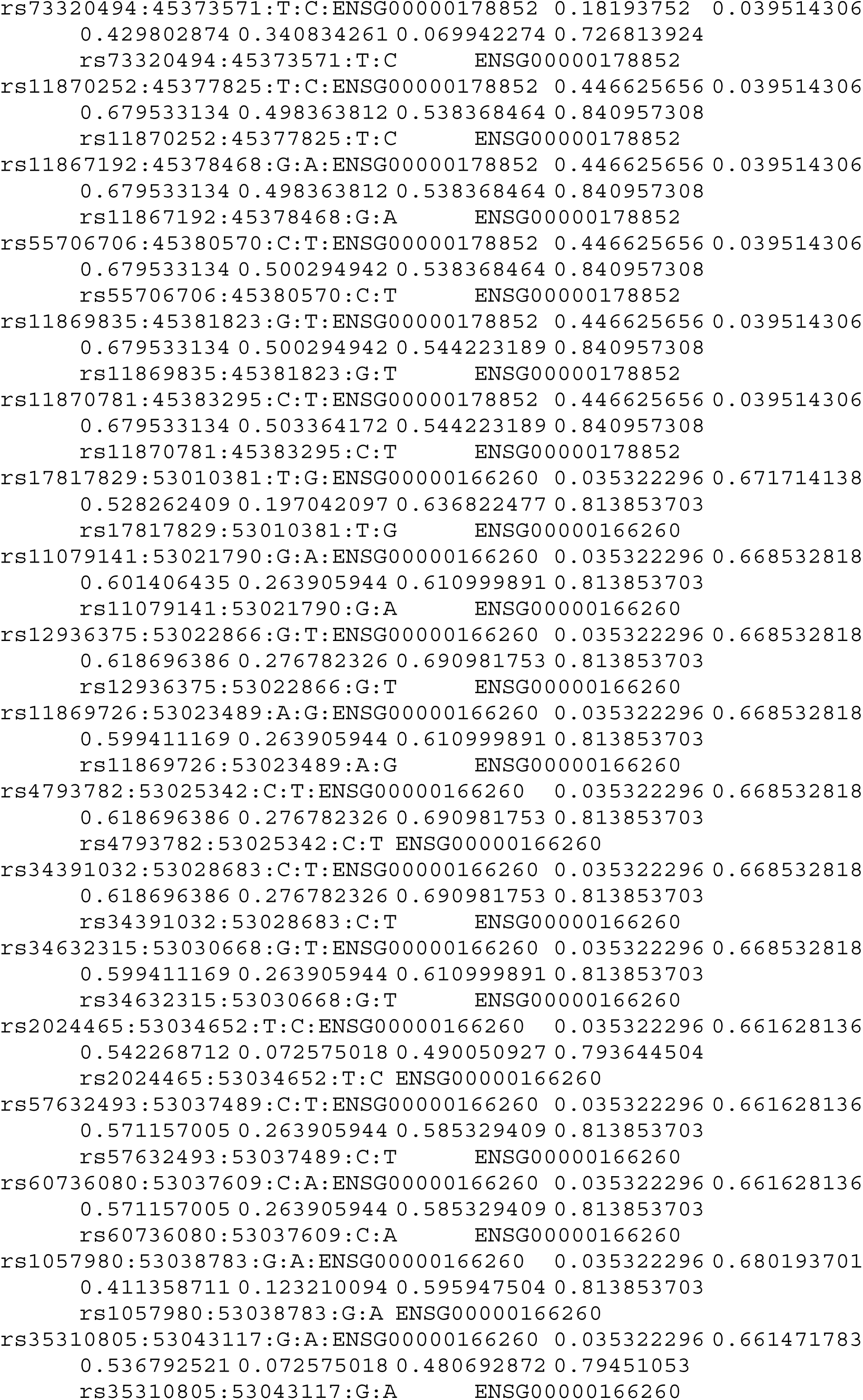

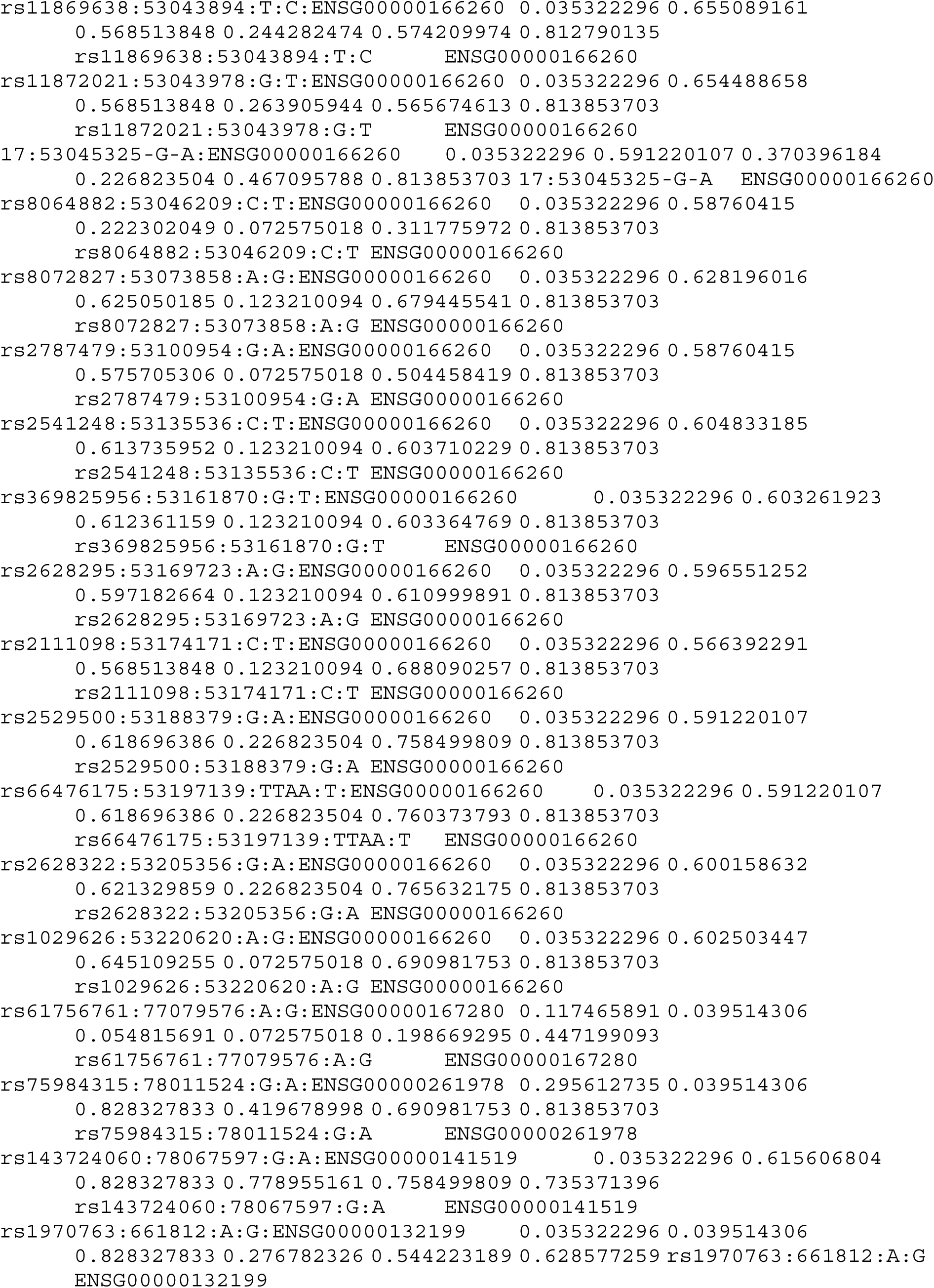

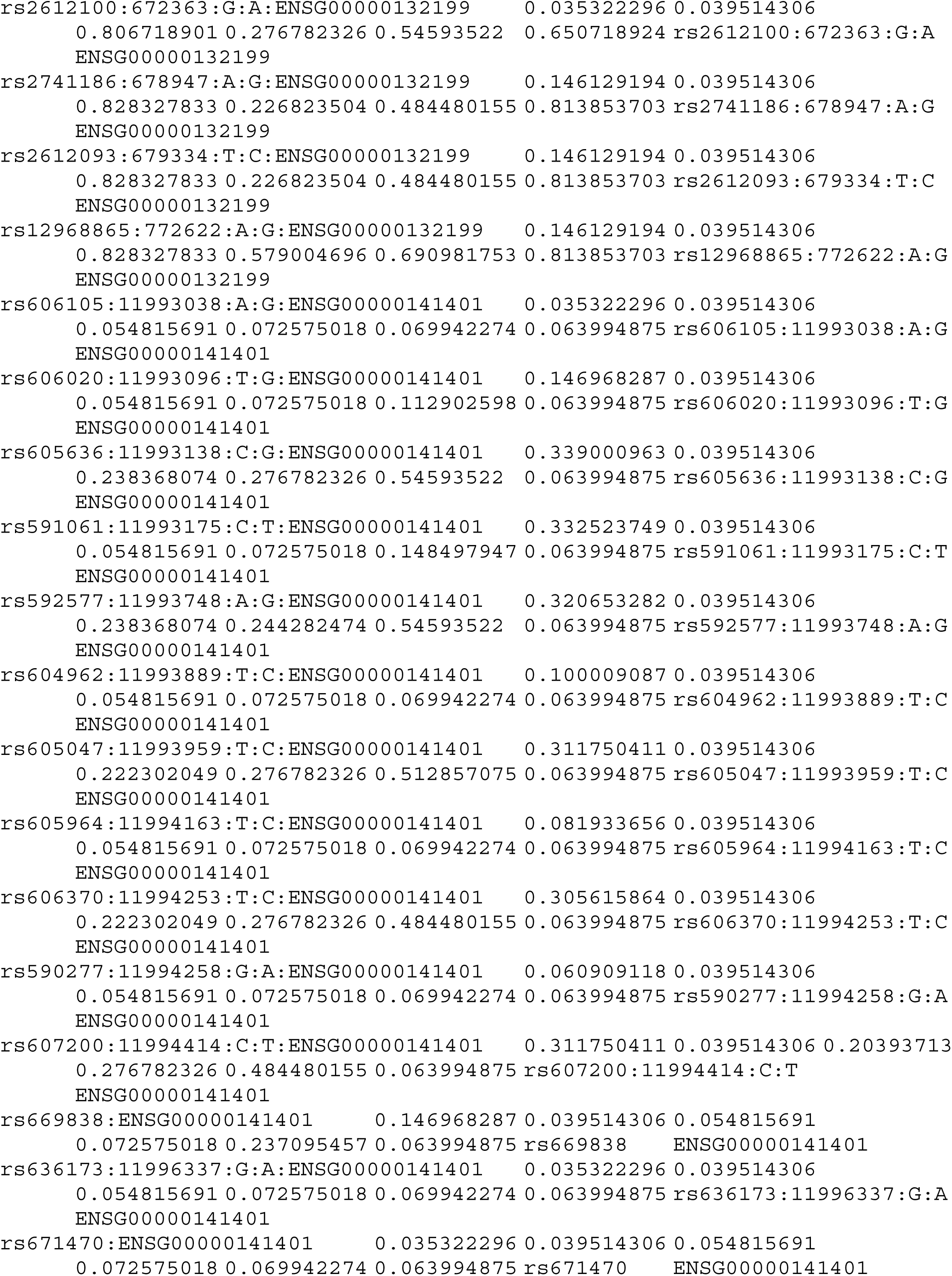

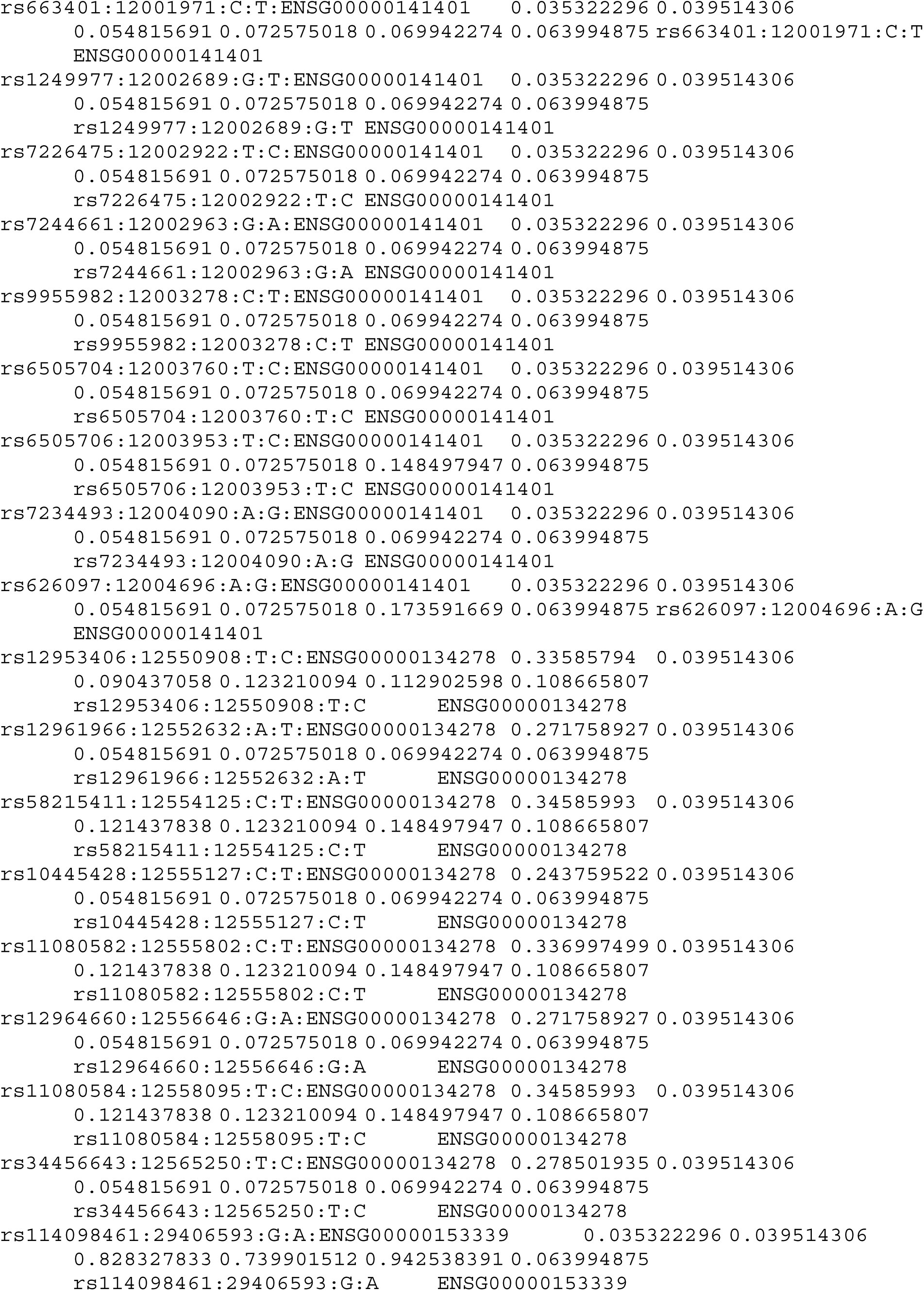

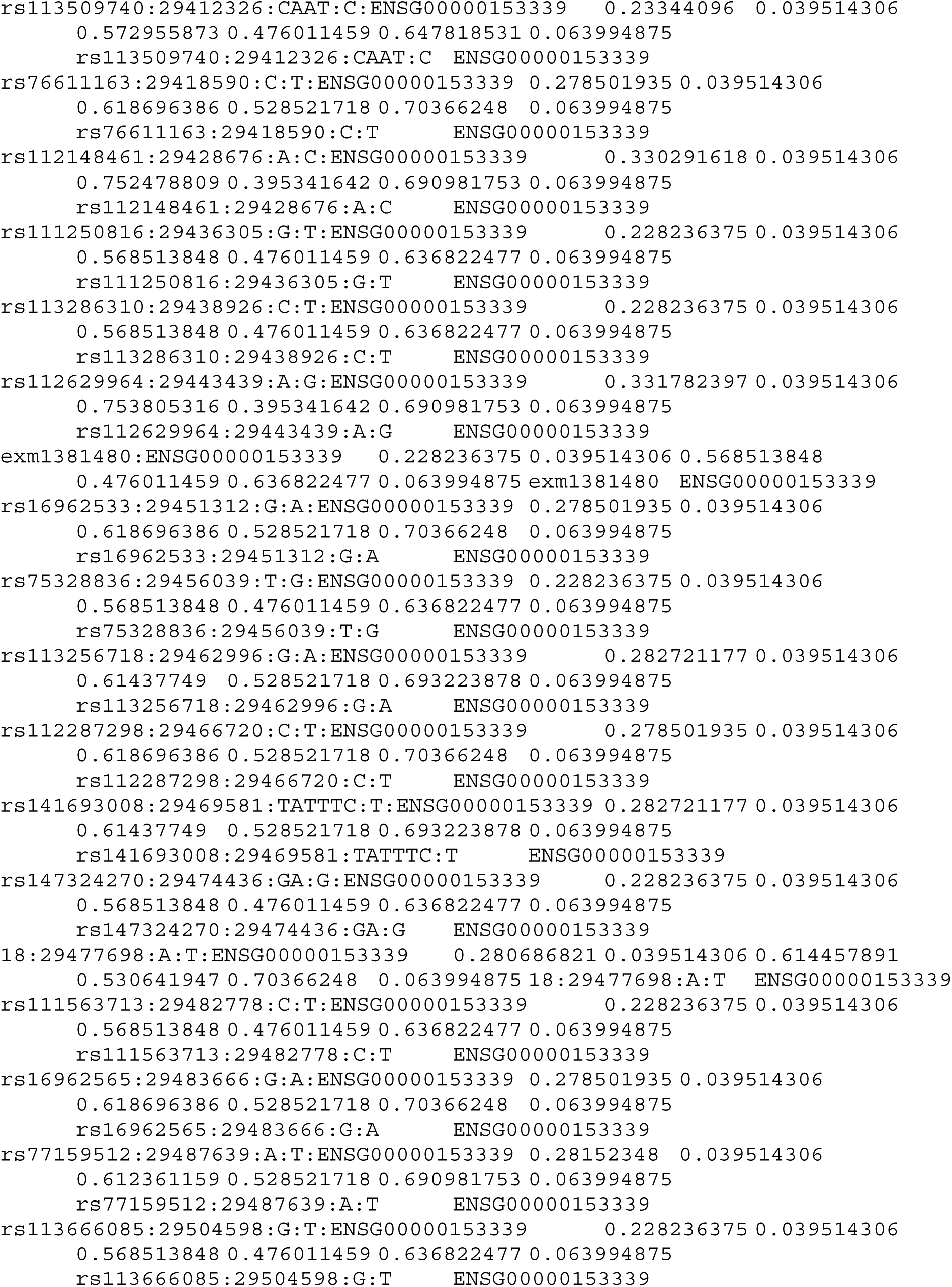

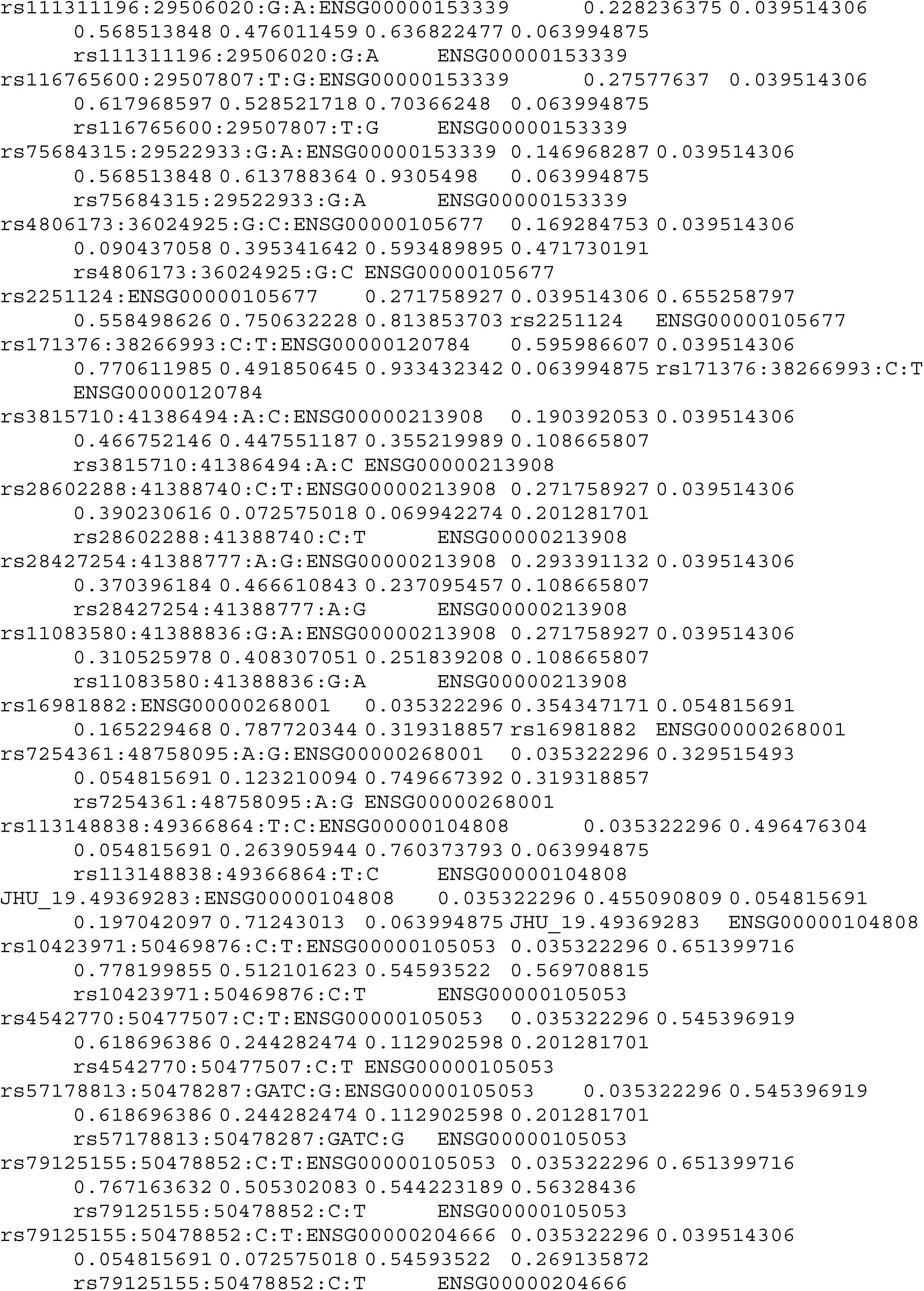

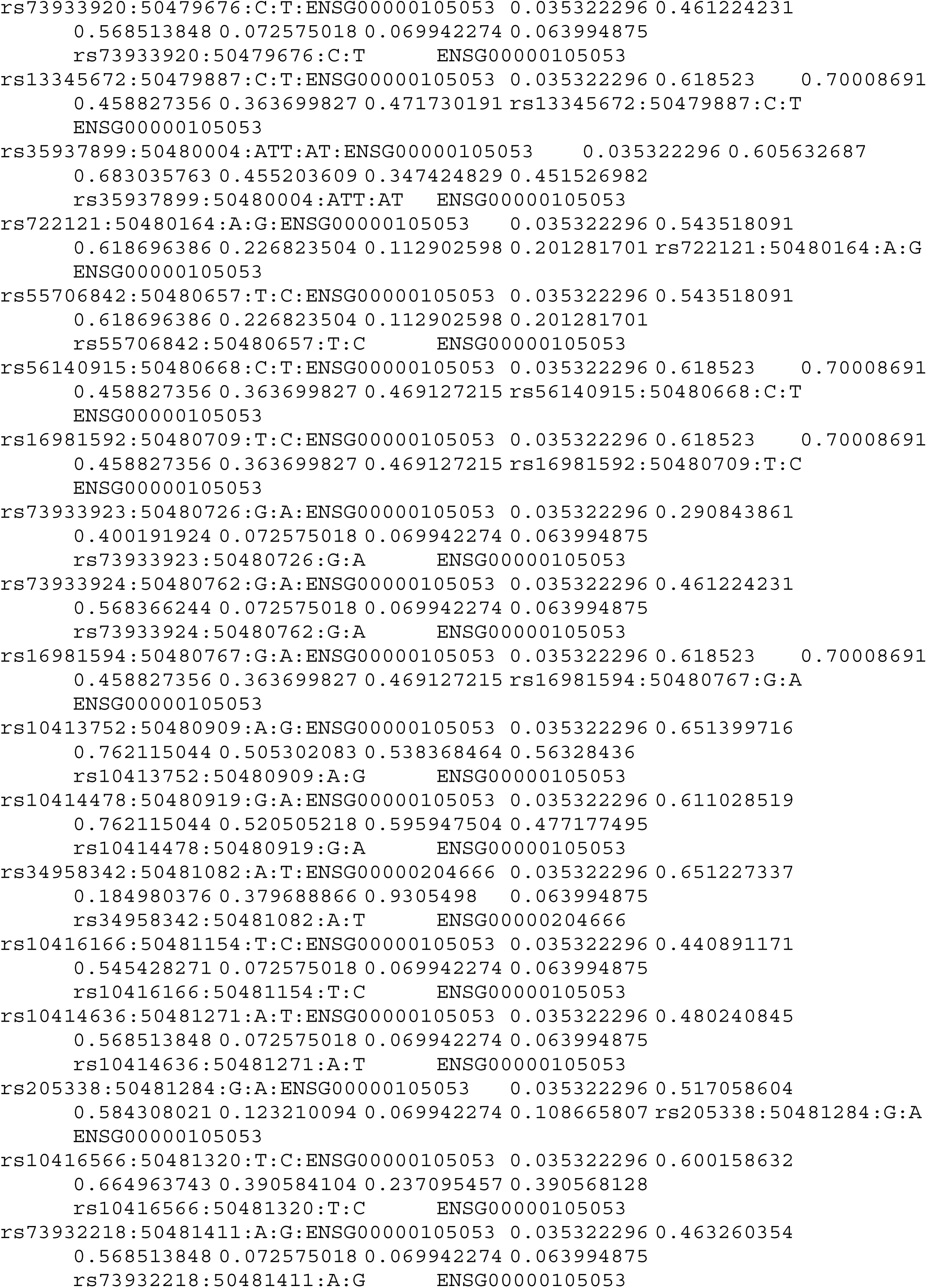

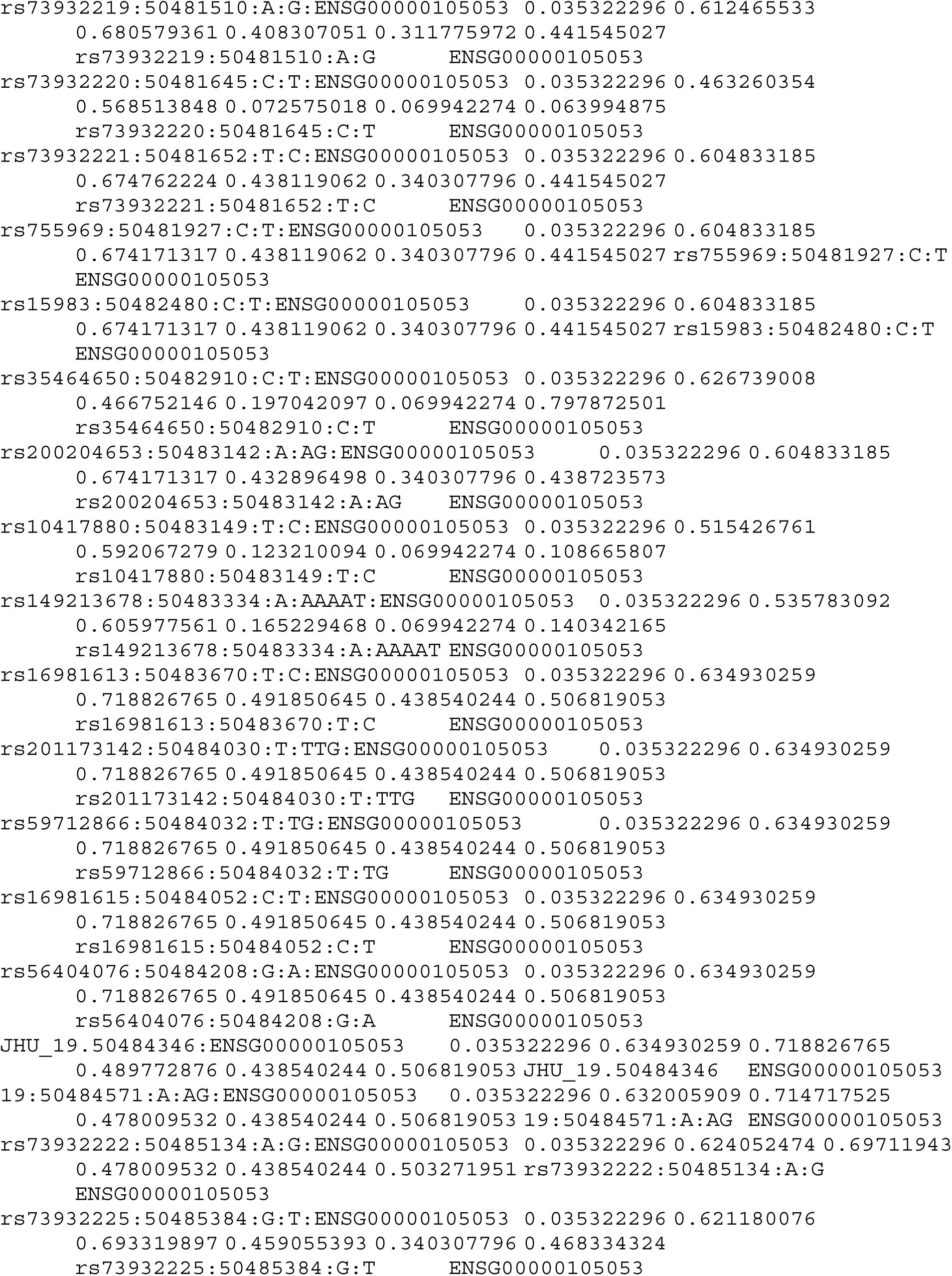

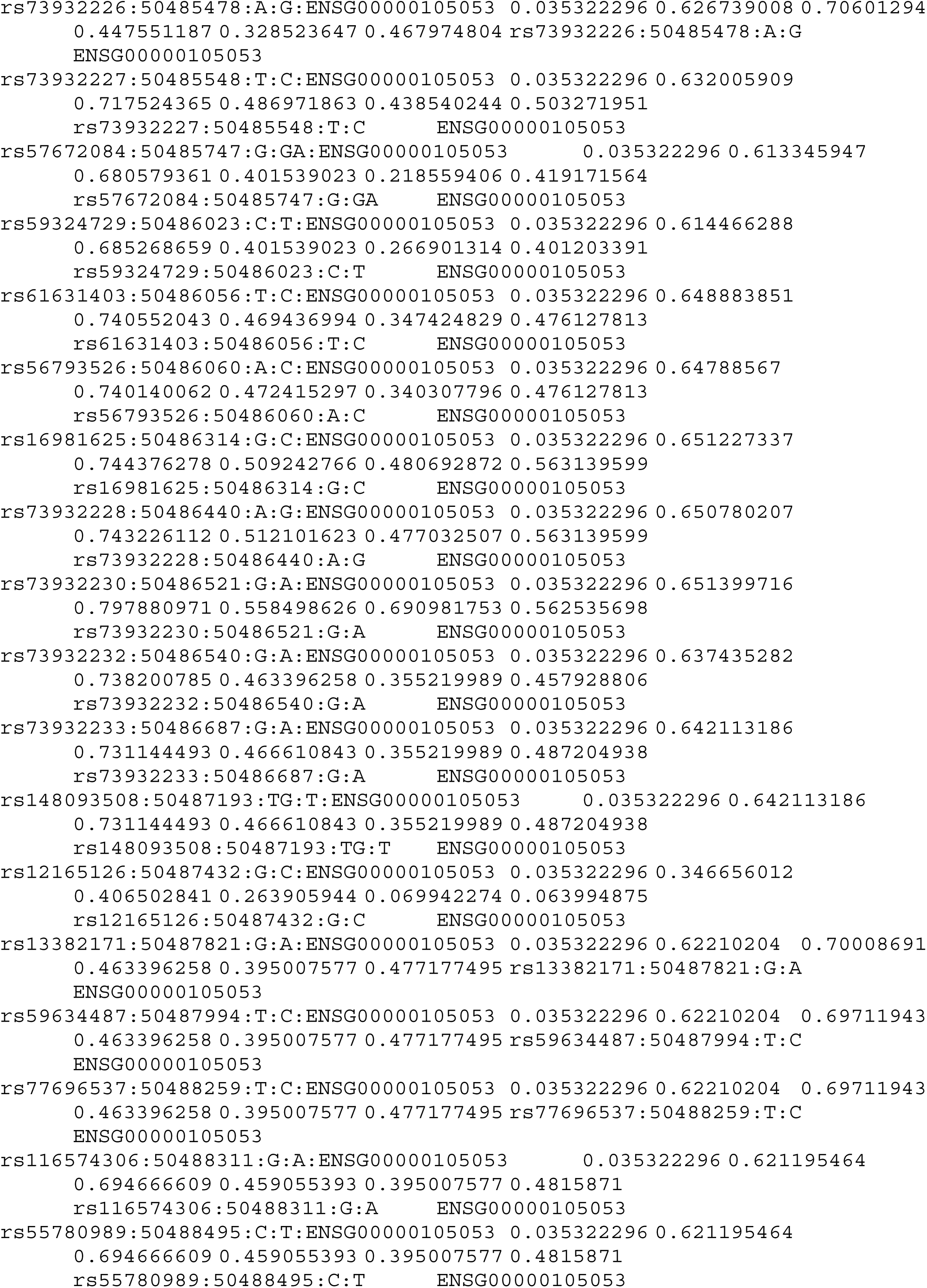

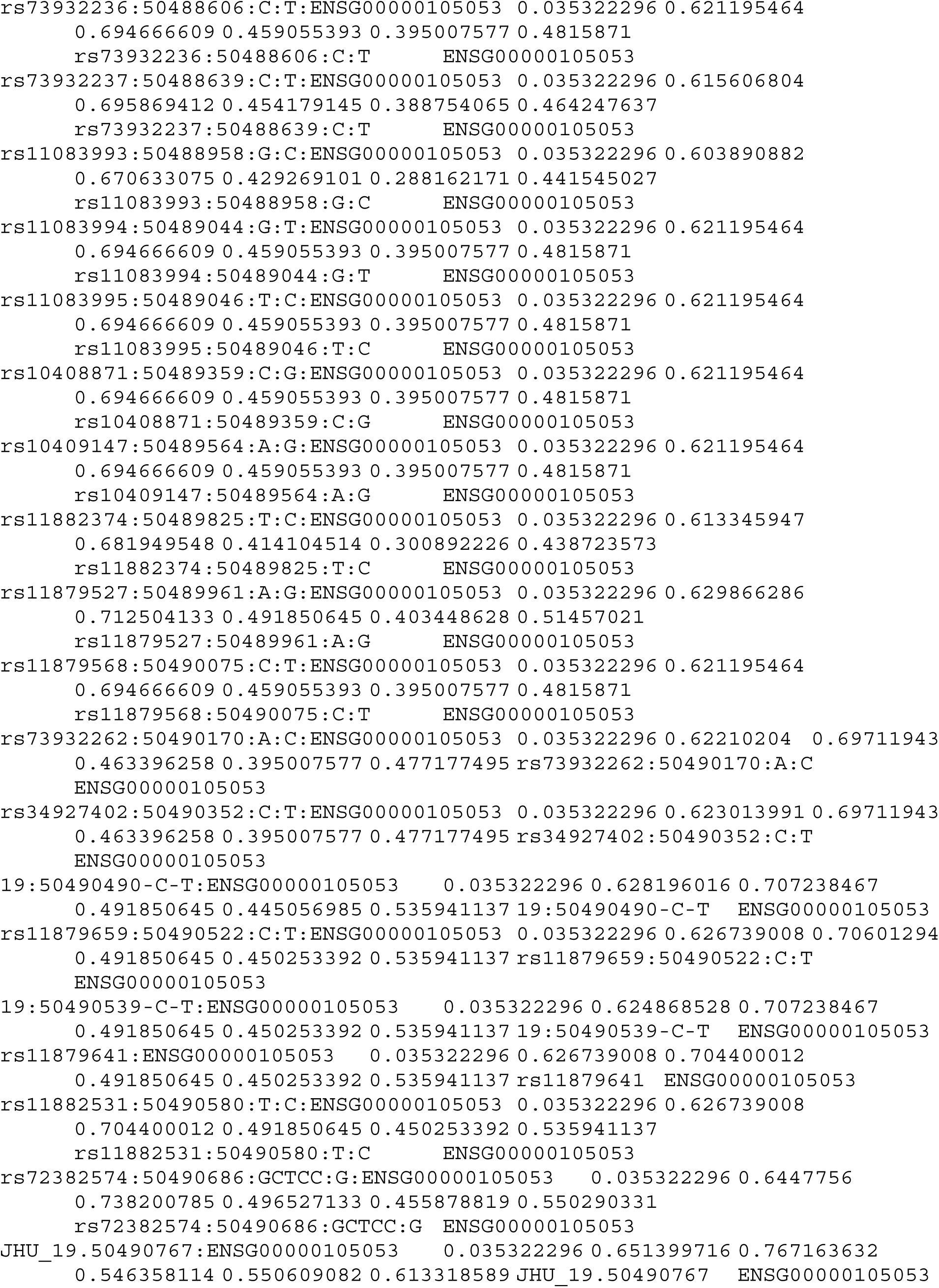

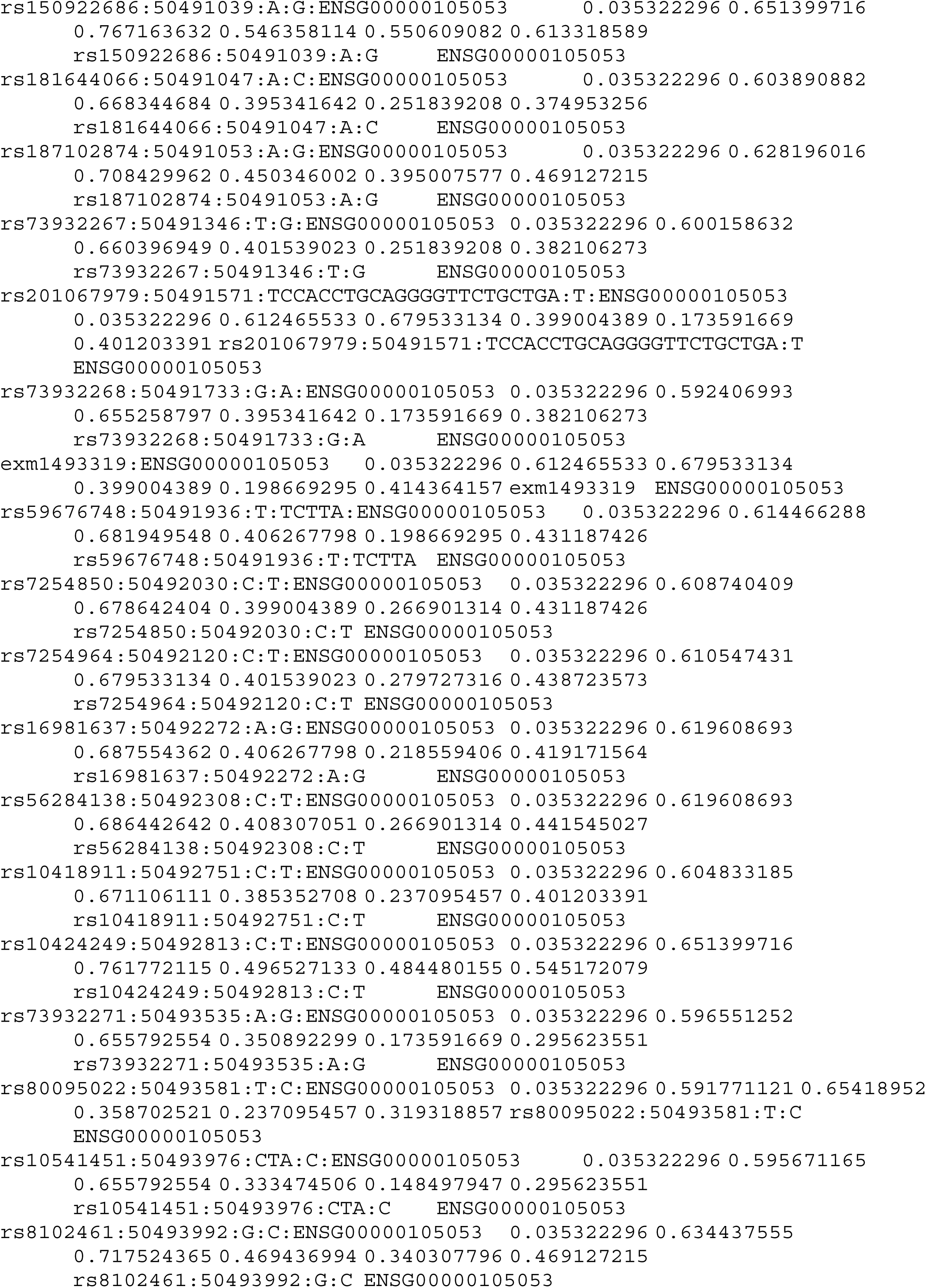

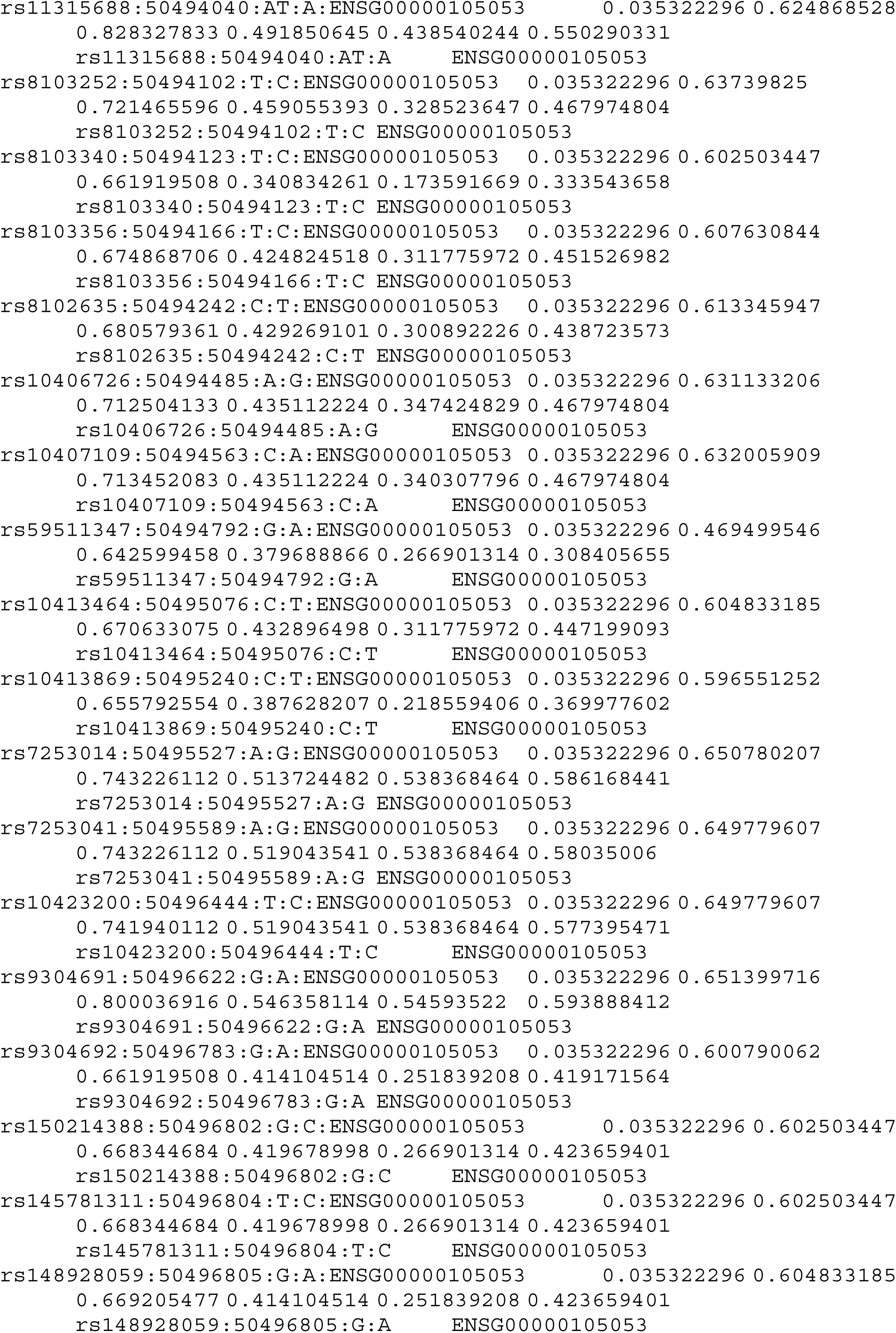

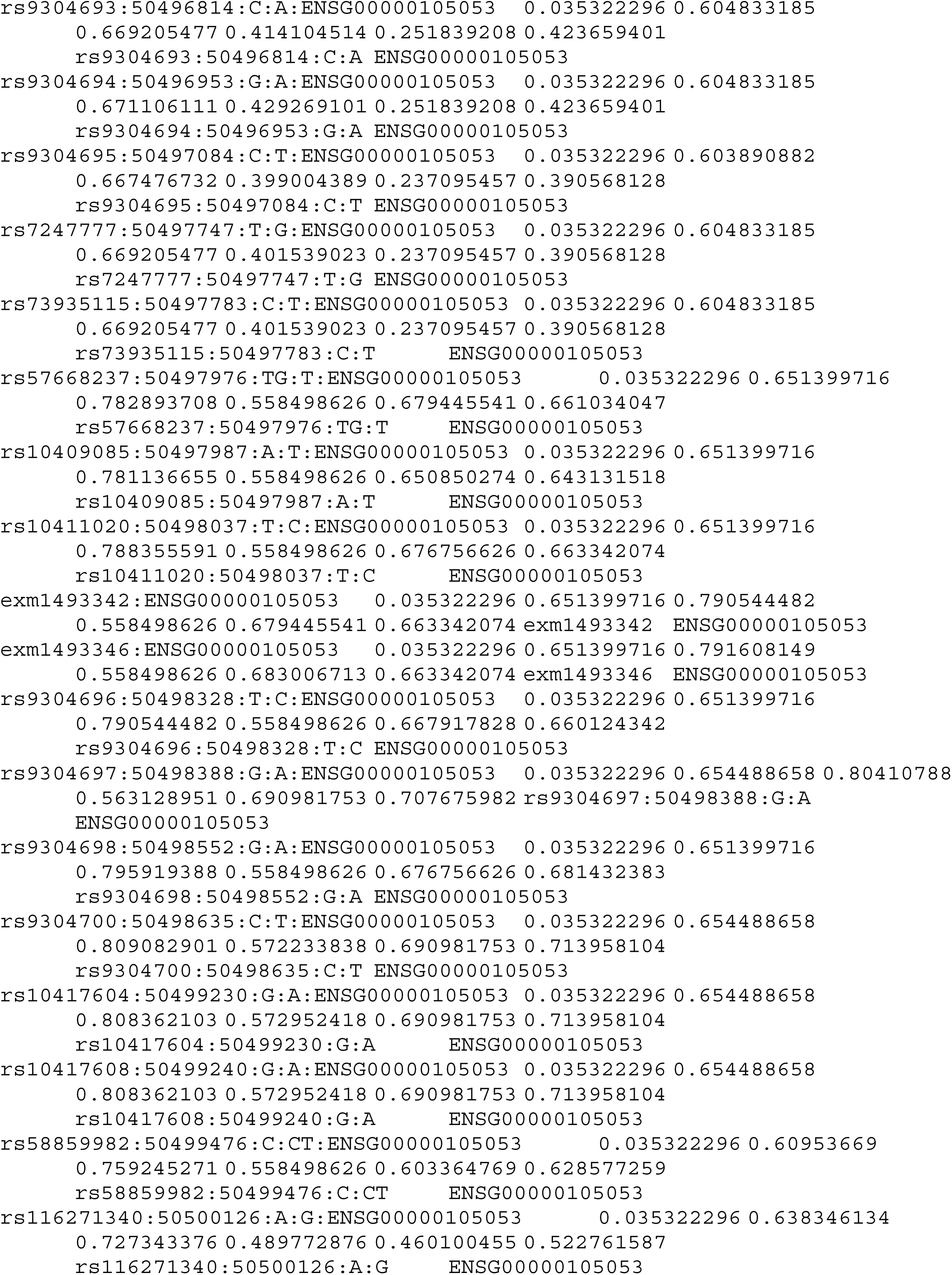

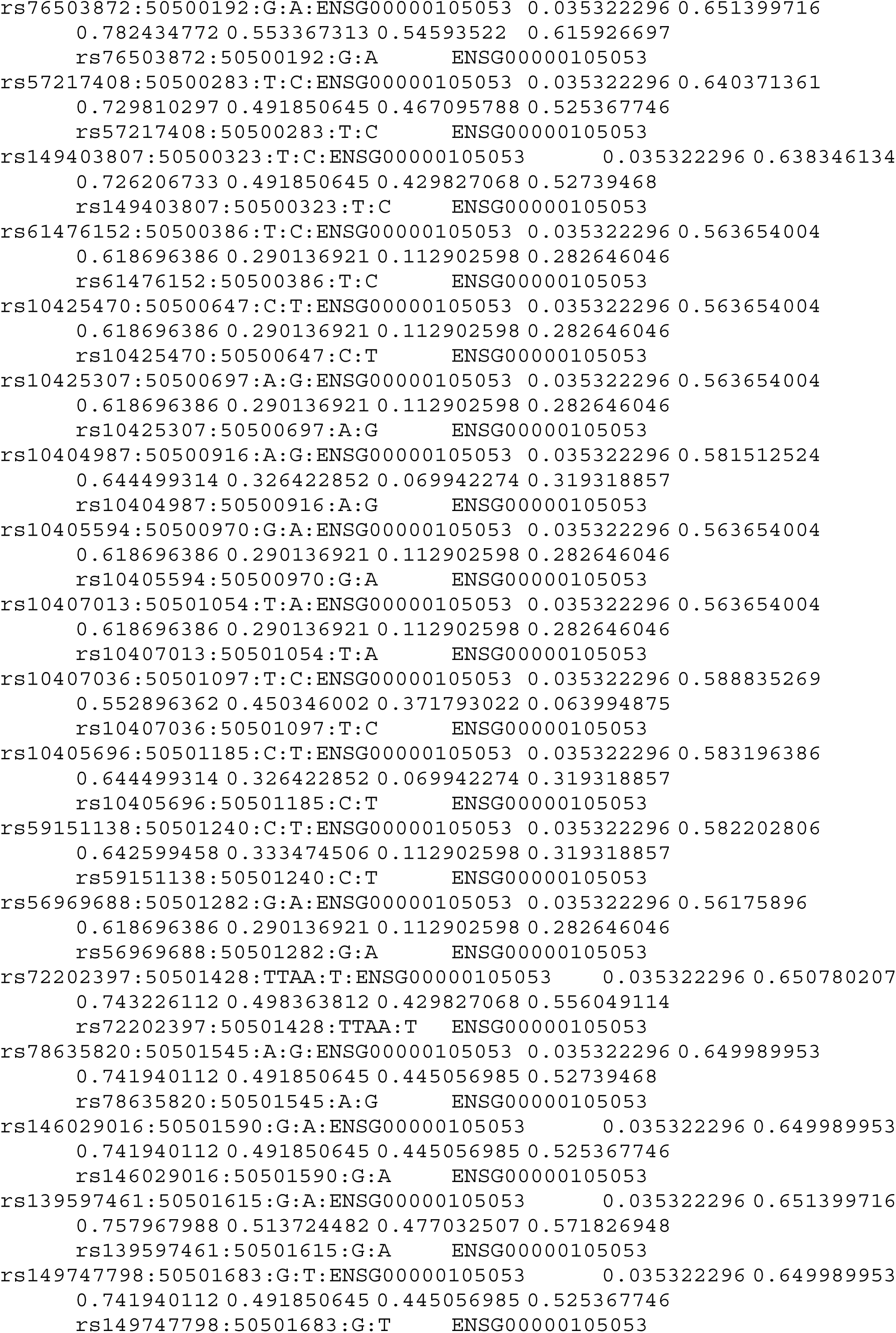

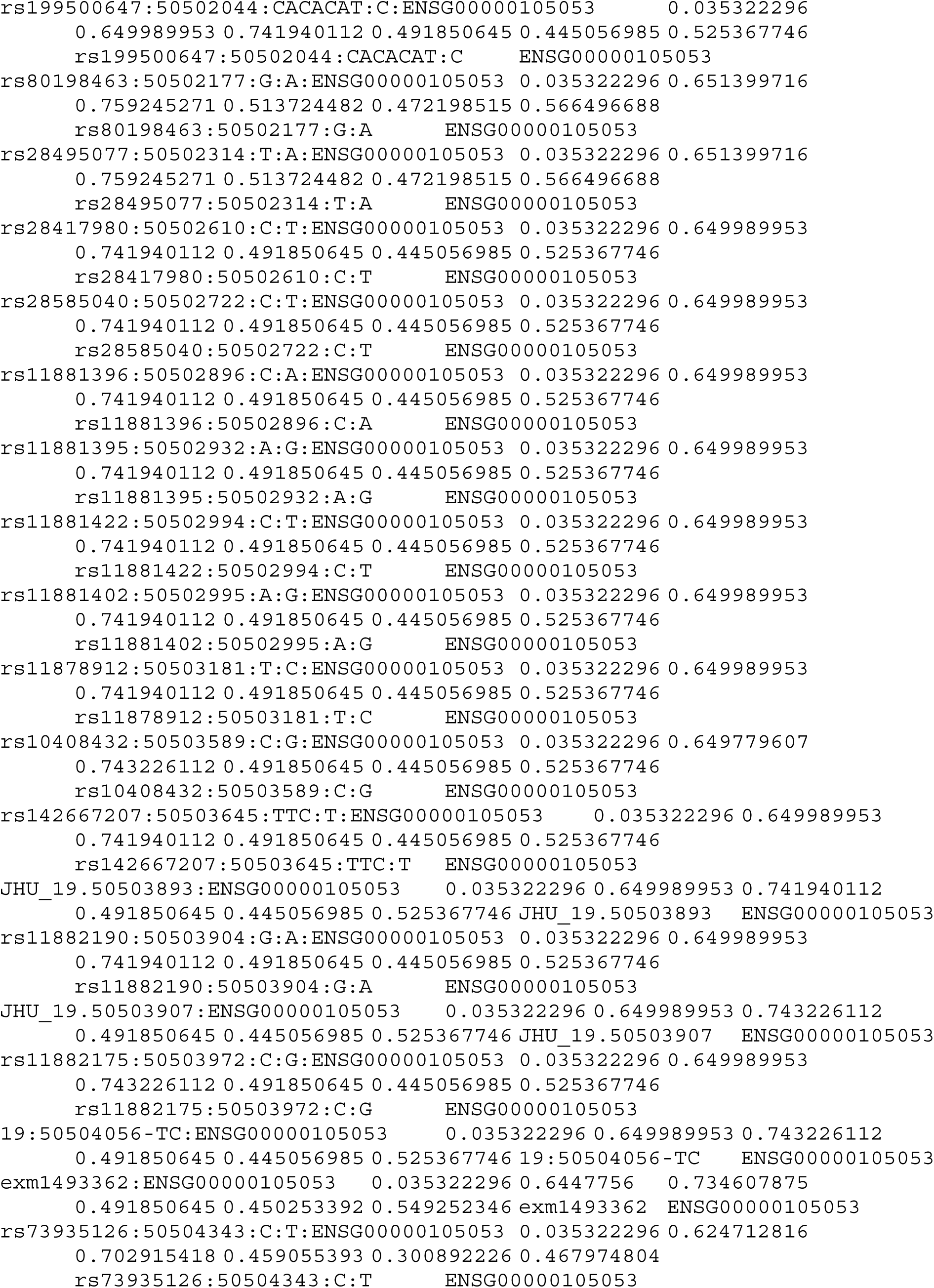

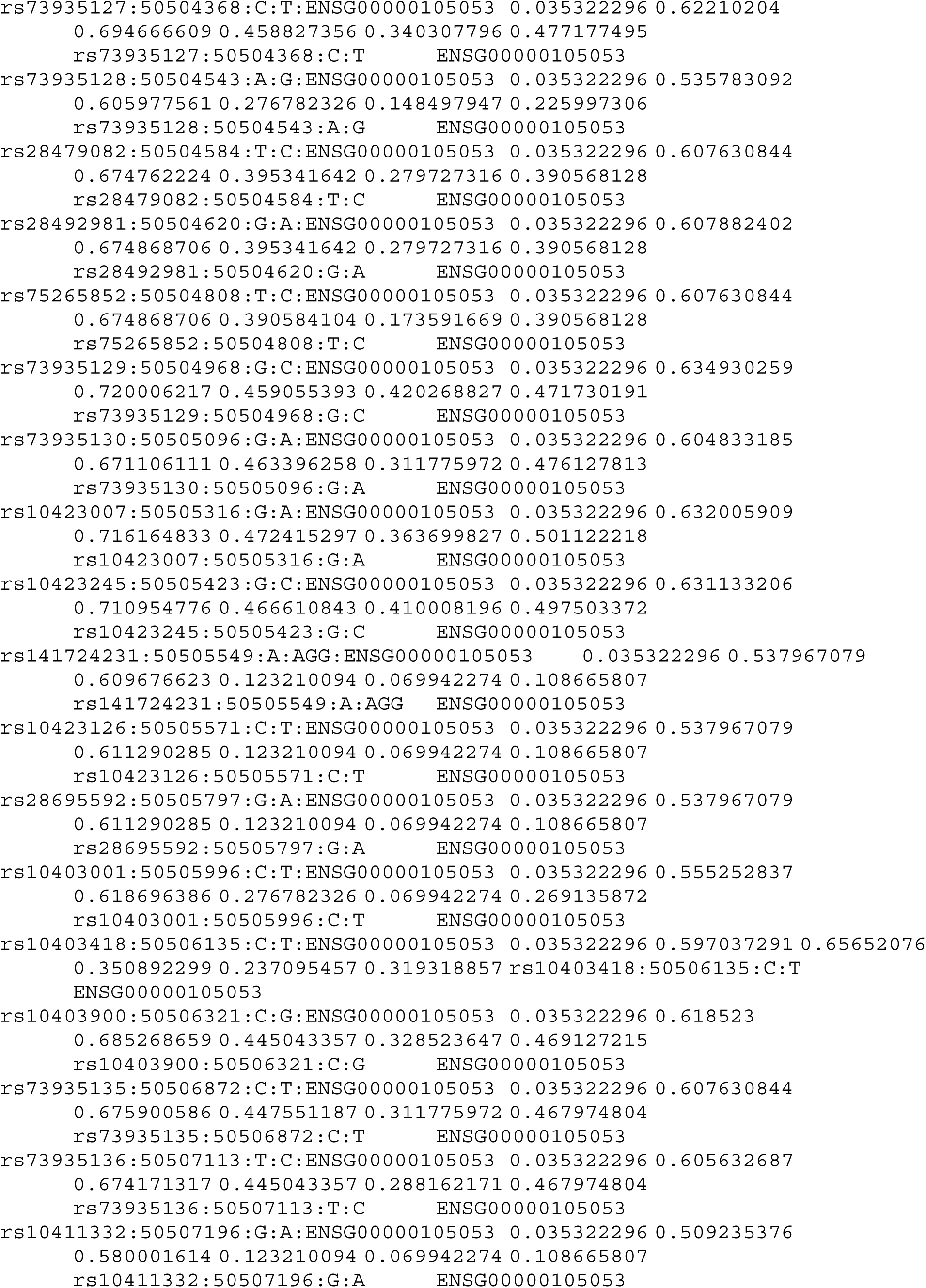

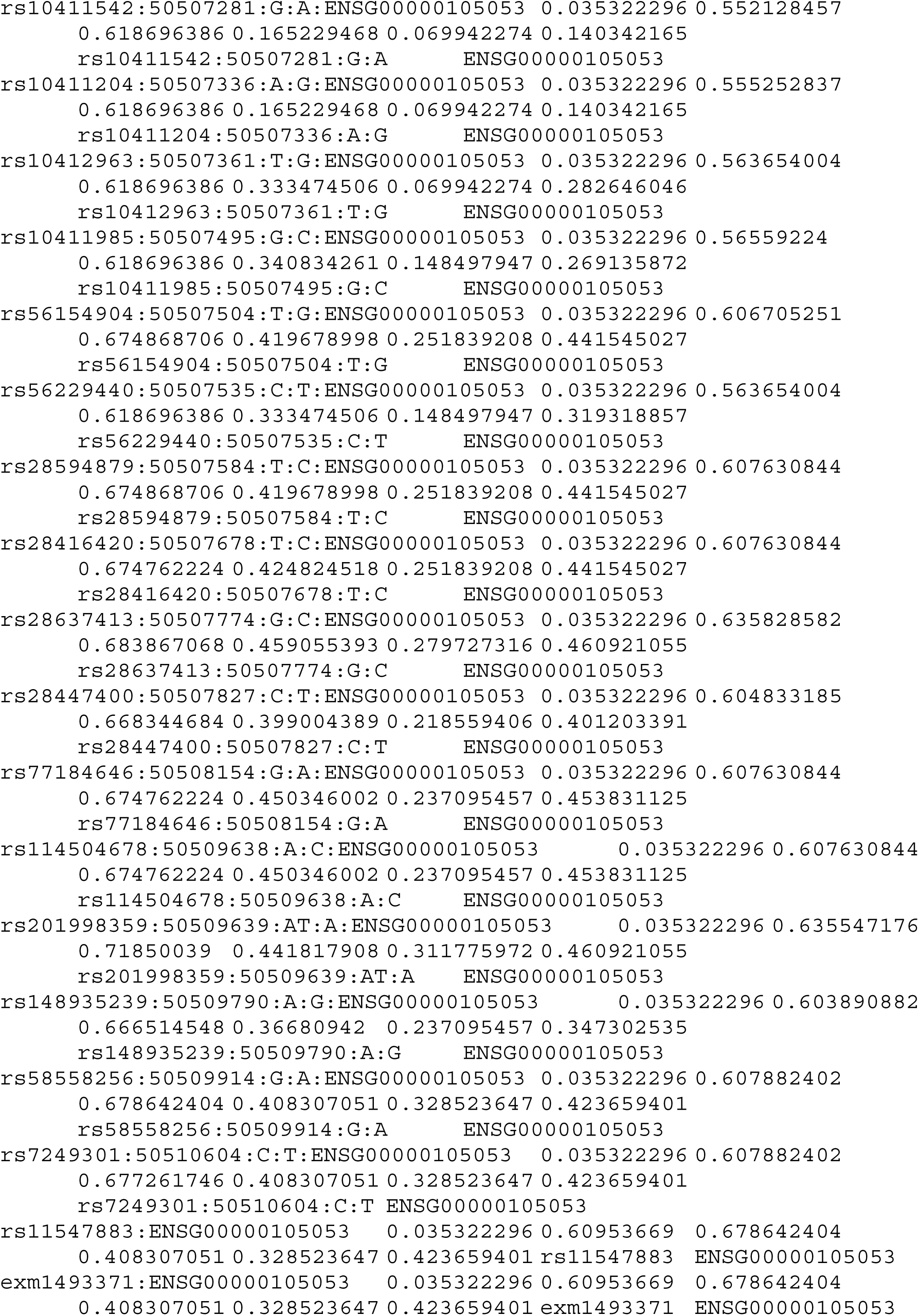

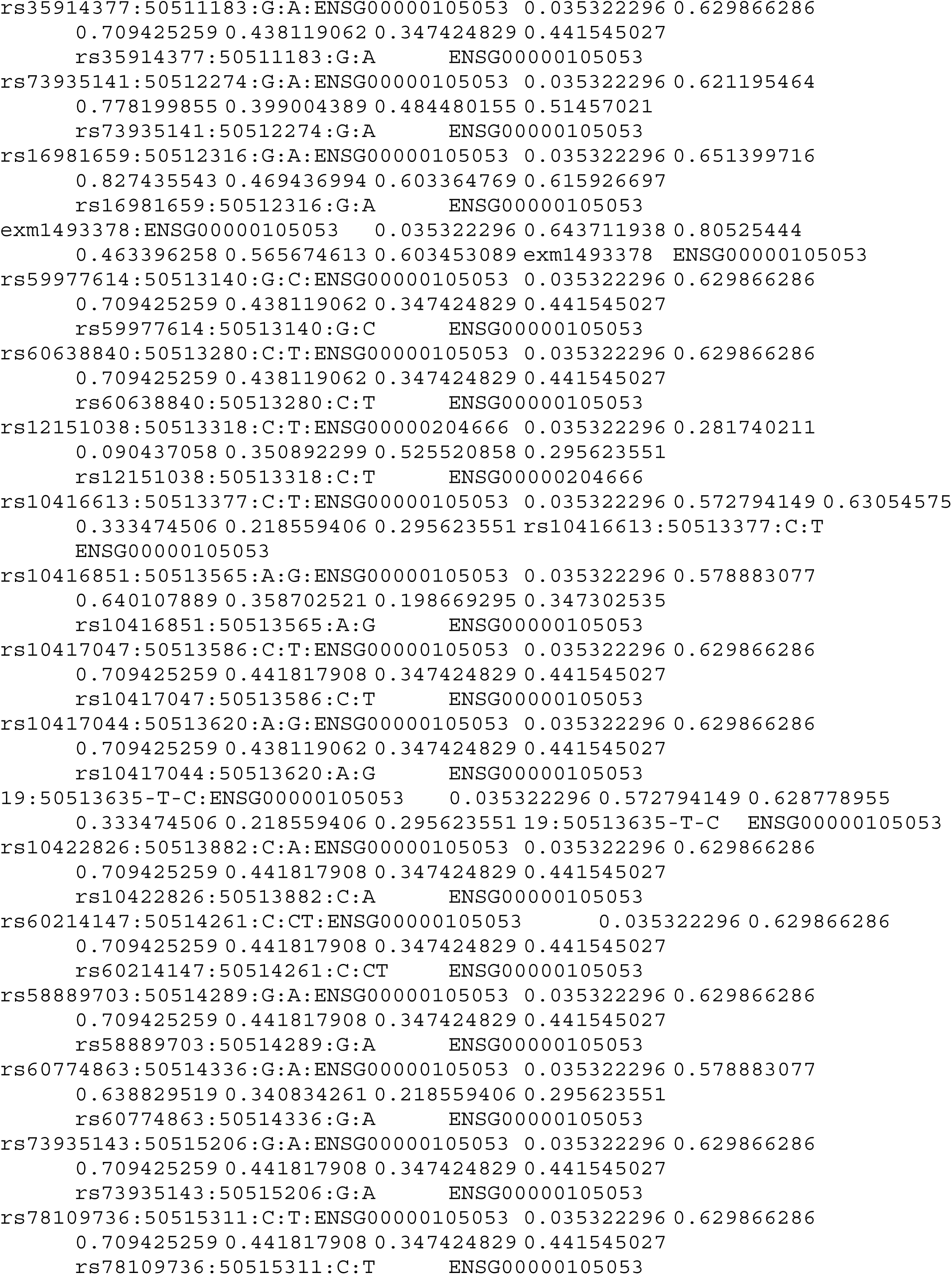

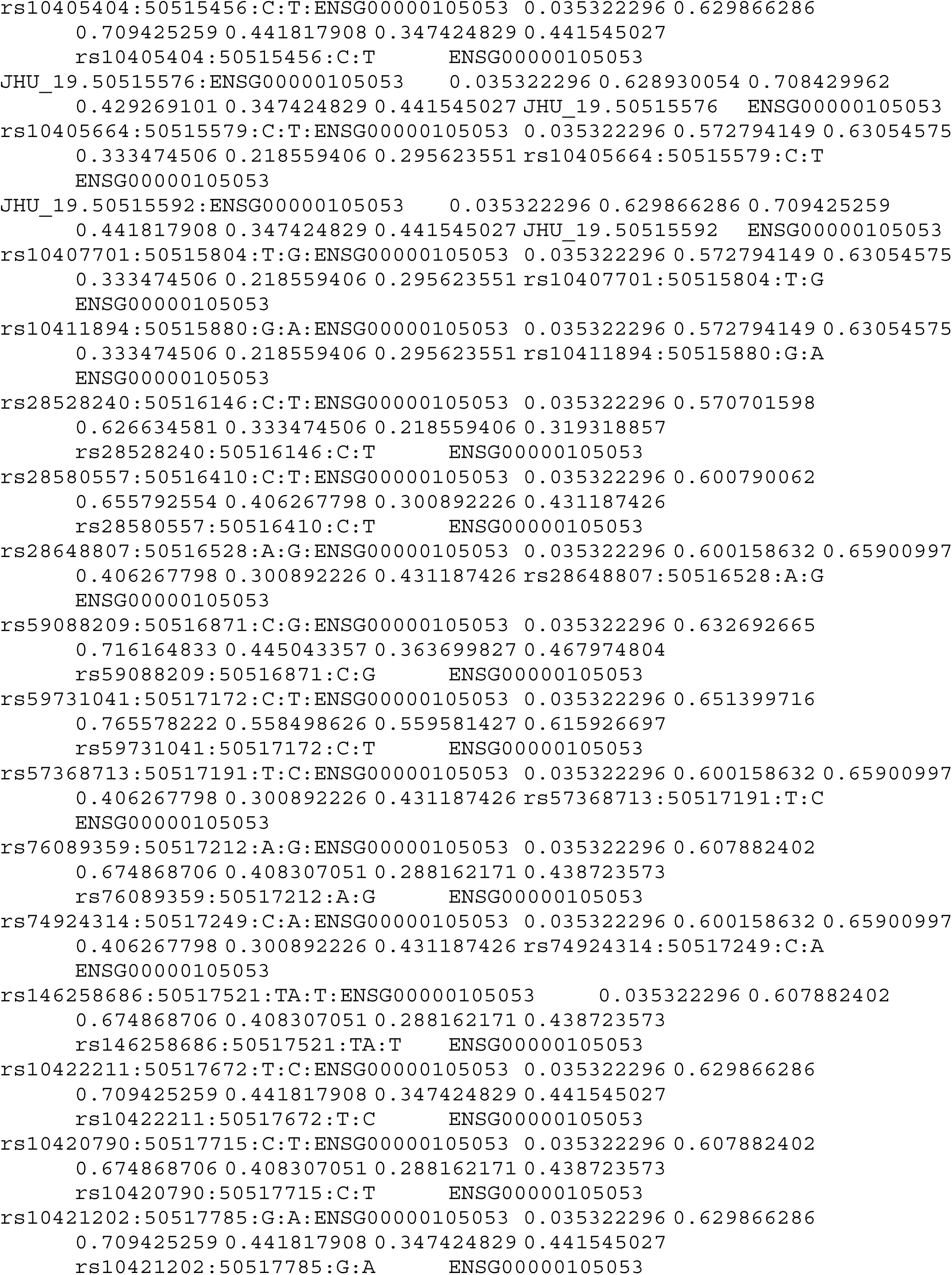

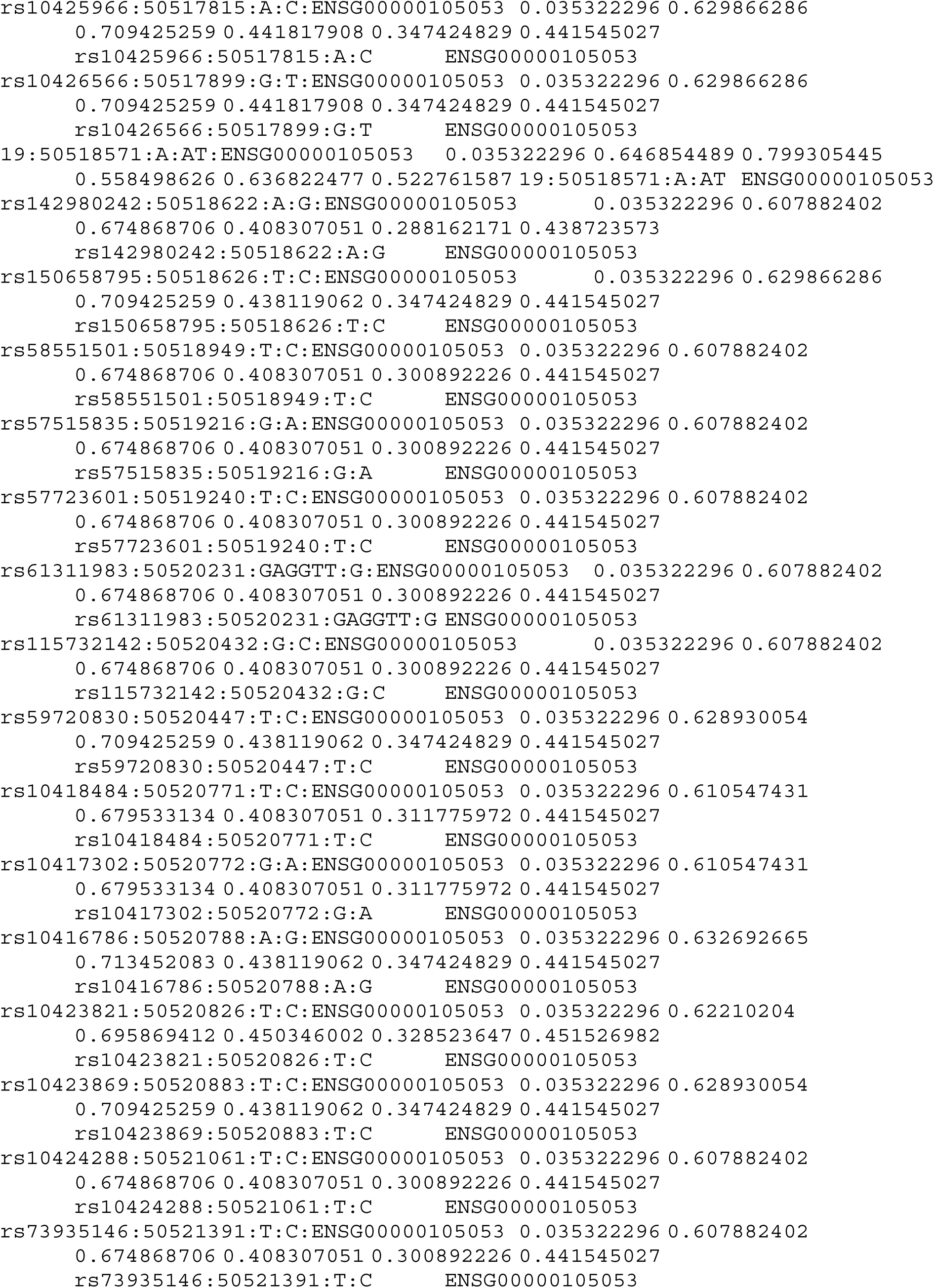

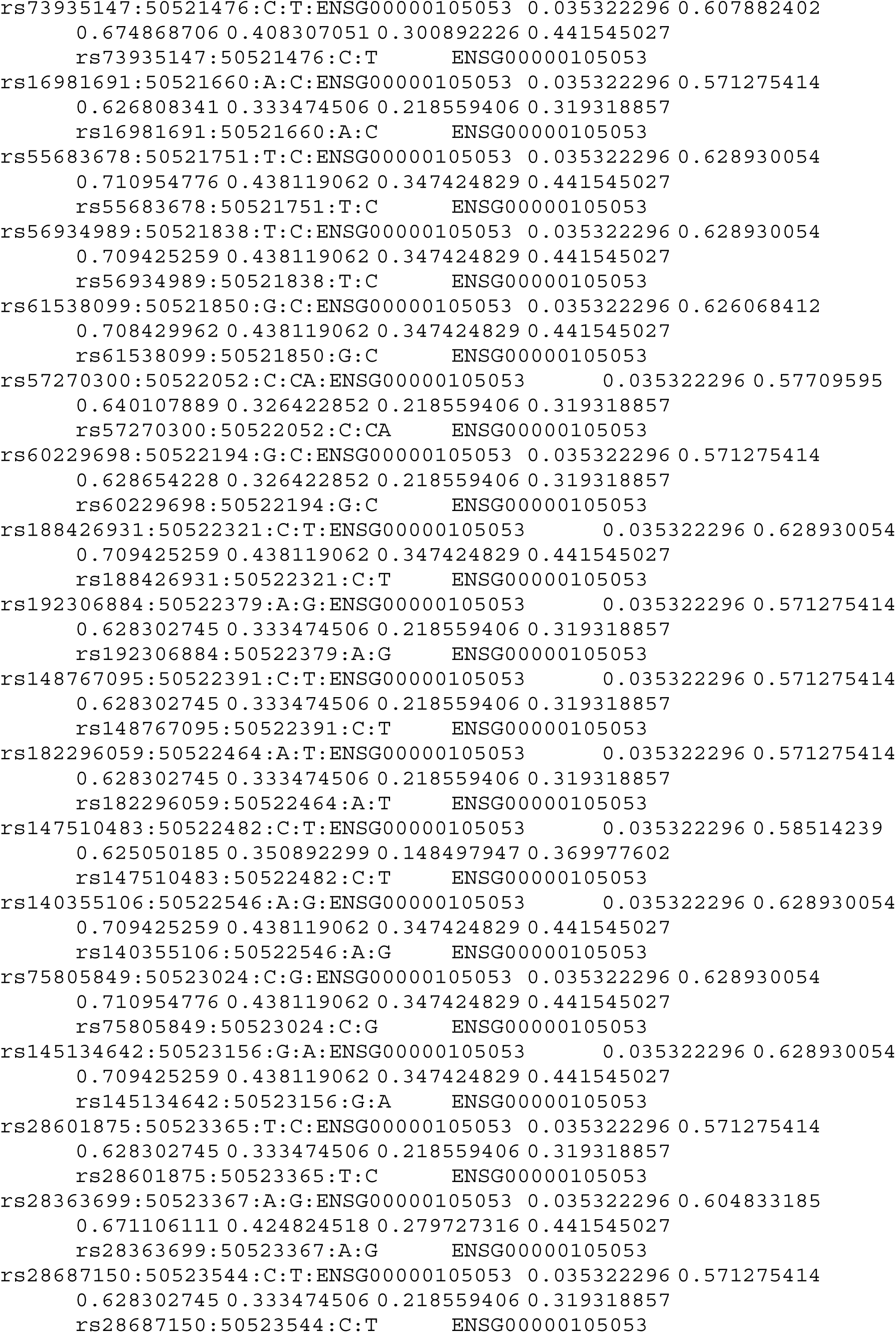

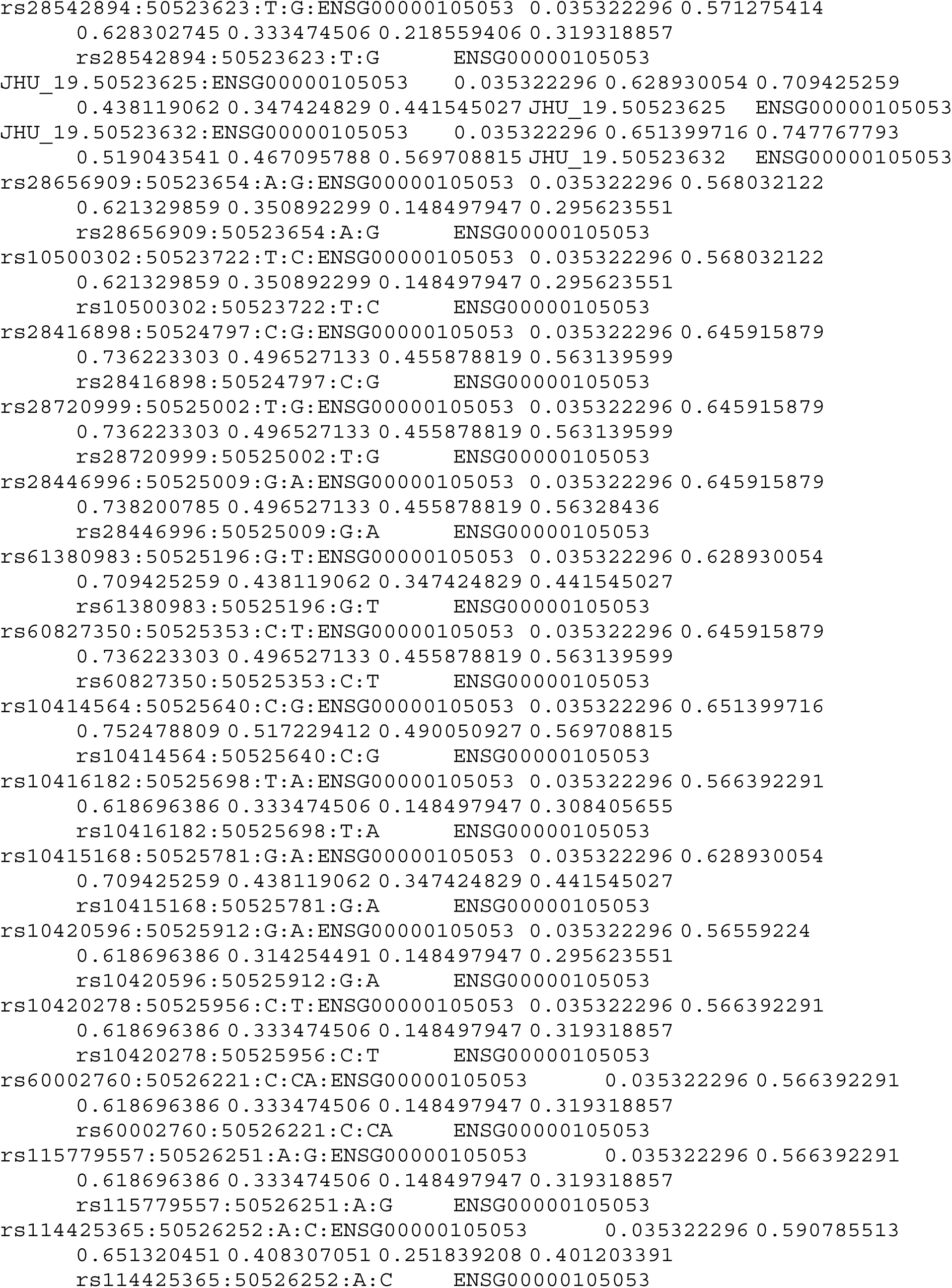

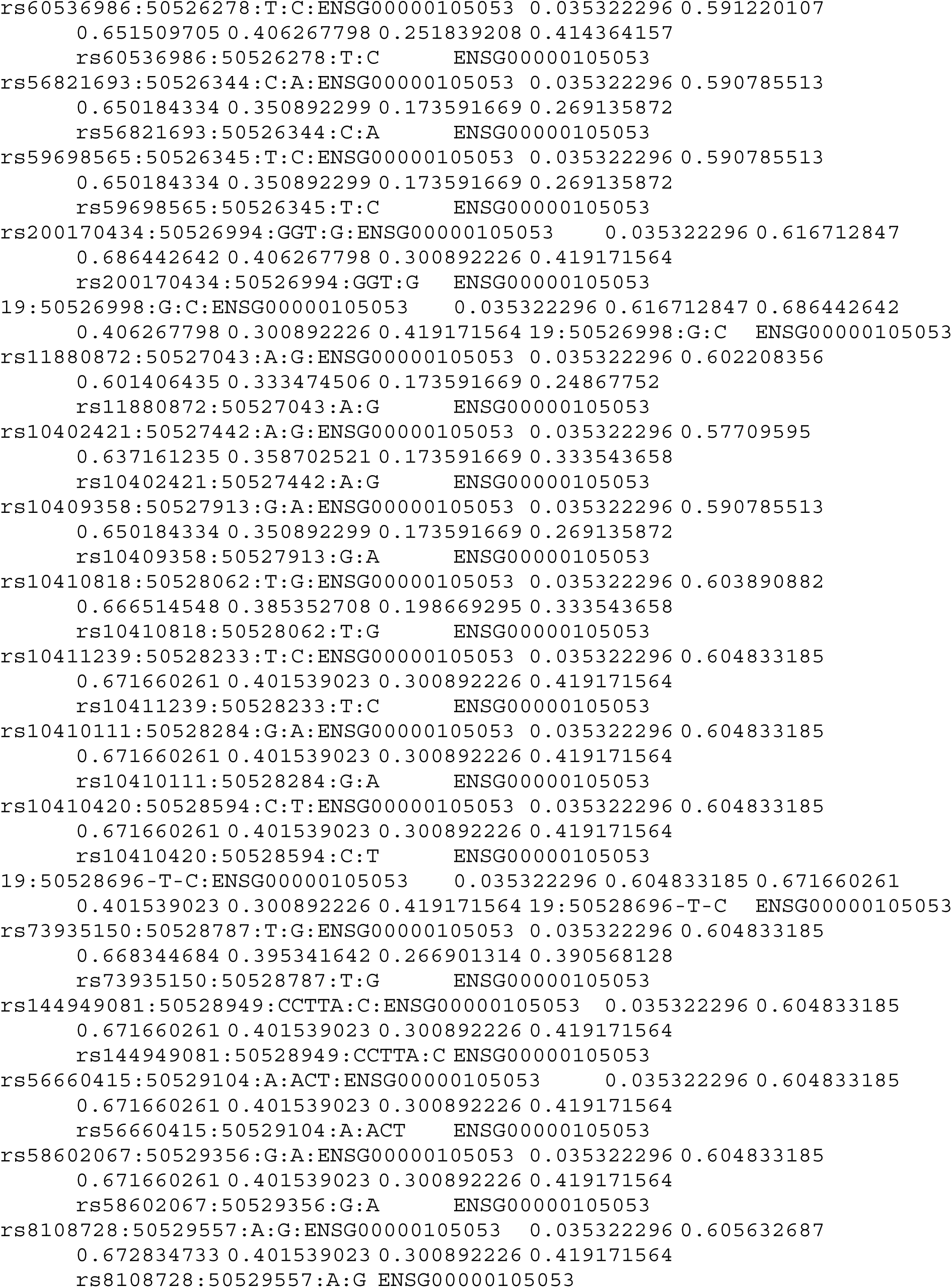

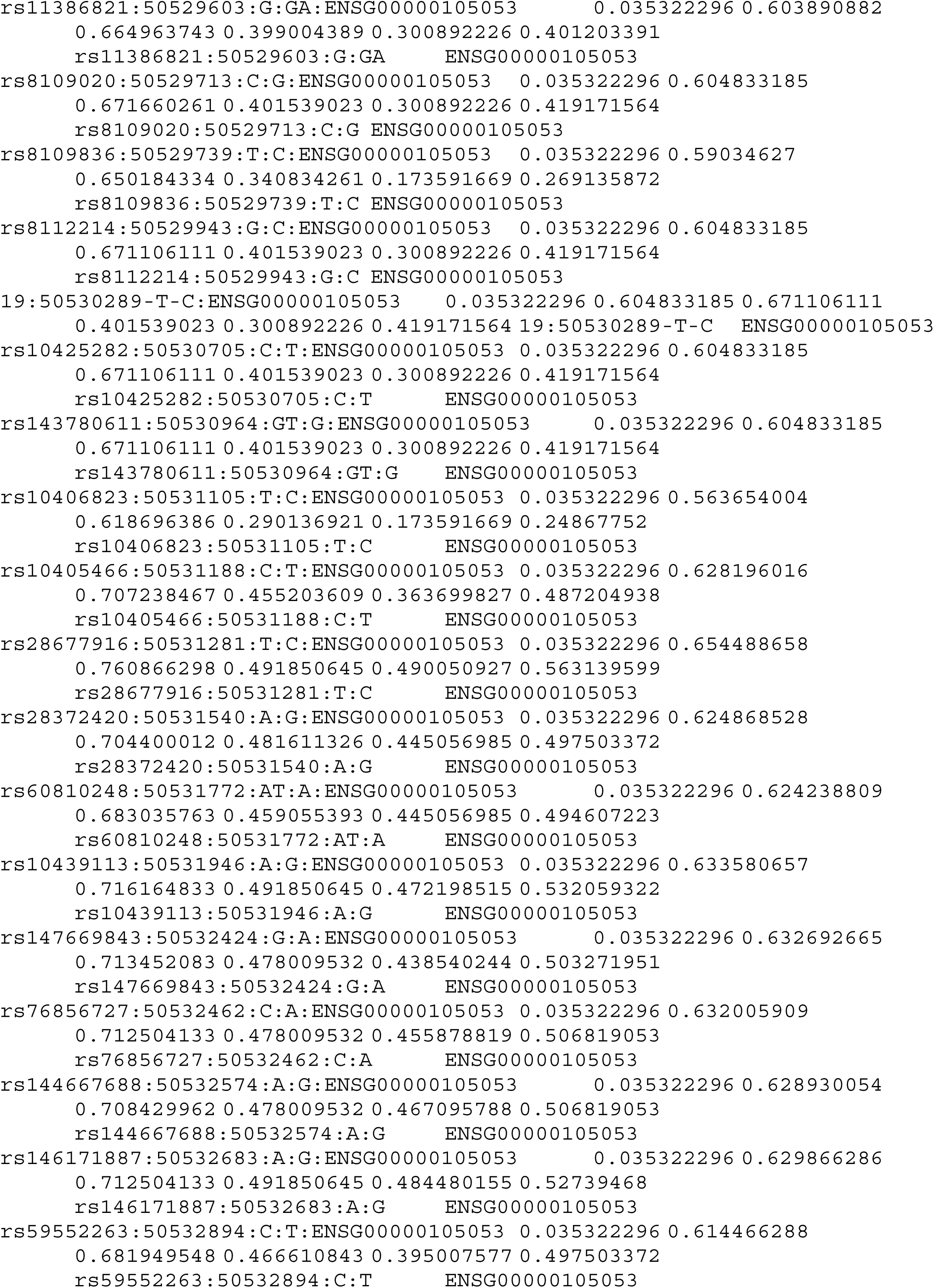

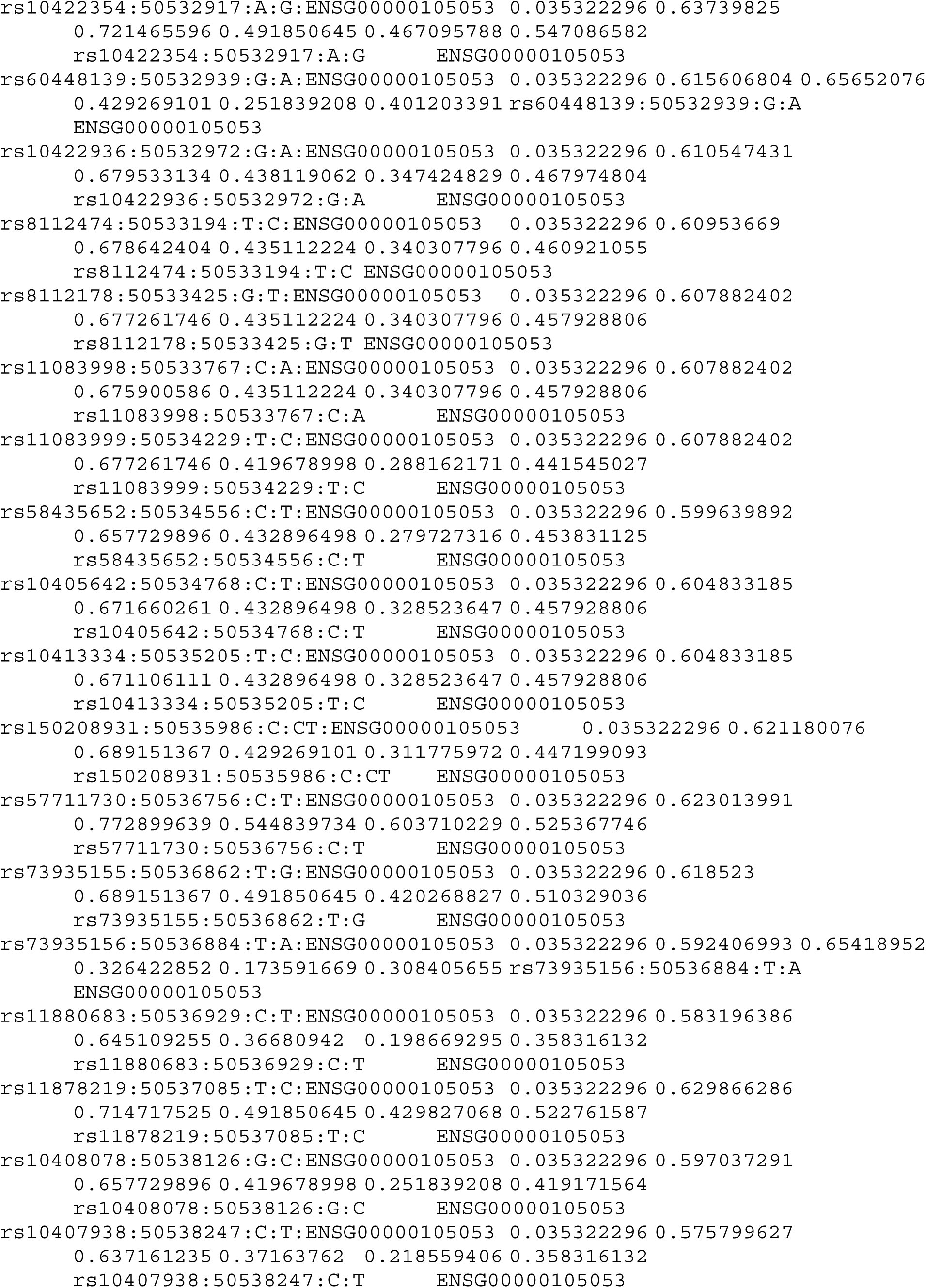

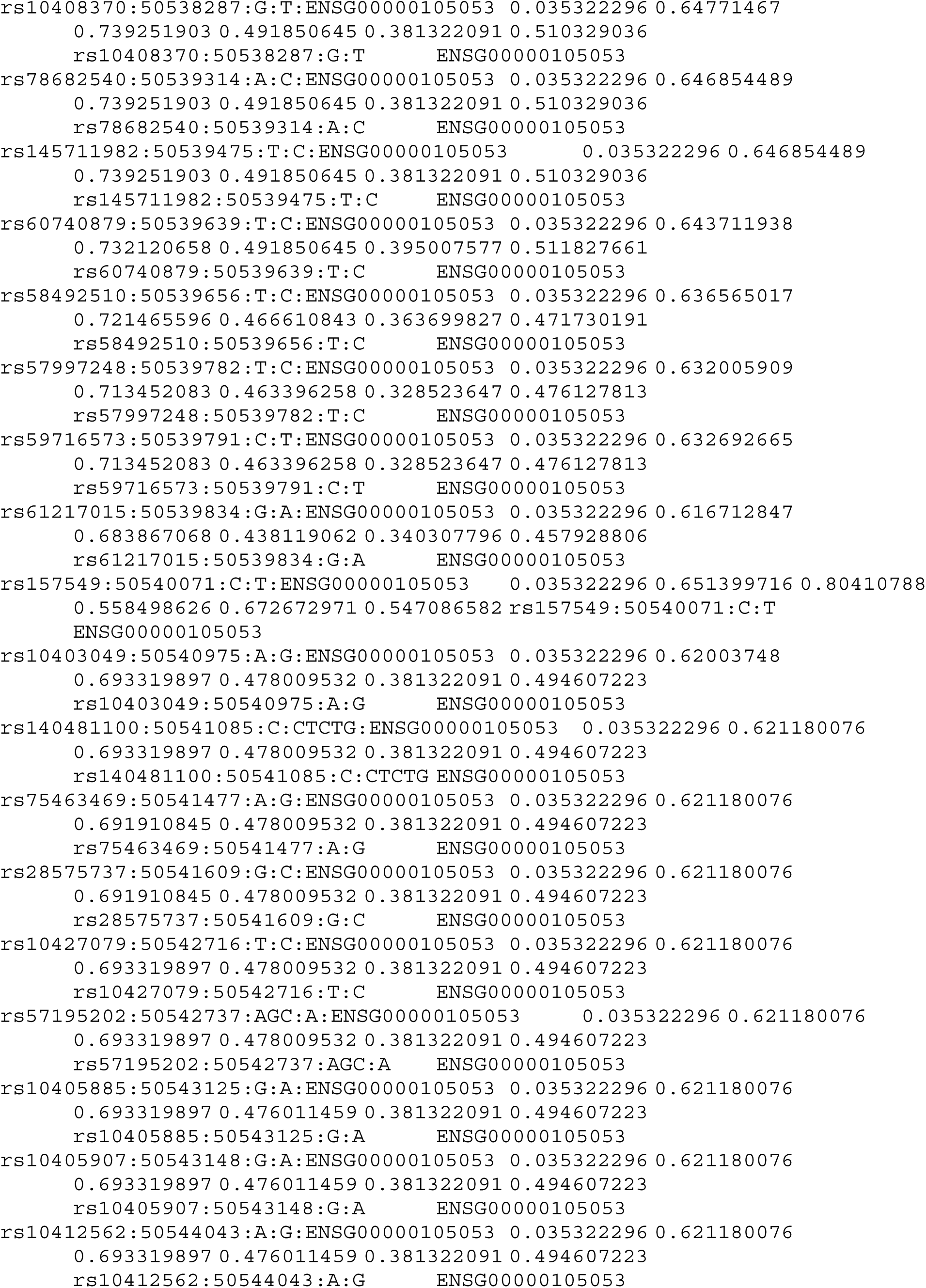

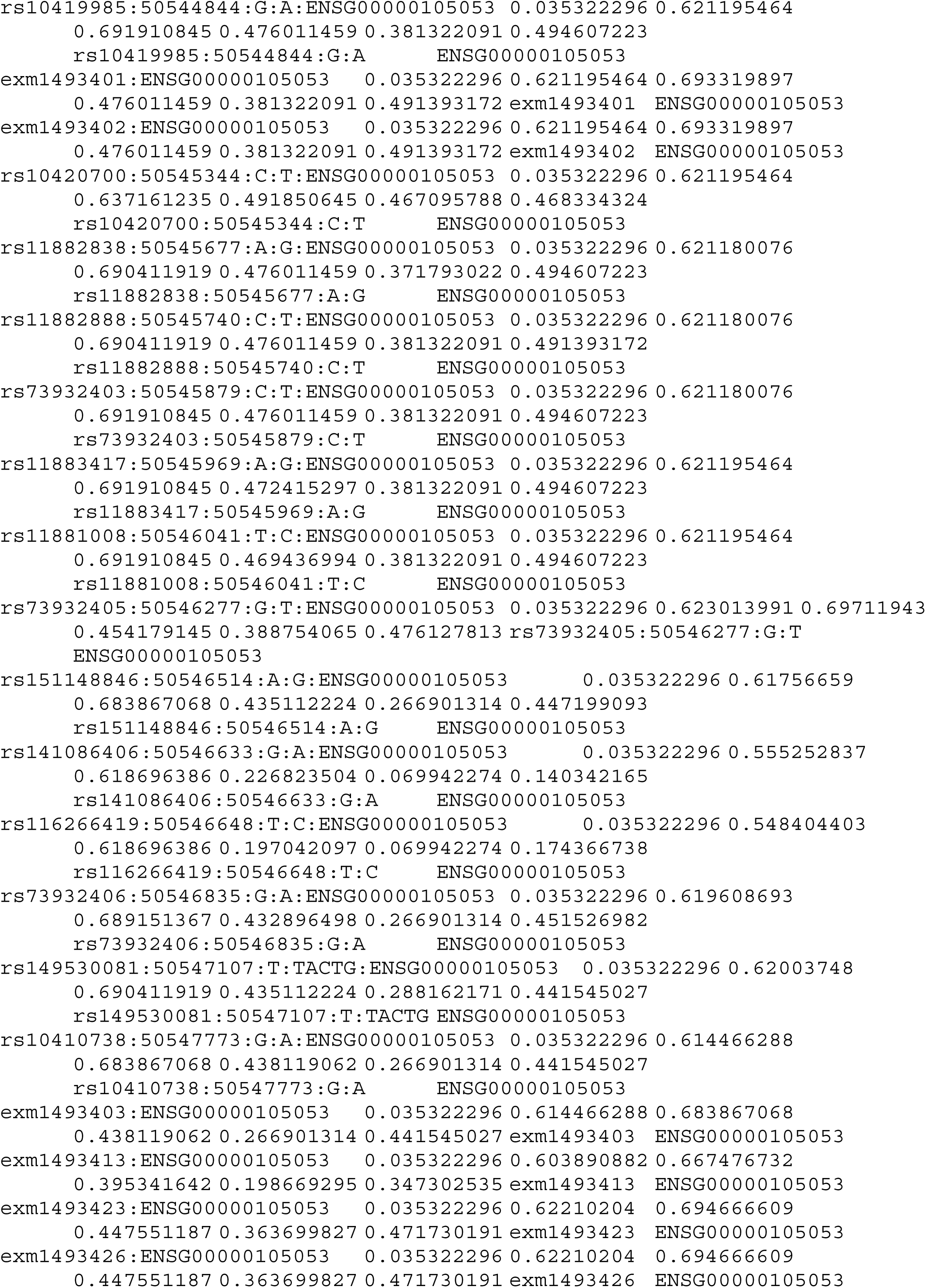

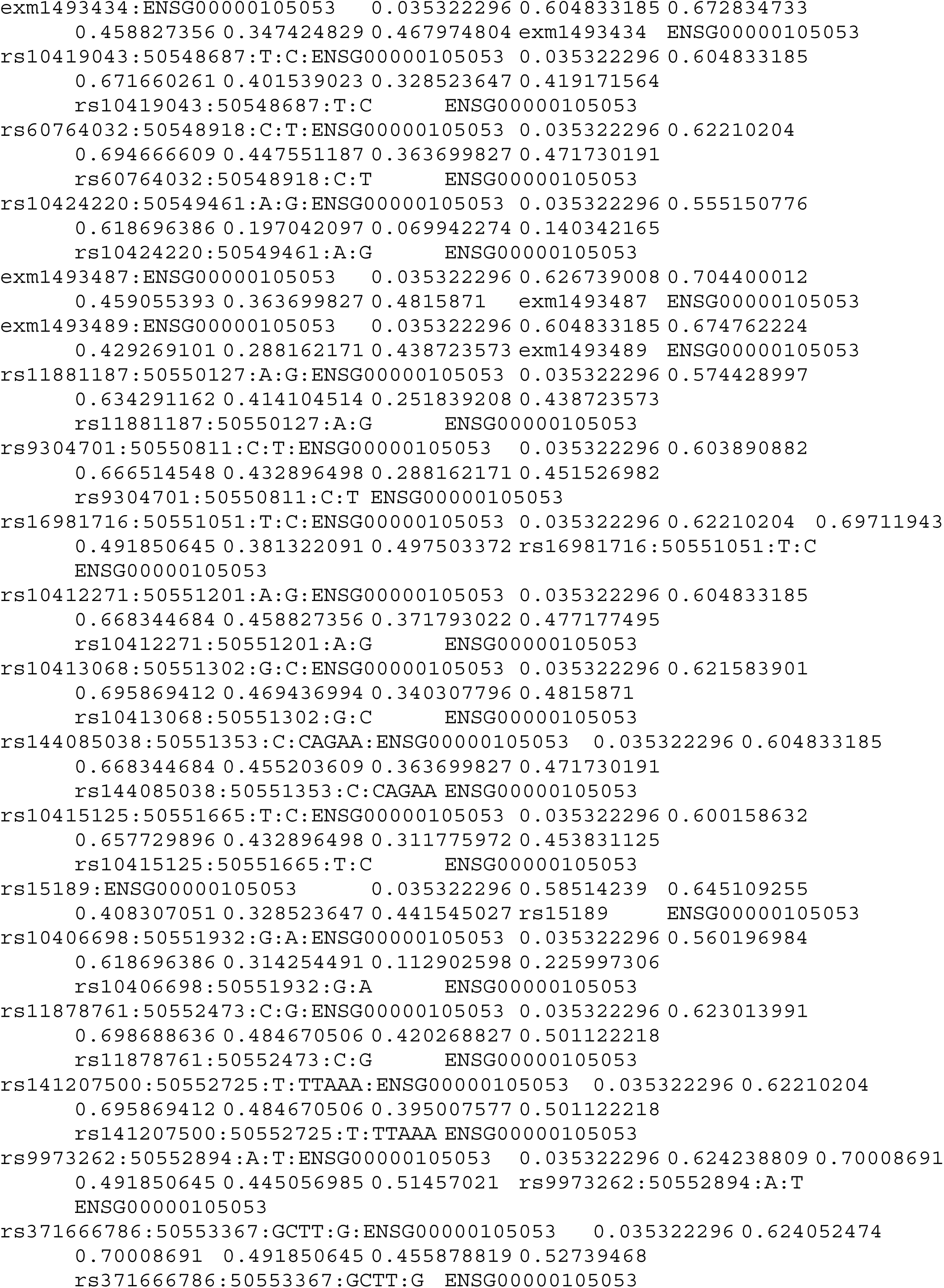

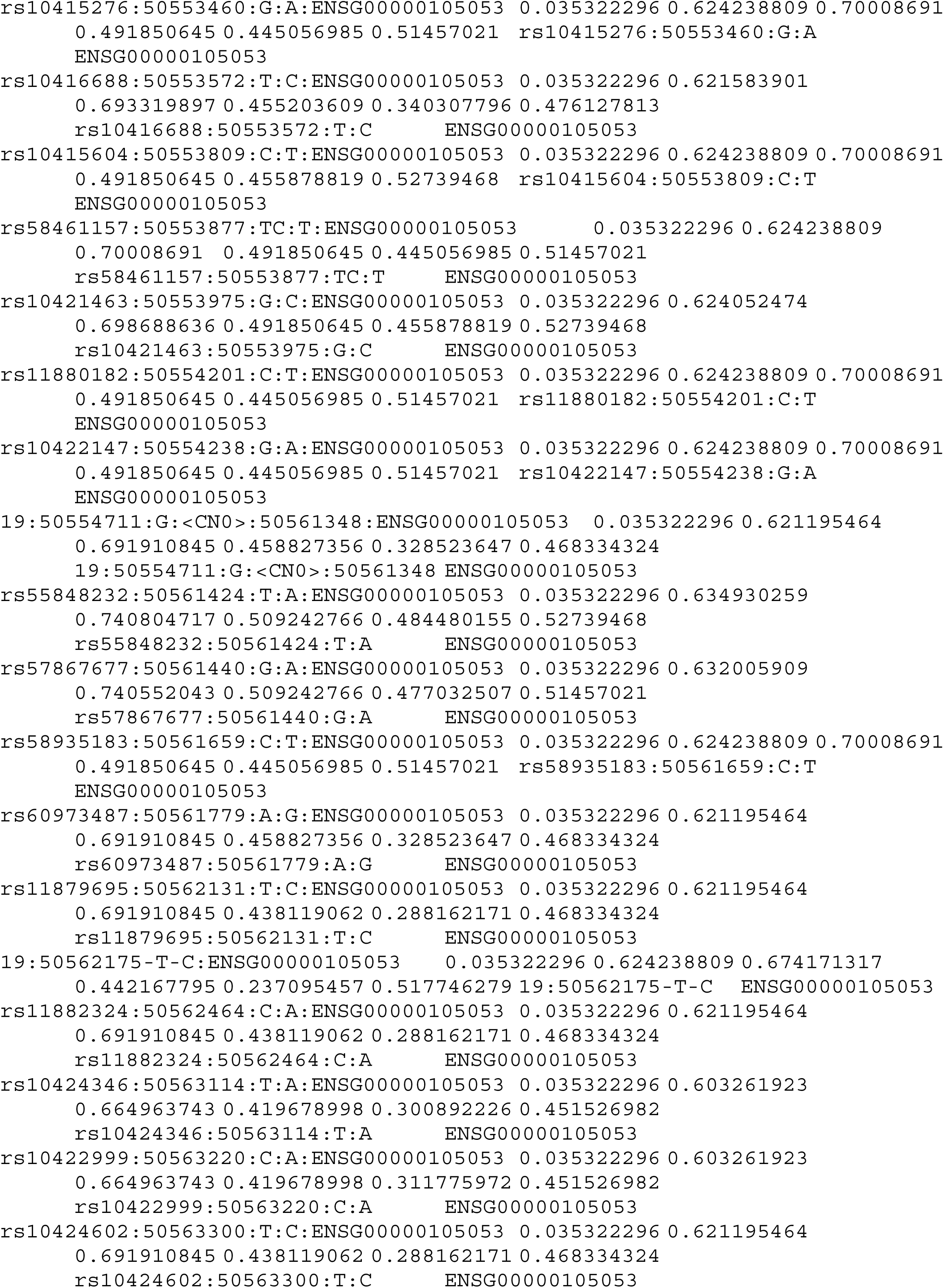

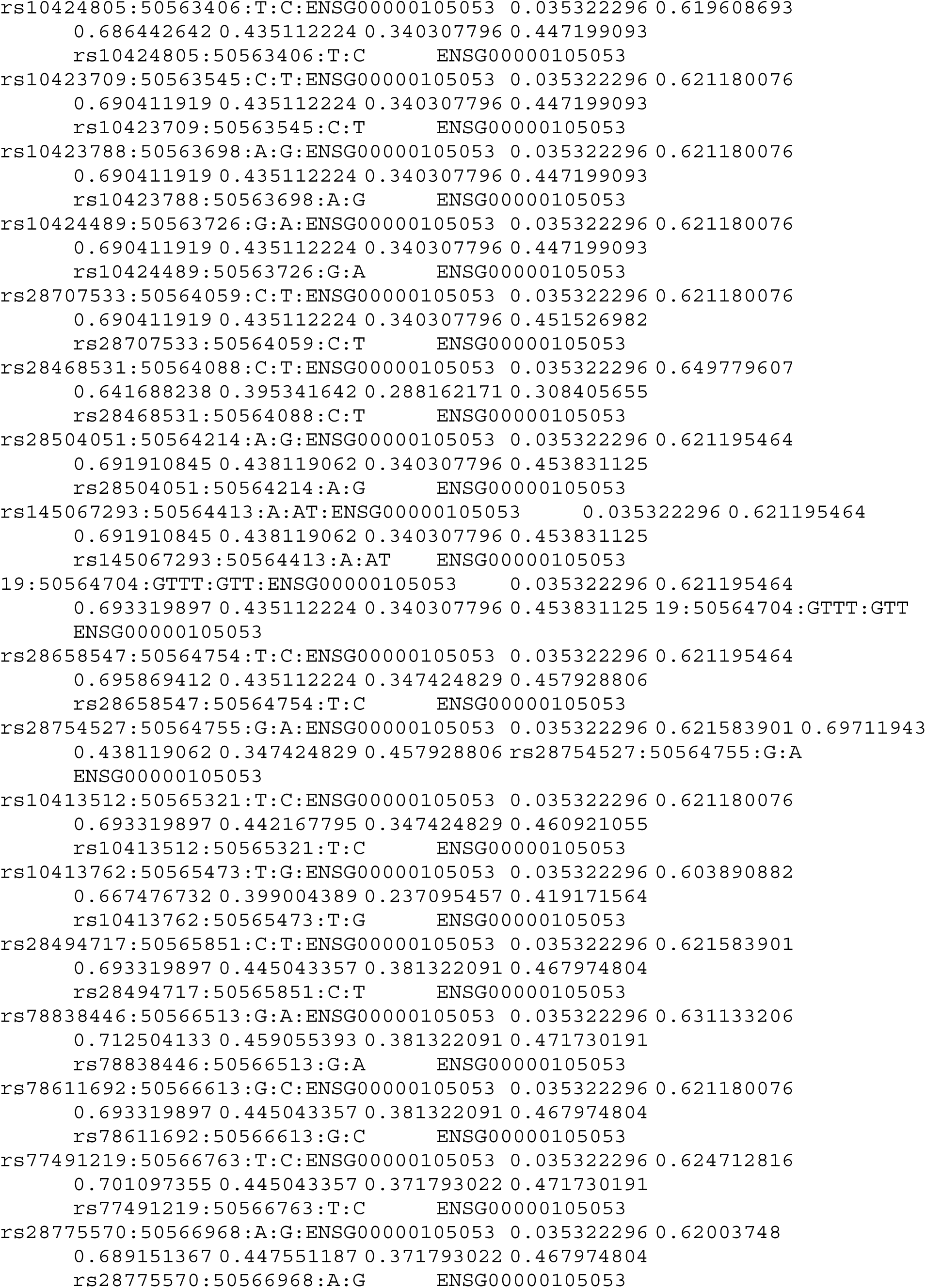

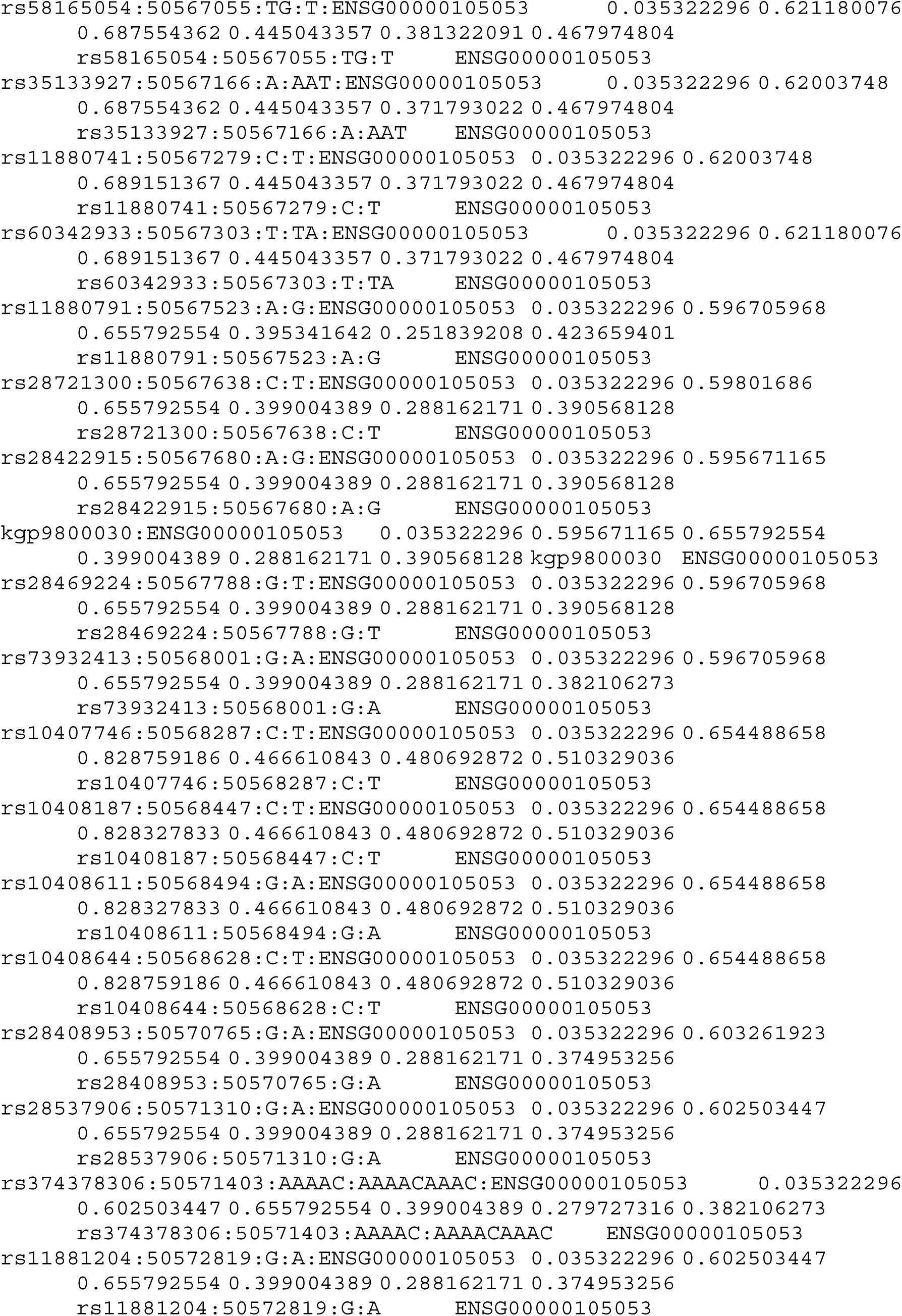

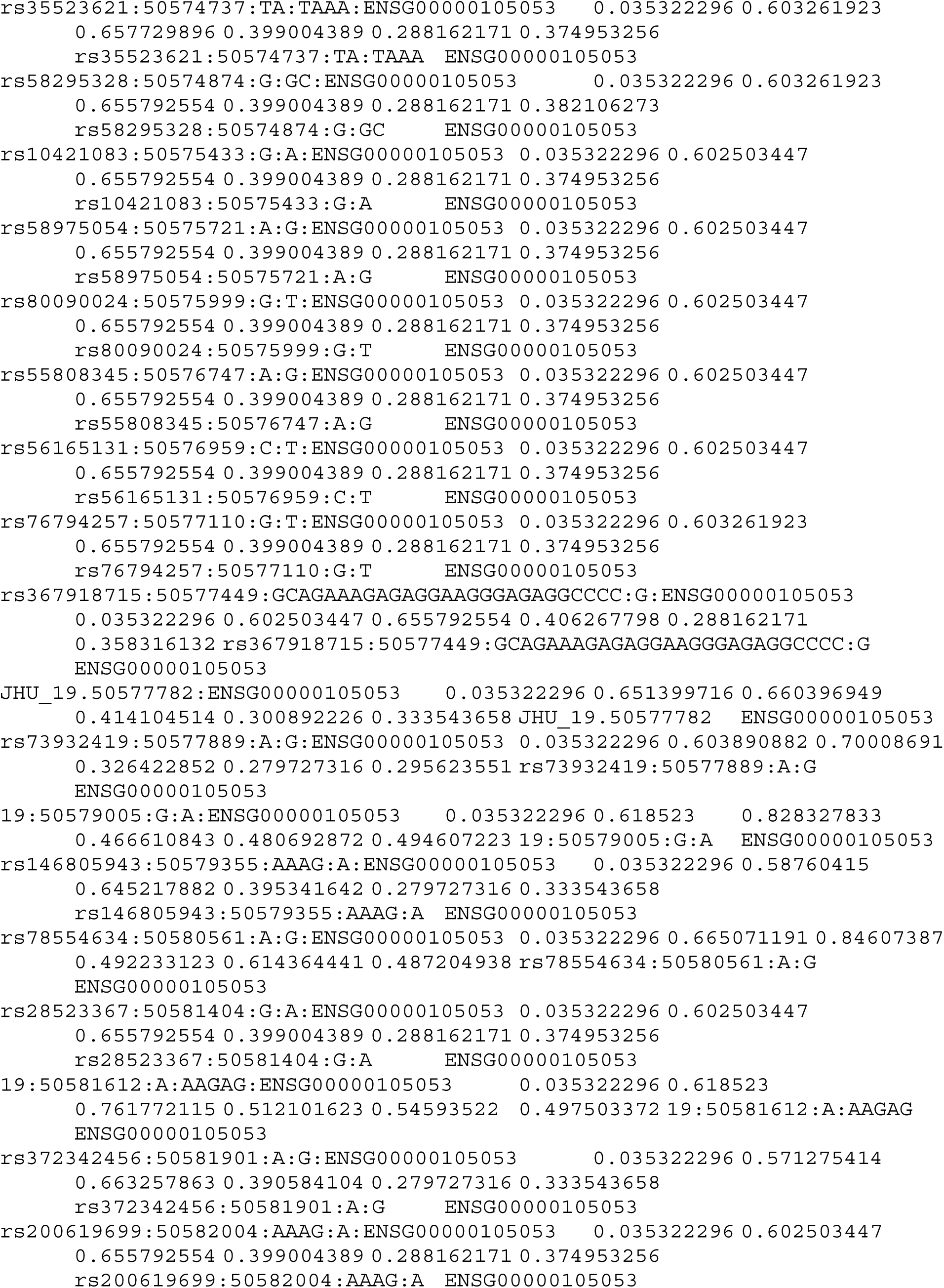

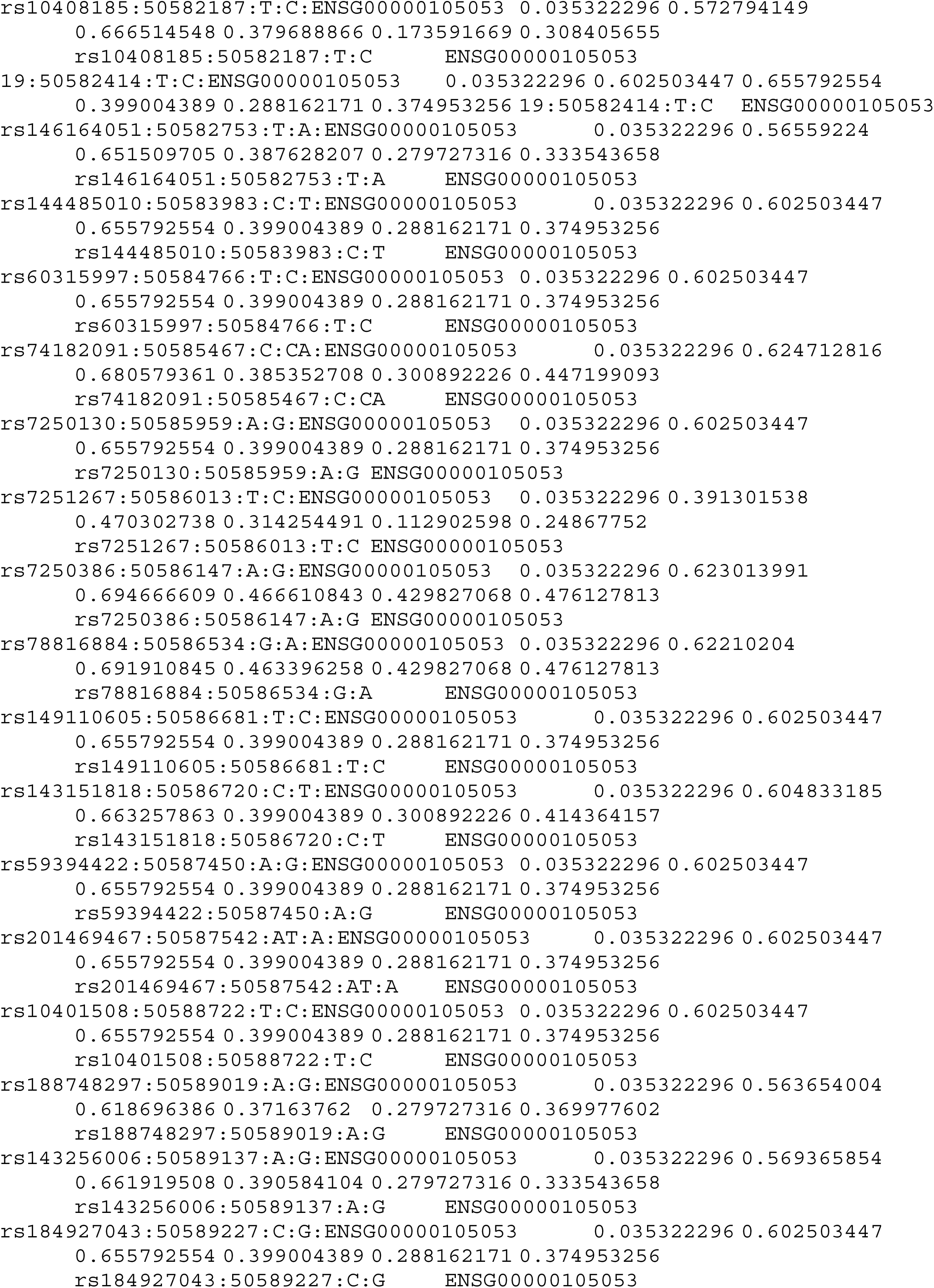

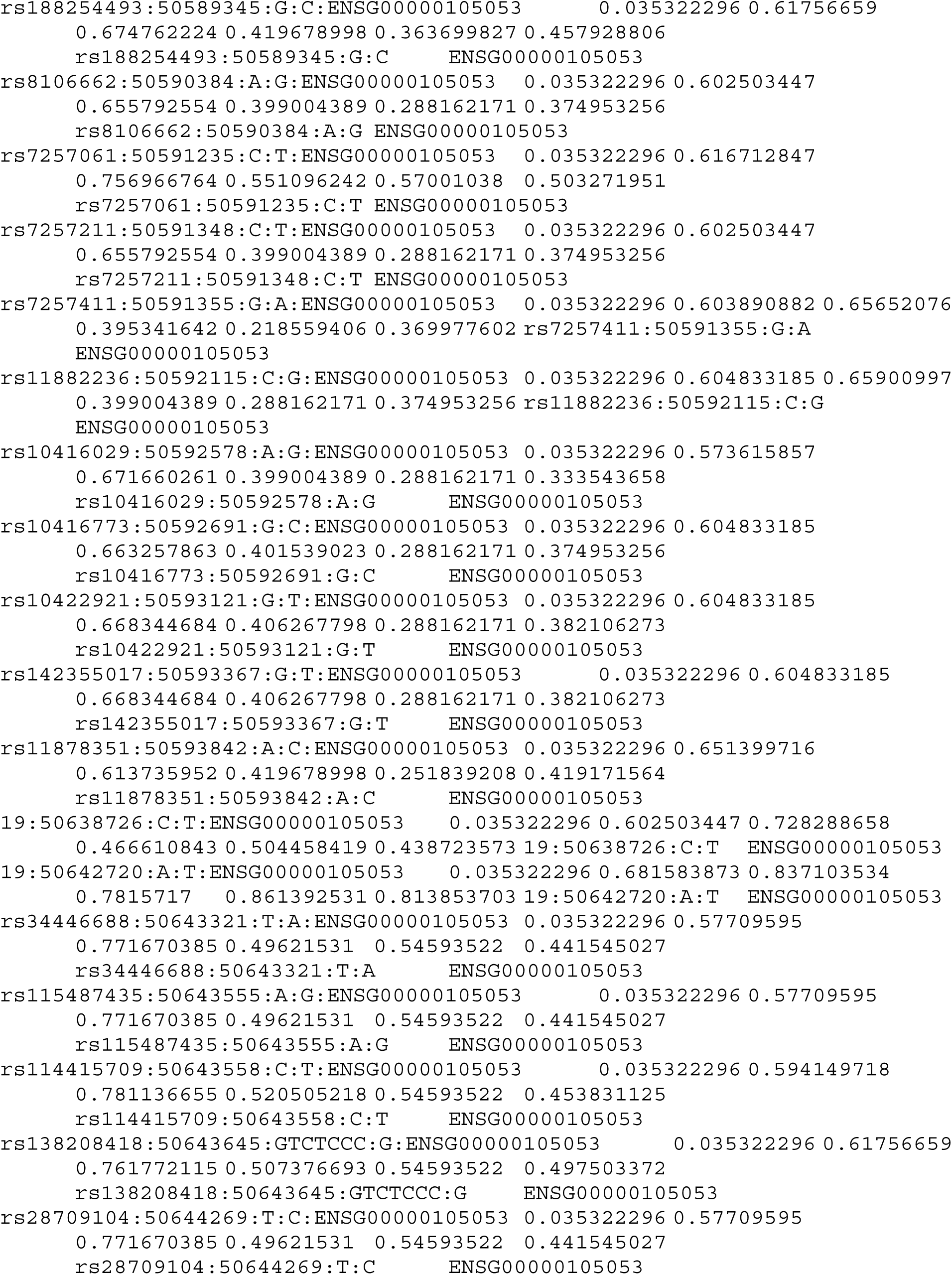

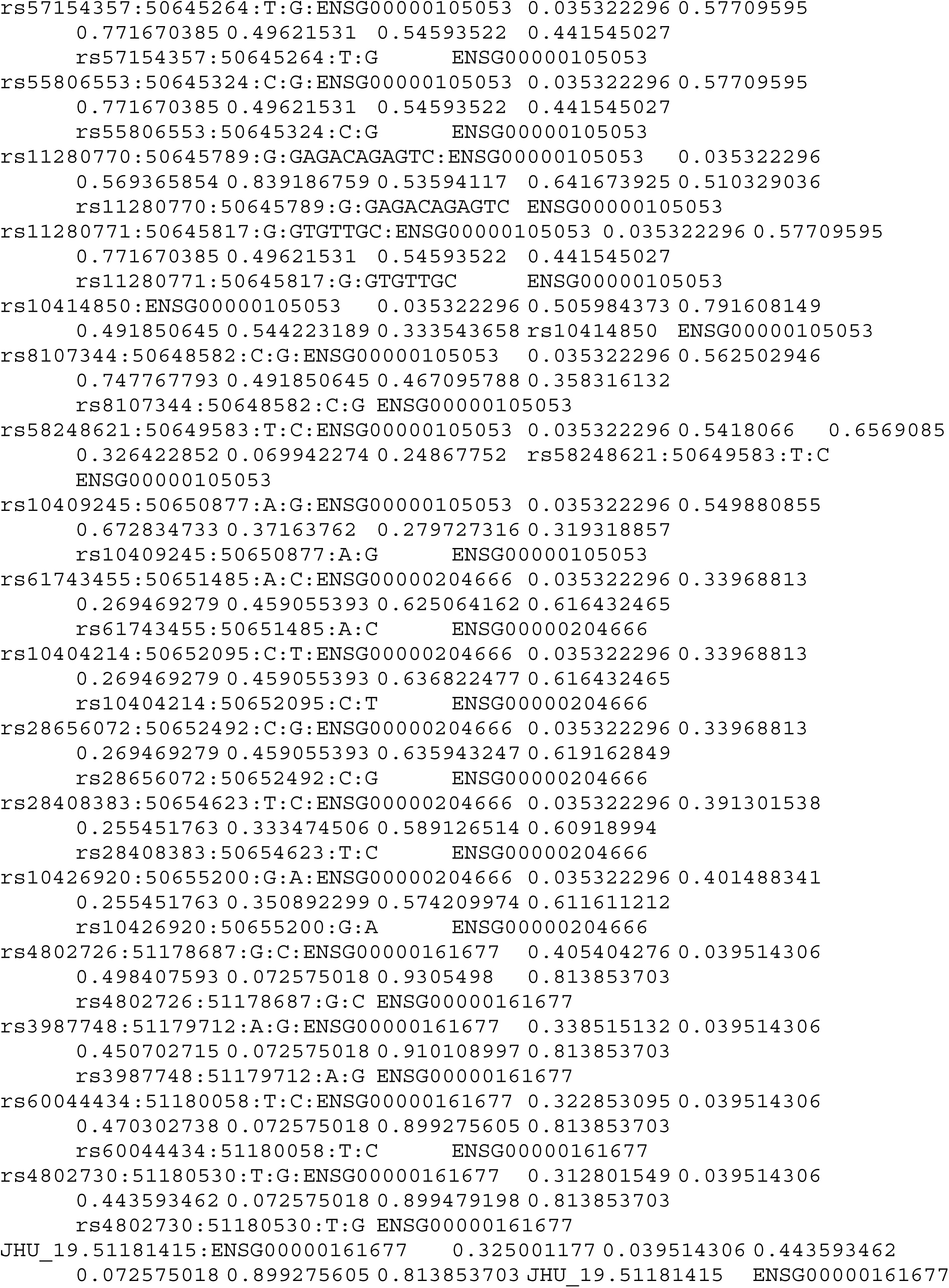

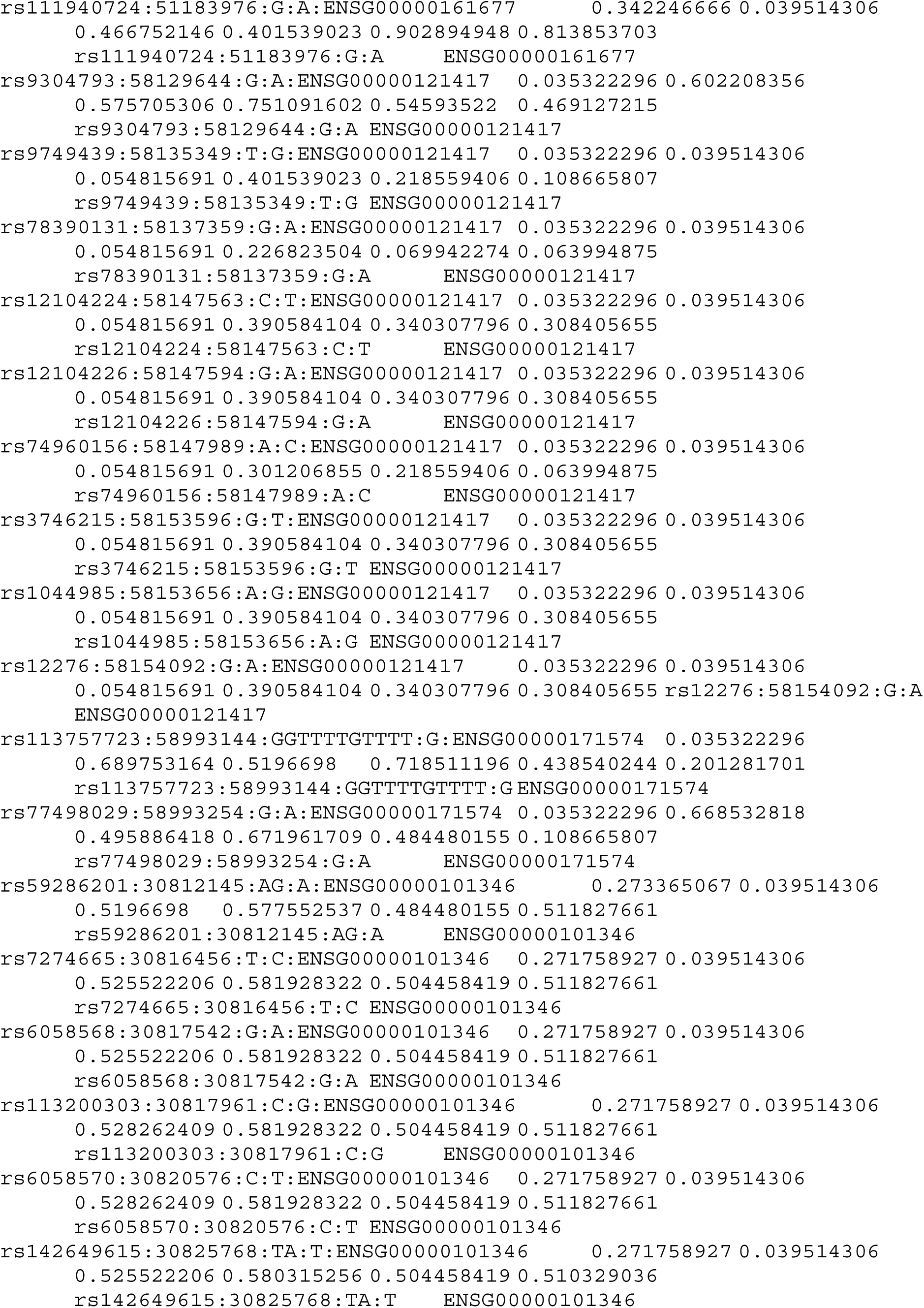

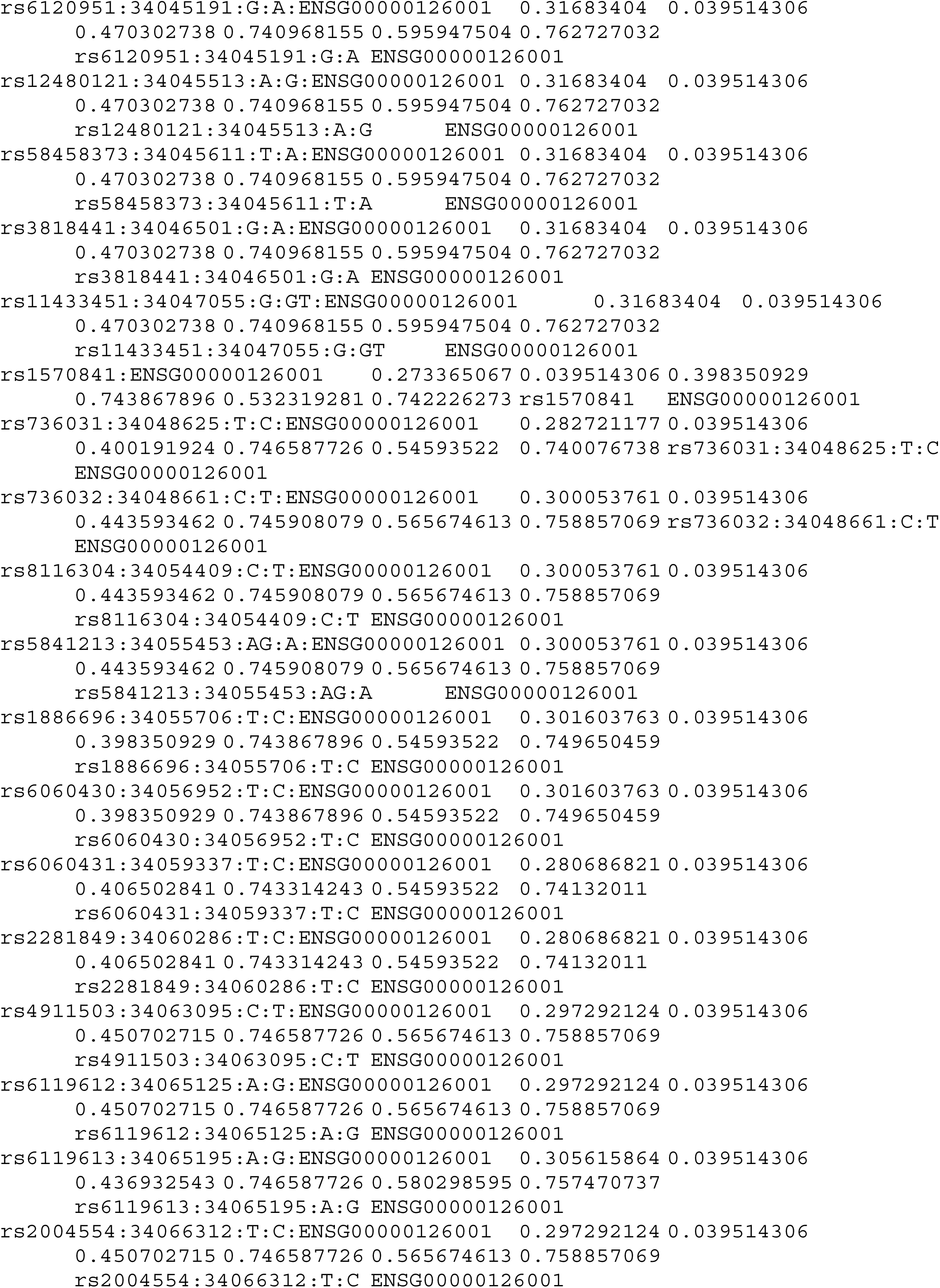

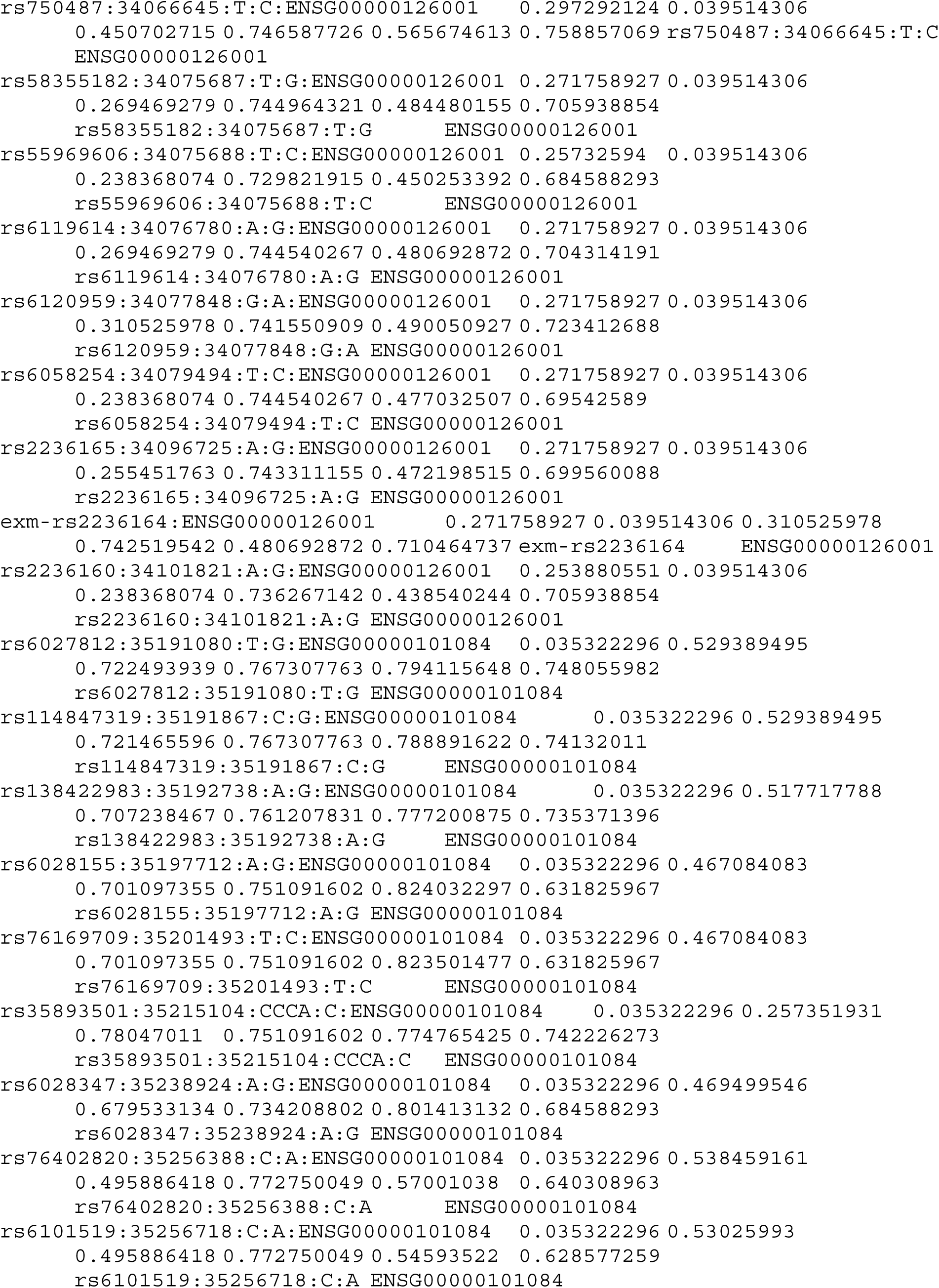

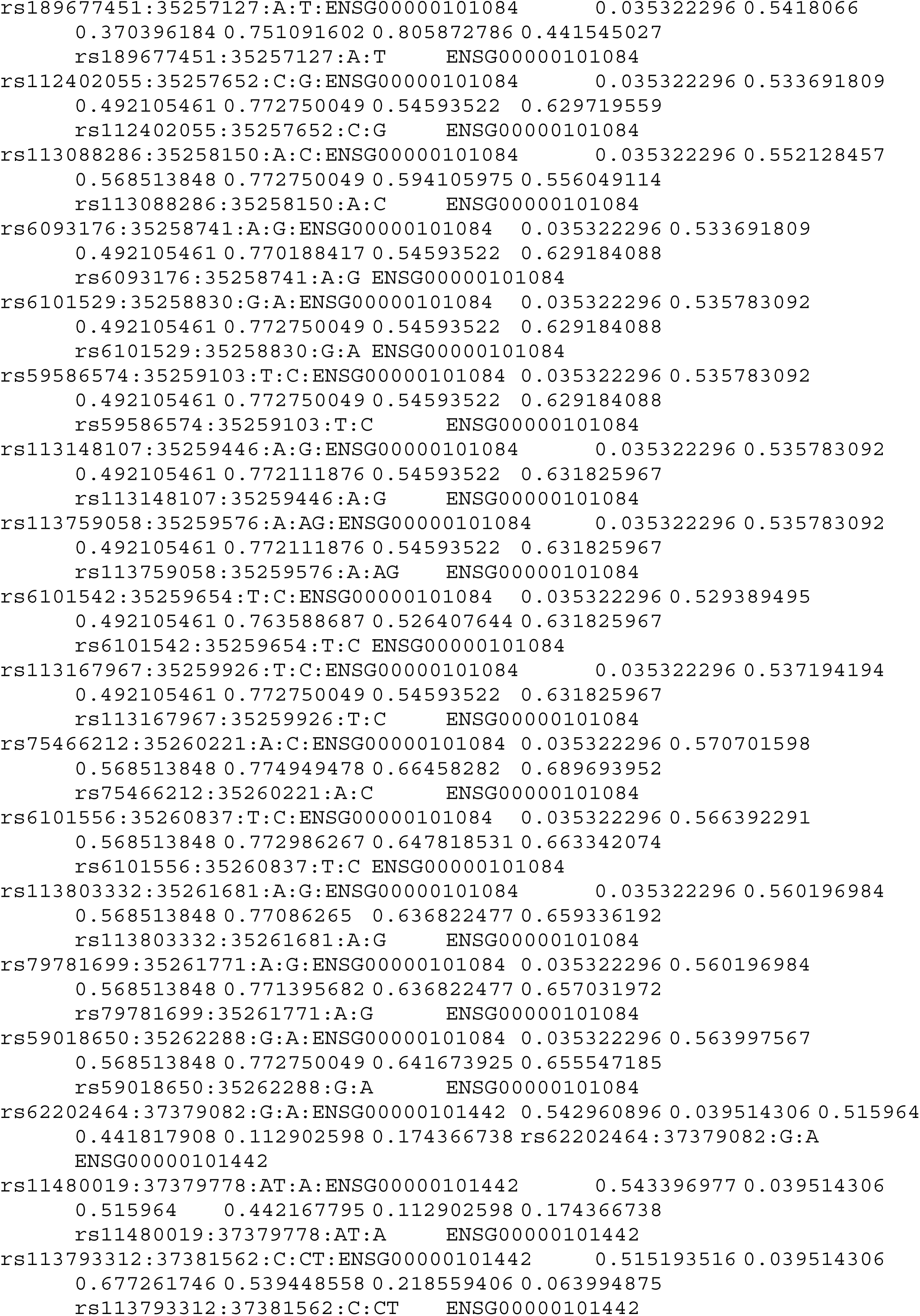

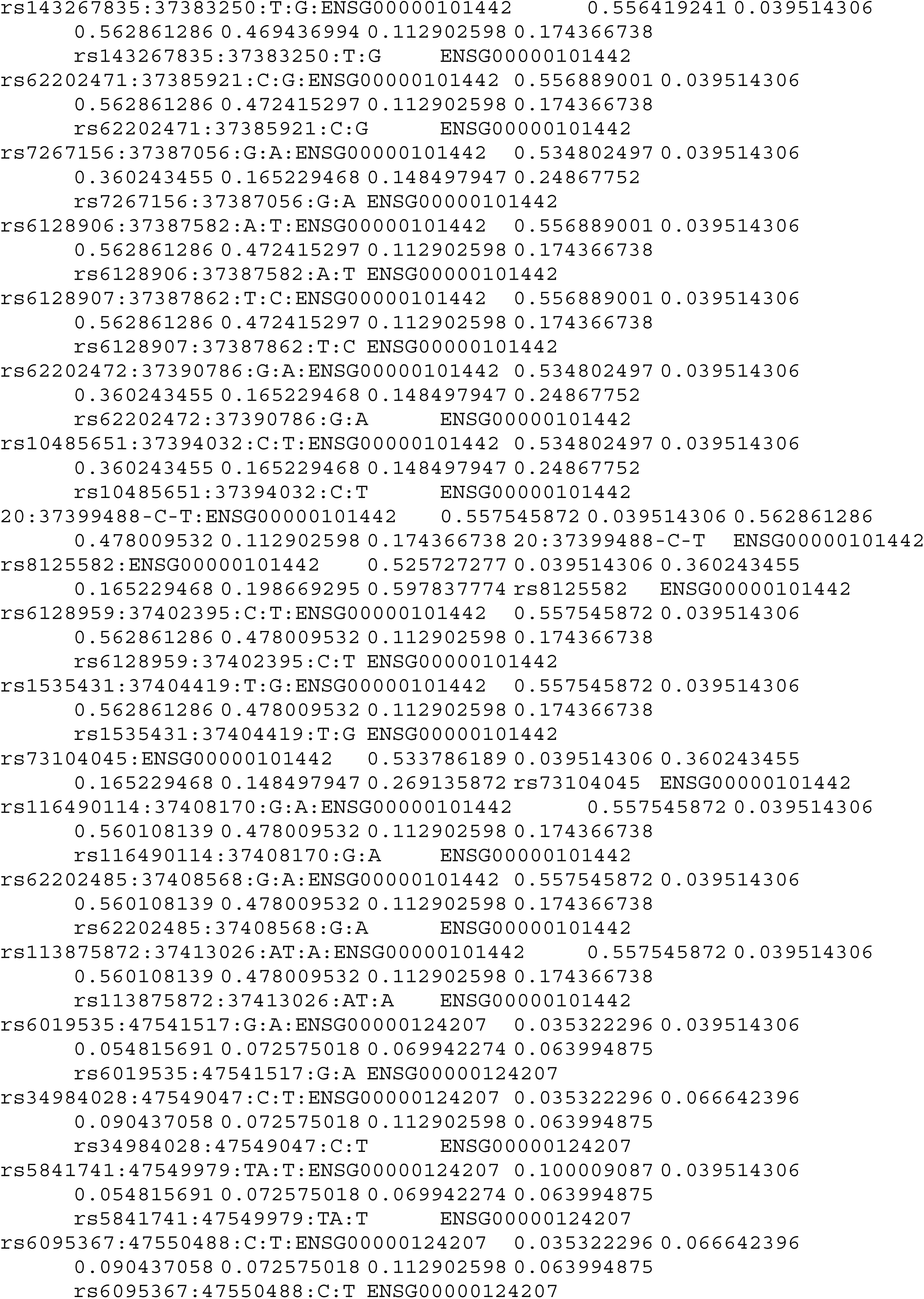

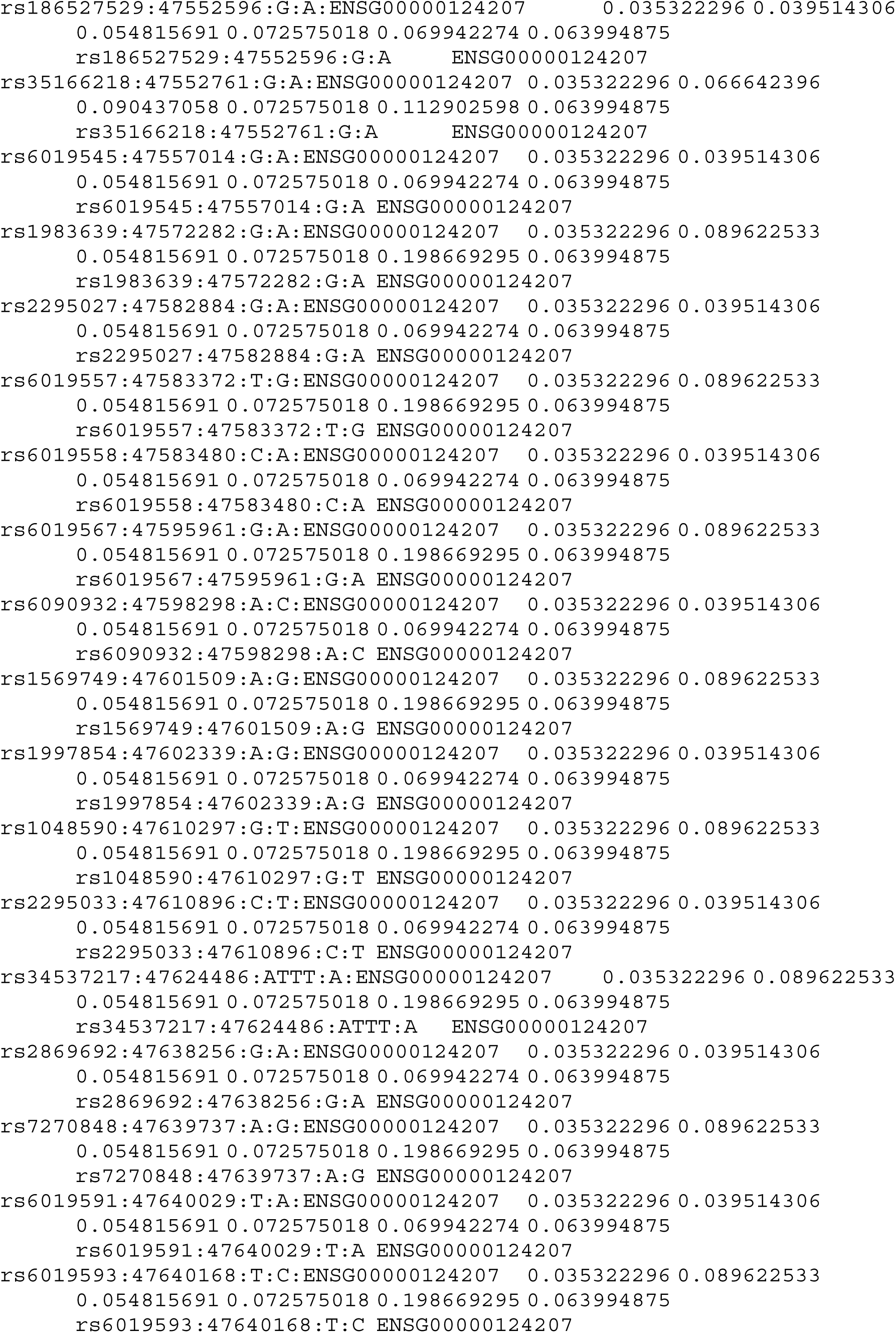

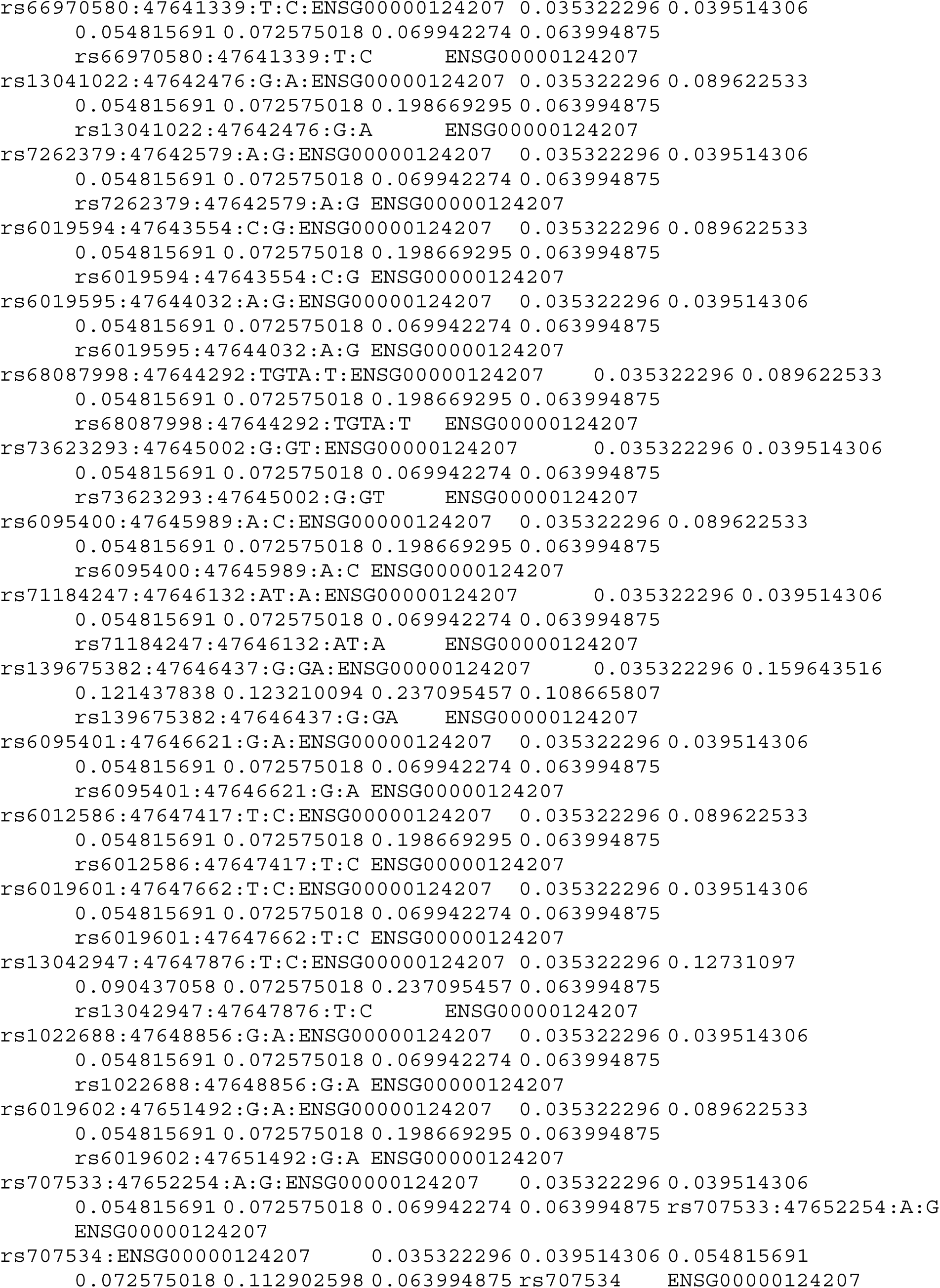

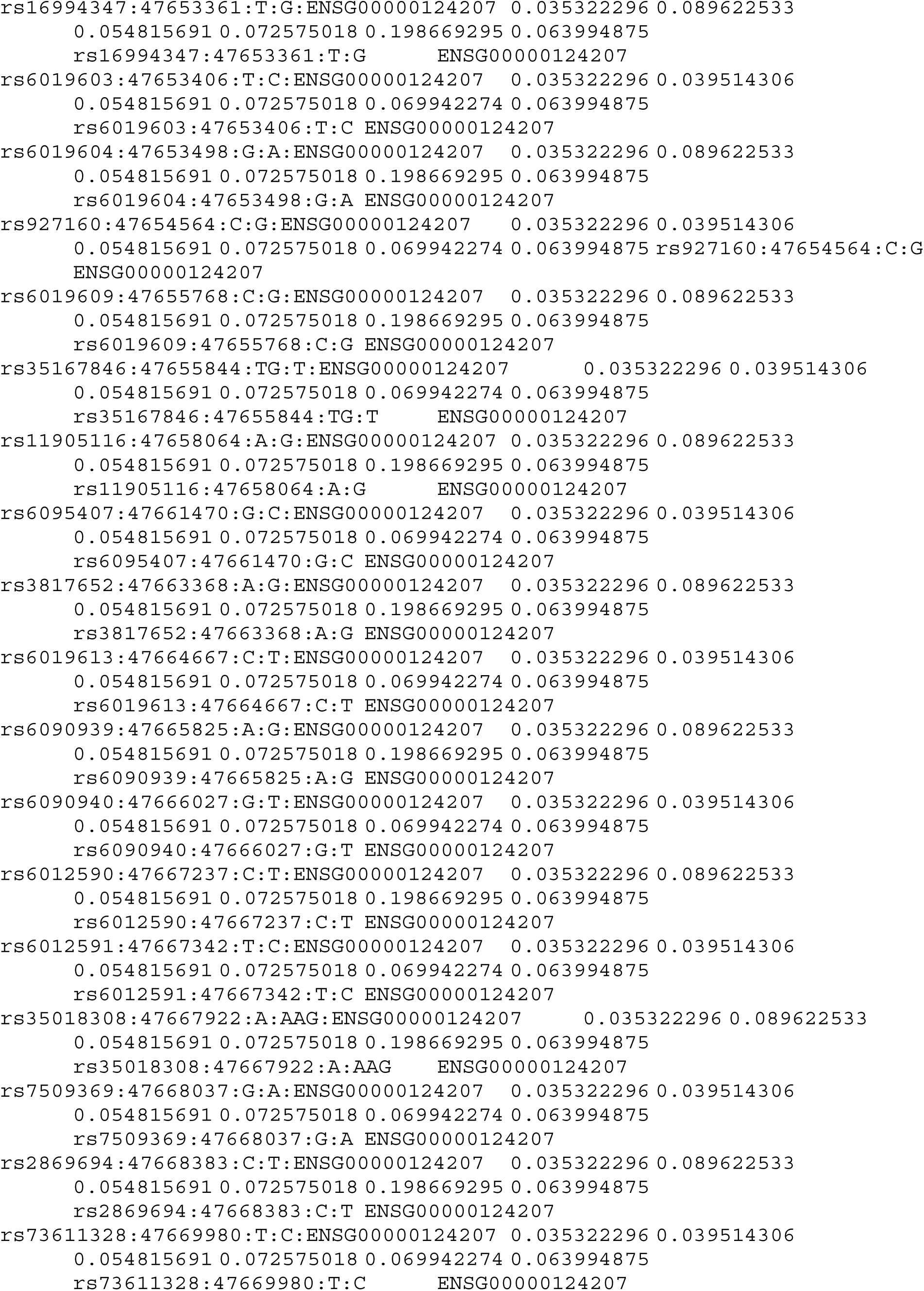

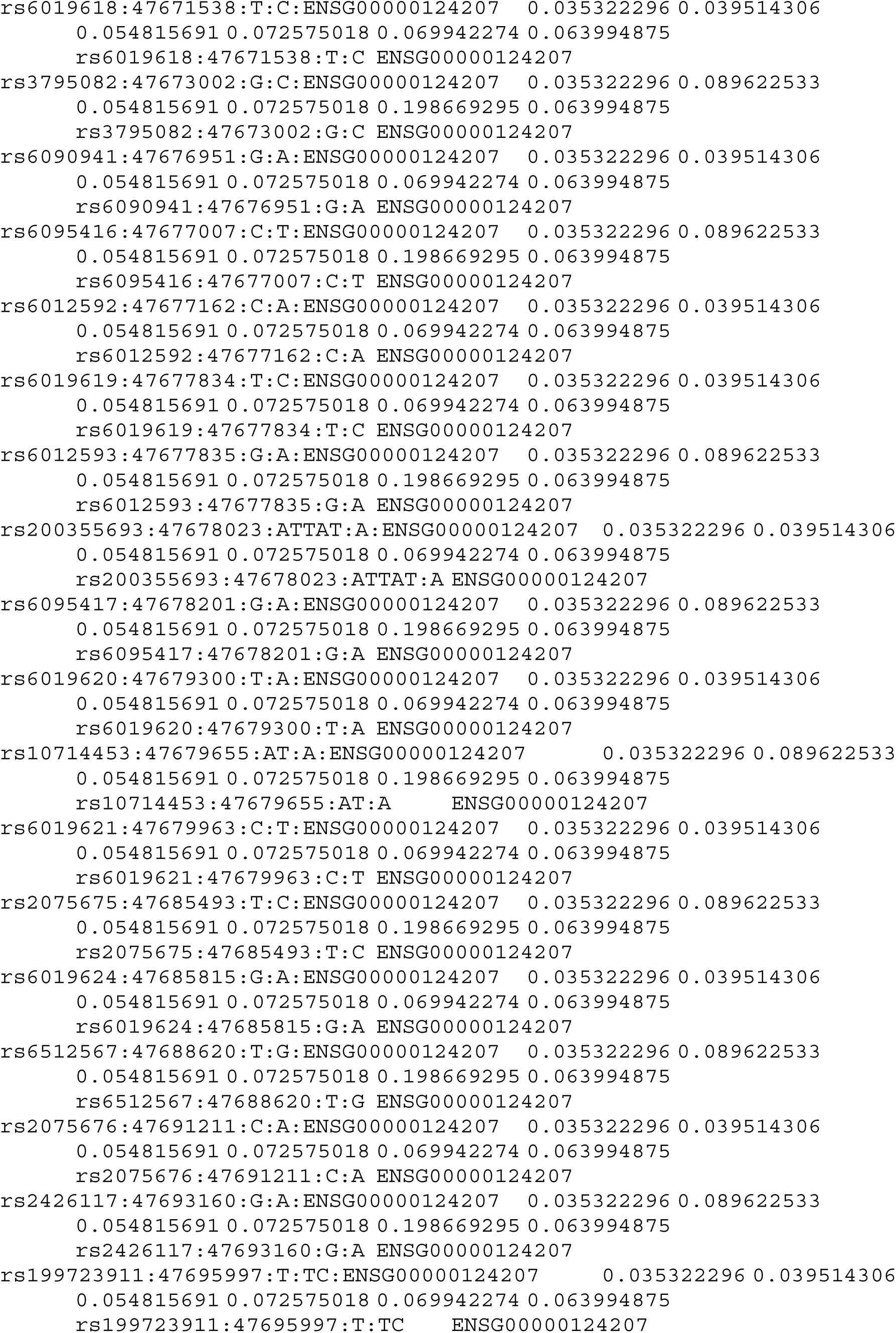

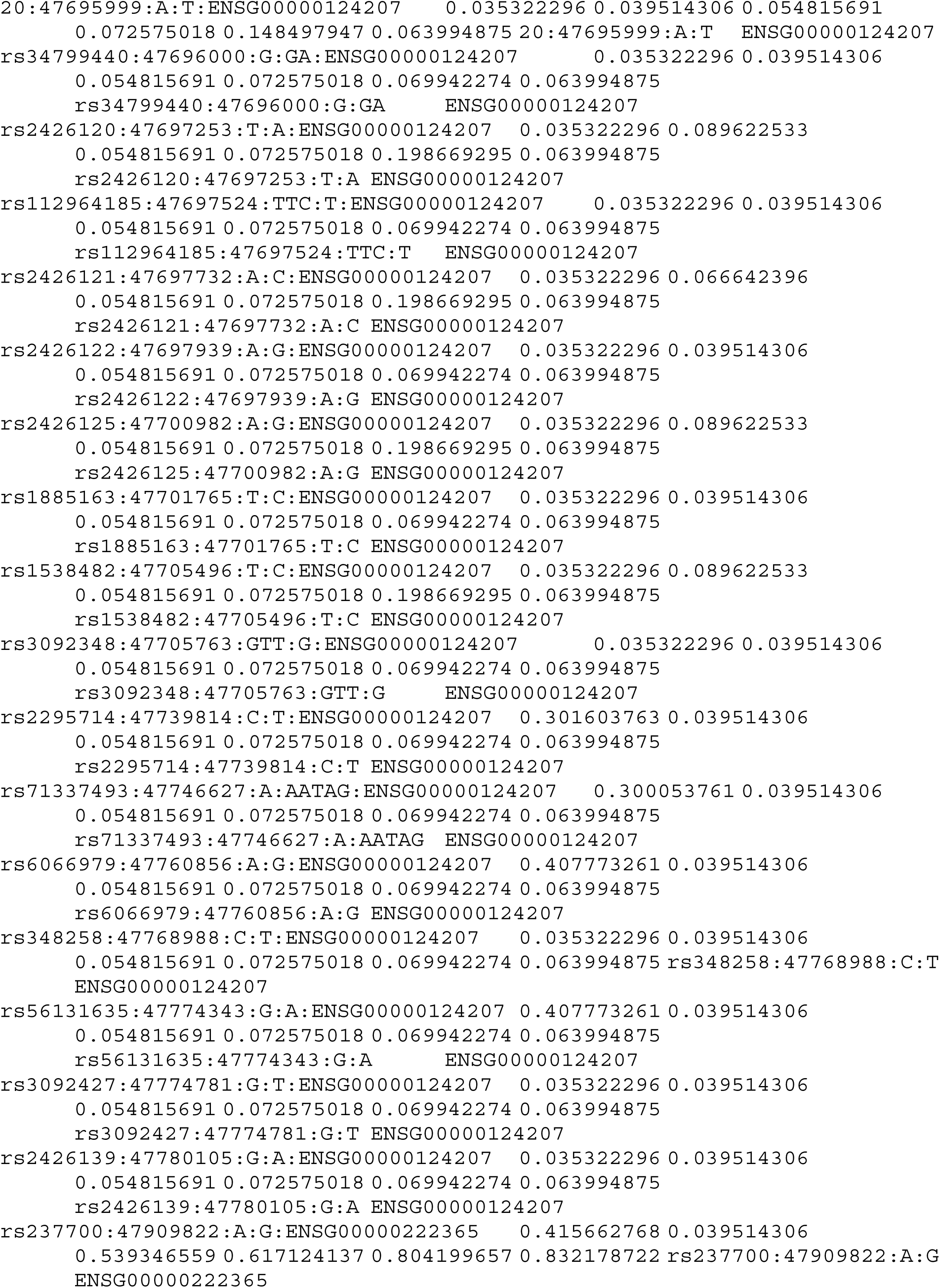

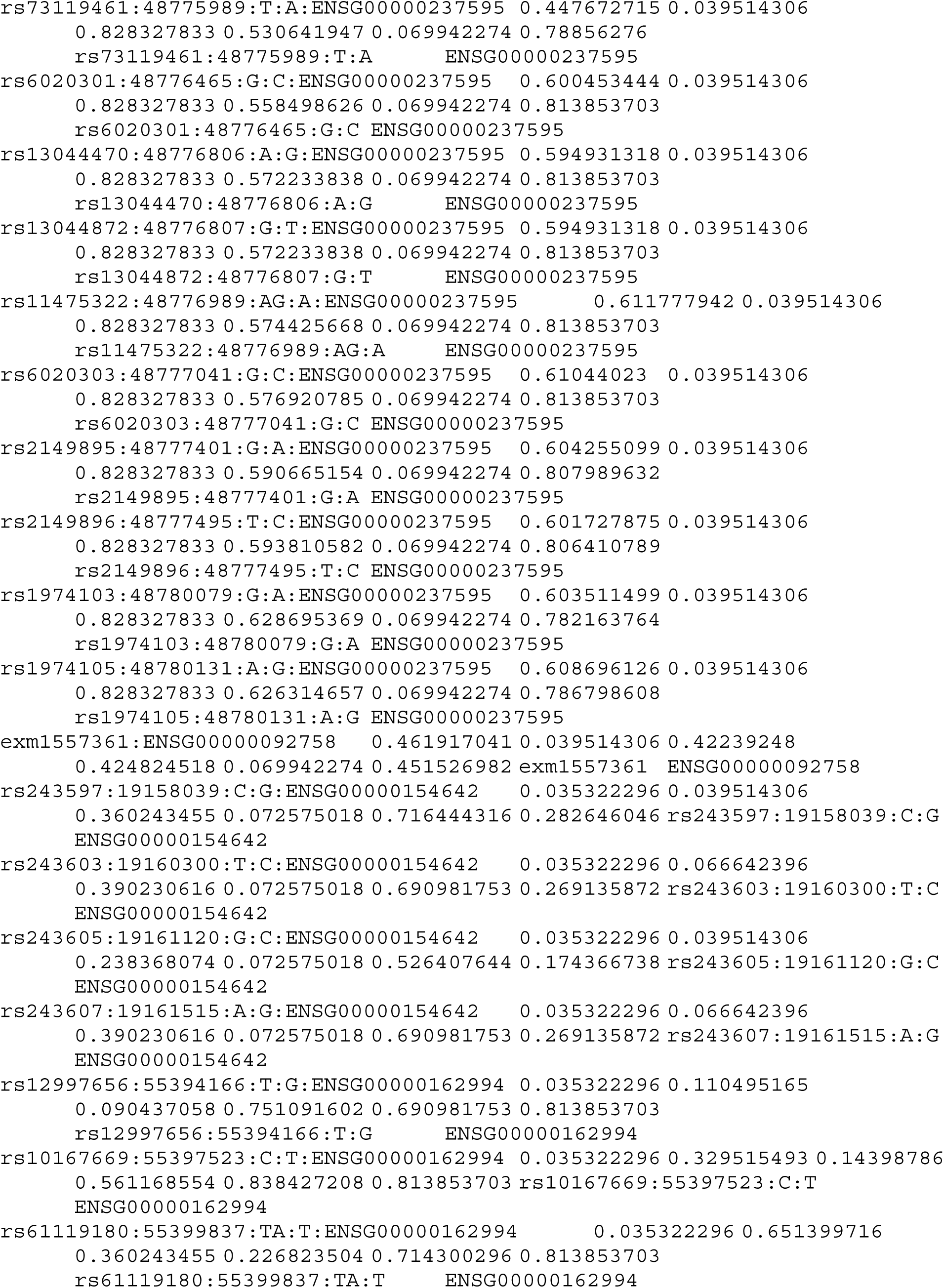

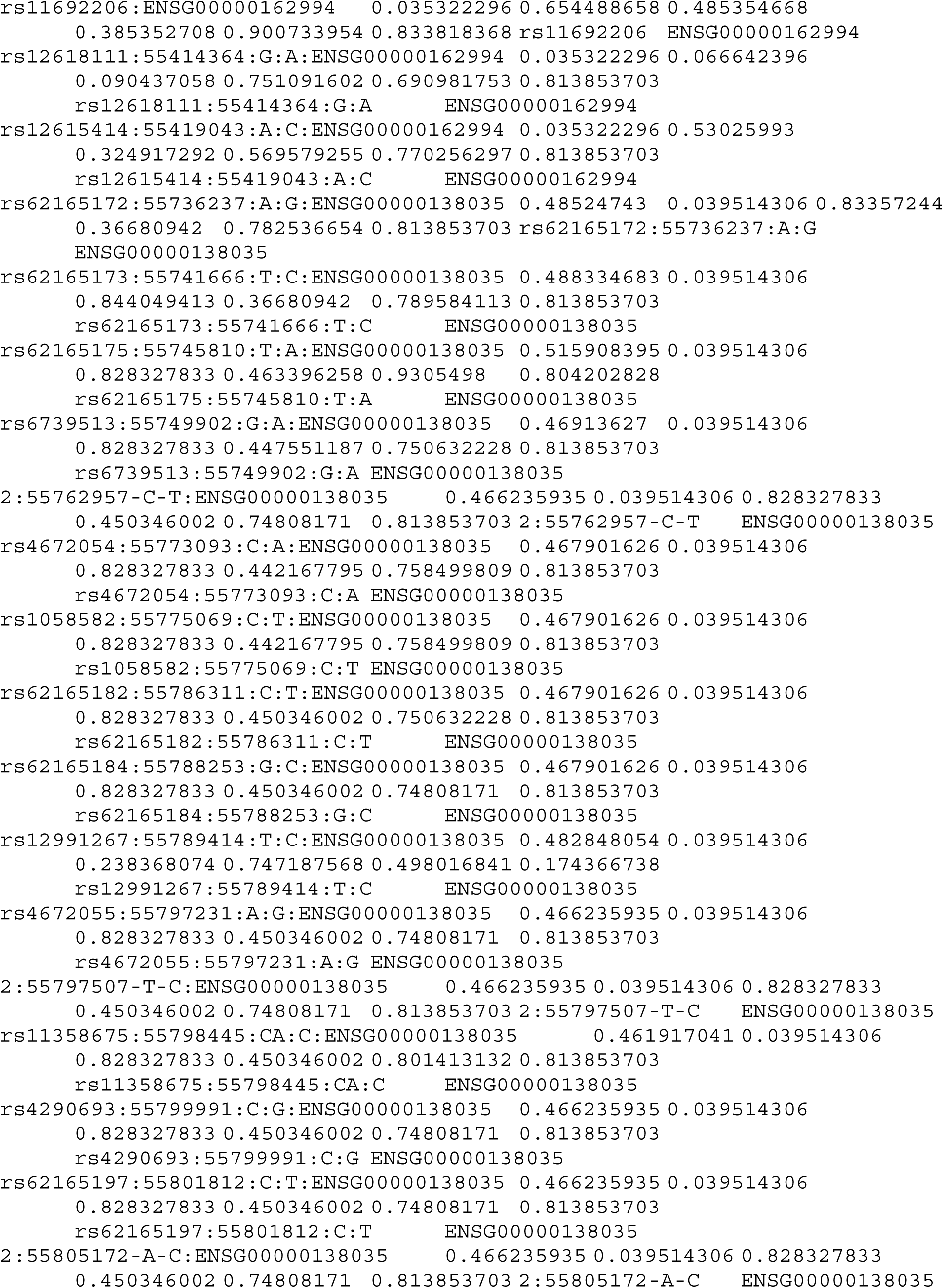

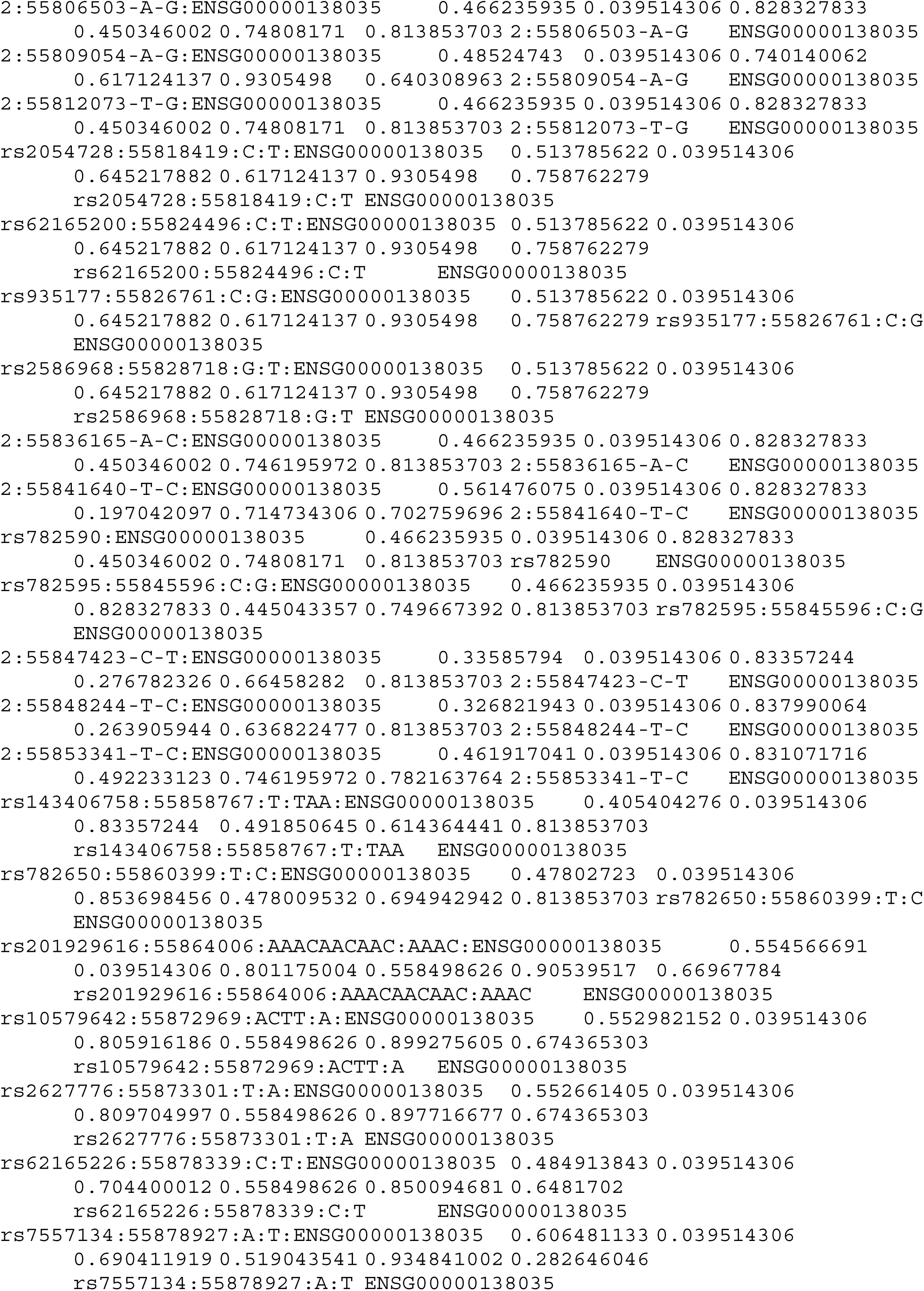

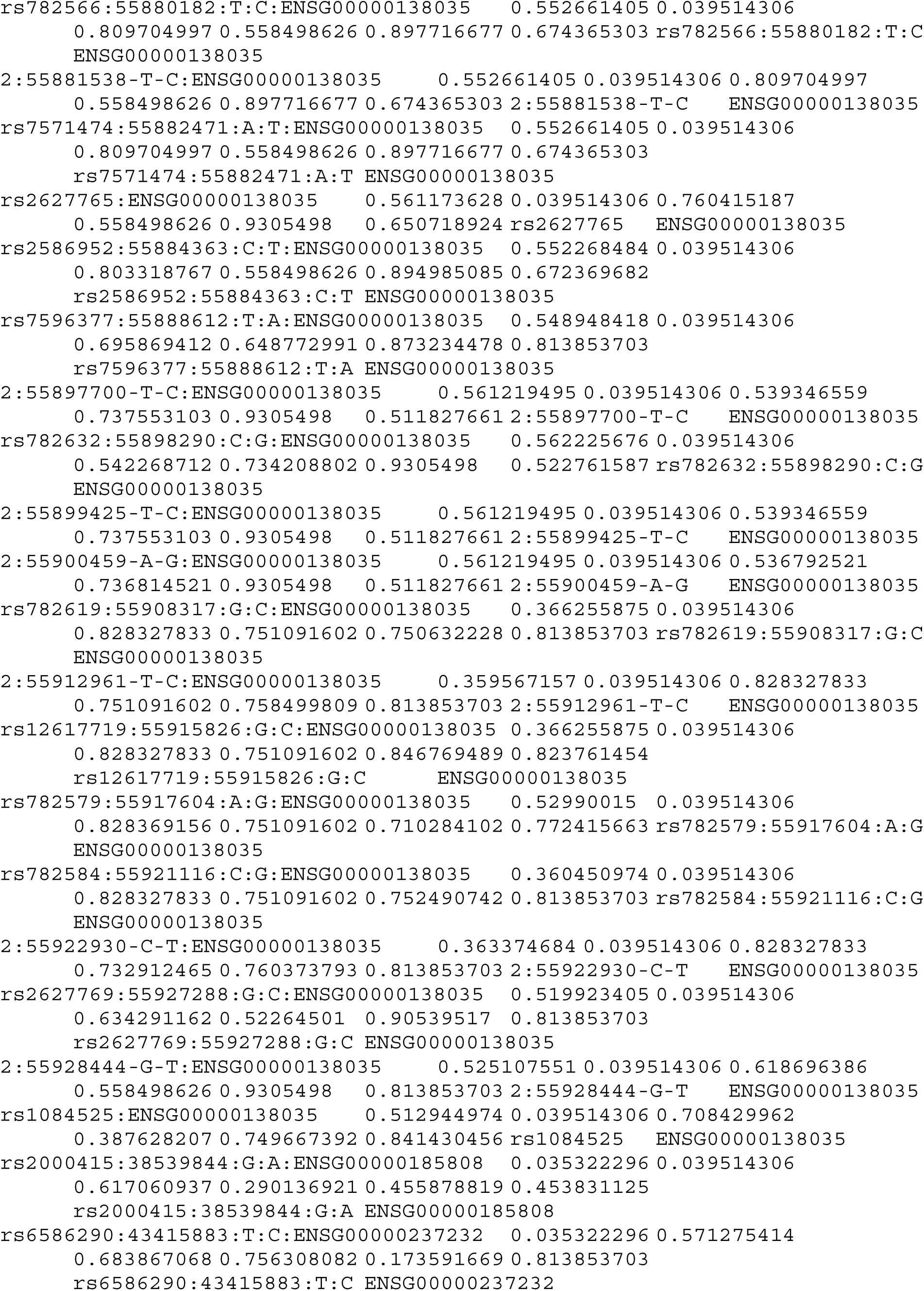

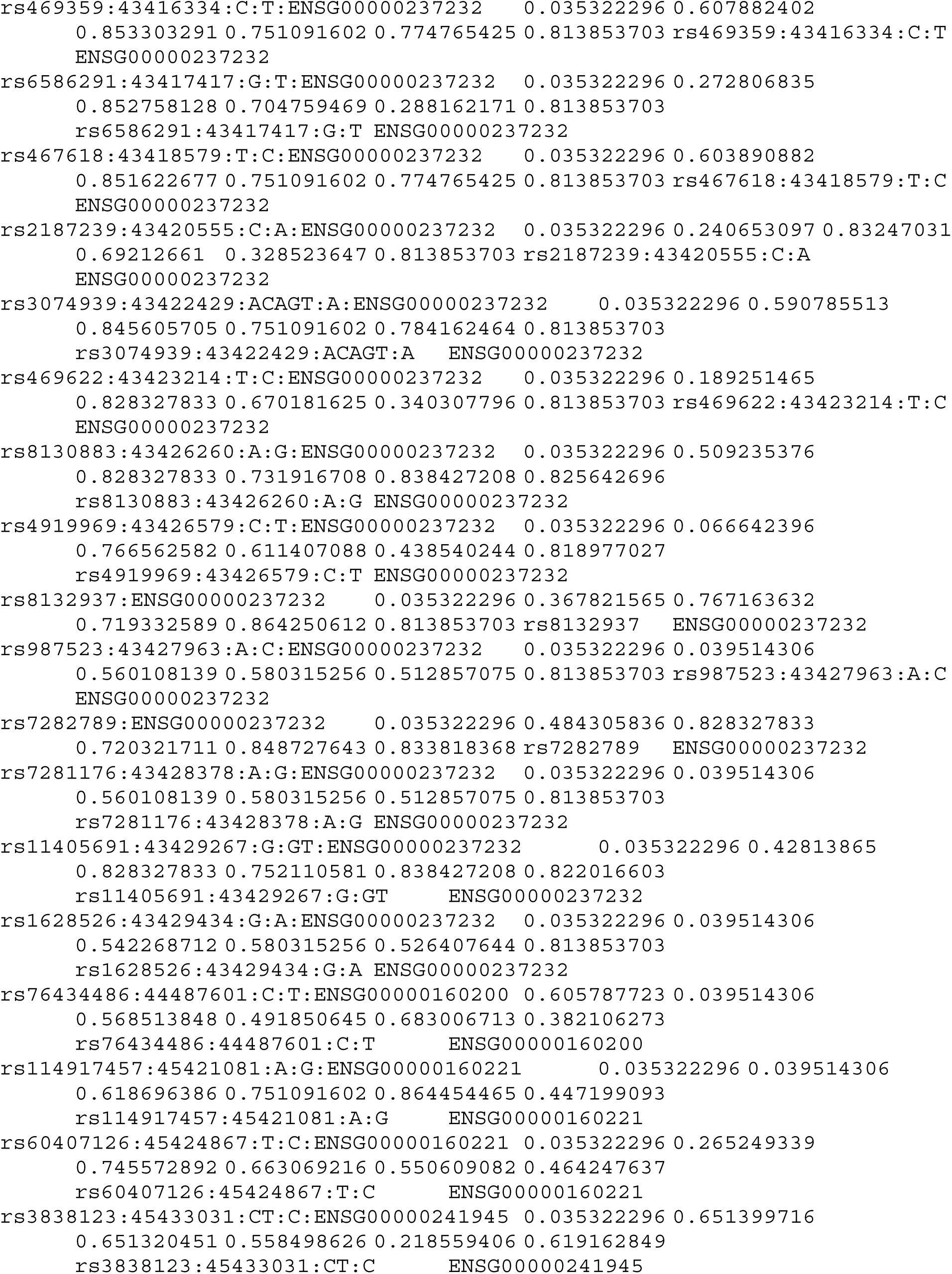

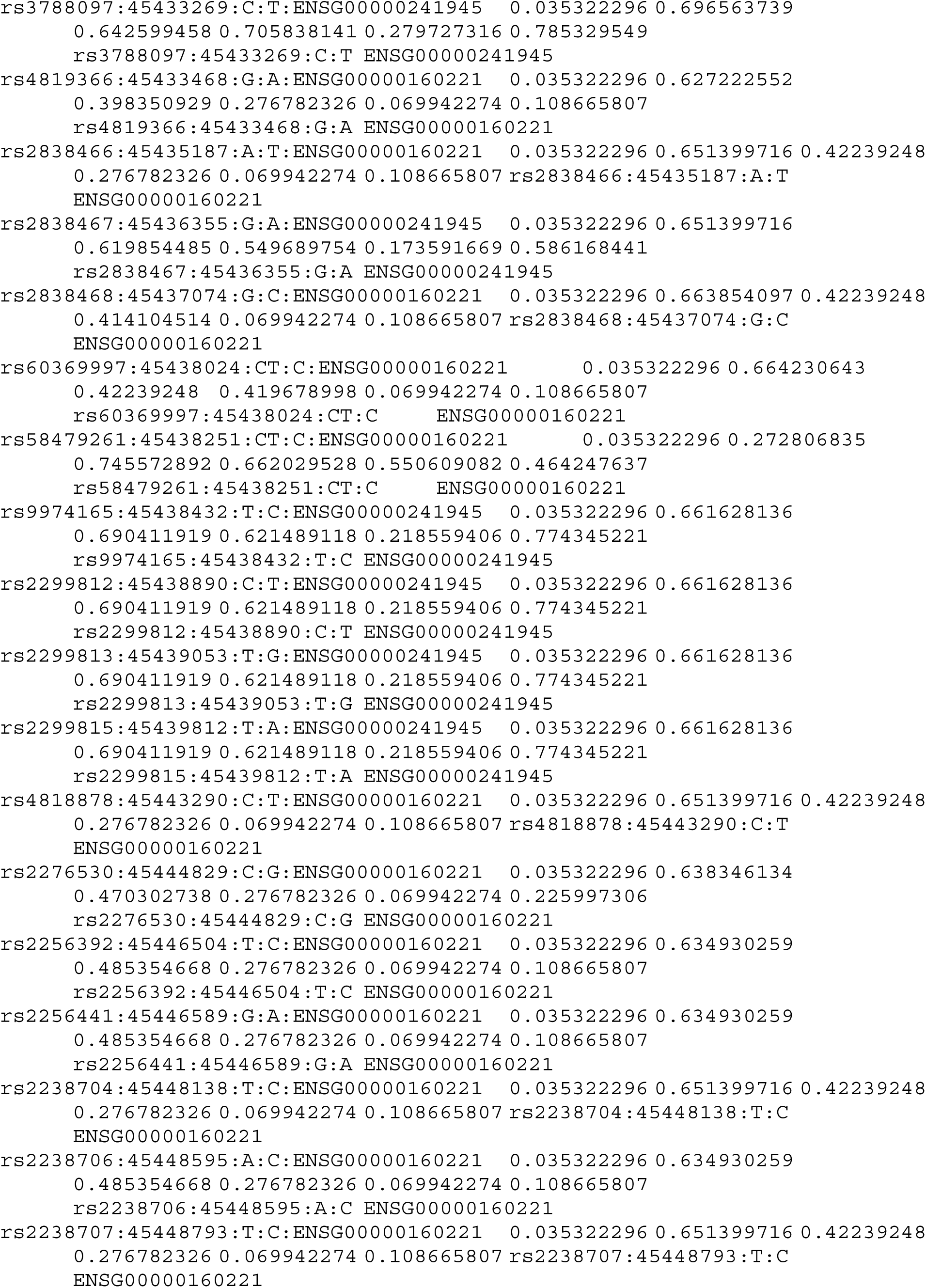

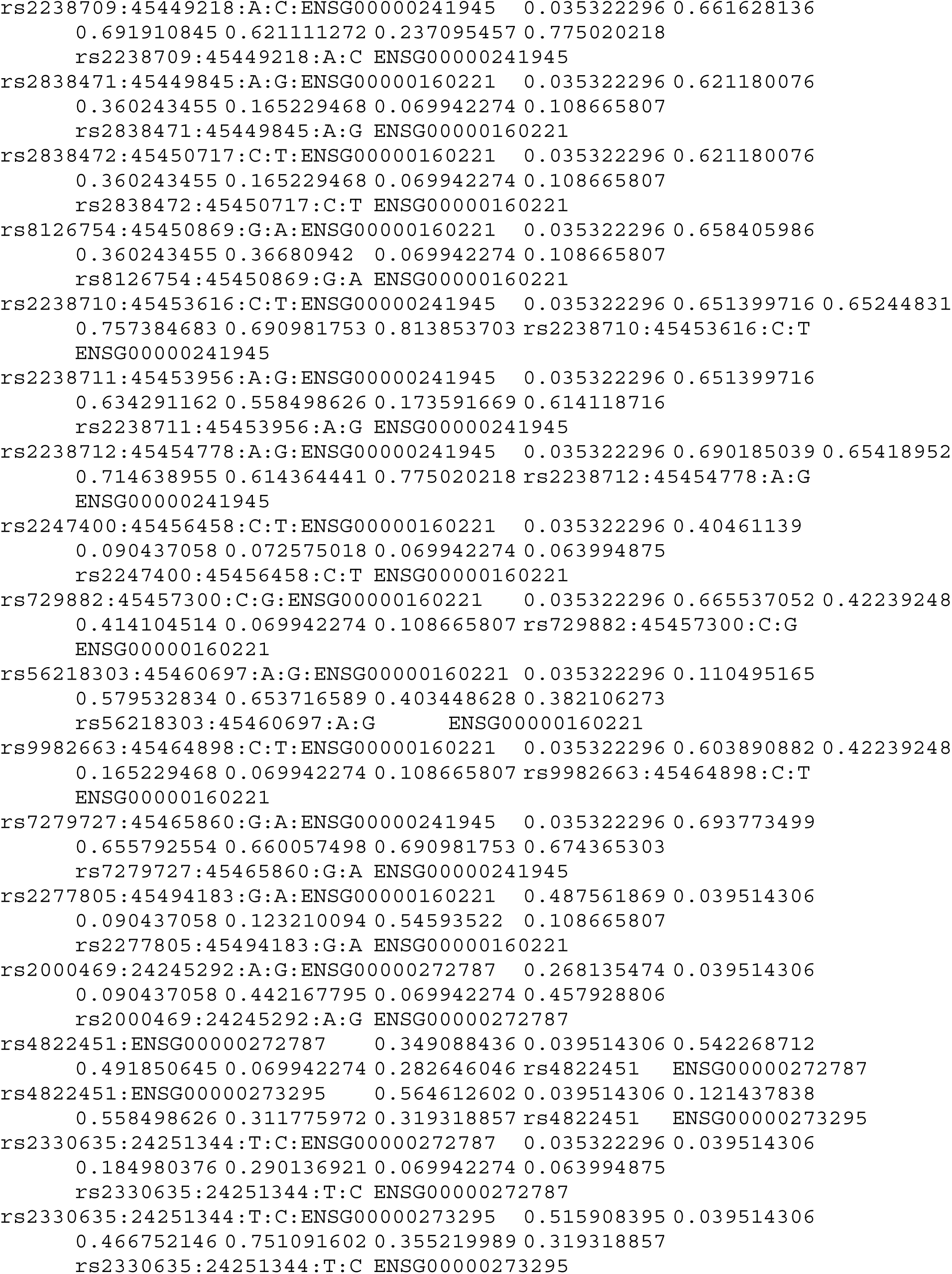

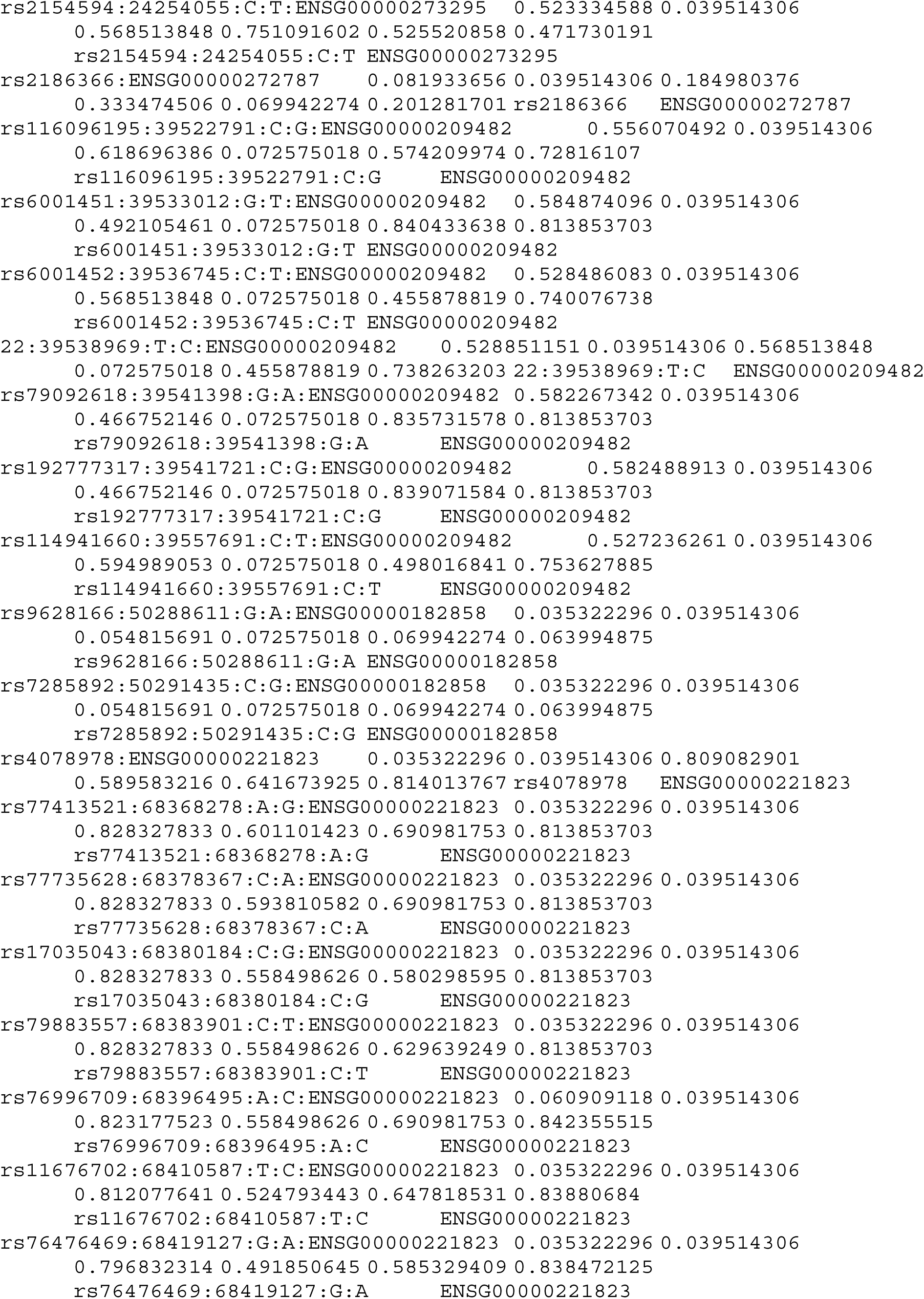

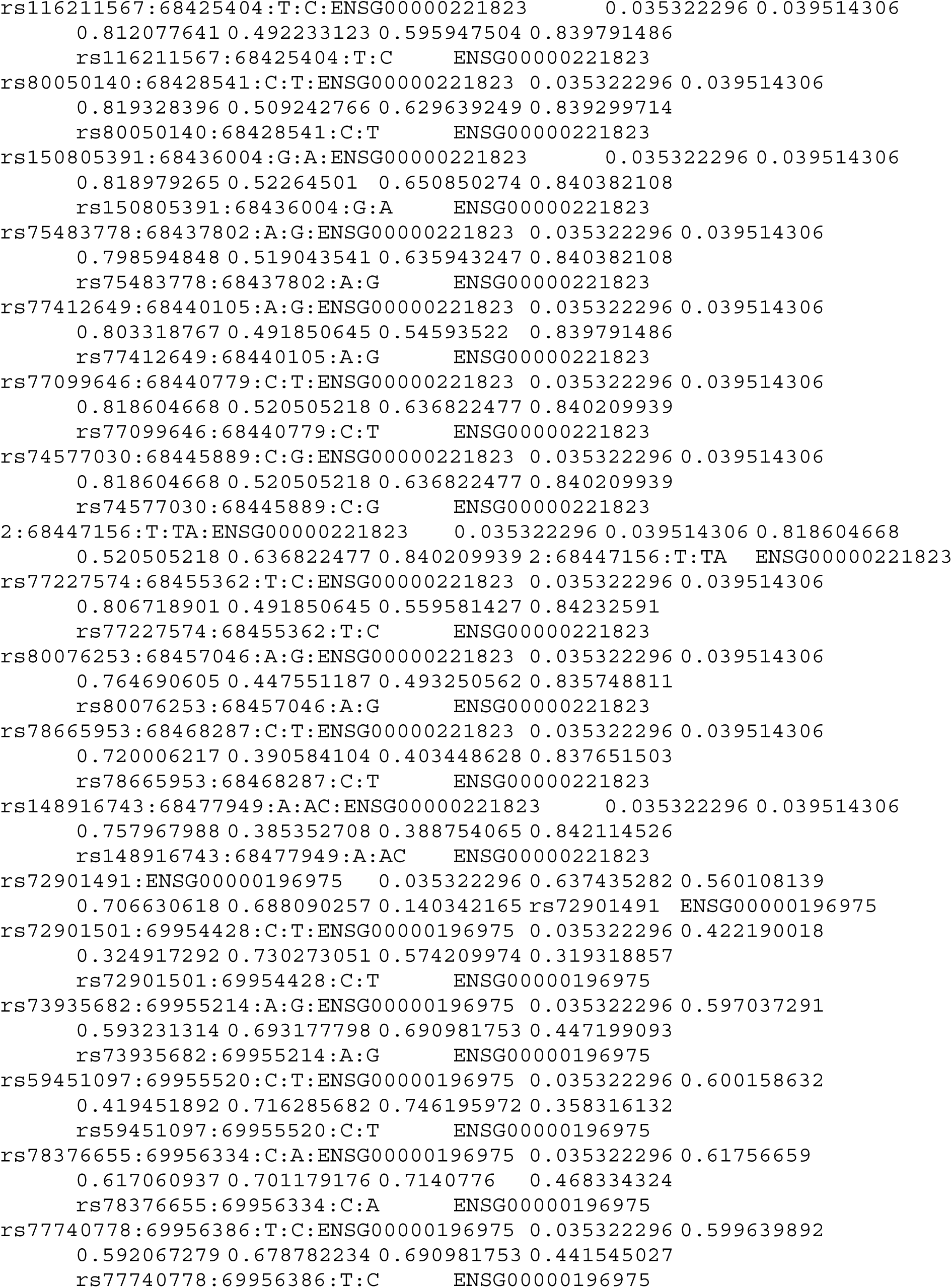

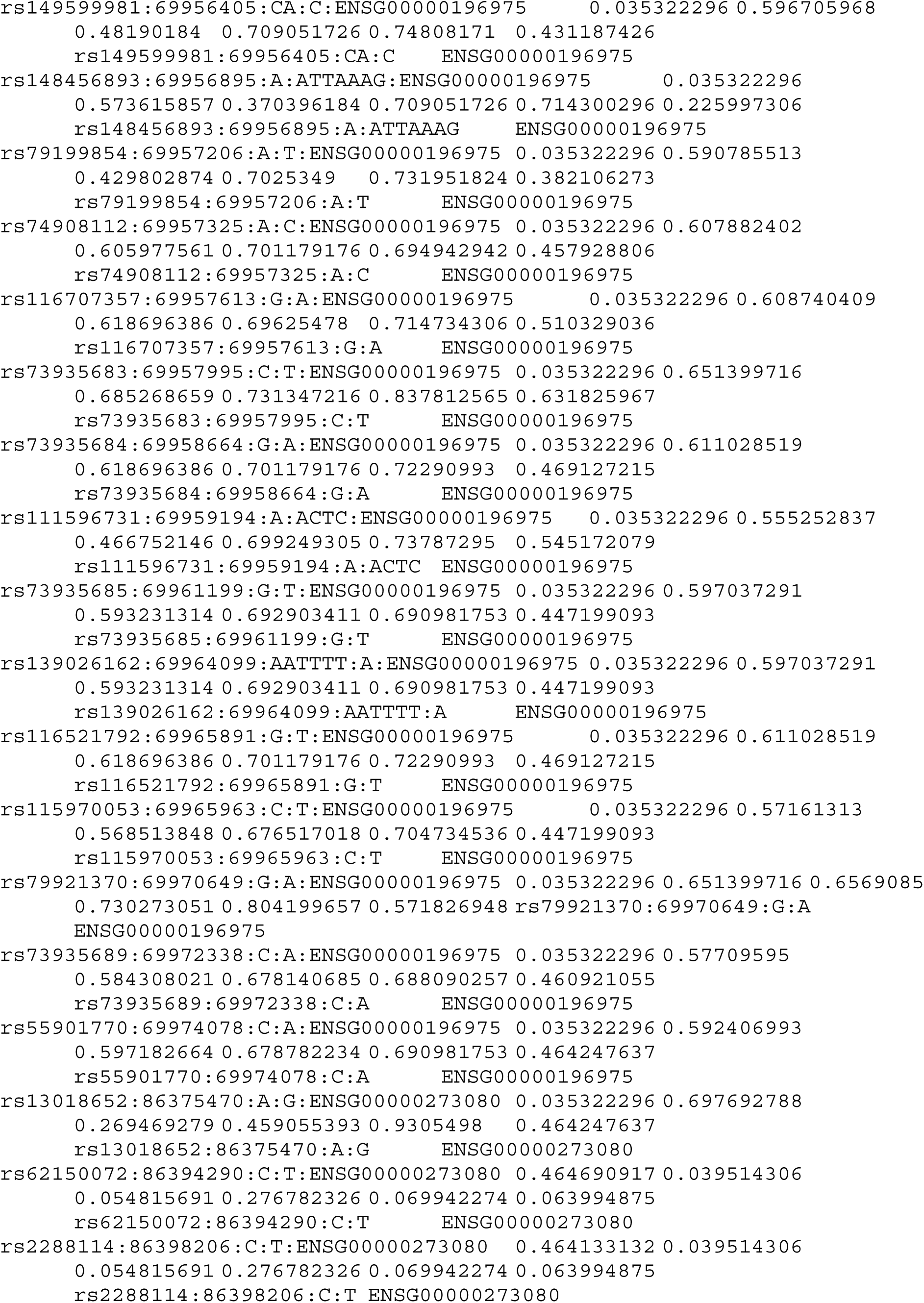

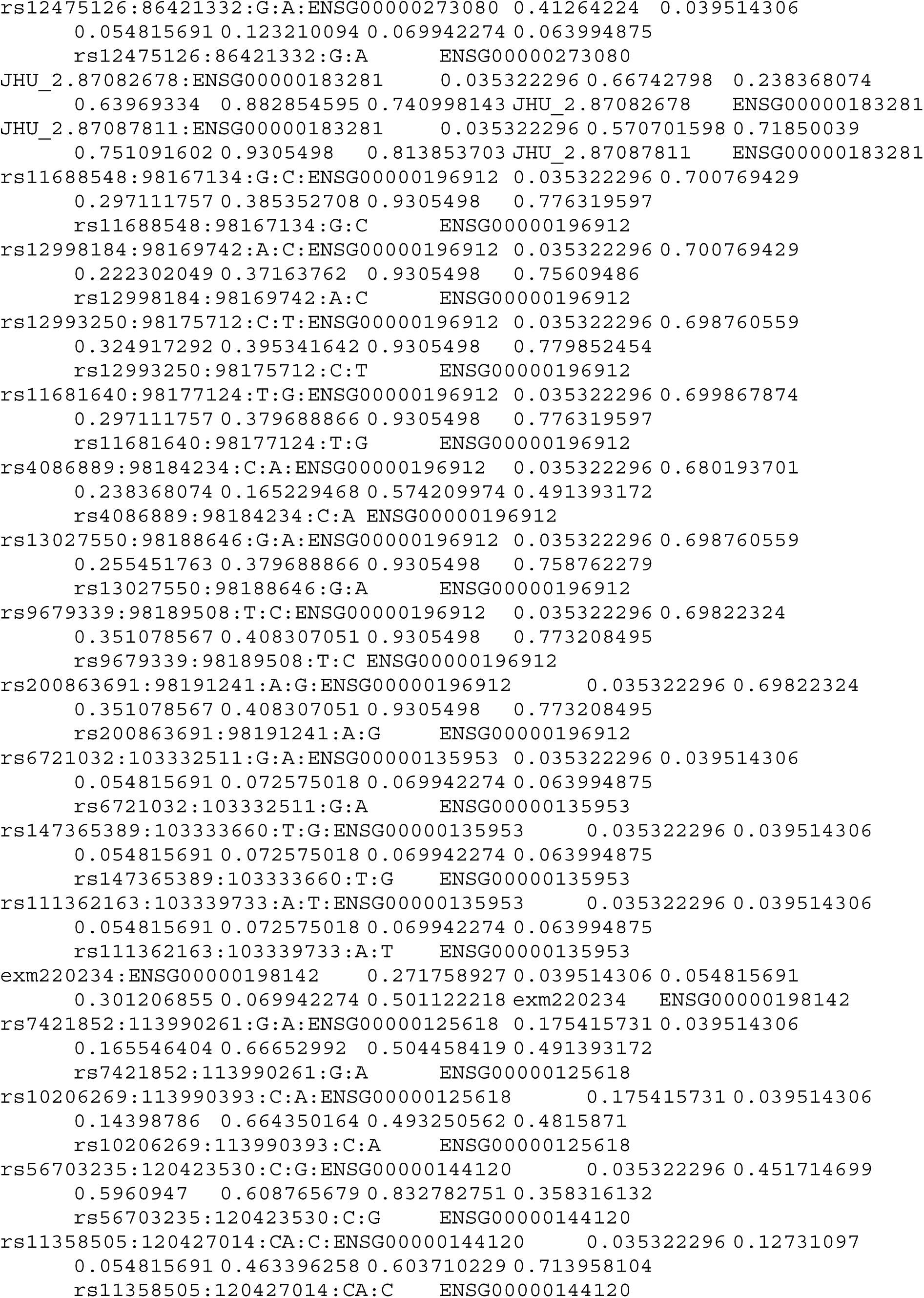

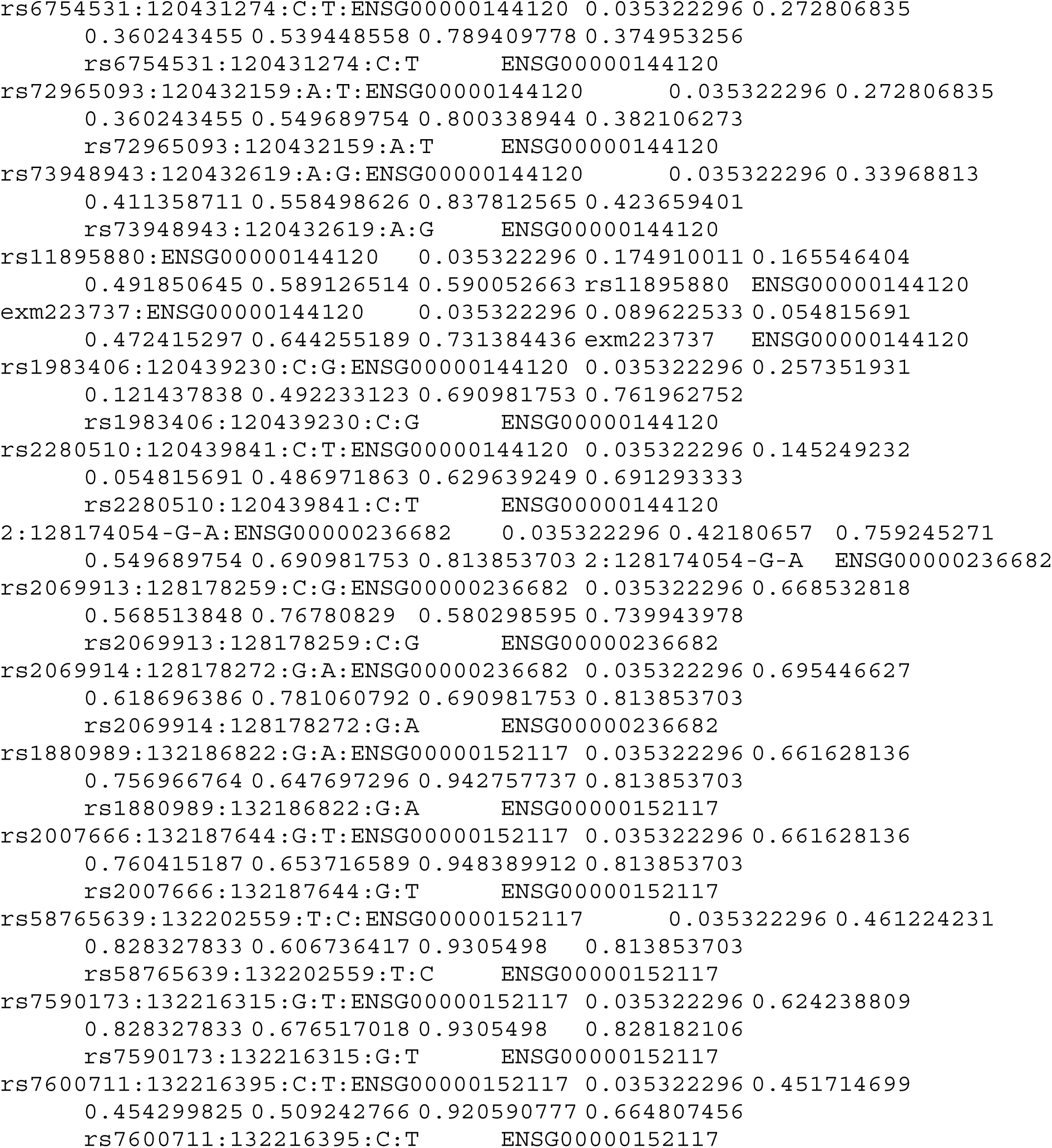

