## Supplementary Figures for "Leverage drug perturbation to reveal genetic regulators of hepatic gene expression in African Americans"

**Supplemental Figures**

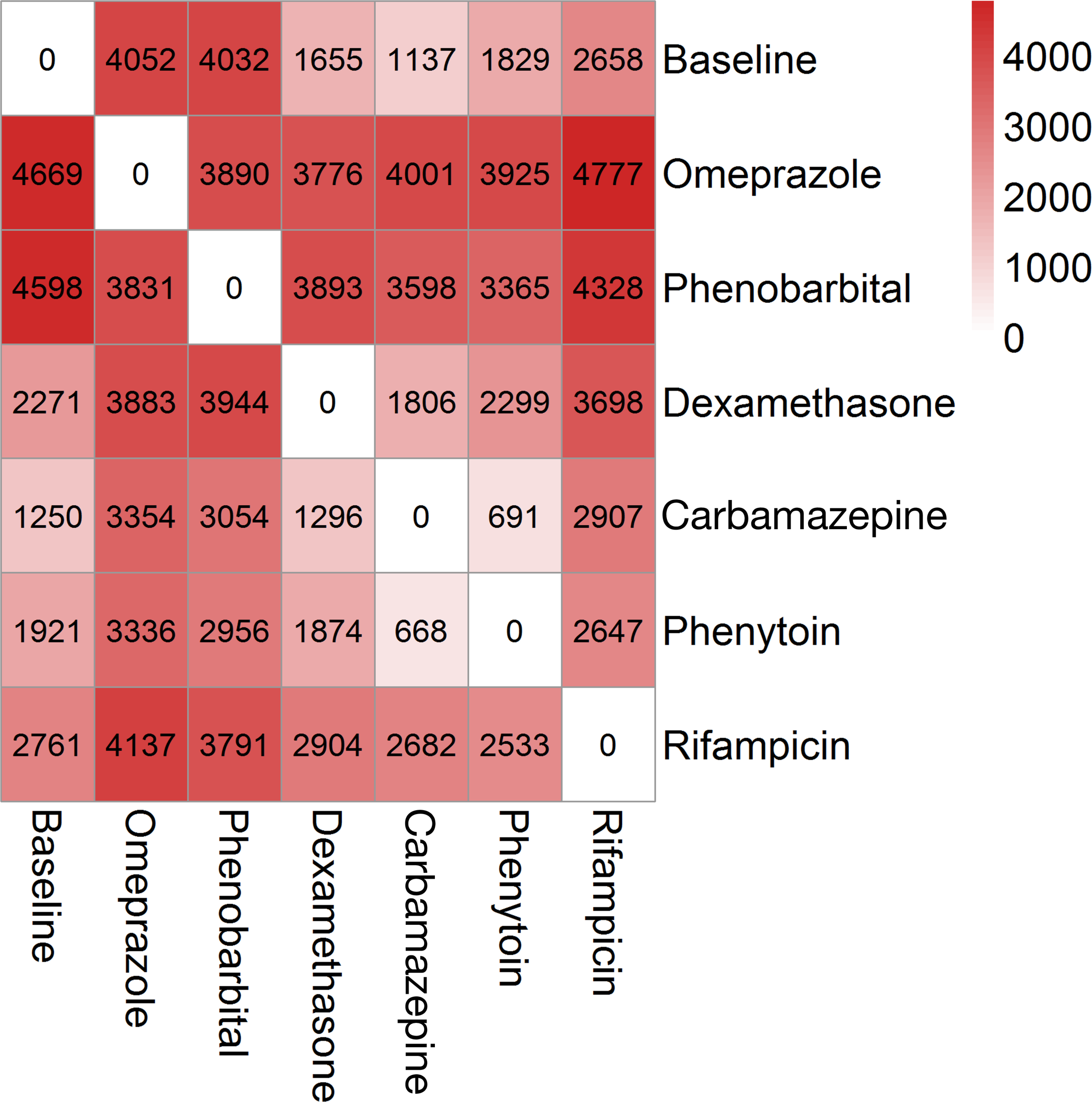

**Supplemental Figure 1: Differentially expressed gene between drug conditions.**

Number of DE genes with FDR<0.01. The values above the diagonal show the number of up-regulated genes and the values below the diagonal show the number of down-regulated genes.

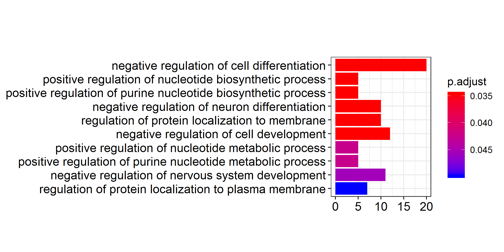

**Supplemental Figure 2: Enrichment of commonly downregulated genes**

Pathways represented by commonly downregulated genes (genes that were significantly downregulated in all conditions at FDR<0.01) The x-axis shows the number of genes in each ontology category with the color scale representing the adjusted p-value of the enrichment.

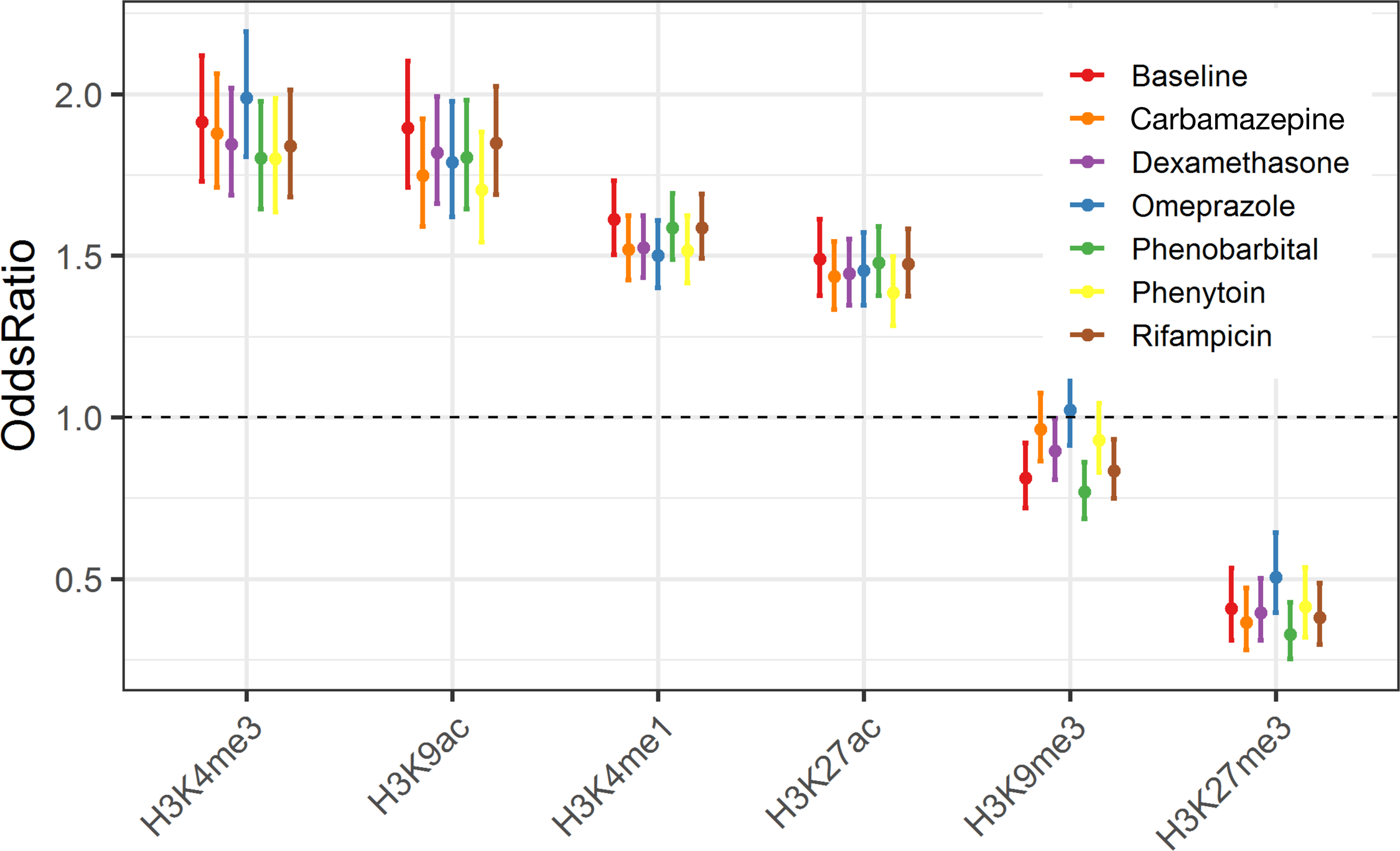

**Supplemental Figure 3: Enrichment of TBT eQTLs in Roadmap Histone Modifications**

We tested the enrichment of TBT eQTLs in each treatment with the Roadmap Histone Modification annotations mapped in liver. We compared each eQTL set to a randomly sampled set of 1000 null SNPs that were matched for LD score, minor allele frequency and distance to the nearest gene for each of the eQTL. We found enrichment across all drug treatments for the active histone markers, H3K4me3, H3K9ac, H3K4me1 and H3K27ac. These eQTLs were also less likely to be found in the repressed histone marker H3K27me3.

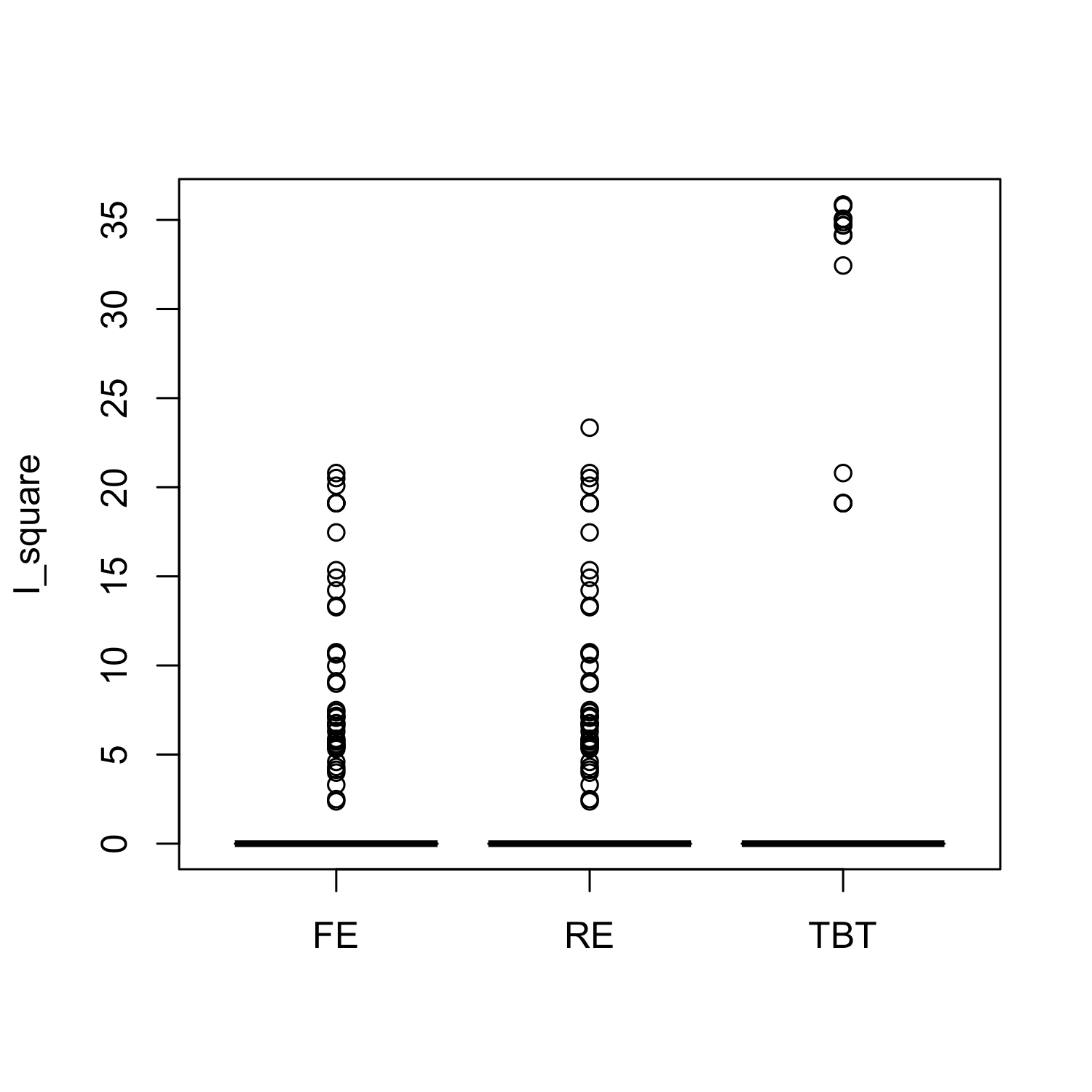

**Supplemental Figure 4: heterogeneity of eQTLs effects across conditions**

The I^2^ statistics estimates the percentage of variation in eQTL effect size explained by heterogeneity across all conditions. TBT eQTLs had more extreme I^2^ than the meta-tissue FE model and the meta-tissue RE model eQTLs.

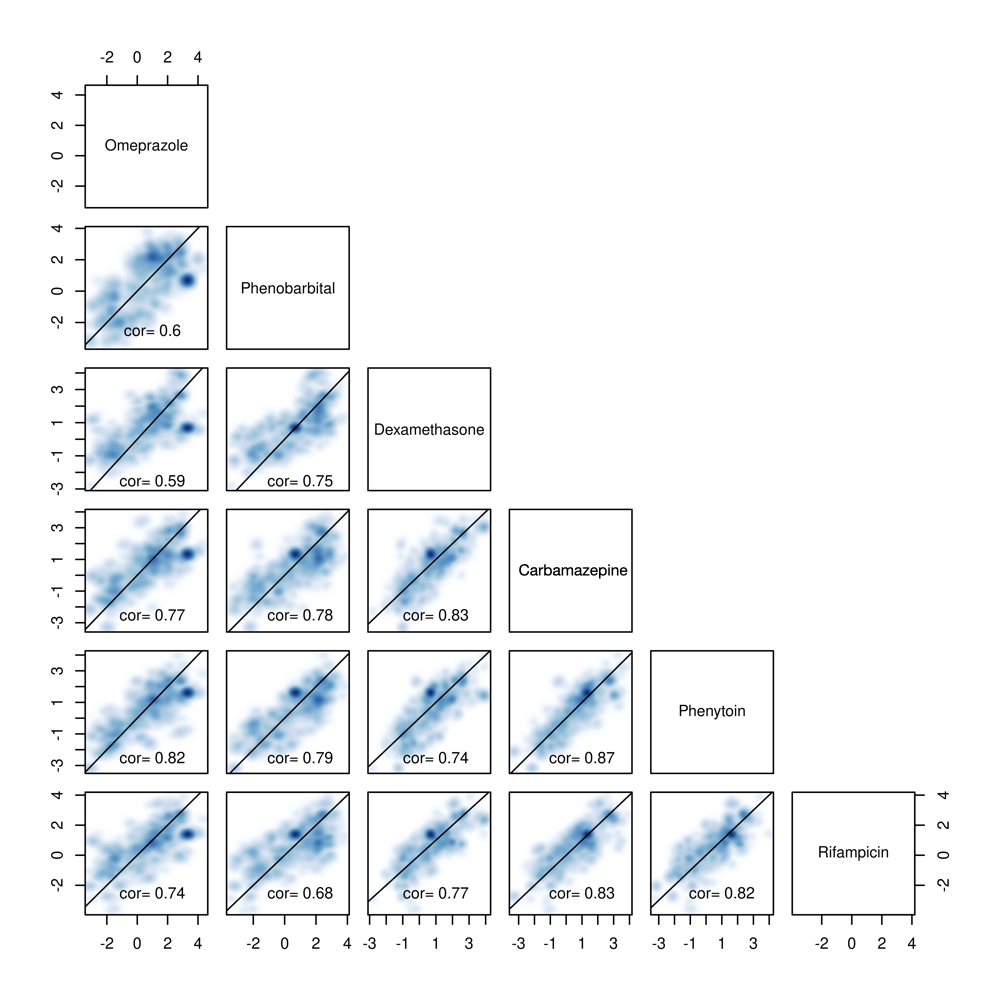

**Supplemental Figure 5: Correlation between TBT eQTLs Z scores across treatments**

The pairwise correlation of z-scores for TBT eQTLs in each treatment are shown as a smoothed scatter plot. Color denotes higher density and the Pearson R^2^ for each comparison is shown in the box.

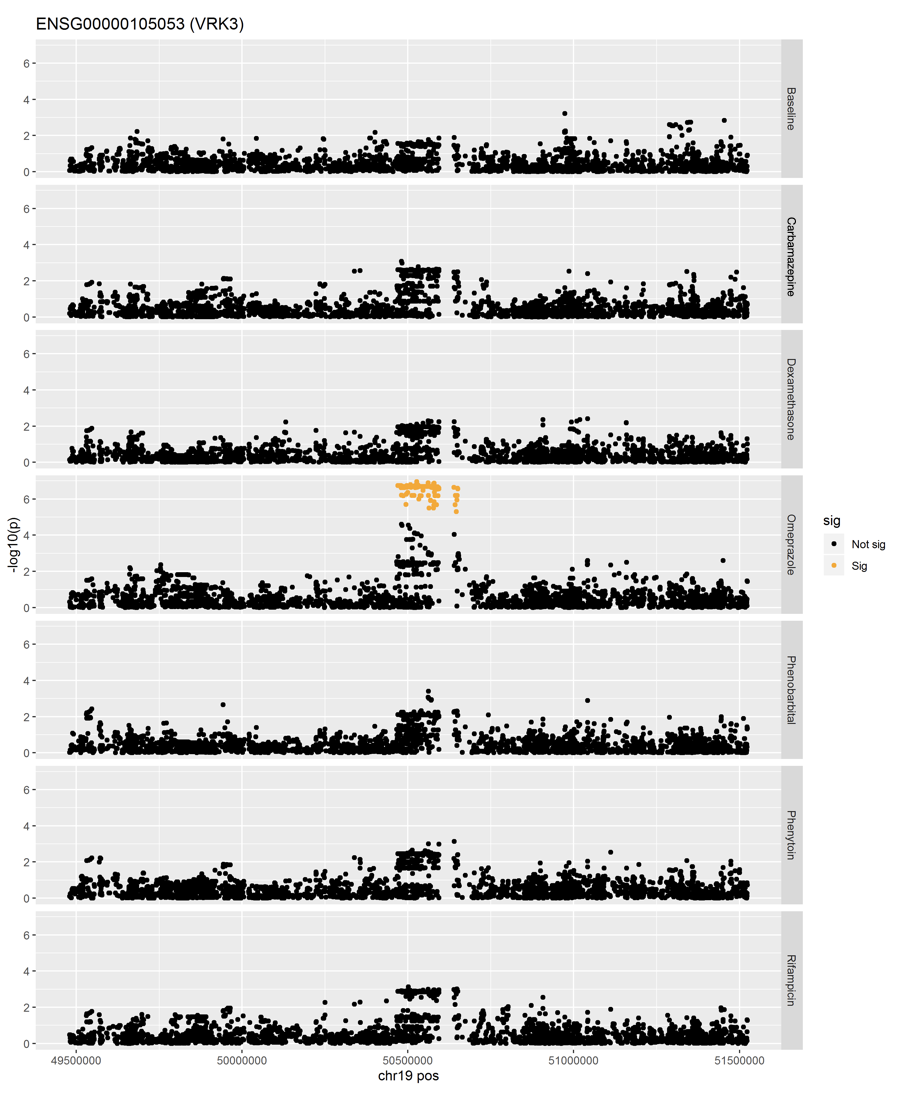

**Supplementary Figure 6: Drug specific eQTL for VRK3 found only after Omeprazole treatment.**

LocusZoom plots for cis-eQTLs of *VRK3* in each drug treatment. Dots along the X-axis represent tested SNPs within the cis-window by chromosomal location and the Y axis is the log_10_ p-value for the association. Yellow dots show statistically significant association after multi-test correction.

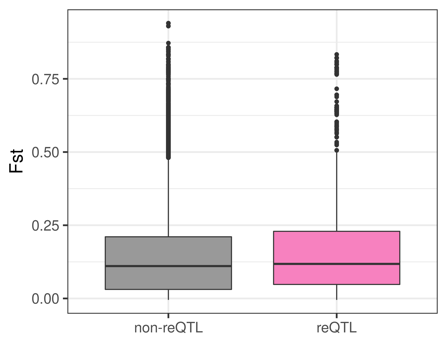

**Supplementary Figure 7: Average F_st_ in reQTL and non-reQTLs**

We calculated F_st_ values using the 1000 Genomes phase 3 data using the YRI and CEU populations with GCTA. The reQTLs and non-reQTLs were defined in Methods. (Wilcoxon rank sum one-sided test p=3.62e-10, median F_st_ of reQTL = 0.118; median F_st_ of non-reQTL: 0.111). This finding suggests allele frequency differences in regulatory variants may underly population differences in drug response.

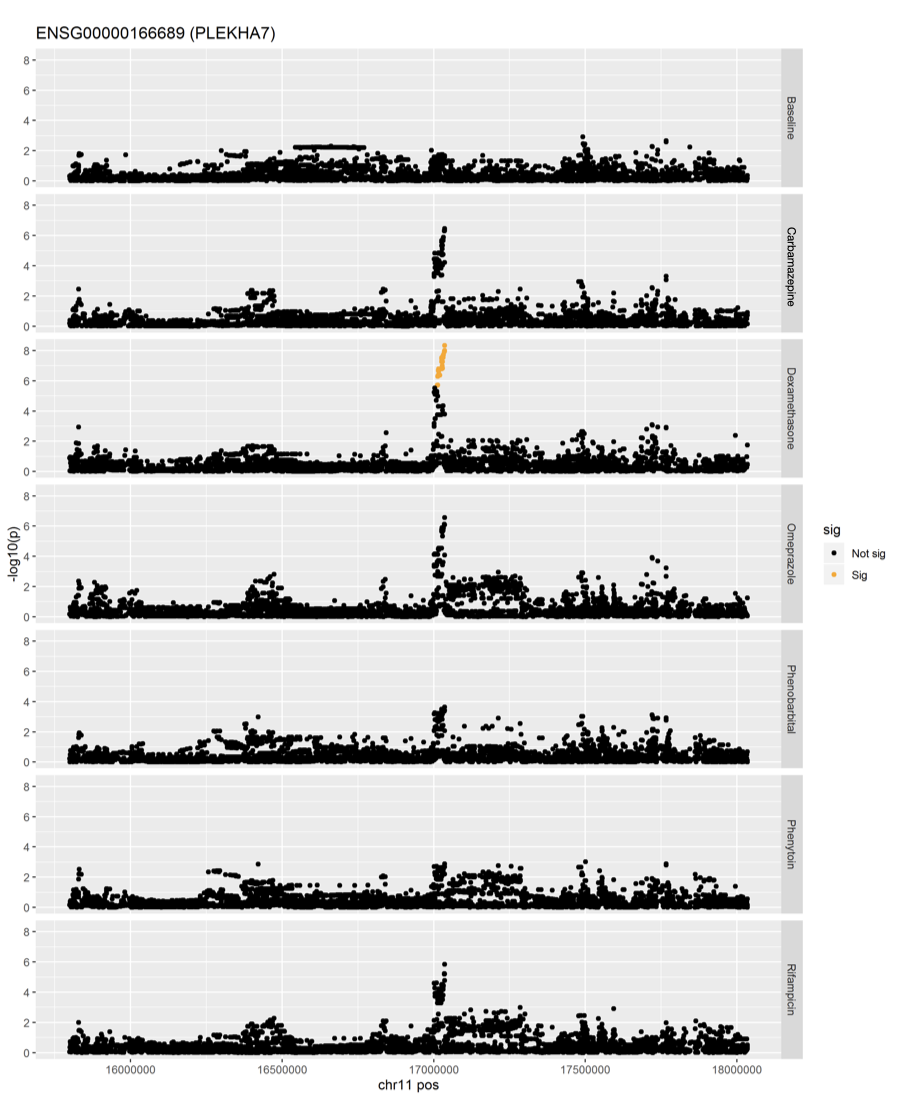

**Supplementary Figure 8: Drug specific eQTL for *PLEKHA7* found only after Dexamethasone treatment.**

LocusZoom plots for cis-eQTLs of *PLEKHA7* in each drug treatment. Dots along the X-axis represent tested SNPs within the cis-window by chromosomal location and the Y axis is the log_10_ p-value for the association. Yellow dots show statistically significant association after multi-test correction. While PLEKHA7 was not differentially expressed between any two drugs, significant eQTLs were found after Dexamethasone treatment.

**Supplementary Figure 9: Correlation between lfc QTL and reQTL z-scores**

Scatterplots of each treatments show low correlation between lfc QTLs and reQTLs. Pearson correlation R^2^ shown for each treatment.
